# Evolutionary conserved multifunctional nitric oxide synthesis proteins responding to bacterial MAMPs are located at the endoplasmic reticulum

**DOI:** 10.1101/2020.07.12.191361

**Authors:** Wenhui Zheng, Hongchen Li, Wenqin Fang, Simon Ipcho, Rosanna C. Hennessy, Bjoern Oest Hansen, Guodong Lu, Zonghua Wang, Mari-Anne Newman, Stefan Olsson

## Abstract

Most Eukaryotic organisms produce nitric oxide (NO); however, the mechanisms underpinning NO’s biosynthesis are only known in animals. In animals, there seems to be a non-described additional system for producing NO in many cell types, including blood vessels where NO is essential for blood pressure control. NO is known to be a signalling molecule of the innate immunity system in plants and fungi although no NO generation has yet been described. In the plant pathogenic fungus *Fusarium graminearum,* we demonstrate an extra NO-producing system in fungi that seems also present in mammals and plants and, thus, likely the evolutionary original. The discovered NO-producing enzymes are already well-known sterol-producing enzymes with more than one function. Both these enzymes are targets for statins and the major fungicides; thus, the NO production of the new system has consequences for agriculture (pathogen resistance and control) and medicine (blood pressure control, immunity and sepsis).

## Introduction

Organisms rely on NO as a signalling and effector molecule for numerous physiological and cellular processes. NO is a diffusible, short-lived ^1^, reactive free radical that acts as a secondary messenger ^2,3^ and NO production is a hallmark of the basal defence response against bacterial infections, also known as innate immunity. Plants and animals produce NO when they detect Microbial-Associated Molecular Patterns (MAMPs) ^2,4,5^. MAMPs are conserved molecular patterns shed from microbes recognized and responded to by host organisms ^6^.

The NO production is mainly catalyzed by the enzyme Nitric Oxide Synthase (NOS), which converts L-arginine to citrulline and NO through the intermediate N-hydroxy-L-arginine (NOHA), as reviewed by ^7^. The archetypal NOS consists of two parts; an N-terminal oxygenase containing the arginine binding site with heme and a tetrabiopterin domain, a central calmodulin (CaM), and the C-terminal reductase part contains FAD, FMN and an NADPH binding site. The enzyme is active in its homodimer form and is regulated by Ca^2+^ and CaM ^7,8^.

NOSs are thus proteins containing two main parts, a Cytochrome P450 (CYP) heme-binding part and an NADP Dependent Cytochrome P450 (NCP) reductase part. To become active, they dimerize aided by CaM and Ca^2+,^ and the Heme-reductase part (the NCP) transports electrons to the heme in the heme-binding CYP in the paired proteins ^8^. NO synthesis is initiated with heme-binding and activated by O_2,_ and the reductase domain transfers NADPH-derived electrons to the oxygenase domain. NOSs are subjected to NO-self-regulation since NO binds to the iron in the heme and blocks NO synthesis at high NO concentrations ^8,9^. NOSs are similar to cytochrome P450s, with the heme and reductase domains present in both enzymes _8._

In mammals, NO is generated by the very well-characterized mammalian three different isoforms of NOS^10^. Two are constitutively expressed in the brain (neuronal NOS, nNOS) and within the endothelial cells (eNOS) the latter involved in regulating the arteries’ vasodilation. During microbial infection, the inducible NOS (iNOS) located within macrophages produces NO and mediates bacteria elimination ^10^. In humans, the iNOS is part of a structural complex that contains the cationic amino acid transporter (CAT1), arginosuccinate synthetase (ASS), arginosuccinate lyase (ASL), the NOS and the heat shock protein 90 (HSP90). ASL and ASS are enzymes responsible for producing the substrate, arginine, fed directly to the NOSs. Disruption of the complex’s structural integrity is detrimental to NO production ^11^.

Although shreds of evidence of NO-production have been reported in fungi and plants, their NOS orthologs have still not been found using sequence homology ^12–14^. In Arabidopsis, a NOS-associated enzyme (AtNOA1) was discovered and found to contribute to the induction of a NO burst in response to the MAMP lipopolysaccharides (LPS) ^15,4,16,17,3^. The AtNOA1 sequence did not resemble the archetypal NOS but was similar to a snail protein exhibiting NOS-like activity ^15^. It was later found to be a cGTPase and not to have anything to do directly with NO production ^18^.

In fungi, genes encoding enzymes responsible for NO production from arginine, like NOSs, are unknown. While the *Aspergillus* species are reported to harbour genes with sequence homology to the mammalian NOS, functional analysis is lacking ^8^. Previous studies have shown that yeast NOS-like proteins can be detected in filamentous fungi using antibodies; however, the sequence of the detected proteins has not yet been reported ^19,20^, nor are there any similarities between the NOS-sequences to the yeast genome sequence ^13^. Nevertheless, NO has been shown to affect various stages of the fungal lifecycle ranging from germination to sporulation ^14,21–23,13^

In addition to arginine, nitrite is a substrate for NO-formation in animals and plants, illustrating redundancies for this radical production ^24–27^. The nitrate reduction pathway is involved in NO production in plants, it is, however, not alone responsible for NO production in plants ^28^. The same was found when nitrate and nitrite reductase were deleted in the fungus *Magnaporthe oryzae*. The deletion mutants still produced NO and showed no altered phenotypes ^14^. Nitrite and nitrate have also been proven to be a source of NO in *F. graminearum* triggered by host perception in the rhizosphere during the precontact phase using the model monocot *Brachyopodium distachyon*. These authors also concluded that nitrite/nitrate reductase only partially contributes to NO production in *F. graminearum* ^29^. Results pointing to additional NO souces than nitrite/nitrate reduction was also found for NO formed by *Aspergillus nidulans* niaD. NO formation was not abolished in niaD deletion mutants but roughly halved, and nitrite and nitrate accumulated in ammonium media free from nitrate/nitrite, especially in the niaD deletion mutant ^30^, further indicating there should be another source of NO getting oxidized to nitrite and nitrate.

We had selected the model strain *Fusarium graminearum* strain PH1 (teleomorph *Gibberella zeae*) to investigate ^31^ whether fungi possess an immune system response to bacteria. The fungus is a severe pathogen of many kinds of grass, including wheat and causes Fusarium head blight (FHB), crown rot (FCR) and root rot (FRR) ^32^. Simultaneously, it is a soil-borne fungus active in the plant rhizosphere ^32^, where it has to co-exist with many bacteria, neutral, beneficial and detrimental to the fungus ^33^. We have previously shown that the transcriptome of *F. graminearum* responds specifically and rapidly to purified bacterial MAMPs by upregulating genes involved in Eukaryotic innate immunity responses ^33^. Our former study served as a starting point to determine whether NO formation is part of a fungal innate immune response and if so identify the proteins responsible for synthesizing NO and if they could have similarities to known systems of NO formation.

Hypotheses: Our first hypothesis is that NO is produced in response to purified bacterial MAMPs as part of an innate immune response ^33^. The second hypothesis is that NO is produced primarily from arginine, like normal NOSs, and not nitrite/nitrate through the nitrate reduction system. If the first and second hypotheses that NO is produced from arginine in response to bacterial MAMPs, NO might be produced by a non-conventional NOS system (hypothesis 3) where the heme protein and the heme-reductase belong to separate proteins that together produce NO since orthologues of mammalian type NOSes are not present in fungi or plants ^14,28^.

## Results

### NO production in response to bacterial MAMPs

NO is known to be elicited in response to bacterial MAMPS in plants and animals. We thus exposed the fungus to various MAMPs under the asme type of nongrowing conditions we had previously used for exposing fungal biomass to bacterial MAMPs ^33^. The level of accumulated NO by the fluorescent NO probe dye (DAF-FM DA) was measured after 10 hours of elicitation. Similar to in mammals, full-length flagellin FFLG, but not the short FLG22 used to elicit NO responses in plants ^34^, was found to be the most consistent elicitor of NO in the fungus (**Fig. 1A**) and was used as a reference in the subsequent investigations even though the commercial flagellin preparation contained small non-declared amounts of sucrose ^33^. We performed a control experiment under the same conditions to be sure that we detected gaseous NO developing as a response to the bacterial MAMP flagellin. For the control experiment, we used the Luminol-H_2_O_2_ fluorescence method ^35^. With that, we developed a technique for head-space detection, detecting the chemiluminescence in a scintillation vial with the reactants in an open Eppendorf tube insert and the fungal biomass in water with or without flagellin added to the scintillation vial (See **Supplementary file SF2** for the experimental setup and results). The NO donor NONOATE was used in a preceeding control experiment instead of the fungus and flagellin to test whether the technique worked and whether the amount of reactants in the Eppendorf tube were enough. Although very selective for NO, this technique was not used further since the DAF-FM DA can be used in microtiter wells and analysed by a microplate reader so that several measurements and replicates can simultaneously be run and that also can be used in microscopy.

**Figure 1.**
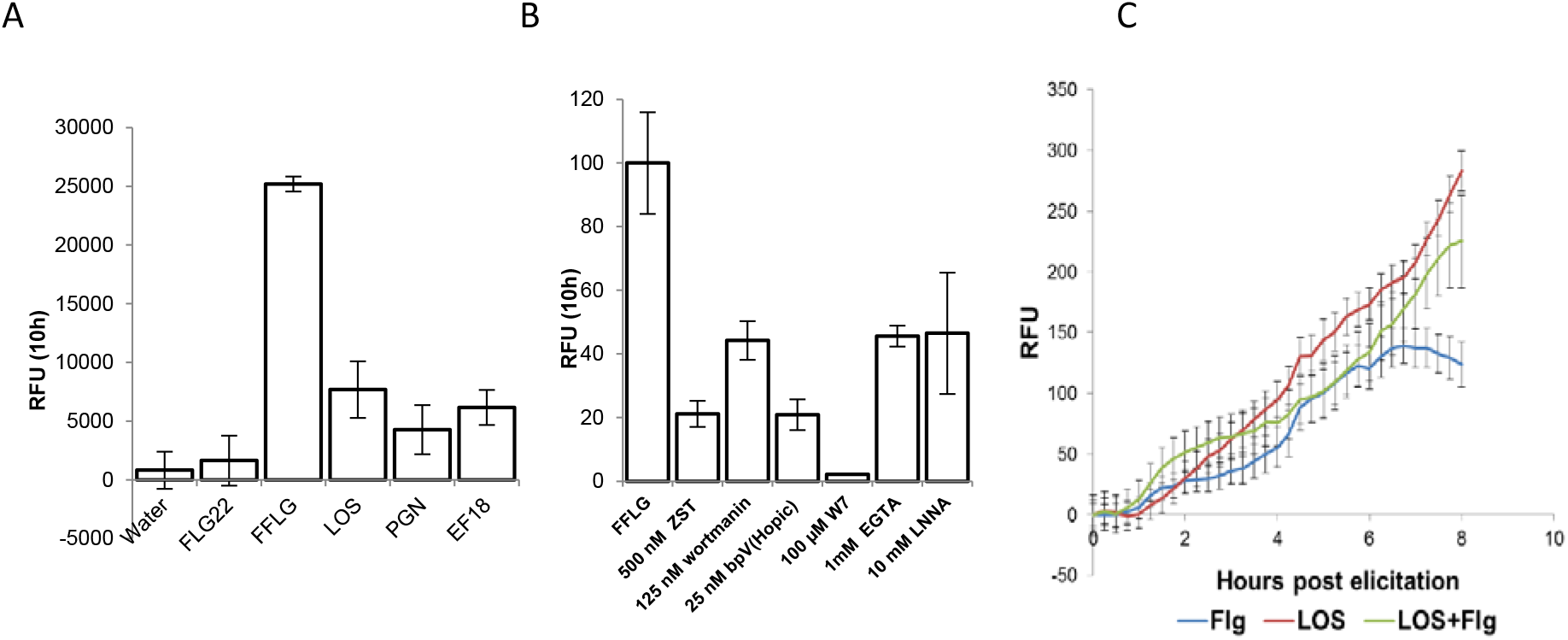
(**A**) Accumulated increased NO production measured as an increase of the NO probe DAF-FM DA’s fluorescence during 10h post-stimulation by adding different bacterial MAMPs (Microbe Associated Molecular Patterns). Water was used as a negative control to test if the disturbance by addition triggers NO production. FLG22=the short flagellin peptide, plant flagellin receptors senses. FFLG=full length flagellin, LOS=lipo-oligosaccharides, PGN=peptidoglycan, EF18= Elongation factor 18. Three biological replicates were used, and error bars show 95% confidence intervals for the mean values. Thus, the P-value for the null hypothesis for the same average of bars with non-overlapping error bars <<0.05. (**B**) Accumulated relative increased NO production was measured as an increase of the NO probe DAF-FM DA fluorescence during 10 h post-stimulation by adding full-length flagellin (FFLG) as positive control with different inhibitors together with FFLG as treatments. FFLG=Full length flagellin, ZST=ZSTK474 PI3K inhibitor, wortmannin=PI3K inhibitor, bpV(Hopic)=PTEN inhibitor, W7=calmodulin inhibitor, EGTA=Calcium chelator, LNNA=L-N^G^-nitro-L-arginine NOS inhibitor. Three biological replicates were used, and error bars show 95% confidence intervals for the mean values. Thus, the P-value for the null hypothesis for the same average of bars with non-overlapping error bars <<0.05. (**C**) The relative increase in fluorescence of the intracellular calcium probe Fluo4, that was directly added to the fungal biomass in the wells (1μg/ml), was subtracted for fluorescence in the negative control with only water. Error bars show the standard error of means (N=3).

### Test if NO production is sensitive to standard NOS inhibitors

In mammals, iNOS is regulated by signalling components such as phosphatidylinositol-3-kinases (PI3K), Phosphatase and tensin homolog (PTEN), and Calcium influx and Calmodulin ^36,37^. It is also well-documented that NOSs use arginine as a substrate to generate NO ^38^. A series of signalling inhibitors were applied to query if similar characteristics governed the NO-producing enzyme in *F. graminearum* as in the mammalian system. PI3K was inhibited with ZSTK474 and wortmannin, and PTEN was inhibited with BpV(HOpic). Inhibition of both signalling components showed a decreased level of NO production (**Fig. 1A**). Extra-cellular calcium and intracellular calmodulin were inhibited with EGTA and W7, respectively. Chelation of calcium ions showed a moderately decreased NO response (**Fig. 1B**), whereas inhibition of calmodulin completely inhibited NO production. The arginine analogue L-NNA, which an iNOS cannot metabolize, was added to elicited cultures as a competitive inhibitor (**Fig. 1B**). The results showed that the accumulation of NO signal was decreased compared to the control indicating that the NOS-system response to bacterial MAMPs in *F. graminearum* uses arginine as substrate as mammalian NOSes and that is also what seems to be the case for NO production during spore germination of the relatively closely related fungus *Magnaporthe oryzae* ^14^.

### Bacterial MAMPs increase intracellular calcium levels

When Ca^2+^ levels increase intracellularly, calmodulin is known to interact with and activate NOS ^39,40,37,41^. To test if calcium signalling is involved in the MAMPs detection, intracellular Ca^2+^ levels following bacterial MAMPs challenges were investigated, and intracellular calcium increases above water controls using the intracellular fluorescence calcium indicator Fluo4 AM that can be loaded into cells. Our results show increasing green fluorescence indicating increasing intracellular Ca^2+^ concentration triggered by the bacterial MAMPs treatments (**Fig. 1C**) that also triggered the NO responses (**Fig. 1A,B**).

### Standard substrates for NO production by NOSes are used for NO production

The ability of *F. graminearum* to utilize exogenous chemicals as substrates were tested. Arginine, the established substrate in mammalian systems, was added into bacterial MAMPs challenged cultures. Interestingly, the NO production rate increased very fast due to arginine addition and then completely stopped (**Fig. 2A**). To assess the possibility of NO production from nitrite (**Fig. 2B**), this was exogenously injected into MAMPs challenged fungal cultures. Interestingly, in both experiments with exogenous arginine and nitrite, the NO produced approximately doubled at 4h compared to at 2h. However, contrary to the arginine experiment, NO accumulation continued post-spiking. To test if the effects were additive exogenous nitrite and arginine were added in different sequential combinations. The results showed that adding nitrite promoted a higher NO production rate when arginine was added later. A typical arrest of NO production post arginine addition followed. When arginine was added before nitrite exogenously, a NO burst was observed, followed by NO production’s complete arrest. The addition of nitrites following the arrest did not promote further NO production (**Fig. 2C and D**).

**Figure 2.**
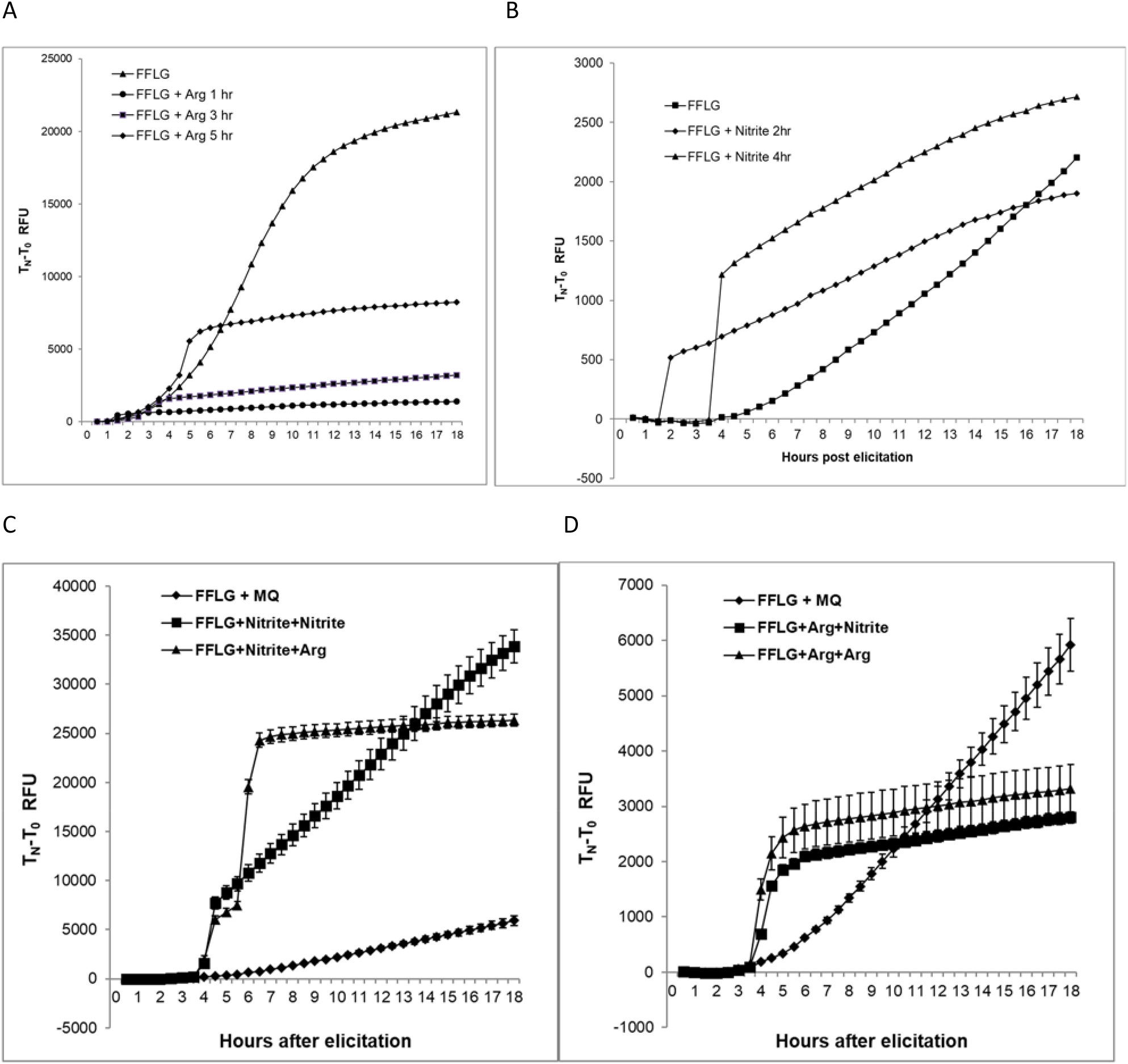
(**A**) Effect on NO production in treatment minus control of adding exogenous arginine or (**B**) nitrite on NO production. Both were added to a final concentration of 5mM. The NO production rate increased very quickly after addition, and in the case of arginine, the rapid increase stopped accumulating soon after addition. The inhibition was less in the nitrite additions. (**C**) Sequential spiking nitrite and arginine to flagellin-induced fungal cultures. First, spiking by nitrite followed by a second nitrite spiking or an arginine spiking. (**D**) First, spiking arginine followed by arginine spiking or nitrite spiking. Arginine spiking limits total NO accumulation, but this does not happen with nitrite spiking. Error bars are standard errors of the mean (N=3). In A and B, no error bars are shown since they are small and hide the shapes of the curves.

In conclusion, our results indicate that both arginine and nitrite can be substrates for the oxidative or reductive formation of NO ^14^, but only the arginine additions result in rapid cumulative increases in the DAF-FM fluorescence and indicate a high effective NO concentration since NO is very short-lived in cell systems ^1^. These results indicate that mainly arginine is responsible for a high transient concentrations of NO that can be used for fast signalling. The rapid increases seemed to self-inhibit NO formation from arginine as expected if it is formed by a heme protein similar to iNOS in mammals ^8,9^, supporting that arginine is the primary substrate for quick NO formation in response to bacterial MAMPs.

### Finding candidate NO production genes in *F. graminearum*

Our experiments thus indicated that the NO-generating system in *F. graminearum* must be very similar to the mammalian system ^8^. Nevertheless, there are no classic NOS candidate genes in fungi ^42^. The NOS-domain PF02898.15 is usually used as a signature for finding nitric oxide synthetase enzymes ^43^. Since no usual NOS genes encoding a protein with a PF02898.15 domain is present in any predicted *F. graminearum* proteins, there must be a different domain organization in the responsible protein protein or proteins. We know from the literature that NOSes need to dimerize for NO to be produced, and this dimerization is Ca^2+^-dependent. It also means that electron transport goes from the heme-reductase domains in one protein to the heme domain in the other ^8^. Thus, there could be another way to bring together a NOS-like heme domain-containing protein and NOS-like heme reductase containing all the other domains of a classic NOS. The heme domain alone in a NOS has similarities to a **CY**tochrome **P**450 heme (**CYP**), and there are many CYP-proteins in *F. graminearum*. We thus turned our interest to find a possible separate **N**ADPH-**C**ytochrome **P**450 reductase (**NCP**) containing all the other domains necessary for a NOS-NCP involved in NO-production ^8^.

The enzyme we are looking for is an ***F****. **g**raminearum* **NCP** involved in NO production, an **FgNCP_NO_**. (from now onwards, we write gene and protein abbreviations in bold where they are first introduced in the text and the proposed cellular function as a subscript). We used the amino acid sequence of the C terminal non-heme part of the Mouse iNOS (NP_035057.1) as bait and made a BLAST search against the *F. graminearum* proteome. We only found 2 candidates FgNCP_NO_ (FGSG_ FGSG_08413 and FGSG_09786), with high similarity and containing all necessary domains ^8^ (**Supplementary File SF2**). Both genes contain regions that indicate NADPH-dependent Cytochrome P450 heme reductases (NCPs) that could be involved in transporting electrons from NADPH to a heme-containing protein that produces the NO from a reaction between arginine and O_2_.

We decided to use our previous transcriptome data to prioritize which gene to study in detail (**Supplementary Data1**) to test if any of the two candidate genes are co-regulated with a putative *F. graminearum* **N**itric **O**xide **D**ioxygenase (**FgNOD**) needed to protect a NO producing cell from NO ^44^. We calculated the correlation coefficients (Pearson) for transcription of the putative *FgNCP_NO_* candidates of the two putative *FgNODs* for all *F. graminearum* genes in 113 transcriptomes from data from our previous experiments where *F. graminearum* has been exposed to different bacterial MAMPs (see methods) and sorted the genes in order of highest to lowest correlation (**Supplementary File SF3**). We found that only FGSG_09786 was highly correlated in expression with the two putative NODs, FgNOD1 and FgNOD2 (FGSG_00765 and FGSG_04458). Both NOD genes are similar to NO dioxygenases in *Candida albicans* that are experimentally verified ^45^ and also to a NOD annotated for *Fusarium sporothrichiodes* (**Supplementary File SF2**) FgNOD1 is almost identical to the *F. sporothricioides* protein (100% cover E-value 0 and identity 98%), and FgNOD2 is similar to FgNOD1 (90% cover, E value 2e-117 and identity 45.61%). FgNOD1 and FgNOD2 have recently been confirmed to be responsive to nitrosative stress and have the suggested functions in removing NO ^46^. The correlation of expression of the two NODs and FGSG_09786 suggests that FGSG_09786 is a putative FgNCP_NO_ (**Fig. 3**). The other possible *FgNCP_NO_*gene, FGSG_08413, was not highly expressed in our bacterial MAMPs data and even showed a slight negative correlation with the two putative FgNODs (**Fig. 3**). It is thus not likely encoding a FgNCP_NO_. FgNOD1 is also more closely co-regulated with FgNCP_NO,_ suggesting that FgNOD1 is the primary NOD responsible for limiting NO stress, assuming FgNCP_NO_ directly participates in NO-generating activities.

**Figure 3.**
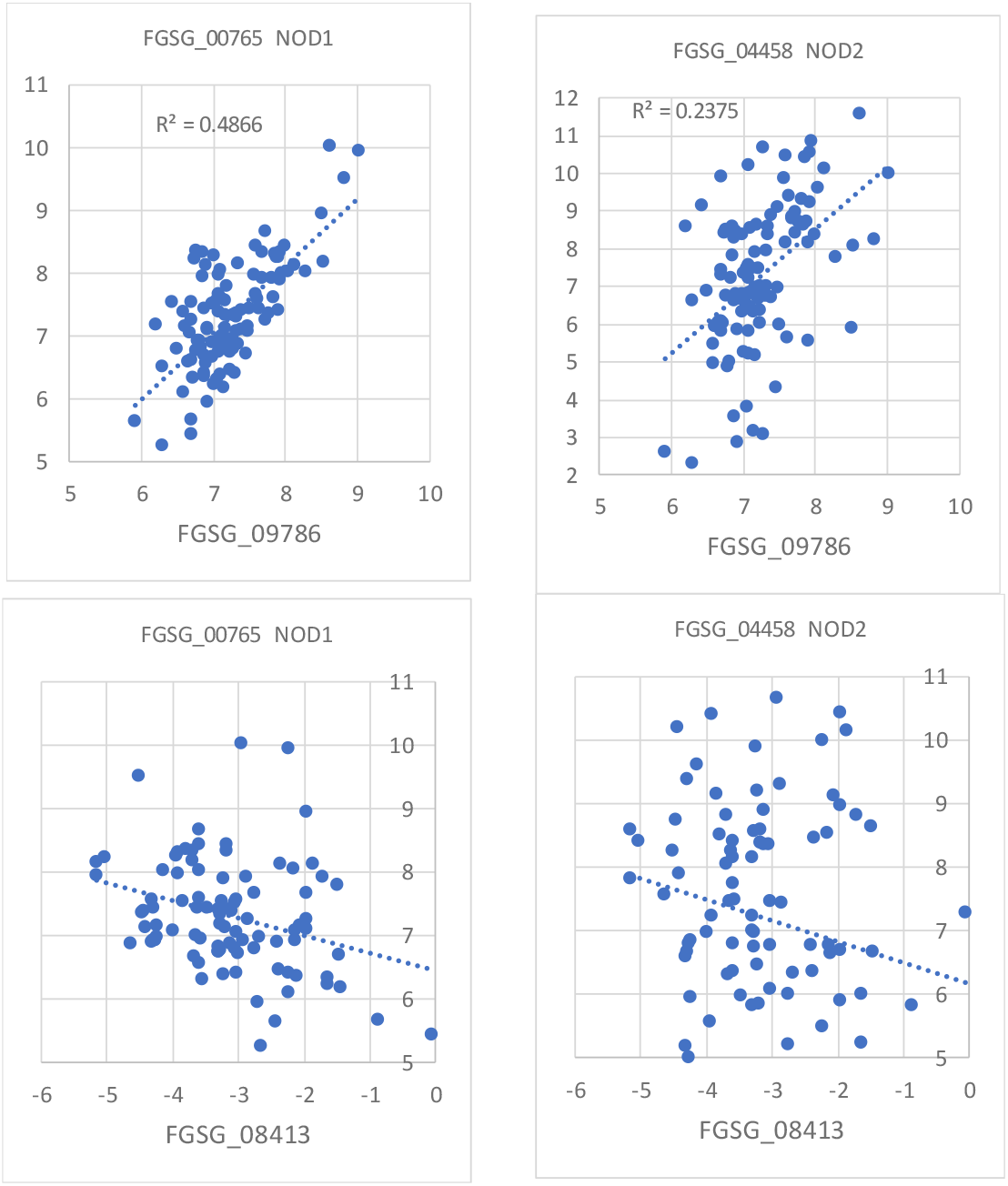
Correlations of expression between the two putative *FgNCP_NO_* (FGSG_09786 and FGSG_08413) and the two putative *FgNODs* (FGSG_00765 NOD1 and FGSG_04458 NOD2) in the transcriptome dataset for fungal interactions with bacterial MAMPs (see methods). FGSG_09786 is positively correlated with the *NOD* genes and is thus more likely to take part in producing NO. Log2 values for the relative expression are used for both axes, and scales for both axes are set the same to aid visual comparisons. N=108.

### Deletions of putative *FgNCP_NO_*, *FgNOD1* and *FgNOD2* and the response to bacterial MAMPs

#### NO production of mutants

Deletion and GFP complementation strains for *FgNCP_NO_*, *FgNOD1* and *FgNOD*2 were constructed (see methods (**Fig. S1**)) and tested for NO production. We found that the *ΔFgNCP_NO_* had a radically lower accumulation rate of DAF-FM fluorescence than the WT, indicating a reduced NO production rate (**Fig. 4A**). When focusing on the difference between controls and induced responses using bacterial free washings from washed bacteria (see methods), the NO response to the MAMPs was markedly diminished (**Fig. 4B**). Of the two NOD deletions, only *ΔFgNOD1* showed a tendency to respond to NO-signals faster than WT suggesting that FgNOD1 could be the main counterpart to the NO generating system limiting NO stresses in *F. graminearum*.

**Figure 4.**
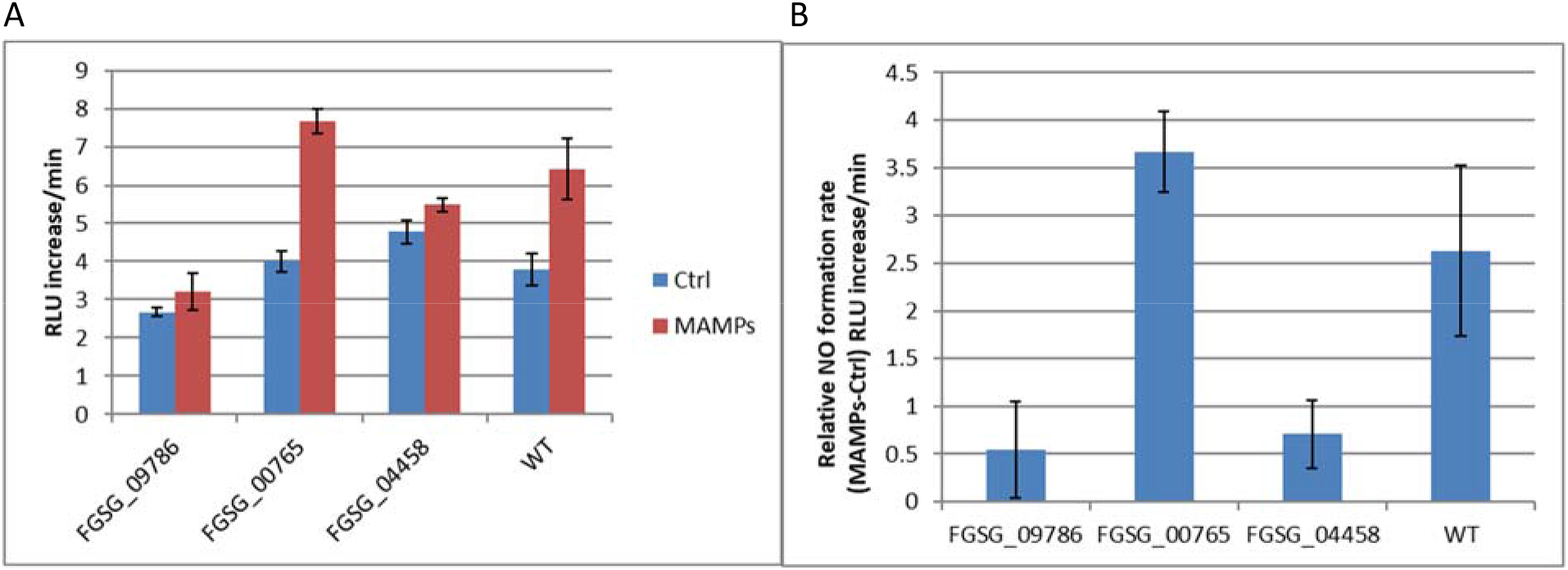
**(A)** Relative increase rate in fluorescence units of the NO probe DAF-FM per minute in water controls hyphae treated with bacterial MAMPs (sterile bacterial washings, see methods). (**B**) The same data was normalized by their respective water controls. The results show that FGSG_09786 is the sought for *FgNCP_NO_* and, of the two putative FgNODs, FGSG_00765 is more likely the NOD counterpart limiting NO production. FGSG_09786=*ΔFgNCP_NO_*, FGSG_00765= *ΔFgNOD1* and FGSG_04458=*ΔFgNOD2.* Four replicates were used, and error bars show 95% confidence intervals for the mean values. Thus, the P-value for the null hypothesis for the same average of bars with non-overlapping error bars <<0.05.

#### Subcellular localization of FgNCP_NO_, FgNOD1 and FgNOD2 and NO production

FgNCP_NO_-GFP localizes to endoplasmic reticulum (ER)-like structures in spores and hyphae (**Fig. S2 I**) and could also be shown to co-localize with an ER-tracker (**Fig. 5A**), and an mCherry labelled ER-marker protein (**Fig. 5B**). In contrast, FgNOD1-GFP localizes to mainly to the cytoplasm in germinating conidia, and FgNOD2-GFP localizes to cytoplasm and dots (**Fig S2 II and III**). Since the FgNCP_NO_-GFP localizes to ER, we tested if NO production also occurs preferentially in the ER to check further the likelihood that it is involved in NO production. DAF-FM is a non-fluorescent fluorescein derivative that only becomes strongly fluorescent after reacting with NO. DAF-FM is loaded in the diacetate form, DAF-FM DA, in the same way as FDA. The two acetate groups are cleaved off by always-present cytosolic esterases leaving strongly fluorescent and relatively hydrophilic fluorescein, accumulating in the cytosol if the plasma membrane is intact. However, the membranes are slightly permeable to fluorescein; thus, the external medium and intracellular compartments, like vacuoles, will be coloured within minutes. Consequently, we needed to load the fungus with DAF-FM DA as fast as possible and see where the first green signal appeared after it had been taken up and reacted with NO. We did not need to use MAMPs to elicit NO as the background NO formation rate was nearly too fast to get good images without green signals leaking out from the fungus. Good signals appeared already minutes after the addition of the NO probe. As can be seen, NO is produced in the ER but also in non-ER dots (**Fig. 5C**).

**Figure 5.**
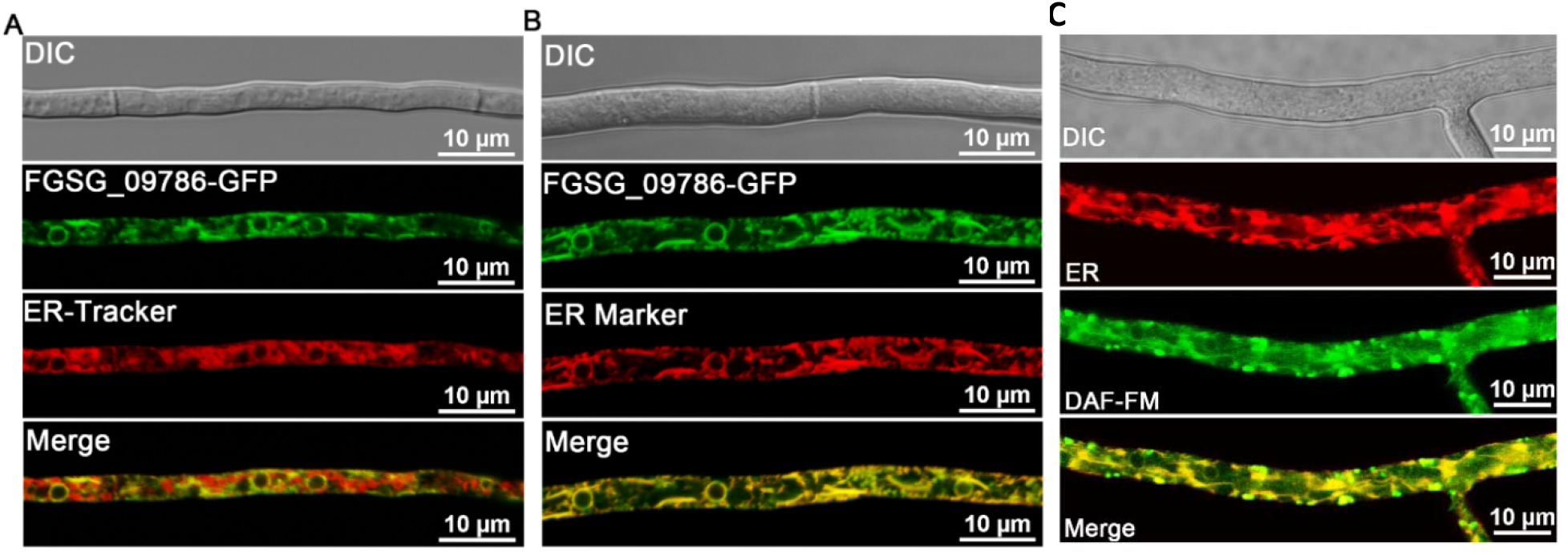
(**A**) Cellular localization of FgNCP_NO_ (FGSG_09786) in hyphae of *F. graminearum* compared with the localization of an ER-tracker often used to label ER but with less specificity than (**B**) mCherry labelled ER Marker protein (see methods). (**C**) Localization of NO signal (DAF-FM) compared with the localization of the ER Marker protein (ER). The results show that the FgNCP_NO_ (FGSG_09786) localizes to the ER, where the NO is mainly produced.

We can thus conclude that the heme reductase FgNCP_NO_ is involved in NO production, and the NO production occurs in the ER where the heme reductase is located and as wella as in dots that might be peroxisomes where NO_2_ and NO_3_ reduction is known to take place. FgNOD1 is likely the FgNOD that could be the main responsible for limiting NO in the nucleus and cytoplasm, and FgNOD2 might be active in the dots (that are maybe peroxisomes) (**Fig S2 II and III**). Our transcriptome data thus indicated that FgNOD1 is the primary NOD since it is most co-regulated with FgNCP_NO_ (**Fig. 3**).

### Phenotypic effects of deletion of NO generating and NO protecting proteins

Phenotype analysis showed that deletion of the FgNCP_NO_ had little effect on growth rate (**Fig. 6A and Fig. S3A**), but conidia formation was inhibited (**Fig. 6B**), and there was no formation of sexual spores (**Fig. S3B**). Pathogenicity was also reduced in the FgNCP_NO_ (**Fig. 6C** and **Fig. S4**. That is probably mainly caused by the lack of production of the phytotoxic metabolite deoxynivalenol (DON) (**Fig. 6D**) since the trichothecene DON is needed for full pathogenicity ^47^. NO signalling is known to be involved in conidial production^30^, so the lowered conidia production could result from this lack of NO (**Fig. 6B**). There was otherwise minor or no effect of the *FgNCP_NO_*mutation on stress phenotypes indicating cell wall and membrane changes, except the *ΔFgNCP_NO_* mutant was somewhat more sensitive to Na^+^ and K^+^ ionic stresses (data not shown). The *NOD* deletions had minimal effect on growth and infection phenotypes, indicating that NO protection is not very important for pathogenicity or growth on traditional sterile lab media (**Fig. 6**).

**Figure 6.**
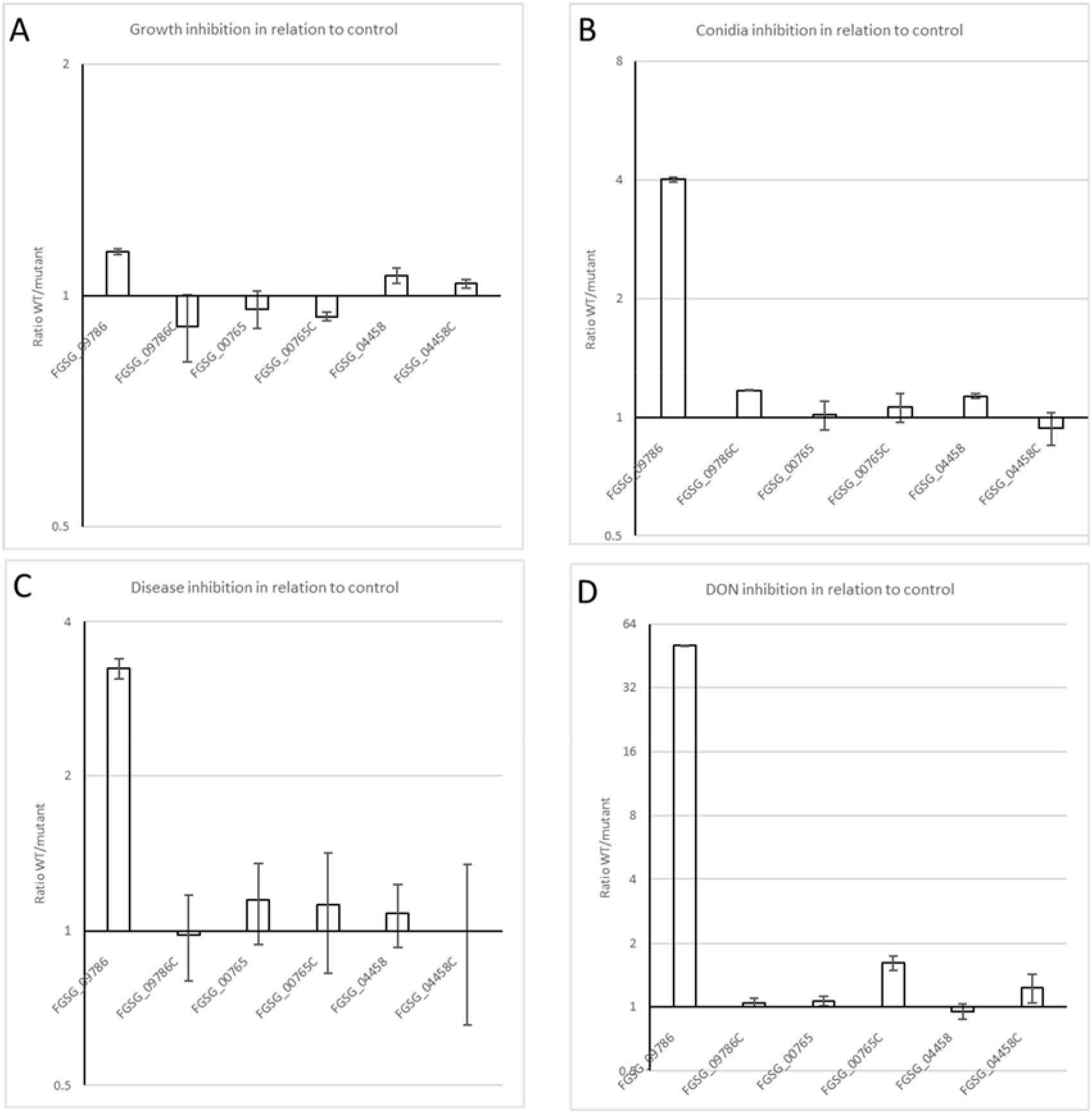
Phenotypic characterization of the mutants. Inhibition effect of deletions and complementations (C) compared with intact PH1 WT genes. Y=(PH1/Mutant or complement). FGSG_09786=*ΔFgNCP_NO_*, FGSG_00765= *ΔFgNOD1* and FGSG_04458=*ΔFgNOD2.* (A) Colony diameter on CM agar. (B) Inhibition of conidiation. (C) Disease inhibition. (D) Inhibition of plant mycotoxin formation DON. Radial growth was little affected but conidia and diseases disease and especially the formation of DON was severely inhibited by deleting FGSG_09786. The deletion of the other genes showd no or little inhibition. Three replicates were used, and error bars show 95% confidence intervals for the mean values. Thus, the P-value for the null hypothesis for the same average for bars with non-overlapping error bars <<0.05.

### Finding a putative heme protein that works together with FgNCP_NO_ to produce NO

Since we found the NO-producing heme-reductase (FgNCP_NO_) and knew where it is located, we turned our interest towards the heme protein that should be the one making the NO using electrons delivered from the FgNCP_NO_. The expected heme-binding protein should have the following characteristics except predicted heme binding. 1. It should be co-regulated with the FgNCP_NO_ in transcriptome data when the fungus is exposed to bacterial MAMPs. 2. It should be predicted to be located at the ER membrane.

One heme protein was on top of our list for genes co-regulated with FgNCP_NO_ when exposed to bacterial MAMPs, FGSG_07925 (**Supplementary File SF5**). The only problem was that it was not predicted to be localized to the ER but so was not FgNCP_NO_ either iosing PSORTII. We knocked out FGSG_07925 and complemented it with FGSG_07925::GFP, but this protein did not localize to the ER (**Fig. S6**). Neither did the mutant show any notable phenotype except a decrease in conidiation (**Fig. S7**).

We later found that the localization prediction server DeepLoc 1.0 ^48^ correctly localized FgNCP_NO_ to ER, as we had found experimentally. Using DeepLoc and our transcriptomic data from the MAMPs experiments we found that only *FGSG_01000* was predicted to encode an ER-localized heme-containing protein that is also co-expressed with FgNCP_NO,_ and that expression is in a close to 1/1 ratio of max expression compared to the max expression of FgNCP_NO_ (**Table S8 and Supplementary File SF6**) making it a better candidate for an ***F****. **g**raminearum* **CY**tochrome **P**450 involved in **NO** production, a **FgCYP_NO_**

In this case, we could obviously not trust PSORTII for localization prediction since it wrongly predicts the localization of the ER-localized FgNCP_NO_. (**Table S5**).

### Deletion of the putative FgCYP_NO_: Localization, NO production rate and effect on phenotype

We decided to knock out *FgCYP_NO_* (FGSG_01000) and test for NO-production. We successfully knocked it out (**Fig S9**) and could show a significantly decreased NO production for Δ*FgCYP_NO_* to the same level as for the Δ*FgNCP_NO_*(**Fig. 7A**) using our newly developed technique employing confocal microscopy to get cleaner data and avoid fluorescent specks not related to the NO-production (See methods). As predicted, the FgCYP_NO_ localizes to ER-like structures (**Fig.7 B**). We can conclude that the putative FgCYP_NO_ is ER located and responsible for NO production.

**Figure 7.**
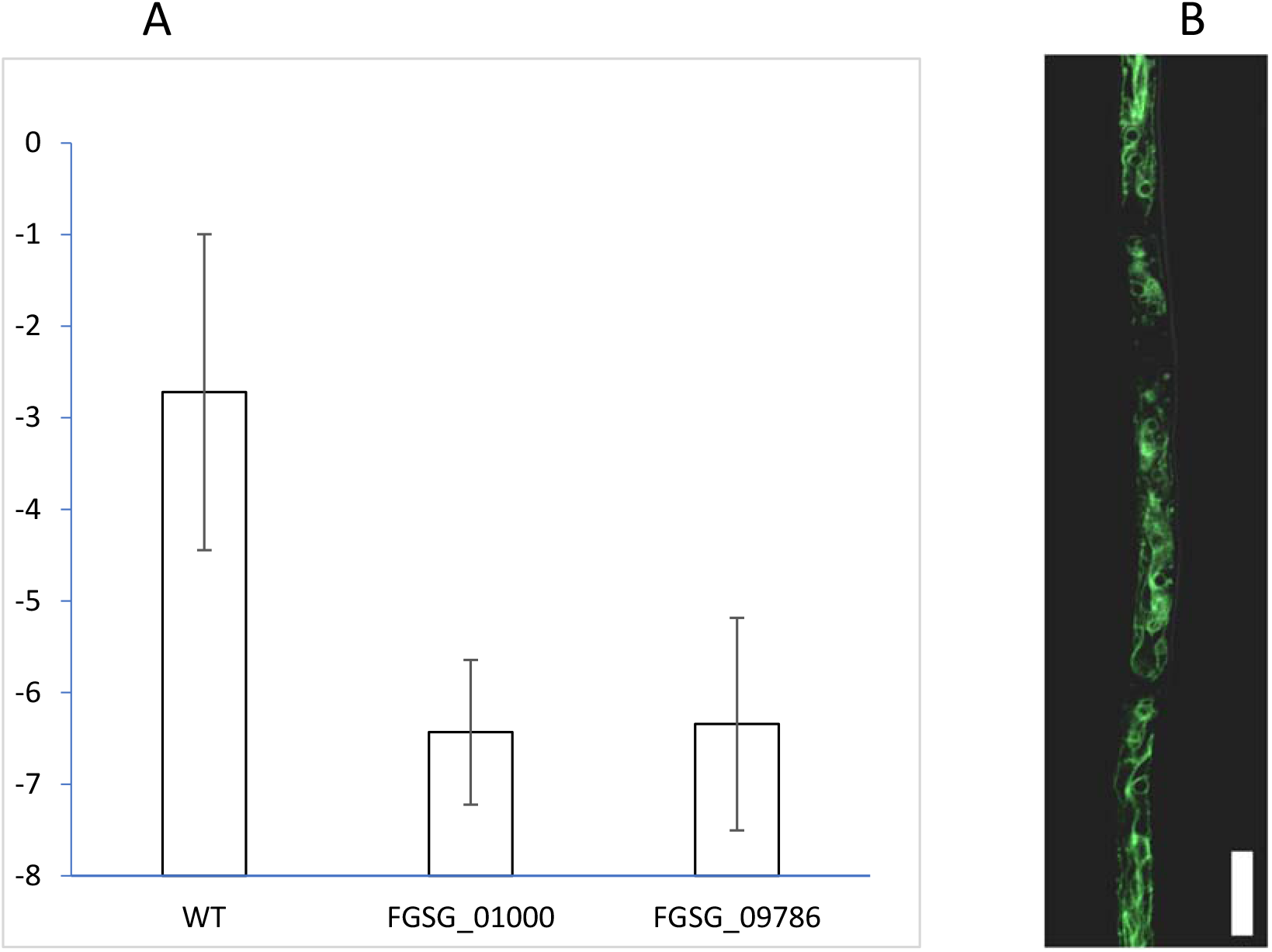
NO production in the FGSG_01000=*ΔFgCYP_NO_*mutant. *ΔFgCYP_NO_* mutant compared to WT (PH-1) and the FGSG_09786 as response to added bacteria free washings (containing bacterial MAMPs). (**A**) NO production was measured as LOG2 (NO signal/biomass signal) using confocal microscopy of large areas with many hyphae in WT=PH1, FGSG_01000= *ΔFgCYP_NO_* FGSG_09786= *ΔFgNCP_NO_*deletions and WT for NO formation rate (see methods). The FGSG_09786 was used as negative control for NO responses. Three replicates were used, and error bars show 95% confidence intervals for the mean values. Thus, the P-value for the null hypothesis for the same average of bars with non-overlapping error bars <<0.05. (**B**) FgCYP_NO_-GFP (FGSG_01000-GFP) localizes to ER-like structures. Size bar 10μm. The results indicate that FGSG-01000 encodes FgCYP_NO_ that receives electrons from FgNCP_NO_ and produces NO.

Phenotype analysis of the *ΔFgCYP_NO_* showed some effect on growth (**Fig. 8A**) but more effect on stress tolerance (**Fig. 8B and Fig. S10A**) and pathogenicity (**Fig. S10 B and C**) and a decrease in DON production (**Fig. 8C**). Although a notable decrease in DON production was registered, DON production was not absent, as for the Δ*FgNCP_NO_*. The stress phenotypes indicate also that **FgCYP_NO_** plays a role in the cell wall and plasma membrane integrity. We knew that sexual spore formation was reduced in the Δ*FgNCP_NO_* (**Fig. S3B**), so we decided to also look at the effects of ascospore formation for both mutants. Both Δ*FgNCP_NO_* and *ΔFgCYP_NO_* formed abnormal asci, even if the latter formed similar amounts of perithecia as the WT (**Fig. S13**).

**Figure 8.**
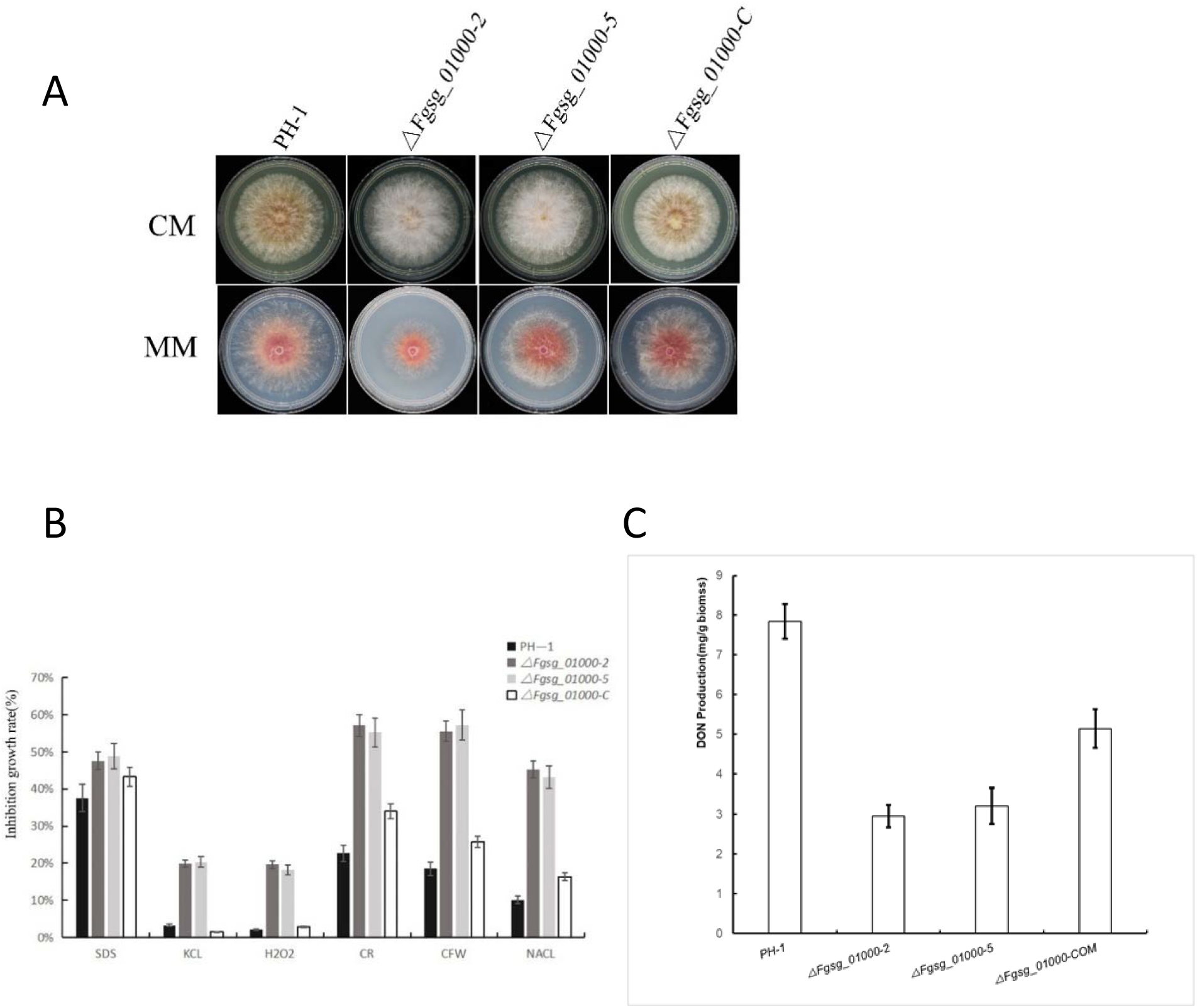
Phenotypic characterization of the *ΔFgCYP_NO_*mutant strains Fgsg_01000-2 and Fgsg_01000-5. (**A)** Growth of WT (PH-1), mutants and complement strains on complete medium (CM) and minimal medium (MM). (**B**) Phenotypic effects of stress treatments by addition of stressors to CM-medium. SDS=0.01%, KCL=1M, H_2_O_2_=5 mM, Congo red CR=1 mg/ml, CFW=200μl/ml, NaCl=1M. Three replicates were used, and error bars show 95% confidence intervals for the mean values. Thus, the P-value for the null hypothesis for the same average of bars with non-overlapping error bars <<0.05. (**C**) Effect of FGSG_01000= *ΔFgCYP_NO_* on DON mycotoxin production. There was little effect on growth caused by the deletion but more effects on stress phenotypes linked to cell membrane and cell wall effects as well as a pronounced effect on DON mycotoxin production. Three replicates were used, and error bars show 95% confidence intervals for the mean values. Thus, the P-value for the null hypothesis for the same average of bars with non-overlapping error bars <<0.05.

### Phylogenetic analysis of the two proteins involved in NO production

Phylogenetic analysis showed a high degree of conservation of both the NCP_NO_ and the CYP_NO_ in a wide range of Eukaryotes (**Fig. 9**). Most of the putative CYP_NO_-proteins were, to our surprise, annotated as CYP51 (ERG11 yeast) involved in sterol synthesis. In contrast, the NCP_NO_-proteins were annotated as NADP-dependent cytochrome P450 protein reductases as expected (**Supplementary File SF2 and SF6**). The domain organization for all these othrologous proteins was almost identical to the mouse iNOS but separated into two proteins with a predicted N-terminal hydrophobic attachment to the ER membrane (**Fig. 10 and Supplementary File SF7 and SF8**). The FgCYP_NO_ contain a CaM-binding domain that could indicate its activity is positively regulated by calcium, as our result indicates (**Fig. 2 and 3**). FgNCP_NO_ is involved in DON production necessary for full plant pathogenicity, and this production might also be Ca-regulated, as shown in the literature ^49^. We find that the FgNCP_NO_ orthologues contain a signal for a putative CaM-binding region using the CaMELS web tool ^50^, possibly explaining why DON-toxin production ^49^ and pathogenicity are dependent on Ca.

**Figure 9.**
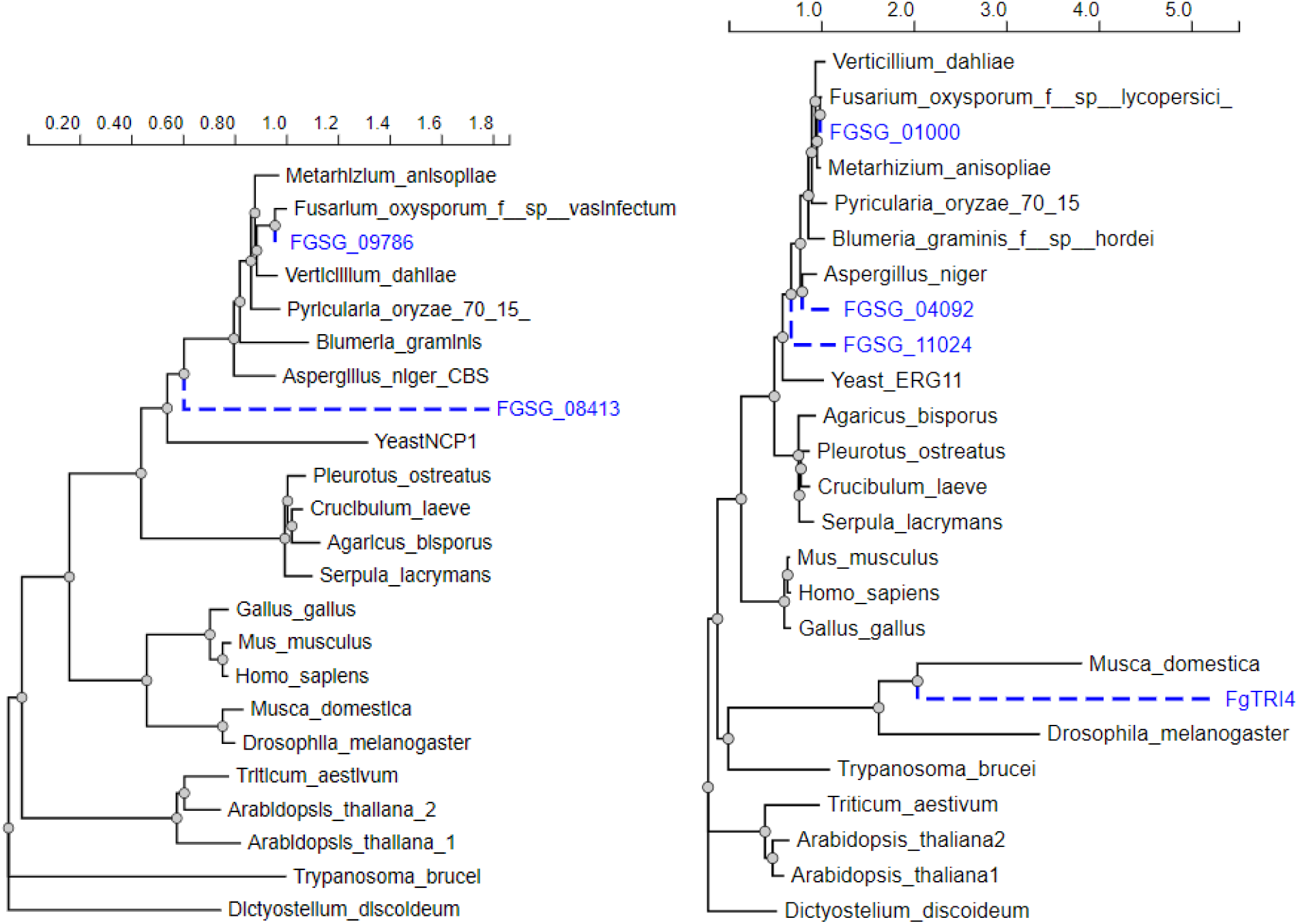
Comparing othologues of the two proteins (FGSG_09786 and FGSG_01000) involved in NO production in orthologous proteins from other eucaryotes at progressively larger taxonomic distance . (A) All proteins are annotated as NADPH-cytochrome-P450 oxidoreductases (NCPs) in the NIH database, and some are known to be involved in sterol synthesis. (B) All are CYPs in the database, and except FgTRI4 involved in DON synthesis, all are annotated as lanosterol 14-alpha-demethylases (=ERG11 in yeast) involved in converting lanosterol to the next step in sterol synthesis. The FGSG_04092 and FGSG_11024 only become activated late during infection, where Tri4 is highly activated *in planta* according to transcriptome data (not shown). For a complete analysis and protein sequences, see **Supplementary files SF7 and SF8**. Our analysis indicate that there are putative orthologues of the two NO generating proteins in all included species from amoeba to human and can also be involved in sterol synthesis.

**Figure 10.**
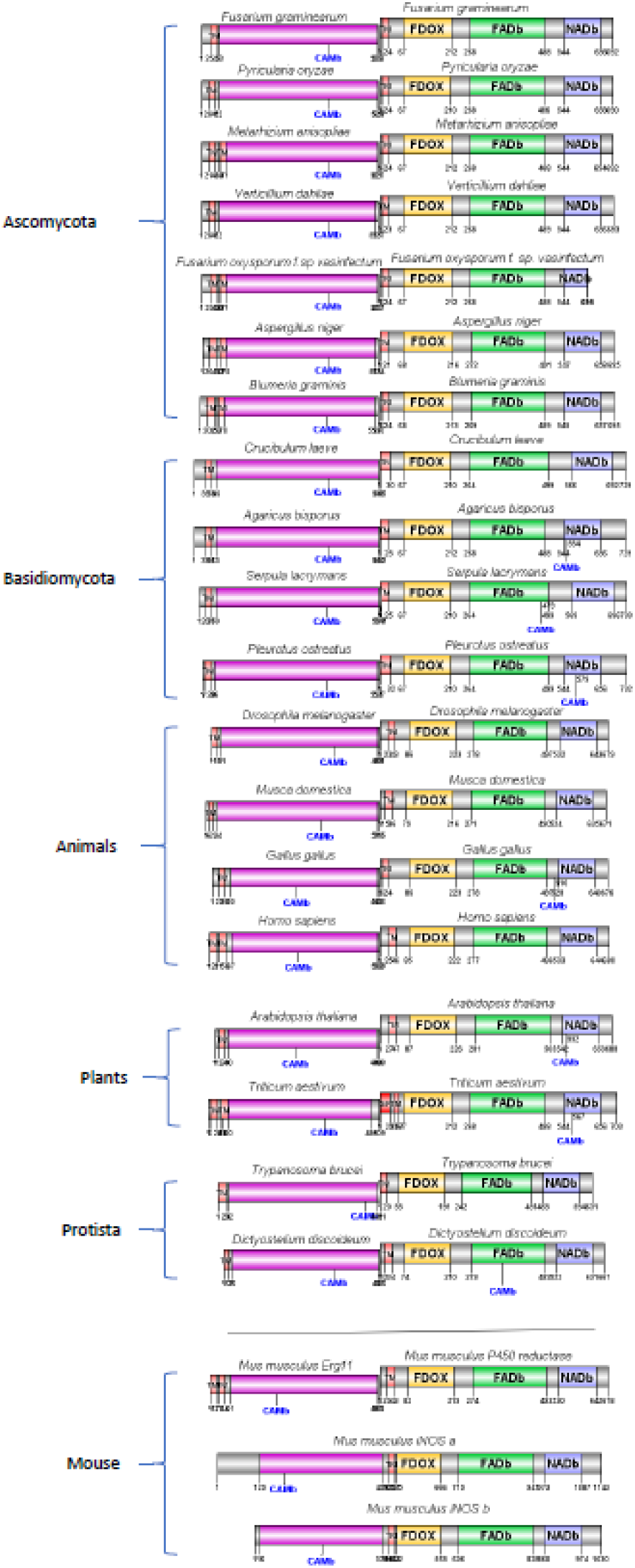
Domain structure of mouse iNOS (bottom) compared to the homologues to the CYP-part of the iNOS left of the hydrophobic helix (TM-marked) region and the heme-reductase part (NCP) to the right according to HMMER. The NCP Erg11 involved in ergosterol synthesis is the protein most like the FgCYP_NO_ (FGSG_01000), and a P450 mouse reductase (NCP) is similar to the FgNCP_NO_ involved in NO production. All these pairs except the iNOses have N-terminal TM regions predicted to be associated and inserted in the ER membrane by DeepLoc. All CYP-like proteins are predicted to be strongly Cam binding, and a few of the NCPs to the right are also predicted to be strongly CaM-binding. The rest of the NCPs are predicted to have cam-binding sites but are predicted to themselves be Cam-binding to a lesser degree (under the confident threshold level 1 ^50^ but might still have Cam binding possibility. For the complete analysis and protein sequences, see **Supplementary file SF6**. Our analysis show a remarkable similarity in domain structure and even distance between domains of the orthologous CYP-NCP pairs with the mouse iNOS. All CYP-NCP proteins contains the N-teminal part that is predicted to be an ER-anchor.

### Can FgCYP_NO_ also be involved in ergosterol biosynthesis?

The FgCYP_NO_ is well conserved in fungi but mainly annotated as a cytochrome P450 (CYP51) having a role in sterol synthesis, converting lanosterol to the next step in the sterols synthesis on the way to ergosterol in fungi and cholesterol in animals and other sterols in other eukaryotes ^51^. The final products, ergosterol or cholesterol, are necessary for endocytosis. Lack of the final product ergosterol in the membrane and accumulating lanosterol should inhibit endocytosis previously as shown in the literature ^52^. Thus, if the FgCYP_NO_ is also involved in ergosterol synthesis, endocytosis should be decreased in the *ΔFgCYP_NO_*strain ^53^. Indeed, endocytosis was shut off as the standard endocytosis probe FM4-64 was not taken up and quickly internalized (**Fig. 11A**) in the *ΔFgCYP_NO_*strain, indicating that the deleted heme protein (FgCYP_NO_) is likely involved in sterol production as many of its orthologues are, and it is consequently an **FgCYP_NO,ERG_** with at least these two functions.

**Figure 11.**
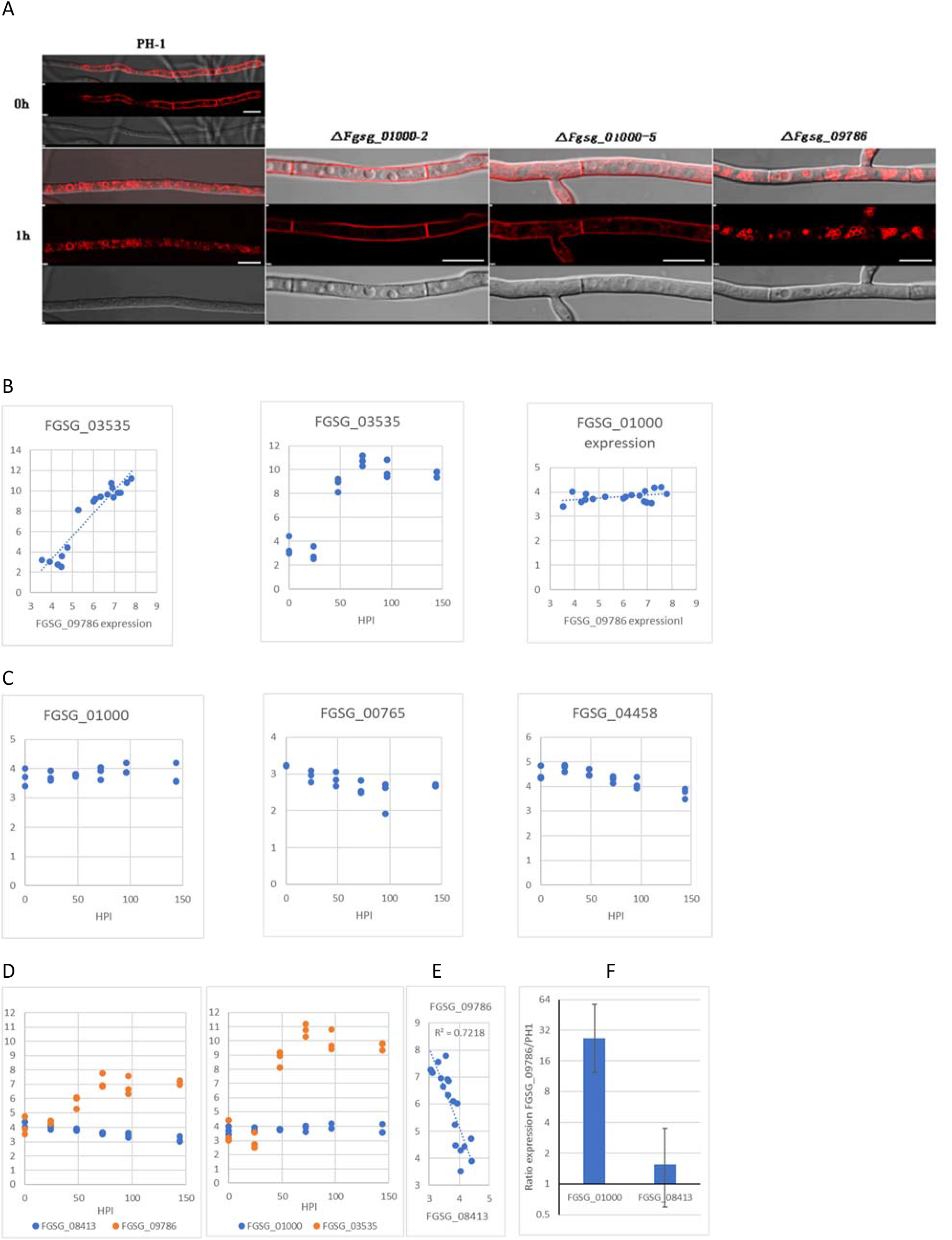
Effect of the *ΔFgCYP_NO,ERG_* on ergosterol biosynthesis and compensatory regulations when the NCP_NO,TRI,ERG_ is used to transfer electrons to the TRI4 heme protein (FGSG_FGSG_03535. (**A**) Staining endocytosis uptake using the endocytosis probe FM4-64 for 1h. Wt =PH1 takes up the stain by endocytosis, and the stain accumulates at the ER around the nucleus. *ΔFgCYP_NO,ERG_*(FGSG_01000) is completely blocked for endocytosis, and the stain stays at the plasma membrane in normal-looking hyphae, and FGSG *ΔFgNCP_NO,TRI,ERG_* (FGSG_09786) is not severely blocked in endocytosis. DIC image to show that FGSG_01000 appears normal in intracellular structures. Size bars 10μm. (**B-E**) Co-regulation and expression during the course of infection of the two NO-producing genes compared with the primary DON-producing genes in a large transcriptome dataset with transcription during plant infection (Experiment FG1 with 3 replicates for each of 6 timepoints and a total N=18) from a previous study ^54^. Expression axes are log2 expressions, and each dot represents data for the genes from one transcriptome. (**B**) Correlation between DON producing *FgCYP_TRI_* (FGSG_03535, Tri4) and *FgNCP_NO,TRI,ERG_* (FGSG_09786)(left). Expression of *FgCYP_TRI_* during the course of infection 0-144 hours post-infection (HPI) (middle). Correlation between DON producing *FgCYP_TRI_* (FGSG_03535, Tri4) and *FgCYP_NO,TRI,ERG_* (FGSG_09786)(right). (**C**) Expression of *FgCYP_NO,ERG_* (*FgNOD1* and *FgNOD2* 0-144 HPI showing NO upregulation of the genes linked to NO-production. (**D-E**) Regulation of *FgNCP_NO,TRI,ERG_*(FGSG_09786) compared to *FgNCP_ERG_* (FGSG_08413) and *FgCYP_TRI_*(FGSG_03535) and *FgCYP_NO,ERG_* (FGSG_01000) during the course of infection. (D) Expression of *FgNCP_NO,TRI,ERG_* increases during the course of infection (Left) as do *FgCYP_TRI_*right) while *FgNCP_ERG_* slightly decreases (left) and *FgCYP_NO,ERG_* does not change (right). (**E**) The negative correlation between FgNCP_NO,TRI_ and FgNCP_ERG_. (**F**) RT-qPCR test if *FgCYP_NO,ERG_* (FGSG_01000) and *FgNCP_ERG_* (FGSG_08413) are compensatory upregulated to aid ergosterol synthesis in the *ΔFgNCP_NO,TRI,ERG_* (FGSG_09786). Y-axis represents the ratio of expression of the respective genes in the *FgNCP_NO,TRI,ERG_*(FGSG_09786) mutant compared to the WT (PH1). Both genes potentially involved in ergosterol synthesis are upregulated, explaining why ergosterol-dependent endocytosis was expected in the *ΔFgNCP_NO,TRI,ERG_*. Three replicates were used, and error bars show 95% confidence intervals for the mean values. Thus, the P-value for the null hypothesis for the same average with non-overlapping error bars <<0.05.

### Regulation of the genes involved in ergosterol, NO and DON production during infection in published transcriptomes from *in planta* experiments

We have previously downloaded publicly available transcriptomic *in planta* data from plant infection experiments with *F. graminearum* as a pathogen ^54^. The first dataset from a time course of infection is interesting since the deletion of FgNCP_NO_ completely stopped DON production (**Fig. 6D**). From the literature, it is known that the *FGSG_03535* gene in the co-regulated trichothecene gene cluster making DON (the TRI cluster) is a P450 heme protein named TRI4 ^55^. It also localizes to the ER ^56^, could named as a FgCYP_TRI_ since it is a CYP necessary for making a reduction step in the DON synthesis. If FgNCP_NO_ is also a reductase delivering electrons to the FgCYP_TRI_ protein, it should be co-regulated with FgCYP_TRI_ in the *in-planta* data and upregulated during different phases of infection (Hours Post Infection, HPI). In plant infection transcriptome dataset, the FgCYP_TRI_ is strongly co-regulated with the FgNCP_NO_ during infection, and FgCYP_TRI_ is upregulated after 24HPI. However, the NO-producing FgCYP_NO_ is not co-regulated with FgNCP_NO_ during infection (**Fig. 11B**), indicating that FgNCP_NO_ should be named **FgNCP_NO,TRI_** since it is involved in reducing heme proteins needed for both NO formation as well as DON synthesis.

Sterol formation is essential and there are only two NCPs with the correct structure (FGSG_09786 and FGSG_08413) (**Fig. 3**), and we could not find any or a minimal effect on endocytosis when deleting FgNCP_NO,TRI_ (**Fig. 11A**), the other heme reductase (FGSG_08413) could be involved in ergosterol production, also delivering electrons to FgCYP_NO,ERG_. Consequently, both heme reductases should be able to complement each other for ergosterol synthesis. Therefore, the remaining heme reductase (FGSG_08413) could have a function as a reductase not involved in DON or NO production but only in ergosterol production, a **FgNCP_ERG_,** and the FgNCP_NO,TRI_ would then be a reductase involved in ergosterol production as well, a **FgNCP_NO,TRI,ERG_**. If this is the case, and since there is probably not any increased NO formed during the first 144HPI indicated by the absence of upregulation of the FgCYP_NO,ERG_, FgNOD1 and FgNOD2 (**Fig. 11C**), there should be an inverse relationship between FgNCP_NO,TRI,ERG_ and FgNCP_ERG_ expression during infection since ergosterol synthesis is necessary under aerobic conditions ^53^ and the FgNCP_NO,TRI,ERG_ get increasingly occupied with the FgCYP_TRI_ for producing DON. This inverse relationship is indeed the case (**Fig. 11D-E**), suggesting that both proteins are involved in ergosterol synthesis providing FgCYP_NO,ERG_ with electrons.

A way to directly test this conclusion is to investigate *FgCYP_NO,ERG_* and *FgNCP_NO,TRI,ERG_* expression in WT (PH1) and compare it with the expression of the same genes in the *ΔFgNCP_NO,TRI,ERG_* strain that does not make DON but is capable of limited endocytosis (**Fig. 11A**) that needs ergosterol to work. We made this comparison using RT-qPCR and can show that FgCYP_NO,ERG_ is strongly and significantly upregulated, and FgNCP_ERG_ is slightly, but not significantly upregulated in the *ΔFgNCP_NO,TRI,ERG_* compared to the WT(PH1), indicating that these genes are compensatory regulated in the mutant (**Fig. 11F**) explaining why the Δ*FgNCP_NO,TRI,ERG_* still show some endocytosis (**Fig. 11A**).

### FgNCP_NO,TRI,ERG_ can be complemented by an orthologues NCP from mouse

The FgNCP_NO,TRI,ERG_ protein is similarly organized in *F. graminearum* and mammals (**Fig. 10**) but annotated as involved in the sterol synthesis, we complemented the *FgNCP_NO,TRI,ERG_* with the heterologous gene from mouse (NM_008898). The mouse gene localized to ER as predicted and restored colony growth rate, but not conidia formation or the trichothecene mycotoxin DON’s production. However, the trichothecene-dependent pathogenicity on wheat coleoptiles was restored, possibly indicating that another *Fusarium graminearum* species complex trichothecene than DON ^57^ could have become the main product when electrons were delivered from the MouseNCP to the CYP_TRI_ (TRI4 gene) (**Supplementary Fig. S11**). Most importantly, the MouseNCP could restore the NO production (**Fig.12**), indicating that in mammals, the CYP and NCP involved in converting lanosterol in the cholesterol synthesis pathway should be able to produce NO also in mice.

**Figure 12.**
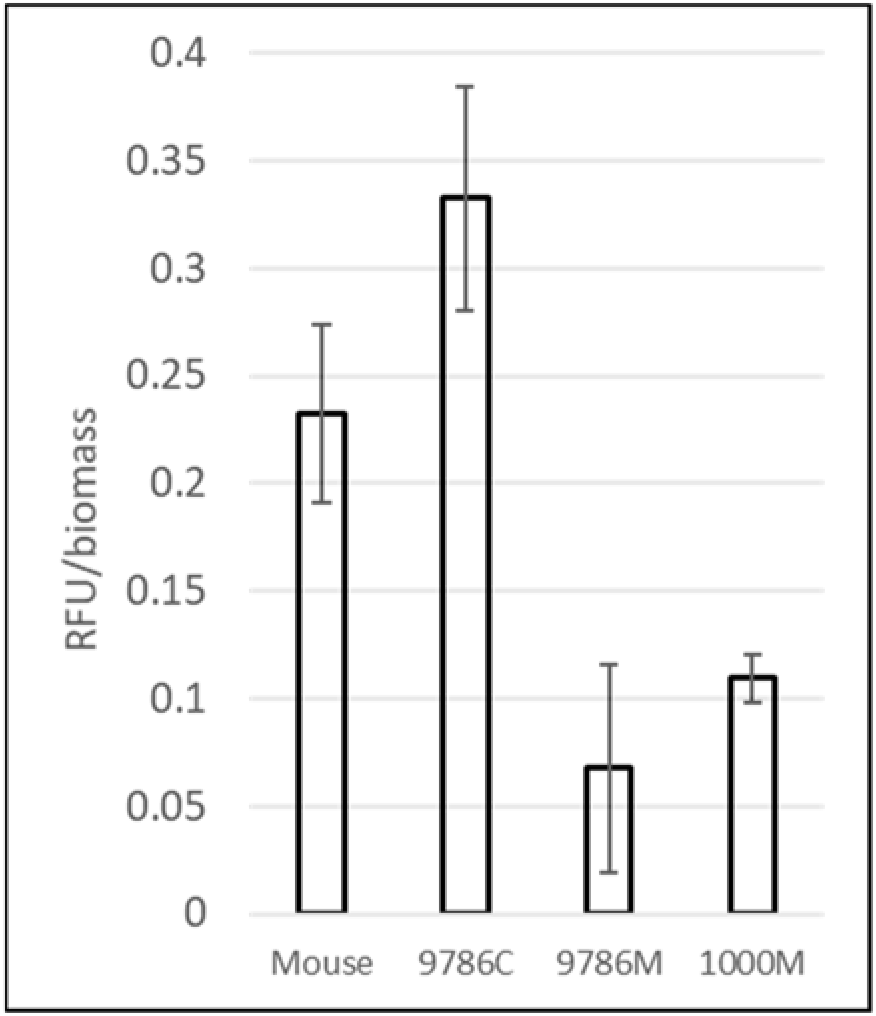
Complementing the *ΔNCP_NO,TRI,ERG_* (9786M) with the orthologue gene from mouse (Mouse) restores NO production to levels similar to the NCP_NO,TRI,ERG_ complemented by the native gene (9786C). The *ΔNCP_NO,TRI,ERG_* (9786M) and the *ΔCYP_NO,ERG_* (1000M) were used as negative controls. Three replicates were used, and error bars show 95% confidence intervals for the mean values. Thus, the P-value for the null hypothesis for the same average of bars with non-overlapping error bars <<0.05.

### Interactions between the FgCYP_NO,ERG_ and FgNCP_ERG_

Both proteins localize to the cytosolic side of ER according to our results and the literature for orthologous proteins in other organisms. At the ER they should interact to form NO or the ergosterol intermediate. We used the yeast 2 hybrid technique (Y2H) to express the proteins in yeast without their N-terminal membrane anchor and introns to detect eventual protein interactions but could not detect any physical interaction (data not shown). We then tried bimolecular fluorescence complementation (BiFC) since that can be used to detect if the proteins are close in vivo in *F. graminearum* but found no positive interaction (data not shown). For both techniques a negative result is no evidence for no interaction. We have thus not been able to confirm that these proteins interact closely and long enough to be recorded as positive with any of these techniques. As an electron transport is needed for the essential sterol synthesis, we must assume that they interact in some way, maybe for short times and at relatively long distances. According to the literature direct contact is not needed for efficient electron transport; on the contrary, that could even be counterproductive ^58,59^ (further discussed in the discussion below).

## Discussion

### Similarities and differences to NOS in mammals

There are apparent similarities between the ER-located NO-generating and lanosterol-reducing (**Fig. 9 and 10**) proteins and mammalian NOS. Calcium-dependent dimerisation is needed for NOS to be activated, at least for nNOS and eNOS ^8^. As we have demonstrated, NO production in *F. graminearum* is stimulated by MAMPs. This stimulation leads to increases in the concentration of cytosolic Ca^2+^.

The ER-located proteins involved in NO production in *F. graminearum* are predicted to be CaM-interacting (**Fig. 10**) and Ca^2+^ increases intracellularly induced by bacterial MAMPs (**Fig. 1C**) that triggers NO production (**Fig. 1A-B,2**). CaM changes in conformation most likely help bring the two NO generating proteins closer together on the ER membrane, activating NO production (**Fig. 13A**). The ER-located CYP_NO,ERG_ and NCP_NO, ERG,TRI_ reducing system is remarkably conserved from Protista to Mammals (**Fig. 9-10**).

**Figure 13.**
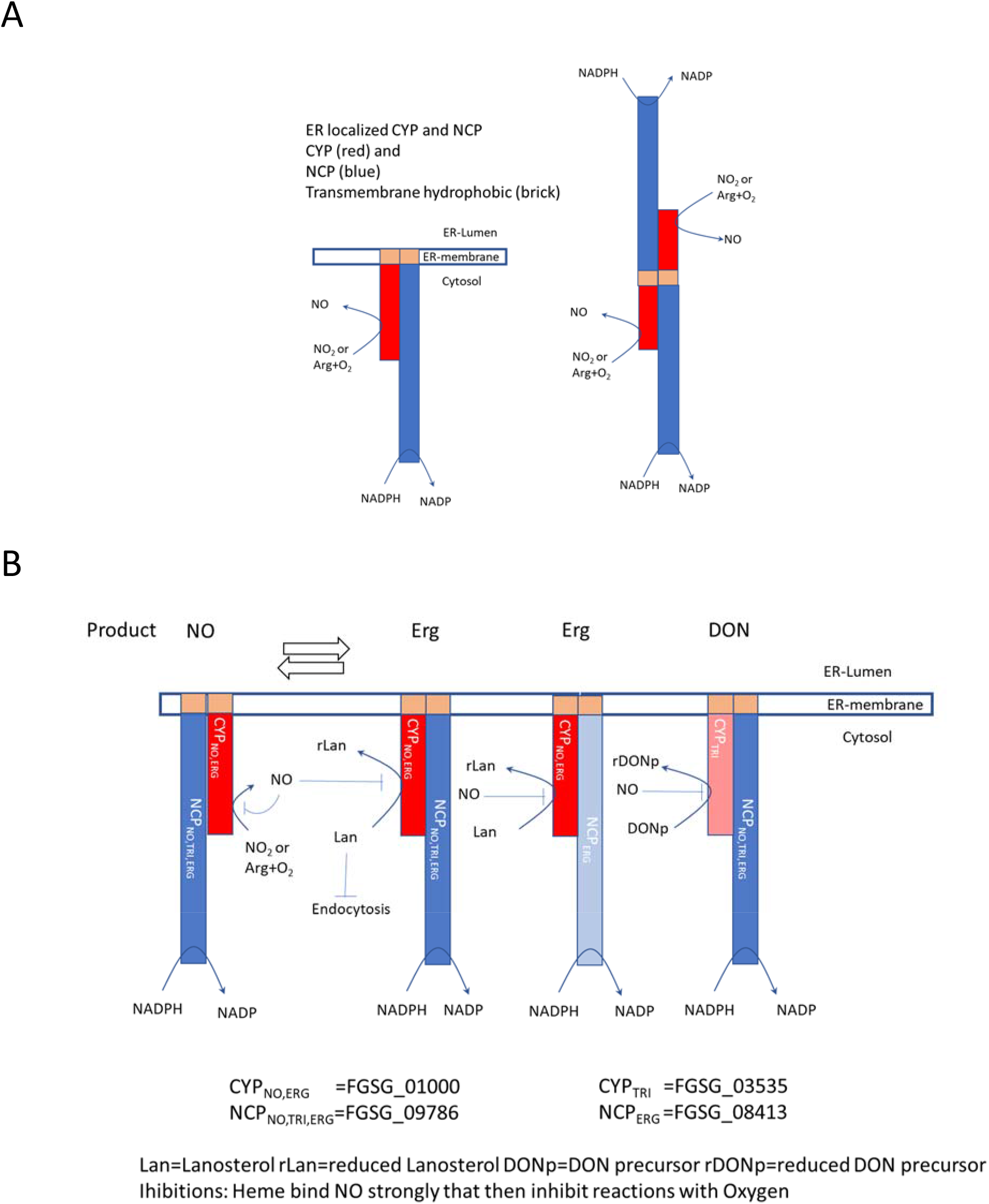
Conceptual models. (**A**) Model for the arrangement of domains in the ER-localized CYP_NO_ and NCP_NO_ implied in this study compared with the localization of domains in a NOS-dimer ^8^. (**B**) A conceptual model for the dynamic arrangements and re-arrangements of the CYPs and NCPs at the ER membrane needed for NO, ergosterol and DON production. Note that interactions needed for electron transfer between proteins do not need physical contact, only brief closeness (100nm seems to be enough) ^58,59,65,108^. The relative abundance of the proteins and more shapes more favourable to electron transfers brought about by conformational changes due to Ca^2+^ binding to CAM binding sites (**Fig. 10**) could favour such interactions at a distance, making it very flexible and quick to change the products’ proportions by gene expression and calcium signalling. The model, however, depicts the symbols for the proteins as if they need to be in direct contact. This study’s main proteins, FgCYP_NO,ERG_ and FgNCP_NO,TRI,ERG_, appear to have 2, respective 3, functions in *F. graminearum*. NO binds to FgCYP_NO,ERG_ heme, and self inhibits NO ^9^ and ergosterol formation at lanosterol ^109^ an effect known to reduce endocytosis^110,111^, and FgCYP_TRI_ since DON formation seems to be shut off by NO-donors ^29^.

Electron transfer proteins where electrons are transferred between domains in the same proteins seem to be made efficient by quantum tunnelling electrons between precisely spaced domains ^58,59^. Quantum tunnelling imposes severe constraints on essential proteins’ evolution and can explain the high conservation of the involved proteins. This need for conservation can also explain why there are double and triple functions for the involved proteins with different roles in combinations with different proteins. Functionally conserved NCPs producing NO is supported by our successful complementation of the *F. graminearum* gene by the mouse’s orthologue (**Fig. 12**).

Previously, in the literature it has been pointed out that the three classical NOS systems are not present quantitatively enough to generate NO in human vascular tissues, pointing towards additional NO generating systems also present in the microvasculature ^60^ and a p450 CYP is a good candidate for an additional system ^61–64^. The conserved CYP-NCP pair that are responsible for sterol synthesis is a good candidate for such a system, since the mouse NCP could successfully complement the fungal NCP for NO production. Electron tunnelling between from NCPs to the CYPs at the ER membrane could also explain the flexibility in the CYPs (FgCYP_NO,ERG_, FgCYP_ERG_, FgCYP_TRI_) and NCP (FgNCP_NO,TRI,ERG_, FgNCP_ERG_) used since there is no need for physical contact for electron transfer since electrons can tunnel between separate protein domains ^58,59^ even over as long distances as 100 nm in the case of heme proteins ^65^. Long distance might not even be energetically problematic. Rather the opposite as close contact is less efficient for tunnelling^66^. If physical protein-protein contact is not needed the relative concentration of the involved proteins at the ER membrane will decide what is produced and how much,. What is mainly needed is changes in transcription and translation of the different proteins and that can be changed quickly.

Electron transfer between proteins attached to a membrane without needing physical contact offers an elegant and quick mode for changing product output while always keeping necessary sterol synthesis going. Since all proteins are membrane-attached and their orientation should be strongly affected by the membrane potential, the protein approaches are more likely to be productive since the proteins will be correctly oriented in a way that can result in an electron transfer. In combination with calcium binding to the predicted calmodulin sites (**Fig. 10**), the proteins’ shape can also be changed further to even better promote electron transfer.

### Plant pathogenicity and NO generation

It has recently been shown that NO is formed in *F. graminearum* as a response to-plant molecules’ during pre-plant contacts in the rhizosphere bot not inside the plant. A plant-sensing receptor triggers downstream responses resulting in fungal NO-formation, and further downstream NO responses have been identified ^29^. Our results (**Fig. 10**) indicate that fungal FgCYP_NO,ERG_ needed for NO production appear to decrease when *in planta,* which would meet the challenge of not producing NO inside a plant since that can activate plant defences ^14,29^. Instead, *in planta,* the electron transfer from the FgNCP_NO,TRI,ERG_ takes part in DON production partly using the NO-generating enzymes (see above). The FgNCP_NO,TRI,ERG_ has also been well described enzymatically under its UniProt entry name I1RZE7_GIBZE as a protein having all the necessary activities to take part in both DON and ergosterol synthesis ^67^. DON is an inhibitor of translation ^68^, and the innate immune responses in Eukaryotes depend on the quick upregulation of defence proteins^69^. A shift from NO to DON production *in planta* (**Fig. 11B**) to attenuate the eventually turned-on plant defences could be an evolutionary adaptation of *F. graminearum*. In native North American grasses, *F. graminearum* seems to have evolved to colonize as an endophyte, disease with accompanying high DON production is not triggered ^70^. Production of large amounts of NO in planta would trigger the plant SA defences and recruit plant PR proteins, as would exposure of ergosterol to the plant. Limiting plant access to ergosterol, a known fungal MAMP, that plants react to should also avoid trigging plant defences ^71,72^. The changing roles of the proteins *in planta* could might indicate an evolutionary adaptation to an endophytic lifestyle, where plant defences stressing the fungus and counteracting DON responses from the fungus balance the interaction with the plant allowing the fungus to grow inside the plant without plant disease development. The higher levels of DON production produced in wheat and other Eurasian grasses might be caused by the fungus being mistaken for a pathogen by these grasses, making the fungus a *de facto* pathogen ^70^ responding with necrotrophic pathogenicity to the stresses imposed by the plant’s pathogen defence system..

### NO generation and essential sterol synthesis are intimately linked, and the system seems present in all eukaryotes

Our results unexpectedly revealed that NO generation and ergosterol synthesis are intimately connected. The genes we found that are responsible for nitric oxide generation have been well characterized as part of sterol synthesis. Sterol synthesis has even been well characterized previously in *F. graminearum* ^73^. There are 3 FgERG11/CYP51 genes responsible for critical sterol synthesis. The primary gene involved (FGSG_01000, CYP51B) is the one we now find is responsible for NO production. For other organisms (fungi, plants and animals), the genes catalysing this step in sterol synthesis are necessary under aerobic conditions ^53^ and the human gene HsCYP51A can replace the ScER11 counterpart ^74^. In *F. graminearum*, that step is catalyzed by 3 paralogues ^73^, where 2 of the proteins complement each other’s function for ergosterol synthesis, as we also found (**Fig. 11**). The orthologous mouse NCP can replace *FgNCP_NO,TRI,ERG_* suggesting that both mammalian genes (**Fig 12, Supplementary Fig. S11**) needed for that specific step in ergosterol synthesis are also involved in nitric oxide production as they are in *F. graminearum*.

Thus, it was sheer luck that we chose to investigate fungal NO production in *F. graminearum* since it has many paralogues of this essential gene and that these can stand in for each other regarding essential sterol synthesis, ^73^. Genes encoding proteins catalysing the same chemical reaction have been assigned different names in different organisms, although they have similar chemical functions. **Table 1** illustrates these names compared to the names used in this paper. Finally, FGSG_08786, the FgNCP_NO,TRI,ERG_ has been chemically characterized and called FgCPR and suggested to take part in DON-synthesis or ergosterol synthesis ^67^ as we also can confirm (**Fig. 6B**).

**Table 1.** Conversion table of gene names.

| ORF id F. Graminearum paralogues | Suggested name based on 1 chemical property and 2 cellular functions | Previously assigned chemical function | Names used in F. graminearum and (other filamentous) | Saccaromyces cereviseae | Mammals |
| --- | --- | --- | --- | --- | --- |
| FGSG_01000<sup>1</sup> | *FgCYP(NO,ERG)* | C14-α-demethylase | *Fgcyp51B*<sup>2</sup> | *Erg11* | *cyp51* |
| FGSG_04092<sup>1</sup> | - | C14-α-demethylase | *Fgcyp51A*<sup>2</sup> | - | - |
| FGSG_11024<sup>1</sup> | - | C14-α-demethylase | *FgCyp 51C*<sup>2</sup><br>(unique for *Fg*) | - | - |
| FGSG_03535 | *FgCYP(TRI)* | trichodiene oxygenase | *FgTri4*<sup>3</sup> | - | - |
| FGSG_09786<sup>1</sup> | *FgNCP(NO,TRI,ERG)* | NADPH-cytochrome P450 reductase | *FgCPR*<sup>4</sup> | YeastNCP1 | *P450R* |
| FGSG_08413<sup>1</sup> | *FgNCP(ERG)* | NADPH-cytochrome P450 reductase | - |  |  |

1. According to this study 2. References<sup>73,112</sup> 3. Reference<sup>113</sup> 4. Reference<sup>67</sup>

### Endocytosis and NO possible consequences

Heme interaction with O_2_ in the oxidative formation of NO from arginine is inhibited at high NO concentrations since NO binds heme better than O_2_ and blocks further NO production ^9^. Thus iNOSes are known to be self-inhibited ^8,9^. Since the heme in the FgCYP_NO,ERG_ is also used for reducing lanosterol in the ergosterols synthesis, NO should also halt ergosterol synthesis ^75^ at lanosterol, inhibiting endocytosis ^52,76,77^. Interestingly, a complete stop in endocytosis was previously observed by us in *Rhizoctonia solani* hyphae confronted with a bacterial biological control strain of *Burkholderia vietnamiensis* ^78–80^ while nitric oxide was produced ^78^. Those observations of NO formation inspired this study, together with the study on fungal innate immunity ^81^ in *F. graminearum* but it is first now with this work we understand why endocytosis became blocked.

A shutting down of endocytosis due to high NO concentration interacting with heme triggered by bacterial challenges is likely an ancient system protecting primitive eukaryotes from pathogenic bacteria utilizing the endocytosis mechanism to gain entry into the interior of the cells ^82^. This general effect infection/inflammation has on endocytosis should have broader consequences since, for example, LDL is taken up by endocytosis ^83^ and depends on CYP51 function ^84^. Since the heme-containing CYP51 (homologue to FgCYP_NO,ERG_) is likely self-inhibited by NO formed during inflammation in the same way as iNOSES this might explain inflammation-induced decreased LDL uptake and resulting health problems linked to chronic inflammation ^83^.

### NO and ergosterol as antioxidants

Both NO and membrane sterols are antioxidants that protect unsaturated membrane lipids from a runaway chain reaction of lipid peroxidation with the formation of lipid peroxide radicals, lipox ^85,86^. NO and membrane sterols might be the original antioxidants protecting membranes in early eukaryotic evolution when oxygen levels were low and the early eucaryotes were anaerobic. That could explain why they are intimately connected in their synthesis and appear to be present in basically all Eukaryotes. Lipid peroxidation is a severe problem for all aerobic life since life originated anaerobically and has never fully adapted to high oxygen levels. Different protective measures have evolved, but a fundamental redesign of the membrane components has not evolved outside bacteria. Eukaryotes use unsaturated membrane lipids sensitive to oxidation to achieve necessary membrane fluidity. Some bacteria can instead produce branched saturated fatty and use them in for the same purpose ^87^, offering better protection against oxidation.

### Nitrogen limitations and NO production, possible consequences

Fungi are generally nitrogen limited in nature. It also applies to *F. graminearum,* which is nitrogen and sometimes carbon-limited inside the plant and dependent on autophagy for mobilizing carbon and nitrogen from stores ^88^. During the N-limited conditions both fungi and plants generally live under, they should have evolved mechanisms to save nitrogen. The NOS formation of NO from arginine is less problematic if the NO_2_ or even NO_3_ formed in that process could also be used to generate NO while making NH_4_^-^ through assimilatory reductions so that arginine can be replenished. However, our result with the spiking experiments show that the effective concentrations of NO should be higher after spiking with arginine since the rate of the increase indicates the effective NO concentration (**Fig. 2C**). This could mean that the NOS-like arginine oxidation is the primary source of NO and the reductive pathway through nitrite reductase is a secondary system needed for prolonged NO production and recycling of N. Thus assimilatory nitrite/nitrate reduction in both plants and fungi could have evolved mainly as a mechanism to save N and recharge the cells with arginine since a large amount of arginine is needed to make NO when challenged by bacteria ^89^. That could also explain why assimilatory nitrate/nitrite reduction is under the control of the GATA factor AreA in fungi that regulates genes needed to use secondary nitrogen sources also regulates secondary metabolite production since those metabolites are needed to interact with other organisms ^90^.

### Conclusion and overall conceptual model for what happens at the ER

Our results show that the evolutionary ancient and very conserved sterol synthesis system in Eukaryotic cells is involved in NO production. In *F. graminearum,* part of the system has a role in producing the mycotoxin deoxynivalenol (DON) that inhibits translation ^68^ in the plant host cell and should lower plant immune responses. *In vitro,* the fungus makes ergosterol and, but if severely challenged with bacteria NO forms, likely stopping ergosterol synthesis at lanosterol, shutting down endocytosis and preparing the fungus for bacterial defences. If attacked by plant defences *in planta,* the part of the system shift again but to the production of DON to inhibit plant translation to attenuate plant defences (**Fig. 13B**) ^68^. Our result further imply that the cholesterol synthesis system in mammals also generates NO when receiving inflammation signals which should have negative effects on LDL uptake and metabolism as well as increased blood pressure linked to chronic inflammation ^83^.

## Materials and Methods

### Maintenance and growth of *Fusarium graminearum* for plate-reader assays

The wild type strain of *Gibberella zeae* ((Schwein) Petch) (anamorph *Fusarium graminearum* (Schwabe) PH1 (NRRL31084) was originally obtained from the Agricultural Research Service Culture Collection, National Center for Agricultural Utilization Research, Peoria, IL, USA. The wild-type and mutant strains were stored as spores in 10% glycerol at −70°C. Cultures were maintained on minimal Defined Fusarium Medium (DFM) at 21°C, in the dark and shaking at 150 rpm if necessary. The strains were cultured in YPG liquid medium to get fungal biomass for DNA extraction ^91^. Spore production was induced by fungal growth in liquid RA medium (succinic acid, 50 g L-1; sodium nitrate, 12.1 g L-1; glucose, 1 g L-1 and 20 mL L-1 50X Vogel’s salts without nitrogen or carbon source) for 3 days. The spore suspension was filtered through Miracloth, washed, and centrifuged twice at 5000 x g for 15 min.

### NO elicitation from *F. graminearum* using purified MAMPs in a plate reader

NO production was measured using a 96 well plate assay with various potential elicitors, including MAMPs. In each well, 2000 spores were cultured overnight in 100 µl of DFM (diluted 1:50) at 21°C in an incubator without light. The growth medium was removed from the overnight culture to produce a thin mycelial mat (network) attached to the bottom of the wells, and sterile Milli-Q water was added for overnight equilibration (starvation). The water was removed from the equilibrated culture and replaced by 100µl elicitor, supplemented with 50ng diaminofluorescein-FM diacetate (DAF-FM DA) (CAS: 254109-22-3) (D23844; Invitrogen, CA, USA) to detect NO production. The elicitors used were 0.01% w/v glucose; 0.01% w/v sucrose; 0.01% w/v glutamate; 0.01% w/v arginine; 0.01% w/v potassium nitrate; 0.01% w/v potassium nitrite; sterile Milli-Q water; 100 ng mL-1 full-length flagellin (FFLG) (containing 0.01% sucrose; tlrl-pstfla; Invivogen, CA, USA); 50 µg mL-1 lipooligosaccharides (LOS; LPS without the O-antigen), ^92,93^; 50 µg mL-1 peptidoglycan (PGN) ^94^; 100 nM flagellin 22 (Peptron Inc., South Korea); 100 nM elongation factor 18 (Peptron Inc., South Korea). We chose to use the 22 amino acid peptide from flagellin (FLG22) and the 18 amino acid from elongation factor Tu (EF-Tu; EF18), because they are known to be the epitopes that elicit innate immunity in plants ^6^. Fluorescence per well was measured in a 4×4 matrix (1400 gain) using a FLUOstar OPTIMA plate reader (BMG Labtech, Ortenberg, Germany) with emission filters at 485 nm and an excitation filter at 520 nm. Measurements were taken every 30 minutes for the duration of the experiment. The basal fluorescence reading (T0) at the start of the experiment was used as the control unless stated otherwise. The accumulation of NO as histograms is reported with readings at 10hr (T10) post elicitation or as graphs of accumulated fluorescence from produced NO. FFLG containing 0.01% sucrose was the most consistent inducer of NO and was used as a positive control in all the subsequent experiments. The germination time of the spores was challenging to replicate between experiments even if the culture producing the spores was grown similarly. Thus, the mycelial mats could be of different densities between the plates. All experiment treatments with controls (N=3 or more) and replicates (N=3 or more) were accommodated on one 96-welled healthy plate to compensate for that. In addition, the plates were first investigated for uniformity of the mycelial well development in all wells due to eventual pipetting handling mistakes (it should be the same absorbance from the fungal mat in all wells at the start of the experiment). The same was done for the other plate-reader-based assays below.

### NOS inhibition assays

The assays were performed as described above, but before elicitation, the culture was exposed to the inhibitors for 2 hours. Inhibitors used was [10mM L-NNA, NG-nitro-L-Arginine;L-N^G^-Nitroarginine (80220, Cayman Chemicals, MI, USA), 100µM W7, N-(6-Aminohexyl)-5-chloro-1-naphthalenesulfonamide hydrochloride (A3281, Sigma, MO, USA); 500nM ZSTK474 (S1072, Selleckchem, TX, USA); 100nM Wortmannin (W1628, Sigma, MO, USA); 25nM BpV(HOpic) (203701, Calbiochem-Millipore, MA, USA)] and elicitation was done with flagellin (containing 0.01% sucrose). Calcium ions in the flagellin solution were chelated with EGTA (E3889, Sigma, MO, USA) before elicitation. After 10 hours post elicitation, the NO accumulated level was compared, using FFLG-untreated cultures as negative controls.

### NOS spike assays

*F. graminearum* was cultured and elicited with flagellin (FFLG) as described above. Post elicitation, the chemical of choice [Arginine (A8094); Potassium Nitrate (P6030); Potassium Nitrite (60417) from Sigma] was added to a final concentration of 5mM per well. The increase or decrease in the production of NO was monitored. For sequential spiking assays with arginine and potassium nitrite, the first substrate was added at 4 hours and the second substrate 2 hours later, (i.e. at 6 hours).

### Procedure to find a putative FgNCP_NO_

This procedure is described in detail (**Supplementary File SF2)** and outlined here. A mouse iNOS was looked up at NCBI. The protein domains were identified using HMMER, and the NCP-part of the protein was used for a protein blast (NCBI) against the *F. graminearum* PH1 predicted proteins. Two good hits (FGSG_09786 and FGSG_08413) were found for putative NCP_NO_ proteins. If any of these produce NO, the NO producing gene is likely co-regulated with a nitric oxide dioxygenase (NOD) to protect against too much NO formed. Two putative NODs were found in *F. graminearum* PH1 (**Supplementary File SF2**). A transcriptome dataset for *F. graminearum* with complete transcriptomes obtained for many different bacterial MAMPs exposure conditions was used to investigate for co-regulation with the identified putative NODs to indicate which of the two putative NCP_NO_ proteins are most likely an NCP_NO_. This dataset was previously partly published ^81^ and is now available in full (**Supplementary Data1** https://figshare.com/s/4a02adb12cb48dba6d7a (reserved DOI: 10.6084/m9.figshare.12361823)). The gene FGSG_09786 but not FGSG_08413 was co-regulated with both NODs (**Fig.3 and Supplementary File SF4**) and is most likely the FgNCP_NO_ taking part in NO production.

### Procedure to find a putative FgCYP_NO_

When the NCP responsible for NO production (FGSG_09786) had been knocked out, found to be responsible for NO production (an NCP_NO_) and localized to the ER, it was possible to look for a heme-containing FgCYP_NO_ that can work together with NCP_NO_ at the ER membrane. The genome of *F. graminearum* PH1 contains numerous genes annotated as coding for heme-binding proteins. To get a complete list, we searched the whole genome for heme-domain-containing proteins using the Batch Conserved Domain search tool to identify all occurrence of putative domains (https://www.ncbi.nlm.nih.gov/Structure/bwrpsb/bwrpsb.cgi). These search results (**Supplementary Data2** https://figshare.com/s/a853e4ff951355eb0228 (reserved DOI: 10.6084/m9.figshare.12386501)) was then searched for proteins predicted to be heme-binding, and these proteins were listed. All heme proteins were then sorted after their co-regulation with the FgNCP_NO_ (FGSG_09786) in the transcriptome data set for *F. graminearum* exposed to bacterial MAMPs (**Supplementary File SF5**). FGSG_07925 was found to be most correlated with FgNCP_NO_ but was not predicted to localize to ER or had a deletion and complementation phenotype that could be expected for a true FgCYP_NO_ (**Fig. S6 and S7**). However, the ER localized FgNCPNO was not correctly predicted by PSORTII (**Table S5**) but correctly predicted to localize to the ER membrane by DeepLoc. Thus, we used DeepLoc to predict the protein localization of the most correlated proteins predicted to be heme-binding (**Supplementary File SF5**) and found that FGSG_01000 is the most likely FgCYP_NO_. This procedure is described in more detail in the supplementary file (**Supplementary File SF6**).

### Knockout and complementation of FgNCP_NO_ (FGSG_09786) and FgCYP_NO_ (FGSG_01000)

A 932-bp fragment upstream of FGSG_09786 and an 1176-bp fragment downstream of FGSG_09786 were amplified with specific primers to replace the FgNCP_NO_ (FGSG_09786) gene (**Table S14**). FGSG_09786 replacement constructs were finally generated by a split-marker approach ^95^. Subsequently, the resulting constructs were transformed into protoplasts of the wild-type strain PH-1. Hygromycin-resistant transformants were screened by PCR using two pairs of primers (**Table S14**). Deletion of FGSG_09786 was further verified by Southern blot with the Digoxigenin High Prime DNA Labeling and Detection Starter Kit I (Roche, Mannheim, Germany). Using a similar strategy, FgCYP_NO_ (FGSG_01000), two FgNCP_NO_ (FGSG_00765, FGSG_04458) and the first suspected FgCYP_NO_ (FGSG_07925) deletion mutants were constructed. All primers used in this study are listed in **Table S14**. At least two positive transformants for every gene were used for phenotypic analyses. The primers FGSG_09786CF and FGSG_09786CR (**Table S14**) were used to amplify the full-length sequence of the FGSG_09786 gene and its native promoter from the WT genomic DNA to generate a pFGSG_09786-GFP fusion vector. It was then cloned into a pKNTG vector using the One Step Cloning Kit (Vazyme Biotech Co., Ltd, China). The same method was used to construct pFGSG_00765-GFP and pFGSG_04458-GFP and pFGSG_07925-GFP vectors using their respective primers. The primers Mouse-09786-F and Mouse-09786-R (**Table S14**) were used to amplify the cDNA sequence of the NM_008898 gene, and the primers FGSG_09786 promoter-F and FGSG_09786 promoter-R were used to amplify the promoter of the FGSG_09786 gene to generate a heterologous complementary vector from the mouse genome. Subsequently, the two fragments were fused and cloned into a pKNTG vector using the One Step Cloning Kit (Vazyme Biotech Co., Ltd, China).

### Preparation of OMVs as MAMPs source for NO production

Outer membrane vesicles carrying MAMPs are shed by Gram-negative bacteria ^96,97^. OMVs were harvested in the following way: An inoculum of *Escherichia coli* DH5α was inoculated to 10 ml 1/10 DFM medium ^81^ in a 50ml in a sterilized centrifuge tube and incubated overnight at 28°C. The bacterial cell culture was harvested by centrifugation at 4200g for 5 minutes. The supernatant was removed, and the pellet washed by being resuspended in 10 ml sterile MilliQ water. This washing was repeated twice. Finally, the pellet was resuspended in 1 ml Milli Q water and incubated at 28°C for 1 hour to allow OMVs to form. After incubation, the suspension was harvested for OMVs by sterile filtration using a sterile 0.2 μm pore size filter. The resulting liquid containing OMVs was used for MAMP treatments.

### NO measurements using mycelial plugs

Milli Q water agar ^81^ was poured into a sterile Omnitray (Nunc), which is in principle a microtiter plate with one 8×12 cm big rectangular well ^98^. Fungal cultures of strains to be tested were grown on 1/10 DFM medium in 9 cm diameter Petri dishes containing 10 ml DFM agar medium (Ipcho *et al.*, 2016). When fungal mycelium was tested for NO production, 5 mm diameter plugs were cut away from the edge of the mycelium to avoid getting DFM nutrients interfering with test ^99^. A DAF-FM DA (ThermoFisher, D23844) staining solution was prepared as 2 μl/ml of a stock solution (2 μg/μl in DMSO) was added to the bacterial MAMPs suspension of OMVs or Milli Q water, giving a final concentration of 4 μg/ml DAF-FM DA. Droplets (10 μl) of the DAF-FM DA staining solution with or without bacterial MAMPs were added on top of the water agar in the Omnitray to perform the assay. Agar plugs were immediately put mycelium down on top of the droplets, one agar plug per droplet. In each Omnitray, there was space for 4 rows of 4 replicate plugs. *F. graminearum* PH1 was always used as positive and negative control taking up 2 rows to be compared with mutant or complement strains with or without bacterial MAMPs in the remaining two rows.

The amount of fluorescence from DAF-FM resulting from its reaction with NO was recorded for the whole plate automatically (using the GFP-filter of the instrument) at roughly 0.5 h intervals. Recordings were made through the plate’s bottom, and a lid was kept on to avoid the plate’s drying. The plate reader was set up to produce one big image for the whole plate that could be exported as a TEXT image and imported into the freeware ImageJ to produce a TIFF Z-stack that can be measured for the fluorescence development over time in any defined area ^98,100^. The fluorescence increase from each agar plug’s hyphae could then be analyzed and compared. For the first hours, the increase in fluorescence was linear, indicating a linear accumulation of fluorescent DAF-FM products due to a constant NO production rate. The curves’ slope was used as relative measures of the NO formation rates to compare MAMPs treatments with controls. The slopes for the fluorescence MAMPs/Water increase over time were used to compare the strains’ relative NO response to the MAMP treatments.

### Subcellular localization of NO production and measurements of NO production over time using confocal microscopy using the NO probe DAF-FM DA

NO was produced by *F. graminearum* PH1 mycelium also without MAMPs addition only a bit slower. DAF-FM DA was added directly to a microscope slide to localise the intracellular sites of NO production. Preliminary experiments showed that colour development was rapid. That was good since fluorescein derivates like the resulting DAF-FM will quickly leak out or move to the vacuole as typical for fluorescein. Therefore, we tried to minimize the time from adding DAF-FM DA to microscopy to get as good localization as possible while still getting a good signal. Thus we picked fresh mycelium from a colony grown in a CM agar medium and transferred that to a microscope slide, and added 10 μl water containing 2 μl DAF-FM DA solution to a final concentration of 4 μg/ml and <u>immediately</u> observed for NO production localization under a Nikon A1 confocal microscope for subcellular localization in NO production. Since we knew that FgNCP_NO_ responsible for NO production localizes to the ER (**Fig. 5B**), we used an *F. graminearum* strain carrying a mCherry tagged ER-marker protein (ER-mCherry-HDEL) ^101^ to confirm that NO is formed at the ER where the NO-forming proteins are localized (**Fig. 5C**).

Looking at the microscope images, it was evident that there were specks of autofluorescence that could interfere with the DAF-FM signal. We chose to use the confocal microscope as a fluorimeter to avoid these problems and get an alternative method for measuring NO formation from FgNCPNO and FgCYPNO. Slides were prepared as above. Analysis method: A time series of images of the mycelium was recorded using a 10X objective for the GFP signal and the DIC signal. First, Import the ND2 time series Z-stack into ImageJ as separate Tiff images, then separate GFP images and DIC images into folders and rename images so the number order index comes first in the name, and import these image folders as separate 16-bit TIFF black and white images into the freeware ImageJ (NIH, https://imagej.nih.gov/ij) to make two TIFF Z stacks. Define a measured area (Region of Interest, ROI) in the DIC image that does not contain air bubbles but relatively many hyphae. Import the ROI to the fluorescence image and check that there are no autofluorescence specs (dots that do not fade with time) in this ROI. Do the same for an area without hyphae as close to the previous area as possible to get a background control area. Measure and record the Z-profile for the fluorescence in the measured area and the control area. Use the edge filter to transform the DIC image to an image where the average light intensity is proportional to the biomass ^102^ and measure and record this for the first image in the sequence (there is no change over time for the DIC image). To maximize the analysis’s sensitivity, we calculated the area under the fitted decay curve corrected for the biomass measured from the DIC images as indicated above and used that as a relative measure for the NO formation/biomass for the PH1, FgNCP_NO_ and FgCYP_NO_ (**Fig. 7A**).

Measurements of the NO formation of NO formation in the NCP-complements using the native FgNCP_NO,TRI,ERG_-GFP, or a Mouse-GFP orthologue created new problems. Both the DAF-FM signal and the GFP signal are green. However, the pH-sensitive GFP signal fades very slowly and reproducibly in a buffered environment (pH7) since blue light damage to membranes will not affect the fluorescence by pH changes due to membrane leakages. Under similar conditions, the green DAF-FM signal fades entirely within minutes of observation (**Supplementary Fig. S11 B**). This complete fading of the DAF-FM signal makes it possible to reconstruct both the DAF-FM signal and the GFP signal at the start of the time sequence and show these as full images (**Supplementary Fig. S11 A**). We made these new measurements directly on agar slide cultures, making it easier to avoid air bubbles. The relative specific activity (NO signal per protein GFP signal) of the two NCPs could thus be compared (**Supplementary Fig. S11 C**).

### Colony growth, conidiation, sexual reproduction, pathogenicity and DON production phenotypes

Infection assays on flowering wheat heads and coleoptiles of cultivar XiaoYan 22 were conducted as described previously ^103,104^. For assaying DON production, fungal cultures were grown for 7 days at 25°C in liquid trichothecene biosynthesis (LTB) medium ^105^. Subsequently, a competitive enzyme-linked immunosorbent assay (ELISA)-based DON detection plate kit (Beacon Analytical Systems, Saco, ME, USA) was used to measure DON production. Growth rates were measured on complete medium (CM) plates and conidiation assayed in liquid carboxymethyl cellulose (CMC) medium as described previously ^106^. The formation of perithecia was tested on carrot agar medium ^107^.

### Endocytosis assay using FM4-64

Agar blocks (5 mm in diameter) were transferred from CM solid medium to 50ml centrifuge tubes containing CM liquid medium. After 24h shaking, 10 μl of the fungal hyphae suspension was mixed with FM4-64 to a final concentration of 4 μg/ml. Endocytosis of the stain was observed using a Nikon A1 confocal microscope using settings for TRITC (**Fig. 11A**). Endocytosis was estimated 1h after stain addition when most of the FM4-64 stain had been endocytosed and distributed to intracellular organelles if endocytosis is not blocked.

### Comparing FgCYP_NO_ and FgNCP_NO_ like proteins in eukaryotes, describing the domain structure and predicting CaM binding

These comparisons are described in detail (**Supplementary File SF7 and SF8)** and outlined here. First, NCBI was blast-searched for all predicted proteins in Eukaryotes, similar to FgNCP_NO_. From this result, a set of organisms were selected that had good hits for the protein’s whole length. Representative species were chosen taxonomically closely related to *F. graminearum* and progressively further away as long as good orthologues was found. The best hits were chosen, and phylogenetic trees for both proteins were generated (**Fig. 9**). DeepLoc was in addition used to check that all similar proteins are predicted to localize to ER and CaMELS (CalModulin intEraction Learning System) https://camels.pythonanywhere.com/) was used to find likely CaM binding sites, while HMMER was used to predict domain structure and membrane attachement for all these proteins (**Supplementary File SF7 and SF8).** All proteins were found to attach to the the ER membrane with their N-terminal, have similar domains, domain order, very similar domain sizes, and even distances between domains (**Fig. 10**).

### Statistics

MS Excel was used to calculate the statistics presented in the figures. We have, were relevant, chosen to use 95% confidence interval bars (SEM times 1.96) instead of standard t-tests to compare with controls since it simplifies for the reader and is more stringent. Thus, bars with non-overlapping 95% confidence intervals are significantly different (P-value for null hypothesis < 0.05).

## Authors contributions divided per authors

**WZ** Deletion and complementation work on the heme reductase the NODs and the first Heme protein we deleted and phenotype analysis. Manuscript writing and correction

**HL** Deletion and complementation of Heme protein and phenotype analysis including of NO measurements for Confocal microscopy. Manuscript writing and correction

**WF** Complementation of the heme reductase gene with a mouse heme reductase and phenotype analysis including growth, conidiation and pathogenicity. Manuscript correction.

**SI** MAMPS stimulation and NO measurements including inhibitor analysis, methods development for carrying out these analyses and experiments and performed the transcriptomic experiments, Manuscript writing and correction.

**RH** NO measurements of effect of HemeR and NOD deletions. Manuscript correction.

**BOH** Preparation of transcriptomic data for co-regulation analysis. Manuscript correction.

**GD** Acquisition of financial support for final research. Manuscript correction.

**ZW** Acquisition of financial support for final research. Manuscript correction.

**MAN** Initial idea, discussions, and preparation of MAMPs for testing. Acquisition of financial support for initial research. Manuscript correction.

**SO** Initial idea. Acquisition of financial support for initial research. Hypotheses generation from data and from literature as well as from bioinformatics, co-regulation analysis, phylogenetic and structure analysis, overall responsible for driving the work forward and methods development. Suggesting confirmatory analyses and experiments for additional functions. Main responsible for manuscript writing and coordination of the manuscript writing and corrections.

## Acknowledgements

Thanks to Professor Peter Stougaard, Department of Environmental Science. Environmental Microbiology and Circular Resource Flow, Aalborg University, Denmark, for the initial discussions at the start of the work. The Minnesota Supercomputing Institute is kindly acknowledged for computing resources and support for producing the transcriptomic data for the fungus interacting with bacterial MAMPs ^81^ that was essential for the study. We also thank The Velux Foundation (Denmark), which financed the initial Copenhagen part of the research. We want to thank Dr Qiurong Xie from Fujian University of Traditional Chinese Medicine for providing us with the cDNA for the mouse gene.

## Data availability statement

All data used for this study are available as images or in supplemental files.

## Supplementary material

**Figure S1.**
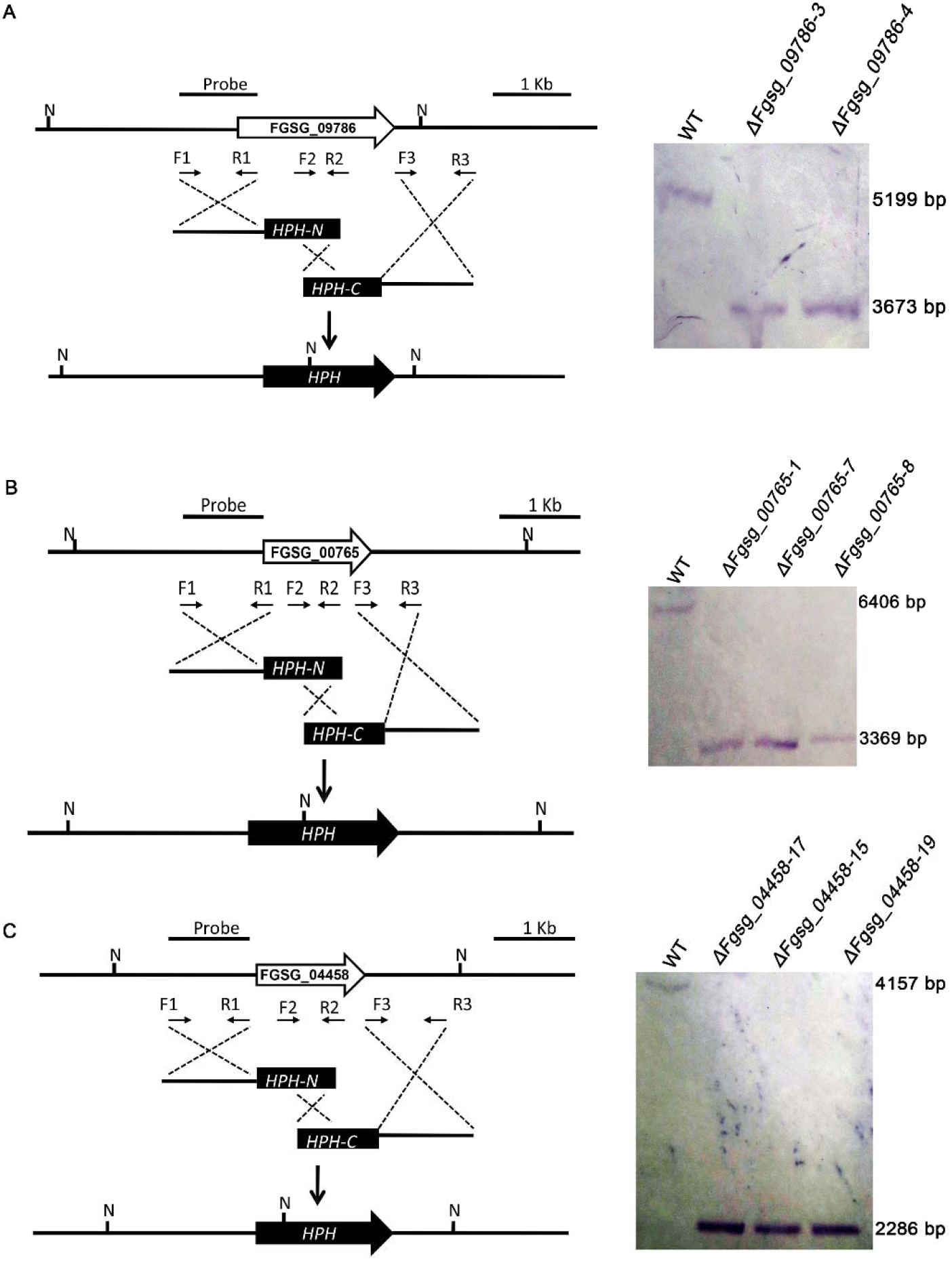
Targeted gene-replacement construct and Southern blot assays. Each targeted gene-replacement construct is shown in the left panel. The primer pairs F1 and R1, F3 and R3 were used to generate the gene replacement constructs. Primers F2 and R2 were used for mutant screening and identification. (A) Targeted deletion of FGSG_09786. Nde I-digested genomic DNAs showed a 5.20 kb band in the WT and a 3.67 kb band in the mutants. (B) Targeted deletion of FGSG_00765. Nde I-digested genomic DNAs showed a 6.41 kb band in the WT and a 3.37 kb band in the mutants. (C) Targeted gene deletion of FGSG_04458. Nde I-digested genomic DNAs showed a 4.16 kb band in the WT and a 2.29 kb band in the mutants.

**Figure S2.**
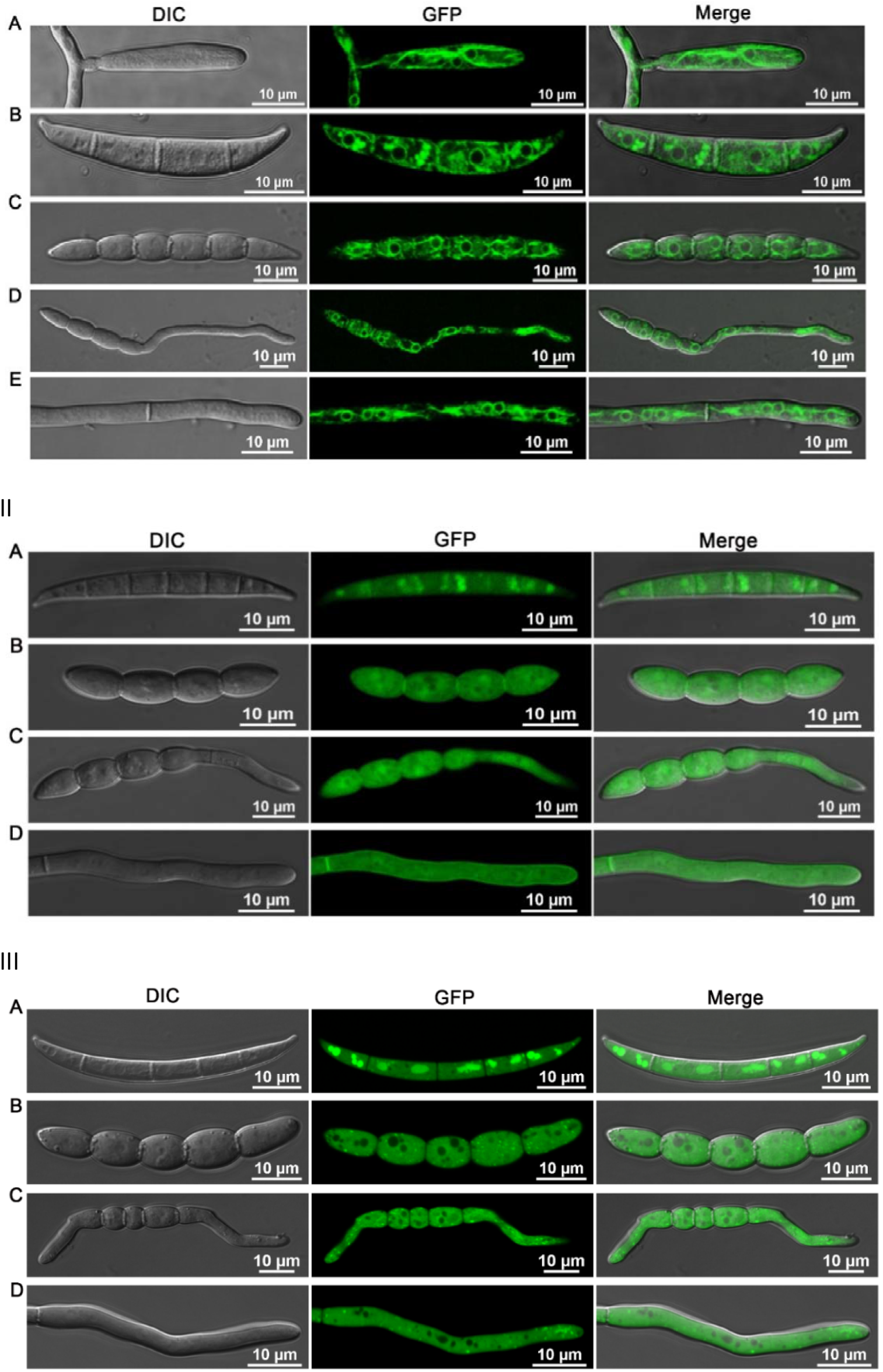
Localization of complements labelled with GFP. (I) FgNCP_NO_ (FGSG_09786) localizes to ER like structures. (II) FgNOD1 (FGSG_00765) principally localizes to the cytoplasm. It also is observed at the septum between cellular compartments in growing hypha. (III) FgNOD2 (FGSG_04458) localizes to the cytoplasm and small dot-like structures in hyphae and germinated conidia. The clustered fluorescence in conidia from FgNOD1 and FgNOD2 mutants may be GFP proteins undergoing degradation at the vacuoles. (A) Conidia (B) Conidia just before germination (C) Germinated conidia (D) Hyphae.

**Figure S3.**
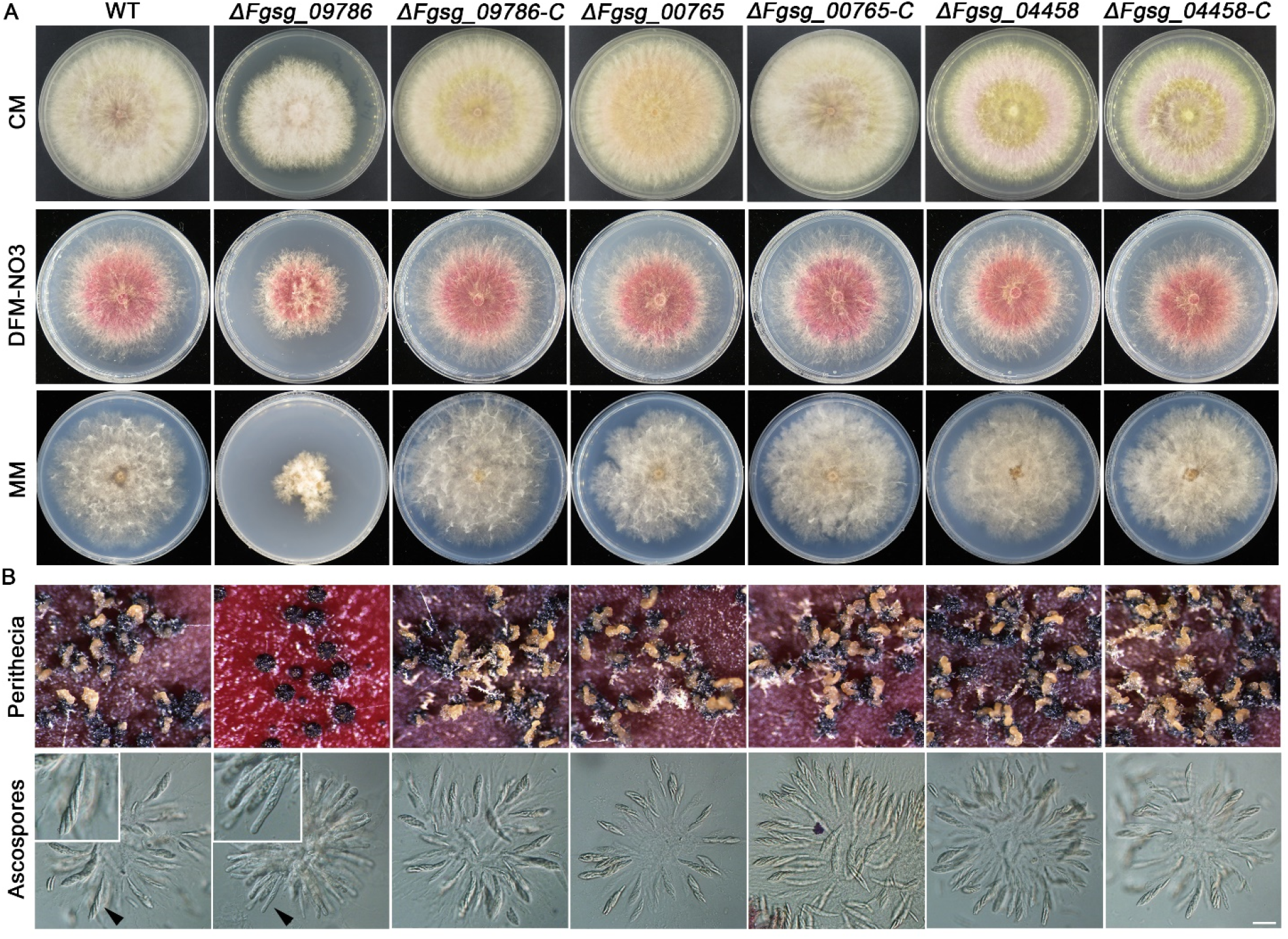
(A) Growth of WT (PH-1), mutant and complementation strains on CM, DFM-NO_3_ containing NO*_3_* as sole nitrogen source and MM medium. FgNCP_NO_ (FGSG_09786) is required for the hyphal growth in CM, DFM-NO3 and MM medium. (B) Sexual reproduction of WT, mutants and complementation strains. FgNCP_NO_ (FGSG_09786) is essential for forming normal ascospores and their ejection from the perithecia (yellow masses). The black arrowhead indicates ascus zoomed in and shown in the top left corner showing a changed morphology of asci. It is mainly the deletion of FGSG_09786 that show phenotypic colony effects and effects on sexual spore formation

**Figure S4.**
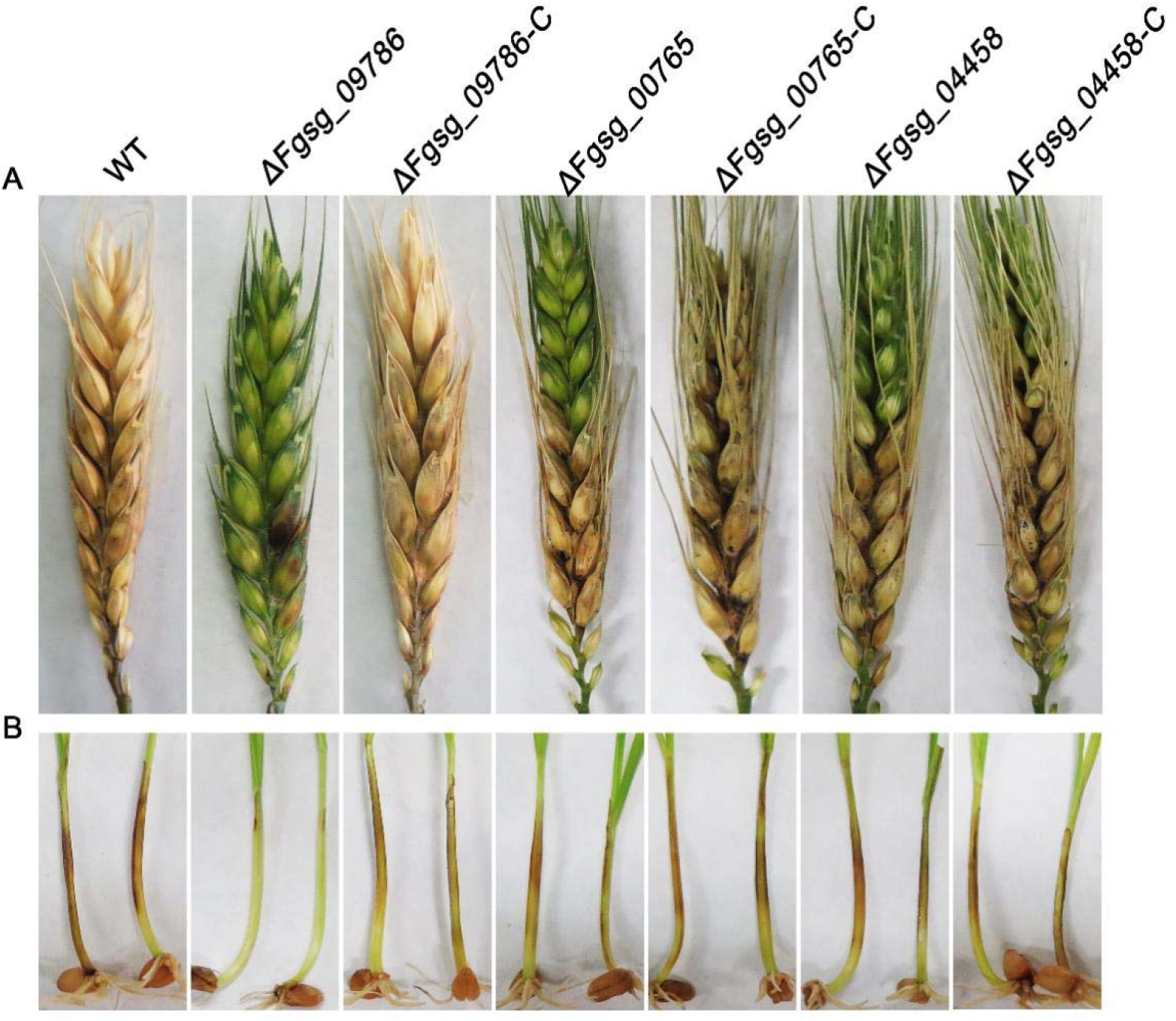
Pathogenicity of each strain on (A) Wheat heads and (B) seedlings. *ΔFgsg_09786* showed reduced virulence in both assays. Loss of chlorophyll green colour and browning lesions indicate infection. The FGSG_09786 has completely lost the ability to spread in the tissue, while the other NOD mutant strains were similar to their respective complement strains.

**Table S5.**
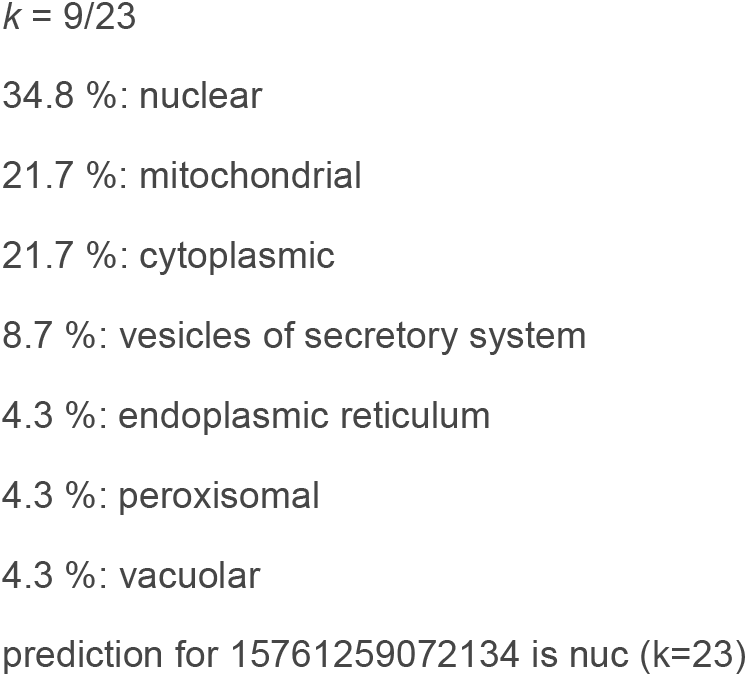
PSORTII predicted localization output for the ER-membrane localized FgNCP_NO_.

**Figure S6.**
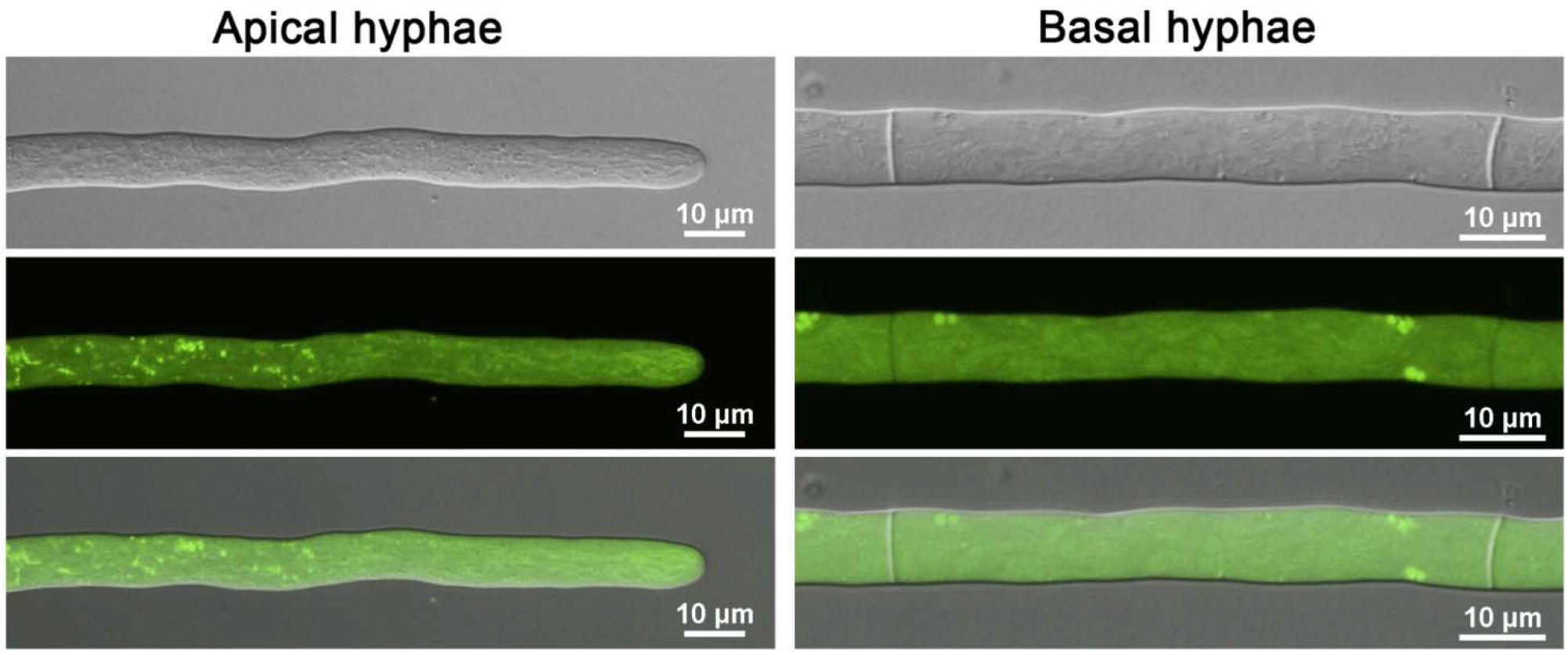
Localization of a potential FgCYP_NO_ (FGSG_07925) labelled with GFP. Localization is not to ER as expected, so this is not the FgCYP_NO_. (Top) DIC images (Middle) GFP images (Bottom) Merge.

**Figure S7.**
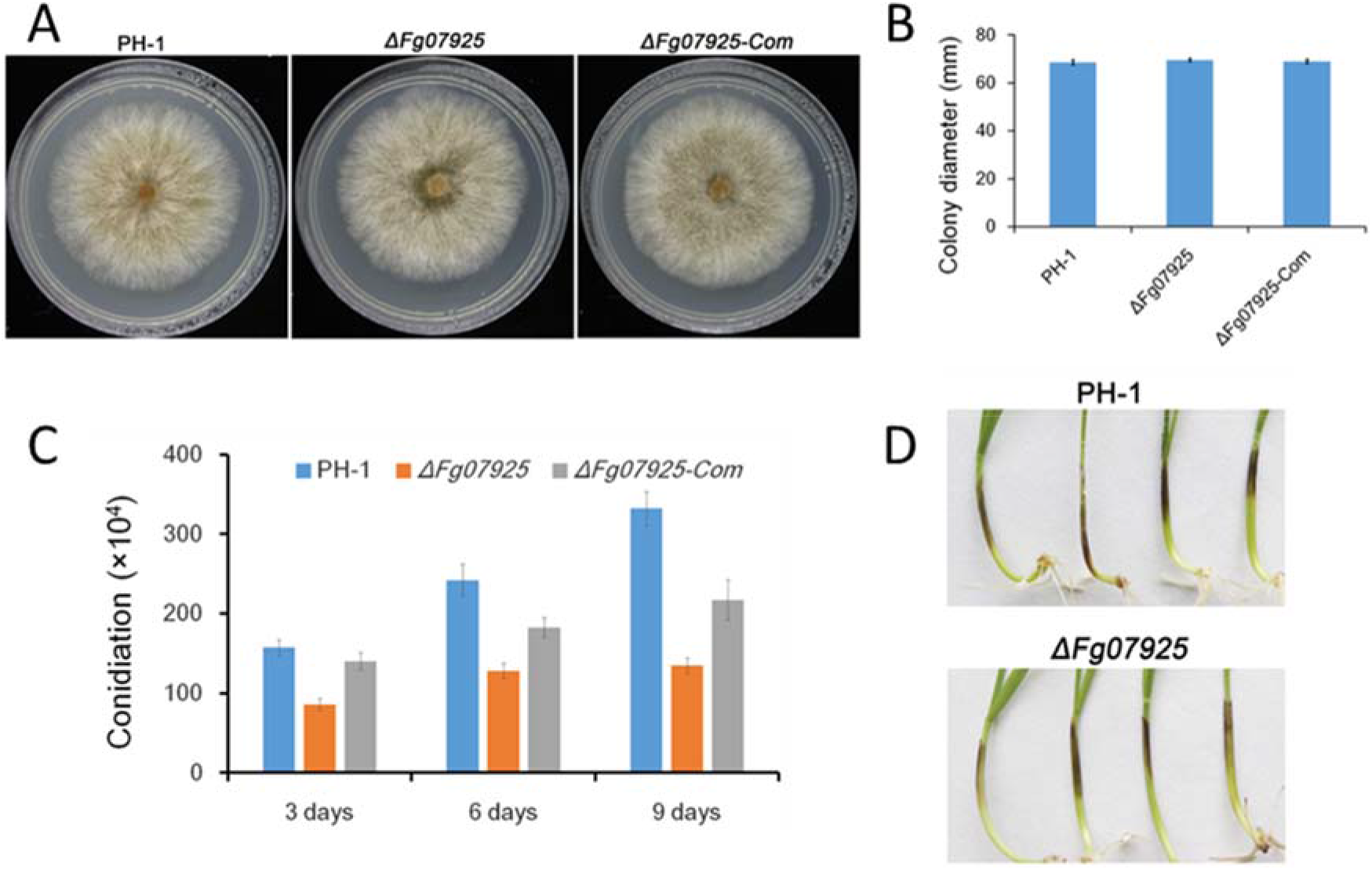
Phenotypic characterization of a CYP gene deletion mutant (FGSG_07925), although this gene cannot encode for the CYP_NO_ since it is not ER-localized (A and B). Deletion of FGSG_07925 does not affect fungal growth on a CM medium. (C) FGSG_07925 appear to be required for conidiation. (D) Deletion of FGSG_07925 does not either affect fungal pathogenicity on wheat coleoptiles. In B and C error bars are SEM and N=3

**Table S8.** List of genes encoding for predicted heme-containing proteins (*FgCYP_NO_*) sorted after gene expression most correlated with *FgNCP_NO_* expression (FGSG_09786) as target gene (T). We first tried to knock out and localize FGSG_07925, but that was not localized to the ER as FGSG_09786. As shown in the table, FGSG_01000 is the most likely *FgCYP_NO_* taking all the following 5 criteria into account since it is (1) a predicted heme-containing protein that is not a predicted NOD. (2) It is transcriptionally highly correlated with *FgNCP_NO_* expression in the dataset from short-term responses to purified bacterial MAMPs. (3) It is predicted to localize to the ER membrane where the FgCYP_NO_ can be found. (4) The maximum expression of the FGSG_01000 is of the same magnitude as the *FgNCP_NO_* expression, which would be expected if they work together as a heterodimer to make NO (5) similar to the homo-dimer iNOS assembly.

| id | Corr | MAX/MaxT* | Annot/location (ER),predictor,experimental/comment |
| --- | --- | --- | --- |
| FGSG_09786 | 1 | 1 | HemeR/located to ER, DeepLoc, Our data |
| FGSG_07925 | 0.786385 | 14.42007 | Heme/Not in ER, DeepLoc, Our data |
| FGSG_00765 | 0.767517 | 2.010113 | NOD1/Not in ER, DeepLoc, Our data/removes NO toxicity |
| FGSG_04458 | 0.517216 | 5.949739 | NOD2/Not in ER, DeepLoc, Our data/removes NO toxicity |
| FGSG_01000 | 0.516057 | 1.00908 | Heme/ER-membrane, DeepLoc/GOOD CANDIDATE |
| FGSG_09373 | 0.425022 | 5.08457 | Heme/Not in ER, Deeploc |
| FGSG_01972 | 0.295199 | 0.264709 | Heme/Not in ER, Deeploc |
| FGSG_10305 | 0.286719 | 2.102463 | Heme/Not in ER, Deeploc |
| FGSG_10881 | 0.269646 | 6.574407 | Heme/Not in ER, Deeploc |
| FGSG_11195 | 0.247331 | 0.008291 | Heme/Not in ER, Deeploc |
| FGSG_02982 | 0.237979 | 0.019898 | Heme/Heme pred. to ER, Deeploc/Expression too low |
| FGSG_07977 | 0.237186 | 0.000886 | Heme/Not in ER, Deeploc |
| FGSG_12534 | 0.234465 | 0.185611 | Heme/Heme pred. to ER, Deeploc/Expression too low |

\*MAX expression of gene/ Max expression of Target gene (FGSG\_09786).

**Figure S9.**
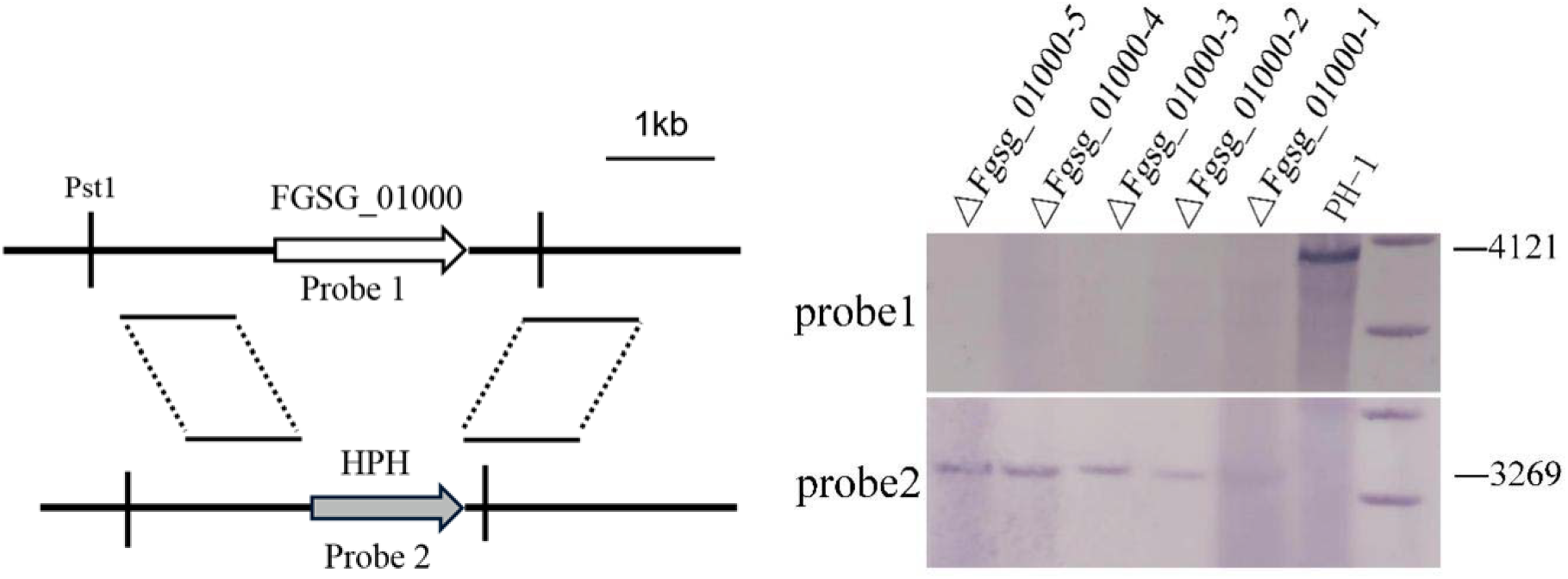
Targeted gene-replacement construct and Southern blot assays for the deletion of FGSG_01000. The targeted gene-replacement construct is shown (left panel), and a Southern blot of digested genomic DNAs showed a 4.12 kb band in the WT and a predicted 3.27 kb band in the deletion mutants.

**Figure S10.**
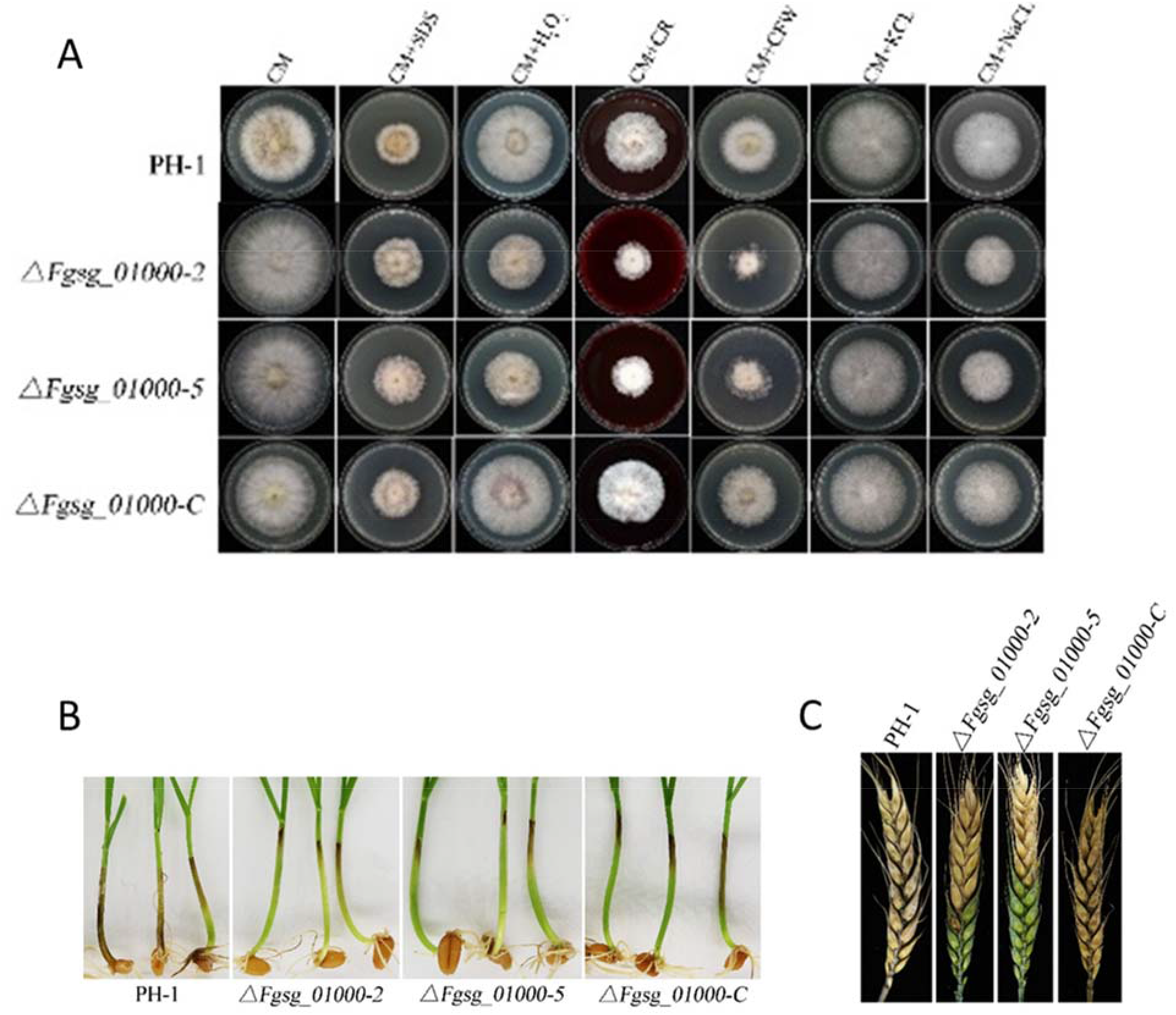
Phenotypic effects of the FGSG_01000 deletion (A) Effect of stress treatment by adding stressors to CM-medium WT (PH-1), mutant strains and complement strain. SDS=0.01%, H_2_O_2_=5 mM, Congo red CR=1 mg/ml, CFW=200μl/ml, KCL=1M, NaCl=1M. (B) Effect deletion and complementation on wheat seedling infection and (C) wheat head infection. As for the deletion of FGSG_09786 (the NCP taking part in making NO), the stress treatments affecting the integrity of the cell wall and cell membrane are most affected, as well as infection of both wheat seedlings and wheat heads.

**Figure S11.**
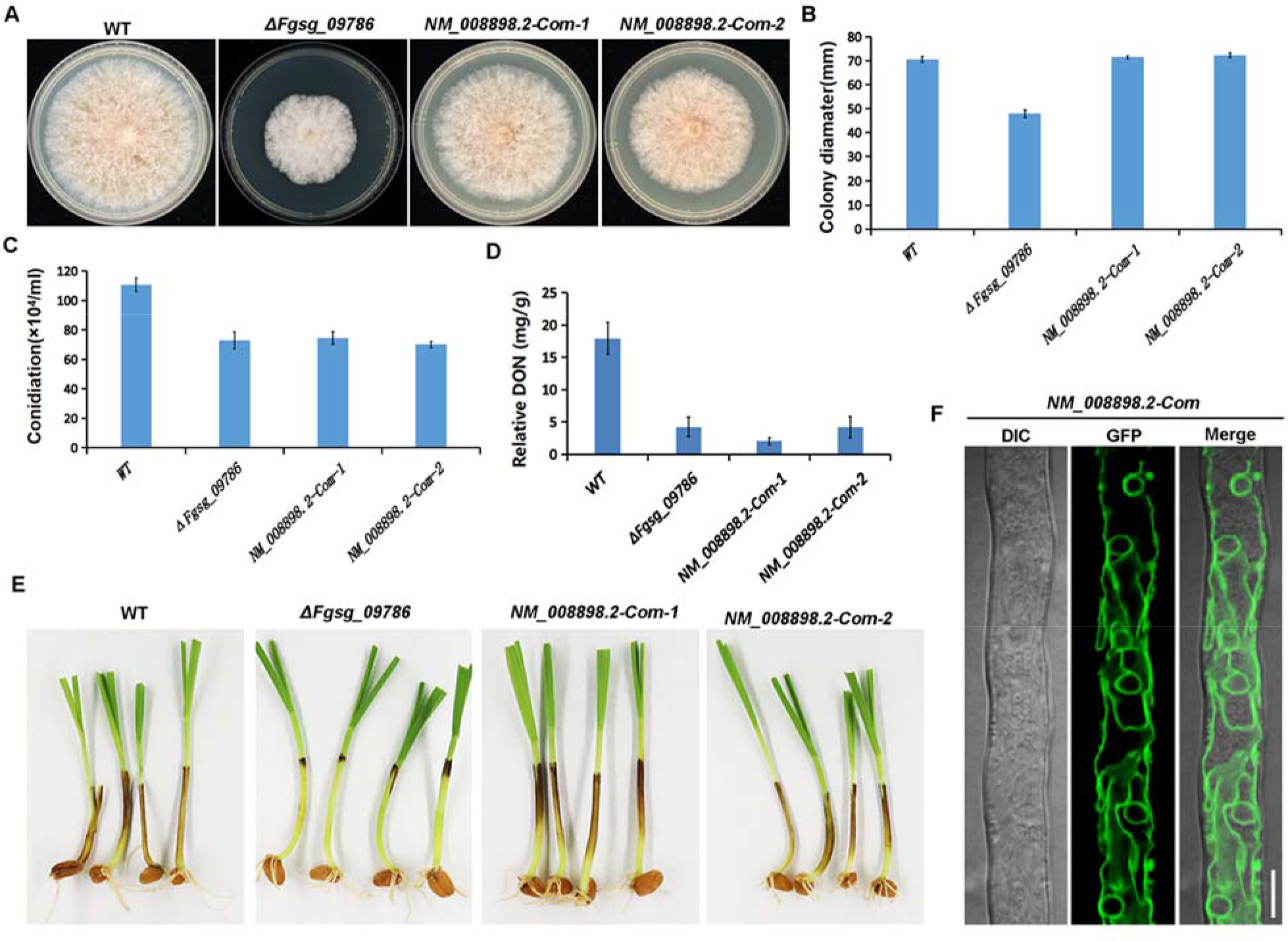
Complementation of the NCP_NO,TRI,ERG_ deletion with the orthologous gene from mouse NM_008898. Effect on colony morphology (A), colony growth (B), conidiation (C), DON production (D) and pathogenicity (E). The mouse gene used for complementation was tagged with GFP and showed typical ER localization (F) (sizebar 10μm). The deleted NCP_NO,TRI,ERG_ showed similar phenotypes as the WT (PH-1) except for conidiation and for DON-production, showing that the mouse gene cannot completely complement all functions. In B-D error bars are SEM and N=3.

**Figure S12.**
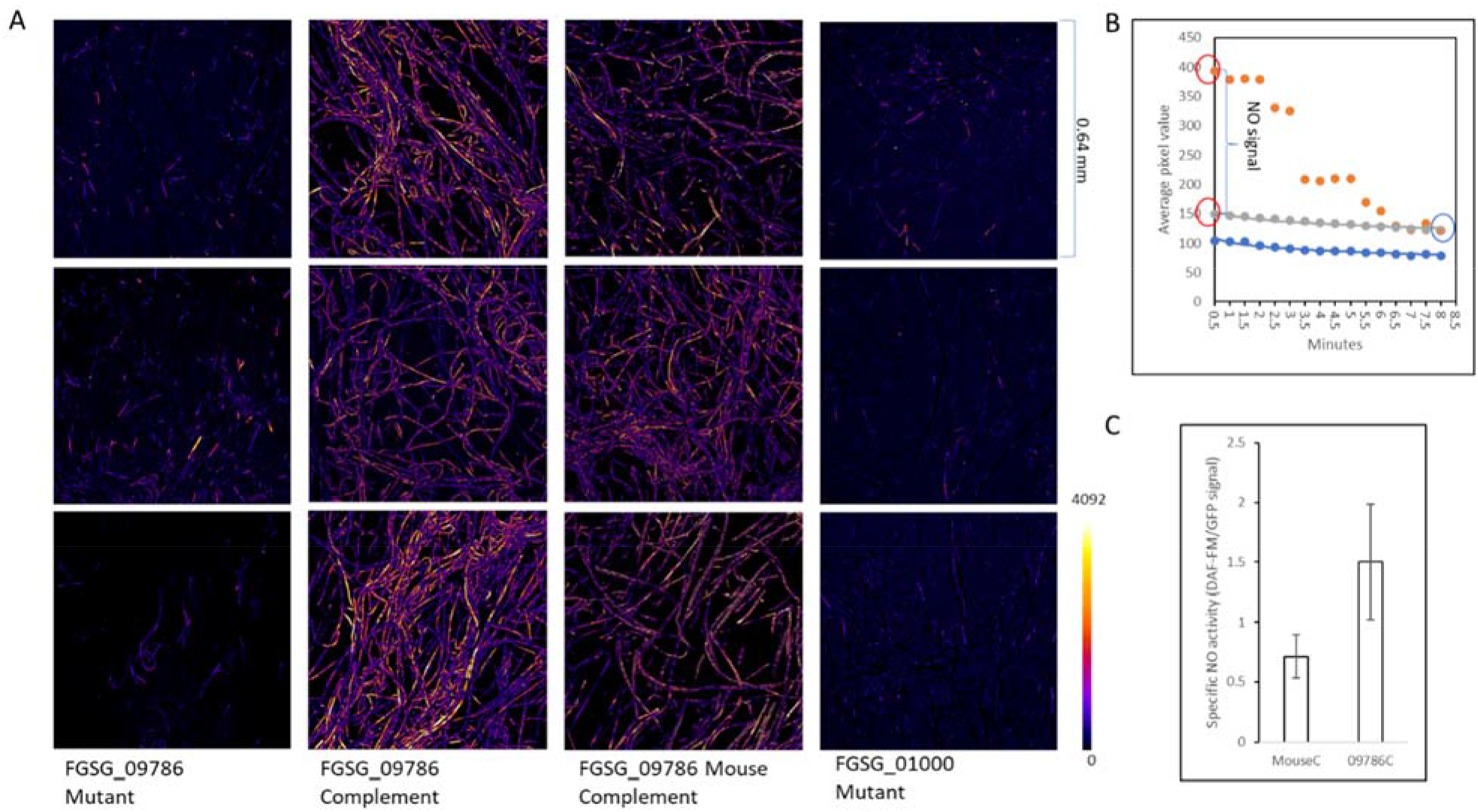
NO formation by the FgNCP_NO,TRI,ERG_ mutant (ΔFGSG_09786), complementation with the native gene complement (FGSG_09586-GFP, FgNCP-GFP) encoding the native NCP protein, and the complementation with a mouse orthologous gene (Mouse-GFP) encoding the mouse NCP protein. The mutant of the NO forming gene for the CYP_NO,ERG_ (FGSG_01000) the NCP_NO,TRI,ERG_ delivers electrons to and makes NO was used as an extra negative control. All images have been handled in the same way. NO forms immediately in DAF FM DA solution (0.5 minutes) and then bleaches under the blue light laser. Both the fluorescent NO derivative of DAF-FM and the GFP-fusion complement fluorescence green excited by the blue laser. However, the stronger DAF-FM NO signal fades quickly, and the fading of the GFP signal is much slower and very reproducible. Thus, the 0.5-minute signal from the NO can be reconstructed using the decay function or the GFP signal that decreases exponentially with a constant percentage per time unit. (**A**) The reconstructed 0.5-minute NO signal images are shown using a “fire” colour scale between 0-4092 (colour scale in the lower right corner). (**B**) Illustration of how the reconstruction was done. Blue curves show that the bleach signal of the GFP decreases with the same percentage for each time unit (like radioactive decay). The reconstructed grey curve is based on the end value (marked by a blue oval) where only GFP fluorescence is left using the blue curve as a reference for the loss of GFP fluorescence. The NO signal at 0.5 minutes equals the difference between the reconstructed 0.5-minute signal for the GFP and the 0.5 signal for GFP+NO. (**C)** Specific NO forming activity by the two complement proteins as DAF-FM signal/GFP signal. Three replicates were used, and error bars show 95% confidence intervals for the mean values. Thus, the P-value for the null hypothesis for the same average of bars with non-overlapping error bars <<0.05. The results show that the mouse NCP can successfully complement the deleted FgNCP_NO,TRI,ERG_ for most NO formations. Note: We had to mount the fungus in an isotonic pH6-7 phosphate buffer for the GFP signal not to fade quickly. That indicates that blue light can compromise membrane permeability and lower cytosolic pH, leading to lower fluorescence as GFP fluorescence is pH sensitive.

**Figure S13.**
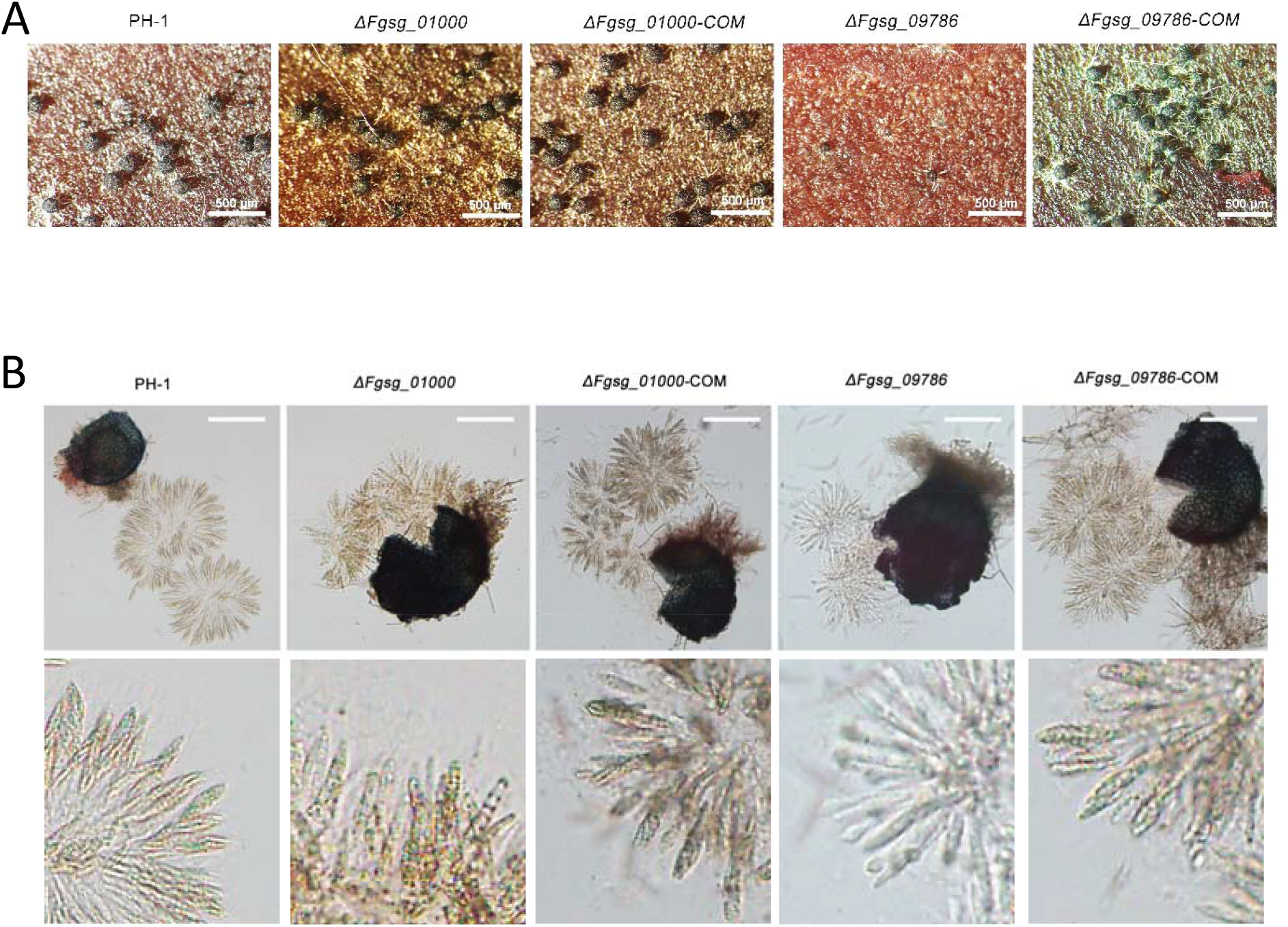
Perithecia formation and ascospore formation on carrot agar. (**A**) Perithecia (size bars 100 μm). Only Δ*Fgsg_09786* show a reduced number of perithecia. (**B**) Asci and ascospores (Size bars 50 μm). Both mutants show abnormal asci. Δ*Fgsg_01000* has septate asci that appear empty of spores. Also, Δ*Fgsg_09786* asci appear empty of normal-looking ascospores.

**Table S14.**
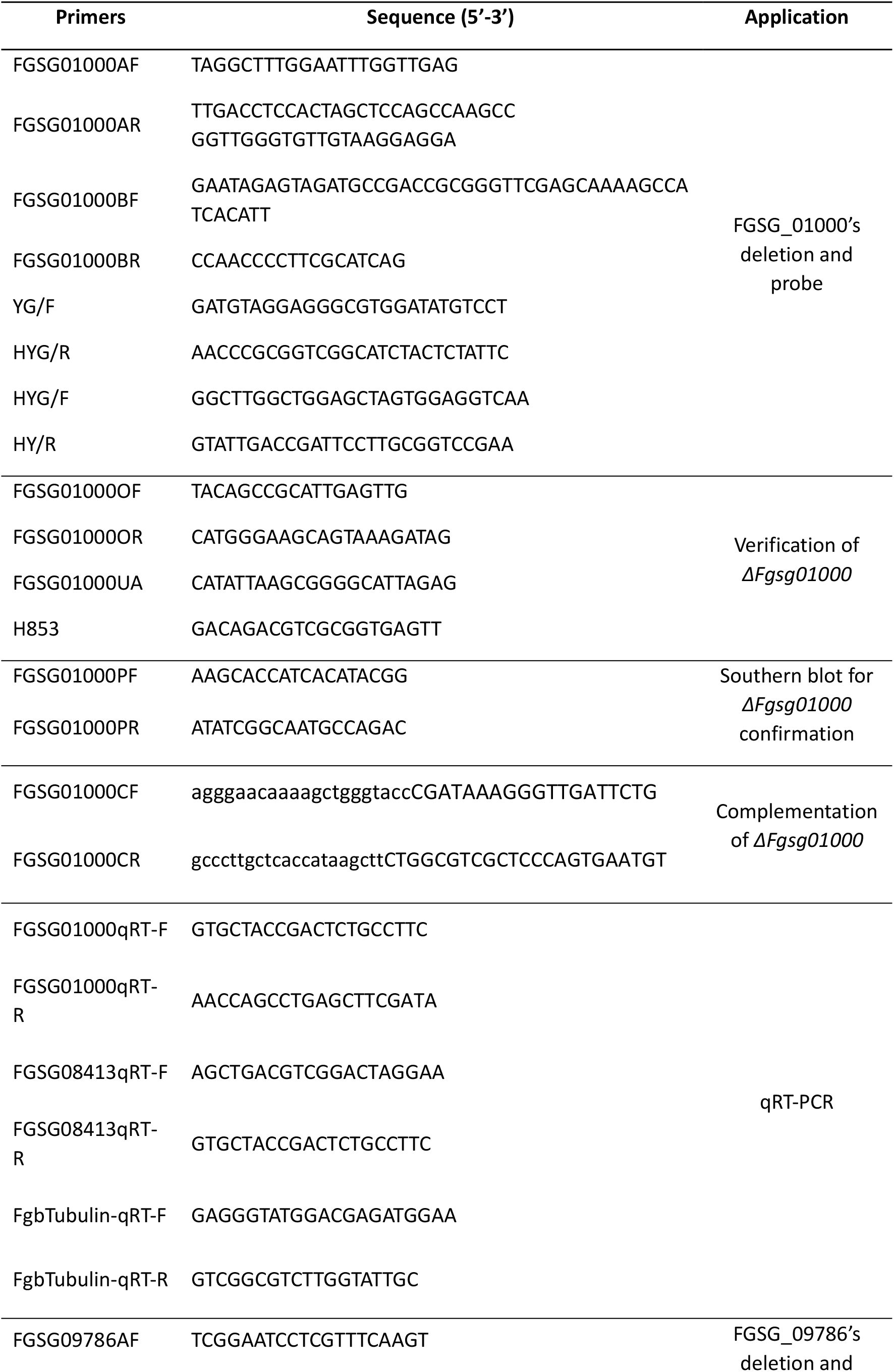
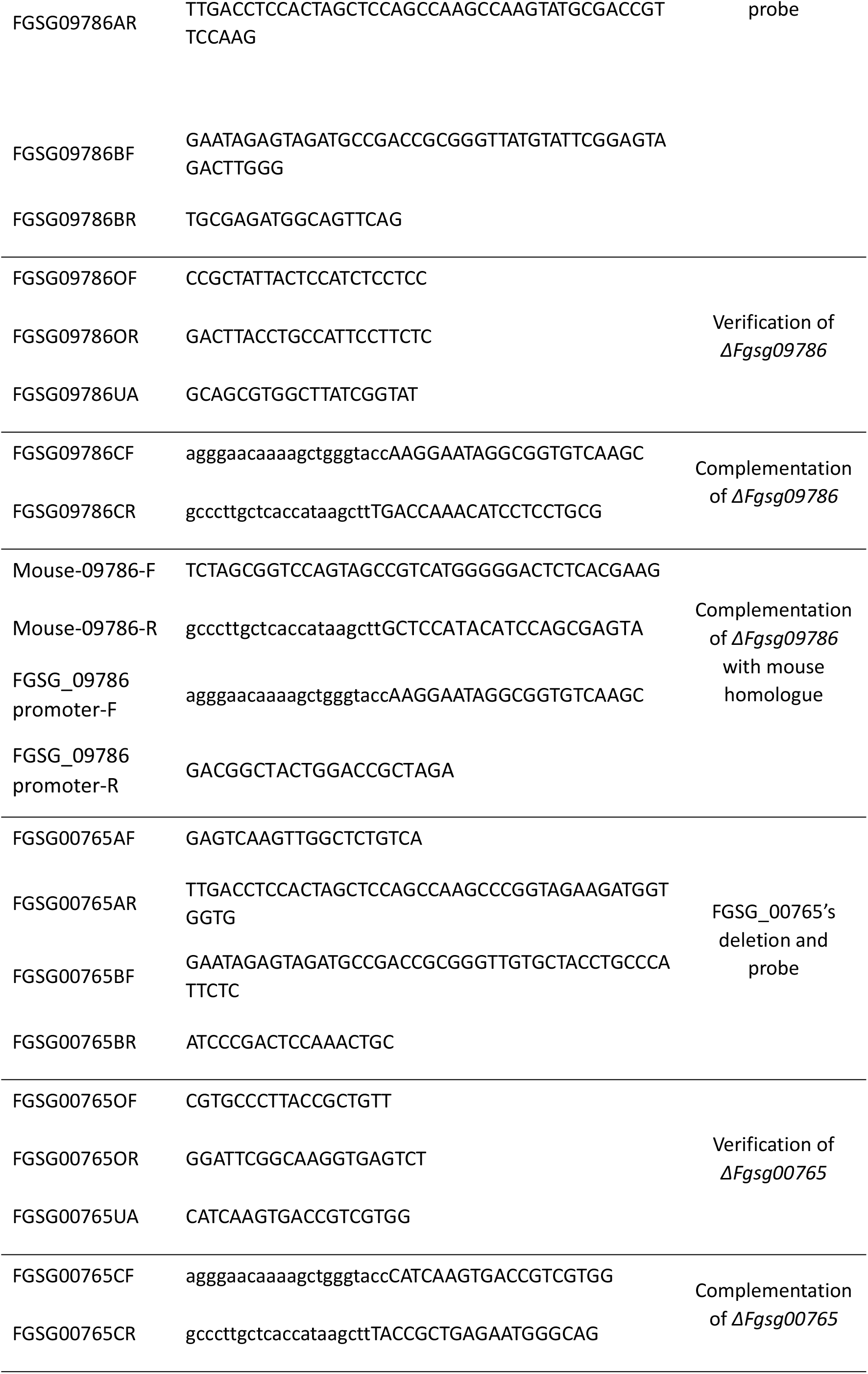
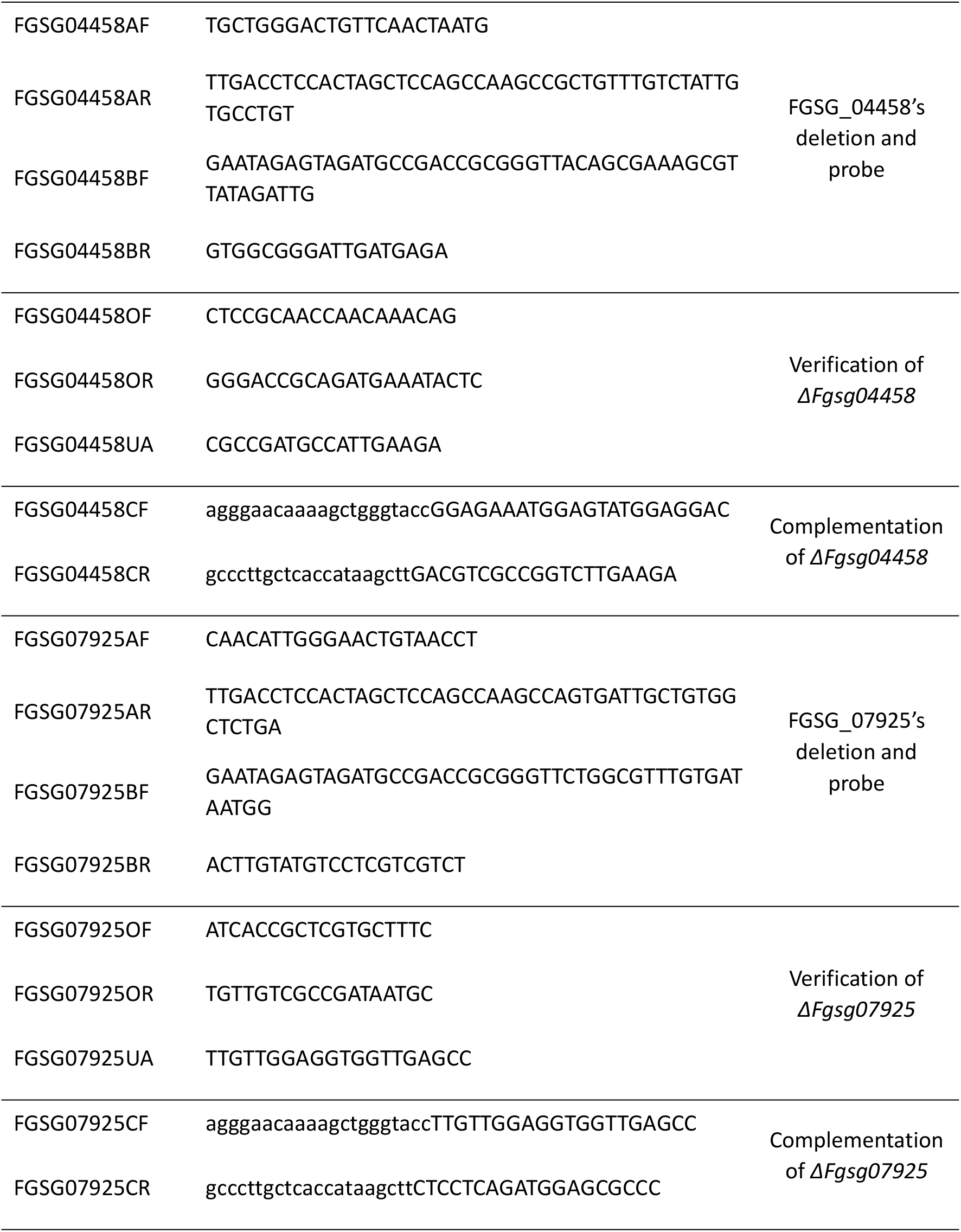
Primers used in this study.

