## Supplemental Files and Data for "Evolutionary conserved multifunctional nitric oxide synthesis proteins responding to bacterial MAMPs are located at the endoplasmic reticulum": Supplementary file SF1 Luminol measurements.pptx

### Slide 1
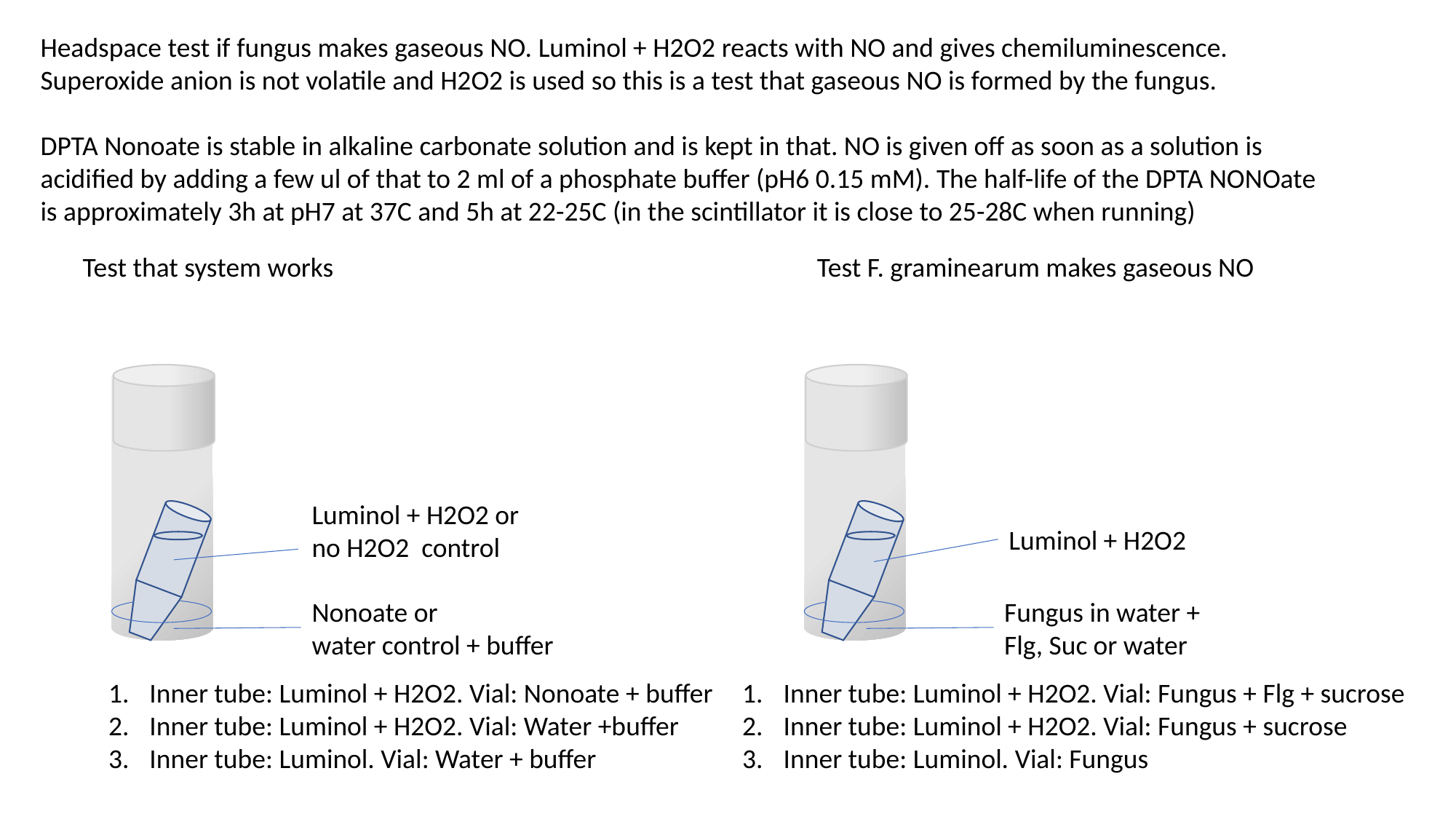

Headspace test if fungus makes gaseous NO. Luminol + H2O2 reacts with NO and gives chemiluminescence. Superoxide anion is not volatile and H2O2 is used so this is a test that gaseous NO is formed by the fungus.
DPTA Nonoate is stable in alkaline carbonate solution and is kept in that. NO is given off as soon as a solution is acidified by adding a few ul of that to 2 ml of a phosphate buffer (pH6 0.15 mM). The half-life of the DPTA NONOate is approximately 3h at pH7 at 37C and 5h at 22-25C (in the scintillator it is close to 25-28C when running)
Test that system works
Test F. graminearum makes gaseous NO
Luminol + H2O2 or
no H2O2 control
Nonoate or
water control + buffer
Luminol + H2O2
Fungus in water +
Flg, Suc or water
Inner tube: Luminol + H2O2. Vial: Nonoate + buffer
Inner tube: Luminol + H2O2. Vial: Water +buffer
Inner tube: Luminol. Vial: Water + buffer
Inner tube: Luminol + H2O2. Vial: Fungus + Flg + sucrose
Inner tube: Luminol + H2O2. Vial: Fungus + sucrose
Inner tube: Luminol. Vial: Fungus

### Slide 2
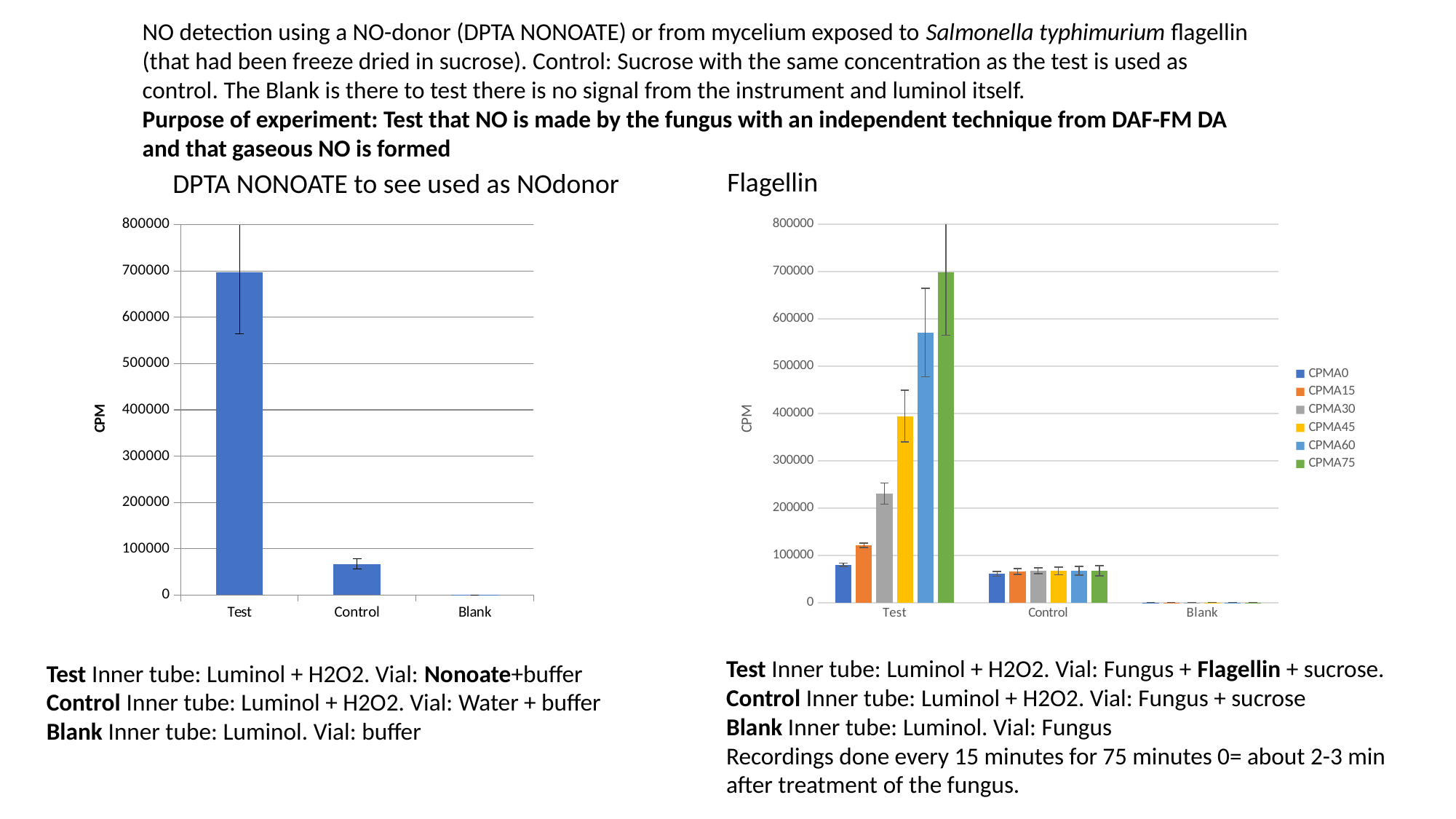

NO detection using a NO-donor (DPTA NONOATE) or from mycelium exposed to Salmonella typhimurium flagellin (that had been freeze dried in sucrose). Control: Sucrose with the same concentration as the test is used as control. The Blank is there to test there is no signal from the instrument and luminol itself.
Purpose of experiment: Test that NO is made by the fungus with an independent technique from DAF-FM DA and that gaseous NO is formed
Flagellin
DPTA NONOATE to see used as NOdonor
#### Chart
| Category | CPMA0 | CPMA15 | CPMA30 | CPMA45 | CPMA60 | CPMA75 |
|---|---|---|---|---|---|---|
| Test | 79919.0 | 121322.33333333333 | 230380.33333333334 | 393937.3333333333 | 570846.6666666666 | 697867.6666666666 |
| Control | 60640.666666666664 | 66078.33333333333 | 67314.66666666667 | 66952.66666666667 | 67162.66666666667 | 67303.66666666667 |
| Blank | 53.333333333333336 | 49.0 | 53.0 | 60.333333333333336 | 58.0 | 67.66666666666667 |
#### Chart
| Category | |
|---|---|
| Test | 697867.6666666666 |
| Control | 67303.66666666667 |
| Blank | 67.66666666666667 |Test Inner tube: Luminol + H2O2. Vial: Fungus + Flagellin + sucrose.
Control Inner tube: Luminol + H2O2. Vial: Fungus + sucrose
Blank Inner tube: Luminol. Vial: Fungus
Recordings done every 15 minutes for 75 minutes 0= about 2-3 min after treatment of the fungus.
Test Inner tube: Luminol + H2O2. Vial: Nonoate+buffer
Control Inner tube: Luminol + H2O2. Vial: Water + buffer
Blank Inner tube: Luminol. Vial: buffer

### Slide 3
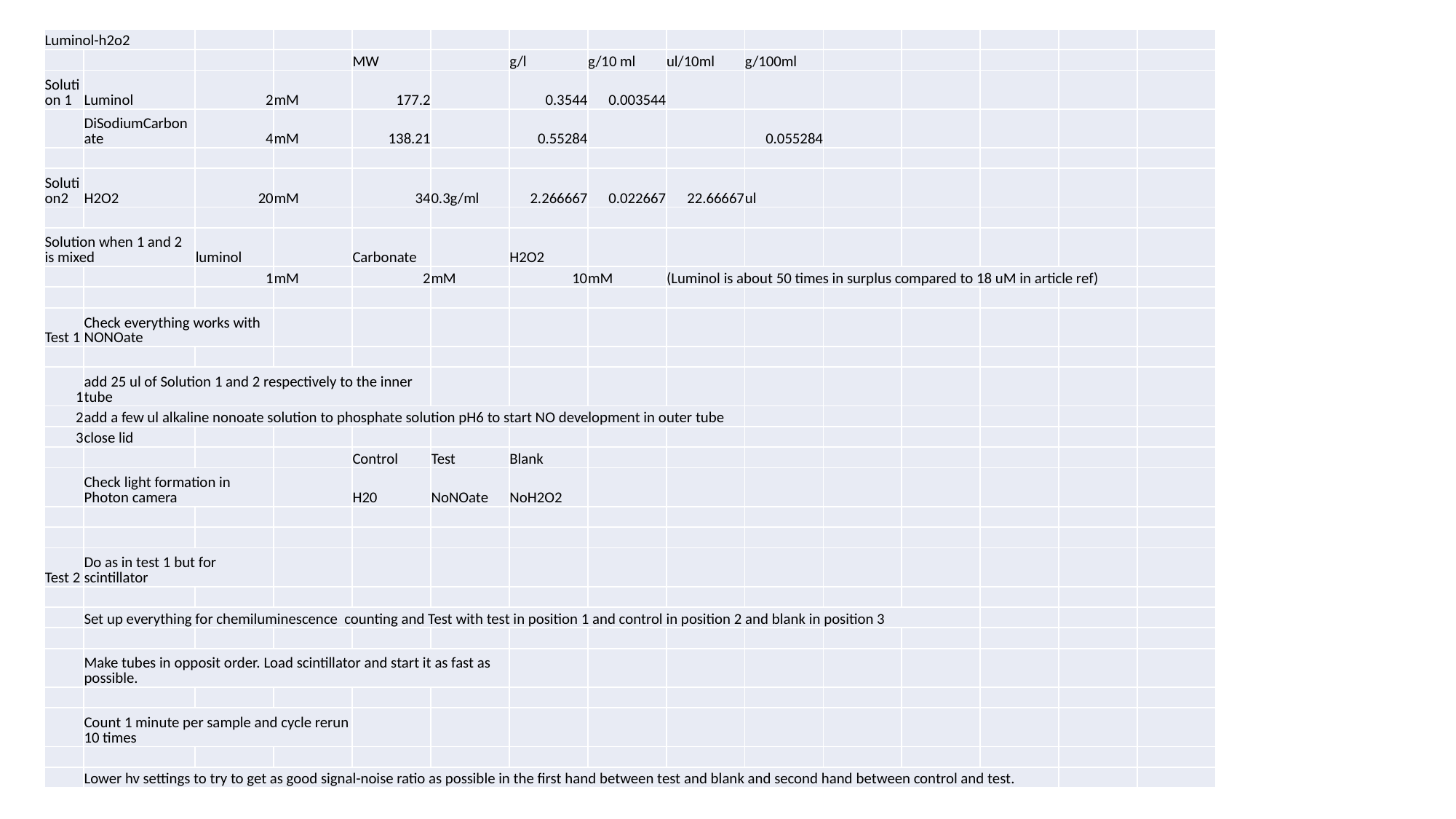

| Luminol-h2o2 | | | | | | | | | |
| --- | --- | --- | --- | --- | --- | --- | --- | --- | --- |
| | | | | MW | | g/l | g/10 ml | ul/10ml | g/100ml |
| Solution 1 | Luminol | 2 | mM | 177.2 | | 0.3544 | 0.003544 | | |
| | DiSodiumCarbonate | 4 | mM | 138.21 | | 0.55284 | | | 0.055284 |
| Solution2 | H2O2 | 20 | mM | 34 | 0.3g/ml | 2.266667 | 0.022667 | 22.66667 | ul |
| Solution when 1 and 2 is mixed | | luminol | | Carbonate | | H2O2 | | | |
| | | 1 | mM | 2 | mM | 10 | mM | (Luminol is about 50 times in surplus compared to 18 uM in article ref) | |
| Test 1 | Check everything works with NONOate | | | | | | | | |
| 1 | add 25 ul of Solution 1 and 2 respectively to the inner tube | | | | | | | | |
| 2 | add a few ul alkaline nonoate solution to phosphate solution pH6 to start NO development in outer tube | | | | | | | | |
| 3 | close lid | | | | | | | | |
| | | | | Control | Test | Blank | | | |
| | Check light formation in Photon camera | | | H20 | NoNOate | NoH2O2 | | | |
| Test 2 | Do as in test 1 but for scintillator | | | | | | | | |
| | Set up everything for chemiluminescence counting and Test with test in position 1 and control in position 2 and blank in position 3 | | | | | | | | |
| | Make tubes in opposit order. Load scintillator and start it as fast as possible. | | | | | | | | |
| | Count 1 minute per sample and cycle rerun 10 times | | | | | | | | |
| | Lower hv settings to try to get as good signal-noise ratio as possible in the first hand between test and blank and second hand between control and test. | | | | | | | | |

### Slide 4
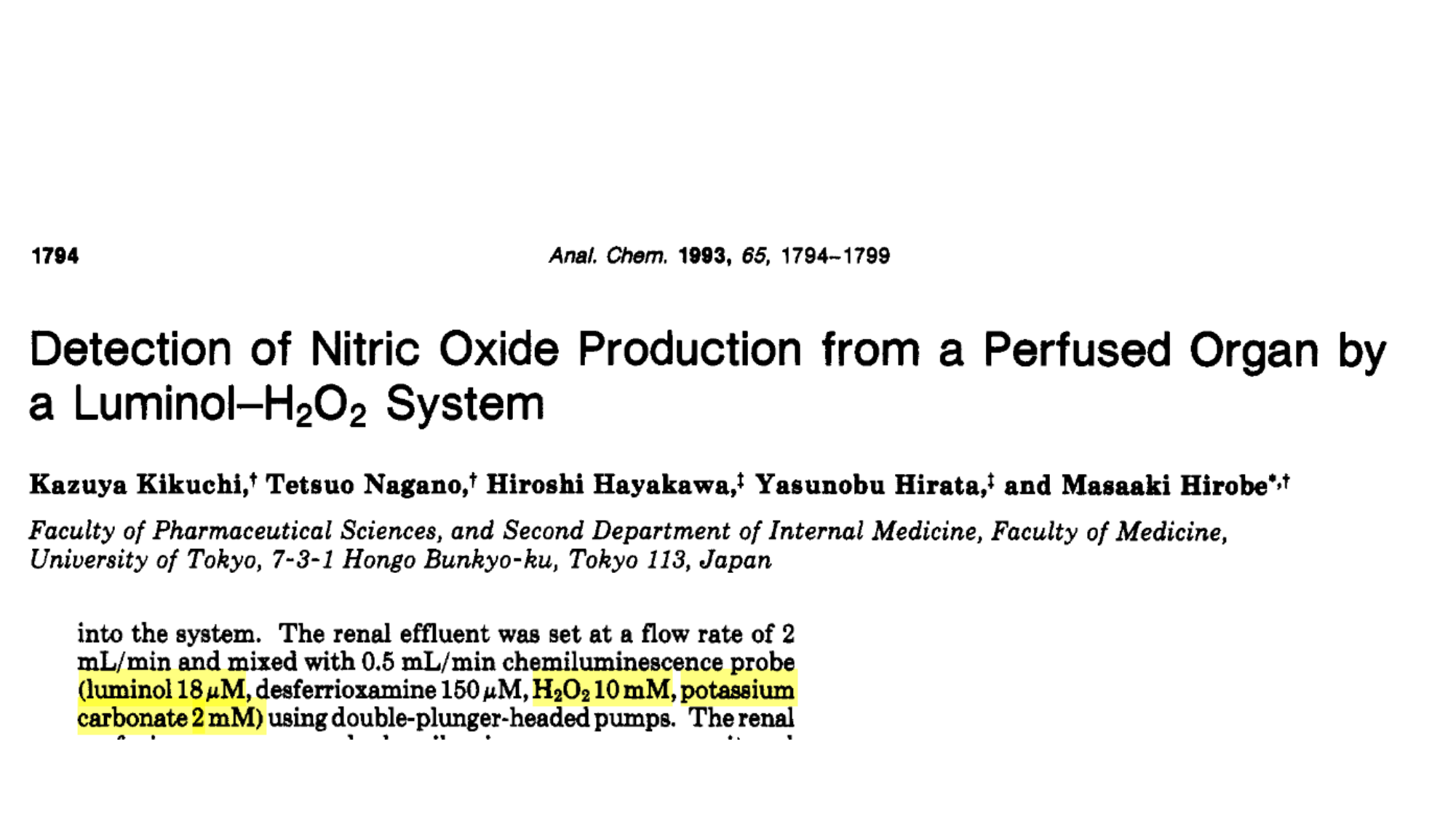
