## Supplemental Files and Data for "Evolutionary conserved multifunctional nitric oxide synthesis proteins responding to bacterial MAMPs are located at the endoplasmic reticulum": Supplementary file SF2 Finding a iNOS type protein containing the NCP(NO) reductase part of the enzyme in Fg PH1.docx

**Procedure to find a nitric oxide synthase type protein containing the** **NADPH dependent Cytochrome P450 (NCP) part of an iNOS**

**1. Mouse iNOS sequence was downloaded from NCBI**

>NP_035057.1 nitric oxide synthase, inducible isoform a [Mus musculus]

MACPWKFLFKVKSYQSDLKEEKDINNNVKKTPCAVLSPTIQDDPKSHQNGSPQLLTGTAQNVPESLDKLH

VTSTRPQYVRIKNWGSGEILHDTLHHKATSDFTCKSKSCLGSIMNPKSLTRGPRDKPTPLEELLPHAIEF

INQYYGSFKEAKIEEHLARLEAVTKEIETTGTYQLTLDELIFATKMAWRNAPRCIGRIQWSNLQVFDARN

CSTAQEMFQHICRHILYATNNGNIRSAITVFPQRSDGKHDFRLWNSQLIRYAGYQMPDGTIRGDAATLEF

TQLCIDLGWKPRYGRFDVLPLVLQADGQDPEVFEIPPDLVLEVTMEHPKYEWFQELGLKWYALPAVANML

LEVGGLEFPACPFNGWYMGTEIGVRDFCDTQRYNILEEVGRRMGLETHTLASLWKDRAVTEINVAVLHSF

QKQNVTIMDHHTASESFMKHMQNEYRARGGCPADWIWLVPPVSGSITPVFHQEMLNYVLSPFYYYQIEPW

KTHIWQNEKLRPRRREIR

FRVLVKVVFFASMLMRKVMASRVRATVLFATETGKSEALARDLATLFSYAFN

TKVVCMDQYKASTLEEEQLLLVVTSTFGNGDCPSNGQTLKKSLFMLRELNHTFRYAVFGLGSSMYPQFCA

FAHDIDQKLSHLGASQLAPTGEGDELSGQEDAFRSWAVQTFRAACETFDVRSKHHIQIPKRFTSNATWEP

QQYRLIQSPEPLDLNRALSSIHAKNVFTMRLKSQQNLQSEKSSRTTLLVQLTFEGSRGPSYLPGEHLGIF

PGNQTALVQGILERVVDCPTPHQTVCLEVLDESGSYWVKDKRLPPCSLSQALTYFLDITTPPTQLQLHKL

ARFATDETDRQRLEALCQPSEYNDWKFSNNPTFLEVLEEFPSLHVPAAFLLSQLPILKPRYYSISSSQDH

TPSEVHLTVAVVTYRTRDGQGPLHHGVCSTWIRNLKPQDPVPCFVRSVSGFQLPEDPSQPCILIGPGTGI

APFRSFWQQRLHDSQHKGLKGGRMSLVFGCRHPEEDHLYQEEMQEMVRKRVLFQVHTGYSRLPGKPKVYV

QDILQKQLANEVLSVLHGEQGHLYICGDVRMARDVATTLKKLVATKLNLSEEQVEDYFFQLKSQKRYHED

IFGAVFSYGAKKGSALEEPKATRL

Colored=Heme reductase domain including hydrophobic transmembrane signal peptide or membrane anchor section identified after HMMER (below)

**2. Use HMMER For identifying domain structure and domain borders**

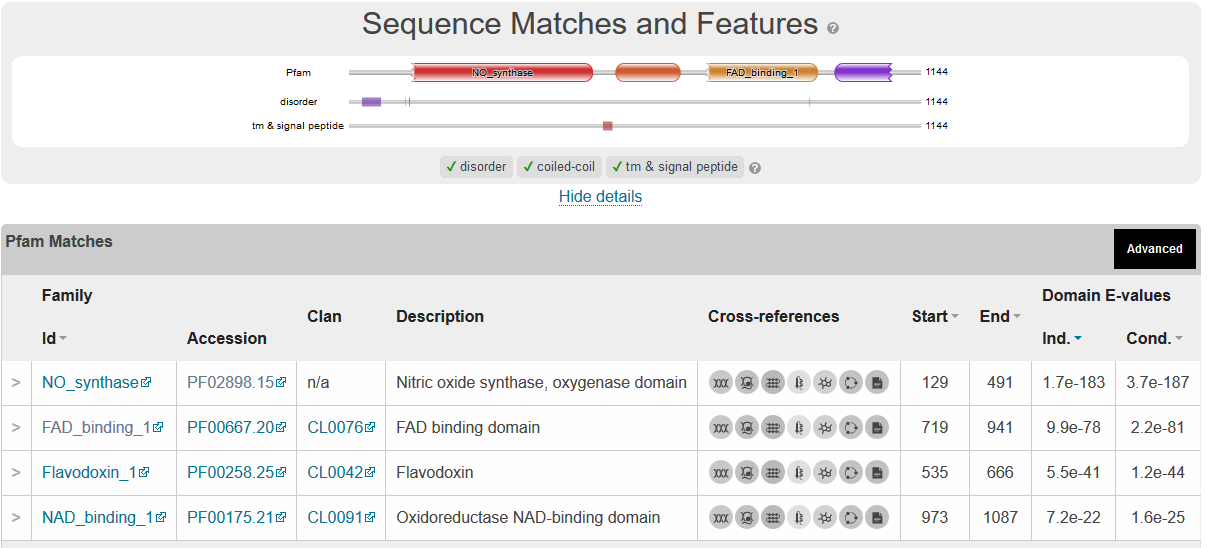

**TM=509-527**

**Disorder=27-64**

**3. NCBI Blast search the yellow-coloured sequence in 1 against the predicted proteins in the F. graminearum PH1 genome**

Blast results of the last 508 residues of the Mouse NOS against Fusarium graminearum gave two hits with high cover.

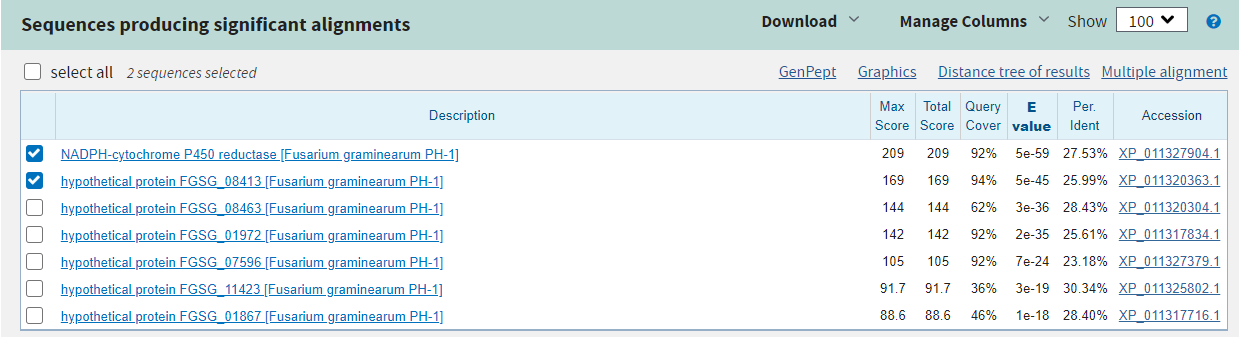

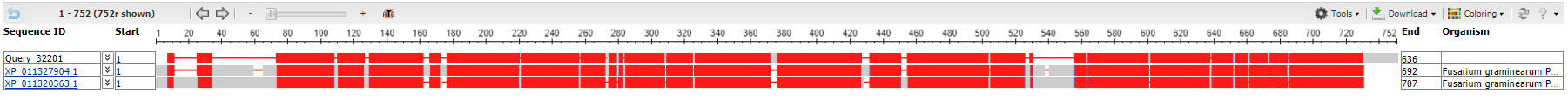

**3. Confirm with HMMER that the found sequences have a similar predicted domain structure as the mouse heme reductase part of the iNOS**

**FGSG_09786**

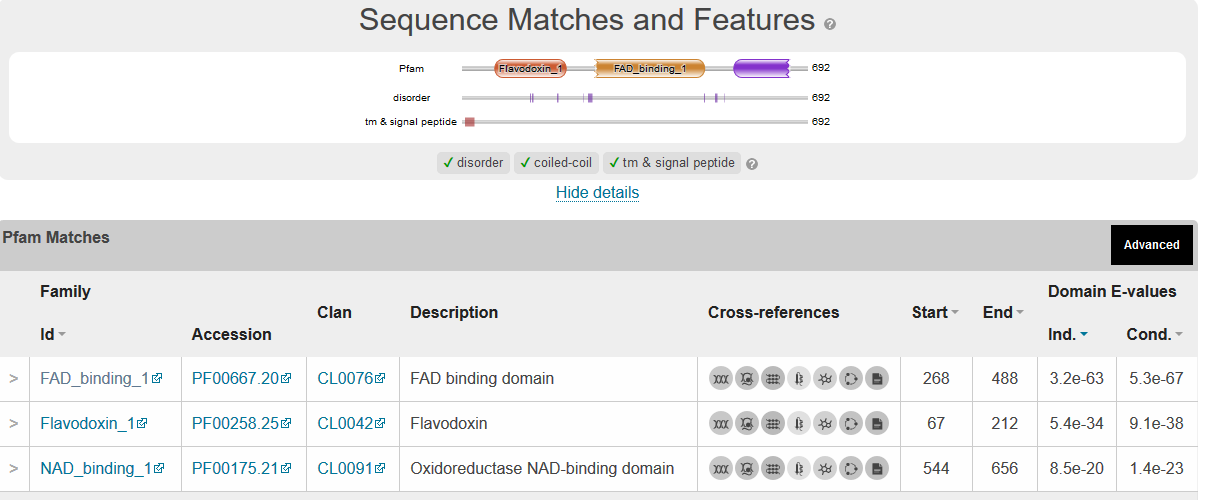

**FGSG_08413**
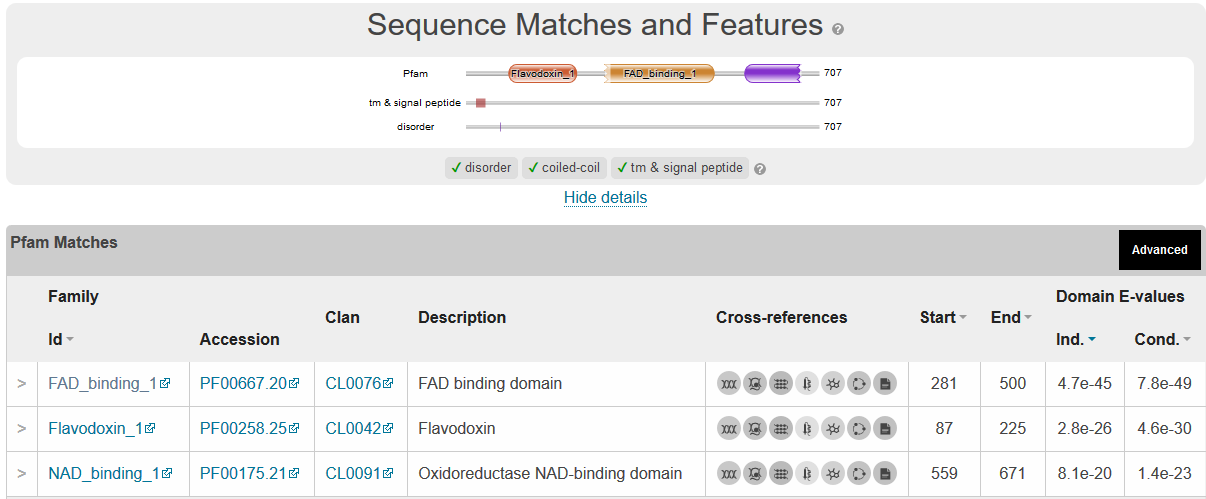

**Conclusion**

Two proteins similar to the NCP part of the mouse inducible NOS was found based on sequence similarities, FGSG_09786 and FGSG_08413. Both have the same domain structure as mouse iNOS, interestingly also including an N-terminal transmembrane part that in the iNOS lies between the CYP-like heme protein and the NCP.
