## Supplemental Files and Data for "Evolutionary conserved multifunctional nitric oxide synthesis proteins responding to bacterial MAMPs are located at the endoplasmic reticulum": Supplementary file SF3 Finding Nitric Oxide Dioxygenase proteins NODs in Fg PH1.docx

**Finding Nitric Oxide Dioxygenase proteins NODs in Fusarium graminearum**

**1. Searching NCBI for fungi there are one experimentally verified protein**

>sp|Q59MV9.1|FHP_CANAL RecName: Full=Flavohemoprotein; AltName: Full=Flavohemoglobin; AltName: Full=Hemoglobin-like protein; AltName: Full=Nitric oxide dioxygenase; Short=NO oxygenase; Short=NOD

MTVEYETKQLTPAQIKIILDTVPILEEAGETLTQKFYQRMIGNYDEVKPFFNTTDQKLLRQPKILAFALL

NYAKNIEDLTPLTDFVKQIVVKHIGLQVLPEHYPIVGTCLIQTMVELLPPEIANKDFLEAWTIAYGNLAK

LLIDLEAAEYAKQPWRWFKDFKVTRIVQECKDVKSVYFTPVDKDLLPLPKPERGQYLCFRWKLPGEEFEI

SREYSVSEFPKENEYRISVRHVPGGKISGYIHNNLKVGDILKVAPPAGNFVYDPATDKELIFVAGGIGIT

PLLSMIERALEEGKNVKLLYSNRSAETRAFGNLFKEYKSKFGDKFQAIEYFSEDNNTDDKIVIDKAFNRK

LTTDDLDFIAPEHDVYLVGPREFMKDIKEHLGKKNVPVKLEYFGPYDP

2. The Candida NOD was used to blast the Fusarium graminearum species complex (taxid:569360) for similar proteins.

One protein previously annotated as a NOD was found with high sequence cover, and similarity was found in the *F. graminearum* species complex.

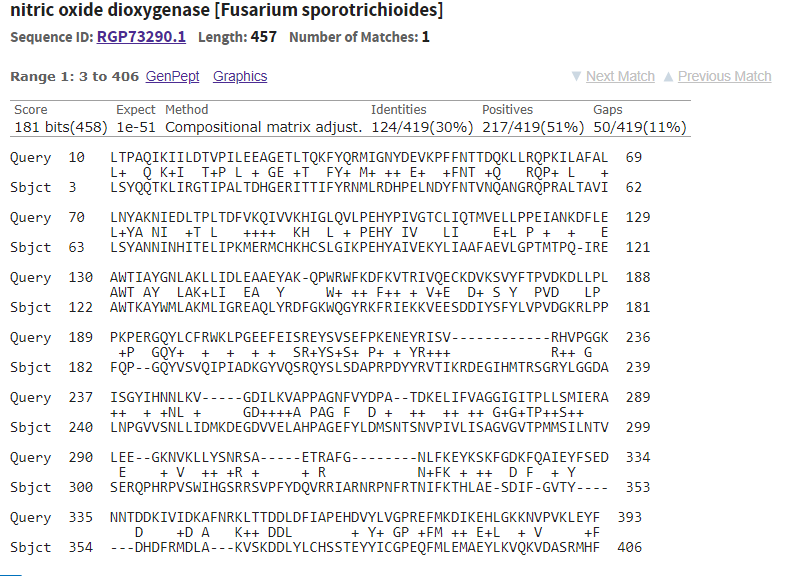

>RGP73290.1:3-406 nitric oxide dioxygenase [Fusarium sporotrichioides]

LSYQQTKLIRGTIPALTDHGERITTIFYRNMLRDHPELNDYFNTVNQANGRQPRALTAVILSYANNINHI

TELIPKMERMCHKHCSLGIKPEHYAIVEKYLIAAFAEVLGPTMTPQIREAWTKAYWMLAKMLIGREAQLY

RDFGKWQGYRKFRIEKKVEESDDIYSFYLVPVDGKRLPPFQPGQYVSVQIPIADKGYVQSRQYSLSDAPR

PDYYRVTIKRDEGIHMTRSGRYLGGDALNPGVVSNLLIDMKDEGDVVELAHPAGEFYLDMSNTSNVPIVL

ISAGVGVTPMMSILNTVSERQPHRPVSWIHGSRRSVPFYDQVRRIARNRPNFRTNIFKTHLAESDIFGVT

YDHDFRMDLAKVSKDDLYLCHSSTEYYICGPEQFMLEMAEYLKVQKVDASRMHF

3. The NOD above belongs to a fungus within the *F. graminearum* species complex. Thus this protein was used for a blast search towards ***F. graminearum*** PH1 to find similar genes in the genome.

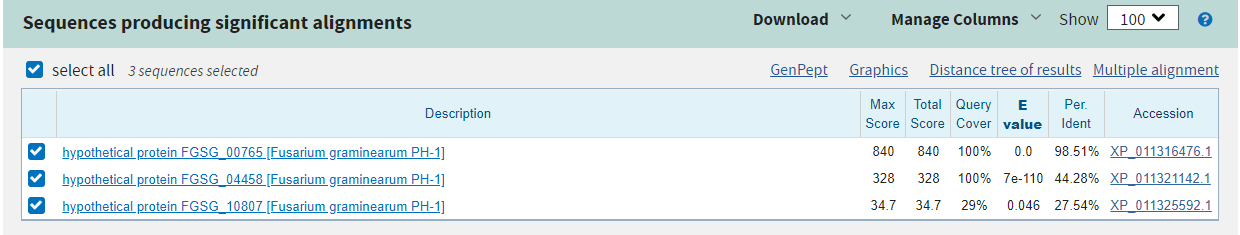

Two good hits were found FGSG_00765 and FGSG_04458

>XP_011316476.1 hypothetical protein FGSG_00765 [Fusarium graminearum PH-1]

MALSYQQTKLIRGTIPALTDHGERITTIFYRNMLRDHPELNDYFNTVNQANGRQPRALTAVILSYANNIN

HITELIPKMERMCHKHCSLGIKPEHYAIVEKYLIAAFAEVLGPTMTTQIREAWTKAYWMLAKMLIGREAQ

LYRDFGKWQGYRKFRIEKKVEESDDIYSFYLVPVDGKRLPSFQPGQYVSVQIPIVDKGYVQSRQYSLSDA

PRPDYYRVTVKRDEGIHMTRSGRYLGGDALNPGVVSNLLIDMKDEGDVVELAHPAGEFYLDMSNTSNVPI

VLISAGVGVTPMMSILNTVSERQPHRPVSWIHGSRRSVPFYDQVRRIARNRSNFRTNIFKTHLAESDIFG

VTYDHDFRMDLAKVSKDDLYLCHSSTEYYICGPEQFMLEMAEYLKAQKVDASRMHFELFSTGDMEFKVDN

LSISSASKSPSIASAEQRCPSTGAVSADGATCPFSAV

>XP_011321142.1 hypothetical protein FGSG_04458 [Fusarium graminearum PH-1]

MALTAAQVAIVKSTAPILKEHGKTITTTFYRNMLGAHPELKNYFSLRNQQTGAQQAALANSVLAYATYID

DLGKLSHAVERIAQKHVSLFIKAEHYPIVGTHLIGAIGEVLGSALTTEIKDAWVAAYGQLADIFIQREGQ

LYDAAGEWNSWRKFKIAKKEAENDSVTSFYLEPLDDKPLPKFLPGQYVSLQIPIPELDGLLQSRQFSLSE

APGSNHYRISVKLQGPEEEPSLEDLSAGKIPGLVCTRLHKRYNVGDEVELSPPAGEFFLNPADTSAAKKP

LVLLSAGVGATPLVSILDSVLESETASRPITWIHGARYSGSTCFVPHVLDSAKKHDNITAKIFLEDVKEG

DQYDFKGEIDLDQLQKDKLLQLDSSDAEYFICGPEDWMVKVRAFLEENGVPRERQHLELFKTGDV

Aligned with the Query next 2 pages

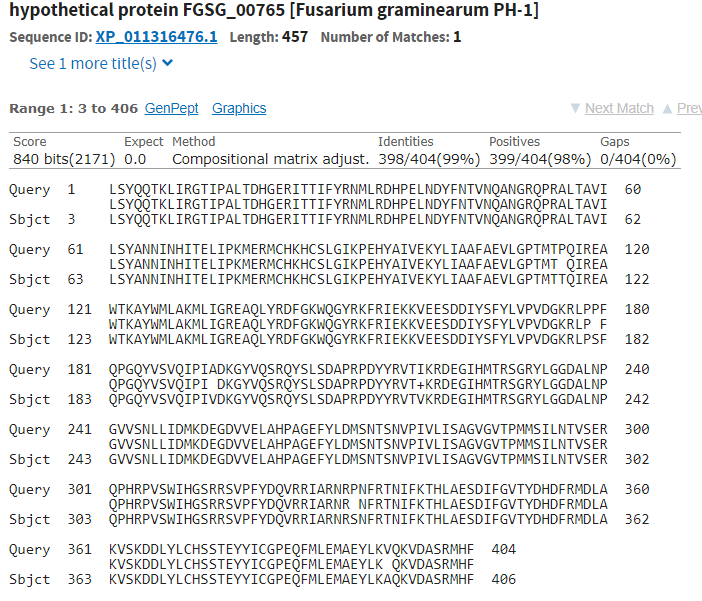

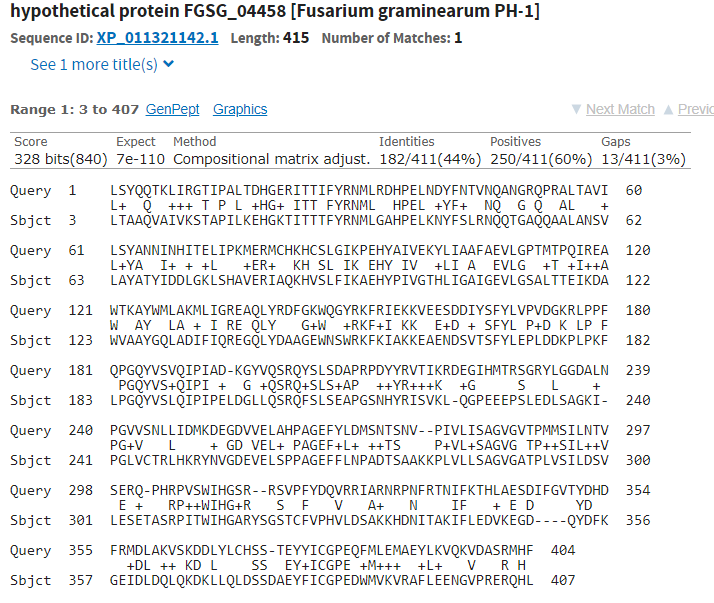

**4. Comparisons of the predicted domain of the NODs** **using HMMER**

Candida albicans NOD
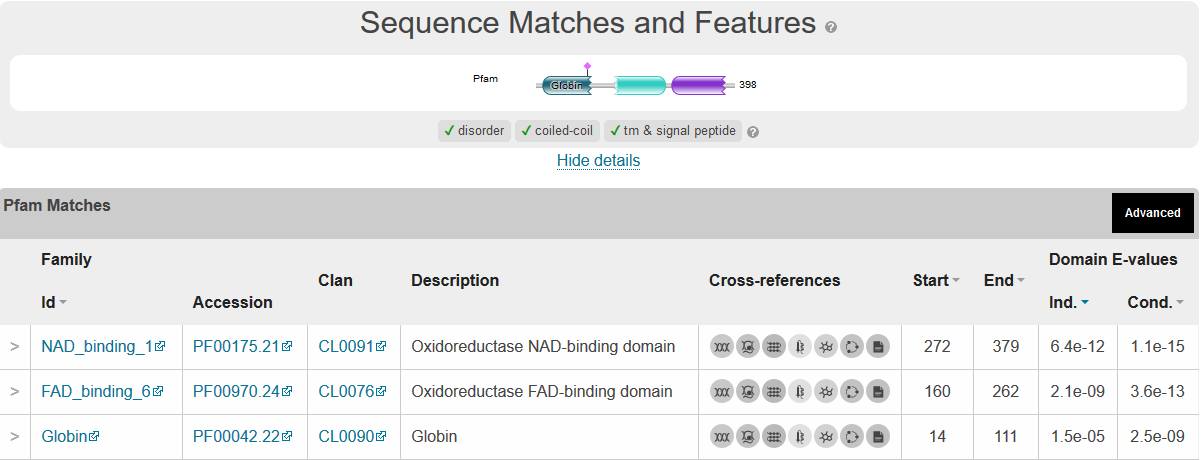

Fusarium sporotrichoides NOD

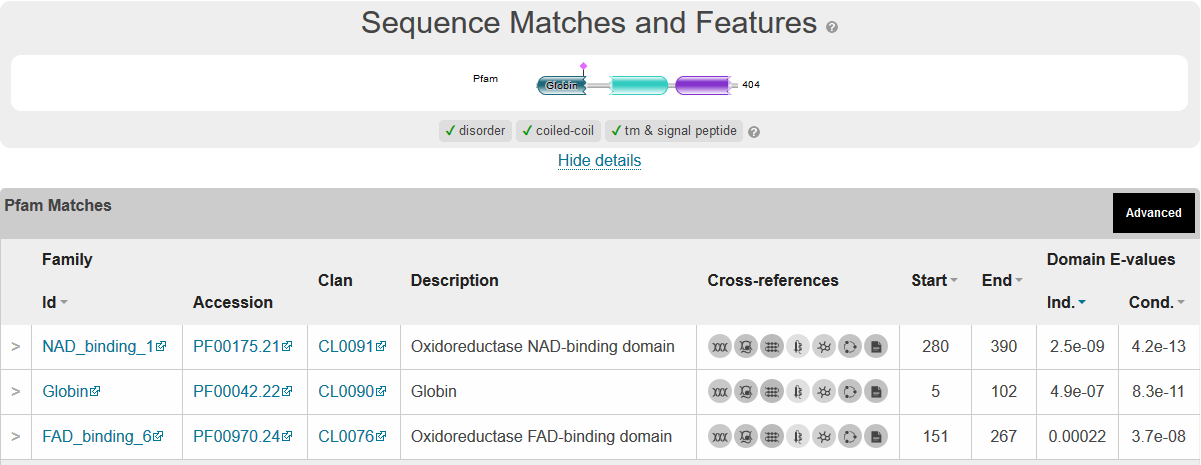

Fusarium graminearum NOD1 (FGSG_00765)
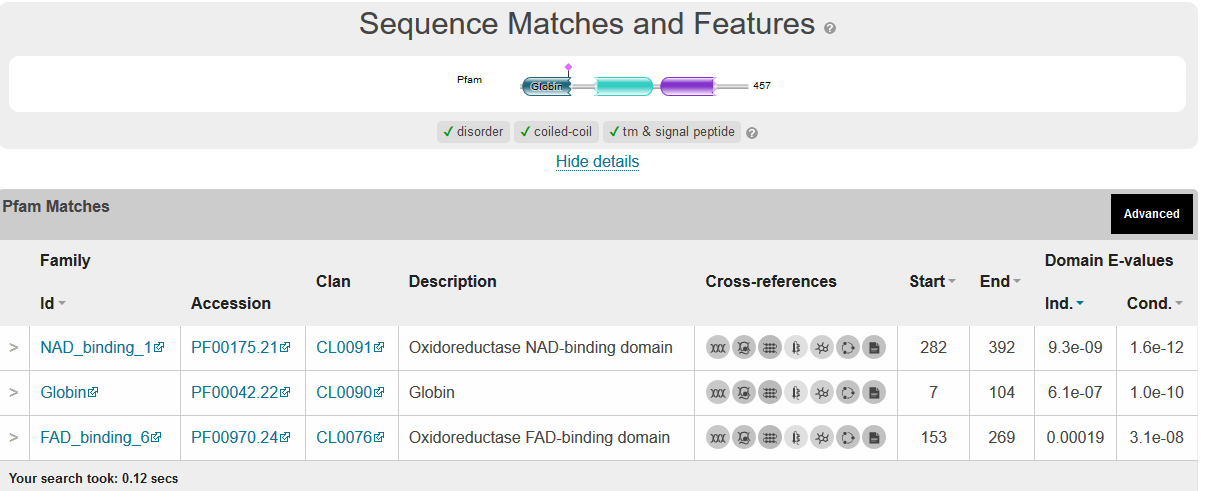

Fusarium graminearum NOD2 (FGSG_04458
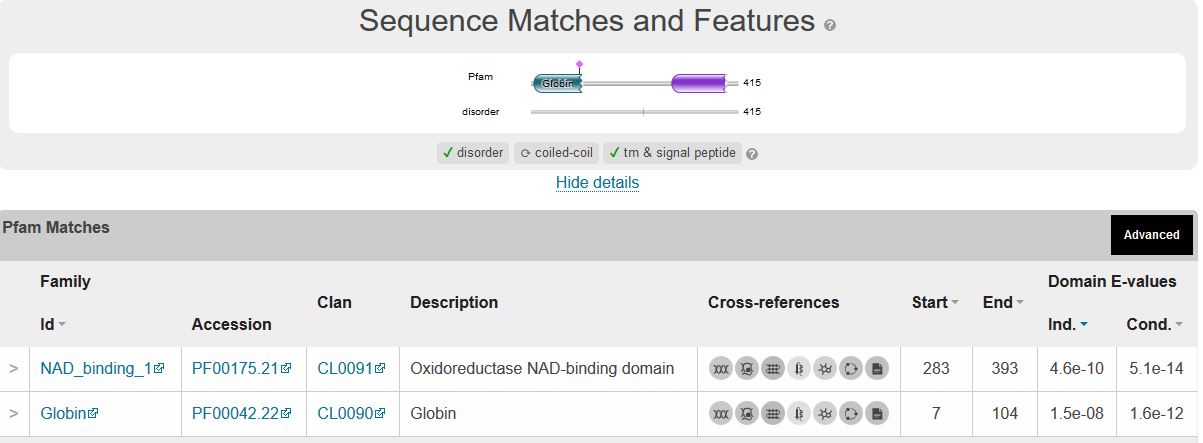
), Seem to lack a FAD binding domain

Since FGSG_04458 is well conserved over the whole length and the Identified domains, the central section of the protein was rerun using HMMER

>XP_011321142.1 hypothetical protein FGSG_04458 [Fusarium graminearum PH-1] AA105-282

GAIGEVLGSALTTEIKDAWVAAYGQLADIFIQREGQ

LYDAAGEWNSWRKFKIAKKEAENDSVTSFYLEPLDDKPLPKFLPGQYVSLQIPIPELDGLLQSRQFSLSE

APGSNHYRISVKLQGPEEEPSLEDLSAGKIPGLVCTRLHKRYNVGDEVELSPPAGEFFLNPADTSAAKKP

LV

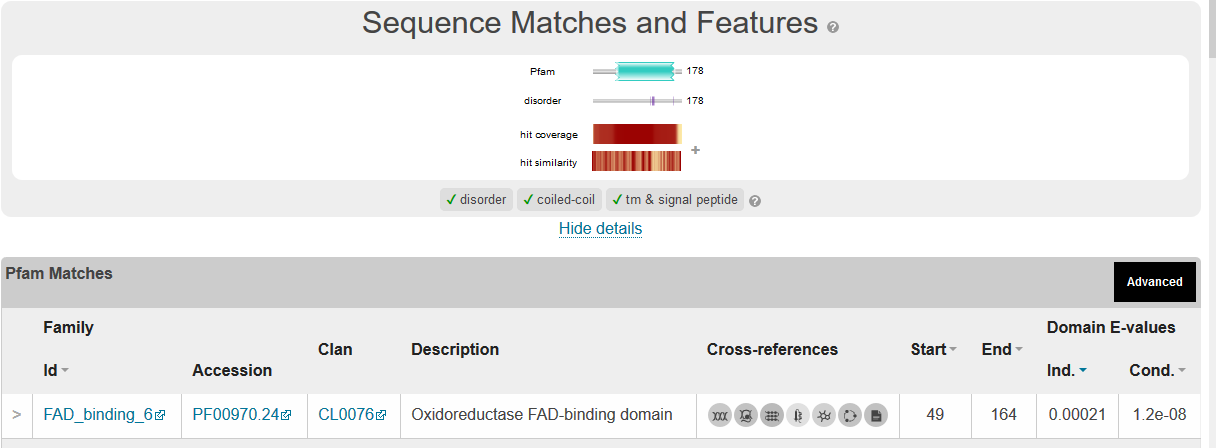

**Conclusion**

FGSG_00765 (FgNOD1) and FGSG_04458 (FgNOD2) are putative Nitric Oxide Dioxygenases based on sequence similarities to *C. albicans* and *F. sporotrichoides* NODs. FgNOD1 is the most similar, while FgNOD2 might have another or complementary function to the NODs in the other species.
