## Supplemental Files and Data for "Evolutionary conserved multifunctional nitric oxide synthesis proteins responding to bacterial MAMPs are located at the endoplasmic reticulum": Supplementary file SF6 Finding an ER localized CYP(NO) that can work together with NCP(NO) in Fg PH1.docx

```
Finding the ER localized HEME containg protein that can work together with an NADPH driven ER localized HEME reductase that we have found to be involved in NO production in the ER
Content										Page
Hypothesis											2
Get a template for the heme binding region					2
Localisation using WolfPsort did not give informative results		   5

Alignements with the heme-binding region in NOS2-mouse			    5

Test if the DeepLoc predictor tool correctly localises the 		   11
4 proteins we already have localised experimentally
since WolfPsort did not localise the NCP (NO) correctly

Localisation using the new DeepLoc-1.0 tool					22

Possible involvement the NO synthesis machinery in DON production?			39
Sterol synthesis is also implied for Cyp51A and Cyp51B and thus			40 (FGSG_01000), the putative heme protein involved in NO production
Conclusion and hypothesis for F. graminearum Er-localised heme protein involvement both in NO-production and ergosterol synthesis						41
```

**Hypothesis:** The heme-binding region reacting with arginine and making NO should have similarities with similar mammals regions. Find genes similar to this in Fg and Arabidopsis. Bot *A. thaliana* and mammals are known to produce NO from arginine. Start with Fg.

1. **Get a template for the heme-binding region**

Mouse sequence of NOS that has a heme-binding region that gets electrons from another similar NOS c-terminal heme reductase domain when two NOSes dimerise. Heme binding region highlighted

>NP_001300851.1 nitric oxide synthase, inducible isoform b [Mus musculus] Highlight Heme binding region

MNPKSLTRGPRDKPTPLEELLPHAIEFINQYYGSFKEAKIEEHLARLEAVTKEIETTGTYQLTLDELIFA

TKMAWRNAPRCIGRIQWSNLQVFDARNCSTAQEMFQHICRHILYATNNGNIRSAITVFPQRSDGKHDFRL

WNSQLIRYAGYQMPDGTIRGDAATLEFTQLCIDLGWKPRYGRFDVLPLVLQADGQDPEVFEIPPDLVLEV

TMEHPKYEWFQELGLKWYALPAVANMLLEVGGLEFPACPFNGWYMGTEIGVRDFCDTQRYNILEEVGRRM

GLETHTLASLWKDRAVTEINVAVLHSFQKQNVTIMDHHTASESFMKHMQNEYRARGGCPADWIWLVPPVS

GSITPVFHQEMLNYVLSPFYYYQIEPW

KTHIWQNEKLRPRRREIRFRVLVKVVFFASMLMRKVMASRVRA

TVLFATETGKSEALARDLATLFSYAFNTKVVCMDQYKASTLEEEQLLLVVTSTFGNGDCPSNGQTLKKSL

FMLRELNHTFRYAVFGLGSSMYPQFCAFAHDIDQKLSHLGASQLAPTGEGDELSGQEDAFRSWAVQTFRA

ACETFDVRSKHHIQIPKRFTSNATWEPQQYRLIQSPEPLDLNRALSSIHAKNVFTMRLKSQQNLQSEKSS

RTTLLVQLTFEGSRGPSYLPGEHLGIFPGNQTALVQGILERVVDCPTPHQTVCLEVLDESGSYWVKDKRL

PPCSLSQALTYFLDITTPPTQLQLHKLARFATDETDRQRLEALCQPSEYNDWKFSNNPTFLEVLEEFPSL

HVPAAFLLSQLPILKPRYYSISSSQDHTPSEVHLTVAVVTYRTRDGQGPLHHGVCSTWIRNLKPQDPVPC

FVRSVSGFQLPEDPSQPCILIGPGTGIAPFRSFWQQRLHDSQHKGLKGGRMSLVFGCRHPEEDHLYQEEM

QEMVRKRVLFQVHTGYSRLPGKPKVYVQDILQKQLANEVLSVLHGEQGHLYICGDVRMARDVATTLKKLV

ATKLNLSEEQVEDYFFQLKSQKRYHEDIFGAVFSYGAKKGSALEEPKATR

2. **Search FG proteomes for heme binding proteins.**
A. Fg's whole genome was searched for 50000 well described domains using the new domain search tool at NCBI.
B. A list of over 200 HEME binding genes was found.
C. All these were investigated for correlation of expression with the HEME reductase known to be involved in NO formation. The correlations were ranked.

A list of Fg proteins with heme-binding regions with the best correlation to the previously identified NOS is a heme reductase attached to ER (FGSG_09786). The 3 red-marked ones have positively been identified as putative ER localised using DeepLoc (see below). These proteins were Identified after a completely new domain search in the entire genome.

| ID | Correlation with heme reductase | Max/max reductaase |
| --- | --- | --- |
| FGSG_01000 | 0.516056849 | 1.009080384 |
| FGSG_09373 | 0.425021553 | 5.084570499 |
| FGSG_01972 | 0.295199455 | 0.264709236 |
| FGSG_10305 | 0.286718586 | 2.102462605 |
| FGSG_10881 | 0.269646267 | 6.57440677 |
| FGSG_11195 | 0.247331425 | 0.00829104 |
| FGSG_02982 | 0.237978559 | 0.019897793 |
| FGSG_07977 | 0.237185522 | 0.000885604 |
| FGSG_12534 | 0.234464981 | 0.185611322 |

Sequences for these proteins

>FGSG_01000T0 | FGSG_01000 | Fusarium graminearum PH-1 cytochrome P450 Cyp51B (527 aa)

MGLLQELAGHPLAQQFQELPLGQQVGIGFAVFLVLSVVLNVLNQLLFRNPNEPPMVFHWFPFVGSTITYG

MDPPTFFRENRAKHGDVFTFILLGKKTTVAVGPAGNDFILNGKLKDVCAEEIYTVLTTPVFGKDVVYDCP

NAKLMEQKKFMKIALTTEAFRSYVPIISSEVRDYFKRSPDFKGKSGIADIPKKMAEITIFTASHALQGSA

IRSKFDESLAALYHDLDMGFTPINFMLHWAPLPWNRKRDHAQRTVAKIYMDTIKERRAKGNNESEHDMMK

HLMNSTYKNGIRVPDHEVAHMMIALLMAGQHSSSSTSSWIMLRLAQYPHIMEELYQEQVKNLGADLPPLT

YEDLAKLPLNQAIVKETLRLHAPIHSIMRAVKSPMPVPGTKYVIPTSHTLLAAPGVSATDSAFFPNPDEW

DPHRWEADSPNFPRMASKGEDEEKIDYGYGLVSKGSASPYLPFGAGRHRCIGEHFANAQLQTIVAEVVRE

FKFRNVDGGHTLIDTDYASLFSRPLEPANIHWERRQ

>FGSG_01972T0 | FGSG_01972 | Fusarium graminearum PH-1 bifunctional P-450:NADPH-P450 reductase (1070 aa)

MAESVPIPEPPGYPLIGNLGEFKTNPLNDLNRLADTYGPIFRLHLGSKTPTFVSSNAFINEVCDEKRFKK

TLKSVLSVVREGVHDGLFTAFEDEPNWGKAHRILIPAFGPLSIRNMFPEMHEIANQLCMKLARHGPHTPV

DASDNFTRLALDTLALCAMDFRFNSYYKEELHPFIEAMGDFLLESGNRNRRPAFAPNFLYRAANDKFYAD

IALMKSVADEVVATRKQNPTDRKDLLAAMLEGVDPQTGEKLSDDNITNQLITFLIAGHETTSGTLSFAMY

HLLKNPEAYNKLQKEIDEVIGRDPVTVEHLTKLPYLSAVLRETLRISSPITGFGVEAIEDTFLGGKYLIK

KGETVLSVLSRGHVDPVVYGPDAEKFVPERMLDDEFARLNKEFPNCWKPFGNGKRACIGRPFAWQESLLA

MALLFQNFNFTQTDPNYELQIKQNLTIKPDNFFFNCTLRHGMTPTDLEGQLAGKGATTSIASHIKAPAAS

KGAKASNGKPMAIYYGSNSGTCEALANRLASDAAGHGFSASVIGTLDQAKQNLPEDRPVVIVTASYEGQP

PSNAAHFIKWMEDLAGNEMEKVSYAVFGCGHHDWVDTFLRIPKLVDTTLEQRGGTRLVPMGSADAATSDM

FSDFEAWEDTVLWPSLKEKYNVTDDEASGQRGLLVEVTTPRKTTLRQDVEEALVVSEKTLTKTGPAKKHI

EIQLPSGMTYKAGDYLAILPLNPRKTVSRVFRRFSLAWDSFLKIQSDGPTTLPINIAISAFDVFSAYVEL

SQPATKRNILALSEATEDKATIQELEKLAGDAYQEDVSAKKVSVLDLLEKYPAVALPISSYLAMLPPMRV

RQYSISSSPFADPSKLTLTYSLLDAPSLSGQGRHVGVATNFLSQLIAGDKLHISVRASSAAFHLPSDPET

TPIICVAAGTGLAPFRGFIQERAAMLAAGRKLAPALLFFGCRDPENDDLYAEELARWEQMGAVDVRRAYS

RATDKSEGCKYVQDRIYHDRADVFKVWDQGAKVFICGSREIGKAVEDICVRLAMERSEATQEGKGATEEK

AREWFERSRNERFATDVFD

>FGSG_02982T0 | FGSG_02982 | Fusarium graminearum PH-1 hypothetical protein (536 aa)

MSKHTLYETSILGPARNIFSLVSPSVFLLLVVPASTLSLWTLASYFTSPLKKYPGPFLAKFTRLWYMYQA

STGDSHLVLERLHKRYGPIVRITPDIIDVDIPEIINTIFSTKDDWLKTPFYHGSSALVNGHIVLNTFSQT

DPVKHKKGRQPIAKLYSSAGVSTLEPHMNKVINQLCDELEKRFTGHNAGQVCSLGQWILFYAWDVVGAIT

FSQPIGYLKKGCDFDGTLKNADKAMDYFTVVGTMPFLDRIFDKNPVFHMGPPGFNTSTEISVKHLIDRYQ

GNDKENHDPAHPDFLDKFIEIKNSKPDEADDAQIISWLMVNMIAGADTTAITIRSVLYFSLKHPRVWKRL

TEEILRAGFQQVPAYKDVKALPYTDAVCREALRMLPGVAMTMERFVPKEGFVLPNGDFLPGGTIVGMNPY

IVARNKSVYGDDADDFRPERWMRSEDETEEQYQIRLLAMNQADLSFGGGSRICIGKYIGLFQTYKVIAIL

LTRFEIELADPNKEWKVTNSWFPRQEGLEARIRKRTGSRLPKSSY

>FGSG_09373T0 | FGSG_09373 | Fusarium graminearum PH-1 hypothetical protein (513 aa)

MSRPQLLFLSCIGFAVALTAMLYSNIIHSSTLTSVLSKSMSSSVRPIVVVGSGLAGLSASYEALQRGAPS

VHLLDRAPKPGGNSIKASSGINGAGTKYQRAAGVESDTTFYSDSVRSAGSRFHLAQPPVDREGLVTKLTT

ESAAAVEWLVDEIGVDLSVVAPLGGHSVARTHRGAGKTPPGAAIIIALLNKLKENKKFSITNLAEVKALL

RENDEVKGVEYEFEGQKHNLEGSVLFASGGFAGDATGLLARYRPDLKGIPSTNDERPGSHDILTSVGAEL

LDMDSVQIHPTGFVDPASPNTMLKFLAAEMLRGEGGILLSPDGSRFVNEMDTREHVSNAIMKLPTATDGD

GVIKQWDITILLDPGASAASANHISFYEWKGLLKKVKVRDLTSAQIAAVDKYAQSVADGSADEFGRTQRG

RWTLPAGEKNRDQDIYIGRVTPITHFTMGGVAIDEKARVLKKSGDKLVPIPGLFAAGEITGGIHGDNRLG

GSSLLECVVYGRTAGAEIVGSA

>FGSG_10305T0 | FGSG_10305 | Fusarium graminearum PH-1 hypothetical protein (105 aa)

MRPTQMLRSGAADPKNGHYIGNWGHFGGEKQRGIITYGLSANRQNPWAGSFNDAIFNTFRRTKGQIFFWL

PPMVAGYWMMSWAIERSEYLNSKAGRAEFGDEEE

>FGSG_10881T0 | FGSG_10881 | Fusarium graminearum PH-1 hypothetical protein (107 aa)

MAGGDIKKGANLFKTRCAQCHTVEKDGGNKIGPALHGLWGRKTGSVEGYSYTDANKQKGIEWNDDTLFEY

LENPKKYIPGTKMAFGGLKKAKDRNDLIAYLKDSTK

>FGSG_11195T0 | FGSG_11195 | Fusarium graminearum PH-1 hypothetical protein (750 aa)

MAAAHRHRKSVAADHPVRSSQNVEFLTDKEISDFLDDLDHDNDGHINYEEVERKLDQEHANLVPKPSAHH

VISTDHSDDDRTRHAFLRRMMGDSGVDQIPRDEFAKMVKEWKIPSLKQAKKEEEEDKSYIKRLPGWRRIR

SYWAVHGPEIVFLGVVISMQLAFGIWQLVKYQTTPGYRAAFGWGVVMAKTCAGALYPTFFFLILSMSRYF

STWLRRSYHISRFFNWDLSQEFHIRISCVAILLATLHAIGHLTGSFVHGSDPANEDAVAEALGPDKVPRP

YIDYVRSLPGFTGITALGLFWILCLLSIPQVRRWNYEVFQLGHLLMFPIIGLMMAHGTAALLQWPMFGYF

LAFPTLLVLVERTVRVGLGFHRIKATMKVLDKETVEVTAIIPSERLWKYKAGQYIFLQVPKISFFQWHPF

TVSFCRGNKMMLHIKTDGNWTAKLRELGGDSGESEIEVGINGPFGAPAQRFYDFNHSIIIGAGIGVTPFS

GILADLQYNDDLDHGGPNHEVDHHRHDSEATAIPQAARRSDSSSSDEATTSDNVPETPTRQGSVGPDLIN

KEKQPQADKAGSFAEDYRRVDFHWMVRERNYLLWLSDLLNDVSMSQDWHREHEDKPHLDIRINTHVTAKQ

KKISTHVYRWLLEMHRTDEHPASPLTGLLNPTHFGRPDFDLILDEHYEEMLKFRASKRTSTRNKEDENYE

EDEELKVGVFYCGAPVVGEILADKCRELTLRGWQDGSKLEYHFMIEVFG

>FGSG_12534T0 | FGSG_12534 | Fusarium graminearum PH-1 hypothetical protein (507 aa)

MALLSIVNVALLGVAYFVACAVWQVVKYRFLHPLAKFPGNFWGSVTRLWITYHNVEADECETFQELHKKH

GPIIRITPTMLLVSDATELPKIYNRHANKSKHYITGSFGKDESLFNMQDSVMHAKYRKVAAPLYSLTNIK

KMEPLIDNNMSAWMSRLQRDFAATNKPFDFAPFSVYLAYDVISEVGFGAPFGFVKEGKDVEGLIQGFHDG

LTPFGIMARLYPFTNWVKSTFLGKYMVASPEQDSGIGILMRFRDRLIEQRFKDIENGSTGGRIDLLQTFI

EARDEDDKPLDINYIKAEILLVLLAGADTTGTAFQAMMVHILTNPSVYKKLLAEIDEATAAGNLSEMPQY

DEVVEHLPYYIACVKEAMRLTPSAPNIFPRIVPQGGLEICGHFVPEGTEVTCNPWLVHRDPNIYGDDAEI

FKPERWLDADKAKIYNKYSMGFGYGPRVCLGQDIARMELYKGPLLFLRSFNVEWVDETKRGKYVINGGVS

YFEDMNIKISRRDAAA

3. **Localisation using WolfPsort did not give informative results. None of the heme proteins was predicted to be localised at the ER membrane**

FGSG_01000T0 [details](https://wolfpsort.hgc.jp/results/fGL505d3a31e2b10991772b1cb33187625e.detailed1.html#FGSG_01000T0) cyto: 12, cyto_nucl: 8, mito: 6, E.R.: 4, nucl: 2, plas: 1, golg: 1, vacu: 1
FGSG_01972T0 [details](https://wolfpsort.hgc.jp/results/fGL505d3a31e2b10991772b1cb33187625e.detailed2.html#FGSG_01972T0) cyto: 14, cyto_nucl: 10, nucl: 4, pero: 4, cysk: 2, mito: 1, golg: 1, vacu: 1
FGSG_02982T0 [details](https://wolfpsort.hgc.jp/results/fGL505d3a31e2b10991772b1cb33187625e.detailed3.html#FGSG_02982T0) extr: 9, mito: 4, plas: 4, E.R.: 4, nucl: 2, cyto: 2, pero: 1, golg: 1
FGSG_09373T0 [details](https://wolfpsort.hgc.jp/results/fGL505d3a31e2b10991772b1cb33187625e.detailed4.html#FGSG_09373T0) extr: 10, mito: 9, cyto: 7.5, cyto_nucl: 4.5
FGSG_10305T0 [details](https://wolfpsort.hgc.jp/results/fGL505d3a31e2b10991772b1cb33187625e.detailed5.html#FGSG_10305T0) cyto: 12, mito: 7, nucl: 4, pero: 3, extr: 1 FGSG_10881T0 [details](https://wolfpsort.hgc.jp/results/fGL505d3a31e2b10991772b1cb33187625e.detailed6.html#FGSG_10881T0) mito: 18, cyto: 7, extr: 1, pero: 1

FGSG_11195T0 [details](https://wolfpsort.hgc.jp/results/fGL505d3a31e2b10991772b1cb33187625e.detailed7.html#FGSG_11195T0) plas: 22, cyto: 3, mito: 1, pero: 1
FGSG_12534T0 [details](https://wolfpsort.hgc.jp/results/fGL505d3a31e2b10991772b1cb33187625e.detailed8.html#FGSG_12534T0) extr: 8, E.R.: 8, cyto: 5, mito: 2, plas: 2, pero: 1, golg: 1

4. **Alignements with the heme binding region in NOS2-mouse**

```
NP_001300851.1      ------MNPKSLTRGPRDKP-------------------TPLEELL---PHAIEFINQYY
FGSG_01000T0        MGLLQELAGHPLAQQFQELPLGQQVGIGFAVFLVLSVVLNVLNQLLFRNPNEPPMVFHWF
                          :  :.*:.  .: *                   . *::**   *:   :: :::

NP_001300851.1      ---GS-----------FKEAKIEEHLARLEAVTKEIETTGTYQLTLDELIFATKMAWRNA
FGSG_01000T0        PFVGSTITYGMDPPTFFRENR-AKHGDVFTFILLGKKTTVAVGPAGNDFILNGKLKDVCA
                       **           *.* .  :*   :  :    :** :   : :::*:  *:    *

NP_001300851.1      PRCIGRIQ---WSNLQVFDARNCS-TAQEMFQHICRHI-LYATNNGNIRSAITVFPQRS-
FGSG_01000T0        EEIYTVLTTPVFGKDVVYDCPNAKLMEQKKFMKIALTTEAFRSYVPIISSEVRDYFKRSP
                          :    :.:  *:*. *..   *: * :*.     : :    * * :  : :**

NP_001300851.1      -----DGKHDFRLWNSQLIRY-AGYQMPDGTIRGD----------AATLEFTQLCIDLGW
FGSG_01000T0        DFKGKSGIADIPKKMAEITIFTASHALQGSAIRSKFDESLAALYHDLDMGFTPINFMLHW
                         .*  *:  . :::  : *.: : ..:**..             : ** : : * *

NP_001300851.1      KPRYGRFDVLPLVLQADGQDPEVFEIPPDLVLEVTME---HPKYEWFQEL-------GLK
FGSG_01000T0        AP-------LPWNRKRDHAQRTVAKIYMDTIKERRAKGNNESEHDMMKHLMNSTYKNGIR
                     *       **   : *  :  * :*  * : *   :    .::: :: *       *:.

NP_001300851.1      WYALPAVANMLLEVGGLEFPACPFNGWYMGTEIGVRDFCDTQRYNILEEVGRRMGLETHT
FGSG_01000T0        VPDHEVAHMMIALLMAGQHSSSSTSSWIM---LRLAQYPHIME-ELYQEQVKNLGADLPP
                         ..  *:  : . :..:.. ..* *   : : ::      :: :*  ..:* :  .

NP_001300851.1      LASLWKDRAVTEINVAVLHSFQKQNVTIMDHHTASESFMKHMQNEY---------RARGG
FGSG_01000T0        LT--YEDLAKLPLNQAIVKETLRLHAPIHSIMRAVKSPMPVPGTKYVIPTSHTLLAAPGV
                    *:  ::* *   :* *:::.  . :..* .   * :* *    .:*          * *

NP_001300851.1      CPADWIWLVPP-----------------------------------VSGSITPVFH----
FGSG_01000T0        SATDSAFFPNPDEWDPHRWEADSPNFPRMASKGEDEEKIDYGYGLVSKGSASPYLPFGAG
                    ..:*  ::  *                                    .** :* :

NP_001300851.1      ---------------------------------QEMLNYVLSPFYYYQIEP----W----
FGSG_01000T0        RHRCIGEHFANAQLQTIVAEVVREFKFRNVDGGHTLIDTDYASLFSRPLEPANIHWERRQ
                                                     : :::   :.::   :**    *


_________________________________________________________________________
NP_001300851.1      MNPKSLTRGPRDKPT--PLEELLPHAIEFINQY---YGSFKEAKIEEHLARL---EAVTK
FGSG_01972T0        MAESVPIPEPPGYPLIGNLGEFKTNPLNDLNRLADTYGPIFRLHLGSKTPTFVSSNAFIN
                    *  .     * . *    * *: .:.:: :*.    **.:   :: .: . :   :*. :

NP_001300851.1      EIETTGTYQLTLDELI--------------FATKMAWRNAPRCI----GRIQWSN-----
FGSG_01972T0        EVCDEKRFKKTLKSVLSVVREGVHDGLFTAFEDEPNWGKAHRILIPAFGPLSIRNMFPEM
                    *:     :: **..::              *  :  * :* * :    * :.  *

NP_001300851.1      -----------------------------------LQVFDAR-NCSTAQEMFQHI-----
FGSG_01972T0        HEIANQLCMKLARHGPHTPVDASDNFTRLALDTLALCAMDFRFNSYYKEELHPFIEAMGD
                                                       * .:* * *.   :*:. .*

NP_001300851.1      --------------CRHILYATNNGNIRSAITVF--------------------------
FGSG_01972T0        FLLESGNRNRRPAFAPNFLYRAANDKFYADIALMKSVADEVVATRKQNPTDRKDLLAAML
                                  . ::** : *.:: : *:::

NP_001300851.1      ----PQRSDGKHDFRLWNSQLIRY--AGYQMPDGT-------------------------
FGSG_01972T0        EGVDPQTGEKLSDDNI-TNQLITFLIAGHETTSGTLSFAMYHLLKNPEAYNKLQKEIDEV
                        ** .:   * .: ..*** :  **:: ..**

NP_001300851.1      IRGDAATLE------------------------FTQLCID---LGWKPRYGRFDVLPLVL
FGSG_01972T0        IGRDPVTVEHLTKLPYLSAVLRETLRISSPITGFGVEAIEDTFLGGKYLIKKGETVLSVL
                    *  *..*:*                        *   .*:   ** *    . :.:  **

NP_001300851.1      QADGQDPEVF-----EIPPDLVLE------------------------------------
FGSG_01972T0        SRGHVDPVVYGPDAEKFVPERMLDDEFARLNKEFPNCWKPFGNGKRACIGRPFAWQESLL
                    . .  ** *:     :: *: :*:

NP_001300851.1      ----------VTMEHPKYE-------------WFQELGLKWYALP---------------
FGSG_01972T0        AMALLFQNFNFTQTDPNYELQIKQNLTIKPDNFFFNCTLRHGMTPTDLEGQLAGKGATTS
                              .*   *:**             :* :  *..   *

NP_001300851.1      ----------------------------------AVANMLL-EVGGLEFP----------
FGSG_01972T0        IASHIKAPAASKGAKASNGKPMAIYYGSNSGTCEALANRLASDAAGHGFSASVIGTLDQA
                                                      *:** *  :..*  *.

NP_001300851.1      ------------------------ACPFNGW---YMGTEI----------GVRDFCDT--
FGSG_01972T0        KQNLPEDRPVVIVTASYEGQPPSNAAHFIKWMEDLAGNEMEKVSYAVFGCGHHDWVDTFL
                                            *. *  *     *.*:          * .*: **

NP_001300851.1      ------------------------------------------------QRYNILEE----
FGSG_01972T0        RIPKLVDTTLEQRGGTRLVPMGSADAATSDMFSDFEAWEDTVLWPSLKEKYNVTDDEASG
                                                                    :.**: ::

NP_001300851.1      ------------------------------------------------------------
FGSG_01972T0        QRGLLVEVTTPRKTTLRQDVEEALVVSEKTLTKTGPAKKHIEIQLPSGMTYKAGDYLAIL


NP_001300851.1      ----------VGRRMGLETHTLASLWKDRAVT---EINVAVLHSFQ----------KQNV
FGSG_01972T0        PLNPRKTVSRVFRRFSLAWDSFLKIQSDGPTTLPINIAISAFDVFSAYVELSQPATKRNI
                              * **:.*   :: .:..* ..*   :* ::.:  *.          *.*:

NP_001300851.1      TIMDHHTASESFMKHMQ----------------------NEYRARGGCPADWIWLVPPV-
FGSG_01972T0        LALSEATEDKATIQELEKLAGDAYQEDVSAKKVSVLDLLEKYPAVALPISSYLAMLPPMR
                      :.  * .:: :: ::                      ::* * .   :.:: ::**:

NP_001300851.1      ----SGSITPVFHQEMLNYV----------------------------------------
FGSG_01972T0        VRQYSISSSPFADPSKLTLTYSLLDAPSLSGQGRHVGVATNFLSQLIAGDKLHISVRASS
                        * * :*.   . *. .

NP_001300851.1      ----------------------LSPF----------------------------------
FGSG_01972T0        AAFHLPSDPETTPIICVAAGTGLAPFRGFIQERAAMLAAGRKLAPALLFFGCRDPENDDL
                                          *:**

NP_001300851.1      YYYQIEPW----------------------------------------------------
FGSG_01972T0        YAEELARWEQMGAVDVRRAYSRATDKSEGCKYVQDRIYHDRADVFKVWDQGAKVFICGSR
                    *  ::  *

NP_001300851.1      --------------------------------------------------
FGSG_01972T0        EIGKAVEDICVRLAMERSEATQEGKGATEEKAREWFERSRNERFATDVFD


_________________________________________________________________________________


NP_001300851.1      MNPKSLTR----GPRDK-----------------------------PTPLEELL-PHAIE
FGSG_02982T0        MSKHTLYETSILGPARNIFSLVSPSVFLLLVVPASTLSLWTLASYFTSPLKKYPGPFLAK
                    *. ::*      **  :                             .:**::   *.  :

NP_001300851.1      FINQYYGSFKEAKIEEHLARLEAVTKEIE-----TTGTYQLTLDEL---IFATKMAWRNA
FGSG_02982T0        FTRLWY-MYQASTGDSHLV-LERLHKRYGPIVRITPDIIDVDIPEIINTIFSTKDDWLKT
                    * . :*  :: :. :.**. ** : *        *..  :: : *:   **:**  * ::

NP_001300851.1      PRCIGRIQWSN----LQVFDARNCSTAQEMFQHICRHILYATN---------NGNIRSAI
FGSG_02982T0        PFYHGSSALVNGHIVLNTFSQTDPVKHKKGRQPIAK--LYSSAGVSTLEPHMNKVINQLC
                    *   *     *    *:.*.  :  . ::  * *..  **::          *  *..

NP_001300851.1      TVFPQRSDGKHD-----------FRLWN-------SQLIRY--AGYQMPDGTIRGDAATL
FGSG_02982T0        DELEKRFTGHNAGQVCSLGQWILFYAWDVVGAITFSQPIGYLKKGCDF-DGTLKNADKAM
                      : :*  *::            *  *:       ** * *   * :: ***:..   ::

NP_001300851.1      EFTQLCIDLGWKPRYGRFDVLP-------------------LVLQADGQDPEVFE-IPPD
FGSG_02982T0        DYFTVVGTMPFLDRI--FDKNPVFHMGPPGFNTSTEISVKHLIDRYQGNDKENHDPAHPD
                    ::  :   : :  *   **  *                   *: . :*:* * .:   **

NP_001300851.1      LV---LEVTMEHPK-------YEWF----------QELGLK---WYAL------PAVANM
FGSG_02982T0        FLDKFIEIKNSKPDEADDAQIISWLMVNMIAGADTTAITIRSVLYFSLKHPRVWKRLTEE
                    ::   :*:. .:*.        .*:            : :.   :::*        :::

NP_001300851.1      LLEVGGLEFPAC------PFNG--------WYMGTEIGVRDFCDTQRYNI----LEEVGR
FGSG_02982T0        ILRAGFQQVPAYKDVKALPYTDAVCREALRMLPGVAMTMERFVPKEGFVLPNGDFLPGGT
                    :* .*  :.**       *:..           *. : :  *  .: : :    :   *

NP_001300851.1      RMGLETHTLASLWKDRAVTEINVAVLHSFQKQNVTIMDHHTASESFMKHM---QNEYRAR
FGSG_02982T0        IVGMNPYIVA---RNKSVYGDDA---DDFRPERWMRSEDETEEQYQIRLLAMNQADLSFG
                     :*::.: :*   .:.:*   :.    .*. :.    :  * .:  :. :   * :

NP_001300851.1      GG---CPADWIWLVPP--VSGSITPVFHQEMLN----YVLSPFYYYQIE-----------
FGSG_02982T0        GGSRICIGKYIGLFQTYKVIAILLTRFEIELADPNKEWKVTNSWFPRQEGLEARIRKRTG
                    **   * ..:* *. .  * . : . *  *: :    : ::  :: . *

NP_001300851.1      ------PW
FGSG_02982T0        SRLPKSSY
                          .:

_________________________________________________________________________________
NP_001300851.1      -MNPKSLTRGPRDKPTPLEELLPHAIEFINQYYGSFKEAK---------IEEHLARLEAV
FGSG_09373T0        MSRPQLLFLSCIGFAVALTAMLYSNIIHSSTLTSVLSKSMSSSVRPIVVVGSGLAGLSAS
                      .*: *  .  . ...*  :*   * . .   . :.::          : . ** *.*

NP_001300851.1      TKEIETTGTYQLTLDEL--------------------------IFATKMAWRNAPRCIG-
FGSG_09373T0        YEALQRGAPSVHLLDRAPKPGGNSIKASSGINGAGTKYQRAAGVESDTTFYSDSVRSAGS
                     : ::  ..    **                            : : .  : :: *. *

NP_001300851.1      RIQWSNLQV--------FDARNCSTAQEMFQHI----------CRHILYATNNGNIRSA-
FGSG_09373T0        RFHLAQPPVDREGLVTKLTTESAAAVEWLVDEIGVDLSVVAPLGGHSVARTHRGAGKTPP
                    *:: ::  *        : : ..::.: :.: *            * :  *:.*  .:.

NP_001300851.1      ----ITVFPQRSDGKHDFRLWNSQLIR---------------YAGYQ--------MPDGT
FGSG_09373T0        GAAIIIALLNKLKENKKFSITNLAEVKALLRENDEVKGVEYEFEGQKHNLEGSVLFASGG
                        * .: :. . ::.* : *   :.               : * :        :..*

NP_001300851.1      IRGDAATLEFTQLCIDLGWKP-----RYGRFDVLPLVLQADGQDPEVFEIPPDLVLEVTM
FGSG_09373T0        FAGDATGL-LARYRPDLKGIPSTNDERPGSHDILTSV-GAELLDMDSVQIHPTGFVDPAS
                    : ***: * ::.   **   *     * * .*:*. *  *:  * : .:* *  .:: :

NP_001300851.1      EHPKYEWFQELGLKWYALPAVANMLLEVGGLEFPACPFNGWYMGTEIGVRDFCDT-----
FGSG_09373T0        PNTMLKFL------------AAEMLRGEGGILLSP---DGSRFVNEMDTREHVSNAIMKL
                     :.  :::            .*:**   **: :..   :*  : .*:..*:. ..

NP_001300851.1      ----------QRYN--ILEEVGRRMGLETHTLASLWKD-------RAVTEINVAVLHSFQ
FGSG_09373T0        PTATDGDGVIKQWDITILLDPGASAASANHISFYEWKGLLKKVKVRDLTSAQIAAVDKYA
                              :.::  ** : *   .  .*     **.       * :*. ::*.: .:

NP_001300851.1      KQ---------------------------------NVTIMDHHTASESFMKHMQNEYRAR
FGSG_09373T0        QSVADGSADEFGRTQRGRWTLPAGEKNRDQDIYIGRVTPITHFTMGGVAIDEKARVLKKS
                    :.                                 .** : *.* .   :.   .  .

NP_001300851.1      GG--CPADWIWLVPPVSGSI---TPVFHQEMLNYVLSPFYYYQIEPW------
FGSG_09373T0        GDKLVPIPGLFAAGEITGGIHGDNRLGGSSLLECVV----YGRTAGAEIVGSA
                    *.   *   :: .  ::*.*   . :  ..:*: *:    * .

____________________________________________________________________________________________

NP_001300851.1      MNPKSLTRGPRDKPTPLEELLPHAIEFINQYYGSFKEAKIEEHLARLEAVTKEIETTGTY
FGSG_10305T0        MRPTQMLRSGAADPKN------------GHYIGNWGHFGGEKQRGII-----------TY
                    *.*..: *.   .*.             .:* *.:     *:: . :           **

NP_001300851.1      QLTLDELIFATKMAWRNAPRCIGRIQWSNLQVFDARNCSTAQEMFQHICRHILYATNNGN
FGSG_10305T0        GLSAN----------RQNP-------WA------------------------------GS
                     *: :          *: *       *:                              *.

NP_001300851.1      IRSAITVFPQRSDGKHDFRLWNSQLIRYAGYQMPDGTIRGDAATLEFTQLCIDLGWKPRY
FGSG_10305T0        FNDAIFNTFRRTKGQIFF--WLPPMV--AGYWM--------------------MSW----
                    :..**    .*:.*:  *  * . ::  ***.*                    :.*

NP_001300851.1      GRFDVLPLVLQADGQDPEVFEIPPDLVLEVTMEHPKYEWFQELGLKWYALPAVANMLLEV
FGSG_10305T0        ------------------------------AIERSE------------------------
                                                  ::*..:

NP_001300851.1      GGLEFPACPFNGWYMGTEIGVRDFCDTQRYNILEEVGRRMGLETHTLASLWKDRAVTEIN
FGSG_10305T0        -------------YLNSKAGRAEFGDEEE-------------------------------
                                 *:.:: *  :* * :

NP_001300851.1      VAVLHSFQKQNVTIMDHHTASESFMKHMQNEYRARGGCPADWIWLVPPVSGSITPVFHQE
FGSG_10305T0        ------------------------------------------------------------


NP_001300851.1      MLNYVLSPFYYYQIEPW
FGSG_10305T0        -----------------


___________________________________________________________________________________
NP_001300851.1      MNPKSLTRGPRDKPTPLEELLPHAIEFINQYYGSFKEAKIEEHLARLEAVTKEIETTGTY
FGSG_10881T0        MAGGDIKKGAN-------------------------------------------------
                    *   .:..*..

NP_001300851.1      QLTLDELIFATKMAWRNAPRCIGRIQWSNLQVFDARNCSTAQEMFQHICRHILYATNNGN
FGSG_10881T0        -------LFKTRCA----------------------QCHTVEK-------------DGGN
                           :* *. *                      :* *.::             :.**

NP_001300851.1      IRSAITVFPQRSDGKHDFRLWNSQLIRYAGYQMPDGTIRGDAATLEFTQLCIDLGWKPRY
FGSG_10881T0        -----------KIGPALHGLWGRKTGSVEGYSYTDANKQK------------GIEWN---
                               . *   . **. :     **. .*.. .             .: *:

NP_001300851.1      GRFDVLPLVLQADGQDPEVFEIPPDLVLEVTMEHPKYEWFQELGLKWYALPAVANMLLEV
FGSG_10881T0        ---------------DDTLFEY---------LENPK--------------KYIPGTKMAF
                                   *  :**          :*:**                :..  : .

NP_001300851.1      GGLEFPACPFNGWYMGTEIGVRDFCDTQRYNILEEVGRRMGLETHTLASLWKDRAVTEIN
FGSG_10881T0        GGLKKA---------------------------------------------KDR------
                    ***: .                                             ***

NP_001300851.1      VAVLHSFQKQNVTIMDHHTASESFMKHMQNEYRARGGCPADWIWLVPPVSGSITPVFHQE
FGSG_10881T0        ---------------------NDLIAYLKDSTK---------------------------
                                         :.:: ::::. .

NP_001300851.1      MLNYVLSPFYYYQIEPW
FGSG_10881T0        -----------------

___________________________________________________________________________________

NP_001300851.1      MNPKSLTRGPRDKPTPLEELLPHAIEF-----INQYY--------GSFKEAKIEEHL---
FGSG_11195T0        MAAAHRHRKSVAADHPVRS--SQNVEFLTDKEISDFLDDLDHDNDGHINYEEVERKLDQE
                    * .    * .     *: .  .: :**     *.::         * ::  ::* :*

NP_001300851.1      -----------------------ARLEAVTKEIETTGTYQLTLDELIFATKMAWR-----
FGSG_11195T0        HANLVPKPSAHHVISTDHSDDDRTRHAFLRRMMGDSGVDQIPRDEFAKMVK-EWKIPSLK
                                           :*   : . :  :*. *:. **:   .*  *.

NP_001300851.1      -------NAPRCIGRIQ-WSNLQVFDARN--------CSTAQEMFQHICRHILYATNNGN
FGSG_11195T0        QAKKEEEEDKSYIKRLPGWRRIRSYWAVHGPEIVFLGVVISMQLAFGIWQLVKYQTTPG-
                           :    * *:  * .:. : * :           : ::   * . : * *. *

NP_001300851.1      IRSAI---TVFPQRSDG-------------KHDFRLW---NSQLIRYAGYQMPD------
FGSG_11195T0        YRAAFGWGVVMAKTCAGALYPTFFFLILSMSRYFSTWLRRSYHISRFFNWDLSQEFHIRI
                     *:*:   .*:.: . *             .. *  *   . :: *: .:::.:

NP_001300851.1      -------------GTIRG------DAATLEFTQLCIDLGWKPR-----------------
FGSG_11195T0        SCVAILLATLHAIGHLTGSFVHGSDPANEDAVAEALGPDKVPRPYIDYVRSLPGFTGITA
                                 * : *      *.*. : .  .:. .. **

NP_001300851.1      YGRFDVLPL--VLQADGQDPEVFEIPPDLVLEVT-----------MEHPKYEWFQEL---
FGSG_11195T0        LGLFWILCLLSIPQVRRWNYEVFQLGHLLMFPIIGLMMAHGTAALLQWPMFGYFLAFPTL
                     * * :* *  : *.  .: ***::   *:: :            ::.* : :*  :

NP_001300851.1      ----------GLKWYALPAVANML----LEVGG-----------------LEFPACPFNG
FGSG_11195T0        LVLVERTVRVGLGFHRIKATMKVLDKETVEVTAIIPSERLWKYKAGQYIFLQVPKISFFQ
                              ** :: : *. ::*    :** .                 *:.*  .*

NP_001300851.1      WY----------------------------------------------------------
FGSG_11195T0        WHPFTVSFCRGNKMMLHIKTDGNWTAKLRELGGDSGESEIEVGINGPFGAPAQRFYDFNH
                    *:

NP_001300851.1      ---MGTEIGVRDF----CDTQRYNILEEVGRRMGLETHTL--------------ASLWKD
FGSG_11195T0        SIIIGAGIGVTPFSGILADLQYNDDLDHGGPNHEVDHHRHDSEATAIPQAARRSDSSSSD
                       :*: ***  *    .* *  : *:  * .  :: *                 *  .*

NP_001300851.1      RAVTEINV---------------------------AVLHSFQKQNVTIMDHHTASESFMK
FGSG_11195T0        EATTSDNVPETPTRQGSVGPDLINKEKQPQADKAGSFAEDYRRVDFHWMVRERNYLLWLS
                     *.*. **                           :.  .:.. :.  * .      ::.

NP_001300851.1      HMQNE-------YRARGGCP-------------------ADWIWLV----------PPVS
FGSG_11195T0        DLLNDVSMSQDWHREHEDKPHLDIRINTHVTAKQKKISTHVYRWLLEMHRTDEHPASPLT
                     : *:       :* . . *                     : **:          .*::

NP_001300851.1      GSITPV-------------FHQEMLN----------------------------YVLSPF
FGSG_11195T0        GLLNPTHFGRPDFDLILDEHYEEMLKFRASKRTSTRNKEDENYEEDEELKVGVFYCGAPV
                    * :.*.             .::***:                            *  :*.

NP_001300851.1      ------------------------YYYQIEPW-
FGSG_11195T0        VGEILADKCRELTLRGWQDGSKLEYHFMIEVFG
                                            *:: ** :

____________________________________________________________________________________


NP_001300851.1      ------------------------------MNP---------KSLTR-------------
FGSG_12534T0        MALLSIVNVALLGVAYFVACAVWQVVKYRFLHPLAKFPGNFWGSVTRLWITYHNVEADEC
                                                  ::*          *:**

NP_001300851.1      ----------GPRDKPTPLEELLPHAIEFINQYYGSFKEAKI--------EEHLARLEAV
FGSG_12534T0        ETFQELHKKHGPIIRITPTMLLVSDATELPKIYNRHANKSKHYITGSFGKDESLFNMQDS
                              **  . **   *:. * *: : *    :::*         :* * .::

NP_001300851.1      TKEIE----TTGTYQLT----LDELIFATKMAW-----RNAPRCIGRIQWSNLQVFDARN
FGSG_12534T0        VMHAKYRKVAAPLYSLTNIKKMEPLIDNNMSAWMSRLQRDFAATNKPFDFAPFSVYLAYD
                    .   :    ::  *.**    :: **  .  **     *: .     :::: :.*: * :

NP_001300851.1      CSTAQEM--------------------------FQHICRHILYAT--NNGNIRSAITVFP
FGSG_12534T0        VISEVGFGAPFGFVKEGKDVEGLIQGFHDGLTPFGIMARLYPFTNWVKSTFLGKYMVASP
                      :   :                          *  :.*   ::.  :.  : . :.. *

NP_001300851.1      QRSDGKHDFRLWNSQLIRYAGYQMPDGTIRGDAATLEFTQLCIDLGWKP---RYGRFDVL
FGSG_12534T0        EQDSGIGILMRFRDRLIEQRFKDIENGSTGGRIDLLQTFIEARDEDDKPLDINYIKAEIL
                    :...*   :  :...**     :: :*:  *    *:    . * . **   .* . ::*

NP_001300851.1      PLVLQADG--------------QDPEVF-----EIPPDLVLEVTMEHPKYEWFQELGLKW
FGSG_12534T0        LVLLAGADTTGTAFQAMMVHILTNPSVYKKLLAEIDEATAAGNLSEMPQYDEVVE-HLPY
                     ::* . .               :*.*:     **    .     * *:*: . *  * :

NP_001300851.1      Y------------ALPAVANMLLEVGGLEFPACPFNGWYM--GTEIGVRDFCDTQRYNIL
FGSG_12534T0        YIACVKEAMRLTPSAPNIFPRIVPQGGLEI--C---GHFVPEGTEVTCNPWLVHRDPNIY
                    *            : * :   ::  ****:  *   *.::  ***:  . :   .  **

NP_001300851.1      EEVGRRMGLETHTLASLWKDRAVTEINVAVLHSFQ-------KQNVTIMDHHTASESFMK
FGSG_12534T0        GDDAEIFKPER------WLDADKAKIYNKYSMGFGYGPRVCLGQDIARMELYKGPLLFLR
                     : .  :  *       * *   ::*      .*         *::: *: :...  *:.

NP_001300851.1      HMQNEY---RARGGCPADWIWLVPPVSGSITPVFHQEMLNYVLSPFYYYQIEPW
FGSG_12534T0        SFNVEWVDETKRGKY---------VINGGVS--YFEDM-NIKISRRDAAA----
                     :: *:     **            :.*.::  :.::* *  :*
```

**Test if the DeepLoc predictor tool correctly localises the 4 proteins we have already localised experimentally**

The 4 proteins we have localised. The heme reductase (NOS), the two NODs and the previously identified putative heme protein. This test, was run to test the new localisation tool. The results agree well with the experiments!!

NOD1 Localises to cytoplasm and nuclei

>FGSG_00765T0 | FGSG_00765 | Fusarium graminearum PH-1 hypothetical protein (458 aa)

MALSYQQTKLIRGTIPALTDHGERITTIFYRNMLRDHPELNDYFNTVNQANGRQPRALTAVILSYANNIN

HITELIPKMERMCHKHCSLGIKPEHYAIVEKYLIAAFAEVLGPTMTTQIREAWTKAYWMLAKMLIGREAQ

LYRDFGKWQGYRKFRIEKKVEESDDIYSFYLVPVDGKRLPSFQPGQYVSVQIPIVDKGYVQSRQYSLSDA

PRPDYYRVTVKRDEGIHMTRSGRYLGGDALNPGVVSNLLIDMKDEGDVVELAHPAGEFYLDMSNTSNVPI

VLISAGVGVTPMMSILNTVSERQPHRPVSWIHGSRRSVPFYDQVRRIARNRSNFRTNIFKTHLAESDIFG

VTYDHDFRMDLAKVSKDDLYLCHSSTEYYICGPEQFMLEMAEYLKAQKVDASRMHFELFSTGDMEFKVDN

LSISSASKSPSIASAEQRCPSTGAVSADGATCPFSAV*

NOD2 Localises to cytoplasm and nuclei

>FGSG_04458T0 | FGSG_04458 | Fusarium graminearum PH-1 hypothetical protein (416 aa)

MALTAAQVAIVKSTAPILKEHGKTITTTFYRNMLGAHPELKNYFSLRNQQTGAQQAALANSVLAYATYID

DLGKLSHAVERIAQKHVSLFIKAEHYPIVGTHLIGAIGEVLGSALTTEIKDAWVAAYGQLADIFIQREGQ

LYDAAGEWNSWRKFKIAKKEAENDSVTSFYLEPLDDKPLPKFLPGQYVSLQIPIPELDGLLQSRQFSLSE

APGSNHYRISVKLQGPEEEPSLEDLSAGKIPGLVCTRLHKRYNVGDEVELSPPAGEFFLNPADTSAAKKP

LVLLSAGVGATPLVSILDSVLESETASRPITWIHGARYSGSTCFVPHVLDSAKKHDNITAKIFLEDVKEG

DQYDFKGEIDLDQLQKDKLLQLDSSDAEYFICGPEDWMVKVRAFLEENGVPRERQHLELFKTGDV*

Heme that localises to the cytoplasm and nucleus

>FGSG_07925T0 | FGSG_07925 | Fusarium graminearum PH-1 hypothetical protein (264 aa)

MNSRPMKHIDREDLYTNLEARVQYLHSFLDFSSRDIEALITGAKYVKALIPAVVNIVYKKLLQYDITARA

FTTRSTSFEGPLDEVPDENSPQILHRKMFLRAYLMKLCSDPSKMEFWEYLDKVGMMHVGLGRKHPLHIEY

VHLGVCLGFIQDIMTEAILSHPRLHIQRKTALVKALNKVIWIQNDLMAKWHVRDGSEFEVGDSDIEIERE

GYLHGKRIIGDNSGSASDDEASDQLPPSRIPHPNGASAGGVCPFSGMGAPSEE*

NCP heme reductase that localises to the ER-membrane

>FGSG_09786T0 | FGSG_09786 | Fusarium graminearum PH-1 NADPH-cytochrome P450 reductase (693 aa)

MAELDTLDVIVLGVIFLGTVAYFTKGKLWGVTKDPYANGFAAGGAAKPGRTRNIVEAMEESGKNCVIFYG

SQTGTAEDYASRLAKEGKSRFGLNTMIADIEDYDFDSLDTVPNDNVVMFVLATYGEGEPTDNAVDFYEFI

TGEDATFNEGNDPPLGNLNYVAFGLGNNTYEHYNAMVRKVDQALEKFGAHRIGEAGEGDDGAGTMEEDFL

AWKDPMWESLAKKMGLEEREAVYEPIFAINERDDLSPESNEVYLGEPNKLHLEGTAKGPFNSHNPYIAPI

AESYELFSAKDRNCLHMEVDISGSNLKYETGDHIAIWPTNPGEEVNRFLDILDLSGKQHSVITVKALEPT

AKVPFPNPTTYDAILRYHLEICAPVSRQFVSTLAAFAPNDSIKAEMNRLGSDKDYFHEKTGPHYYNIARF

LSSVSKGEKWTTIPFSAFIEGLTKLQPRYYSISSSSLVQPKKISITAVVESQQIPGRDDPFRGVATNYLF

ALKQKQNGDPSPAPFGQTYELTGPRNKYDGIHVPVHVRHSNFKLPSDPGKPVIMIGPGTGVAPFRGFVQE

RAKLARDGVEVGKTLLFFGCRKPSEDFMYEKEWQEYKEALGDKFEMITAFSRESAKKVYVQHRLKERAQE

VSDLLSQKAYFYVCGDASNMAREVNTVLAQIIAEGRGVSEAKGEEIVKNMRSANQYQEDVWS*

### **Summary of 4 predicted sequences** DeepLoc-1.0

Table of predicted subcellular localisations. Use the help page for more detailed description of the output page.

**Predicted proteins**

**FGSG_07925T0**

**Prediction: Cytoplasm, Soluble**

| **Localisation** | **Cytoplasm** | | **Nucleus** | | **Golgi apparatus** | **Endoplasmic reticulum** | | **Lysosome/Vacuole** | **Cell membrane** | **Peroxisome** | **Mitochondrion** | **Extracellular** | **Plastid** |
| --- | --- | --- | --- | --- | --- | --- | --- | --- | --- | --- | --- | --- | --- |
| **Likelihood** | 0.5244 | | 0.3933 | | 0.0396 | 0.0202 | | 0.0142 | 0.0037 | 0.0023 | 0.0014 | 0.0007 | 0.0001 |
| **Type** | | **Soluble** | | **Membrane** | | |  |  |  |  |  |  |  |
| **Likelihood** | | 0.9094 | | 0.0906 | | |  |  |  |  |  |  |  |

**Hierarchical Tree. Donwload:** [PNG](http://www.cbs.dtu.dk/services/DeepLoc-1.0/tmp/5B9D4E85000022815DF31AED/tree_66124.png) **/** [EPS](http://www.cbs.dtu.dk/services/DeepLoc-1.0/tmp/5B9D4E85000022815DF31AED/tree_66124.eps)
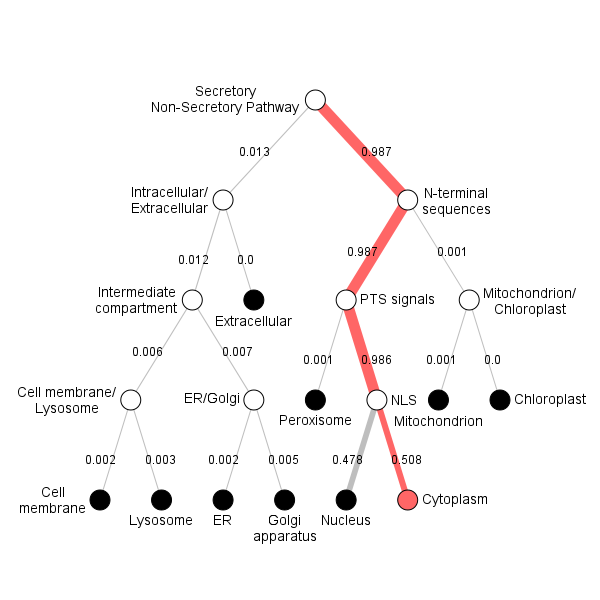

**Position Importance. Donwload:** [PNG](http://www.cbs.dtu.dk/services/DeepLoc-1.0/tmp/5B9D4E85000022815DF31AED/alpha_66124.png) **/** [EPS](http://www.cbs.dtu.dk/services/DeepLoc-1.0/tmp/5B9D4E85000022815DF31AED/alpha_66124.eps) **/** [CSV](http://www.cbs.dtu.dk/services/DeepLoc-1.0/tmp/5B9D4E85000022815DF31AED/alpha_66124.csv)
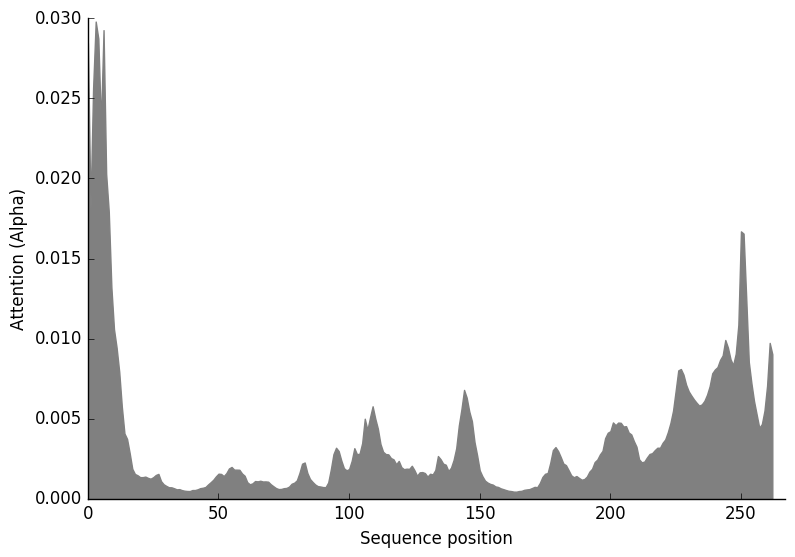

**FGSG_09786T0**

**Prediction: Endoplasmic reticulum, Membrane**

| **Localisation** | **Endoplasmic reticulum** | | **Mitochondrion** | | **Cell membrane** | | **Lysosome/Vacuole** | **Plastid** | **Golgi apparatus** | **Cytoplasm** | **Peroxisome** | **Nucleus** | **Extracellular** |
| --- | --- | --- | --- | --- | --- | --- | --- | --- | --- | --- | --- | --- | --- |
| **Likelihood** | 0.9022 | | 0.0472 | | 0.0231 | | 0.0135 | 0.0091 | 0.0035 | 0.0006 | 0.0006 | 0.0001 | 0 |
| **Type** | | **Soluble** | | **Membrane** | |  |  |  |  |  |  |  |  |
| **Likelihood** | | 0.0012 | | 0.9988 | |  |  |  |  |  |  |  |  |

**Hierarchical Tree. Donwload:** [PNG](http://www.cbs.dtu.dk/services/DeepLoc-1.0/tmp/5B9D4E85000022815DF31AED/tree_66125.png) **/** [EPS](http://www.cbs.dtu.dk/services/DeepLoc-1.0/tmp/5B9D4E85000022815DF31AED/tree_66125.eps)
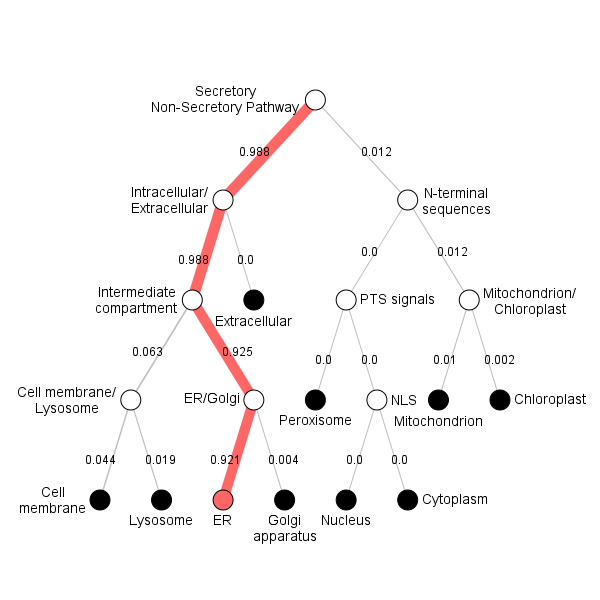

**Position Importance. Donwload:** [PNG](http://www.cbs.dtu.dk/services/DeepLoc-1.0/tmp/5B9D4E85000022815DF31AED/alpha_66125.png) **/** [EPS](http://www.cbs.dtu.dk/services/DeepLoc-1.0/tmp/5B9D4E85000022815DF31AED/alpha_66125.eps) **/** [CSV](http://www.cbs.dtu.dk/services/DeepLoc-1.0/tmp/5B9D4E85000022815DF31AED/alpha_66125.csv)
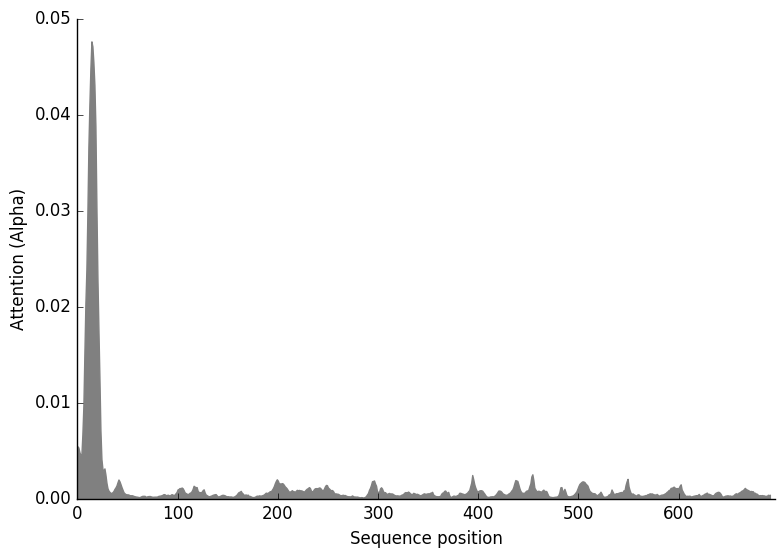

**FGSG_00765T0**

**Prediction: Cytoplasm, Soluble**

| **Localisation** | **Cytoplasm** | | **Peroxisome** | | **Mitochondrion** | | **Plastid** | **Nucleus** | **Extracellular** | **Endoplasmic reticulum** | **Cell membrane** | **Lysosome/Vacuole** | **Golgi apparatus** |
| --- | --- | --- | --- | --- | --- | --- | --- | --- | --- | --- | --- | --- | --- |
| **Likelihood** | 0.5113 | | 0.2989 | | 0.1425 | | 0.0239 | 0.0087 | 0.0084 | 0.0034 | 0.0016 | 0.0009 | 0.0004 |
| **Type** | | **Soluble** | | **Membrane** | |  |  |  |  |  |  |  |  |
| **Likelihood** | | 0.9695 | | 0.0305 | |  |  |  |  |  |  |  |  |

**Hierarchical Tree. Donwload:** [PNG](http://www.cbs.dtu.dk/services/DeepLoc-1.0/tmp/5B9D4E85000022815DF31AED/tree_66123.png) **/** [EPS](http://www.cbs.dtu.dk/services/DeepLoc-1.0/tmp/5B9D4E85000022815DF31AED/tree_66123.eps)
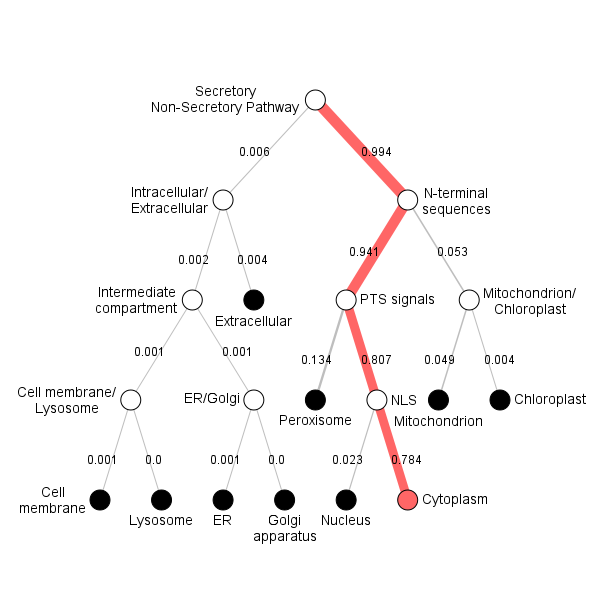

**Position Importance. Donwload:** [PNG](http://www.cbs.dtu.dk/services/DeepLoc-1.0/tmp/5B9D4E85000022815DF31AED/alpha_66123.png) **/** [EPS](http://www.cbs.dtu.dk/services/DeepLoc-1.0/tmp/5B9D4E85000022815DF31AED/alpha_66123.eps) **/** [CSV](http://www.cbs.dtu.dk/services/DeepLoc-1.0/tmp/5B9D4E85000022815DF31AED/alpha_66123.csv)
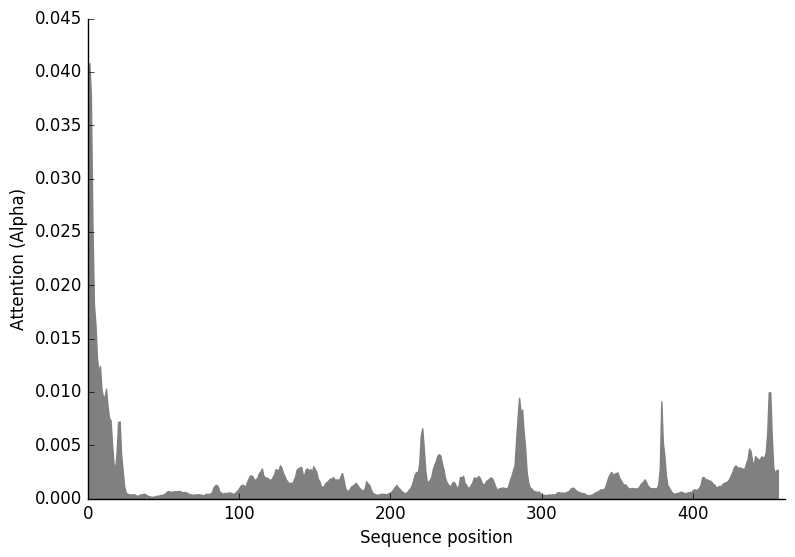

**FGSG_04458T0**

**Prediction: Cytoplasm, Soluble**

| **Localisation** | **Cytoplasm** | | **Mitochondrion** | | **Peroxisome** | | **Plastid** | **Nucleus** | **Extracellular** | **Cell membrane** | **Endoplasmic reticulum** | **Lysosome/Vacuole** | **Golgi apparatus** |
| --- | --- | --- | --- | --- | --- | --- | --- | --- | --- | --- | --- | --- | --- |
| **Likelihood** | 0.4567 | | 0.2614 | | 0.2051 | | 0.0469 | 0.0108 | 0.009 | 0.0042 | 0.0033 | 0.0019 | 0.0006 |
| **Type** | | **Soluble** | | **Membrane** | |  |  |  |  |  |  |  |  |
| **Likelihood** | | 0.9339 | | 0.0661 | |  |  |  |  |  |  |  |  |

**Hierarchical Tree. Donwload:** [PNG](http://www.cbs.dtu.dk/services/DeepLoc-1.0/tmp/5B9D4E85000022815DF31AED/tree_14292.png) **/** [EPS](http://www.cbs.dtu.dk/services/DeepLoc-1.0/tmp/5B9D4E85000022815DF31AED/tree_14292.eps)
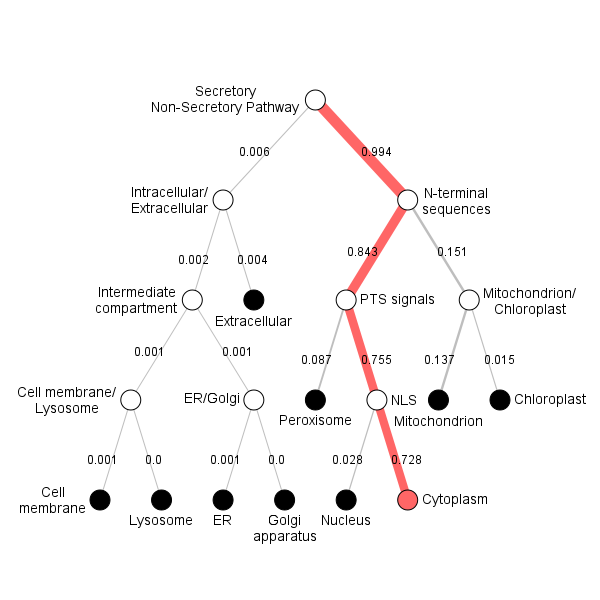

**Position Importance. Donwload:** [PNG](http://www.cbs.dtu.dk/services/DeepLoc-1.0/tmp/5B9D4E85000022815DF31AED/alpha_14292.png) **/** [EPS](http://www.cbs.dtu.dk/services/DeepLoc-1.0/tmp/5B9D4E85000022815DF31AED/alpha_14292.eps) **/** [CSV](http://www.cbs.dtu.dk/services/DeepLoc-1.0/tmp/5B9D4E85000022815DF31AED/alpha_14292.csv)
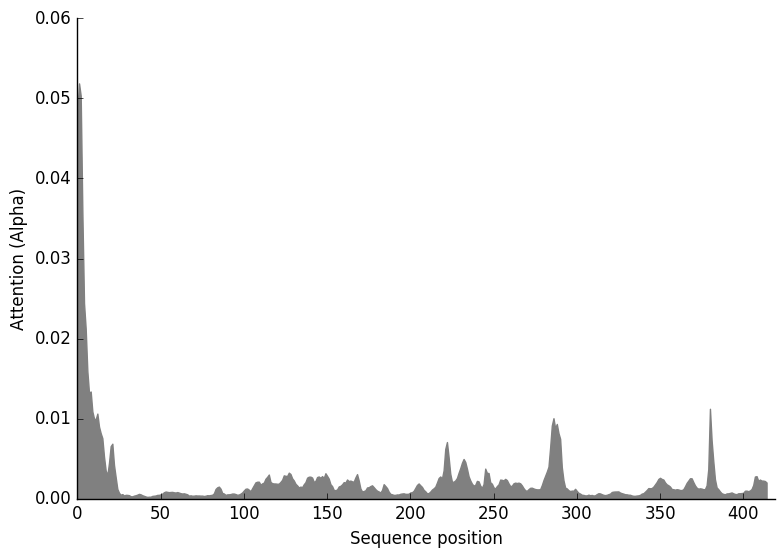

### 1. Localisation using the new DeepLoc-1.0 tool

### This tool was very successful, and we recommend this tool. It gave returned the correct results for the proteins we previously localised (see close to the end)

### **Summary of 8 predicted sequences** DeepLoc-1.0

Table of predicted subcelullar localisations. Use the help page for more detailed description of the output page.

**Predicted proteins**

**FGSG_12534T0**

**Prediction: Endoplasmic reticulum, Membrane**

| **Localisation** | **Endoplasmic reticulum** | | **Mitochondrion** | | **Plastid** | | **Peroxisome** | **Cell membrane** | **Cytoplasm** | **Nucleus** | **Golgi apparatus** | **Lysosome/Vacuole** | **Extracellular** |
| --- | --- | --- | --- | --- | --- | --- | --- | --- | --- | --- | --- | --- | --- |
| **Likelihood** | 0.9021 | | 0.0887 | | 0.0059 | | 0.0022 | 0.0006 | 0.0002 | 0.0001 | 0.0001 | 0 | 0 |
| **Type** | | **Soluble** | | **Membrane** | |  |  |  |  |  |  |  |  |
| **Likelihood** | | 0.0001 | | 0.9999 | |  |  |  |  |  |  |  |  |

**Hierarchical Tree. Donwload:** [PNG](http://www.cbs.dtu.dk/services/DeepLoc-1.0/tmp/5B9D43FC0000227CB34092DD/tree_66115.png) **/** [EPS](http://www.cbs.dtu.dk/services/DeepLoc-1.0/tmp/5B9D43FC0000227CB34092DD/tree_66115.eps)
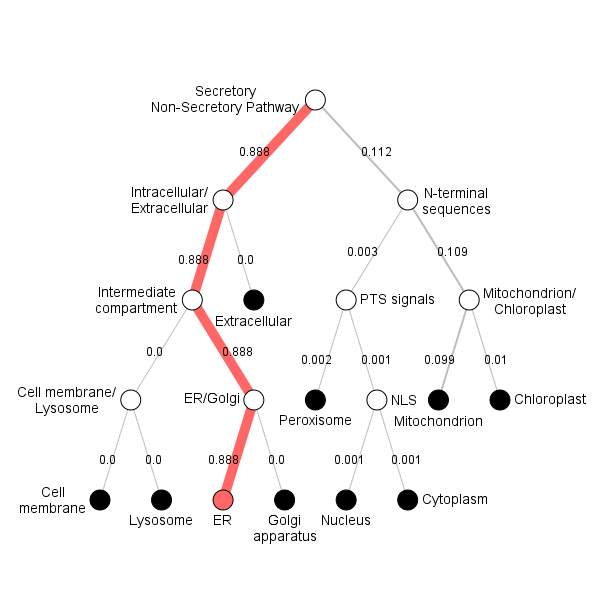

**Position Importance. Donwload:** [PNG](http://www.cbs.dtu.dk/services/DeepLoc-1.0/tmp/5B9D43FC0000227CB34092DD/alpha_66115.png) **/** [EPS](http://www.cbs.dtu.dk/services/DeepLoc-1.0/tmp/5B9D43FC0000227CB34092DD/alpha_66115.eps) **/** [CSV](http://www.cbs.dtu.dk/services/DeepLoc-1.0/tmp/5B9D43FC0000227CB34092DD/alpha_66115.csv)
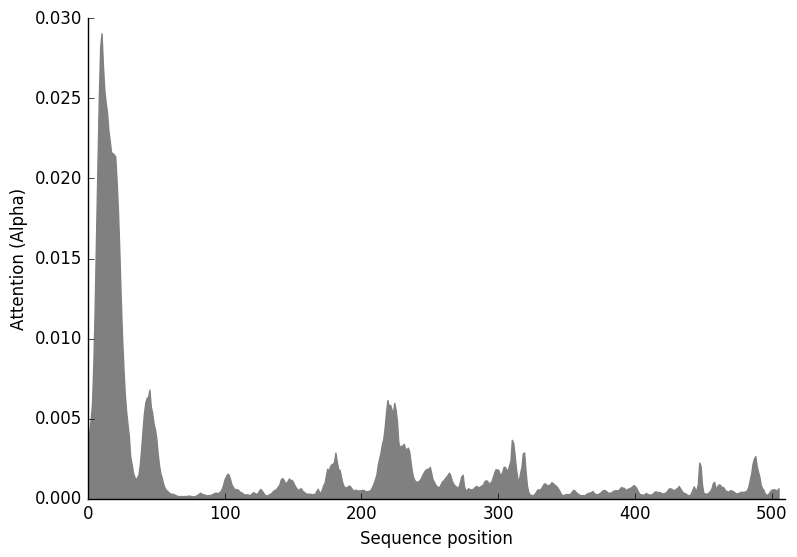

**FGSG_11195T0**

**Prediction: Cell membrane, Membrane**

| **Localisation** | **Cell membrane** | | **Lysosome/Vacuole** | | **Endoplasmic reticulum** | | **Golgi apparatus** | **Mitochondrion** | **Peroxisome** | **Nucleus** | **Plastid** | **Cytoplasm** | **Extracellular** |
| --- | --- | --- | --- | --- | --- | --- | --- | --- | --- | --- | --- | --- | --- |
| **Likelihood** | 0.4641 | | 0.3095 | | 0.1659 | | 0.0465 | 0.0079 | 0.0054 | 0.0004 | 0.0003 | 0 | 0 |
| **Type** | | **Soluble** | | **Membrane** | |  |  |  |  |  |  |  |  |
| **Likelihood** | | 0 | | 1 | |  |  |  |  |  |  |  |  |

**Hierarchical Tree. Donwload:** [PNG](http://www.cbs.dtu.dk/services/DeepLoc-1.0/tmp/5B9D43FC0000227CB34092DD/tree_66114.png) **/** [EPS](http://www.cbs.dtu.dk/services/DeepLoc-1.0/tmp/5B9D43FC0000227CB34092DD/tree_66114.eps)
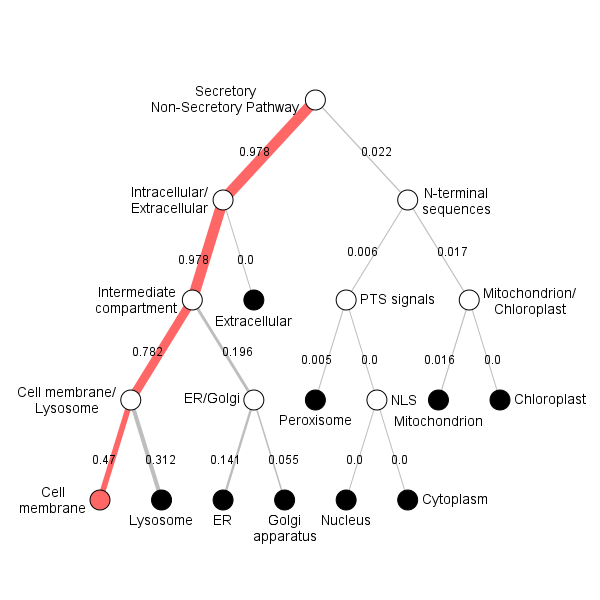

**Position Importance. Donwload:** [PNG](http://www.cbs.dtu.dk/services/DeepLoc-1.0/tmp/5B9D43FC0000227CB34092DD/alpha_66114.png) **/** [EPS](http://www.cbs.dtu.dk/services/DeepLoc-1.0/tmp/5B9D43FC0000227CB34092DD/alpha_66114.eps) **/** [CSV](http://www.cbs.dtu.dk/services/DeepLoc-1.0/tmp/5B9D43FC0000227CB34092DD/alpha_66114.csv)
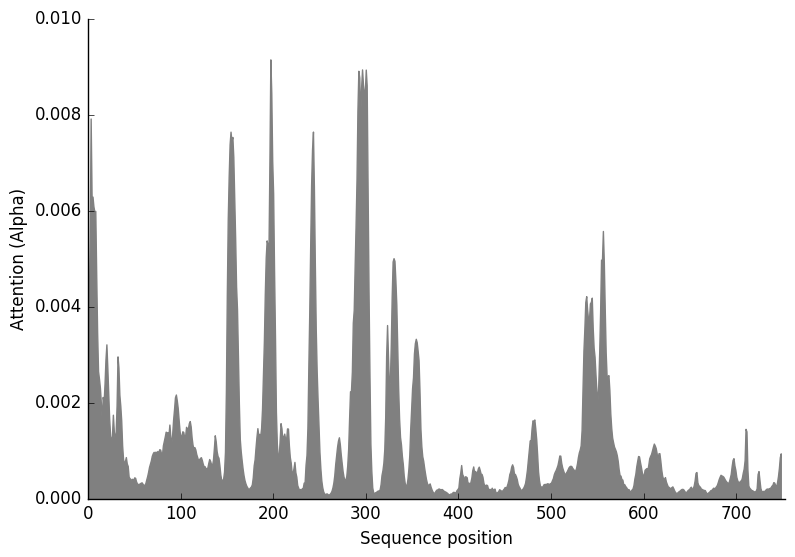

**FGSG_09373T0**

**Prediction: Mitochondrion, Soluble**

| **Localisation** | **Mitochondrion** | | **Endoplasmic reticulum** | | **Extracellular** | | **Peroxisome** | **Plastid** | **Lysosome/Vacuole** | **Cytoplasm** | **Cell membrane** | **Nucleus** | **Golgi apparatus** |
| --- | --- | --- | --- | --- | --- | --- | --- | --- | --- | --- | --- | --- | --- |
| **Likelihood** | 0.7233 | | 0.0908 | | 0.0795 | | 0.0286 | 0.0276 | 0.0227 | 0.0209 | 0.0064 | 0.0001 | 0 |
| **Type** | | **Soluble** | | **Membrane** | |  |  |  |  |  |  |  |  |
| **Likelihood** | | 0.5613 | | 0.4387 | |  |  |  |  |  |  |  |  |

**Hierarchical Tree. Donwload:** [PNG](http://www.cbs.dtu.dk/services/DeepLoc-1.0/tmp/5B9D43FC0000227CB34092DD/tree_66111.png) **/** [EPS](http://www.cbs.dtu.dk/services/DeepLoc-1.0/tmp/5B9D43FC0000227CB34092DD/tree_66111.eps)
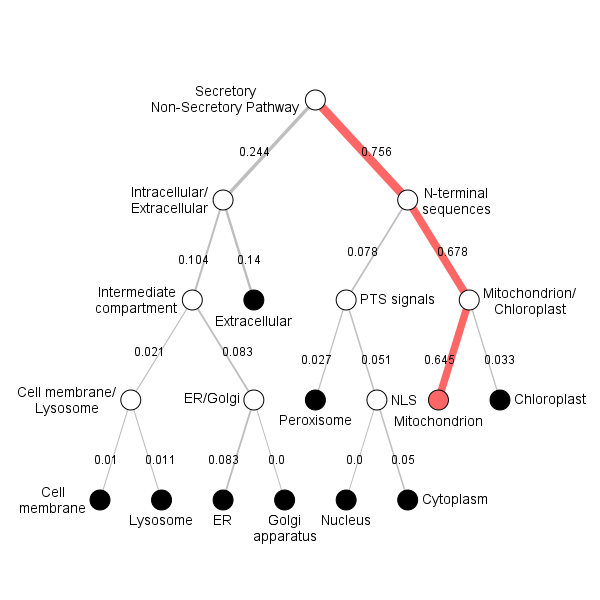

**Position Importance. Donwload:** [PNG](http://www.cbs.dtu.dk/services/DeepLoc-1.0/tmp/5B9D43FC0000227CB34092DD/alpha_66111.png) **/** [EPS](http://www.cbs.dtu.dk/services/DeepLoc-1.0/tmp/5B9D43FC0000227CB34092DD/alpha_66111.eps) **/** [CSV](http://www.cbs.dtu.dk/services/DeepLoc-1.0/tmp/5B9D43FC0000227CB34092DD/alpha_66111.csv)
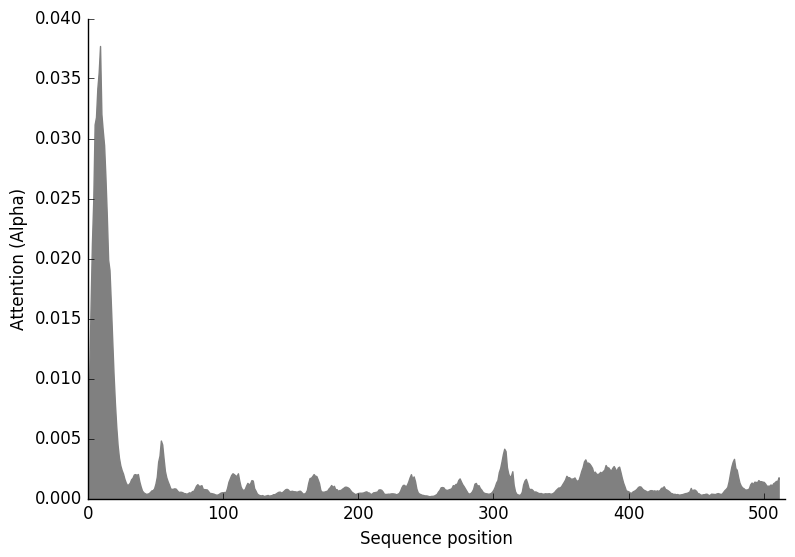

**FGSG_10305T0**

**Prediction: Mitochondrion, Membrane**

| **Localisation** | **Mitochondrion** | | **Peroxisome** | | **Endoplasmic reticulum** | | **Plastid** | **Cytoplasm** | **Nucleus** | **Cell membrane** | **Golgi apparatus** | **Lysosome/Vacuole** | **Extracellular** |
| --- | --- | --- | --- | --- | --- | --- | --- | --- | --- | --- | --- | --- | --- |
| **Likelihood** | 0.5833 | | 0.1956 | | 0.1034 | | 0.092 | 0.0133 | 0.0064 | 0.0038 | 0.001 | 0.0007 | 0.0005 |
| **Type** | | **Soluble** | | **Membrane** | |  |  |  |  |  |  |  |  |
| **Likelihood** | | 0.0104 | | 0.9896 | |  |  |  |  |  |  |  |  |

**Hierarchical Tree. Donwload:** [PNG](http://www.cbs.dtu.dk/services/DeepLoc-1.0/tmp/5B9D43FC0000227CB34092DD/tree_66112.png) **/** [EPS](http://www.cbs.dtu.dk/services/DeepLoc-1.0/tmp/5B9D43FC0000227CB34092DD/tree_66112.eps)
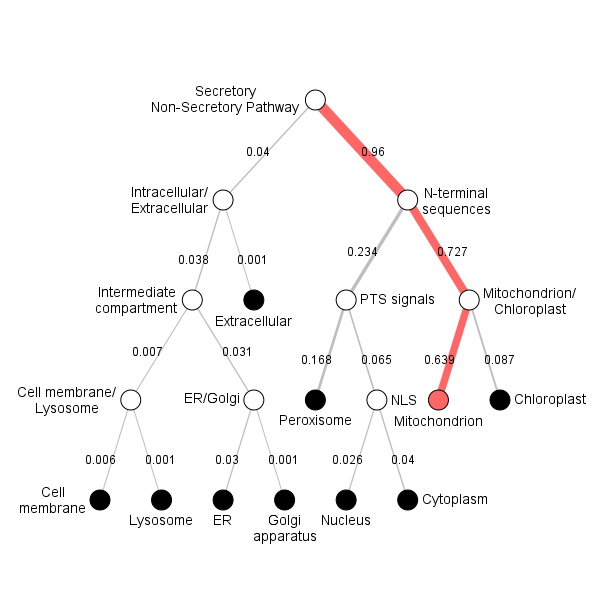

**Position Importance. Donwload:** [PNG](http://www.cbs.dtu.dk/services/DeepLoc-1.0/tmp/5B9D43FC0000227CB34092DD/alpha_66112.png) **/** [EPS](http://www.cbs.dtu.dk/services/DeepLoc-1.0/tmp/5B9D43FC0000227CB34092DD/alpha_66112.eps) **/** [CSV](http://www.cbs.dtu.dk/services/DeepLoc-1.0/tmp/5B9D43FC0000227CB34092DD/alpha_66112.csv)
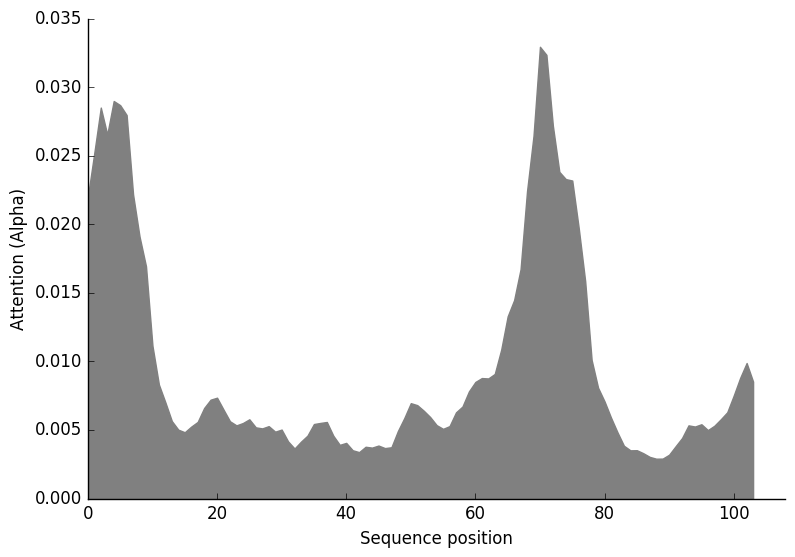

**FGSG_10881T0**

**Prediction: Mitochondrion, Soluble**

| **Localisation** | **Mitochondrion** | | **Extracellular** | | **Cytoplasm** | | **Plastid** | **Peroxisome** | **Cell membrane** | **Endoplasmic reticulum** | **Lysosome/Vacuole** | **Nucleus** | **Golgi apparatus** |
| --- | --- | --- | --- | --- | --- | --- | --- | --- | --- | --- | --- | --- | --- |
| **Likelihood** | 0.6559 | | 0.1856 | | 0.1126 | | 0.0244 | 0.0091 | 0.0063 | 0.0027 | 0.0026 | 0.0007 | 0.0001 |
| **Type** | | **Soluble** | | **Membrane** | |  |  |  |  |  |  |  |  |
| **Likelihood** | | 0.8827 | | 0.1173 | |  |  |  |  |  |  |  |  |

**Hierarchical Tree. Donwload:** [PNG](http://www.cbs.dtu.dk/services/DeepLoc-1.0/tmp/5B9D43FC0000227CB34092DD/tree_66113.png) **/** [EPS](http://www.cbs.dtu.dk/services/DeepLoc-1.0/tmp/5B9D43FC0000227CB34092DD/tree_66113.eps)
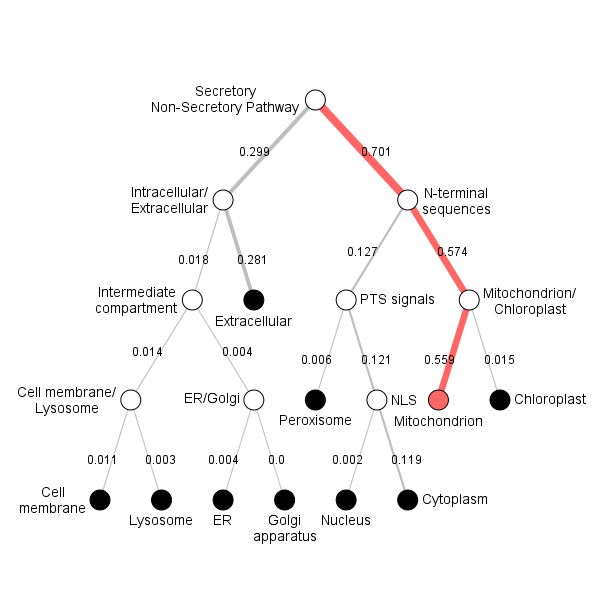

**Position Importance. Donwload:** [PNG](http://www.cbs.dtu.dk/services/DeepLoc-1.0/tmp/5B9D43FC0000227CB34092DD/alpha_66113.png) **/** [EPS](http://www.cbs.dtu.dk/services/DeepLoc-1.0/tmp/5B9D43FC0000227CB34092DD/alpha_66113.eps) **/** [CSV](http://www.cbs.dtu.dk/services/DeepLoc-1.0/tmp/5B9D43FC0000227CB34092DD/alpha_66113.csv)
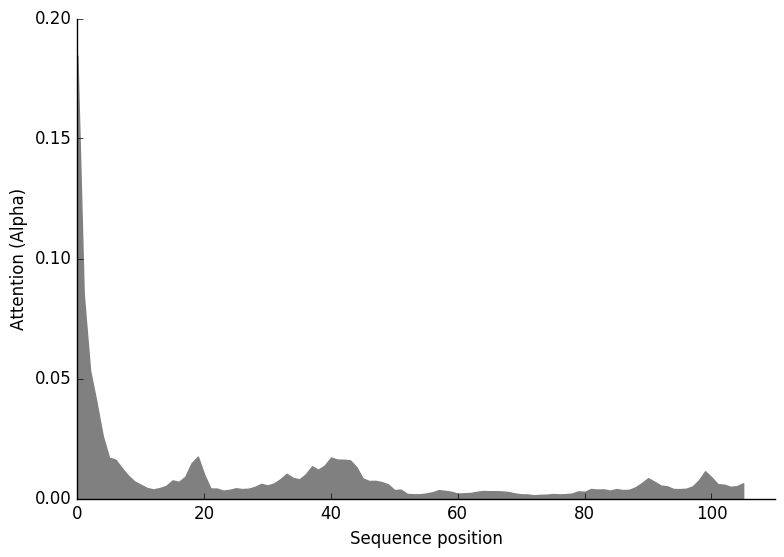

**FGSG_01972T0**

**Prediction: Cytoplasm, Soluble**

| **Localisation** | **Cytoplasm** | | **Peroxisome** | | **Mitochondrion** | | **Plastid** | **Endoplasmic reticulum** | **Extracellular** | **Nucleus** | **Lysosome/Vacuole** | **Cell membrane** | **Golgi apparatus** |
| --- | --- | --- | --- | --- | --- | --- | --- | --- | --- | --- | --- | --- | --- |
| **Likelihood** | 0.5113 | | 0.4509 | | 0.0135 | | 0.0098 | 0.0038 | 0.0034 | 0.0026 | 0.0024 | 0.002 | 0.0003 |
| **Type** | | **Soluble** | | **Membrane** | |  |  |  |  |  |  |  |  |
| **Likelihood** | | 0.9736 | | 0.0264 | |  |  |  |  |  |  |  |  |

**Hierarchical Tree. Donwload:** [PNG](http://www.cbs.dtu.dk/services/DeepLoc-1.0/tmp/5B9D43FC0000227CB34092DD/tree_66109.png) **/** [EPS](http://www.cbs.dtu.dk/services/DeepLoc-1.0/tmp/5B9D43FC0000227CB34092DD/tree_66109.eps)
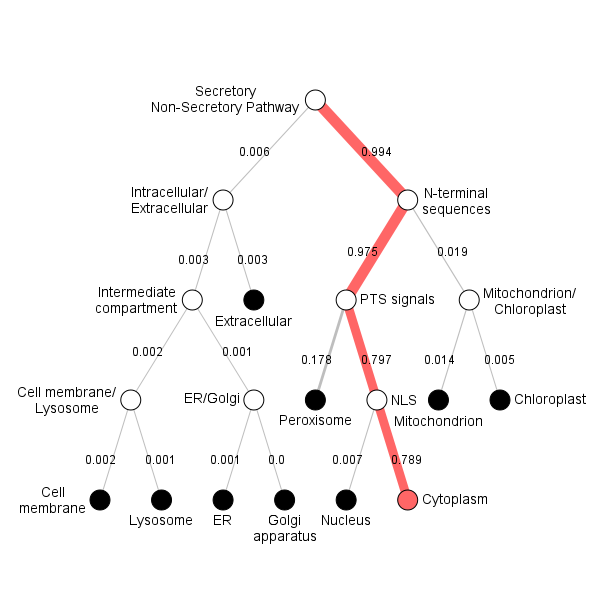

**Position Importance. Donwload:** [PNG](http://www.cbs.dtu.dk/services/DeepLoc-1.0/tmp/5B9D43FC0000227CB34092DD/alpha_66109.png) **/** [EPS](http://www.cbs.dtu.dk/services/DeepLoc-1.0/tmp/5B9D43FC0000227CB34092DD/alpha_66109.eps) **/** [CSV](http://www.cbs.dtu.dk/services/DeepLoc-1.0/tmp/5B9D43FC0000227CB34092DD/alpha_66109.csv)
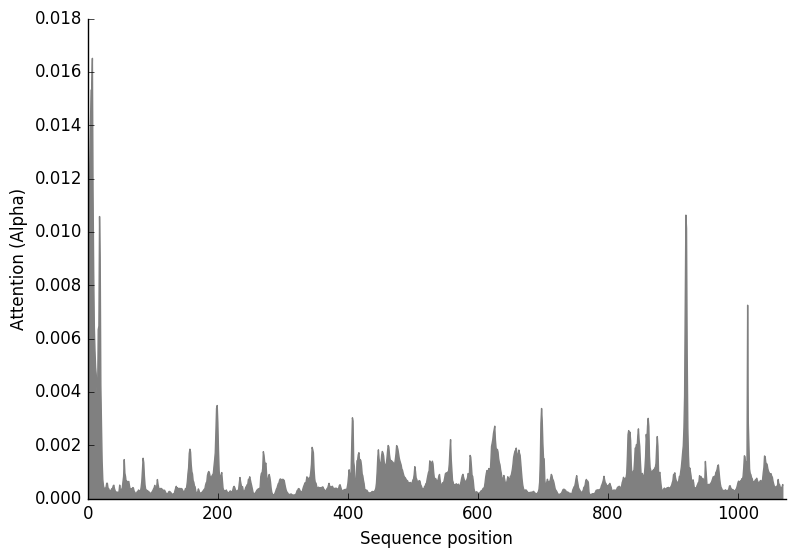

**FGSG_01000T0**

**Prediction: Endoplasmic reticulum, Membrane**

| **Localisation** | **Endoplasmic reticulum** | | **Mitochondrion** | | **Golgi apparatus** | | **Lysosome/Vacuole** | **Plastid** | **Cell membrane** | **Peroxisome** | **Nucleus** | **Cytoplasm** | **Extracellular** |
| --- | --- | --- | --- | --- | --- | --- | --- | --- | --- | --- | --- | --- | --- |
| **Likelihood** | 0.9007 | | 0.0355 | | 0.0317 | | 0.0214 | 0.0072 | 0.0029 | 0.0003 | 0.0003 | 0.0001 | 0 |
| **Type** | | **Soluble** | | **Membrane** | |  |  |  |  |  |  |  |  |
| **Likelihood** | | 0.0001 | | 0.9999 | |  |  |  |  |  |  |  |  |

**Hierarchical Tree. Donwload:** [PNG](http://www.cbs.dtu.dk/services/DeepLoc-1.0/tmp/5B9D43FC0000227CB34092DD/tree_66108.png) **/** [EPS](http://www.cbs.dtu.dk/services/DeepLoc-1.0/tmp/5B9D43FC0000227CB34092DD/tree_66108.eps)
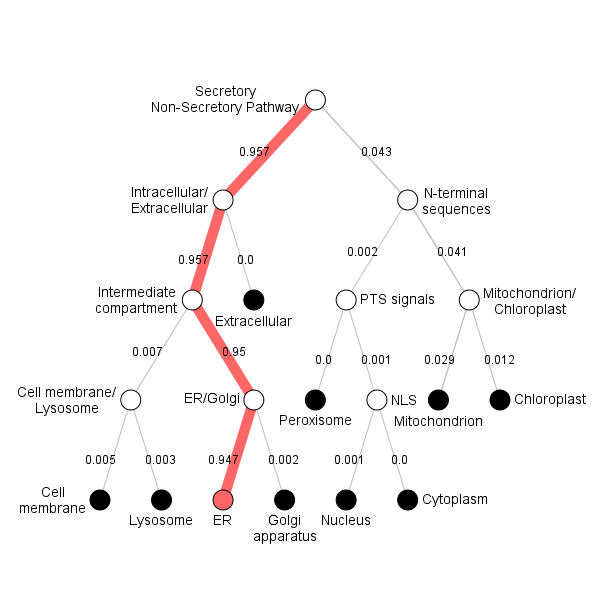

**Position Importance. Donwload:** [PNG](http://www.cbs.dtu.dk/services/DeepLoc-1.0/tmp/5B9D43FC0000227CB34092DD/alpha_66108.png) **/** [EPS](http://www.cbs.dtu.dk/services/DeepLoc-1.0/tmp/5B9D43FC0000227CB34092DD/alpha_66108.eps) **/** [CSV](http://www.cbs.dtu.dk/services/DeepLoc-1.0/tmp/5B9D43FC0000227CB34092DD/alpha_66108.csv)
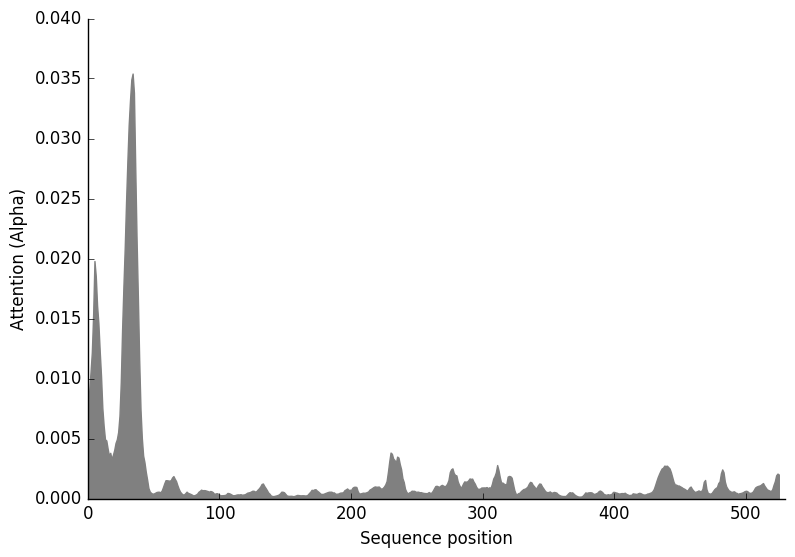

**FGSG_02982T0**

**Prediction: Endoplasmic reticulum, Membrane**

| **Localisation** | **Endoplasmic reticulum** | | **Mitochondrion** | | **Golgi apparatus** | | **Plastid** | **Peroxisome** | **Cell membrane** | **Lysosome/Vacuole** | **Cytoplasm** | **Nucleus** | **Extracellular** |
| --- | --- | --- | --- | --- | --- | --- | --- | --- | --- | --- | --- | --- | --- |
| **Likelihood** | 0.9157 | | 0.0459 | | 0.0175 | | 0.0129 | 0.0033 | 0.0031 | 0.0011 | 0.0002 | 0.0002 | 0.0001 |
| **Type** | | **Soluble** | | **Membrane** | |  |  |  |  |  |  |  |  |
| **Likelihood** | | 0.0006 | | 0.9994 | |  |  |  |  |  |  |  |  |

**Hierarchical Tree. Donwload:** [PNG](http://www.cbs.dtu.dk/services/DeepLoc-1.0/tmp/5B9D43FC0000227CB34092DD/tree_66110.png) **/** [EPS](http://www.cbs.dtu.dk/services/DeepLoc-1.0/tmp/5B9D43FC0000227CB34092DD/tree_66110.eps)
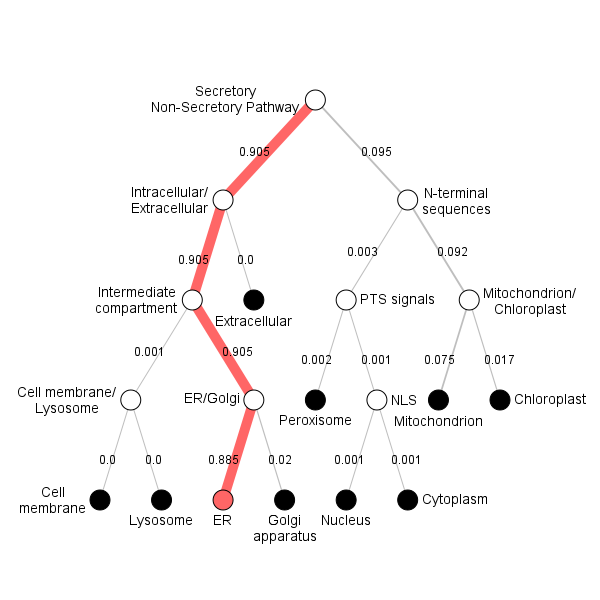

**Position Importance. Donwload:** [PNG](http://www.cbs.dtu.dk/services/DeepLoc-1.0/tmp/5B9D43FC0000227CB34092DD/alpha_66110.png) **/** [EPS](http://www.cbs.dtu.dk/services/DeepLoc-1.0/tmp/5B9D43FC0000227CB34092DD/alpha_66110.eps) **/** [CSV](http://www.cbs.dtu.dk/services/DeepLoc-1.0/tmp/5B9D43FC0000227CB34092DD/alpha_66110.csv)
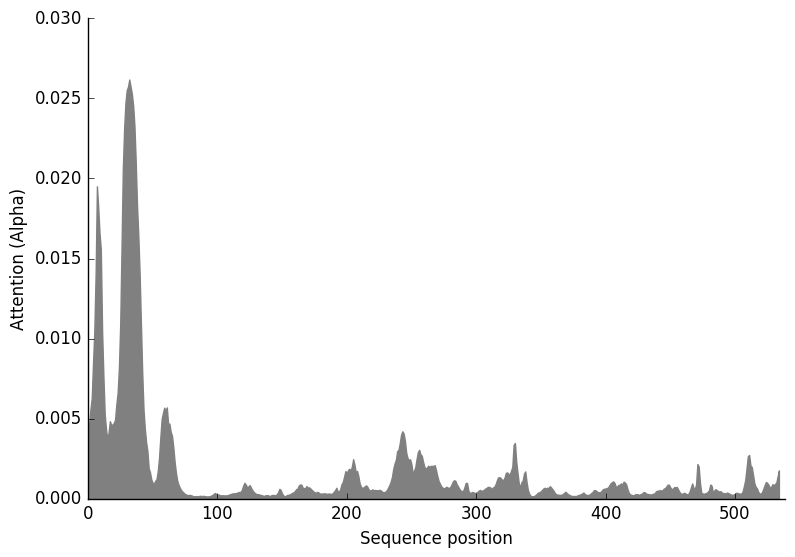

**Possible INVOLVEMENT OF NO synthesis machinery in DON production?**

The Heme reductase (that we previously labelled NOS) could, in principle, deliver electrons for the biosynthesis of Trichothecenes. Tri production is very down when it is deleted. Test if it could have such a double function. That could explain why the heme reductase is not perfectly correlated to the heme protein since it could have double functions.

|  | FGSG_01000 |
| --- | --- |
| 105.814 | 116.985 |
| 103.789 | 112.276 |
| 74.0881 | 78.2851 |
| 159.452 | 170.467 |
| 362.533 | 292.332 |
| 239.945 | 101.704 |
| 78.0037 | 38.2767 |
| 59.7918 | 15.2254 |
| 517.379 | 522.077 |
| 119.968 | 67.9993 |
| 132.006 | 59.3714 |
| 394.684 | 442.054 |
| 127.991 | 280.696 |
| 168.664 | 383.656 |
| 247.965 | 244.405 |
| 103.419 | 200.586 |
| 126.362 | 75.882 |
| 103.5 | 210.093 |
| 163.187 | 87.4043 |
| 167.149 | 127.958 |
| 216.671 | 100.473 |
| 211.886 | 137.476 |
| 143.737 | 82.6768 |
| 260.913 | 137.668 |
| 134.19 | 86.9612 |
| 193.604 | 114.256 |
| 155.01 | 79.5232 |
| 178.297 | 76.2637 |
| 140.038 | 68.9901 |
| 279.094 | 114.518 |
| 97.3659 | 198.356 |
| 117.122 | 151.966 |
| 126.603 | 98.1796 |
| 157.053 | 111.65 |
| 151.161 | 96.2144 |
| 120.102 | 141.774 |
| 117.221 | 98.6742 |
| 148.814 | 117.242 |
| 101.801 | 117.723 |
| 137.59 | 96.831 |
| 160.474 | 82.4114 |
| 143.045 | 162.207 |
| 140.919 | 81.4852 |
| 95.9737 | 119.754 |
| 109.593 | 113.494 |
| 134.257 | 70.8292 |
| 140.441 | 96.8505 |
| 111.148 | 102.438 |
| 120.004 | 69.0561 |
| 117.534 | 80.7569 |
| 134.15 | 147.405 |
| 151.223 | 76.8234 |
| 154.192 | 113.026 |
| 143.011 | 155.79 |
| 145.405 | 19.0682 |
| 118.752 | 15.6191 |
| 212.537 | 27.8689 |
| 132.977 | 32.5972 |
| 137.192 | 15.425 |
| 191.517 | 27.6019 |
| 128.841 | 15.8433 |
| 162.581 | 14.2008 |
| 310.421 | 20.4902 |
| 256.63 | 15.3215 |
| 203.698 | 11.0802 |
| 367.391 | 21.2072 |
| 108.617 | 13.5246 |
| 106.457 | 14.3457 |
| 135.791 | 28.2176 |
| 114.804 | 12.3059 |
| 115.343 | 9.52594 |
| 139.164 | 32.5375 |
| 188.392 | 173.755 |
| 227.615 | 158.216 |
| 199.036 | 156.169 |
| 243.351 | 271.031 |
| 206.464 | 213.656 |
| 236.917 | 297.101 |
| 233.339 | 286.159 |
| 243.046 | 215.32 |
| 226.32 | 192.884 |
| 118.081 | 124.935 |
| 112.802 | 96.431 |
| 135.138 | 80.872 |
| 148.29 | 128.265 |
| 194.3 | 314.505 |
| 177.926 | 176.858 |
| 126.828 | 94.9467 |
| 180.648 | 266.102 |
| 147.893 | 128.613 |
| 99.8174 | 120.757 |
| 134.513 | 140.844 |
| 154.586 | 204.601 |
| 78.408 | 68.7319 |
| 150.558 | 159.839 |
| 129.7 | 163.712 |
| 89.6631 | 73.8577 |
| 174.54 | 238.363 |
| 107.602 | 124.613 |
| 102.673 | 124.211 |
| 137.87 | 144.36 |
| 159.019 | 210.468 |
| 117.326 | 91.279 |
| 103.874 | 62.4403 |
| 86.0486 | 45.5165 |

FGSG_03537 TRI5

|  | FGSG_03537 |
| --- | --- |
| 105.814 | 0.129578 |
| 103.789 | 0.240301 |
| 74.0881 | 0.178242 |
| 159.452 | 1.37706 |
| 362.533 | 0 |
| 239.945 | 0.785223 |
| 78.0037 | 0.0686202 |
| 59.7918 | 0.0637806 |
| 517.379 | 0 |
| 119.968 | 0.264003 |
| 132.006 | 0.519216 |
| 394.684 | 0.447291 |
| 127.991 | 0.11086 |
| 168.664 | 0.267461 |
| 247.965 | 0 |
| 103.419 | 0.172318 |
| 126.362 | 0.83257 |
| 103.5 | 0.698079 |
| 163.187 | 0.225478 |
| 167.149 | 0.262973 |
| 216.671 | 0 |
| 211.886 | 0.59636 |
| 143.737 | 1.2942 |
| 260.913 | 0.868785 |
| 134.19 | 0.301535 |
| 193.604 | 0.146034 |
| 155.01 | 0.133947 |
| 178.297 | 0.0981361 |
| 140.038 | 0.412407 |
| 279.094 | 0.18803 |
| 97.3659 | 0.954073 |
| 117.122 | 0.774211 |
| 126.603 | 0.201862 |
| 157.053 | 0.773051 |
| 151.161 | 0.163079 |
| 120.102 | 1.16623 |
| 117.221 | 0.15179 |
| 148.814 | 0.723476 |
| 101.801 | 0.209818 |
| 137.59 | 0.177171 |
| 160.474 | 0.078553 |
| 143.045 | 0.530218 |
| 140.919 | 0.25518 |
| 95.9737 | 0.398443 |
| 109.593 | 0.290197 |
| 134.257 | 0.565975 |
| 140.441 | 0.140812 |
| 111.148 | 0.445615 |
| 120.004 | 0.180594 |
| 117.534 | 0.06067 |
| 134.15 | 0.167916 |
| 151.223 | 0 |
| 154.192 | 0.137149 |
| 143.011 | 0.173245 |
| 145.405 | 2.15075 |
| 118.752 | 1.62028 |
| 212.537 | 2.25304 |
| 132.977 | 12.9295 |
| 137.192 | 8.39317 |
| 191.517 | 10.9307 |
| 128.841 | 0.273247 |
| 162.581 | 0.242817 |
| 310.421 | 0.035581 |
| 256.63 | 0.445464 |
| 203.698 | 0.619003 |
| 367.391 | 0.365418 |
| 108.617 | 0.175312 |
| 106.457 | 0.159901 |
| 135.791 | 0.228306 |
| 114.804 | 0.266064 |
| 115.343 | 0.381032 |
| 139.164 | 0.24982 |
| 188.392 | 0.154176 |
| 227.615 | 0.249899 |
| 199.036 | 0.443568 |
| 243.351 | 2.95106 |
| 206.464 | 2.65305 |
| 236.917 | 2.5844 |
| 233.339 | 2.15093 |
| 243.046 | 3.67259 |
| 226.32 | 2.31739 |
| 118.081 | 0.175086 |
| 112.802 | 0.118085 |
| 135.138 | 0.416622 |
| 148.29 | 3.07568 |
| 194.3 | 1.88262 |
| 177.926 | 2.49757 |
| 126.828 | 5.25116 |
| 180.648 | 1.34729 |
| 147.893 | 3.0566 |
| 99.8174 | 0.335735 |
| 134.513 | 0.897137 |
| 154.586 | 0.609177 |
| 78.408 | 2.80902 |
| 150.558 | 2.00805 |
| 129.7 | 1.91039 |
| 89.6631 | 1.16121 |
| 174.54 | 0.863593 |
| 107.602 | 2.16111 |
| 102.673 | 0.345339 |
| 137.87 | 0.919529 |
| 159.019 | 0.626647 |
| 117.326 | 0.181156 |
| 103.874 | 0.180944 |
| 86.0486 | 0.240668 |

FIG. In Vitro, short-time exposure to bacterial MAMPs shows the following. X axis the Heme reductase. Correlation with putative ER membrane HEME protein but no correlation with TRI5 (also note relative TRI5 expression low).

As can be seen at the left, the Heme protein FGSG_01000 is correlated to the heme reductase, but other functions are for the reductase. The reductase is not correlated with TRI5 in the DON synthesis in vitro (first step and is correlated with the TRI4 that is also a HEME localised in ER) (we also know that there is no DON formed during these conditions)

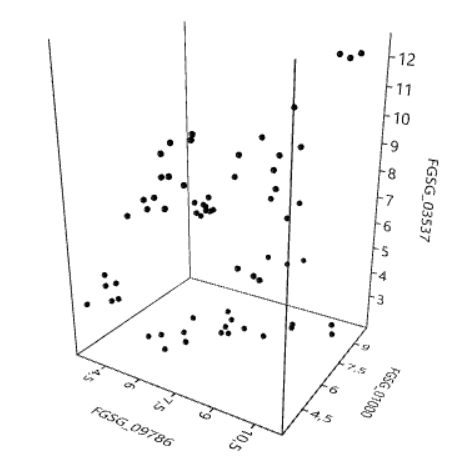

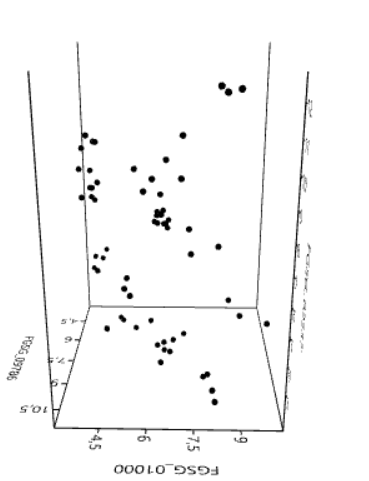

In “in planta” transcriptomic data, we find the following relationship. As can be seen Tri5 (FGSG_03537) (the first step in DON synthesis) and the putative ER Heme (FGSG_01000) both correlates with the heme reductase (FGSG_09786). However, the TRI5 does not correlate with the putative ER heme (FGSG_01000). It seems to be a negative correlation, maybe a competition for electrons (or a difference in inhibition by NO from fungus and/or plant.

**Sterol synthesis is also implied for Cyp51A and Cyp51B and thus (FGSG_01000), the putative heme protein involved in NO production**

Cyp51A <https://bmcgenomics.biomedcentral.com/articles/10.1186/1471-2164-12-52> is probably the one that makes sterol. FGSG_01000 is annotated as Cyp51B and is thus implied for sterol synthesis. The paper shows little response to azoles for FGSG_01000, and it responds a lot to MAMPs in our data and correlates with the heme reductase involved in NO production. The response seen for FGSG_01000 to azoles in the linked paper is probably a stress response since NO production is also part of a general stress response. Alternatively, FGSG_01000 is involved BOTH in sterol synthesis and NO synthesis.

>XP_011321548.1 cytochrome P450 Cyp51A [Fusarium graminearum PH-1]

MFHLLIYPLWVLVALFAVIIANLLYQQLPRRPDEPPLVFHWFPFFGNAVAYGLDPCGFFEKCREKHGDVF

TFILFGRKIVACLGVDGNDFVLNSRLQDANAEEVYGPLTIPVFGSDVVYDCPNSKLMEQKKFVKFGLTQK

ALESHVQLIEREVLDYVETDPSFSGRTSTIDVPKAMAEITIFTASRSLQGEEVRRKLTAEFAALYHDLDL

GFRPVNFLFPWLPLPHNRKRDAAHIKMREVYMDIINDRRKGGIRTEDGTDMIANLMGCTYKNGQPVPDKE

IAHMMITLLMAGQHSSSSASSWIVLHLASSPDITEELYQEQLVNLSVNGALPPLQYSDLDKLPLLQNVVK

ETLRVHSSIHSILRKVKRPMQVPNSPYTITTDKVIMASPTVTAMSEEYFENAKTWNPHRWDNRAKEEVDT

EDVIDYGYGAVSKGTKSPYLPFGAGRHRCIGEKFAYVNLGVIVATLVRNFRLSTIDGRPGVPETDYTSLF

SRPAQPAFIRWERRKKI

Conclusion and hypothesis for F. graminearum

1. FGSG_01000 is a putative ER localised heme protein involved in NO production of NO from arginine.
2. It gets probably its electrons from a heme reductase FGSG_09786 that we already know is involved in making NO and DON, that potentially also delivers electrons from NADPH to both the NO synthesis and the DON synthesis heme (TRI4). This double function could explain why DON is absent and why infection is reduced when the reductase is deleted.
3. The apparent negative correlation between the TRI5 expression (First step in DON production) could indicate a feed-back inhibition by some compound in the DON-synthesis caused by lack of reduction by the heme reductase (TRI4) of some intermediary in the DON synthesis when there is a high NO synthesis. NO is known to self-inhibit the heme in the NOS enzymes of mammals, so other hemes are involved in NO production is likely self-inhibited or or inhibited by NO produced by the plant innate immunity system getting stressed by the fungus presence.
4. This self inhibition might also self-inhibit the FGSG_01000 involvement in ergosterol synthesis if FGSG_01000 is indeed ivolved in sterol production and should lead to accumulation of lanosterol and inhibition of endocytosis since ergosterol is needed for endocytosis.
