## Supplemental Files and Data for "Evolutionary conserved multifunctional nitric oxide synthesis proteins responding to bacterial MAMPs are located at the endoplasmic reticulum": Supplementary file SF7 Phylogenetic and domain analysis of the CYP(NO) for production of NO at the ER membrane.docx

Phylogenetic analysis of the Cytochrome (CYP) responsible for NO production at the ER membrane.

>1. FGSG_01000T0 | FGSG_01000 | Fusarium graminearum PH-1 cytochrome P450 Cyp51B (527 aa)

MGLLQELAGHPLAQQFQELPLGQQVGIGFAVFLVLSVVLNVLNQLLFRNPNEPPMVFHWFPFVGSTITYG

MDPPTFFRENRAKHGDVFTFILLGKKTTVAVGPAGNDFILNGKLKDVCAEEIYTVLTTPVFGKDVVYDCP

NAKLMEQKKFMKIALTTEAFRSYVPIISSEVRDYFKRSPDFKGKSGIADIPKKMAEITIFTASHALQGSA

IRSKFDESLAALYHDLDMGFTPINFMLHWAPLPWNRKRDHAQRTVAKIYMDTIKERRAKGNNESEHDMMK

HLMNSTYKNGIRVPDHEVAHMMIALLMAGQHSSSSTSSWIMLRLAQYPHIMEELYQEQVKNLGADLPPLT

YEDLAKLPLNQAIVKETLRLHAPIHSIMRAVKSPMPVPGTKYVIPTSHTLLAAPGVSATDSAFFPNPDEW

DPHRWEADSPNFPRMASKGEDEEKIDYGYGLVSKGSASPYLPFGAGRHRCIGEHFANAQLQTIVAEVVRE

FKFRNVDGGHTLIDTDYASLFSRPLEPANIHWERRQ

>2. XP_011325340.1 cytochrome P450 51 [Fusarium graminearum PH-1] FGSG_11024

MESLYETLRTLPLSVSIPLTTSIIIILSIVTNVVKQLWFPNPHRPPVVFHIFPFIGSTVQYGIDPYAFFF

DCRDKYGDCFTFILLGKSTTVFLGPKGNDFILNGKHADLNAEDVYGKLTTPVFGEEVVYDCSNARFMDQK

RLLKLGLTTDSLRCYIPKFVKEVEDYVKNSPYFKGDTGIVNITEVMAEITIYTASGSLLGNEVRSMFDST

FATLYRHLDDGFQPINFVMPGLPLPQNFRRNHARKVMEKLFSDIISKRRETGNQGDETDMIWMLMNAQYK

DGEPLPDHHAARMLIAILMGGQHNTAVSGAWLLLNLAHKPHLVQELYEEQTQVLGSPQEPLTWENLQKLT

LNGQVIKETLRLHSPIHSILRQVKSPMRVPGTEWVVPPSHTLLSSPGTMARSEEFFPRPSEWDPHRWDKI

EPLVKTAEDGQTVDYGFGVMSKSVSSPYLPFGAGRHRCVGENYAYAQLGAIVATFIRLVHIEQPDPKAPL

PAPDYSSMFSRPMNPAEIRWRRRETVE

>3. XP_011321548.1 cytochrome P450 51 [Fusarium graminearum PH-1] FGSG_04092

MFHLLIYPLWVLVALFAVIIANLLYQQLPRRPDEPPLVFHWFPFFGNAVAYGLDPCGFFEKCREKHGDVF

TFILFGRKIVACLGVDGNDFVLNSRLQDANAEEVYGPLTIPVFGSDVVYDCPNSKLMEQKKFVKFGLTQK

ALESHVQLIEREVLDYVETDPSFSGRTSTIDVPKAMAEITIFTASRSLQGEEVRRKLTAEFAALYHDLDL

GFRPVNFLFPWLPLPHNRKRDAAHIKMREVYMDIINDRRKGGIRTEDGTDMIANLMGCTYKNGQPVPDKE

IAHMMITLLMAGQHSSSSASSWIVLHLASSPDITEELYQEQLVNLSVNGALPPLQYSDLDKLPLLQNVVK

ETLRVHSSIHSILRKVKRPMQVPNSPYTITTDKVIMASPTVTAMSEEYFENAKTWNPHRWDNRAKEEVDT

EDVIDYGYGAVSKGTKSPYLPFGAGRHRCIGEKFAYVNLGVIVATLVRNFRLSTIDGRPGVPETDYTSLF

SRPAQPAFIRWERRKKI

>XP_011322174.1 trichodiene oxygenase [Fusarium graminearum PH-1] FGSG_03535

MIDQDWIKSLLNIPVSHVAGIFAASTVIYFLSSCFYNLYLHPLRKIPGPKLAAIGPYLEFYHEVIRDGQY

LWEISKMHDKYGPIVRVNAREVHIRDSSYYTTIYTAGSRKTNKDPATVGAFDVPSATAATVDHDHHRSRR

GYLNPYFSKRTITNLEPFIHERVTKLLTRFQQHLDDDQVLSLDGAFCALTADVITNRFYGKHNDYLSLPD

FHFVVRDGFLGLTKIYHLARFLPGLVTILKRLPYSCIRMIAPSVCDLLQMRDEIQDRGGEEFLSNKSHEA

KSSILFGALADSHIPSHERTVERMLDEGTVILFAGTETTSRTLAITVLYLLTHPECLKKLREELNSLPPV

KDGQYSLATLENLPYLNGVIHEGFRLAFGPISRSGRVATQENLKYKEHVIPKGTPISQSTYFMHTDPKNF

PEPEKFKPERWIEAQQKGIPLKKYITNFSQGSRQCIGYTMAFAEMYLALSRIARAYDIELYDTTKADIDM

THARIVGYPKAIPGKKEHLGEVRVKVLKAL

>pdb|5EQB|A Chain A, Lanosterol 14-alpha Demethylase

MSATKSIVGEALEYVNIGLSHFLALPLAQRISLIIIIPFIYNIVWQLLYSLRKDRPPLVFYWIPWVGSAVVYGMKPYEFFEECQKKYGDIFSFVLLGRVMTVYLGPKGHEFVFNAKLADVSAEAAYAHLTTPVFGKGVIYDCPNSRLMEQKKFVKGALTKEAFKSYVPLIAEEVYKYFRDSKNFRLNERTTGTIDVMVTQPEMTIFTASRSLLGKEMRAKLDTDFAYLYSDLDKGFTPINFVFPNLPLEHYRKRDHAQKAISGTYMSLIKERRKNNDIQDRDLIDSLMKNSTYKDGVKMTDQEIANLLIGVLMGGQHTSAATSAWILLHLAERPDVQQELYEEQMRVLDGGKKELTYDLLQEMPLLNQTIKETLRMHHPLHSLFRKVMKDMHVPNTSYVIPAGYHVLVSPGYTHLRDEYFPNAHQFNIHRWNNDSASSYSVGEEVDYGFGAISKGVSSPYLPFGGGRHRCIGEHFAYCQLGVLMSIFIRTLKWHYPEGKTVPPPDFTSMVTLPTGPAKIIWEKRNPEQKIGGRHHHHHH

>XP_003713527.1 cytochrome P450 51 [Pyricularia oryzae 70-15]

MGLLQDTTGPLVDAFYQLGTGAQVGVAFVSFIFLSVFFHVAQQIFFKNPHEPPVVFSWFPVVGSTVTYGK

DPPQFFRDMAKKYGNIFTFILLGKKTTVYIGTEGNEFILNGKLRDVNAEEIYGPMTTPVFGKDVVYDCPN

AKLMEQKKFMKIALTTEAFRSYVPIIADEVSSYLKRTPAFKGPSGVVNIPPKMAEITIFTASHALQGKEI

RDQFDETLADLYHDLDMGFHPVNFKLHWLPLPRNIRRDKAQKTIAKIYMDTIQRRRAKGKDSEAKDMMYH

LMNSTYKNGTPVPDHEIAHMMIALLMAGQHSSSSTSSWIMLRLASRPDIMEELYQEQVRALGADLPPLRY

EDLANLPLHLAVIKETLRLHAPINSILRAVKQDLPVPGTNYVIAKDTTVLAAPGYSAGDPNHFPEPELWE

PHRWEADSRLAPRISMSNDNDEEEKIDYGYGLVSKGTTSPYLPFGAGRHRCIGEHFANVQLQTIVAMIVR

EFKFRNVDGSGKVVGTNYASLFSRPEEPAKIYWERR

>KFG87779.1 cytochrome P450 51B [Metarhizium anisopliae]

MGVLQQVASHPLSQQFQTLGLGSQIAVALGGFIALAVALNVASQILFKNPNEPPLVFHWFPFIGSTVTYG

MDPPKFFKENRAKFGDVFTFVLLGKKTTVAVGPAGNDFILNGKLKDVNAEEIYTVLTTPVFGRDVVYDCP

NAKLMEQKKFMKIALTTEAFRSYVPIISGEVQSYFKRDPGFKGKSGIVDIPKKMAEITIFTASHALQGSA

IRGKFDESLAALYHDLDMGFTPINFMLHWAPLPWNRKRDHAQRTVAKIYMDTIKERRAKGDDDKELDIMK

HLMNSTYKNGTPVPDHEVAHMMIALLMAGQHSSSSTSSWIMLRLAQNPHLVEELYQEQIKALGADLPPLT

YEDLSKLPLNQAIIKETLRLHAPIHSIMRAVKQPMPVPGTKYVIPTTHTLLAAPGVSASDPTYFPSPESW

DPHRWDTDSPNAPTIVRNVAEEEEKVDYGYGLVSKGAASPYLPFGAGRHRCIGEHFANVQLQTIVAETVR

LFKLSNVDGSNTIIGTDYASLFSRPLEPARIRWERRD

>RXG47959.1 hypothetical protein VDGE_04380 [Verticillium dahliae]

MGLLQEITGPAAQQFQTLGTASQVGVVVFGVVLLSVVLHVANQLLFKNPNEPPVVFSWFPFIGNTITYGM

DPPAFFEANKKKYGDVFTFILLGSKTTVAVGPQGNDFILNGKQRDVNAEEIYSVLTTPVFGRDVVYDCPN

AKLMEQKKFMKIALTTEAFRSYVPIISHEVQNFFKTSKYFKEQSGVVNIPPRMAEITIFTASHALQGSEI

RNKFDTSLAELYHDLDMGFSAINFALHWAPLPWNRKRDAAQKAVAQIFMDTIKDRRESGRTDGLDMISHL

MRSTYKNGINVPDHEIAHMMIALLMAGQHSSSSTSSWIMARLAQNPSVIDDLYKEQVEALGADLPPLTYE

DLSKLPLNQAIIKETLRLHAPIHSILRKVKSPMPVPGTKYVVPTSHNLLSAPGTSATDPTYFPNPAIWDP

YRWLPDSPNAPTFSRAENADEEKIDYGYGLVSKGANSPYLPFGAGRHRCIGEQFANVQLQTITAEIVRLV

KFSNADGSSNINGTDYASLFSRPLEPVNIRYERREKA

>XP_018232321.1 eburicol 14-alpha-demethylase [Fusarium oxysporum f. sp. lycopersici 4287]

MGLLQELASHPLAQQYQELPLGQQIGIGFGAFIILSVVLNVLNQLLFKNRNEPPLVFHWFPFVGSTITYG

MDPPKFFKENRAKHGDVFTFVLLGKKTTVAVGPTGNDFILNGKLKDVSAEEIYTVLTTPVFGKDVVYDCP

NAKLMEQKKFMKIALTTEAFRSYVPIISAEVRDYFKKSPDFKGKSGIVDIPKKMAEITIFTASHALQGSV

IRNKFDESLAALYHDLDMGFTPINFMLHWAPLPWNRKRDHAQRTVAKIYMDTIKERRAKDNDDTEHDMMK

HLMNSTYKNGTPVPDHEVAHMMIALLMAGQHSSSSTSSWIMLRLAQYPHIMEELYQEQVRELGADLPPLT

YDNLAKLPLNQAIIKETLRLHAPIHSIMRAVKSPMPVPGTKYTIPTSHTLLAAPGVSATDSAYFPNPDEW

DPHRWEVDSPNFPRMATRGDDEEKIDYGYGLVSKGSASPYLPFGAGRHRCIGEHFANAQLQTIVAEVVRE

FKFRNVDGGNTLIDTDYASLFSRPLEPANIHWERRQQ

>XP_001396151.2 sterol 14-alpha-demethylase [Aspergillus niger CBS 513.88]

MGLLAVVLDSLCERCSNTSVLVVLGFGLLSLLAVSVIFNILRQLFFKNPNEPPLVFHWFPFIGSTISYGM

DPYKFFFDCRAKYGDIFTFILLGKKTTVYLGTRGNDFILNGKLRDVCAEEVYSPLTTPVFGRNVVYDCPN

AKLMEQKKFVKFGLTSDALRSYVRLITGEVENFVEHSAAFKGSSGVFDVCKTIAEITIYTASRSLQGKEV

RSRFDSTFAELYHDLDMGFAPINFMLPWAPLPHNRKRDAAQKKMTETYMEIIKERRNGGNKKDSEDMVWN

LMSCIYKDGTPVPDEEIAHMMIALLMAGQHSSSSTASWIVLHLARNPQIMEELYAEQIRVLGSDLPPLTY

DNLQKLDLHAKVIKETLRIHAPIHSIIRAVKNPMPVDGTPYVIPNSHNVLSSPGVTARSEEYFPNPLKWD

PHRWDETIAASAEEEDQIDYGYGLVSKGTNSPYLPFGAGRHRCIGEQFAYVQLGAITAALVRLFKFRNLP

NVKDIPETDYSSLFSKPAGKSIIQFEKRATTIKS

>CAC85622.1 eburicol 14alpha demethylase [Blumeria graminis f. sp. hordei]

MGISESFMFPYLQPLLQLGFGIALASGIISLLLLLTFLNVLKQLLFKNPNEPPIVFHWIPIIGSTISYGM

NPYKFFHESQAKYGNIFTFILLGKKTTVYLGRQGNNFILNGKLRDVNAEEIYTVLTTPVFGTDVVYDCPN

SKLMEQKKFMKAALTTEAFRSYVPIIQNEVKSFIEKCDDFRKSKGIINIDAVMAEITIYTASHTLQGKEV

RDRFDSSLAVLYHDLDMGFTPINFMLHWAPLPHNRARDHAQRTVAKIYMEIINSRRTQKETDDSNLDIMW

QLMRSSYKDGTPVPDKEIAHMMIALLMAGQHSSSSSSTWIMLWLAARPDITEELYQEQLELLGSELPPLK

YEDLSKLSLHQNVLKEVLRLHAPIHSILRKVKNPMPVPGTSYVIPKTHSLLAAPGWTSRDASYFPNPLKW

DPHRWDTGSGGVIGTDMEDEKFDYGYGLISTGAASPYLPFGAGRHRCIGEQFATVQLVTIMATMVRSFKF

HNLDGRNSVAETDYSSMFSRPMAPATIAWEKRDKKDKTEC

Basidiomycota

>TFK33208.1 lanosterol 14-alpha-demethylase [Crucibulum laeve]6E-164

MSFGAFNSSSIPLPDGWSGYLAHAQEHWLPENSRTLLILLINAPVLAILLNALRQVIMPRDPSLPPEVFH

WLPIIGSAVQYGNDPLNFFFKCQEKYGDVFTFILFGRRVTVALGPKGNNFILGGKSTVFNAEDAYTHLTT

PIFGKDVVYDVPNEKFMEQKRFVKVGLSTDNLRAYVGMIEEEVDEFLKIDPSFRVYQTNDINEWGTFDVI

MALQEITILTASRTLQGKEVREGLNKTFAHLYSDLDGGFTPLNFLFPNLPLESYRKRDRAHQKISQFYIE

IIQKRREGGEDHEHDMIAALMQQTYRNGEHLKDHEIAHIMIALLMAGQHTSSATGSWLLLHIANNPDVAE

ALYQEQVKFFSTPDGKLRSPTYEELRQLPLMDAVIRETLRMHPPIHSIMRYVRDDVPIPGTLSAPSKDTT

YVVPKGHYVLASPAVSQMDPKVWKDTSKWDPYRWADPEGMAARAFDTYADESGEKIDYGFGAVSKGTESP

YQPFGAGKHRCIGEQFAYLQLGTIVATVIRRMEMRIEKVPEHNYHTMITMPKKPRIISYRRRNFD

>XP_006459056.1 hypothetical protein AGABI2DRAFT_191124 [Agaricus bisporus var. bisporus H97] 2e-155

MSYAANGTTPILDAWSGYVAYAQTQLTPWSSRMTILSIATIPVLVVVLNVLSQLLQFRKSSEPPVVFHWL

PFIGSAIGYGTDPLNFFNRCRDKYGDVFTFVLFGRRVTVALGSKGNNFILGGKSIVFNAEDAYTHLTTPV

FGKDVVYDIPNDLFMEQKKFVKVGLSSENLRAYVGMIEEEVEEYLGSDSAFSAFQSNDANEWGHFDVLKV

MSGITILTASRTLQGKEVRAGLDKTFAEIFNDLDGGFTPLNFLFPNLPLASYKKRDVAHKKISDFFSSIV

KGRRAETNPDHEYDMIASLMNQKYRDGRQLKDHEIAHIMIALLMAGQHTSSATGSWVLCHLARRPDVCDA

LYKEQVKNFSNPDGSFRSMTHEELRQLPILDSVIRETLRLHPPIHSVMRHVRDDMPVPATLAAPGKDTTY

VIPKGHYVLASPLVSQLDRTIWKDAETWEPLRWSDPEGIAAQANKLYEDVQGEKIDYGFGAVSKGTESPY

QPFGAGKHRCIGEQFAYLQLGTIITTFIRHMELKMDKLPEHNYQTLITLPKAPREVYYRRRK

>XP_007318458.1 hypothetical protein SERLADRAFT_449218 [Serpula lacrymans var. lacrymans S7.9]

MSAFLDYVLNYVVTSPGRAILLALLYTPIAAIVLNVLRQLVVPRDPSLPPDVFHWIPIVGSAIEYGNDPL

KFFFKCRDKYGDVFTFVLLGRKVTVALGAKGNNFILGGKSTAFSAEDAYTHLTTPVFGKDVVYDVPNEVF

MEQKKFVKVGLSTENFRSYVGMIEDEVDEFMRNDPSFLIYQMNDINEWGRFDAATAMSEITILTASRTLQ

GKEVRSNLDKSFSELYNALDGGFTPLNFMFPNLPLDSYRRRDEAHKKMSEFYVNIIRKRKEGNNDHEHDM

IAALRDQTYRNGRPLPDHEIAHIMIALLMAGQHTSSASGSWTLLHLAADPAVQEALYQEQVKNFRTPDGK

LRSMTYEDIRDLPVLDSVIRETLRIHPPIHSIMRKVRSDVPVPTTLSAPSKDSTYVVPKGYFVLASPAVS

QMDPRVWLNPGKWDPSRWSDPEGEAARAYDTYQDENGEKIDYGFGAVSKGTESPYQPFGAGRHRCIGEQF

AYLQLGAVITTIIRRVELRLEADTFPANNYHTMITMPKKPRNICYRRRAFD

>KDQ24437.1 hypothetical protein PLEOSDRAFT_1047295 [Pleurotus ostreatus PC15] 7e-158

MSNTRLALIALINAPVIIILLNAVWQLIAPRDPSKPPTVFHWLPIIGSAVGYGNDPLNFFLACREKYGDV

FTFILFGRRVTVALGAKGNNFVLGGKSTVLNAEDAYTHLTTPVFGKDVVYDVPNEKFMEQKRFVKVGLST

ENLRAYVGMIEDEVEEYMNNDANFRVYGMNDINDWGMFDVVSVLQEITILTASRTLQGKEVRSKLDKTFS

QLYNDLDGGFTPLNFLFPNLPLESYRKRDRANVKMTEFYVDIIQKRRQAGADSDHDMIAALIKQTYRDGT

PLKDHEIAHILIALLMAGQHTSSATGSWALLHIAENPDVGEALYQEQVKYFGTPDGNLRSMTYEELRELP

VLDAVIRETLRLHPPIHSIMRYVRDDVSVPPSLAAPSKDGVFVVPKGHYVLASPSVSQMDPRVWKDAQRF

DVSRWTDPDGVAAQAFKAYADENGEKIDYGFGAVSKGTESPYQPFGAGRHRCIGEQFAYLQLGTLISTMI

RKVELRLKDPVPAHNYHTMIVMPKKPRNILYRRRKFD

Animals

>sp|Q27589.2|CP4D2_DROME RecName: Full=Cytochrome P450 4d2; AltName: Full=CYPIVD2 Drosophila melanogaster 5E-22

MLGVVGVLLLVAFATLLLWDFLWRRRGNGILPGPRPLPFLGNLLMYRGLDPEQIMDFVKKNQRKYGRLYR

VWILHQLAVFSTDPRDIEFVLSSQQHITKNNLYKLLNCWLGDGLLMSTGRKWHGRRKIITPTFHFKILEQ

FVEIFDQQSAVMVEQLQSRADGKTPINIFPVICLTALDIIAETAMGTKINAQKNPNLPYVQAVNDVTNIL

IKRFIHAWQRVDWIFRLTQPTEAKRQDKAIKVMHDFTENIIRERRETLVNNSKETTPEEEVNFLGQKRRM

ALLDVLLQSTIDGAPLSDEDIREEVDTFMFEGHDTTTSAISFCLYEISRHPEVQQRLQQEIRDVLGEDRK

SPVTLRDLGELKFMENVIKESLRLHPPVPMIGRWFAEDVEIRGKHIPAGTNFTMGIFVLLRDPEYFESPD

EFRPERFDADVPQIHPYAYIPFSAGPRNCIGQKFAMLEMKSTVSKLLRHFELLPLGPEPRHSMNIVLRSA

NGVHLGLKPRA

>NP_001292049.1 cytochrome P450 6a9 [Musca domestica]2E-19

MWLSGILVGLVVTLISYLVLLMKRRLNYWHSRNVPCERPSLLLGNFKGMRTKYSFPEIWMNYYKKFKGSG

PFAGFFWFSHPAVFVLDLELIKNILTRDFNKFMDRGFFHNEQDDPLTGHLFFLDGLKWKSLRQKLTPTFS

SGKIAKMFPMVKNLTGRLMETMEEKLEDSEKQEDENAKHVLEMKDLLARFGTDVIGCCAFGIDCNSLSDP

EAKFHIMGQRLFSEPRNGQLGNALAFNFPELVQKLHMKVIPDEISEFFMDLVKKTIQSREENPTERDDFL

ALLMELRESKQIKTEDGEETKSLTLEEIAAQIVLFFLAGYETSSTTVGFALYELARHQEIQNRLRQEVNE

IWVKYGKDFTYESVKDMTYLQQVIQETLRLYIPVPVLNRKCLEDYPVPGHDEKYLIKKGMNVIIPVLAIQ

RDEEFFPQPEEFNPDNFEASRCKDRESVVYMPFGEGPRNCIGKRFGEMQTGLVLATLIKKFKFSTCPQTQ

IPVIFNKETYFLGAGHGIHLRVEKI

>XP_015136877.1 lanosterol 14-alpha demethylase isoform X1 [Gallus gallus] correct

MLSLLEVGGSLLERAAVGNPLSLLLAASAFALSLGYLFQLGYRRHVGADRTNHPPHIPSSIPFLGHAIAF

GKSPIEFLENAYDKYGPVFSFTMVGKTFTYLLGSDAAALLFNSKNEDLNAEDVYSRLTTPVFGKGVAYDV

PNVVFLEQKKMLKTGLNIAQFKQHVTLIEEETKEYFKAWGESGERNLFEAFSELIILTASHCLHGKEIRS

LLNEKVAQLYADLDGGFTHAAWLLPAWLPLPSFRRRDRAHRAIKNIFYKVIQKRRSSEEKEDDMLQTLLD

ASYKDGRPLTDDEIAGMLIGLLLAGQHTSSTTSAWLGFFIARDKAIQEQCYAEQKAVCGDDLPPLTYDQL

KDLSLLDRCLKETLRLRPPIMTIMRLAKTPQTVAGYNIPPGHQVCVSPTVNQRLKDSWKDALDFKPDRYL

RDNPAAGEKFAYIPFGAGRHRCIGENFAYVQIKTIWSTLLRLYEFDLVDGYFPSINYTTMIHTPNNPVIR

YKRRSL

>NP_000777.1 lanosterol 14-alpha demethylase isoform 1 precursor [Homo sapiens] Correct

MAAAAGMLLLGLLQAGGSVLGQAMEKVTGGNLLSMLLIACAFTLSLVYLIRLAAGHLVQLPAGVKSPPYI

FSPIPFLGHAIAFGKSPIEFLENAYEKYGPVFSFTMVGKTFTYLLGSDAAALLFNSKNEDLNAEDVYSRL

TTPVFGKGVAYDVPNPVFLEQKKMLKSGLNIAHFKQHVSIIEKETKEYFESWGESGEKNVFEALSELIIL

TASHCLHGKEIRSQLNEKVAQLYADLDGGFSHAAWLLPGWLPLPSFRRRDRAHREIKDIFYKAIQKRRQS

QEKIDDILQTLLDATYKDGRPLTDDEVAGMLIGLLLAGQHTSSTTSAWMGFFLARDKTLQKKCYLEQKTV

CGENLPPLTYDQLKDLNLLDRCIKETLRLRPPIMIMMRMARTPQTVAGYTIPPGHQVCVSPTVNQRLKDS

WVERLDFNPDRYLQDNPASGEKFAYVPFGAGRHRCIGENFAYVQIKTIWSTMLRLYEFDLIDGYFPTVNY

TTMIHTPENPVIRYKRRSK

>NP_064394.2 lanosterol 14-alpha demethylase [Mus musculus]

MVLLGLLQSGGWVLGQAMEQVTGGNLLSTLLIACAFTLSLVYLFRLAVGHMVQLPAGAKSPPHIYSPIPF

LGHAIAFGKSPIEFLENAYEKYGPVFSFTMVGKTFTYLLGSDAAALLFNSKNEDLNAEEVYGRLTTPVFG

KGVAYDVPNAIFLEQKKIIKSGLNIAHFKQYVPIIEKEAKEYFQSWGESGERNVFEALSELIILTASHCL

HGKEIRSQLNEKVAQLYADLDGGFTHAAWLLPAWLPLPSFRRRDRAHREIKNIFYKAIQKRRLSKEPAED

ILQTLLDSTYKDGRPLTDEEISGMLIGLLLAGQHTSSTTSAWMGFFLAKDKPLQEKCYLEQKAVCGEDLP

PLTYDQLKDLNLLDRCIKETLRLRPPIMTMMRMAKTPQTVAGYTIPPGHQVCVSPTVNQRLKDSWAERLD

FNPDRYLQDNPASGEKFAYVPFGAGRHRCVGENFAYVQIKTIWSTMLRLYEFDLINGYFPTVNYTTMIHT

PENPVIRYKRRSK

Plants

>NP_172633.1 CYTOCHROME P450 51G1 [Arabidopsis thaliana]1E-71 correct

MELDSENKLLKTGLVIVATLVIAKLIFSFFTSDSKKKRLPPTLKAWPPLVGSLIKFLKGPIIMLREEYPK

LGSVFTVNLVHKKITFLIGPEVSAHFFKASESDLSQQEVYQFNVPTFGPGVVFDVDYSVRQEQFRFFTEA

LRVNKLKGYVDMMVTEAEDYFSKWGESGEVDIKVELERLIILTASRCLLGREVRDQLFDDVSALFHDLDN

GMLPISVLFPYLPIPAHRRRDRAREKLSEIFAKIIGSRKRSGKTENDMLQCFIESKYKDGRQTTESEVTG

LLIAALFAGQHTSSITSTWTGAYLMRYKEYFSAALDEQKNLIAKHGDKIDHDILSEMDVLYRCIKEALRL

HPPLIMLMRASHSDFSVTARDGKTYDIPKGHIVATSPAFANRLPHIFKDPDTYDPERFSPGREEDKAAGA

FSYIAFGGGRHGCLGEPFAYLQIKAIWSHLLRNFELELVSPFPEIDWNAMVVGVKGNVMVRYKRRQLS

>OAP10887.1 CYP51G2 [Arabidopsis thaliana]7E-69

MDWDYYTLLKTSVAIIIVFVVAKLITSSKSKKKTSVVPLPPVLKAWPPFIGSLIRFMKGPIVLLREEYPK

LGSVFTVKLLHKNITFLIGPEVSSHFFNAYESELSQKEIYKFNVPTFGPGVVFDVDYPVRMEQFRFFSSA

LKVNKLRGYVDQMTKETEDYFSKWGESGEVDLKAELERLITLTASRCLLGREVRDQLFDDVAPLFHDLDK

GMQPISVIFPKLPIPAHNCRDRARGKIAKIFSNIIATRKRSGDKSENDMLQCFIDSKYKDGRETTESEVT

GLLIAGLFAGQHTSSITATWTGAYLIQNKHWWSAALDEQKKLIGKHGDKIDYDVLSEMDFLFRSAKEALR

LHPPKILLLRTVHSDFTVTTPEGKQYEIPKGHIVATSPAFANRLPHVYKDPENFDPDRFSKEREEDKAAG

SCSYISLGAGRHECPGGSFAFLQIKAVWCHLLRNFELELVSPFPEINWNALVVGAKGNVMVRYKRRPLS

>AML47779.1 putative cytochrome P450 [Triticum aestivum] 1 correct

MEVATVNWVAFVVPIAVATIIIFISAMMARGRRRTERSPRARPPPVAAGAPLVGVLPWLLAKGPLQVIRD

AHAELGSVFTVRLLHREVTFLVGPDVSSHFYQGLDSEVSQDEVSRFTVPTFGPGVAFDVDLATRREQIRF

FGDAMKPAKLRTYAGLMVREVEEYFTRWGEMGTVDLKQELGHLVTLVASRCLFGEEVRSKMLREAATHLR

ELNDGMRLVTILFPHLPIPAHRRRDRARARLGEIFSDMVRSRRESGRPVDDMLQCLIDSRYKDGRATTDT

ELVGMLVSALFAGQHTSSSTGTWTGARLLAGANAEHLRAAVREQERVVARHGDRVDYEVLQEMETLHRSV

KEALRLHPPAMMLLRHARRSFVVRTREGDEYEVPEGRTVASPMVIHNRLPHVYRDPERYEPGRFGPGRGE

DGAGGALSYTAFGGGRHACVGEAFAYMQIKVIWSHLLRNFEMEMVSPFPETDWNVVMPGPKGKVMLRYKR

RKKMSPTAQITNPHTRYVL

>XP_828695.1 lanosterol 14-alpha-demethylase [Trypanosoma brucei brucei TREU927]3E-46

MLLEVAIFLLTALALYSFYFVKSFNVTRPTDPPVYPVTVPILGHIIQFGKSPLGFMQECKRQLKSGIFTI

NIVGKRVTIVGDPHEHSRFFLPRNEVLSPREVYSFMVPVFGEGVAYAAPYPRMREQLNFLAEELTIAKFQ

NFVPAIQHEVRKFMAANWDKDEGEINLLEDCSTMIINTACQCLFGEDLRKRLDARRFAQLLAKMESSLIP

AAVFLPILLKLPLPQSARCHEARTELQKILSEIIIARKEEEVNKDSSTSDLLSGLLSAVYRDGTPMSLHE

VCGMIVAAMFAGQHTSSITTTWSMLHLMHPANVKHLEALRKEIEEFPAQLNYNNVMDEMPFAERCARESI

RRDPPLLMLMRKVMADVKVGSYVVPKGDIIACSPLLSHHDEEAFPEPRRWDPERDEKVEGAFIGFGAGVH

KCIGQKFGLLQVKTILATAFRSYDFQLLRDEVPDPDYHTMVVGPTASQCRVKYIRRKAAAA

>XP_001134568.1 cytochrome P450 family protein [Dictyostelium discoideum AX4]2E-71

MIGTIAVLVIAILVIFAFKKSPSNIPPIVETIPFIGCFYQFAKNPLQLVRNSYDRLGEIFTLHLMGFKMT

FVLGPEAQALFFRGTDEELSPKEAYRFVTPVFGKGVVYDSETEIMYEQLRFVKNGLVLSQLKKAVGIIQE

ETEKYFETKWGDSGEIDLLYEMNKLTILTASRCLMGKSINKSLGQSGQLADLYHELEEGLNPISFFFPNL

PLPSFKKRDAARAKVAAIFHSIIQERRRSTDDSVDDVLYTLMNSKYKDGSVLEDEQIVGLMIGLLFAGQH

TSSITLTYTIFYLLNNLEYFDETQKDINDIVQKENQGEINFDGLKRMNRLETVIREVLRLHPPLIFLMRK

VMTPMEYKGKTIPAGHILAVSPQVGMRLPTVYKNPDSFEPKRFDVEDKTPFSFIAFGGGKHGCPGENFGI

LQIKTIWTVLSTKYNLEVGPVPPTDFTSLVAGPKGPCMVKYSKKQK

>FGSG_01000

MGLLQELAGHPLAQQFQELPLGQQVGIGFAVFLVLSVVLNVLNQLLFRNPNEPPMVFHWFPFVGSTITYG

MDPPTFFRENRAKHGDVFTFILLGKKTTVAVGPAGNDFILNGKLKDVCAEEIYTVLTTPVFGKDVVYDCP

NAKLMEQKKFMKIALTTEAFRSYVPIISSEVRDYFKRSPDFKGKSGIADIPKKMAEITIFTASHALQGSA

IRSKFDESLAALYHDLDMGFTPINFMLHWAPLPWNRKRDHAQRTVAKIYMDTIKERRAKGNNESEHDMMK

HLMNSTYKNGIRVPDHEVAHMMIALLMAGQHSSSSTSSWIMLRLAQYPHIMEELYQEQVKNLGADLPPLT

YEDLAKLPLNQAIVKETLRLHAPIHSIMRAVKSPMPVPGTKYVIPTSHTLLAAPGVSATDSAFFPNPDEW

DPHRWEADSPNFPRMASKGEDEEKIDYGYGLVSKGSASPYLPFGAGRHRCIGEHFANAQLQTIVAEVVRE

FKFRNVDGGHTLIDTDYASLFSRPLEPANIHWERRQ

>FGSG_11024

MESLYETLRTLPLSVSIPLTTSIIIILSIVTNVVKQLWFPNPHRPPVVFHIFPFIGSTVQYGIDPYAFFF

DCRDKYGDCFTFILLGKSTTVFLGPKGNDFILNGKHADLNAEDVYGKLTTPVFGEEVVYDCSNARFMDQK

RLLKLGLTTDSLRCYIPKFVKEVEDYVKNSPYFKGDTGIVNITEVMAEITIYTASGSLLGNEVRSMFDST

FATLYRHLDDGFQPINFVMPGLPLPQNFRRNHARKVMEKLFSDIISKRRETGNQGDETDMIWMLMNAQYK

DGEPLPDHHAARMLIAILMGGQHNTAVSGAWLLLNLAHKPHLVQELYEEQTQVLGSPQEPLTWENLQKLT

LNGQVIKETLRLHSPIHSILRQVKSPMRVPGTEWVVPPSHTLLSSPGTMARSEEFFPRPSEWDPHRWDKI

EPLVKTAEDGQTVDYGFGVMSKSVSSPYLPFGAGRHRCVGENYAYAQLGAIVATFIRLVHIEQPDPKAPL

PAPDYSSMFSRPMNPAEIRWRRRETVE

>FGSG_04092

MFHLLIYPLWVLVALFAVIIANLLYQQLPRRPDEPPLVFHWFPFFGNAVAYGLDPCGFFEKCREKHGDVF

TFILFGRKIVACLGVDGNDFVLNSRLQDANAEEVYGPLTIPVFGSDVVYDCPNSKLMEQKKFVKFGLTQK

ALESHVQLIEREVLDYVETDPSFSGRTSTIDVPKAMAEITIFTASRSLQGEEVRRKLTAEFAALYHDLDL

GFRPVNFLFPWLPLPHNRKRDAAHIKMREVYMDIINDRRKGGIRTEDGTDMIANLMGCTYKNGQPVPDKE

IAHMMITLLMAGQHSSSSASSWIVLHLASSPDITEELYQEQLVNLSVNGALPPLQYSDLDKLPLLQNVVK

ETLRVHSSIHSILRKVKRPMQVPNSPYTITTDKVIMASPTVTAMSEEYFENAKTWNPHRWDNRAKEEVDT

EDVIDYGYGAVSKGTKSPYLPFGAGRHRCIGEKFAYVNLGVIVATLVRNFRLSTIDGRPGVPETDYTSLF

SRPAQPAFIRWERRKKI

>FgTRI4

MIDQDWIKSLLNIPVSHVAGIFAASTVIYFLSSCFYNLYLHPLRKIPGPKLAAIGPYLEFYHEVIRDGQY

LWEISKMHDKYGPIVRVNAREVHIRDSSYYTTIYTAGSRKTNKDPATVGAFDVPSATAATVDHDHHRSRR

GYLNPYFSKRTITNLEPFIHERVTKLLTRFQQHLDDDQVLSLDGAFCALTADVITNRFYGKHNDYLSLPD

FHFVVRDGFLGLTKIYHLARFLPGLVTILKRLPYSCIRMIAPSVCDLLQMRDEIQDRGGEEFLSNKSHEA

KSSILFGALADSHIPSHERTVERMLDEGTVILFAGTETTSRTLAITVLYLLTHPECLKKLREELNSLPPV

KDGQYSLATLENLPYLNGVIHEGFRLAFGPISRSGRVATQENLKYKEHVIPKGTPISQSTYFMHTDPKNF

PEPEKFKPERWIEAQQKGIPLKKYITNFSQGSRQCIGYTMAFAEMYLALSRIARAYDIELYDTTKADIDM

THARIVGYPKAIPGKKEHLGEVRVKVLKAL

>Yeast ERG11

MSATKSIVGEALEYVNIGLSHFLALPLAQRISLIIIIPFIYNIVWQLLYSLRKDRPPLVFYWIPWVGSAVVYGMKPYEFFEECQKKYGDIFSFVLLGRVMTVYLGPKGHEFVFNAKLADVSAEAAYAHLTTPVFGKGVIYDCPNSRLMEQKKFVKGALTKEAFKSYVPLIAEEVYKYFRDSKNFRLNERTTGTIDVMVTQPEMTIFTASRSLLGKEMRAKLDTDFAYLYSDLDKGFTPINFVFPNLPLEHYRKRDHAQKAISGTYMSLIKERRKNNDIQDRDLIDSLMKNSTYKDGVKMTDQEIANLLIGVLMGGQHTSAATSAWILLHLAERPDVQQELYEEQMRVLDGGKKELTYDLLQEMPLLNQTIKETLRMHHPLHSLFRKVMKDMHVPNTSYVIPAGYHVLVSPGYTHLRDEYFPNAHQFNIHRWNNDSASSYSVGEEVDYGFGAISKGVSSPYLPFGGGRHRCIGEHFAYCQLGVLMSIFIRTLKWHYPEGKTVPPPDFTSMVTLPTGPAKIIWEKRNPEQKIGGRHHHHHH

>Pyricularia oryzae 70-15

MGLLQDTTGPLVDAFYQLGTGAQVGVAFVSFIFLSVFFHVAQQIFFKNPHEPPVVFSWFPVVGSTVTYGK

DPPQFFRDMAKKYGNIFTFILLGKKTTVYIGTEGNEFILNGKLRDVNAEEIYGPMTTPVFGKDVVYDCPN

AKLMEQKKFMKIALTTEAFRSYVPIIADEVSSYLKRTPAFKGPSGVVNIPPKMAEITIFTASHALQGKEI

RDQFDETLADLYHDLDMGFHPVNFKLHWLPLPRNIRRDKAQKTIAKIYMDTIQRRRAKGKDSEAKDMMYH

LMNSTYKNGTPVPDHEIAHMMIALLMAGQHSSSSTSSWIMLRLASRPDIMEELYQEQVRALGADLPPLRY

EDLANLPLHLAVIKETLRLHAPINSILRAVKQDLPVPGTNYVIAKDTTVLAAPGYSAGDPNHFPEPELWE

PHRWEADSRLAPRISMSNDNDEEEKIDYGYGLVSKGTTSPYLPFGAGRHRCIGEHFANVQLQTIVAMIVR

EFKFRNVDGSGKVVGTNYASLFSRPEEPAKIYWERR

>Metarhizium anisopliae

MGVLQQVASHPLSQQFQTLGLGSQIAVALGGFIALAVALNVASQILFKNPNEPPLVFHWFPFIGSTVTYG

MDPPKFFKENRAKFGDVFTFVLLGKKTTVAVGPAGNDFILNGKLKDVNAEEIYTVLTTPVFGRDVVYDCP

NAKLMEQKKFMKIALTTEAFRSYVPIISGEVQSYFKRDPGFKGKSGIVDIPKKMAEITIFTASHALQGSA

IRGKFDESLAALYHDLDMGFTPINFMLHWAPLPWNRKRDHAQRTVAKIYMDTIKERRAKGDDDKELDIMK

HLMNSTYKNGTPVPDHEVAHMMIALLMAGQHSSSSTSSWIMLRLAQNPHLVEELYQEQIKALGADLPPLT

YEDLSKLPLNQAIIKETLRLHAPIHSIMRAVKQPMPVPGTKYVIPTTHTLLAAPGVSASDPTYFPSPESW

DPHRWDTDSPNAPTIVRNVAEEEEKVDYGYGLVSKGAASPYLPFGAGRHRCIGEHFANVQLQTIVAETVR

LFKLSNVDGSNTIIGTDYASLFSRPLEPARIRWERRD

>Verticillium dahliae

MGLLQEITGPAAQQFQTLGTASQVGVVVFGVVLLSVVLHVANQLLFKNPNEPPVVFSWFPFIGNTITYGM

DPPAFFEANKKKYGDVFTFILLGSKTTVAVGPQGNDFILNGKQRDVNAEEIYSVLTTPVFGRDVVYDCPN

AKLMEQKKFMKIALTTEAFRSYVPIISHEVQNFFKTSKYFKEQSGVVNIPPRMAEITIFTASHALQGSEI

RNKFDTSLAELYHDLDMGFSAINFALHWAPLPWNRKRDAAQKAVAQIFMDTIKDRRESGRTDGLDMISHL

MRSTYKNGINVPDHEIAHMMIALLMAGQHSSSSTSSWIMARLAQNPSVIDDLYKEQVEALGADLPPLTYE

DLSKLPLNQAIIKETLRLHAPIHSILRKVKSPMPVPGTKYVVPTSHNLLSAPGTSATDPTYFPNPAIWDP

YRWLPDSPNAPTFSRAENADEEKIDYGYGLVSKGANSPYLPFGAGRHRCIGEQFANVQLQTITAEIVRLV

KFSNADGSSNINGTDYASLFSRPLEPVNIRYERREKA

>Fusarium oxysporum f. sp. lycopersici]

MGLLQELASHPLAQQYQELPLGQQIGIGFGAFIILSVVLNVLNQLLFKNRNEPPLVFHWFPFVGSTITYG

MDPPKFFKENRAKHGDVFTFVLLGKKTTVAVGPTGNDFILNGKLKDVSAEEIYTVLTTPVFGKDVVYDCP

NAKLMEQKKFMKIALTTEAFRSYVPIISAEVRDYFKKSPDFKGKSGIVDIPKKMAEITIFTASHALQGSV

IRNKFDESLAALYHDLDMGFTPINFMLHWAPLPWNRKRDHAQRTVAKIYMDTIKERRAKDNDDTEHDMMK

HLMNSTYKNGTPVPDHEVAHMMIALLMAGQHSSSSTSSWIMLRLAQYPHIMEELYQEQVRELGADLPPLT

YDNLAKLPLNQAIIKETLRLHAPIHSIMRAVKSPMPVPGTKYTIPTSHTLLAAPGVSATDSAYFPNPDEW

DPHRWEVDSPNFPRMATRGDDEEKIDYGYGLVSKGSASPYLPFGAGRHRCIGEHFANAQLQTIVAEVVRE

FKFRNVDGGNTLIDTDYASLFSRPLEPANIHWERRQQ

>Aspergillus niger

MGLLAVVLDSLCERCSNTSVLVVLGFGLLSLLAVSVIFNILRQLFFKNPNEPPLVFHWFPFIGSTISYGM

DPYKFFFDCRAKYGDIFTFILLGKKTTVYLGTRGNDFILNGKLRDVCAEEVYSPLTTPVFGRNVVYDCPN

AKLMEQKKFVKFGLTSDALRSYVRLITGEVENFVEHSAAFKGSSGVFDVCKTIAEITIYTASRSLQGKEV

RSRFDSTFAELYHDLDMGFAPINFMLPWAPLPHNRKRDAAQKKMTETYMEIIKERRNGGNKKDSEDMVWN

LMSCIYKDGTPVPDEEIAHMMIALLMAGQHSSSSTASWIVLHLARNPQIMEELYAEQIRVLGSDLPPLTY

DNLQKLDLHAKVIKETLRIHAPIHSIIRAVKNPMPVDGTPYVIPNSHNVLSSPGVTARSEEYFPNPLKWD

PHRWDETIAASAEEEDQIDYGYGLVSKGTNSPYLPFGAGRHRCIGEQFAYVQLGAITAALVRLFKFRNLP

NVKDIPETDYSSLFSKPAGKSIIQFEKRATTIKS

>Blumeria graminis f. sp. hordei

MGISESFMFPYLQPLLQLGFGIALASGIISLLLLLTFLNVLKQLLFKNPNEPPIVFHWIPIIGSTISYGM

NPYKFFHESQAKYGNIFTFILLGKKTTVYLGRQGNNFILNGKLRDVNAEEIYTVLTTPVFGTDVVYDCPN

SKLMEQKKFMKAALTTEAFRSYVPIIQNEVKSFIEKCDDFRKSKGIINIDAVMAEITIYTASHTLQGKEV

RDRFDSSLAVLYHDLDMGFTPINFMLHWAPLPHNRARDHAQRTVAKIYMEIINSRRTQKETDDSNLDIMW

QLMRSSYKDGTPVPDKEIAHMMIALLMAGQHSSSSSSTWIMLWLAARPDITEELYQEQLELLGSELPPLK

YEDLSKLSLHQNVLKEVLRLHAPIHSILRKVKNPMPVPGTSYVIPKTHSLLAAPGWTSRDASYFPNPLKW

DPHRWDTGSGGVIGTDMEDEKFDYGYGLISTGAASPYLPFGAGRHRCIGEQFATVQLVTIMATMVRSFKF

HNLDGRNSVAETDYSSMFSRPMAPATIAWEKRDKKDKTEC

>Crucibulum laeve

MSFGAFNSSSIPLPDGWSGYLAHAQEHWLPENSRTLLILLINAPVLAILLNALRQVIMPRDPSLPPEVFH

WLPIIGSAVQYGNDPLNFFFKCQEKYGDVFTFILFGRRVTVALGPKGNNFILGGKSTVFNAEDAYTHLTT

PIFGKDVVYDVPNEKFMEQKRFVKVGLSTDNLRAYVGMIEEEVDEFLKIDPSFRVYQTNDINEWGTFDVI

MALQEITILTASRTLQGKEVREGLNKTFAHLYSDLDGGFTPLNFLFPNLPLESYRKRDRAHQKISQFYIE

IIQKRREGGEDHEHDMIAALMQQTYRNGEHLKDHEIAHIMIALLMAGQHTSSATGSWLLLHIANNPDVAE

ALYQEQVKFFSTPDGKLRSPTYEELRQLPLMDAVIRETLRMHPPIHSIMRYVRDDVPIPGTLSAPSKDTT

YVVPKGHYVLASPAVSQMDPKVWKDTSKWDPYRWADPEGMAARAFDTYADESGEKIDYGFGAVSKGTESP

YQPFGAGKHRCIGEQFAYLQLGTIVATVIRRMEMRIEKVPEHNYHTMITMPKKPRIISYRRRNFD

>Agaricus bisporus

MSYAANGTTPILDAWSGYVAYAQTQLTPWSSRMTILSIATIPVLVVVLNVLSQLLQFRKSSEPPVVFHWL

PFIGSAIGYGTDPLNFFNRCRDKYGDVFTFVLFGRRVTVALGSKGNNFILGGKSIVFNAEDAYTHLTTPV

FGKDVVYDIPNDLFMEQKKFVKVGLSSENLRAYVGMIEEEVEEYLGSDSAFSAFQSNDANEWGHFDVLKV

MSGITILTASRTLQGKEVRAGLDKTFAEIFNDLDGGFTPLNFLFPNLPLASYKKRDVAHKKISDFFSSIV

KGRRAETNPDHEYDMIASLMNQKYRDGRQLKDHEIAHIMIALLMAGQHTSSATGSWVLCHLARRPDVCDA

LYKEQVKNFSNPDGSFRSMTHEELRQLPILDSVIRETLRLHPPIHSVMRHVRDDMPVPATLAAPGKDTTY

VIPKGHYVLASPLVSQLDRTIWKDAETWEPLRWSDPEGIAAQANKLYEDVQGEKIDYGFGAVSKGTESPY

QPFGAGKHRCIGEQFAYLQLGTIITTFIRHMELKMDKLPEHNYQTLITLPKAPREVYYRRRK

>Serpula lacrymans

MSAFLDYVLNYVVTSPGRAILLALLYTPIAAIVLNVLRQLVVPRDPSLPPDVFHWIPIVGSAIEYGNDPL

KFFFKCRDKYGDVFTFVLLGRKVTVALGAKGNNFILGGKSTAFSAEDAYTHLTTPVFGKDVVYDVPNEVF

MEQKKFVKVGLSTENFRSYVGMIEDEVDEFMRNDPSFLIYQMNDINEWGRFDAATAMSEITILTASRTLQ

GKEVRSNLDKSFSELYNALDGGFTPLNFMFPNLPLDSYRRRDEAHKKMSEFYVNIIRKRKEGNNDHEHDM

IAALRDQTYRNGRPLPDHEIAHIMIALLMAGQHTSSASGSWTLLHLAADPAVQEALYQEQVKNFRTPDGK

LRSMTYEDIRDLPVLDSVIRETLRIHPPIHSIMRKVRSDVPVPTTLSAPSKDSTYVVPKGYFVLASPAVS

QMDPRVWLNPGKWDPSRWSDPEGEAARAYDTYQDENGEKIDYGFGAVSKGTESPYQPFGAGRHRCIGEQF

AYLQLGAVITTIIRRVELRLEADTFPANNYHTMITMPKKPRNICYRRRAFD

>Pleurotus ostreatus

MSNTRLALIALINAPVIIILLNAVWQLIAPRDPSKPPTVFHWLPIIGSAVGYGNDPLNFFLACREKYGDV

FTFILFGRRVTVALGAKGNNFVLGGKSTVLNAEDAYTHLTTPVFGKDVVYDVPNEKFMEQKRFVKVGLST

ENLRAYVGMIEDEVEEYMNNDANFRVYGMNDINDWGMFDVVSVLQEITILTASRTLQGKEVRSKLDKTFS

QLYNDLDGGFTPLNFLFPNLPLESYRKRDRANVKMTEFYVDIIQKRRQAGADSDHDMIAALIKQTYRDGT

PLKDHEIAHILIALLMAGQHTSSATGSWALLHIAENPDVGEALYQEQVKYFGTPDGNLRSMTYEELRELP

VLDAVIRETLRLHPPIHSIMRYVRDDVSVPPSLAAPSKDGVFVVPKGHYVLASPSVSQMDPRVWKDAQRF

DVSRWTDPDGVAAQAFKAYADENGEKIDYGFGAVSKGTESPYQPFGAGRHRCIGEQFAYLQLGTLISTMI

RKVELRLKDPVPAHNYHTMIVMPKKPRNILYRRRKFD

>Drosophila melanogaster

MLGVVGVLLLVAFATLLLWDFLWRRRGNGILPGPRPLPFLGNLLMYRGLDPEQIMDFVKKNQRKYGRLYR

VWILHQLAVFSTDPRDIEFVLSSQQHITKNNLYKLLNCWLGDGLLMSTGRKWHGRRKIITPTFHFKILEQ

FVEIFDQQSAVMVEQLQSRADGKTPINIFPVICLTALDIIAETAMGTKINAQKNPNLPYVQAVNDVTNIL

IKRFIHAWQRVDWIFRLTQPTEAKRQDKAIKVMHDFTENIIRERRETLVNNSKETTPEEEVNFLGQKRRM

ALLDVLLQSTIDGAPLSDEDIREEVDTFMFEGHDTTTSAISFCLYEISRHPEVQQRLQQEIRDVLGEDRK

SPVTLRDLGELKFMENVIKESLRLHPPVPMIGRWFAEDVEIRGKHIPAGTNFTMGIFVLLRDPEYFESPD

EFRPERFDADVPQIHPYAYIPFSAGPRNCIGQKFAMLEMKSTVSKLLRHFELLPLGPEPRHSMNIVLRSA

NGVHLGLKPRA

>Musca domestica

MWLSGILVGLVVTLISYLVLLMKRRLNYWHSRNVPCERPSLLLGNFKGMRTKYSFPEIWMNYYKKFKGSG

PFAGFFWFSHPAVFVLDLELIKNILTRDFNKFMDRGFFHNEQDDPLTGHLFFLDGLKWKSLRQKLTPTFS

SGKIAKMFPMVKNLTGRLMETMEEKLEDSEKQEDENAKHVLEMKDLLARFGTDVIGCCAFGIDCNSLSDP

EAKFHIMGQRLFSEPRNGQLGNALAFNFPELVQKLHMKVIPDEISEFFMDLVKKTIQSREENPTERDDFL

ALLMELRESKQIKTEDGEETKSLTLEEIAAQIVLFFLAGYETSSTTVGFALYELARHQEIQNRLRQEVNE

IWVKYGKDFTYESVKDMTYLQQVIQETLRLYIPVPVLNRKCLEDYPVPGHDEKYLIKKGMNVIIPVLAIQ

RDEEFFPQPEEFNPDNFEASRCKDRESVVYMPFGEGPRNCIGKRFGEMQTGLVLATLIKKFKFSTCPQTQ

IPVIFNKETYFLGAGHGIHLRVEKI

>Gallus gallus

MLSLLEVGGSLLERAAVGNPLSLLLAASAFALSLGYLFQLGYRRHVGADRTNHPPHIPSSIPFLGHAIAF

GKSPIEFLENAYDKYGPVFSFTMVGKTFTYLLGSDAAALLFNSKNEDLNAEDVYSRLTTPVFGKGVAYDV

PNVVFLEQKKMLKTGLNIAQFKQHVTLIEEETKEYFKAWGESGERNLFEAFSELIILTASHCLHGKEIRS

LLNEKVAQLYADLDGGFTHAAWLLPAWLPLPSFRRRDRAHRAIKNIFYKVIQKRRSSEEKEDDMLQTLLD

ASYKDGRPLTDDEIAGMLIGLLLAGQHTSSTTSAWLGFFIARDKAIQEQCYAEQKAVCGDDLPPLTYDQL

KDLSLLDRCLKETLRLRPPIMTIMRLAKTPQTVAGYNIPPGHQVCVSPTVNQRLKDSWKDALDFKPDRYL

RDNPAAGEKFAYIPFGAGRHRCIGENFAYVQIKTIWSTLLRLYEFDLVDGYFPSINYTTMIHTPNNPVIR

YKRRSL

>Homo sapiens

MAAAAGMLLLGLLQAGGSVLGQAMEKVTGGNLLSMLLIACAFTLSLVYLIRLAAGHLVQLPAGVKSPPYI

FSPIPFLGHAIAFGKSPIEFLENAYEKYGPVFSFTMVGKTFTYLLGSDAAALLFNSKNEDLNAEDVYSRL

TTPVFGKGVAYDVPNPVFLEQKKMLKSGLNIAHFKQHVSIIEKETKEYFESWGESGEKNVFEALSELIIL

TASHCLHGKEIRSQLNEKVAQLYADLDGGFSHAAWLLPGWLPLPSFRRRDRAHREIKDIFYKAIQKRRQS

QEKIDDILQTLLDATYKDGRPLTDDEVAGMLIGLLLAGQHTSSTTSAWMGFFLARDKTLQKKCYLEQKTV

CGENLPPLTYDQLKDLNLLDRCIKETLRLRPPIMIMMRMARTPQTVAGYTIPPGHQVCVSPTVNQRLKDS

WVERLDFNPDRYLQDNPASGEKFAYVPFGAGRHRCIGENFAYVQIKTIWSTMLRLYEFDLIDGYFPTVNY

TTMIHTPENPVIRYKRRSK

>Mus musculus

MVLLGLLQSGGWVLGQAMEQVTGGNLLSTLLIACAFTLSLVYLFRLAVGHMVQLPAGAKSPPHIYSPIPF

LGHAIAFGKSPIEFLENAYEKYGPVFSFTMVGKTFTYLLGSDAAALLFNSKNEDLNAEEVYGRLTTPVFG

KGVAYDVPNAIFLEQKKIIKSGLNIAHFKQYVPIIEKEAKEYFQSWGESGERNVFEALSELIILTASHCL

HGKEIRSQLNEKVAQLYADLDGGFTHAAWLLPAWLPLPSFRRRDRAHREIKNIFYKAIQKRRLSKEPAED

ILQTLLDSTYKDGRPLTDEEISGMLIGLLLAGQHTSSTTSAWMGFFLAKDKPLQEKCYLEQKAVCGEDLP

PLTYDQLKDLNLLDRCIKETLRLRPPIMTMMRMAKTPQTVAGYTIPPGHQVCVSPTVNQRLKDSWAERLD

FNPDRYLQDNPASGEKFAYVPFGAGRHRCVGENFAYVQIKTIWSTMLRLYEFDLINGYFPTVNYTTMIHT

PENPVIRYKRRSK

>Arabidopsis thaliana1

MELDSENKLLKTGLVIVATLVIAKLIFSFFTSDSKKKRLPPTLKAWPPLVGSLIKFLKGPIIMLREEYPK

LGSVFTVNLVHKKITFLIGPEVSAHFFKASESDLSQQEVYQFNVPTFGPGVVFDVDYSVRQEQFRFFTEA

LRVNKLKGYVDMMVTEAEDYFSKWGESGEVDIKVELERLIILTASRCLLGREVRDQLFDDVSALFHDLDN

GMLPISVLFPYLPIPAHRRRDRAREKLSEIFAKIIGSRKRSGKTENDMLQCFIESKYKDGRQTTESEVTG

LLIAALFAGQHTSSITSTWTGAYLMRYKEYFSAALDEQKNLIAKHGDKIDHDILSEMDVLYRCIKEALRL

HPPLIMLMRASHSDFSVTARDGKTYDIPKGHIVATSPAFANRLPHIFKDPDTYDPERFSPGREEDKAAGA

FSYIAFGGGRHGCLGEPFAYLQIKAIWSHLLRNFELELVSPFPEIDWNAMVVGVKGNVMVRYKRRQLS

>Arabidopsis thaliana2

MDWDYYTLLKTSVAIIIVFVVAKLITSSKSKKKTSVVPLPPVLKAWPPFIGSLIRFMKGPIVLLREEYPK

LGSVFTVKLLHKNITFLIGPEVSSHFFNAYESELSQKEIYKFNVPTFGPGVVFDVDYPVRMEQFRFFSSA

LKVNKLRGYVDQMTKETEDYFSKWGESGEVDLKAELERLITLTASRCLLGREVRDQLFDDVAPLFHDLDK

GMQPISVIFPKLPIPAHNCRDRARGKIAKIFSNIIATRKRSGDKSENDMLQCFIDSKYKDGRETTESEVT

GLLIAGLFAGQHTSSITATWTGAYLIQNKHWWSAALDEQKKLIGKHGDKIDYDVLSEMDFLFRSAKEALR

LHPPKILLLRTVHSDFTVTTPEGKQYEIPKGHIVATSPAFANRLPHVYKDPENFDPDRFSKEREEDKAAG

SCSYISLGAGRHECPGGSFAFLQIKAVWCHLLRNFELELVSPFPEINWNALVVGAKGNVMVRYKRRPLS

>Triticum aestivum

MEVATVNWVAFVVPIAVATIIIFISAMMARGRRRTERSPRARPPPVAAGAPLVGVLPWLLAKGPLQVIRD

AHAELGSVFTVRLLHREVTFLVGPDVSSHFYQGLDSEVSQDEVSRFTVPTFGPGVAFDVDLATRREQIRF

FGDAMKPAKLRTYAGLMVREVEEYFTRWGEMGTVDLKQELGHLVTLVASRCLFGEEVRSKMLREAATHLR

ELNDGMRLVTILFPHLPIPAHRRRDRARARLGEIFSDMVRSRRESGRPVDDMLQCLIDSRYKDGRATTDT

ELVGMLVSALFAGQHTSSSTGTWTGARLLAGANAEHLRAAVREQERVVARHGDRVDYEVLQEMETLHRSV

KEALRLHPPAMMLLRHARRSFVVRTREGDEYEVPEGRTVASPMVIHNRLPHVYRDPERYEPGRFGPGRGE

DGAGGALSYTAFGGGRHACVGEAFAYMQIKVIWSHLLRNFEMEMVSPFPETDWNVVMPGPKGKVMLRYKR

RKKMSPTAQITNPHTRYVL

>Trypanosoma brucei

MLLEVAIFLLTALALYSFYFVKSFNVTRPTDPPVYPVTVPILGHIIQFGKSPLGFMQECKRQLKSGIFTI

NIVGKRVTIVGDPHEHSRFFLPRNEVLSPREVYSFMVPVFGEGVAYAAPYPRMREQLNFLAEELTIAKFQ

NFVPAIQHEVRKFMAANWDKDEGEINLLEDCSTMIINTACQCLFGEDLRKRLDARRFAQLLAKMESSLIP

AAVFLPILLKLPLPQSARCHEARTELQKILSEIIIARKEEEVNKDSSTSDLLSGLLSAVYRDGTPMSLHE

VCGMIVAAMFAGQHTSSITTTWSMLHLMHPANVKHLEALRKEIEEFPAQLNYNNVMDEMPFAERCARESI

RRDPPLLMLMRKVMADVKVGSYVVPKGDIIACSPLLSHHDEEAFPEPRRWDPERDEKVEGAFIGFGAGVH

KCIGQKFGLLQVKTILATAFRSYDFQLLRDEVPDPDYHTMVVGPTASQCRVKYIRRKAAAA

>Dictyostelium discoideum

MIGTIAVLVIAILVIFAFKKSPSNIPPIVETIPFIGCFYQFAKNPLQLVRNSYDRLGEIFTLHLMGFKMT

FVLGPEAQALFFRGTDEELSPKEAYRFVTPVFGKGVVYDSETEIMYEQLRFVKNGLVLSQLKKAVGIIQE

ETEKYFETKWGDSGEIDLLYEMNKLTILTASRCLMGKSINKSLGQSGQLADLYHELEEGLNPISFFFPNL

PLPSFKKRDAARAKVAAIFHSIIQERRRSTDDSVDDVLYTLMNSKYKDGSVLEDEQIVGLMIGLLFAGQH

TSSITLTYTIFYLLNNLEYFDETQKDINDIVQKENQGEINFDGLKRMNRLETVIREVLRLHPPLIFLMRK

VMTPMEYKGKTIPAGHILAVSPQVGMRLPTVYKNPDSFEPKRFDVEDKTPFSFIAFGGGKHGCPGENFGI

LQIKTIWTVLSTKYNLEVGPVPPTDFTSLVAGPKGPCMVKYSKKQK

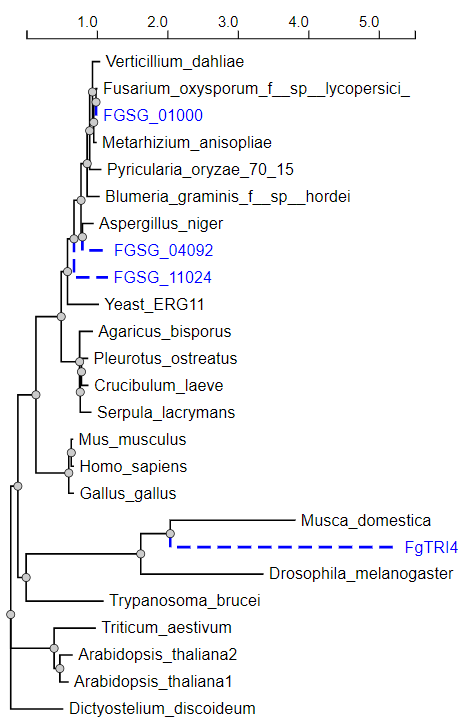

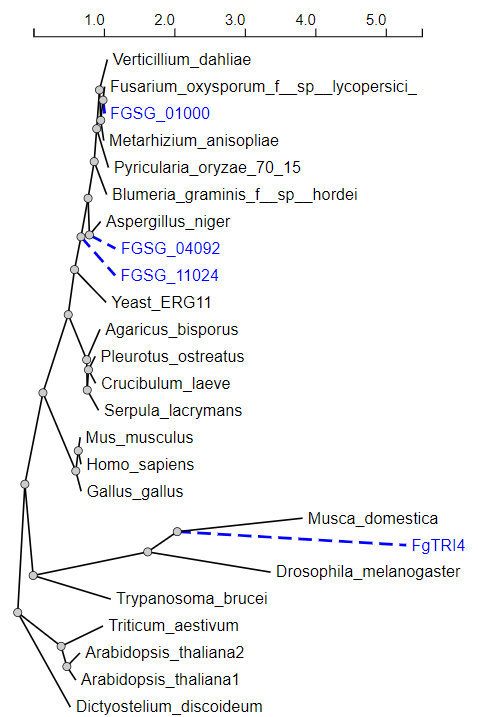

**Deeploc predictions.**

**Summary of 24 predicted sequences**

Table of predicted subcelullar localizations. Use the help page for more detailed description of the output page.

**Predicted proteins**

**Fusarium**

**Prediction: Endoplasmic reticulum, Membrane**

| **Localization** | **Endoplasmic reticulum** | | **Golgi apparatus** | | **Lysosome/Vacuole** | | **Mitochondrion** | **Cell membrane** | **Plastid** | **Peroxisome** | **Nucleus** | **Cytoplasm** | **Extracellular** |
| --- | --- | --- | --- | --- | --- | --- | --- | --- | --- | --- | --- | --- | --- |
| **Likelihood** | 0.9199 | | 0.0364 | | 0.0206 | | 0.0105 | 0.007 | 0.0053 | 0.0002 | 0.0001 | 0 | 0 |
| **Type** | | **Soluble** | | **Membrane** | |  |  |  |  |  |  |  |  |
| **Likelihood** | | 0.0001 | | 0.9999 | |  |  |  |  |  |  |  |  |

**Hierarchical Tree. Donwload:** [PNG](http://www.cbs.dtu.dk/services/DeepLoc-1.0/tmp/5D01F176000064A5AFD60F1E/tree_7.png) **/** [EPS](http://www.cbs.dtu.dk/services/DeepLoc-1.0/tmp/5D01F176000064A5AFD60F1E/tree_7.eps)
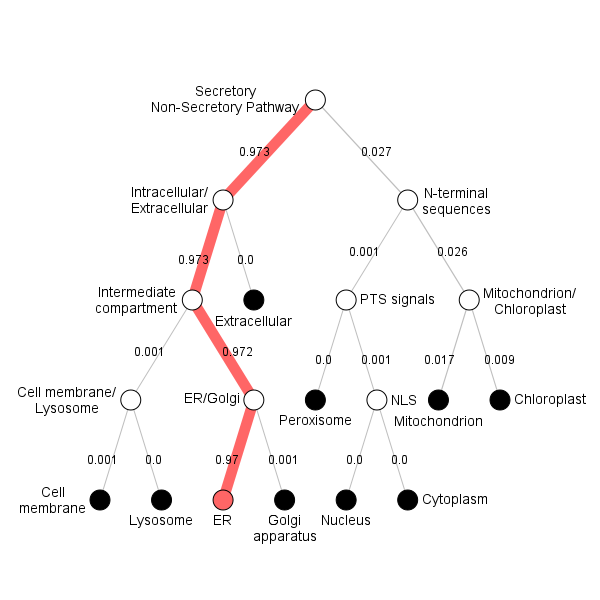

**Position Importance. Donwload:** [PNG](http://www.cbs.dtu.dk/services/DeepLoc-1.0/tmp/5D01F176000064A5AFD60F1E/alpha_7.png) **/** [EPS](http://www.cbs.dtu.dk/services/DeepLoc-1.0/tmp/5D01F176000064A5AFD60F1E/alpha_7.eps) **/** [CSV](http://www.cbs.dtu.dk/services/DeepLoc-1.0/tmp/5D01F176000064A5AFD60F1E/alpha_7.csv)
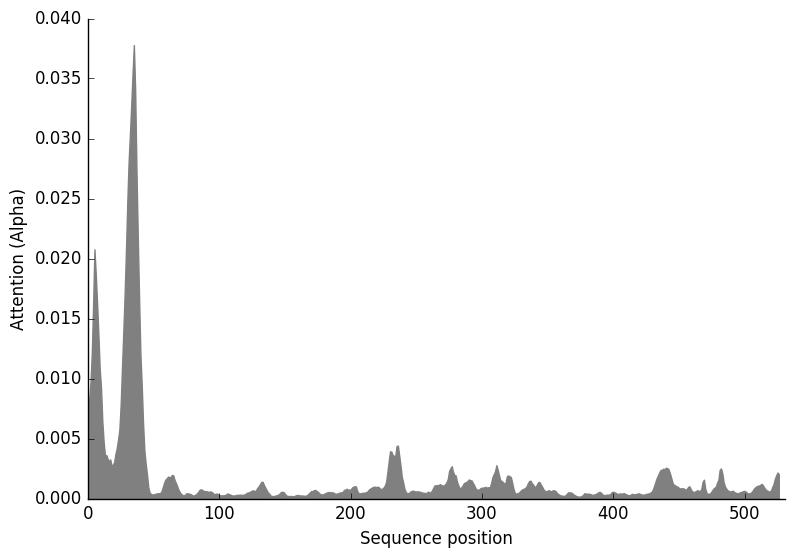

**Blumeria**

**Prediction: Endoplasmic reticulum, Membrane**

| **Localization** | **Endoplasmic reticulum** | | **Golgi apparatus** | | **Mitochondrion** | | **Plastid** | **Cell membrane** | **Lysosome/Vacuole** | **Peroxisome** | **Nucleus** | **Extracellular** | **Cytoplasm** |
| --- | --- | --- | --- | --- | --- | --- | --- | --- | --- | --- | --- | --- | --- |
| **Likelihood** | 0.9962 | | 0.0013 | | 0.0011 | | 0.0006 | 0.0005 | 0.0002 | 0 | 0 | 0 | 0 |
| **Type** | | **Soluble** | | **Membrane** | |  |  |  |  |  |  |  |  |
| **Likelihood** | | 0 | | 1 | |  |  |  |  |  |  |  |  |

**Hierarchical Tree. Donwload:** [PNG](http://www.cbs.dtu.dk/services/DeepLoc-1.0/tmp/5D01F176000064A5AFD60F1E/tree_9.png) **/** [EPS](http://www.cbs.dtu.dk/services/DeepLoc-1.0/tmp/5D01F176000064A5AFD60F1E/tree_9.eps)
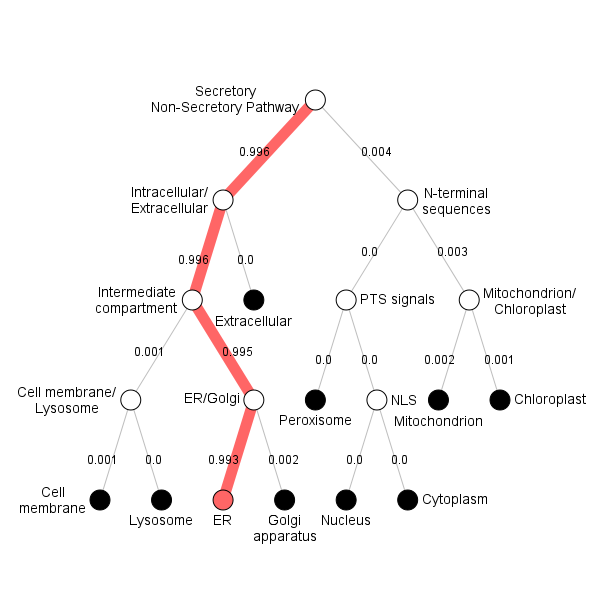

**Position Importance. Donwload:** [PNG](http://www.cbs.dtu.dk/services/DeepLoc-1.0/tmp/5D01F176000064A5AFD60F1E/alpha_9.png) **/** [EPS](http://www.cbs.dtu.dk/services/DeepLoc-1.0/tmp/5D01F176000064A5AFD60F1E/alpha_9.eps) **/** [CSV](http://www.cbs.dtu.dk/services/DeepLoc-1.0/tmp/5D01F176000064A5AFD60F1E/alpha_9.csv)
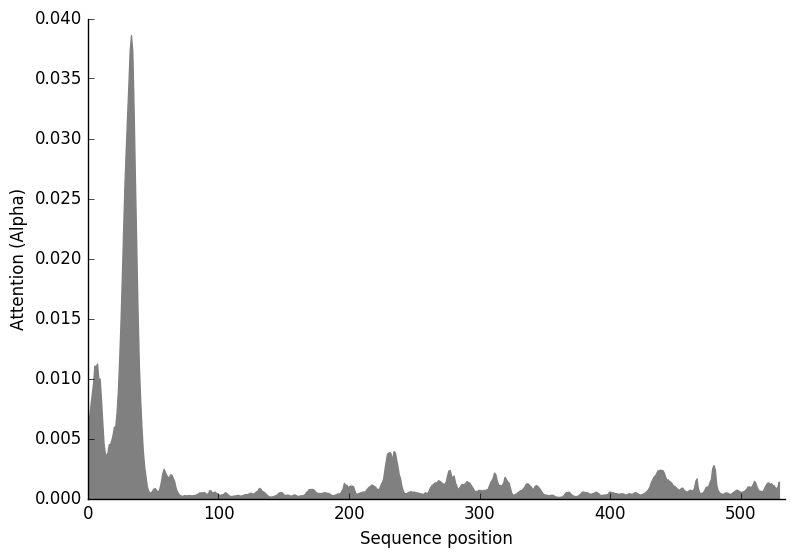

**Aspergillus**

**Prediction: Endoplasmic reticulum, Membrane**

| **Localization** | **Endoplasmic reticulum** | | **Golgi apparatus** | | **Mitochondrion** | | **Lysosome/Vacuole** | **Plastid** | **Cell membrane** | **Peroxisome** | **Nucleus** | **Cytoplasm** | **Extracellular** |
| --- | --- | --- | --- | --- | --- | --- | --- | --- | --- | --- | --- | --- | --- |
| **Likelihood** | 0.9688 | | 0.0189 | | 0.0064 | | 0.0024 | 0.0022 | 0.0012 | 0.0001 | 0 | 0 | 0 |
| **Type** | | **Soluble** | | **Membrane** | |  |  |  |  |  |  |  |  |
| **Likelihood** | | 0.0001 | | 0.9999 | |  |  |  |  |  |  |  |  |

**Hierarchical Tree. Donwload:** [PNG](http://www.cbs.dtu.dk/services/DeepLoc-1.0/tmp/5D01F176000064A5AFD60F1E/tree_8.png) **/** [EPS](http://www.cbs.dtu.dk/services/DeepLoc-1.0/tmp/5D01F176000064A5AFD60F1E/tree_8.eps)
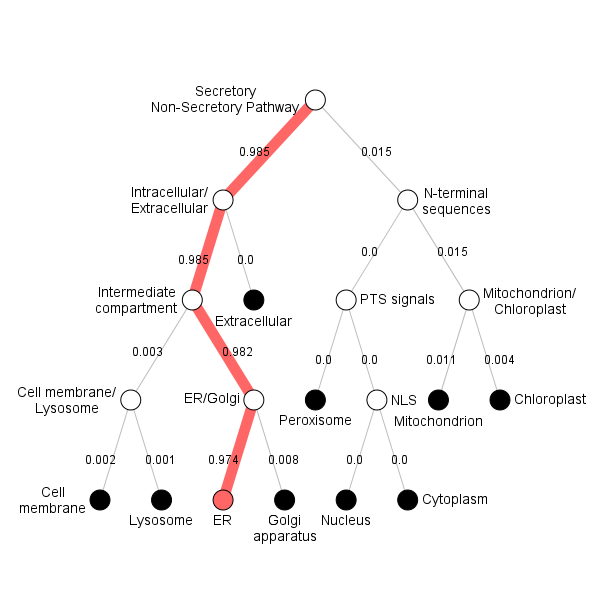

**Position Importance. Donwload:** [PNG](http://www.cbs.dtu.dk/services/DeepLoc-1.0/tmp/5D01F176000064A5AFD60F1E/alpha_8.png) **/** [EPS](http://www.cbs.dtu.dk/services/DeepLoc-1.0/tmp/5D01F176000064A5AFD60F1E/alpha_8.eps) **/** [CSV](http://www.cbs.dtu.dk/services/DeepLoc-1.0/tmp/5D01F176000064A5AFD60F1E/alpha_8.csv)
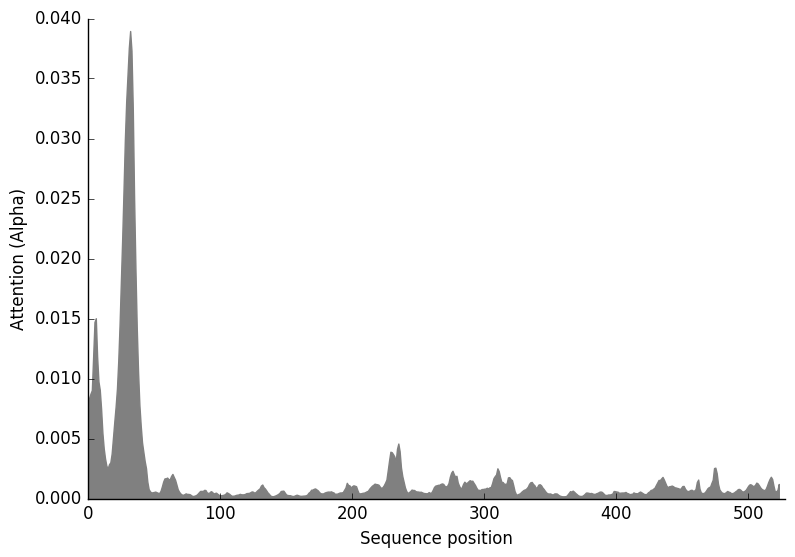

**Crucibulum**

**Prediction: Endoplasmic reticulum, Membrane**

| **Localization** | **Endoplasmic reticulum** | | **Plastid** | | **Mitochondrion** | | **Golgi apparatus** | **Cell membrane** | **Lysosome/Vacuole** | **Nucleus** | **Peroxisome** | **Cytoplasm** | **Extracellular** |
| --- | --- | --- | --- | --- | --- | --- | --- | --- | --- | --- | --- | --- | --- |
| **Likelihood** | 0.7622 | | 0.1273 | | 0.0853 | | 0.0074 | 0.0072 | 0.0071 | 0.0018 | 0.0016 | 0.0001 | 0.0001 |
| **Type** | | **Soluble** | | **Membrane** | |  |  |  |  |  |  |  |  |
| **Likelihood** | | 0.0002 | | 0.9998 | |  |  |  |  |  |  |  |  |

**Hierarchical Tree. Donwload:** [PNG](http://www.cbs.dtu.dk/services/DeepLoc-1.0/tmp/5D01F176000064A5AFD60F1E/tree_10.png) **/** [EPS](http://www.cbs.dtu.dk/services/DeepLoc-1.0/tmp/5D01F176000064A5AFD60F1E/tree_10.eps)
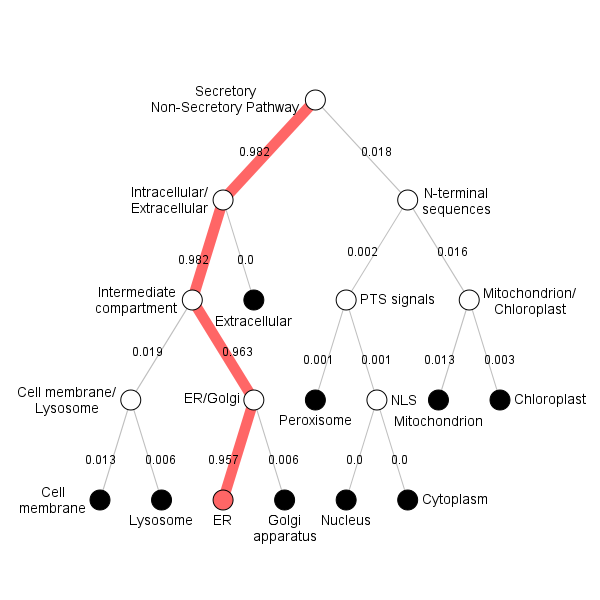

**Position Importance. Donwload:** [PNG](http://www.cbs.dtu.dk/services/DeepLoc-1.0/tmp/5D01F176000064A5AFD60F1E/alpha_10.png) **/** [EPS](http://www.cbs.dtu.dk/services/DeepLoc-1.0/tmp/5D01F176000064A5AFD60F1E/alpha_10.eps) **/** [CSV](http://www.cbs.dtu.dk/services/DeepLoc-1.0/tmp/5D01F176000064A5AFD60F1E/alpha_10.csv)
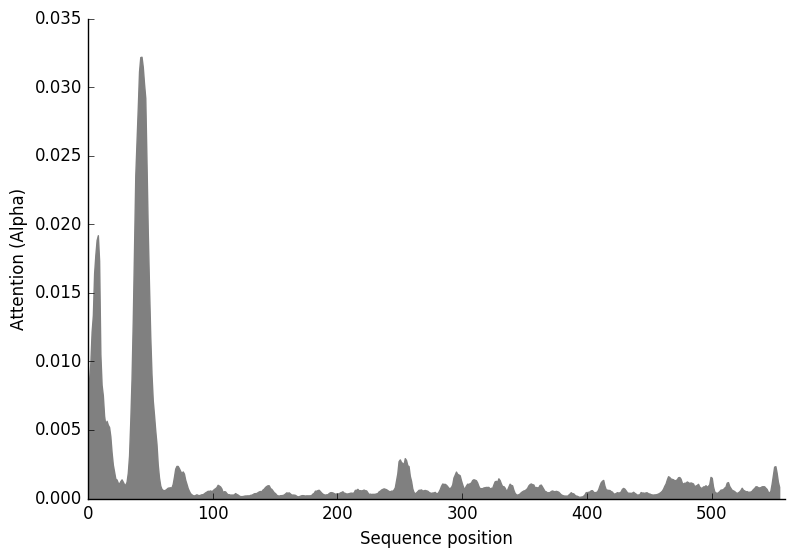

**Homo_sapiens1**

**Prediction: Endoplasmic reticulum, Membrane**

| **Localization** | **Endoplasmic reticulum** | | **Mitochondrion** | | **Plastid** | | **Cell membrane** | **Lysosome/Vacuole** | **Golgi apparatus** | **Peroxisome** | **Extracellular** | **Nucleus** | **Cytoplasm** |
| --- | --- | --- | --- | --- | --- | --- | --- | --- | --- | --- | --- | --- | --- |
| **Likelihood** | 0.613 | | 0.1272 | | 0.1196 | | 0.0979 | 0.0298 | 0.0079 | 0.0033 | 0.0006 | 0.0005 | 0.0004 |
| **Type** | | **Soluble** | | **Membrane** | |  |  |  |  |  |  |  |  |
| **Likelihood** | | 0.0006 | | 0.9994 | |  |  |  |  |  |  |  |  |

**Hierarchical Tree. Donwload:** [PNG](http://www.cbs.dtu.dk/services/DeepLoc-1.0/tmp/5D01F176000064A5AFD60F1E/tree_17.png) **/** [EPS](http://www.cbs.dtu.dk/services/DeepLoc-1.0/tmp/5D01F176000064A5AFD60F1E/tree_17.eps)
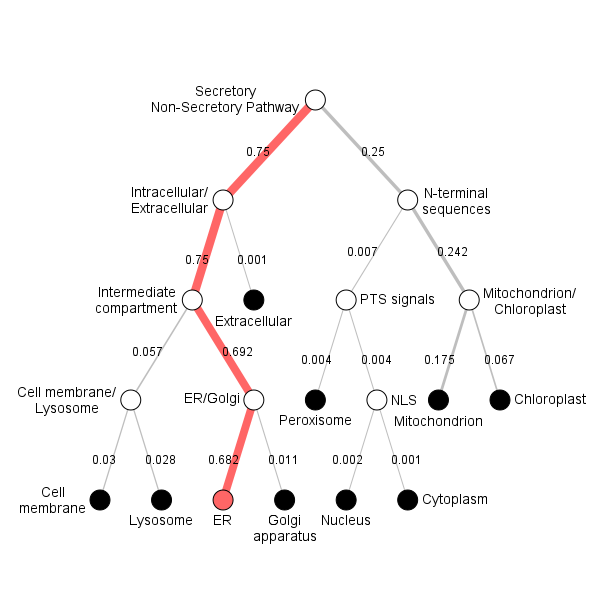

**Position Importance. Donwload:** [PNG](http://www.cbs.dtu.dk/services/DeepLoc-1.0/tmp/5D01F176000064A5AFD60F1E/alpha_17.png) **/** [EPS](http://www.cbs.dtu.dk/services/DeepLoc-1.0/tmp/5D01F176000064A5AFD60F1E/alpha_17.eps) **/** [CSV](http://www.cbs.dtu.dk/services/DeepLoc-1.0/tmp/5D01F176000064A5AFD60F1E/alpha_17.csv)
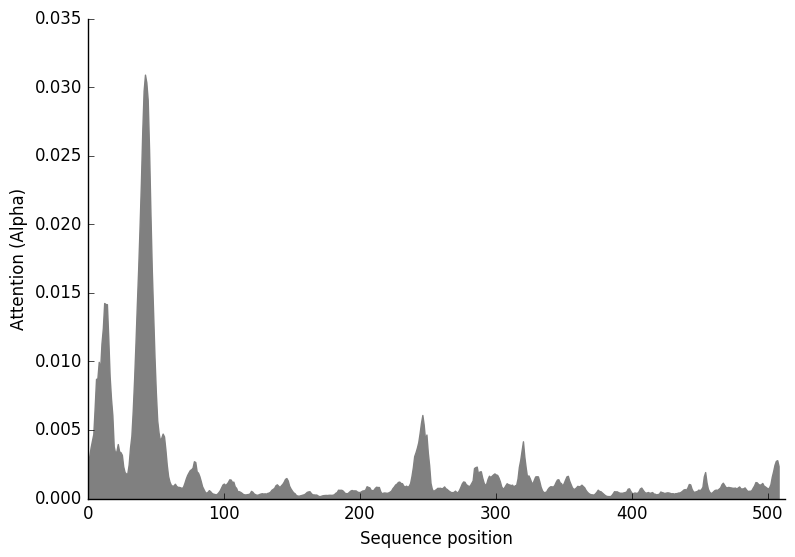

**Homo_sapiens**

**Prediction: Mitochondrion, Soluble**

| **Localization** | **Mitochondrion** | | **Peroxisome** | | **Plastid** | | **Cytoplasm** | **Endoplasmic reticulum** | **Extracellular** | **Nucleus** | **Cell membrane** | **Lysosome/Vacuole** | **Golgi apparatus** |
| --- | --- | --- | --- | --- | --- | --- | --- | --- | --- | --- | --- | --- | --- |
| **Likelihood** | 0.3018 | | 0.2509 | | 0.1678 | | 0.1661 | 0.0588 | 0.0266 | 0.0106 | 0.0103 | 0.0043 | 0.0026 |
| **Type** | | **Soluble** | | **Membrane** | |  |  |  |  |  |  |  |  |
| **Likelihood** | | 0.5626 | | 0.4374 | |  |  |  |  |  |  |  |  |

**Hierarchical Tree. Donwload:** [PNG](http://www.cbs.dtu.dk/services/DeepLoc-1.0/tmp/5D01F176000064A5AFD60F1E/tree_16.png) **/** [EPS](http://www.cbs.dtu.dk/services/DeepLoc-1.0/tmp/5D01F176000064A5AFD60F1E/tree_16.eps)
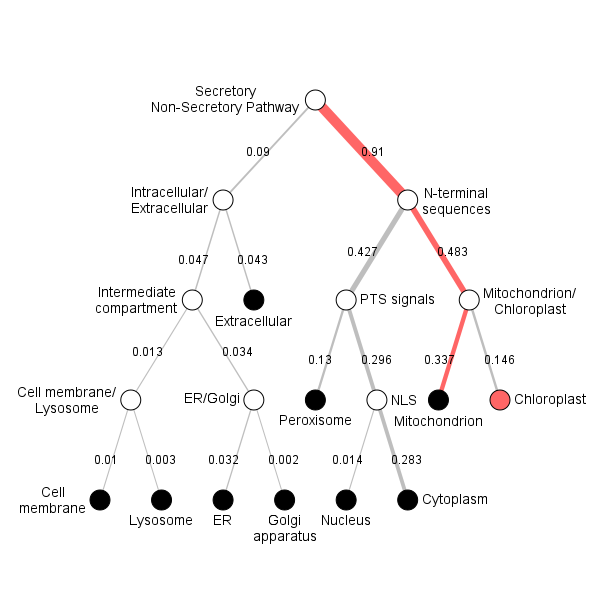

**Position Importance. Donwload:** [PNG](http://www.cbs.dtu.dk/services/DeepLoc-1.0/tmp/5D01F176000064A5AFD60F1E/alpha_16.png) **/** [EPS](http://www.cbs.dtu.dk/services/DeepLoc-1.0/tmp/5D01F176000064A5AFD60F1E/alpha_16.eps) **/** [CSV](http://www.cbs.dtu.dk/services/DeepLoc-1.0/tmp/5D01F176000064A5AFD60F1E/alpha_16.csv)
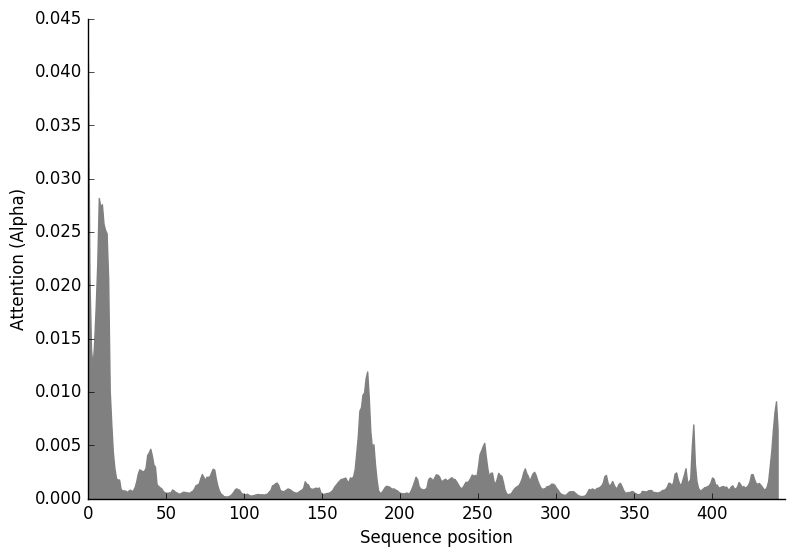

**Pyricularia**

**Prediction: Endoplasmic reticulum, Membrane**

| **Localization** | **Endoplasmic reticulum** | | **Golgi apparatus** | | **Lysosome/Vacuole** | | **Mitochondrion** | **Cell membrane** | **Plastid** | **Peroxisome** | **Nucleus** | **Cytoplasm** | **Extracellular** |
| --- | --- | --- | --- | --- | --- | --- | --- | --- | --- | --- | --- | --- | --- |
| **Likelihood** | 0.9402 | | 0.0352 | | 0.0188 | | 0.0024 | 0.0018 | 0.0015 | 0.0001 | 0 | 0 | 0 |
| **Type** | | **Soluble** | | **Membrane** | |  |  |  |  |  |  |  |  |
| **Likelihood** | | 0.0001 | | 0.9999 | |  |  |  |  |  |  |  |  |

**Hierarchical Tree. Donwload:** [PNG](http://www.cbs.dtu.dk/services/DeepLoc-1.0/tmp/5D01F176000064A5AFD60F1E/tree_4.png) **/** [EPS](http://www.cbs.dtu.dk/services/DeepLoc-1.0/tmp/5D01F176000064A5AFD60F1E/tree_4.eps)
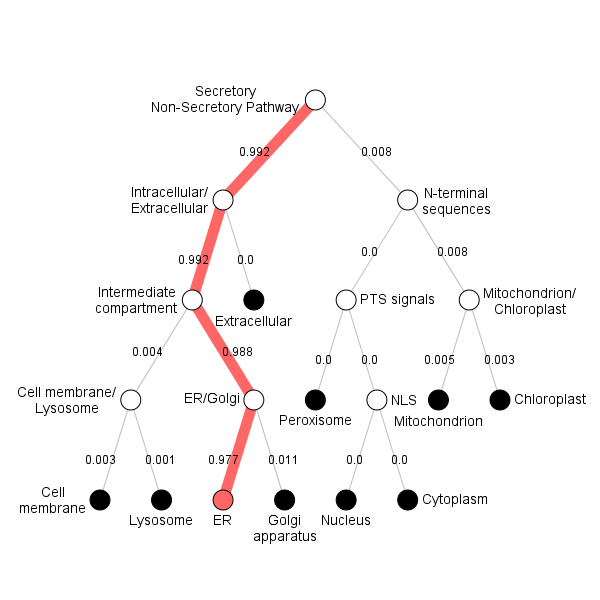

**Position Importance. Donwload:** [PNG](http://www.cbs.dtu.dk/services/DeepLoc-1.0/tmp/5D01F176000064A5AFD60F1E/alpha_4.png) **/** [EPS](http://www.cbs.dtu.dk/services/DeepLoc-1.0/tmp/5D01F176000064A5AFD60F1E/alpha_4.eps) **/** [CSV](http://www.cbs.dtu.dk/services/DeepLoc-1.0/tmp/5D01F176000064A5AFD60F1E/alpha_4.csv)
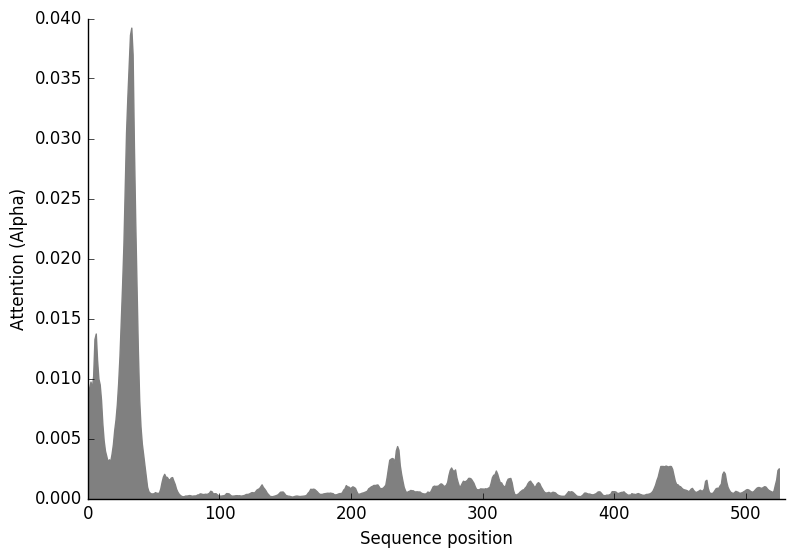

**Pleurotus**

**Prediction: Endoplasmic reticulum, Membrane**

| **Localization** | **Endoplasmic reticulum** | | **Mitochondrion** | | **Cell membrane** | | **Plastid** | **Golgi apparatus** | **Lysosome/Vacuole** | **Peroxisome** | **Cytoplasm** | **Nucleus** | **Extracellular** |
| --- | --- | --- | --- | --- | --- | --- | --- | --- | --- | --- | --- | --- | --- |
| **Likelihood** | 0.9371 | | 0.0587 | | 0.0016 | | 0.0015 | 0.0006 | 0.0003 | 0.0002 | 0.0001 | 0 | 0 |
| **Type** | | **Soluble** | | **Membrane** | |  |  |  |  |  |  |  |  |
| **Likelihood** | | 0.0001 | | 0.9999 | |  |  |  |  |  |  |  |  |

**Hierarchical Tree. Donwload:** [PNG](http://www.cbs.dtu.dk/services/DeepLoc-1.0/tmp/5D01F176000064A5AFD60F1E/tree_13.png) **/** [EPS](http://www.cbs.dtu.dk/services/DeepLoc-1.0/tmp/5D01F176000064A5AFD60F1E/tree_13.eps)
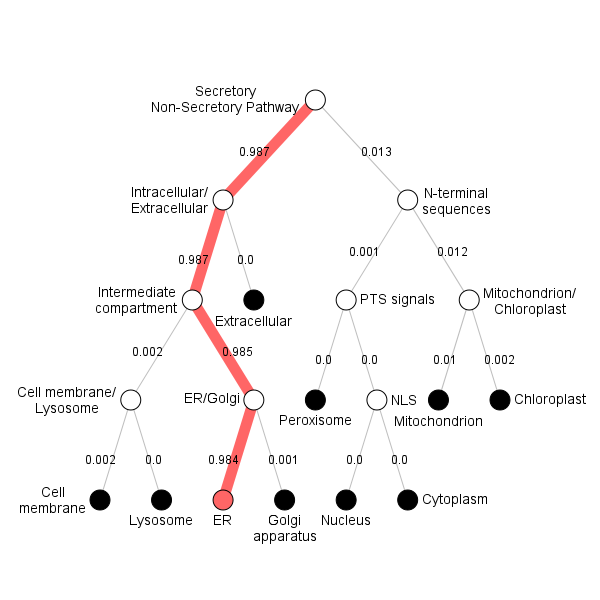

**Position Importance. Donwload:** [PNG](http://www.cbs.dtu.dk/services/DeepLoc-1.0/tmp/5D01F176000064A5AFD60F1E/alpha_13.png) **/** [EPS](http://www.cbs.dtu.dk/services/DeepLoc-1.0/tmp/5D01F176000064A5AFD60F1E/alpha_13.eps) **/** [CSV](http://www.cbs.dtu.dk/services/DeepLoc-1.0/tmp/5D01F176000064A5AFD60F1E/alpha_13.csv)
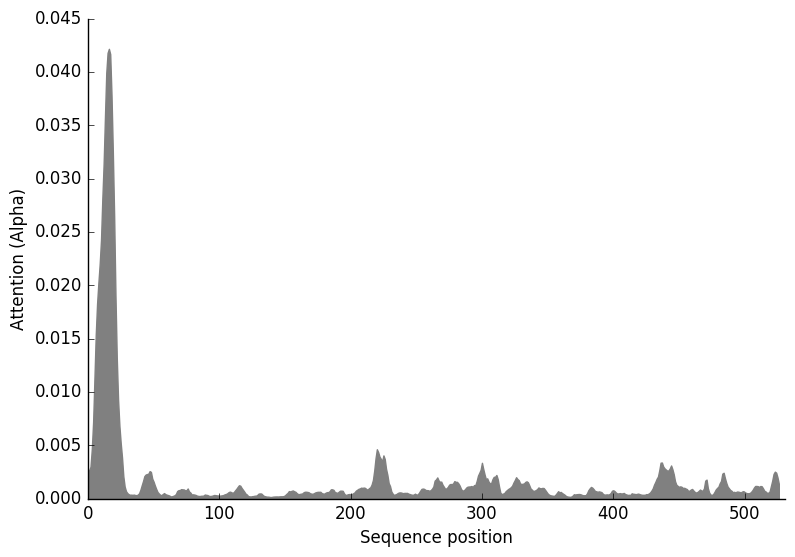

**Gallus**

**Prediction: Endoplasmic reticulum, Membrane**

| **Localization** | **Endoplasmic reticulum** | | **Mitochondrion** | | **Plastid** | | **Cytoplasm** | **Extracellular** | **Peroxisome** | **Cell membrane** | **Nucleus** | **Lysosome/Vacuole** | **Golgi apparatus** |
| --- | --- | --- | --- | --- | --- | --- | --- | --- | --- | --- | --- | --- | --- |
| **Likelihood** | 0.6373 | | 0.1315 | | 0.1241 | | 0.0541 | 0.0321 | 0.0123 | 0.0038 | 0.0021 | 0.002 | 0.0005 |
| **Type** | | **Soluble** | | **Membrane** | |  |  |  |  |  |  |  |  |
| **Likelihood** | | 0.1889 | | 0.8111 | |  |  |  |  |  |  |  |  |

**Hierarchical Tree. Donwload:** [PNG](http://www.cbs.dtu.dk/services/DeepLoc-1.0/tmp/5D01F176000064A5AFD60F1E/tree_15.png) **/** [EPS](http://www.cbs.dtu.dk/services/DeepLoc-1.0/tmp/5D01F176000064A5AFD60F1E/tree_15.eps)
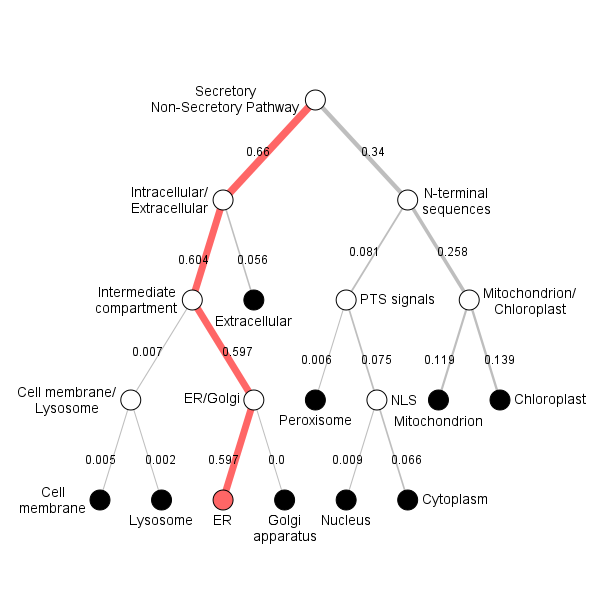

**Position Importance. Donwload:** [PNG](http://www.cbs.dtu.dk/services/DeepLoc-1.0/tmp/5D01F176000064A5AFD60F1E/alpha_15.png) **/** [EPS](http://www.cbs.dtu.dk/services/DeepLoc-1.0/tmp/5D01F176000064A5AFD60F1E/alpha_15.eps) **/** [CSV](http://www.cbs.dtu.dk/services/DeepLoc-1.0/tmp/5D01F176000064A5AFD60F1E/alpha_15.csv)
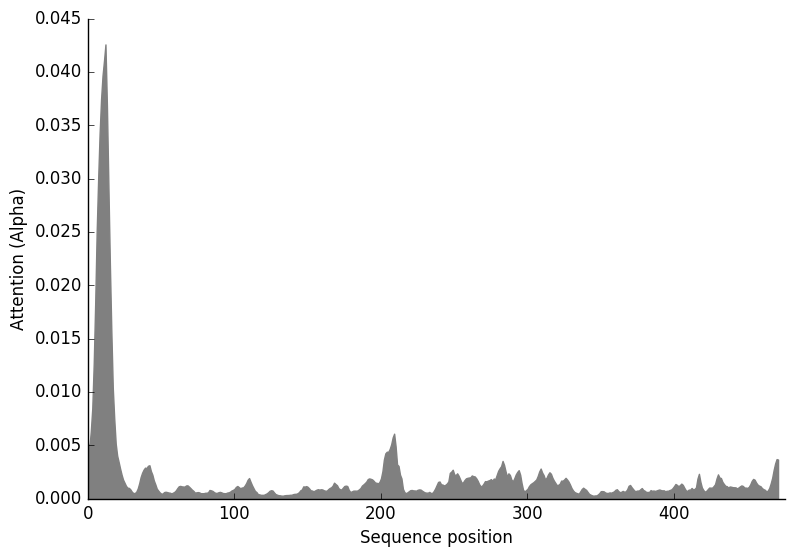

**Serpula**

**Prediction: Endoplasmic reticulum, Membrane**

| **Localization** | **Endoplasmic reticulum** | | **Mitochondrion** | | **Golgi apparatus** | | **Plastid** | **Lysosome/Vacuole** | **Cell membrane** | **Peroxisome** | **Nucleus** | **Extracellular** | **Cytoplasm** |
| --- | --- | --- | --- | --- | --- | --- | --- | --- | --- | --- | --- | --- | --- |
| **Likelihood** | 0.9853 | | 0.0095 | | 0.0022 | | 0.0017 | 0.0007 | 0.0003 | 0.0001 | 0 | 0 | 0 |
| **Type** | | **Soluble** | | **Membrane** | |  |  |  |  |  |  |  |  |
| **Likelihood** | | 0.0003 | | 0.9997 | |  |  |  |  |  |  |  |  |

**Hierarchical Tree. Donwload:** [PNG](http://www.cbs.dtu.dk/services/DeepLoc-1.0/tmp/5D01F176000064A5AFD60F1E/tree_12.png) **/** [EPS](http://www.cbs.dtu.dk/services/DeepLoc-1.0/tmp/5D01F176000064A5AFD60F1E/tree_12.eps)
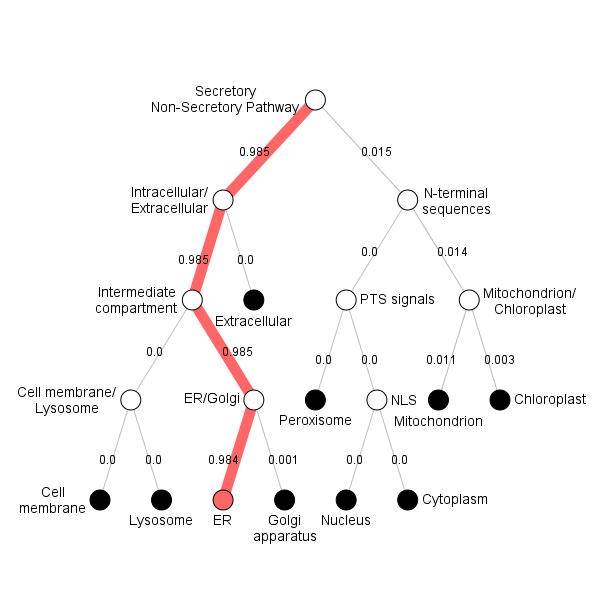

**Position Importance. Donwload:** [PNG](http://www.cbs.dtu.dk/services/DeepLoc-1.0/tmp/5D01F176000064A5AFD60F1E/alpha_12.png) **/** [EPS](http://www.cbs.dtu.dk/services/DeepLoc-1.0/tmp/5D01F176000064A5AFD60F1E/alpha_12.eps) **/** [CSV](http://www.cbs.dtu.dk/services/DeepLoc-1.0/tmp/5D01F176000064A5AFD60F1E/alpha_12.csv)
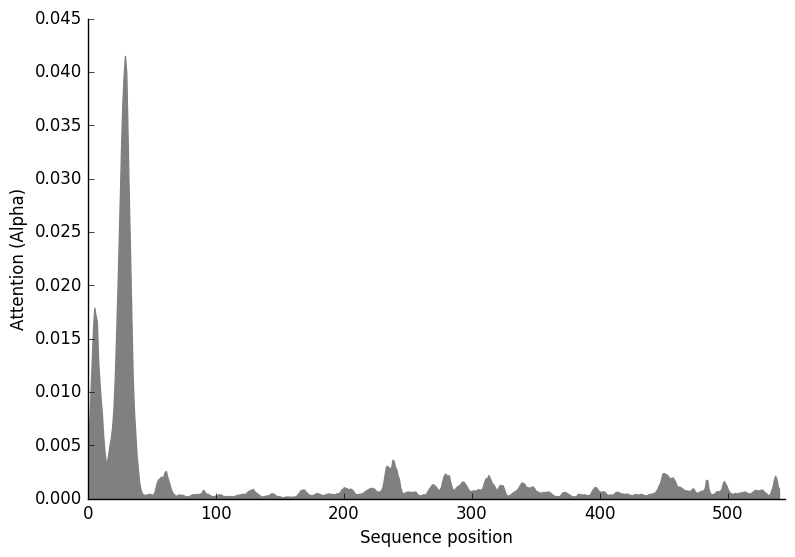

**Arabidopsis_thaliana1**

**Prediction: Endoplasmic reticulum, Membrane**

| **Localization** | **Endoplasmic reticulum** | | **Mitochondrion** | | **Plastid** | | **Cell membrane** | **Golgi apparatus** | **Lysosome/Vacuole** | **Extracellular** | **Peroxisome** | **Cytoplasm** | **Nucleus** |
| --- | --- | --- | --- | --- | --- | --- | --- | --- | --- | --- | --- | --- | --- |
| **Likelihood** | 0.9923 | | 0.0029 | | 0.0023 | | 0.0014 | 0.0007 | 0.0002 | 0.0001 | 0 | 0 | 0 |
| **Type** | | **Soluble** | | **Membrane** | |  |  |  |  |  |  |  |  |
| **Likelihood** | | 0.0004 | | 0.9996 | |  |  |  |  |  |  |  |  |

**Hierarchical Tree. Donwload:** [PNG](http://www.cbs.dtu.dk/services/DeepLoc-1.0/tmp/5D01F176000064A5AFD60F1E/tree_19.png) **/** [EPS](http://www.cbs.dtu.dk/services/DeepLoc-1.0/tmp/5D01F176000064A5AFD60F1E/tree_19.eps)
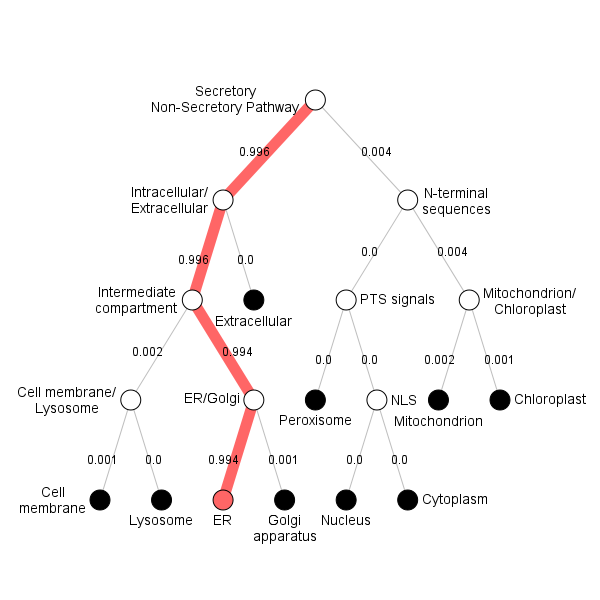

**Position Importance. Donwload:** [PNG](http://www.cbs.dtu.dk/services/DeepLoc-1.0/tmp/5D01F176000064A5AFD60F1E/alpha_19.png) **/** [EPS](http://www.cbs.dtu.dk/services/DeepLoc-1.0/tmp/5D01F176000064A5AFD60F1E/alpha_19.eps) **/** [CSV](http://www.cbs.dtu.dk/services/DeepLoc-1.0/tmp/5D01F176000064A5AFD60F1E/alpha_19.csv)
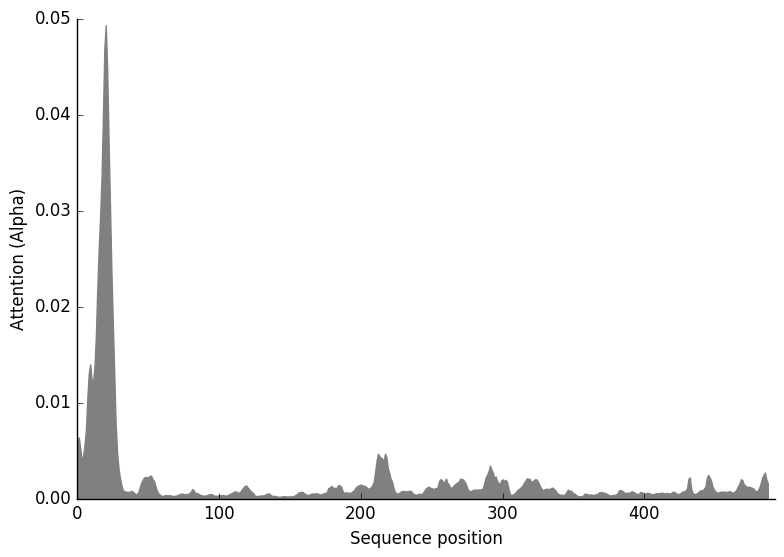

**Metarhizium**

**Prediction: Endoplasmic reticulum, Membrane**

| **Localization** | **Endoplasmic reticulum** | | **Golgi apparatus** | | **Mitochondrion** | | **Lysosome/Vacuole** | **Plastid** | **Cell membrane** | **Peroxisome** | **Nucleus** | **Cytoplasm** | **Extracellular** |
| --- | --- | --- | --- | --- | --- | --- | --- | --- | --- | --- | --- | --- | --- |
| **Likelihood** | 0.9359 | | 0.0273 | | 0.0179 | | 0.0125 | 0.0033 | 0.0029 | 0.0001 | 0.0001 | 0 | 0 |
| **Type** | | **Soluble** | | **Membrane** | |  |  |  |  |  |  |  |  |
| **Likelihood** | | 0.0001 | | 0.9999 | |  |  |  |  |  |  |  |  |

**Hierarchical Tree. Donwload:** [PNG](http://www.cbs.dtu.dk/services/DeepLoc-1.0/tmp/5D01F176000064A5AFD60F1E/tree_5.png) **/** [EPS](http://www.cbs.dtu.dk/services/DeepLoc-1.0/tmp/5D01F176000064A5AFD60F1E/tree_5.eps)
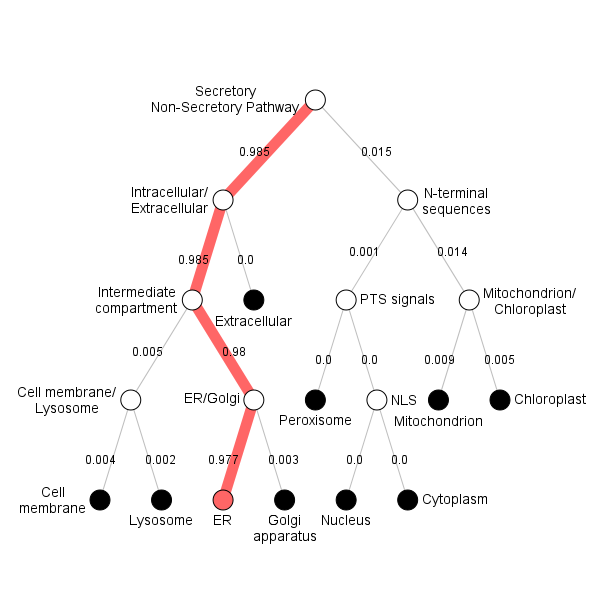

**Position Importance. Donwload:** [PNG](http://www.cbs.dtu.dk/services/DeepLoc-1.0/tmp/5D01F176000064A5AFD60F1E/alpha_5.png) **/** [EPS](http://www.cbs.dtu.dk/services/DeepLoc-1.0/tmp/5D01F176000064A5AFD60F1E/alpha_5.eps) **/** [CSV](http://www.cbs.dtu.dk/services/DeepLoc-1.0/tmp/5D01F176000064A5AFD60F1E/alpha_5.csv)
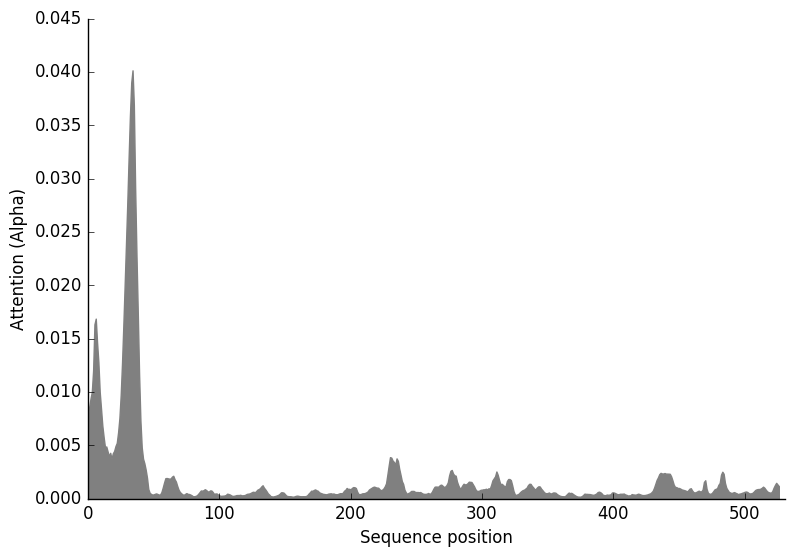

**Triticum_aestivum1**

**Prediction: Mitochondrion, Soluble**

| **Localization** | **Mitochondrion** | | **Peroxisome** | | **Cytoplasm** | | **Plastid** | **Extracellular** | **Endoplasmic reticulum** | **Nucleus** | **Lysosome/Vacuole** | **Cell membrane** | **Golgi apparatus** |
| --- | --- | --- | --- | --- | --- | --- | --- | --- | --- | --- | --- | --- | --- |
| **Likelihood** | 0.3618 | | 0.2106 | | 0.1854 | | 0.1118 | 0.0561 | 0.0426 | 0.0113 | 0.009 | 0.0081 | 0.0033 |
| **Type** | | **Soluble** | | **Membrane** | |  |  |  |  |  |  |  |  |
| **Likelihood** | | 0.6363 | | 0.3637 | |  |  |  |  |  |  |  |  |

**Hierarchical Tree. Donwload:** [PNG](http://www.cbs.dtu.dk/services/DeepLoc-1.0/tmp/5D01F176000064A5AFD60F1E/tree_21.png) **/** [EPS](http://www.cbs.dtu.dk/services/DeepLoc-1.0/tmp/5D01F176000064A5AFD60F1E/tree_21.eps)
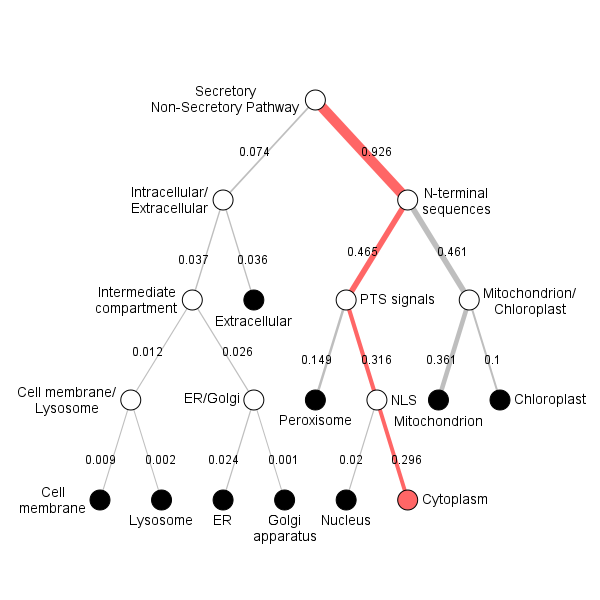

**Position Importance. Donwload:** [PNG](http://www.cbs.dtu.dk/services/DeepLoc-1.0/tmp/5D01F176000064A5AFD60F1E/alpha_21.png) **/** [EPS](http://www.cbs.dtu.dk/services/DeepLoc-1.0/tmp/5D01F176000064A5AFD60F1E/alpha_21.eps) **/** [CSV](http://www.cbs.dtu.dk/services/DeepLoc-1.0/tmp/5D01F176000064A5AFD60F1E/alpha_21.csv)
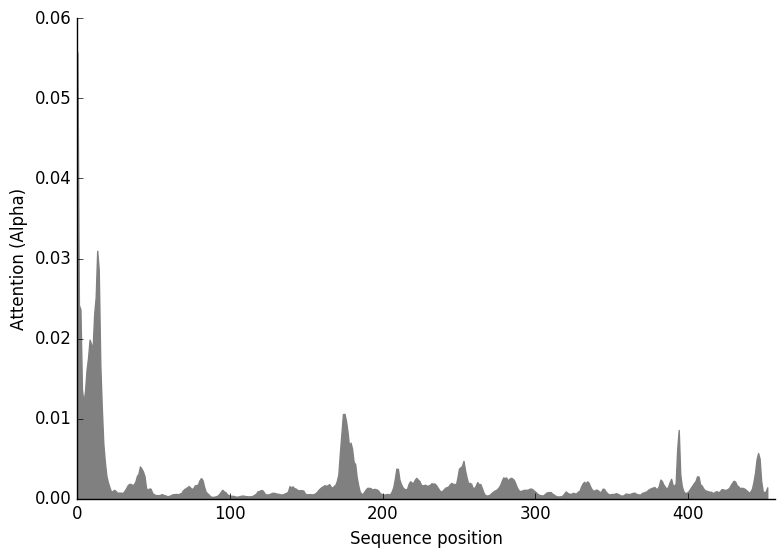

**Triticum_aestivum2**

**Prediction: Endoplasmic reticulum, Membrane**

| **Localization** | **Endoplasmic reticulum** | | **Mitochondrion** | | **Cell membrane** | | **Lysosome/Vacuole** | **Plastid** | **Golgi apparatus** | **Peroxisome** | **Cytoplasm** | **Nucleus** | **Extracellular** |
| --- | --- | --- | --- | --- | --- | --- | --- | --- | --- | --- | --- | --- | --- |
| **Likelihood** | 0.8845 | | 0.0695 | | 0.0256 | | 0.0101 | 0.0059 | 0.0037 | 0.0004 | 0.0001 | 0.0001 | 0.0001 |
| **Type** | | **Soluble** | | **Membrane** | |  |  |  |  |  |  |  |  |
| **Likelihood** | | 0.0002 | | 0.9998 | |  |  |  |  |  |  |  |  |

**Hierarchical Tree. Donwload:** [PNG](http://www.cbs.dtu.dk/services/DeepLoc-1.0/tmp/5D01F176000064A5AFD60F1E/tree_22.png) **/** [EPS](http://www.cbs.dtu.dk/services/DeepLoc-1.0/tmp/5D01F176000064A5AFD60F1E/tree_22.eps)
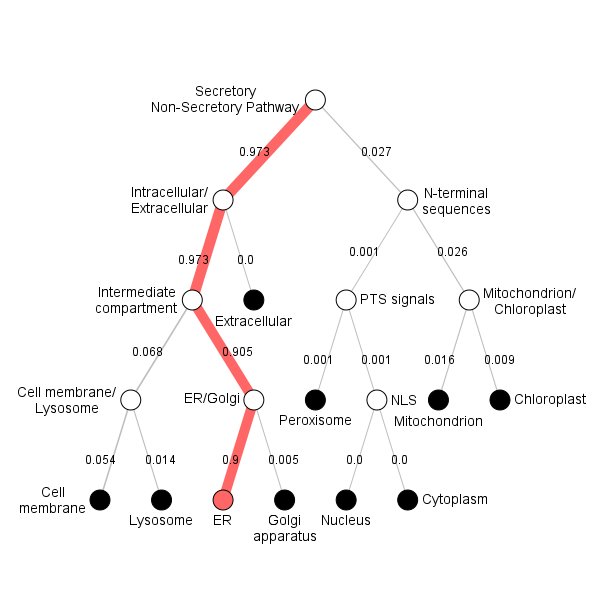

**Position Importance. Donwload:** [PNG](http://www.cbs.dtu.dk/services/DeepLoc-1.0/tmp/5D01F176000064A5AFD60F1E/alpha_22.png) **/** [EPS](http://www.cbs.dtu.dk/services/DeepLoc-1.0/tmp/5D01F176000064A5AFD60F1E/alpha_22.eps) **/** [CSV](http://www.cbs.dtu.dk/services/DeepLoc-1.0/tmp/5D01F176000064A5AFD60F1E/alpha_22.csv)
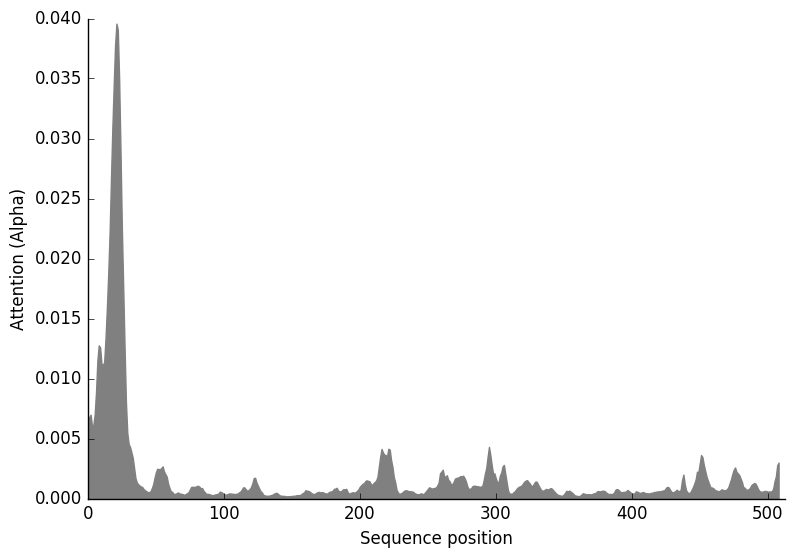

**Verticillium**

**Prediction: Endoplasmic reticulum, Membrane**

| **Localization** | **Endoplasmic reticulum** | | **Golgi apparatus** | | **Lysosome/Vacuole** | | **Mitochondrion** | **Plastid** | **Cell membrane** | **Peroxisome** | **Nucleus** | **Cytoplasm** | **Extracellular** |
| --- | --- | --- | --- | --- | --- | --- | --- | --- | --- | --- | --- | --- | --- |
| **Likelihood** | 0.9245 | | 0.042 | | 0.0212 | | 0.0076 | 0.0031 | 0.0014 | 0.0002 | 0.0001 | 0 | 0 |
| **Type** | | **Soluble** | | **Membrane** | |  |  |  |  |  |  |  |  |
| **Likelihood** | | 0.0001 | | 0.9999 | |  |  |  |  |  |  |  |  |

**Hierarchical Tree. Donwload:** [PNG](http://www.cbs.dtu.dk/services/DeepLoc-1.0/tmp/5D01F176000064A5AFD60F1E/tree_6.png) **/** [EPS](http://www.cbs.dtu.dk/services/DeepLoc-1.0/tmp/5D01F176000064A5AFD60F1E/tree_6.eps)
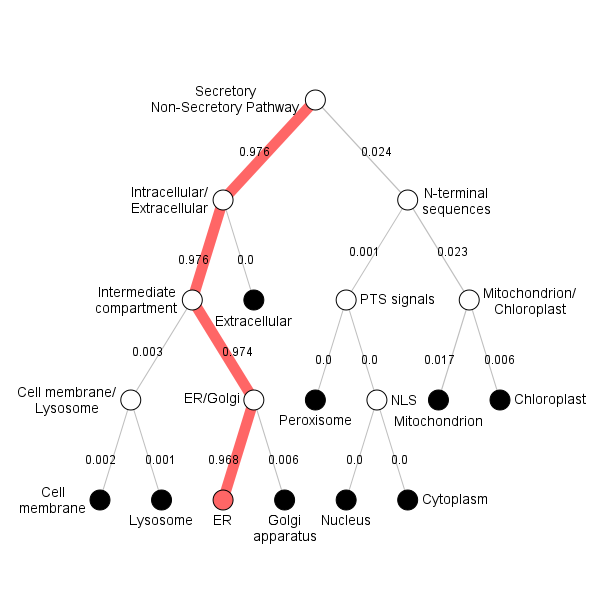

**Position Importance. Donwload:** [PNG](http://www.cbs.dtu.dk/services/DeepLoc-1.0/tmp/5D01F176000064A5AFD60F1E/alpha_6.png) **/** [EPS](http://www.cbs.dtu.dk/services/DeepLoc-1.0/tmp/5D01F176000064A5AFD60F1E/alpha_6.eps) **/** [CSV](http://www.cbs.dtu.dk/services/DeepLoc-1.0/tmp/5D01F176000064A5AFD60F1E/alpha_6.csv)
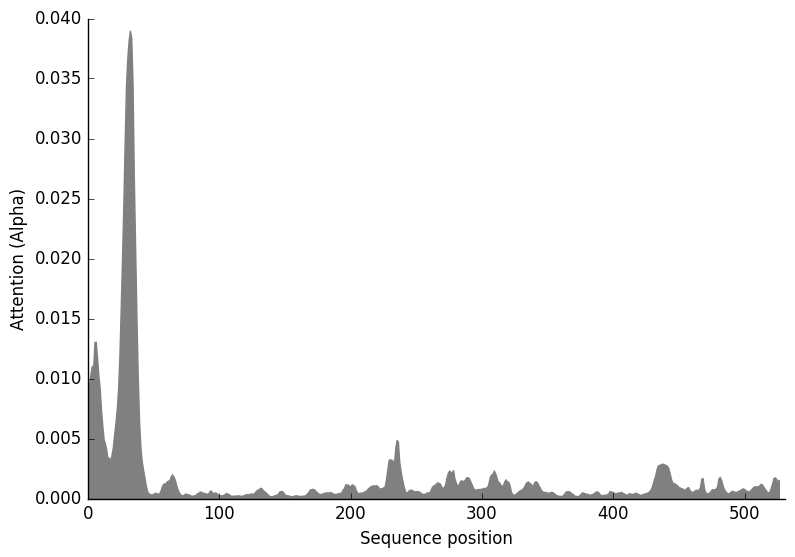

**Arabidopsis_thaliana2**

**Prediction: Endoplasmic reticulum, Membrane**

| **Localization** | **Endoplasmic reticulum** | | **Plastid** | | **Mitochondrion** | | **Cell membrane** | **Golgi apparatus** | **Lysosome/Vacuole** | **Peroxisome** | **Cytoplasm** | **Nucleus** | **Extracellular** |
| --- | --- | --- | --- | --- | --- | --- | --- | --- | --- | --- | --- | --- | --- |
| **Likelihood** | 0.9877 | | 0.0046 | | 0.003 | | 0.0024 | 0.0015 | 0.0007 | 0.0001 | 0.0001 | 0 | 0 |
| **Type** | | **Soluble** | | **Membrane** | |  |  |  |  |  |  |  |  |
| **Likelihood** | | 0.0002 | | 0.9998 | |  |  |  |  |  |  |  |  |

**Hierarchical Tree. Donwload:** [PNG](http://www.cbs.dtu.dk/services/DeepLoc-1.0/tmp/5D01F176000064A5AFD60F1E/tree_20.png) **/** [EPS](http://www.cbs.dtu.dk/services/DeepLoc-1.0/tmp/5D01F176000064A5AFD60F1E/tree_20.eps)
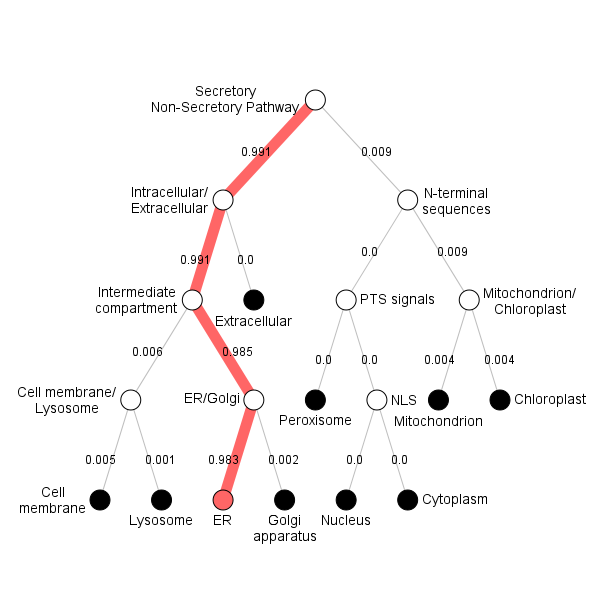

**Position Importance. Donwload:** [PNG](http://www.cbs.dtu.dk/services/DeepLoc-1.0/tmp/5D01F176000064A5AFD60F1E/alpha_20.png) **/** [EPS](http://www.cbs.dtu.dk/services/DeepLoc-1.0/tmp/5D01F176000064A5AFD60F1E/alpha_20.eps) **/** [CSV](http://www.cbs.dtu.dk/services/DeepLoc-1.0/tmp/5D01F176000064A5AFD60F1E/alpha_20.csv)
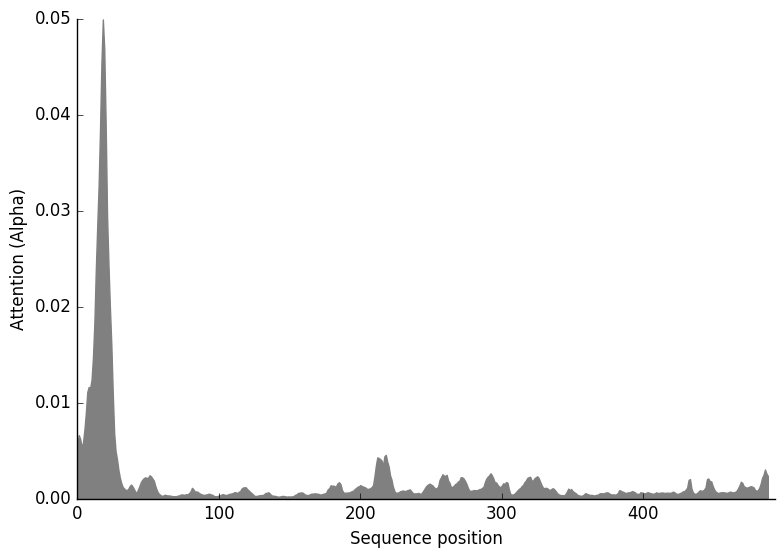

**Trypanosoma**

**Prediction: Endoplasmic reticulum, Membrane**

| **Localization** | **Endoplasmic reticulum** | | **Mitochondrion** | | **Plastid** | | **Cytoplasm** | **Cell membrane** | **Extracellular** | **Golgi apparatus** | **Peroxisome** | **Lysosome/Vacuole** | **Nucleus** |
| --- | --- | --- | --- | --- | --- | --- | --- | --- | --- | --- | --- | --- | --- |
| **Likelihood** | 0.9319 | | 0.0278 | | 0.0163 | | 0.0056 | 0.0053 | 0.0043 | 0.0027 | 0.0023 | 0.0021 | 0.0017 |
| **Type** | | **Soluble** | | **Membrane** | |  |  |  |  |  |  |  |  |
| **Likelihood** | | 0.0384 | | 0.9616 | |  |  |  |  |  |  |  |  |

**Hierarchical Tree. Donwload:** [PNG](http://www.cbs.dtu.dk/services/DeepLoc-1.0/tmp/5D01F176000064A5AFD60F1E/tree_23.png) **/** [EPS](http://www.cbs.dtu.dk/services/DeepLoc-1.0/tmp/5D01F176000064A5AFD60F1E/tree_23.eps)
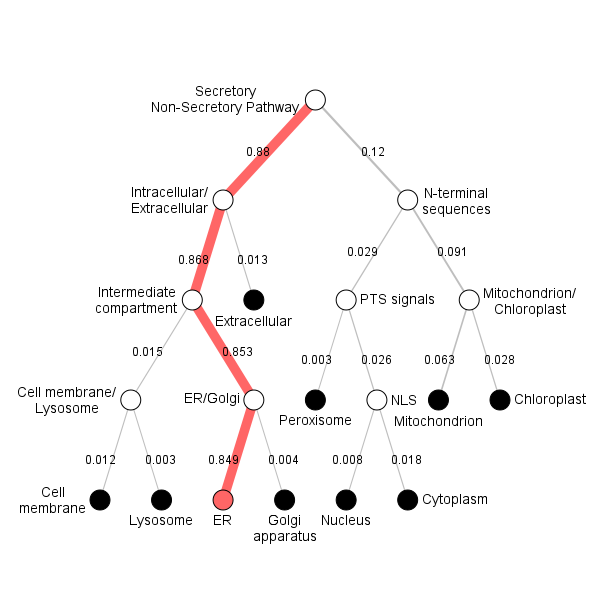

**Position Importance. Donwload:** [PNG](http://www.cbs.dtu.dk/services/DeepLoc-1.0/tmp/5D01F176000064A5AFD60F1E/alpha_23.png) **/** [EPS](http://www.cbs.dtu.dk/services/DeepLoc-1.0/tmp/5D01F176000064A5AFD60F1E/alpha_23.eps) **/** [CSV](http://www.cbs.dtu.dk/services/DeepLoc-1.0/tmp/5D01F176000064A5AFD60F1E/alpha_23.csv)
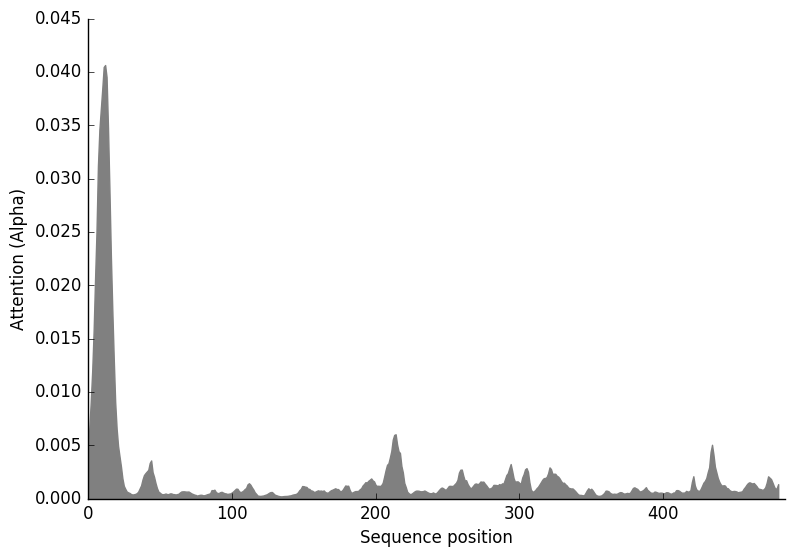

**Musca**

**Prediction: Endoplasmic reticulum, Membrane**

| **Localization** | **Endoplasmic reticulum** | | **Mitochondrion** | | **Golgi apparatus** | | **Cell membrane** | **Lysosome/Vacuole** | **Peroxisome** | **Plastid** | **Cytoplasm** | **Nucleus** | **Extracellular** |
| --- | --- | --- | --- | --- | --- | --- | --- | --- | --- | --- | --- | --- | --- |
| **Likelihood** | 0.9985 | | 0.0007 | | 0.0004 | | 0.0003 | 0.0001 | 0 | 0 | 0 | 0 | 0 |
| **Type** | | **Soluble** | | **Membrane** | |  |  |  |  |  |  |  |  |
| **Likelihood** | | 0 | | 1 | |  |  |  |  |  |  |  |  |

**Hierarchical Tree. Donwload:** [PNG](http://www.cbs.dtu.dk/services/DeepLoc-1.0/tmp/5D01F176000064A5AFD60F1E/tree_18.png) **/** [EPS](http://www.cbs.dtu.dk/services/DeepLoc-1.0/tmp/5D01F176000064A5AFD60F1E/tree_18.eps)
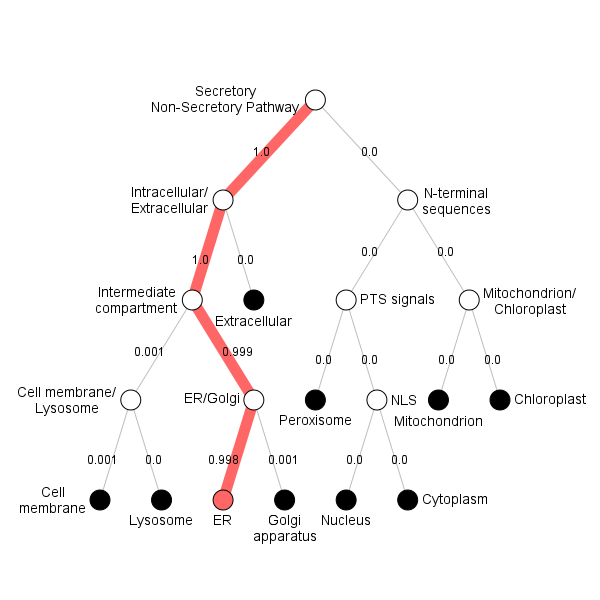

**Position Importance. Donwload:** [PNG](http://www.cbs.dtu.dk/services/DeepLoc-1.0/tmp/5D01F176000064A5AFD60F1E/alpha_18.png) **/** [EPS](http://www.cbs.dtu.dk/services/DeepLoc-1.0/tmp/5D01F176000064A5AFD60F1E/alpha_18.eps) **/** [CSV](http://www.cbs.dtu.dk/services/DeepLoc-1.0/tmp/5D01F176000064A5AFD60F1E/alpha_18.csv)
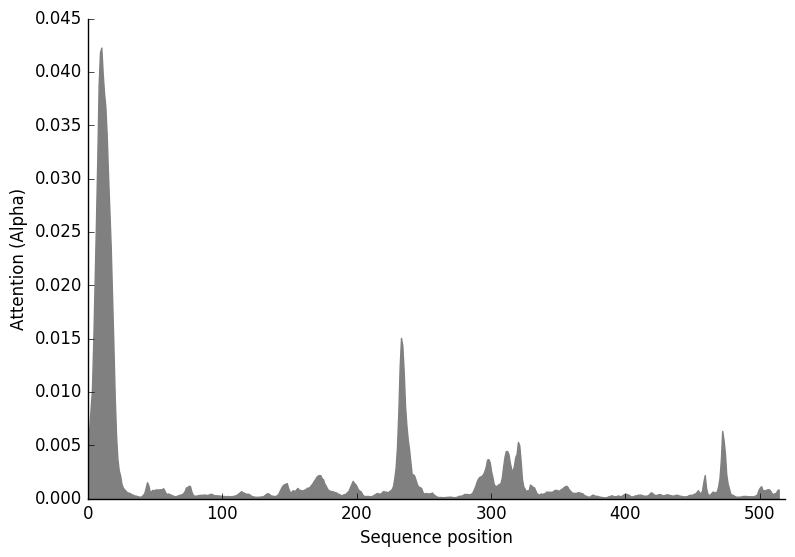

**Drosophila**

**Prediction: Endoplasmic reticulum, Membrane**

| **Localization** | **Endoplasmic reticulum** | | **Mitochondrion** | | **Cell membrane** | | **Golgi apparatus** | **Plastid** | **Lysosome/Vacuole** | **Cytoplasm** | **Peroxisome** | **Extracellular** | **Nucleus** |
| --- | --- | --- | --- | --- | --- | --- | --- | --- | --- | --- | --- | --- | --- |
| **Likelihood** | 0.9997 | | 0.0001 | | 0 | | 0 | 0 | 0 | 0 | 0 | 0 | 0 |
| **Type** | | **Soluble** | | **Membrane** | |  |  |  |  |  |  |  |  |
| **Likelihood** | | 0.0003 | | 0.9997 | |  |  |  |  |  |  |  |  |

**Hierarchical Tree. Donwload:** [PNG](http://www.cbs.dtu.dk/services/DeepLoc-1.0/tmp/5D01F176000064A5AFD60F1E/tree_14.png) **/** [EPS](http://www.cbs.dtu.dk/services/DeepLoc-1.0/tmp/5D01F176000064A5AFD60F1E/tree_14.eps)
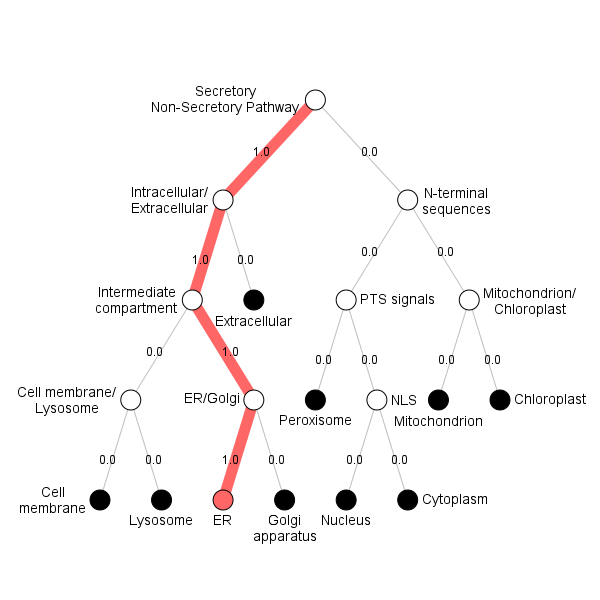

**Position Importance. Donwload:** [PNG](http://www.cbs.dtu.dk/services/DeepLoc-1.0/tmp/5D01F176000064A5AFD60F1E/alpha_14.png) **/** [EPS](http://www.cbs.dtu.dk/services/DeepLoc-1.0/tmp/5D01F176000064A5AFD60F1E/alpha_14.eps) **/** [CSV](http://www.cbs.dtu.dk/services/DeepLoc-1.0/tmp/5D01F176000064A5AFD60F1E/alpha_14.csv)
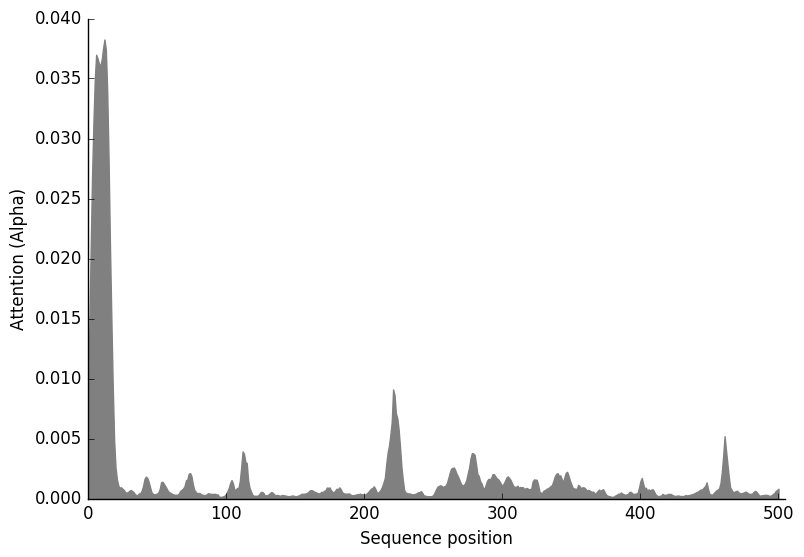

**FGPh1.3**

**Prediction: Endoplasmic reticulum, Membrane**

| **Localization** | **Endoplasmic reticulum** | | **Mitochondrion** | | **Plastid** | | **Cell membrane** | **Golgi apparatus** | **Lysosome/Vacuole** | **Extracellular** | **Peroxisome** | **Cytoplasm** | **Nucleus** |
| --- | --- | --- | --- | --- | --- | --- | --- | --- | --- | --- | --- | --- | --- |
| **Likelihood** | 0.9892 | | 0.0087 | | 0.001 | | 0.0006 | 0.0003 | 0.0001 | 0.0001 | 0 | 0 | 0 |
| **Type** | | **Soluble** | | **Membrane** | |  |  |  |  |  |  |  |  |
| **Likelihood** | | 0.0001 | | 0.9999 | |  |  |  |  |  |  |  |  |

**Hierarchical Tree. Donwload:** [PNG](http://www.cbs.dtu.dk/services/DeepLoc-1.0/tmp/5D01F176000064A5AFD60F1E/tree_3.png) **/** [EPS](http://www.cbs.dtu.dk/services/DeepLoc-1.0/tmp/5D01F176000064A5AFD60F1E/tree_3.eps)
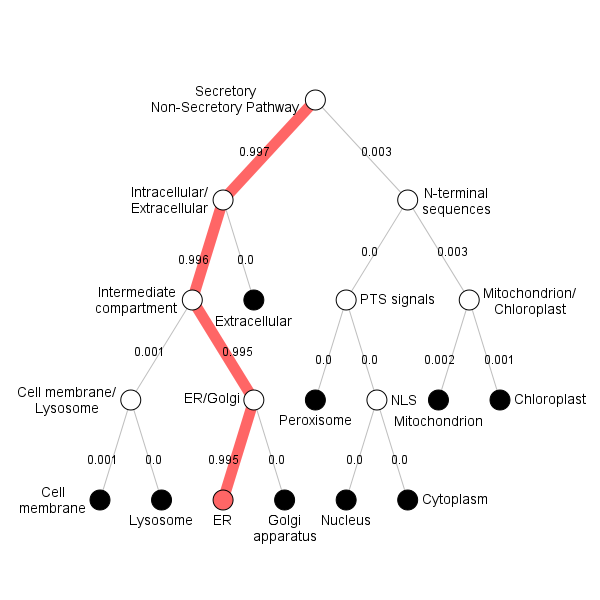

**Position Importance. Donwload:** [PNG](http://www.cbs.dtu.dk/services/DeepLoc-1.0/tmp/5D01F176000064A5AFD60F1E/alpha_3.png) **/** [EPS](http://www.cbs.dtu.dk/services/DeepLoc-1.0/tmp/5D01F176000064A5AFD60F1E/alpha_3.eps) **/** [CSV](http://www.cbs.dtu.dk/services/DeepLoc-1.0/tmp/5D01F176000064A5AFD60F1E/alpha_3.csv)
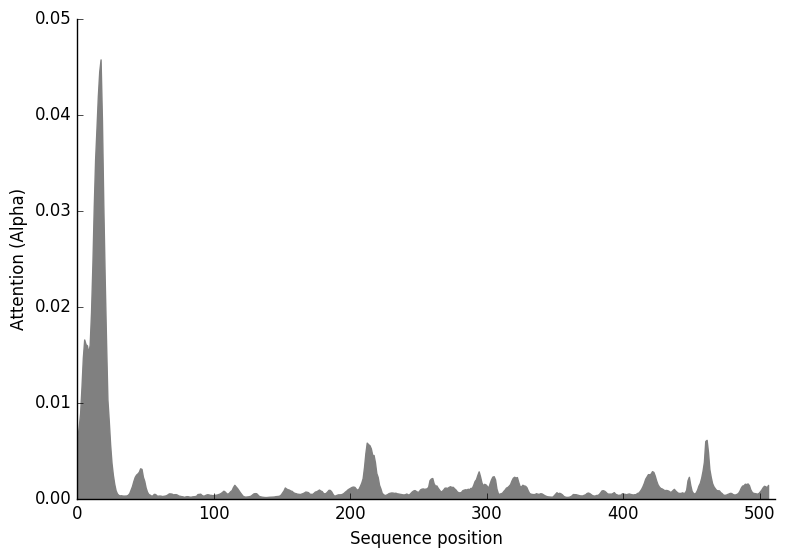

**FGPh1.2**

**Prediction: Endoplasmic reticulum, Membrane**

| **Localization** | **Endoplasmic reticulum** | | **Mitochondrion** | | **Plastid** | | **Golgi apparatus** | **Cell membrane** | **Lysosome/Vacuole** | **Peroxisome** | **Nucleus** | **Cytoplasm** | **Extracellular** |
| --- | --- | --- | --- | --- | --- | --- | --- | --- | --- | --- | --- | --- | --- |
| **Likelihood** | 0.9796 | | 0.0072 | | 0.0057 | | 0.0053 | 0.0016 | 0.0004 | 0.0002 | 0 | 0 | 0 |
| **Type** | | **Soluble** | | **Membrane** | |  |  |  |  |  |  |  |  |
| **Likelihood** | | 0.0001 | | 0.9999 | |  |  |  |  |  |  |  |  |

**Hierarchical Tree. Donwload:** [PNG](http://www.cbs.dtu.dk/services/DeepLoc-1.0/tmp/5D01F176000064A5AFD60F1E/tree_2.png) **/** [EPS](http://www.cbs.dtu.dk/services/DeepLoc-1.0/tmp/5D01F176000064A5AFD60F1E/tree_2.eps)
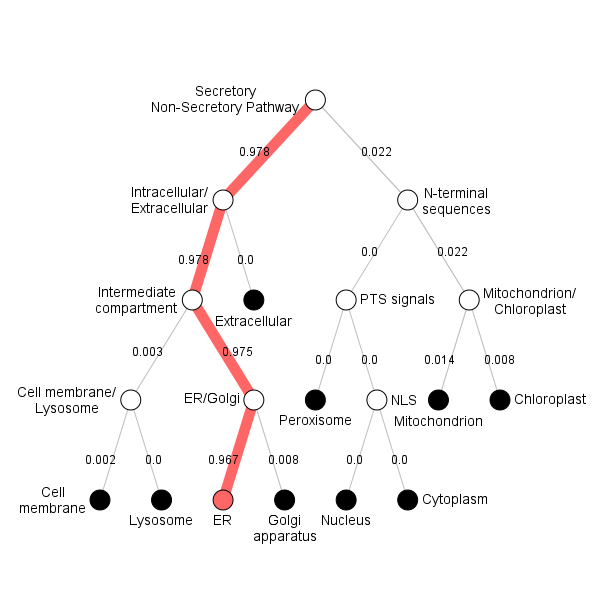

**Position Importance. Donwload:** [PNG](http://www.cbs.dtu.dk/services/DeepLoc-1.0/tmp/5D01F176000064A5AFD60F1E/alpha_2.png) **/** [EPS](http://www.cbs.dtu.dk/services/DeepLoc-1.0/tmp/5D01F176000064A5AFD60F1E/alpha_2.eps) **/** [CSV](http://www.cbs.dtu.dk/services/DeepLoc-1.0/tmp/5D01F176000064A5AFD60F1E/alpha_2.csv)
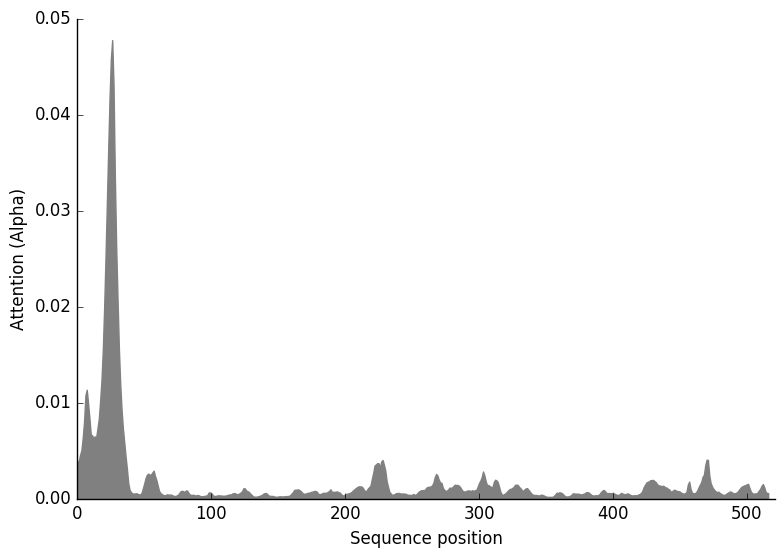

**FGPh1.1**

**Prediction: Endoplasmic reticulum, Membrane**

| **Localization** | **Endoplasmic reticulum** | | **Mitochondrion** | | **Golgi apparatus** | | **Lysosome/Vacuole** | **Plastid** | **Cell membrane** | **Peroxisome** | **Nucleus** | **Cytoplasm** | **Extracellular** |
| --- | --- | --- | --- | --- | --- | --- | --- | --- | --- | --- | --- | --- | --- |
| **Likelihood** | 0.9007 | | 0.0355 | | 0.0317 | | 0.0214 | 0.0072 | 0.0029 | 0.0003 | 0.0003 | 0.0001 | 0 |
| **Type** | | **Soluble** | | **Membrane** | |  |  |  |  |  |  |  |  |
| **Likelihood** | | 0.0001 | | 0.9999 | |  |  |  |  |  |  |  |  |

**Hierarchical Tree. Donwload:** [PNG](http://www.cbs.dtu.dk/services/DeepLoc-1.0/tmp/5D01F176000064A5AFD60F1E/tree_1.png) **/** [EPS](http://www.cbs.dtu.dk/services/DeepLoc-1.0/tmp/5D01F176000064A5AFD60F1E/tree_1.eps)
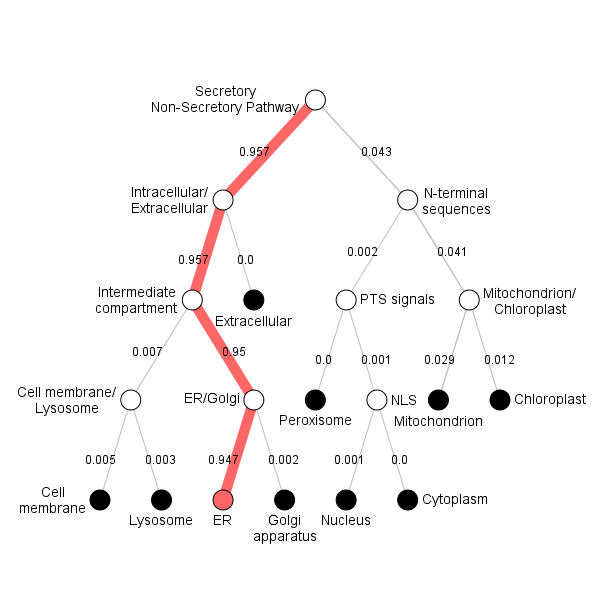

**Position Importance. Donwload:** [PNG](http://www.cbs.dtu.dk/services/DeepLoc-1.0/tmp/5D01F176000064A5AFD60F1E/alpha_1.png) **/** [EPS](http://www.cbs.dtu.dk/services/DeepLoc-1.0/tmp/5D01F176000064A5AFD60F1E/alpha_1.eps) **/** [CSV](http://www.cbs.dtu.dk/services/DeepLoc-1.0/tmp/5D01F176000064A5AFD60F1E/alpha_1.csv)
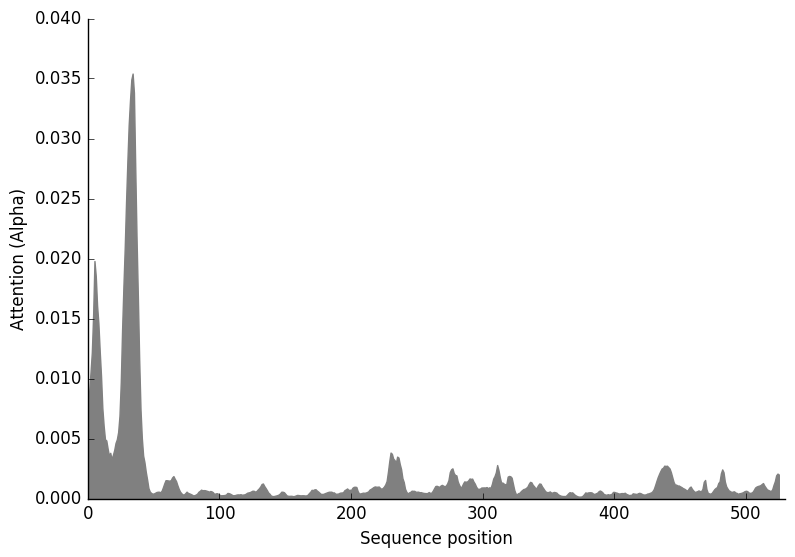

**Dictyostelium**

**Prediction: Endoplasmic reticulum, Membrane**

| **Localization** | **Endoplasmic reticulum** | | **Plastid** | | **Golgi apparatus** | **Cytoplasm** | | **Mitochondrion** | **Extracellular** | **Nucleus** | **Cell membrane** | **Lysosome/Vacuole** | **Peroxisome** |
| --- | --- | --- | --- | --- | --- | --- | --- | --- | --- | --- | --- | --- | --- |
| **Likelihood** | 0.9072 | | 0.0498 | | 0.022 | 0.0116 | | 0.0059 | 0.001 | 0.001 | 0.0009 | 0.0004 | 0.0002 |
| **Type** | | **Soluble** | | **Membrane** | | |  |  |  |  |  |  |  |
| **Likelihood** | | 0.1495 | | 0.8505 | | |  |  |  |  |  |  |  |

**Hierarchical Tree. Donwload:** [PNG](http://www.cbs.dtu.dk/services/DeepLoc-1.0/tmp/5D01F176000064A5AFD60F1E/tree_24.png) **/** [EPS](http://www.cbs.dtu.dk/services/DeepLoc-1.0/tmp/5D01F176000064A5AFD60F1E/tree_24.eps)
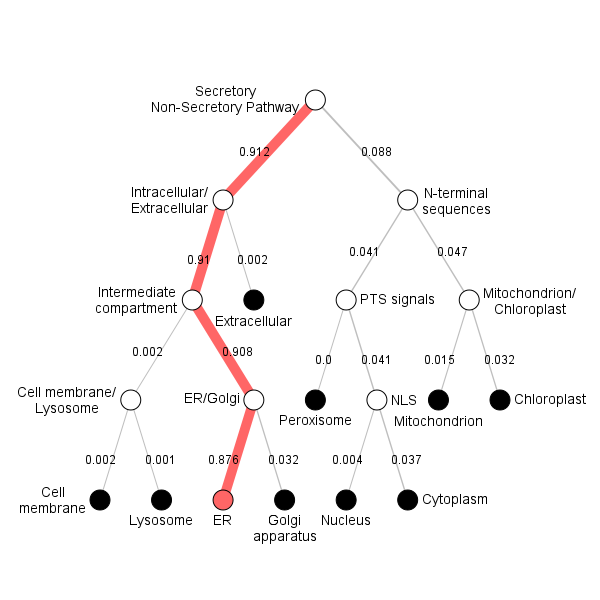

**Position Importance. Donwload:** [PNG](http://www.cbs.dtu.dk/services/DeepLoc-1.0/tmp/5D01F176000064A5AFD60F1E/alpha_24.png) **/** [EPS](http://www.cbs.dtu.dk/services/DeepLoc-1.0/tmp/5D01F176000064A5AFD60F1E/alpha_24.eps) **/** [CSV](http://www.cbs.dtu.dk/services/DeepLoc-1.0/tmp/5D01F176000064A5AFD60F1E/alpha_24.csv)
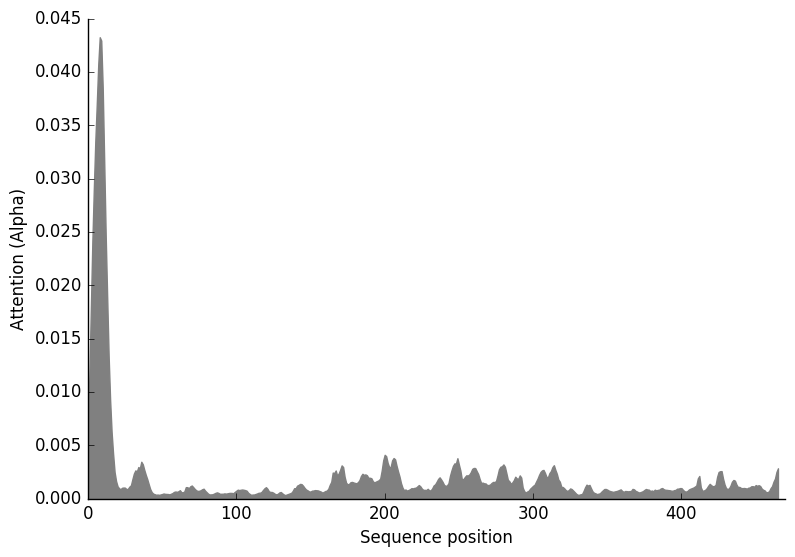

**Agaricus**

**Prediction: Endoplasmic reticulum, Membrane**

| **Localization** | **Endoplasmic reticulum** | | **Cell membrane** | | **Mitochondrion** | | **Plastid** | **Lysosome/Vacuole** | **Golgi apparatus** | **Peroxisome** | **Nucleus** | **Cytoplasm** | **Extracellular** |
| --- | --- | --- | --- | --- | --- | --- | --- | --- | --- | --- | --- | --- | --- |
| **Likelihood** | 0.8645 | | 0.0841 | | 0.02 | | 0.0137 | 0.0128 | 0.0041 | 0.0007 | 0.0001 | 0 | 0 |
| **Type** | | **Soluble** | | **Membrane** | |  |  |  |  |  |  |  |  |
| **Likelihood** | | 0 | | 1 | |  |  |  |  |  |  |  |  |

**Hierarchical Tree. Donwload:** [PNG](http://www.cbs.dtu.dk/services/DeepLoc-1.0/tmp/5D01F176000064A5AFD60F1E/tree_11.png) **/** [EPS](http://www.cbs.dtu.dk/services/DeepLoc-1.0/tmp/5D01F176000064A5AFD60F1E/tree_11.eps)
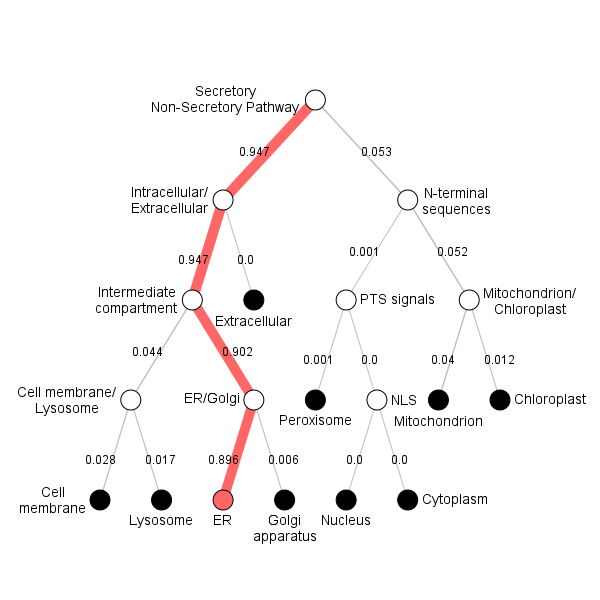

**Position Importance. Donwload:** [PNG](http://www.cbs.dtu.dk/services/DeepLoc-1.0/tmp/5D01F176000064A5AFD60F1E/alpha_11.png) **/** [EPS](http://www.cbs.dtu.dk/services/DeepLoc-1.0/tmp/5D01F176000064A5AFD60F1E/alpha_11.eps) **/** [CSV](http://www.cbs.dtu.dk/services/DeepLoc-1.0/tmp/5D01F176000064A5AFD60F1E/alpha_11.csv)
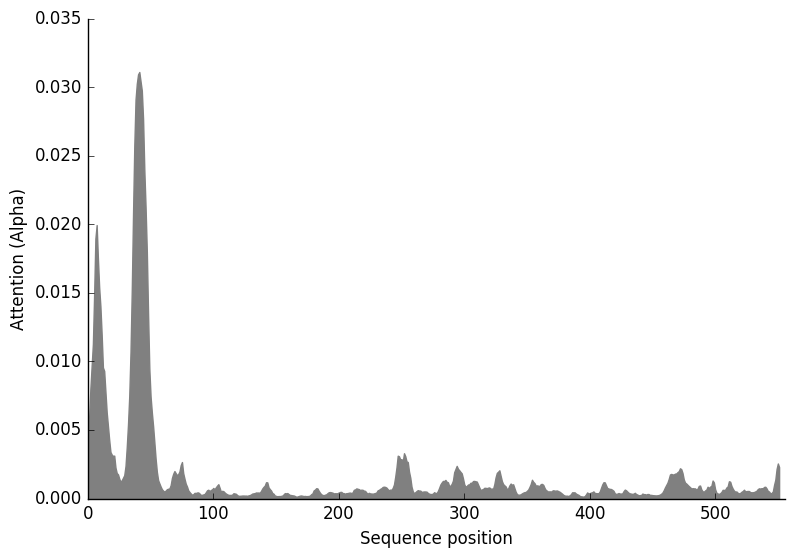

CAM prediction using CaMELS (CalModulin intEraction Learning System) <https://camels.pythonanywhere.com/> and domain structure prediction using HMMER

**Fusarium graminearum FGSG_01000**

Interact pred 1.12 Yes

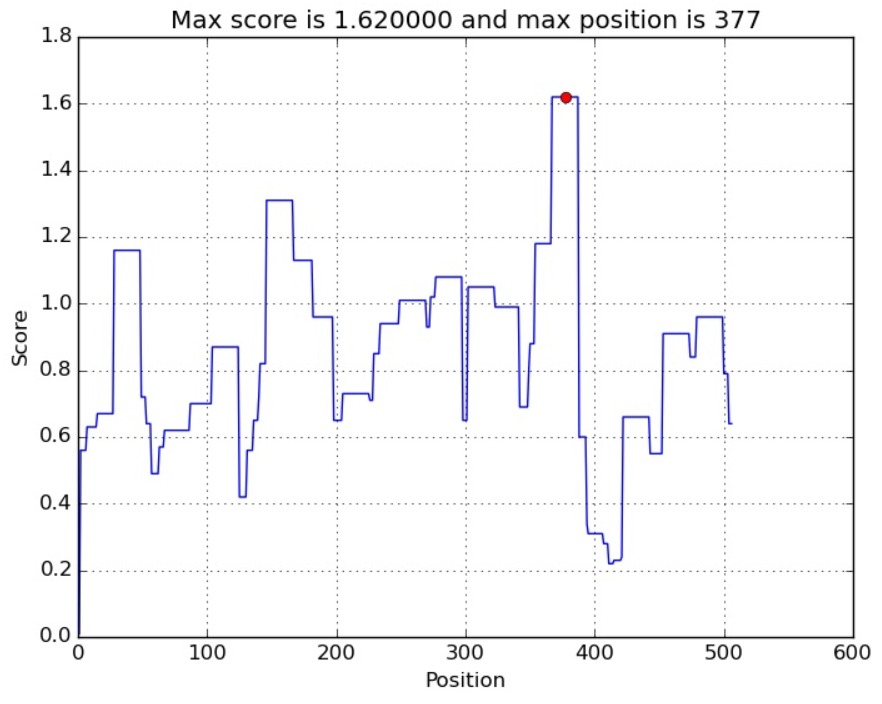

TM25-47

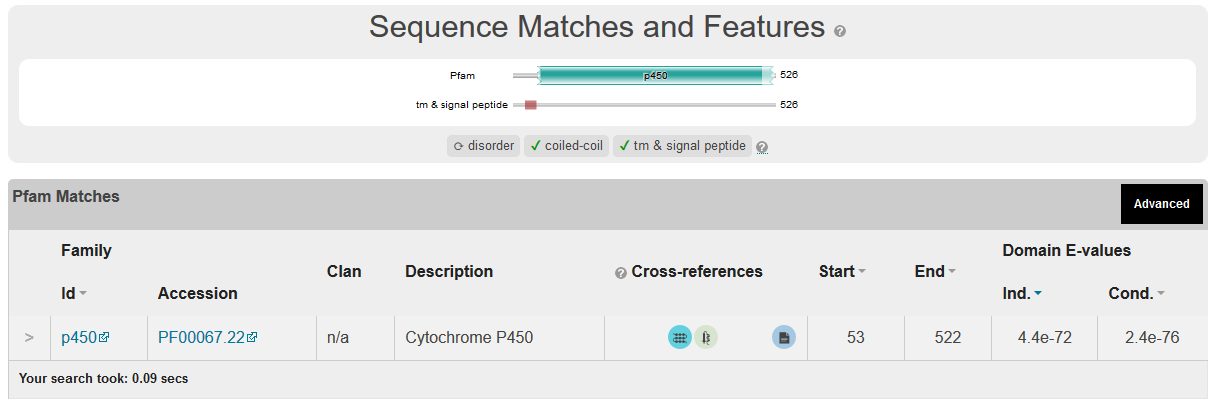

FGSG_11024

Interact pred =6.86

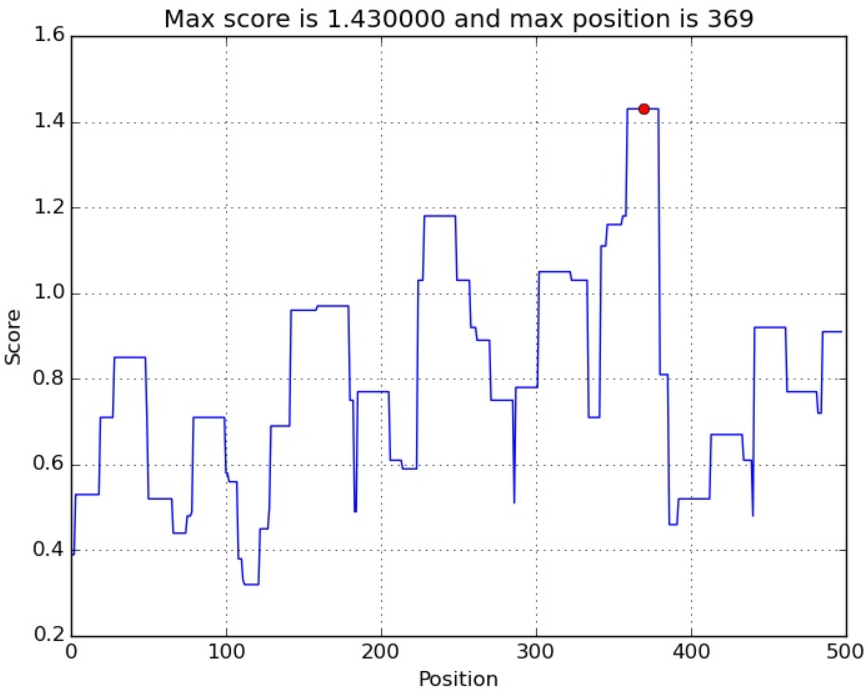

TM=12=34

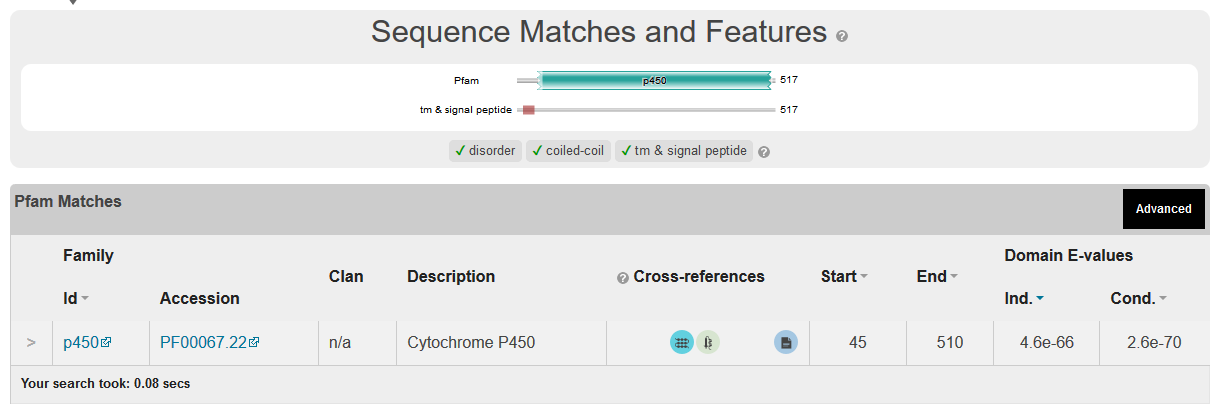

FGSG_04092

Interact pred = 2.35

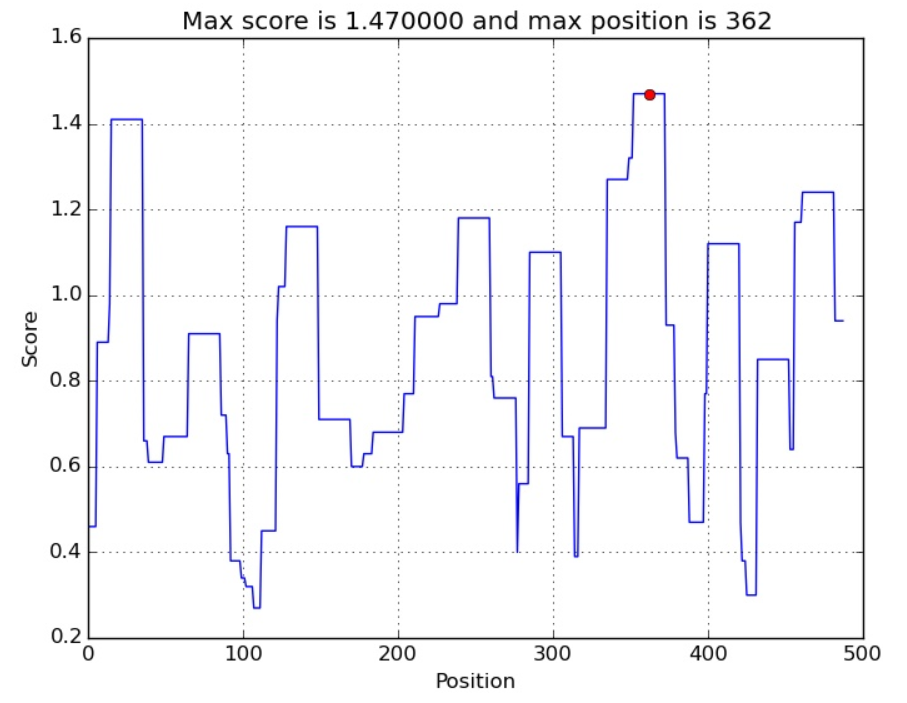

TM= 6-25 and 37-53

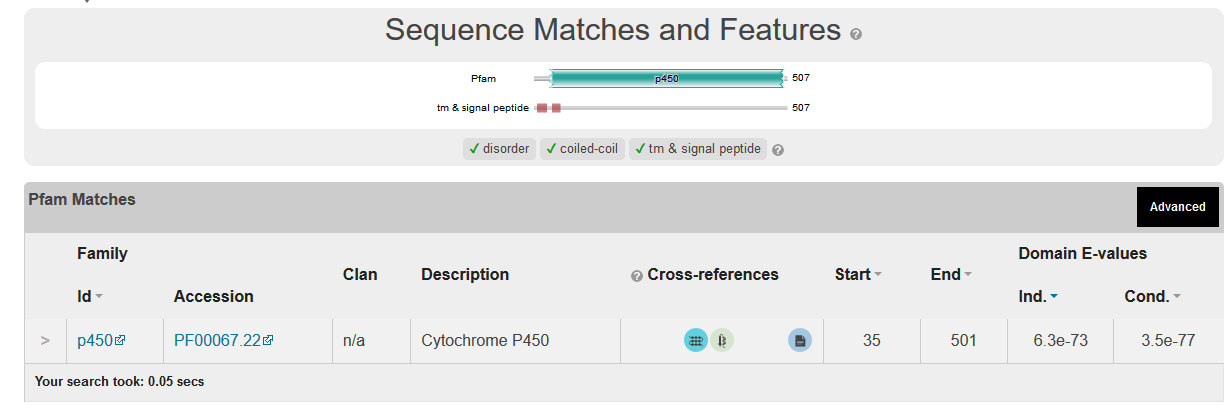

trichodiene oxygenase [Fusarium graminearum PH-1] FGSG_03535

Intereact pred= 1.52 yes

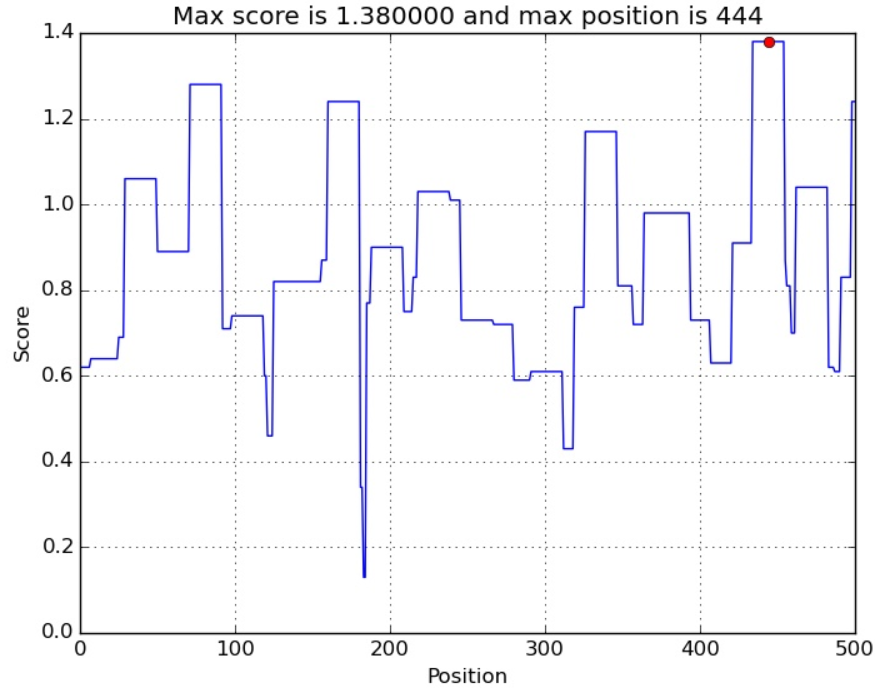

TM=20-40

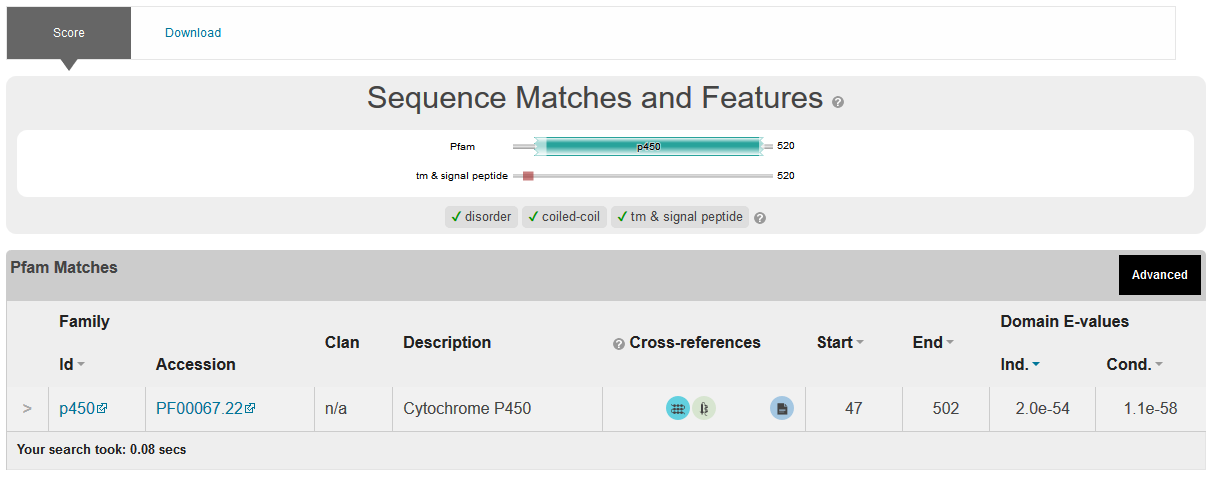

>pdb|5EQB|A Chain A, Lanosterol 14-alpha Demethylase

IPS= 1.06

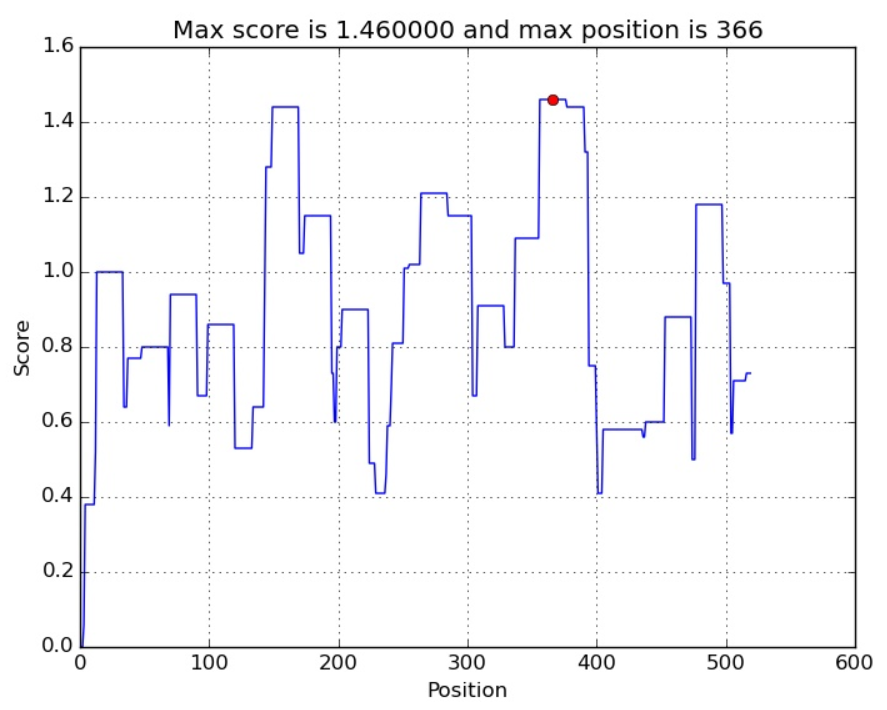

TM 30-51 and 58-74

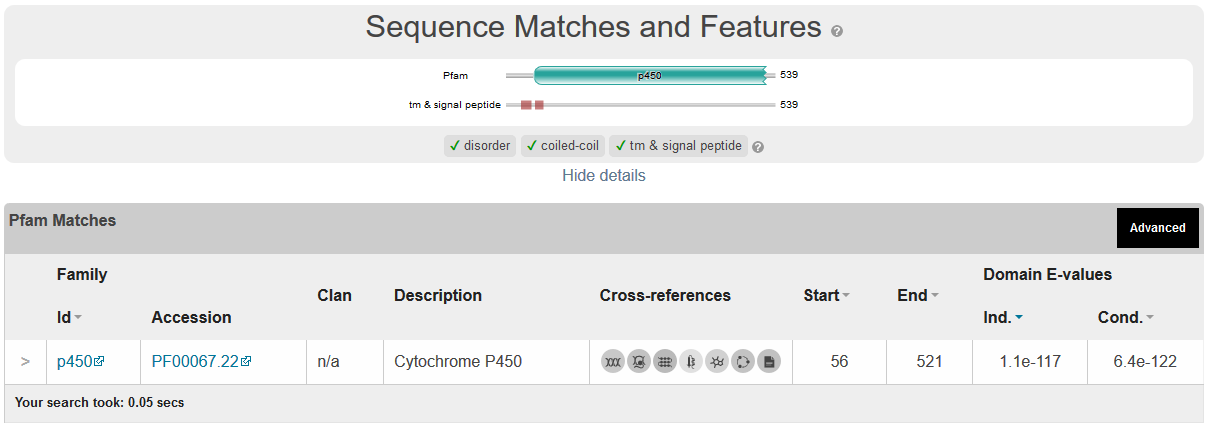

Pyricularia

Pred interact=1.45 yes

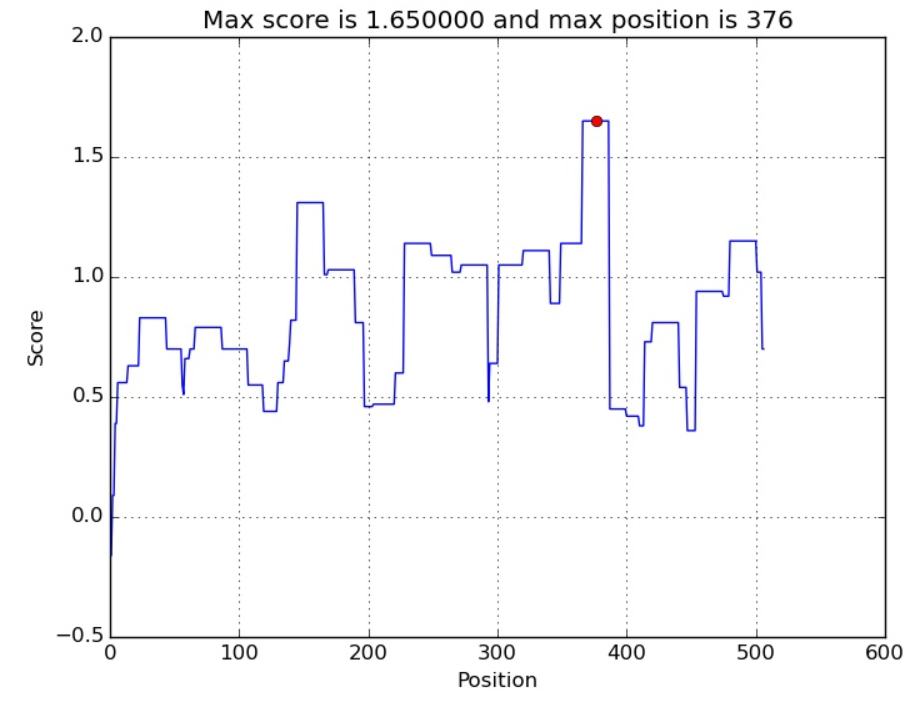

TM=20-41

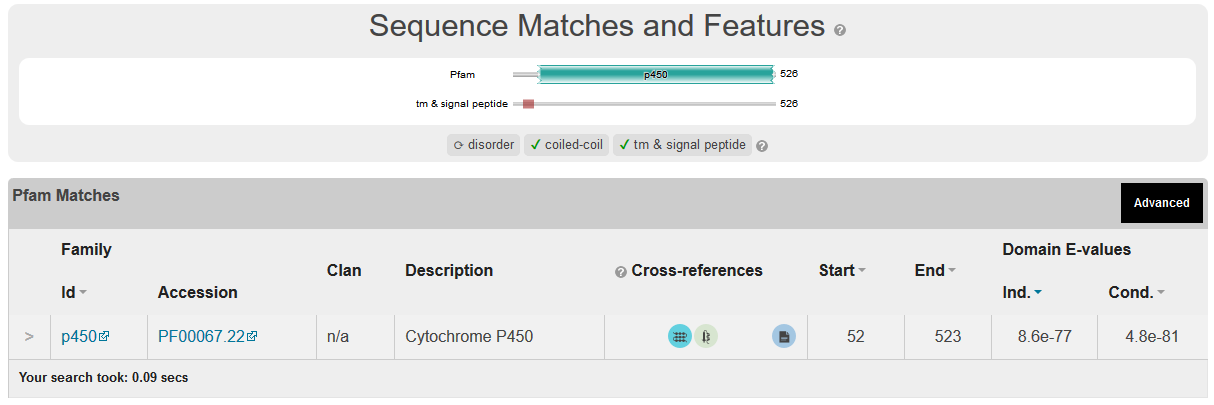

Metarhizium anisopliae

Int pred=1.3 Yes

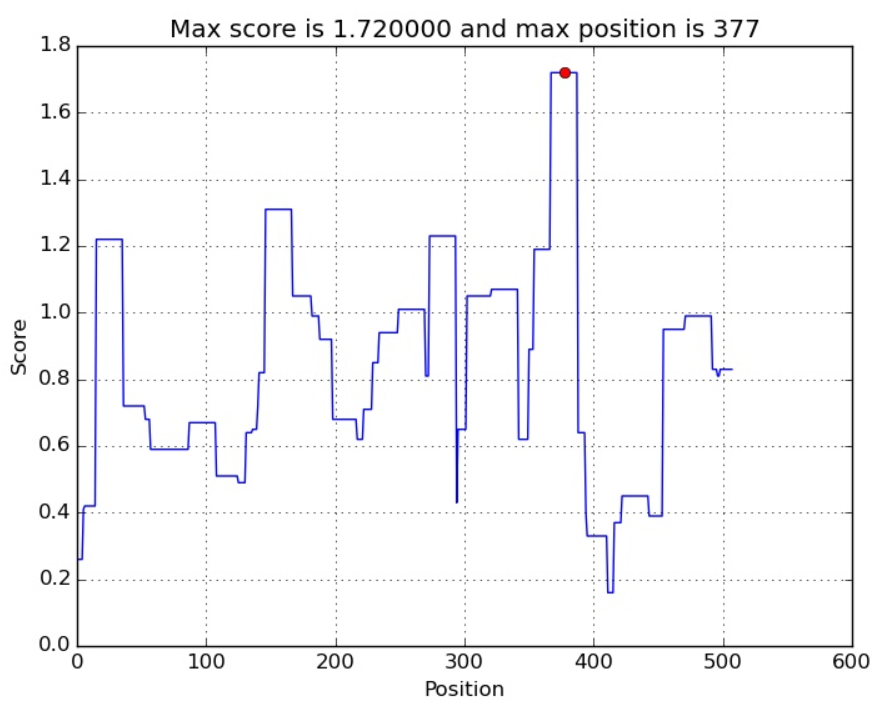

TM= 21-42 and 54-71

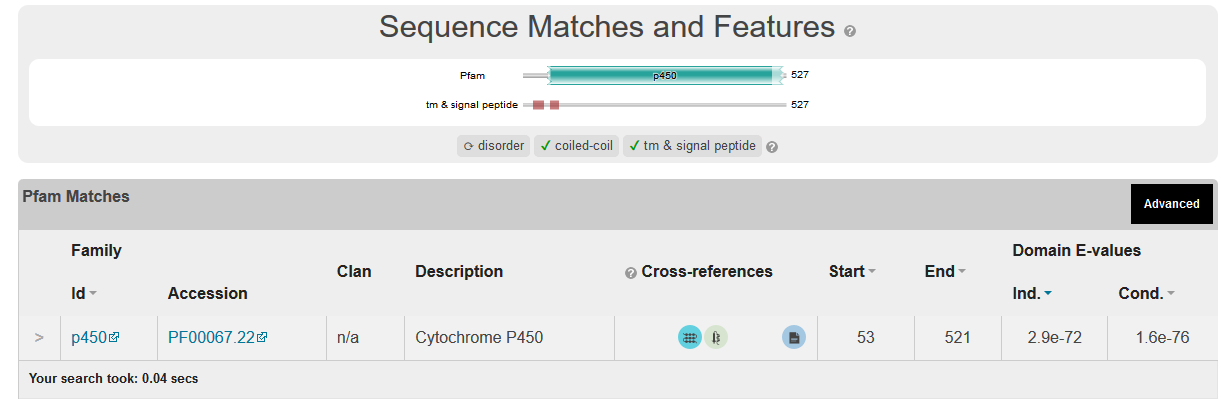

Verticillium dahliae

Int pred 1.02 Yes

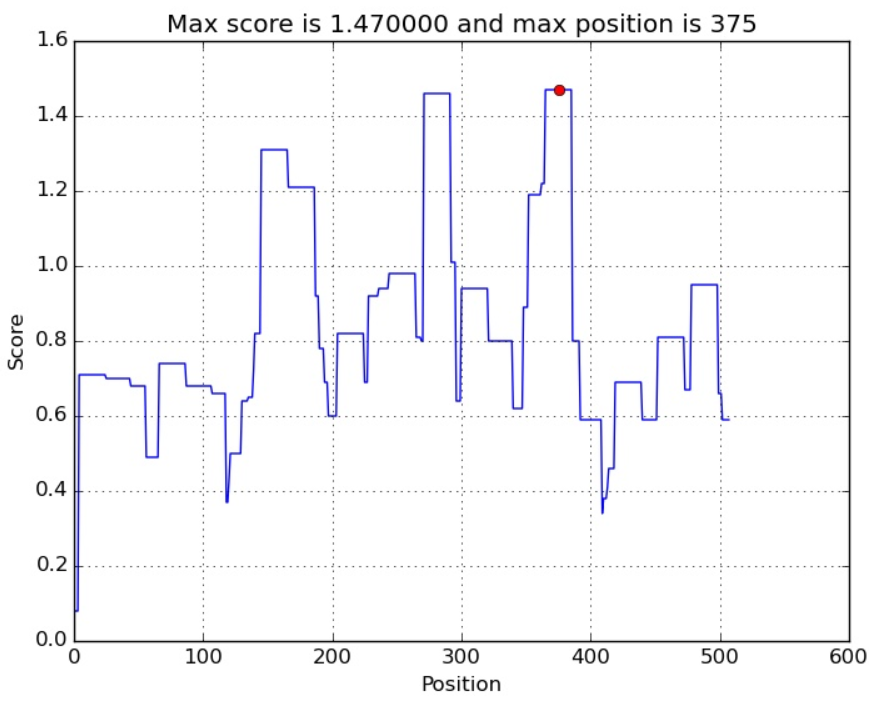

TM=20-41

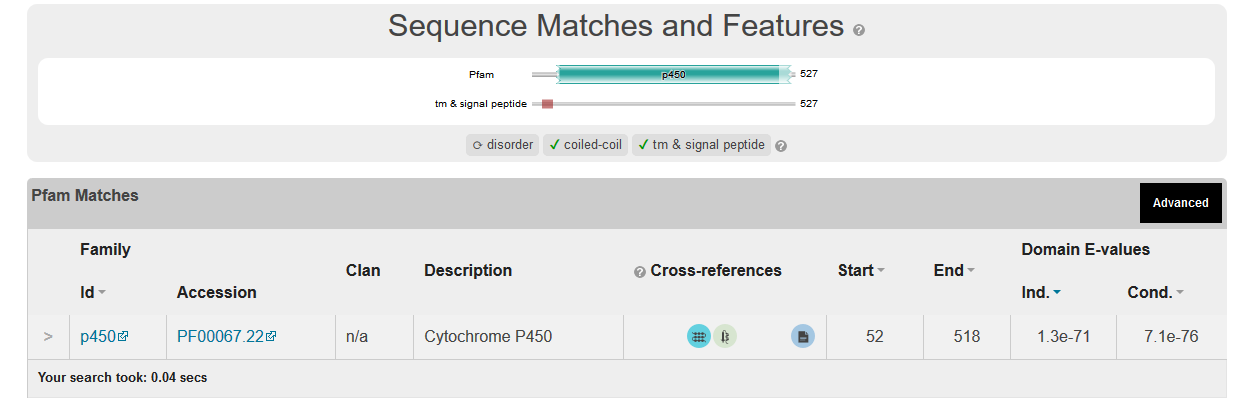

Fusarium oxysporum

IPS 1.43 Yes

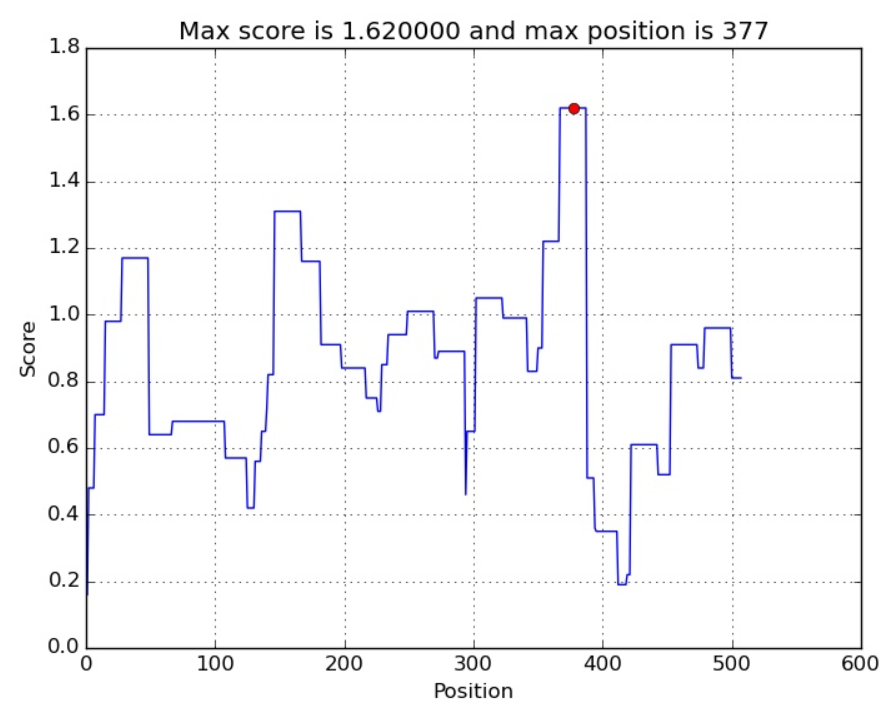

TM 25-43 and 77-71

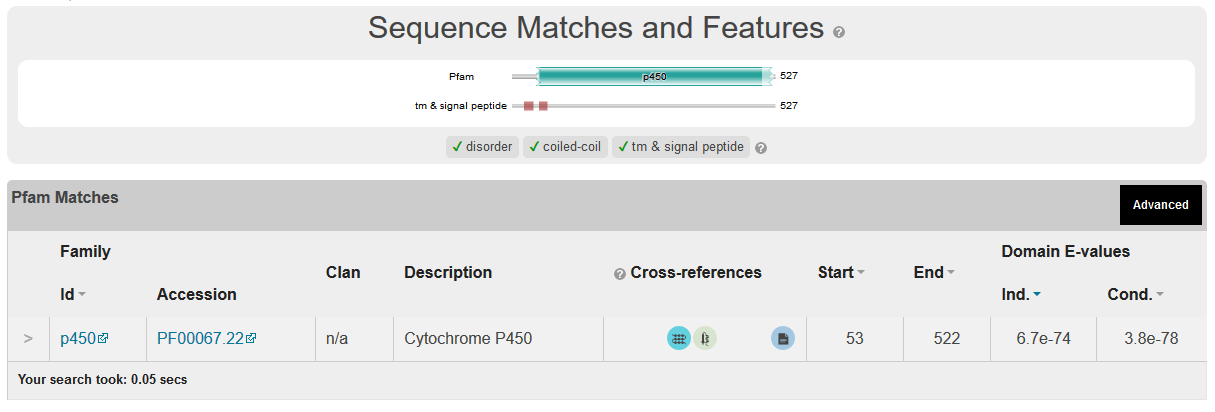

Aspergillus niger

IPS 4.61 Yes

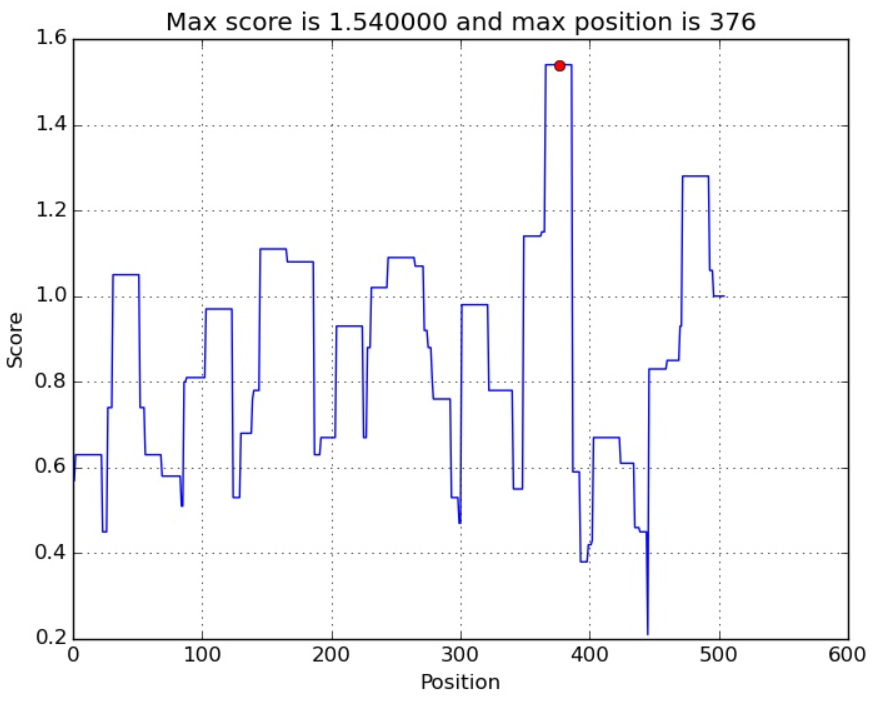
TM

TM 20-41 and 53-70

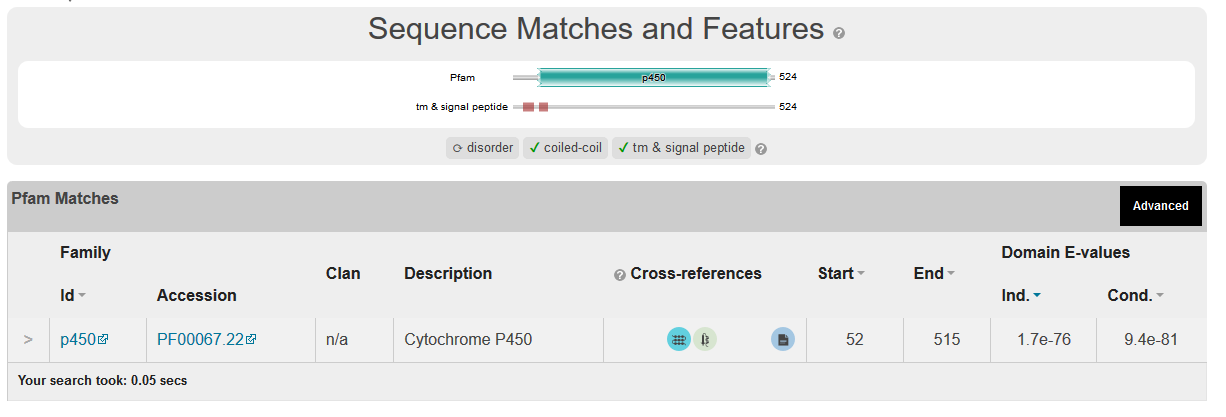

Crucibulum leave

1.64 Yes

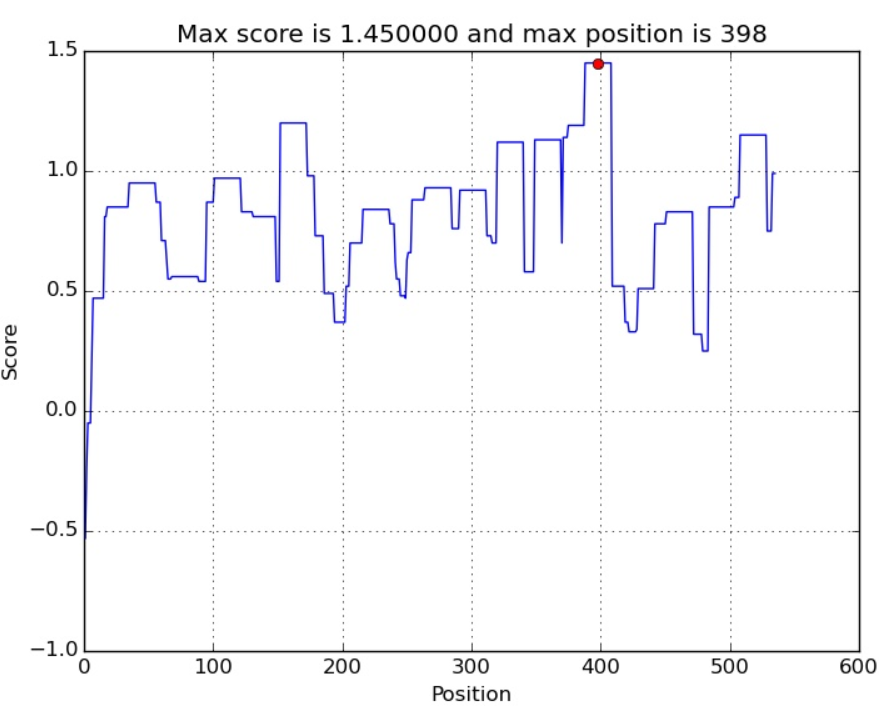

TM=35-53

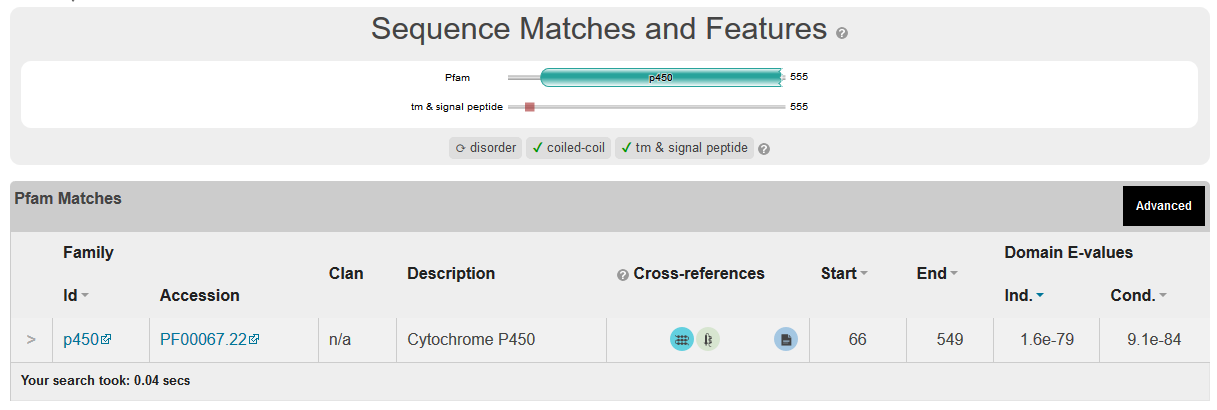

Blumeria graminis

IPS 2.06 Yes

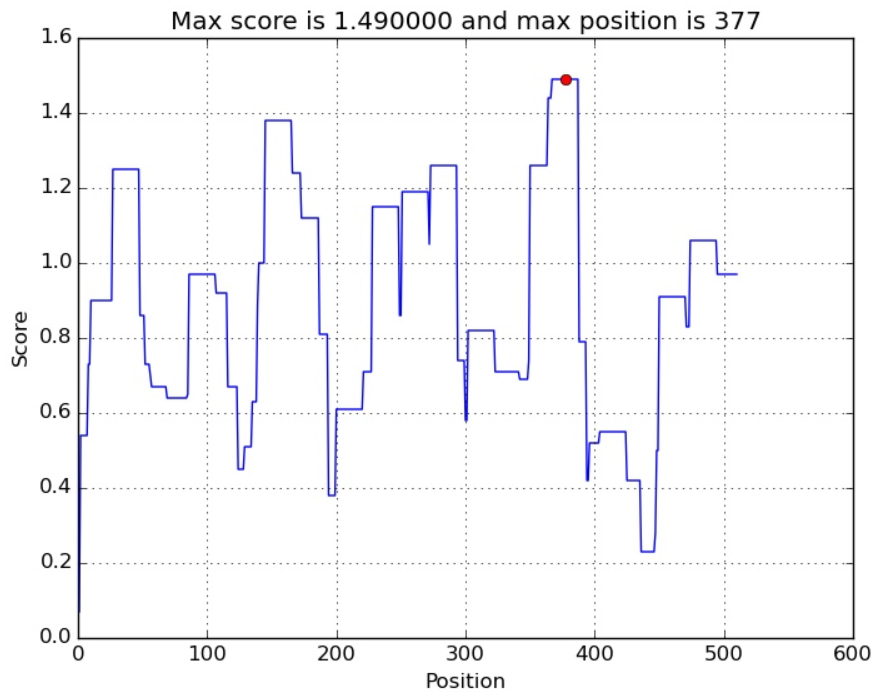

TM 20-41 and 53-70

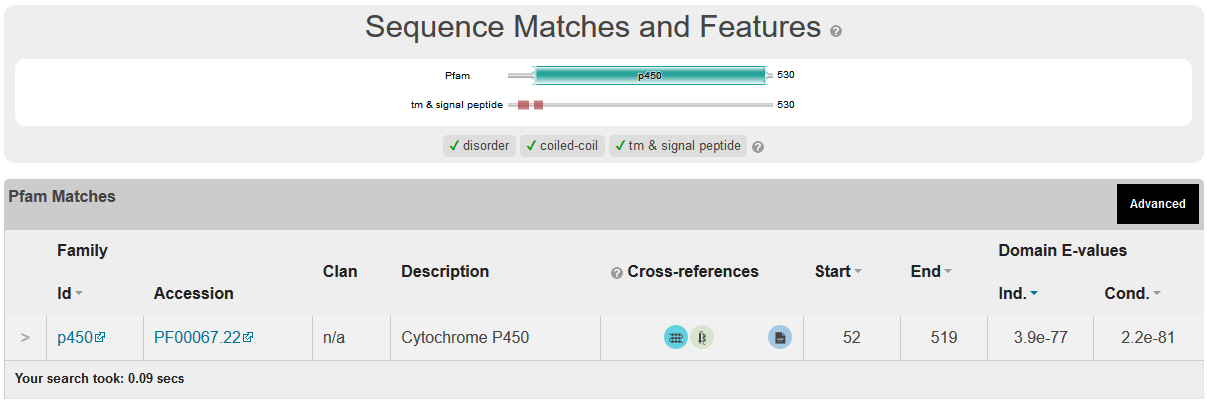

Crucibulum leave

IPS 1.64 Yes

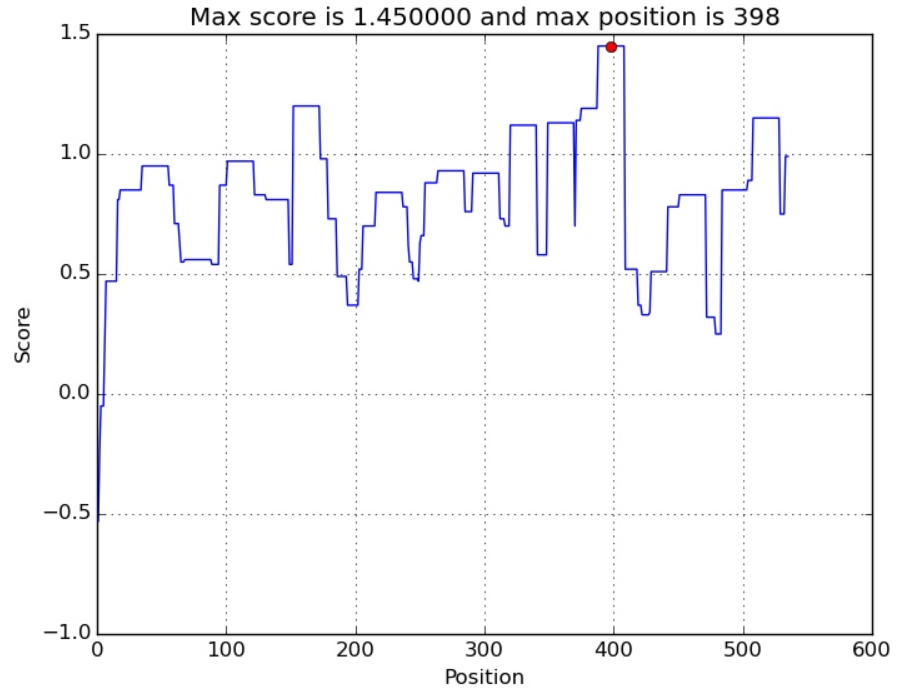

TM=35-53

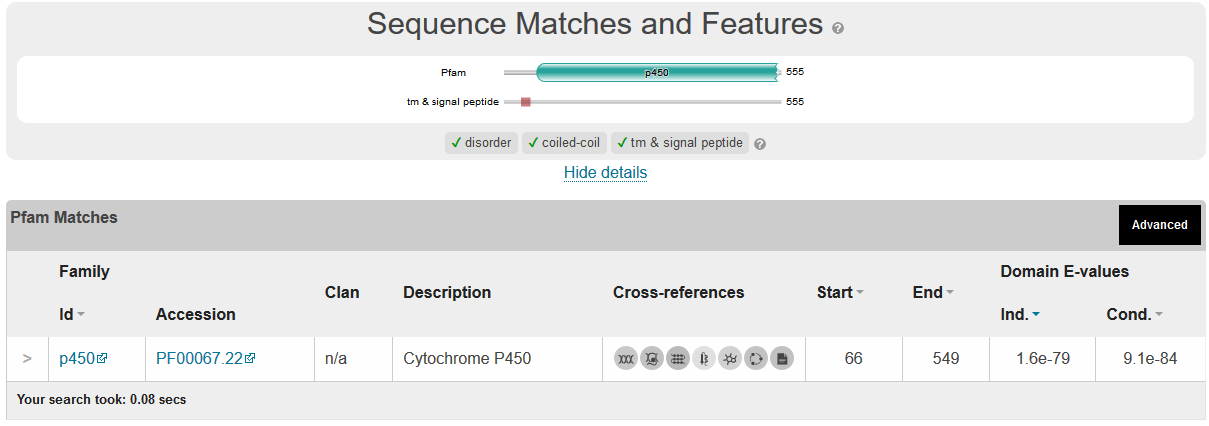

Agaricus bisporus

IPS 1.92 Yes

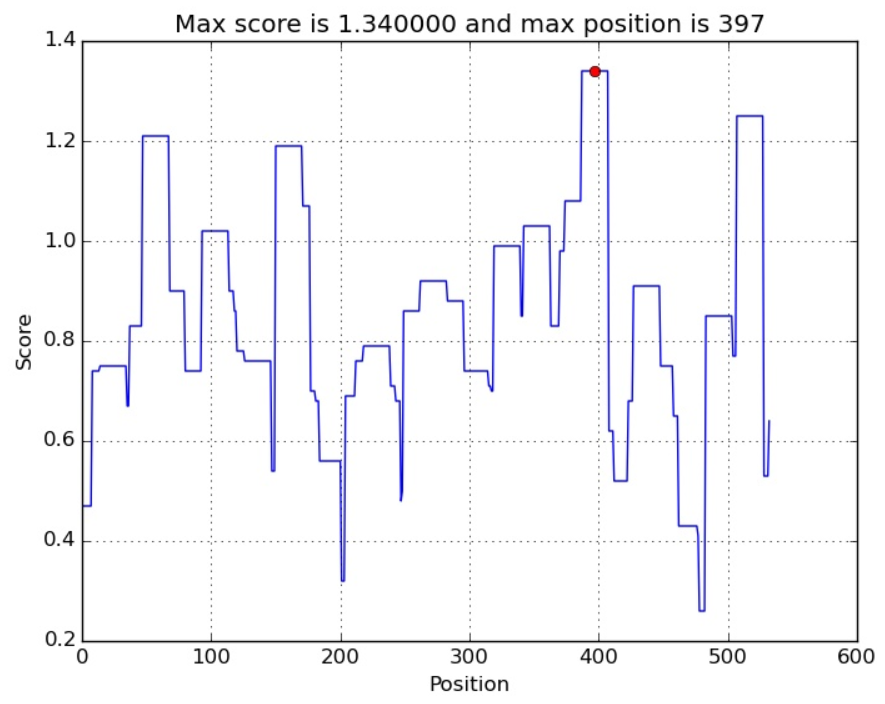

TM=33-51

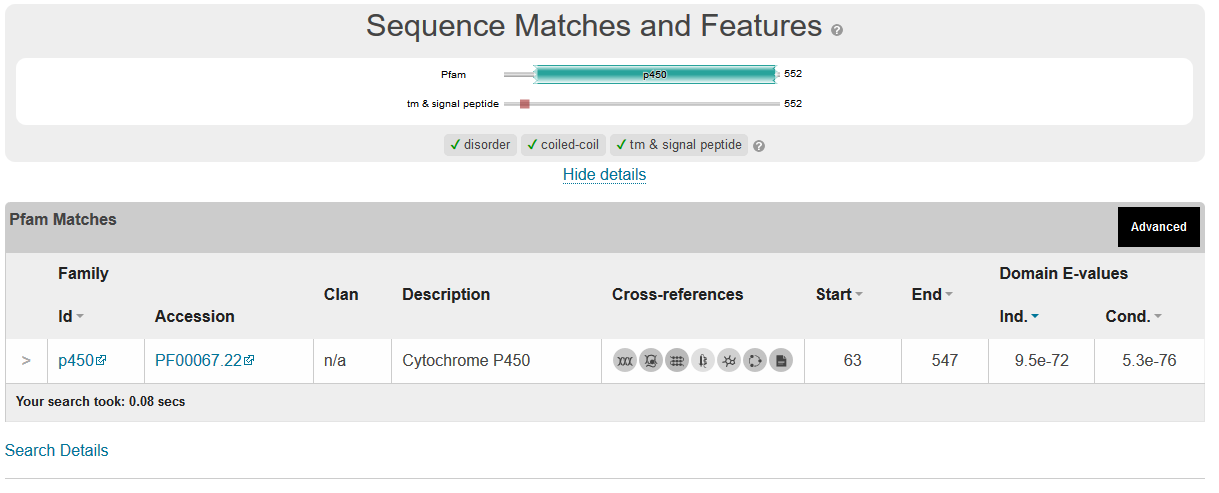

Serpula lacrymans

IPS 2.06 Yes

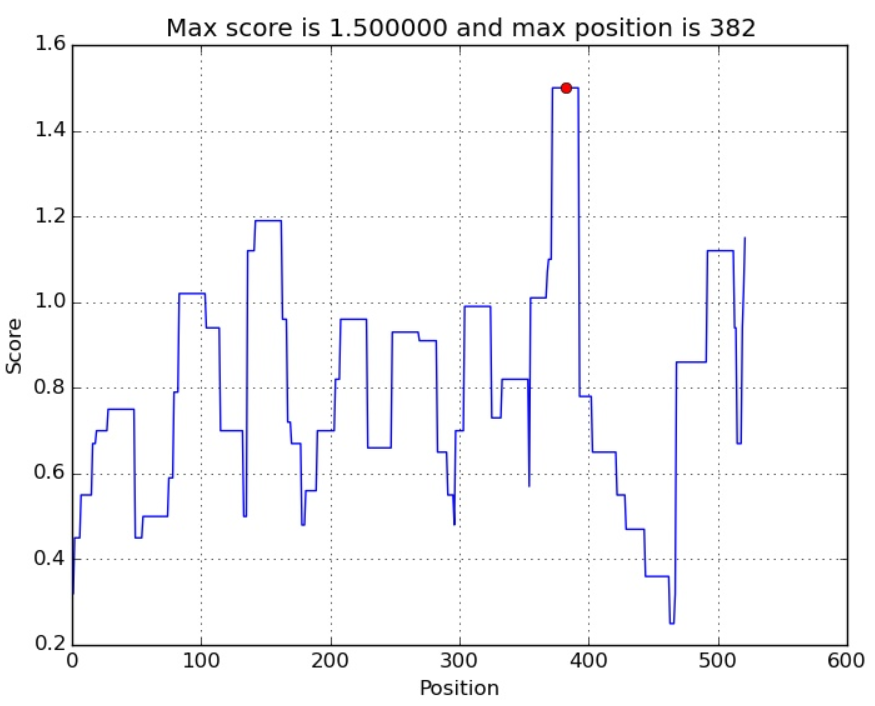

TM=20-37

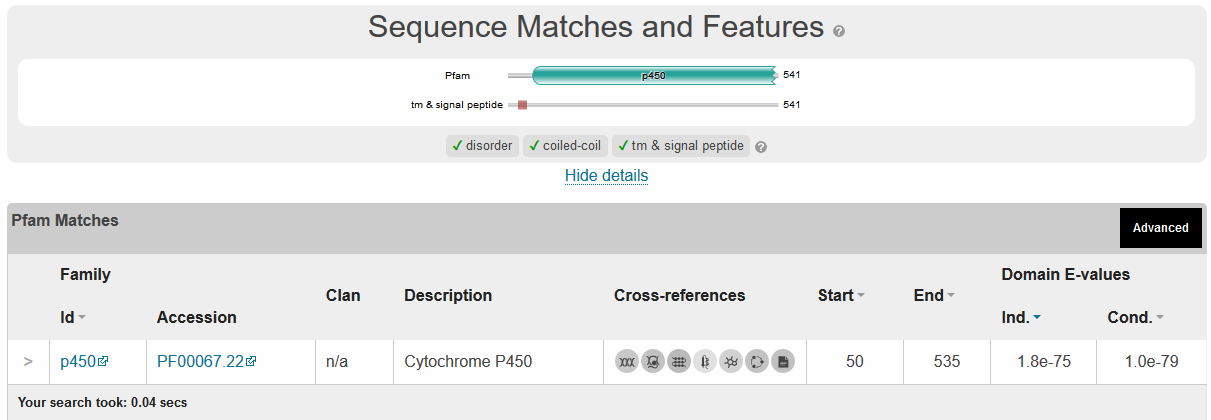

Pleurotus ostreatus

IPS 2.05 Yes

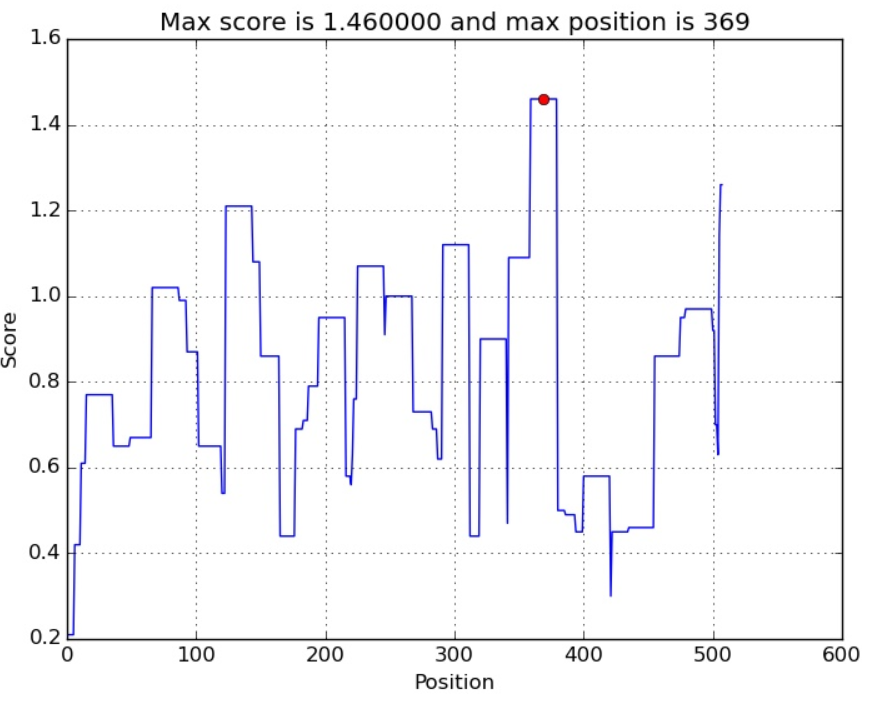

TM 7-28

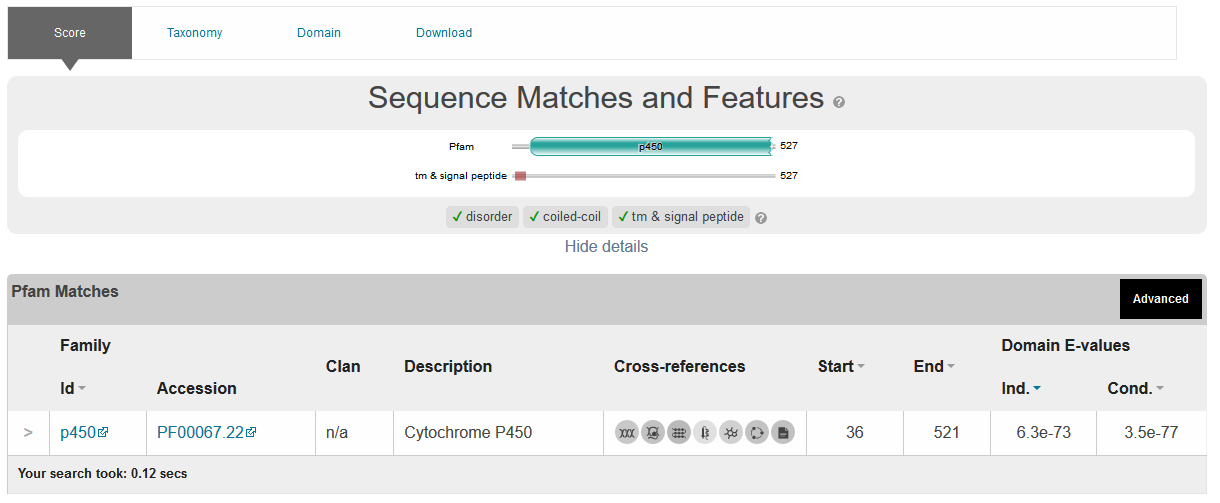

Drosophila melanogaster

IPS 8

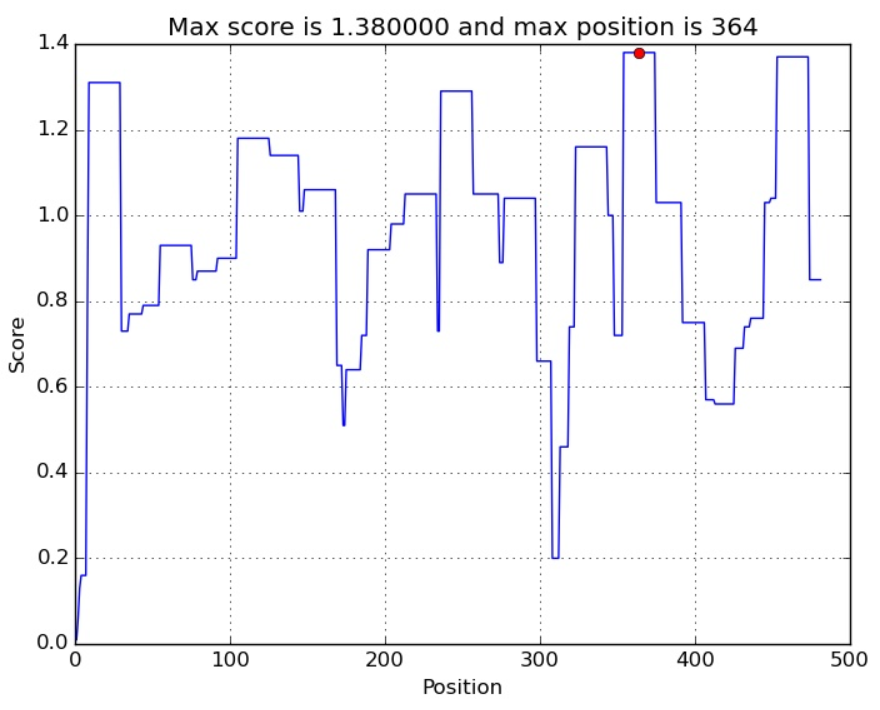

TM 1-18

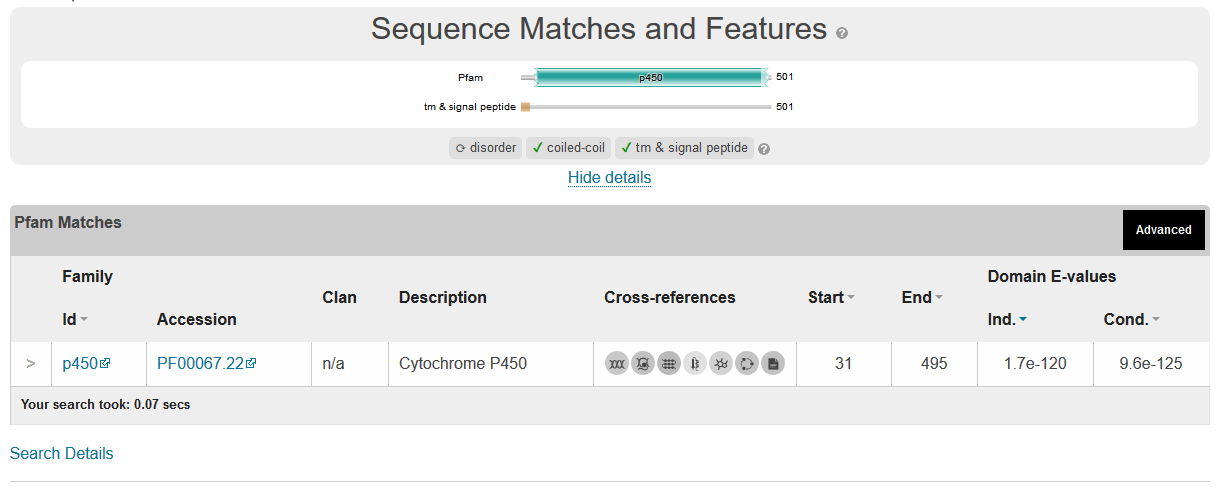

Musca domestica

IPR 2.85

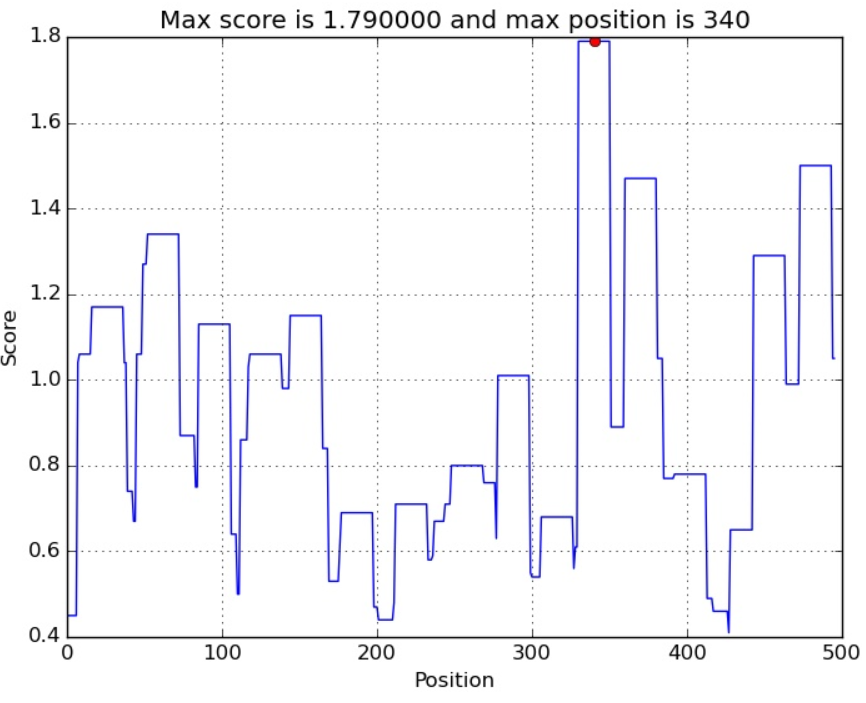

TM 6-22

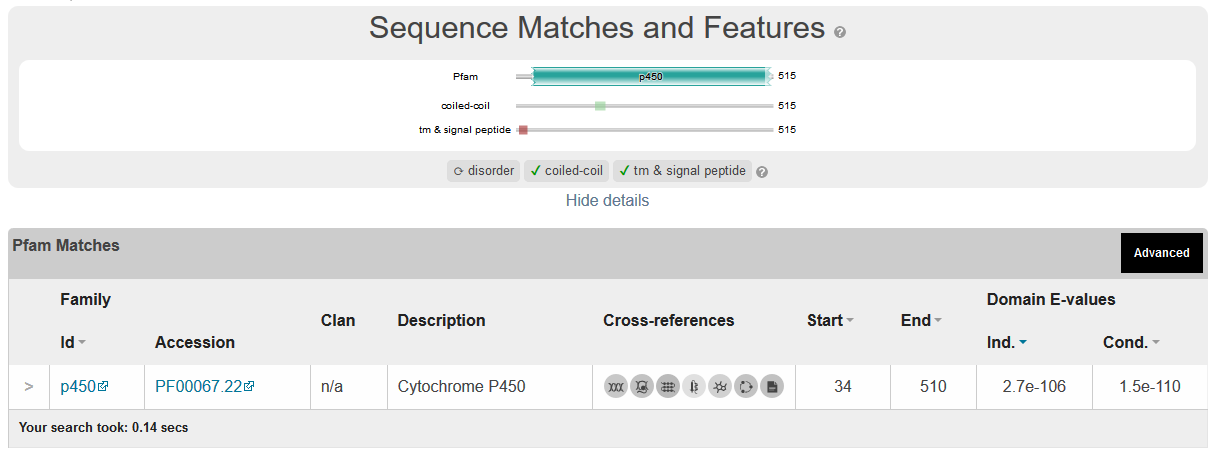

Gallus gallus

IPR 2.77	Position =250

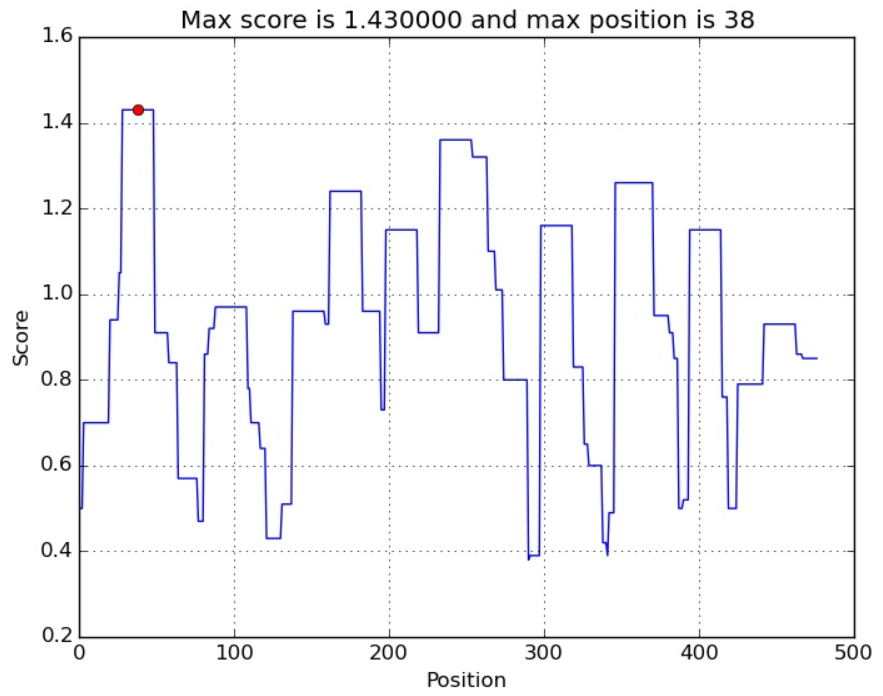

TM 20-38

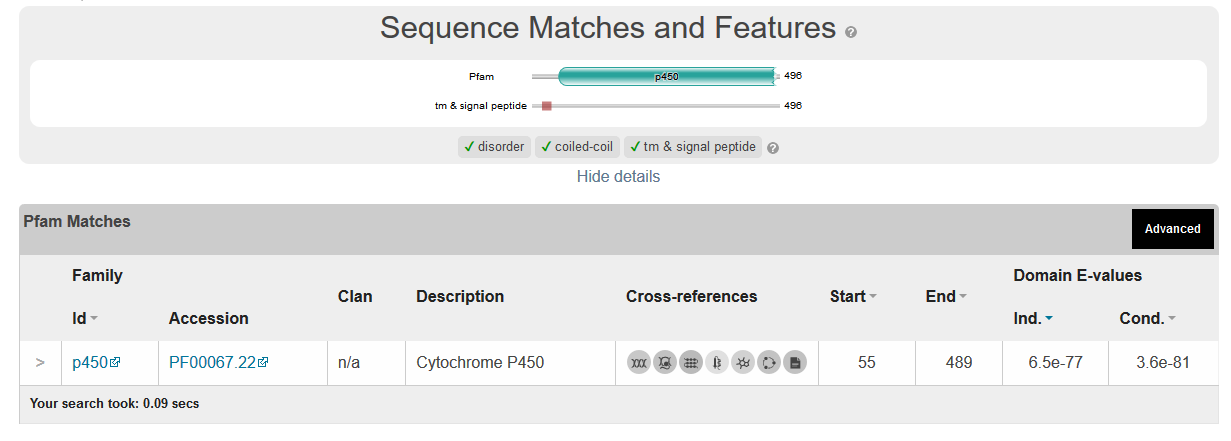

Homo sapiens

IPR 3.58 Yes

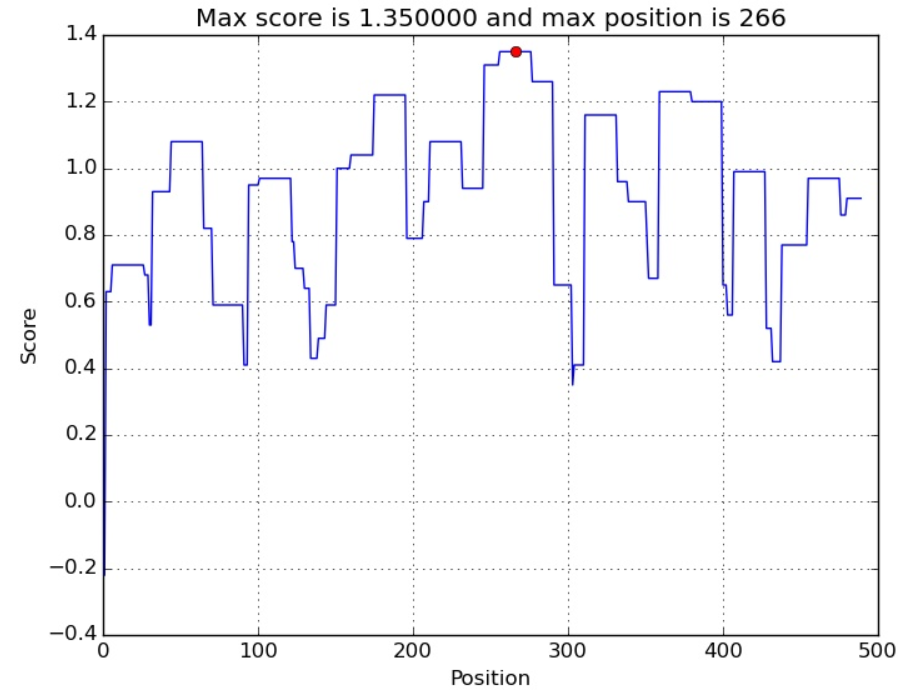

TM1-21 and 31-50

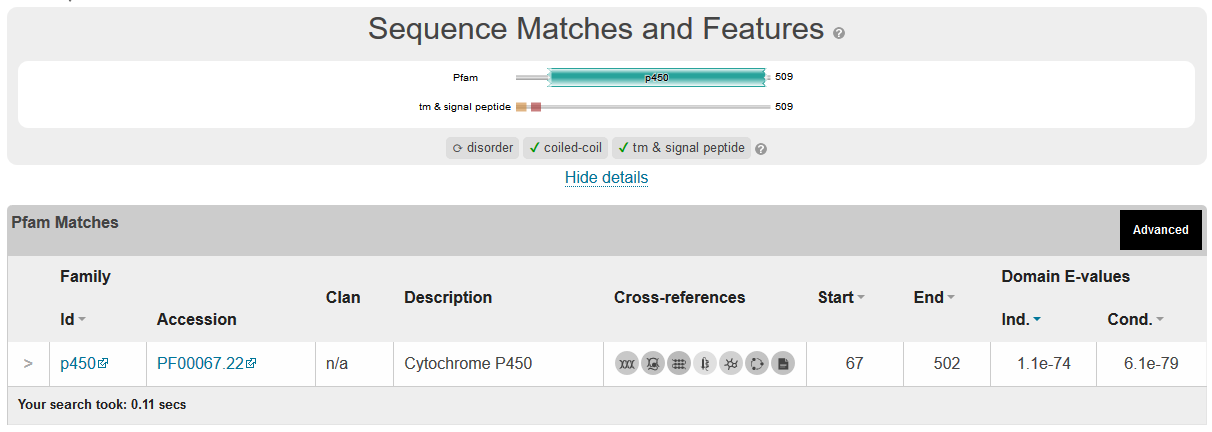

Arabidopsis thaliana

Idp 0.77 On the edge

Position 240 second highest not close to the membrane

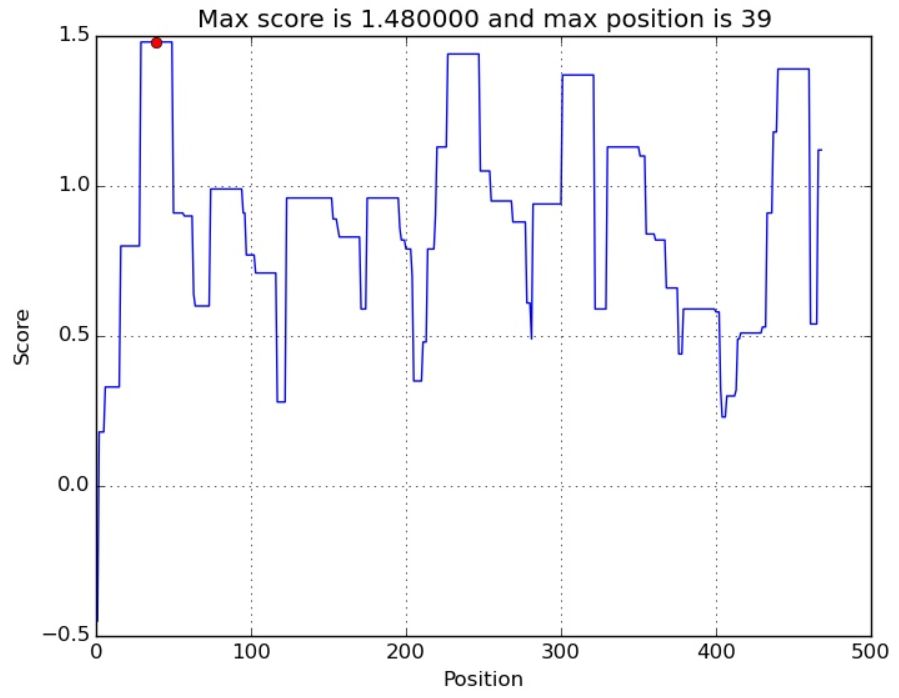

TM 12-30

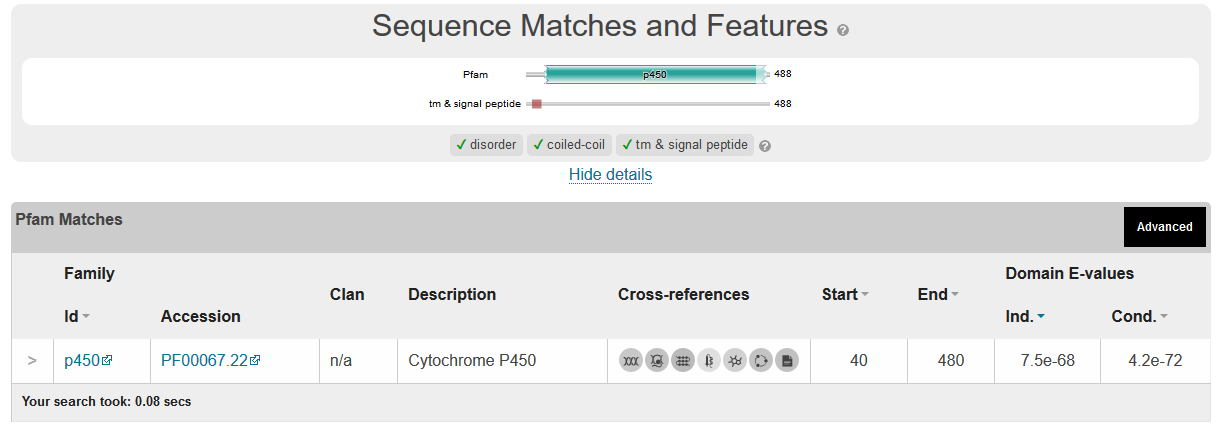

Triticum aestivum

IDP 2.6

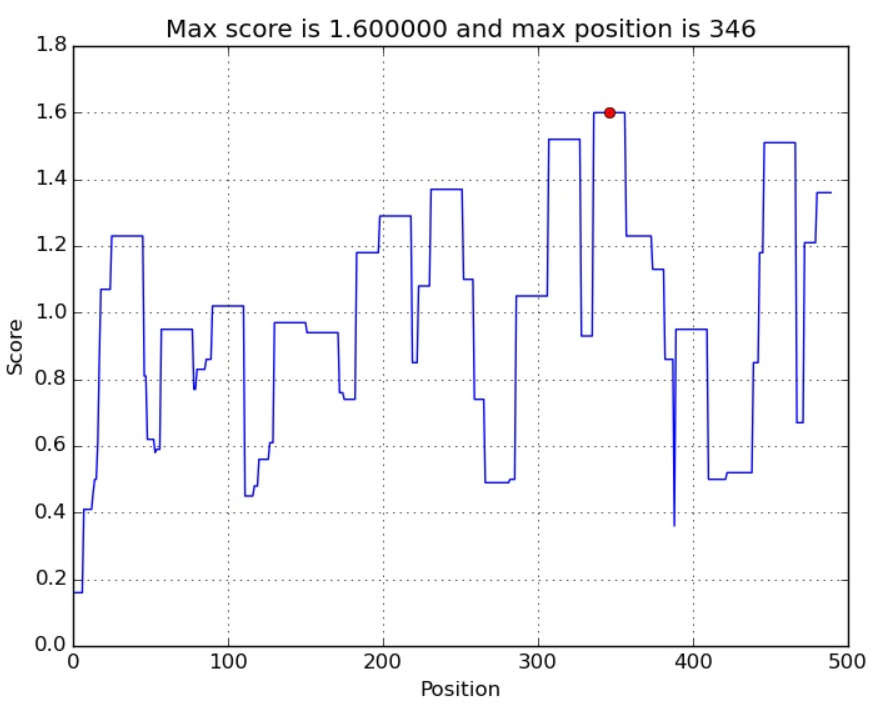

TM 6-29 and 41-60

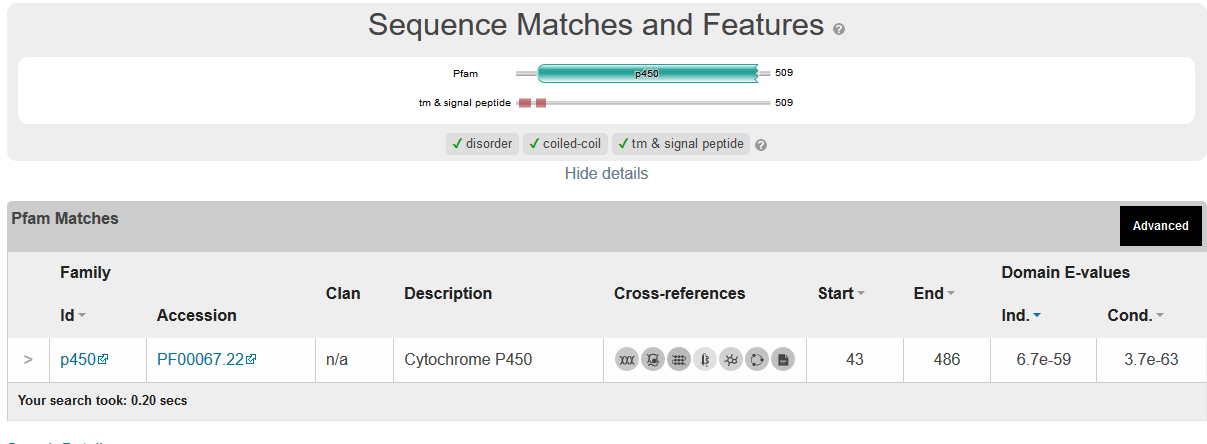

Trypanosoma brucei

IDP 5.95

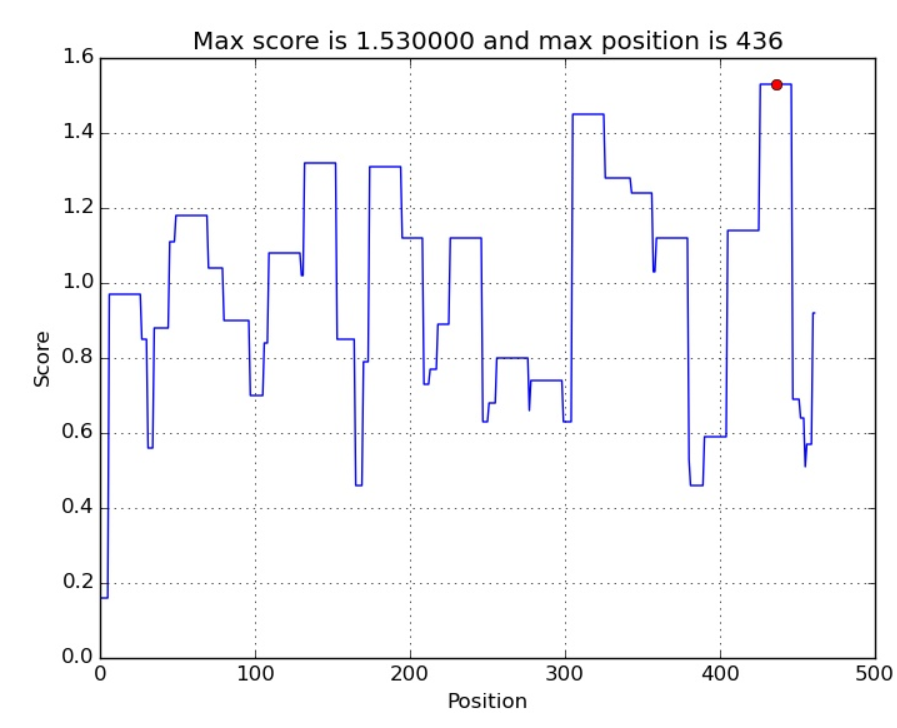

TM 1-23

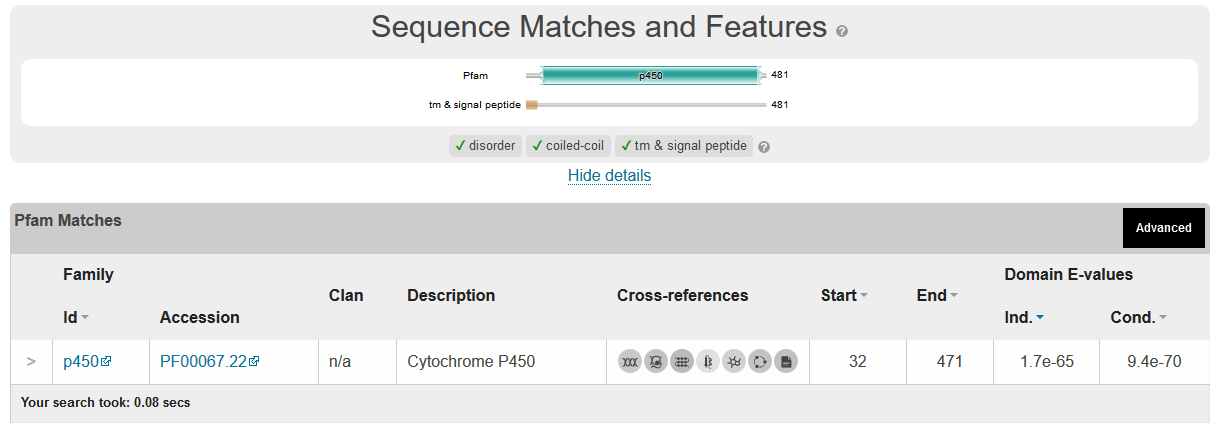

Dictyostelium discoideum

IDP 1.12
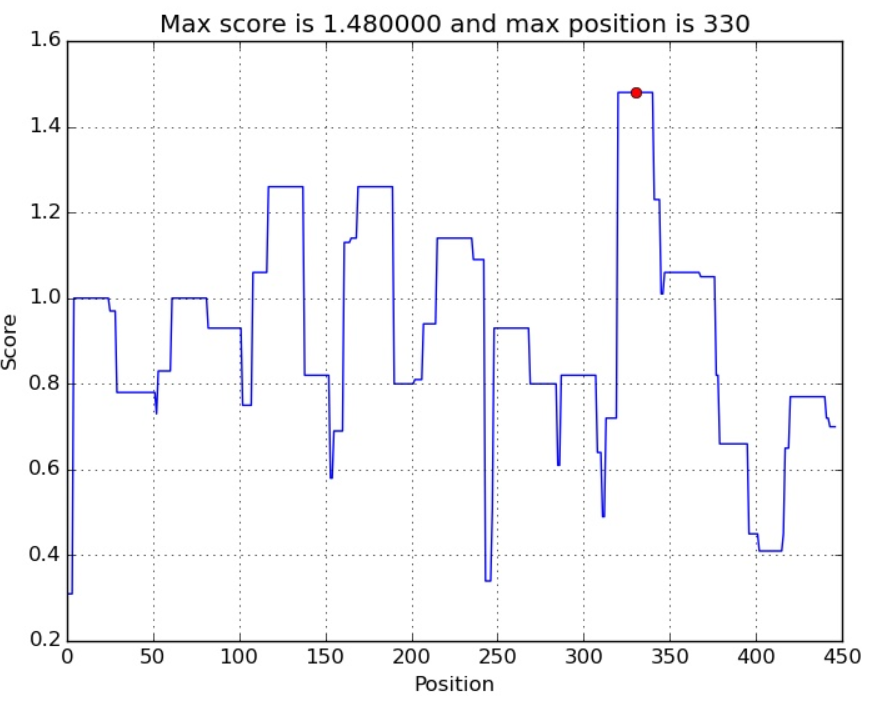

TM 1-17

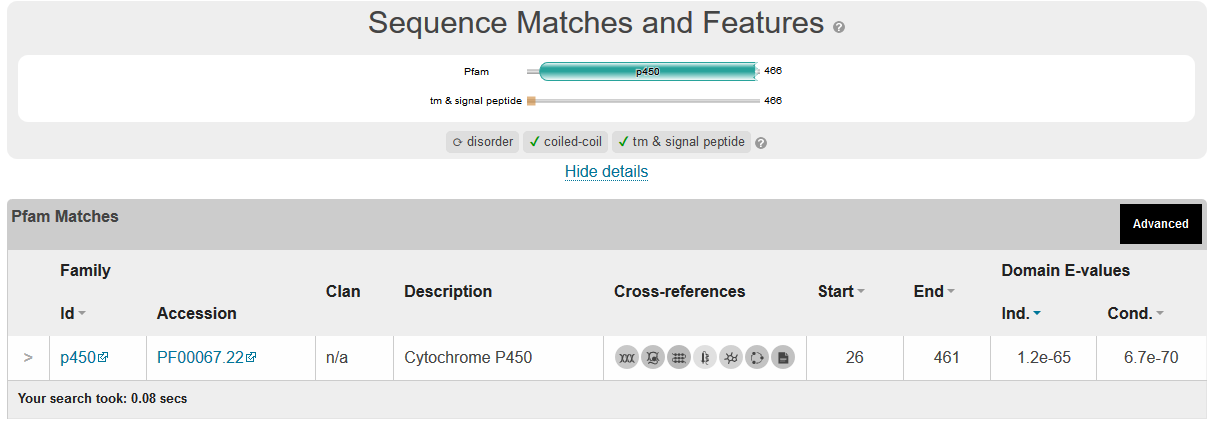

MAMMALS

>NP_064394.2 lanosterol 14-alpha demethylase [Mus musculus]

MVLLGLLQSGGWVLGQAMEQVTGGNLLSTLLIACAFTLSLVYLFRLAVGHMVQLPAGAKSPPHIYSPIPF

LGHAIAFGKSPIEFLENAYEKYGPVFSFTMVGKTFTYLLGSDAAALLFNSKNEDLNAEEVYGRLTTPVFG

KGVAYDVPNAIFLEQKKIIKSGLNIAHFKQYVPIIEKEAKEYFQSWGESGERNVFEALSELIILTASHCL

HGKEIRSQLNEKVAQLYADLDGGFTHAAWLLPAWLPLPSFRRRDRAHREIKNIFYKAIQKRRLSKEPAED

ILQTLLDSTYKDGRPLTDEEISGMLIGLLLAGQHTSSTTSAWMGFFLAKDKPLQEKCYLEQKAVCGEDLP

PLTYDQLKDLNLLDRCIKETLRLRPPIMTMMRMAKTPQTVAGYTIPPGHQVCVSPTVNQRLKDSWAERLD

FNPDRYLQDNPASGEKFAYVPFGAGRHRCVGENFAYVQIKTIWSTMLRLYEFDLINGYFPTVNYTTMIHT

PENPVIRYKRRSK

Pred interact = 2.4 Yes

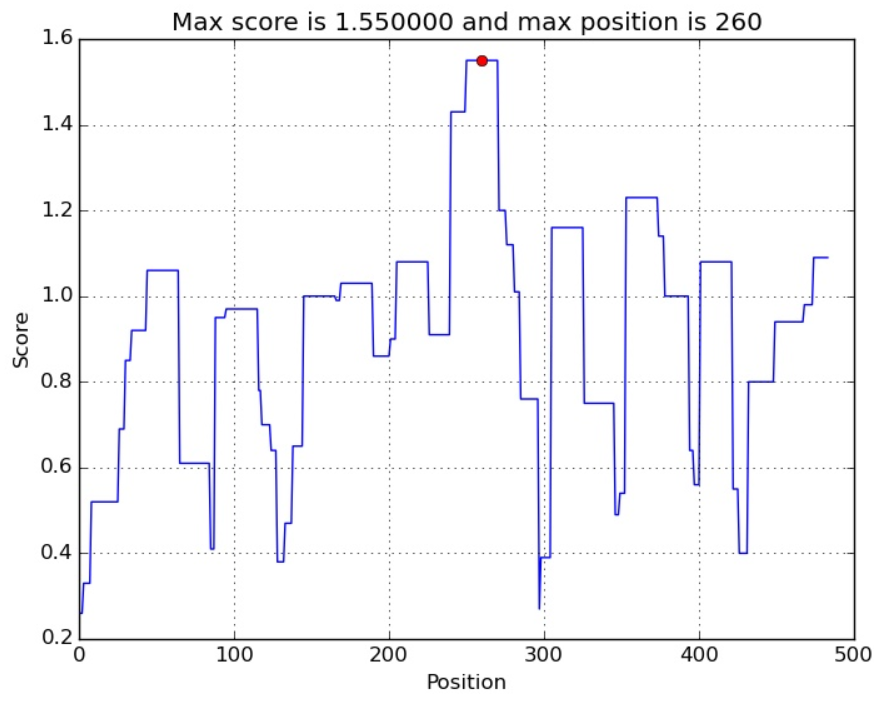

TM 1-17 and 27-44

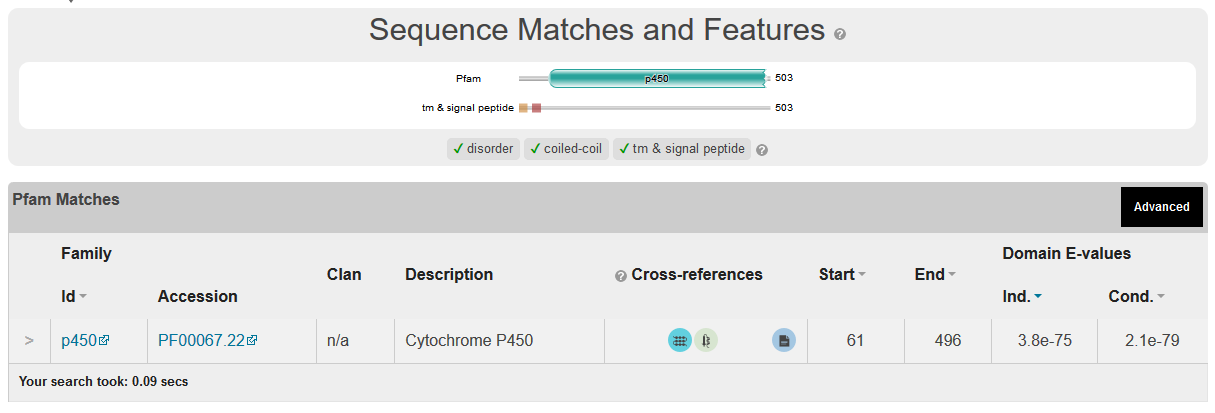
