## Supplemental Files and Data for "Evolutionary conserved multifunctional nitric oxide synthesis proteins responding to bacterial MAMPs are located at the endoplasmic reticulum": Supplementary file SF8 Phylogenetic and domain analysis of the NCP(NO) for production of NO at the ER membrane.docx

**Phylogenetic analysis of the NADPH Cytochrome Reductase responsible for NO production at the ER membrane.**

>FGSG_09786 PH-1

MAELDTLDVIVLGVIFLGTVAYFTKGKLWGVTKDPYANGFAAGGAAKPGRTRNIVEAMEESGKNCVIFYG

SQTGTAEDYASRLAKEGKSRFGLNTMIADIEDYDFDSLDTVPNDNVVMFVLATYGEGEPTDNAVDFYEFI

TGEDATFNEGNDPPLGNLNYVAFGLGNNTYEHYNAMVRKVDQALEKFGAHRIGEAGEGDDGAGTMEEDFL

AWKDPMWESLAKKMGLEEREAVYEPIFAINERDDLSPESNEVYLGEPNKLHLEGTAKGPFNSHNPYIAPI

AESYELFSAKDRNCLHMEVDISGSNLKYETGDHIAIWPTNPGEEVNRFLDILDLSGKQHSVITVKALEPT

AKVPFPNPTTYDAILRYHLEICAPVSRQFVSTLAAFAPNDSIKAEMNRLGSDKDYFHEKTGPHYYNIARF

LSSVSKGEKWTTIPFSAFIEGLTKLQPRYYSISSSSLVQPKKISITAVVESQQIPGRDDPFRGVATNYLF

ALKQKQNGDPSPAPFGQTYELTGPRNKYDGIHVPVHVRHSNFKLPSDPGKPVIMIGPGTGVAPFRGFVQE

RAKLARDGVEVGKTLLFFGCRKPSEDFMYEKEWQEYKEALGDKFEMITAFSRESAKKVYVQHRLKERAQE

VSDLLSQKAYFYVCGDASNMAREVNTVLAQIIAEGRGVSEAKGEEIVKNMRSANQYQEDVWS

>FGSG_08413 XP_011320363.1 hypothetical protein FGSG_08413 [Fusarium graminearum PH-1]

MTNVFAADTLLSQLGPPQSIADIAALSALGVASAAYLLRGITWDKPDHYHHVWFERMGSKGGSSSSHPKA

TRDIAKKLEETGKDVVIFWGSQSGTAETFANRLSKECHLRFGLQALCADLCDYDPESIANLSQSKLAIFI

LSTYGEGDPSDNTAAFWDWLTKTPNIQIPNLRYMAFGLGNTSYRYYNRVIDVVAQHLDKYGAQRLMPVGR

ANDAQGGTEEDFLSWKDDLYTHFQENLGYQERDIPYEPSIQLIQDESLDIMDLHLGEPIQNRNGPAKVVK

QYSPIRPLAIQSSQELYTSPGRNCLHMELDISNQPELRYRTGDHLAIYPINPDYEVQLLLKAFGLEDRAE

KPLLVQTLEEGTSTKIPSPTSALALFRHYLEVAAPVSRETVGQLARFAPLSESVETLTALAKNKEAYATY

IGSNHITMGRLLHLVAPGAIWTKLPLSYVVETLPCIQARYYSISSSSTVSARRLSITVGVDKSPLQQDPS

RVIRGITTNYLYALGNALNCDTSQSVVGSDAPSYALSGPGDTLNGRKVFACIRRSNFKLPTLSSTPIIMI

GAGTGLAPFRGFILERARLQAVGKPIGKMLLFFGCRSPDQDYLYQGELAEVAQKLQGSLEIVTAFSRAEE

EPKKYVQDRVEERKTQVCHLLQEGASIYFCGRAAMARKVGNKVEESMKTQNNWTDAEARSWAESVKKGNK

WLEDVWG

>YeastNCP1 YHR042W SGDID:S000001084

MPFGIDNTDFTVLAGLVLAVLLYVKRNSIKELLMSDDGDITAVSSGNRDIAQVVTENNKNYLVLYASQTGTAEDYAKKFSKELVAKFNLNVMCADVENYDFESLNDVPVIVSIFISTYGEGDFPDGAVNFEDFICNAEAGALSNLRYNMFGLGNSTYEFFNGAAKKAEKHLSAAGAIRLGKLGEADDGAGTTDEDYMAWKDSILEVLKDELHLDEQEAKFTSQFQYTVLNEITDSMSLGEPSAHYLPSHQLNRNADGIQLGPFDLSQPYIAPIVKSRELFSSNDRNCIHSEFDLSGSNIKYSTGDHLAVWPSNPLEKVEQFLSIFNLDPETIFDLKPLDPTVKVPFPTPTTIGAAIKHYLEITGPVSRQLFSSLIQFAPNADVKEKLTLLSKDKDQFAVEITSKYFNIADALKYLSDGAKWDTVPMQFLVESVPQMTPRYYSISSSSLSEKQTVHVTSIVENFPNPELPDAPPVVGVTTNLLRNIQLAQNNVNIAETNLPVHYDLNGPRKLFANYKLPVHVRRSNFRLPSNPSTPVIMIGPGTGVAPFRGFIRERVAFLESQKKGGNNVSLGKHILFYGSRNTDDFLYQDEWPEYAKKLDGSFEMVVAHSRLPNTKKVYVQDKLKDYEDQVFEMINNGAFIYVCGDAKGMAKGVSTALVGILSRGKSITTDEATELIKMLKTSGRYQEDVW

>XP_003715306.1 NADPH-cytochrome P450 reductase, variant [Pyricularia oryzae 70-15]

MAELDTLDIIVLGLILCGTIAYFTKGKYWGVVKDPYAASFANTNGPKTGKTRNIVEKMEETGKNCVIFYG

SQTGTAEDYASRLAKEGKSRFGLETMVADLEDYDYENLDTVPSDKIVMFVLATYGEGEPTDNAVDFYEFI

TGEDVSFSEGSTLDNLNYVAFGLGNNTYEHYNSMVRNVNKALEKLGAHRIGDAGEGDDGAGTMEEDFLAW

KDPMWAALAEKMGLEEREAVYEPVFSVTEREGLTVESPEVYLGEPNKMHLEGTAKGPFNAHNPYIAPIVK

SYELFNVKDRNCLHIDVDVSGSNLTYQTGDHIAVWPTNPGEEVDCLLDVLGLTDKRDTVVSVRPLEPTAK

VPFPAPTTYDAILRYHMEICAPVSRQFIATLAAFAPDEETKAEMTKLGGDKDYFSAKISKHYLNIARVLF

NVGKGKKWNNIPFSAFIEGLTKLQPRYYSISSSSLVQPKVITITAVVEKQEIPGRDDPFRGVTTNYLLAL

KQKQNGEPHPEPFGRTFALAGPRDKYDGIKVPVHVRHSNFKLPSDSTKPIILVGPGTGVAPMRAFVQERA

KQAENGEEVGKTILFFGCRKSTEDFLYKDEWDEYKKVLGDKFELVTAFSREGPKKVYVQHRLKERAQEIN

ELLTKKAYIYVCGDAANMAREVNSVLGQIIAEQRGIPEAKAEEIVKNMRAANQYQEDVWS

>KFG88153.1 cytochrome P450 oxidoreductase [Metarhizium anisopliae]

MAEFDTLDIVVLGVILLGTIAYFTKGTLWGVTKDPYANAFANANGAKAGRSRNILEKMDETGKNCVIFYG

SQTGTAEDYASRLAKEGKSRFGLETMVADLEEYDFDNLDAMPSDKVAMFILATYGEGEPTDNAVEFYEFI

TGDDVSFSEGSDPALQNLNYVAFGLGNNTYEHYNSMVRNVDKALQKLGANRIGEAGEGDDGAGTMEEDFL

AWKDPMWAALAEKMGLEEREAVYEPTFGIVDRENLTVDSPEVYLGEPNKMHLEGTAKGPFNSHNPYIAPI

AESRELFSAKDRNCIHMDVDINGSNLSYQTGDHIAIWPTNSGDEVDRFLDIIGLKEKRNNVISIKALEPT

AKVPFPTPTTYDAIVRYHLEICAPVSRQFVATLAAFAPNDEVKAEMARLGSDKEYFHSKTGPHFYNIARL

LDTVGKGEKWTNIPFSAFIEGLNKLQPRYYSISSSSLVQPKKISITAVIESQAIPGRKDPFRGVATNYLF

ALKQKQNGDPNPSPFGKTYALNGPRNKFDGIHVPVHVRHSNFKLPSDPAKPVIMVGPGTGVAPFRGFIQE

RAKQAQNGATVGRTILFFGCRKRSEDFLYESEWEEYKKALGDSLEIVTAFSRESSKKVYVQHRLKERSKE

IGELLSQKAYFYVCGDAAHMAREVNTVLAQIIAESRGVSETKGEEIVKNMRAANQYQEDVWS

>XP_009658187.1 NADPH-cytochrome P450 reductase [Verticillium dahliae VdLs.17]

MAELDTLDIIVLSVILLGTAAYFTKGKYWAIEKDPYANGFASAGGPKAGKTRNIIEKLDESGKNCVIFYG

SQTGTAEDYASRLAKEGKSRFGLETMVADLEDYDFDNLDAVSTDKVVMFVLATYGEGEPTDNAVEFYEFI

TGEDVSFNEANEPALGNLNFVAFGLGNNTYEHYNSMVRNVTKALEKLGAHRIGEAGEGDDGAGTMEEDFL

AWKDPMWTALAEKMGLEEREAVYEPIFSITERDGLTPESPEVYLGEPNKMHLEGAAKGPFNSHNPYIAPI

AESRELFNVKDRNCLHVEVDVSGSNLSYQTGDHIAIWPTNPGHEVDLFLDVIGLKDKRHTVVSIKALEPT

AKVPFPTPTTYDAIVRYHLEIGAPVSRQFVSTLAAFAPNDTARAEMTRLGGDKDYFHEKTGAHYFNISRF

LSSVSGGAPWANIPFSAWIEGITKLQPRYYSISSSSLVQPKKISITAVVERQAIPGREEDPFRGVATNYL

YALKQKQNGDPNPEAFGQTYEITGPRNKYDGIHVPVHVRHSNFKLPSDPSRPVIMVGPGTGVAPFRAFIQ

ERAKQAEDGATVGPTLLFFGCRKSTEDFVYKEEFESYKKSLGDSFELITAFSREGPKKVYVQHRLRERAQ

QVNDLLQKKAYFYVCGDAANMAREVNTFLGQIIAEQRGVSEAKAEEIVKGMRAANQYQEDVWS

>EXM25379.1 NADPH-cytochrome P450 reductase [Fusarium oxysporum f. sp. vasinfectum 25433]

MAELDTLDIVVLGVIFLGTVAYFTKGKLWGVTKDPYANGFAAGGASKPGRTRNIVEAMEESGKNCVVFYG

SQTGTAEDYASRLAKEGKSRFGLNTMIADLEDYDFDNLDTVPSDNIVMFVLATYGEGEPTDNAVDFYEFI

TGEDASFNEGNDPPLGNLNYVAFGLGNNTYEHYNSMVRNVNKALEKLGAHRIGEAGEGDDGAGTMEEDFL

AWKDPMWEALAKKMGLEEREAVYEPIFAINERDDLTPEANEVYLGEPNKLHLEGTAKGPFNSHNPYIAPI

AESYELFSAKDRNCLHMEIDISGSNLKYETGDHIAIWPTNPGEEVNKFLDILDLSGKQHSVVTVKALEPT

AKVPFPNPTTYDAILRYHLEICAPVSRQFVSTLAAFAPNDDIKAEMNRLGSDKDYFHEKTGPHYYNIARF

LASVSKGEKWTKIPFSAFIEGLTKLQPRYYSISSSSLVQPKKISITAVVESQQIPGRDDPFRGVATNYLF

ALKQKQNGDPNPAPFGQSYELTGPRNKYDGIHVPVHVRHSNFKLPSDPGKPIIMIGPGTGVAPFRGFVQE

RAKQARDGVEVGKTLLFFGCRKSTEDFMYQKEWQVRQHRTQSFEACTNCFSGIQGGSWR

>XP_001392901.1 NADPH--cytochrome P450 reductase [Aspergillus niger CBS 513.88]

MAQLDTLDLVVLAVLLVGSVAYFTKGTYWAVAKDPYASTGPAMNGAAKAGKTRNIIEKMEETGKNCVIFY

GSQTGTAEDYASRLAKEGSQRFGLKTMVADLEEYDYENLDQFPEDKVAFFVLATYGEGEPTDNAVEFYQF

FTGDDVAFESGASADEKPLSKLKYVAFGLGNNTYEHYNAMVRQVDAAFQKLGAQRIGSAGEGDDGAGTME

EDFLAWKEPMWAALSESMDLQEREAVYEPVFCVTENESLSPEDETVYLGEPTQSHLQGTPKGPYSAHNPF

IAPIAESRELFTVKDRNCLHMEISIAGSNLSYQTGDHIAVWPTNAGAEVDRFLQVFGLEGKRDSVINIKG

IDVTAKVPIPTPTTYDAAVRYYMEVCAPVSRQFVATLAAFAPDEESKAEIVRLGSDKDYFHEKVTNQCFN

IAQALQSITSKPFSAVPFSLLIEGITKLQPRYYSISSSSLVQKDKISITAVVESVRLPGASHMVKGVTTN

YLLALKQKQNGDPSPDPHGLTYSITGPRNKYDGIHVPVHVRHSNFKLPSDPSRPIIMVGPGTGVAPFRGF

IQERAALAAKGEKVGPTVLFFGCRKSDEDFLYKDEWKTYQDQLGDNLKIITAFSREGPQKVYVQHRLREH

SELVSDLLKQKATFYVCGDAANMAREVNLVLGQIIAAQRGLPAEKGEEMVKHMRSSGSYQEDVWS

>CCU75332.1 putative NADPH-cytochrome P450 reductase [Blumeria graminis f. sp. hordei DH14]

MAQLDTLDIVVLVGVLIASVAYFTKGTYWGKTEDPFAKYQLNSDKNRALGKTRNIVEKMADTGKNCVVFY

GSQTGTAEDYAARLAKEGKSRYGLETMVADLEDYDFETLDTLPDDKVAMFVVATYGEGEPTDNAVDFYQF

INSDPPEFSLGQDPPLQNLTFVAFGLGNNTYEHYNSMVRNLTSTLEKLGAHKVGAAGEGDDGAGTMEEDF

IAWKDPMWAALAEKMNLEEREAVYEPVFSIKEAENLTNDSPDVYLGEPNRLHLQGQAKGPFNSHNPFIAP

VIASRELFTNTDRNCLHLEIDLSDSGLSYQTGDHVAVWPTNSSQEVDRFLKITGLYSKRNTVVHVKPLDS

TAKIQFPTPTTYDAIARYYMEINAAVSRQFVASLAPFAPTESAKMEMIKLGNNKDYFQSKVGSQYLNVAQ

VLENVSTGVEWKSIPFSIIIEGMNKIQPRYYSISSSSLAEEKKISITATVESKSIPGRSDTLKGVTTNYL

LALKQKQNEEQDSALNGLLYNIDGPRKRYEGIRLPIHLRHSNFKLPSDPSKPIIMVGPGTGVAPFRAFVQ

ERATQAESGTKIGRMILFFGCRKESEDFLYKEEWKAHQKSLGDNFMLVTSFSRDGPKKIYVQDKMQEYSK

EINDLITKKAYFYVCGDAANMAKAVLNALTRVISEQRSIPESNAEEIVKRMRNSSQYQEDVWS

basidio

>TFK40696.1 hypothetical protein BDQ12DRAFT_697373 [Crucibulum laeve]

MASTSSSDVAILALGVILAALYLFRDQLFAASKPKVAPIVASKSANGSGNPRDFIAKMKDGKKRLVIFYG

SQTGTAEEYAIRLAKEAKSKFGLASLVCDPEEYDFENLDQLPEDCAAFFVMATYGEGEPTDNAVQLMQNL

QDDSFEFSNGERKLDGLKYVVFGLGNKTYEHYNSIGRAVDDQLTKMGATRIGERGEGDDDKSMEEDYLEW

KDGMWEAFATAMGVEEGQGGDTADFAVSELESHPAEKVYLGEYSARALTKTKGIHDAKNPFPAPISHARE

LFEDTKDRNCVHIELNTEGSGITYQHGDHVGVWPSNPDVEVNRLLCALGLTGKKDTVIGIESLDPALAKV

PFPVPTTYGTVLRHYIDISAVAGRQILGTLSKFAPSPEAEAFMKNLNTNKEEYHRVIAGGCLKLGEVLQL

AAGNDIRAVPNSENTTAWPIPFDIIVSAIPRLQPRYYSISSSPKLHPNSIHVTVVVLKYQSIPSEHVKEK

WVYGVGSNFLLNLKNAANGETVPLISDAGEERVAMPSYEIEGPRGAYKLDTVYKAPIHVRRSTFRLPTNP

KSPVIMIGPGTGVAPFRGFVQERVALARRSIEKNGADALADWGRISLFYGCRRSTEDFLYKDEWSKYAEE

LHGKFSMHCAFSREKYKPDGSKIYVQDLIWDDREHIADAILNGKGYVYICGEAKNMSKQVEDVLARILGE

AKGGSGDVEGQAEVKLLKERSRLMLDVWS

>XP_006462636.1 hypothetical protein AGABI2DRAFT_193746 [Agaricus bisporus var. bisporus H97]

MSSTSTSDVVILALGVTLAAAYLFRDQIFSSKPKSAPVAAHSKIANGGGNPRDFIAKMKEGKKRLVIFYG

SQTGTAEEYAIRLAKEAKSKFGLASLVCDPEEYDFENLDQLPEDCVAFFVMATYGEGEPTDNAVQLMQNI

EDDSFEFSNGSHRLDGLKYVVFSLGNRTYEHYNFIGRNVDAALTKMGAQRIGERGEGDDDKSMEEDYLEW

KDGMWEAFATALGVEEGQGADSADFAVSELDSHPPEKVYLGEYSARALTKTKGIHDSKNPYPAPLCETKE

LFQSVNDRNCVHAELSIEGSGITYQHGDHLGVWPTNAELEVDRLLCALGLYNKRDTVIGIESLDPALAKV

PFPVPTTYGTVLRHYIDISAVAGRQILGPLSKYAPNPEAEALMKSLNTNKEQYLKVVSEGCLRLGEVLQL

AAGNDSSVVPTPENTTAWSIPFDLIVSAIPRLQPRYYSISSSPKLHPNSVHVTCVVLKYQNIPSDTVRQK

WIYGVSSNFLLNLRLATTNEAAPFMNDNGAQSTSFPKYHIEGPRGAYKIDNMFKAPVHVRRSTFRLPTNP

KSPVIMVGPGTGVAPFRGFVQERVALARRSIEKNGPDALQDWGRISLFYGCRREDEDFLYKDEWPLYKEE

LKGKFEMYCAFSRQNYKPDGSKIYVQDLILEQREHIADAILNGKGYVYICGEAKNMSKQVEDVLAIILGE

AKGGSAAVEGVAEVKLLKERSRLMLDVWS

>XP_007319380.1 hypothetical protein SERLADRAFT_361923 [Serpula lacrymans var. lacrymans S7.9]

MAVASSSDVVILALGVTLAAAYLFRDQIFAPSKPKIVTTAPSKSENGSGNPRDFVAKMKDGKKRLVIFYG

SQTGTAEEYAIRLAKEAKTKFGITSLVCDPEEYDFENLDEVPEDCAVFFVVATYGEGEPTDNAVQLTQNL

TDDSFTFGNGEHKLPGLKYVIFSLGNKTYEHYNAIGRSYDAELTKMGAVRIGERGEGDDDKSMEEDYLEW

KDAMWEAFATAMNVEEGQGGDTADFTVTELDSHPQEKVYLGELSARALTKTKGIHDAKNPYASPVRNSRE

LFQLTGDRNCIHVELDIEGSGITYQHGDHVGVWPSNPDAEVDRLLCVLGLYDKKDSVIGIESLDPALAKV

PFPVPTTYATVLRHYIDISAVVGRQILGAMSKFAPSPEAEAFLKNLNVDKEEYAAVIGQGCFKLGEILQI

AAANDIRVPPTKDNTTPWPIPFDIIVSSIPRLQPRYYSISSSPKLHPTSIHVTAVVLKYQSATNKATQAR

WVYGVGTNFILNVKFAANGEAAPLMSGTSNTDPATVSMPSYAIQGPRGAHKQETIYKIPIHVRRSTFRLP

TNPKSPVIMVGPGTGVAPFRGFVQERVALARRTLDKNGPEALADWGKISLFYGCRKSTEDFLYKDEWPTY

TEELRGKFTMHCAFSREPPYKPDGSKIYVQDLIWQDRENIAEAILNGKGYVYICGDAKAMSKAVEEVLSR

ILGEAKGGSAEVEGAAEMKLLKERSRLMLDVWS

>KDQ25324.1 hypothetical protein PLEOSDRAFT_1057641 [Pleurotus ostreatus PC15]

MASTSSSDVLVLAVGVALAAIYLFRDQLFAQAKPKSVPVPTSKASNGSGNPRDFIAKMKEGKKRLVIFYG

SQTGTAEEYAIRLAKEAKSKFGLASLVCDPEEYDFENLDQLPEDCAVFFVMATYGEGEPTDNAVTLMQNL

EDESFEFSNGEHKLEGLKYVVFSLGNKTYEHYNKIGRDVDNVLTKMGAIRIGERGEGDDDKSMEEDYLEW

KDGMWDAFSTAMGVEEGQGGDTPDFAVTELESHPPEKVYLGELSARALTKTKGIHDAKNPFPAPISVARE

LFQSTHDRNCVHIELNTESSGISYQHGDHVGVWPSNPDVEVTRLLCALGLYEKKDNVIGIESLDPALAKV

PFPVPTTYATVLRHYIDISAVAGRQILGALSKFAPNPEAEAFLKGLSTNKEEYHTLIANGCLKLGEVLQL

AAGNDLSAAPTPENTTAWTIPFDIIVSSIPRLQPRYYSISSSPKLHPNSIHVTAVVLKYESIPNERVNGR

WIFGVGSNFLLNLKYAANGETAPLVATGSESKAASVTIPGYAIEGPRGAYKQETIYKAPIHVRRSTFRLP

TNPKSPVIMIGPGTGVAPFRGFIQERVALARRSIEKNGPDALADWGRISLFYGCRKSTEDFLYKDEWPQY

QEELRGKFTMHCAFSREVYRPDGSKIYVQDLLWDDREQVADAIINGKGYIYICGDAKSMSKAVEETLAKI

LGEAKGGSADVEGNAEVKLLKERSRLMLDVWS

Animals

>CAA63639.1 NADPH--ferrihemoprotein reductase [Drosophila melanogaster]2E-143

MASEQTIDGAAAIPSGGGDEPFLGLLDVALLAVLIGGVTFYFLRTRKKEEEPTRSYSIQPTTVCTTSASD

NSFIKKLKASGRSLVVFYGSQTGTGEEFAGRLAKEGIRYRLKGMVADPEECDMEELLQLKDTDNSLAVFC

LATYGEGDPTDNAMEFYEWITSGDVDLSGLNYAVFGLGNKTYEHYNKVAIYVDKRLEELGANRVFELGLG

DDDANIEDDFITWKDRFWPAVCDHFGIEGGGEEVLIRQYRLLEQPDVQPDRIYTGEIARLHSIQNQRPPF

DAKNPFLAPIKVNRELHKGGGRSCMHIELSIEGSKMRYDAGDHVAMFPVNDKSLVEKLGQLCNADLDTVF

SLINTDTDSSKKHPFPCPTTYRTALTHYLEITAIPRTHILKELAEYCTDEKEKELLRSMASISPEGKEKY

QSWIQDACRNIVHILEDIKSCRPPIDHVCELLPRLQPRYYSISSSAKLHPTDVHVTAVLVEYKTPTGRIN

KGVATTYLKNKQPQGSEEVKVPVFIRKSQFRLPTKPETPIIMVGPGTGLAPFRGFIQERQFLRDEGKTVG

ESILYFGCRKRSEDYIYESELEEWVKKGTLNLKAAFSRDQGKKVYVQHLLEQDADLIWNVIGENKGHFYI

CGDAKNMAVDVRNILVKILSTKGNMSEADAVQYIKKMEAQKRYSADVWS

>NP_001273818.1 NADPH--cytochrome P450 reductase [Musca domestica] 6E-142

MSAEHVEEVVSEEPFLGTLDIALLVVLLVGATWYFMRSRKKEEAPIRSYSIQPTTVSTVSTTENSFIKKL

KASGRSLVVFYGSQTGTAEEFAGRLAKEGLRYRMKGMVADPEECDMEELLQMKDIPNSLAVFCLATYGEG

DPTDNAMEFYEWITNGEVDLTGLNYAVFGLGNKTYEHYNKVAIYVDKRLEELGATRVFELGLGDDDANIE

DDFITWKDRFWPSVCDFFGIEGSGEEVLMRQFRLLEQPDVQPDRIYTGEIARLHSMQNQRPPFDAKNPFL

ASVIVNRELHKGGGRSCMHIELDIDGSKMRYDAGDHIAMYPINDKILVEKLGKLCDANLDTVFSLINTDT

DSSKKHPFPCPTTYRTALTHYLEITAIPRTHILKELAEYCSDEKDKEFLRNMASITPEGKEKYQNWIQNS

SRNIVHILEDIKSCRPPIDHICELLPRLQPRYYSISSSSKLYPTNVHITAVLVQYETPTGRVNKGVATSY

MKEKNPSVGEVKVPVFIRKSQFRLPTKSEIPIIMVGPGTGLAPFRGFIQERQFLRDGGKVVGDTILYFGC

RKKDEDFIYREELEQYVQNGTLTLKTAFSRDQQEKIYVTHLIEQDADLIWKVIGEQKGHFYICGDAKNMA

VDVRNILVKILSTKGNMNESDAVQYIKKMEAQKRYSADVWS

>AAH34277.1 P450 (cytochrome) oxidoreductase [Homo sapiens]2E-139

MINMGDSHVDTSSTVSEAVAEEVSLFSMTDMILFSLIVGLLTYWFLFRKKKEEVPEFTKIQTLTSSVRES

SFVEKMKKTGRNIIVFYGSQTGTAEEFANRLSKDAHRYGMRGMSADPEEYDLADLSSLPEIDNALVVFCM

ATYGEGDPTDNAQDFYDWLQETDVDLSGVKFAVFGLGNKTYEHFNAMGKYVDKRLEQLGAQRIFELGLGD

DDGNLEEDFITWREQFWLAVCEHFGVEATGEESSIRQYELVVHTDIDAAKVYMGEMGRLKSYENQKPPFD

AKNPFLAAVTTNRKLNQGTERHLMHLELDISDSKIRYESGDHVAVYPANDSALVNQLGKILGADLDVVMS

LNNLDEESNKKHPFPCPTSYRTALTYYLDITNPPRTNVLYELAQYASEPSEQELLRKMASSSGEGKELYL

SWVVEARRHILAILQDCPSLRPPIDHLCELLPRLQARYYSIASSSKVHPNSVHICAVVVEYETKAGRINK

GVATNWLRAKEPVGENGGRALVPMFVRKSQFRLPFKATTPVIMVGPGTGVAPFIGFIQERAWLRQQGKEV

GETLLYYGCRRSDEDYLYREELAQFHRDGALTQLNVAFSREQSHKVYVQHLLKQDREHLWKLIEGGAHIY

VCGDARNMARDVQNTFYDIVAELGAMEHAQAVDYIKKLMTKGRYSLDVWS

>NP_032924.1 NADPH--cytochrome P450 reductase [Mus musculus] Similar to the uman gene above and to Ph1

MGDSHEDTSATVPEAVAEEVSLFSTTDIVLFSLIVGVLTYWFIFKKKKEEIPEFSKIQTTAPPVKESSFV

EKMKKTGRNIIVFYGSQTGTAEEFANRLSKDAHRYGMRGMSADPEEYDLADLSSLPEIDKSLVVFCMATY

GEGDPTDNAQDFYDWLQETDVDLTGVKFAVFGLGNKTYEHFNAMGKYVDQRLEQLGAQRIFELGLGDDDG

NLEEDFITWREQFWPAVCEFFGVEATGEESSIRQYELVVHEDMDTAKVYTGEMGRLKSYENQKPPFDAKN

PFLAAVTTNRKLNQGTERHLMHLELDISDSKIRYESGDHVAVYPANDSTLVNQIGEILGADLDVIMSLNN

LDEESNKKHPFPCPTTYRTALTYYLDITNPPRTNVLYELAQYASEPSEQEHLHKMASSSGEGKELYLSWV

VEARRHILAILQDYPSLRPPIDHLCELLPRLQARYYSIASSSKVHPNSVHICAVAVEYEAKSGRVNKGVA

TSWLRTKEPAGENGRRALVPMFVRKSQFRLPFKPTTPVIMVGPGTGVAPFMGFIQERAWLREQGKEVGET

LLYYGCRRSDEDYLYREELARFHKDGALTQLNVAFSREQAHKVYVQHLLKRDKEHLWKLIHEGGAHIYVC

GDARNMAKDVQNTFYDIVAEFGPMEHTQAVDYVKKLMTKGRYSLDVWS

>NP_001182725.1 NADPH--cytochrome P450 reductase [Gallus gallus]6E-142

MGDAGMESTVSPPEGTAQDSFLSMTDVFLISLITGLFTYWFFFRKKKEEIPDLPKIQTVSSPARDSSFIE

KMKKTGRNIVVFYGSQTGTAEEFANRLSKDAHRYGLRGMAADPEEYDLSDLSRLSEIDKSLAVFCMATYG

EGDPTDNAQDFYDWLQEADTDLSGLRFAVFGLGNKTYEHFNAMGKYVDKRLEELGAQRIFELGLGDDDGN

LEEDFITWREQFWPAVCEHFGVEATGEESSIRQYELVVHTDVNMNKVYTGEMGRLKSYENQKPPFDAKNP

FLAVVTENRKLNEGGERHLMHLELDISNSKIRYESGDHVAVYPANDASLVNQLGEILGTDLDTVMSLNNL

DEESNKKHPFPCPTSYRTALTYYLDITNPPRTNVLYELAQYATDTGEQEQLRKMASSSAEGKALYLSWVV

EARRNILAILQDMPSLRPPIDHLCELLPRLQARYYSIASSSKVHPNSIHICAVTVEYETKTGRLNKGVAT

NWLKDKVPNENGRNSLVPMYVRKSQFRLPFKPSTPVIMIGPGTGIAPFIGFIQERAWLKEQGKEVGETVL

YYGCRREREDYLYRQELARFKQEGVLTQLNVAFSRDQAEKVYVQHLLKKNKEHIWKLVNDGNAHIYVCGD

ARNMARDVQNTFYEIVSEYGNMNQSQAVDYVKKLMTKGRYSLDVWS

Planta

>NP_001190823.1 P450 reductase 1 [Arabidopsis thaliana] 6E-112

MTSALYASDLFKQLKSIMGTDSLSDDVVLVIATTSLALVAGFVVLLWKKTTADRSGELKPLMIPKSLMAK

DEDDDLDLGSGKTRVSIFFGTQTGTAEGFAKALSEEIKARYEKAAVKDDYAADDDQYEEKLKKETLAFFC

VATYGDGEPTDNAARFYKWFTEENERDIKLQQLAYGVFALGNRQYEHFNKIGIVLDEELCKKGAKRLIEV

GLGDDDQSIEDDFNAWKESLWSELDKLLKDEDDKSVATPYTAVIPEYRVVTHDPRFTTQKSMESNVANGN

TTIDIHHPCRVDVAVQKELHTHESDRSCIHLEFDISRTGITYETGDHVGVYAENHVEIVEEAGKLLGHSL

DLVFSIHADKEDGSPLESAVPPPFPGPCTLGTGLARYADLLNPPRKSALVALAAYATEPSEAEKLKHLTS

PDGKDEYSQWIVASQRSLLEVMAAFPSAKPPLGVFFAAIAPRLQPRYYSISSSPRLAPSRVHVTSALVYG

PTPTGRIHKGVCSTWMKNAVPAEKSHECSGAPIFIRASNFKLPSNPSTPIVMVGPGTGLAPFRGFLQERM

ALKEDGEELGSSLLFFGCRNRQMDFIYEDELNNFVDQGVISELIMAFSREGAQKEYVQHKMMEKAAQVWD

LIKEEGYLYVCGDAKGMARDVHRTLHTIVQEQEGVSSSEAEAIVKKLQTEGRYLRDVW

>pdb|5GXU|A Chain A, Nadph--cytochrome P450 Reductase 2 5E-112

GRRSGSGNSKRVEPLKPLVIKPREEEIDDGRKKVTIFFGTQTGTAEGFAKALGEEAKARYEKTRFKIVDL

DDYAADDDEYEEKLKKEDVAFFFLATYGDGEPTDNAARFYKWFTEGNDRGEWLKNLKYGVFGLGNRQYEH

FNKVAKVVDDILVEQGAQRLVQVGLGDDDQCIEDDFTAWREALWPELDTILREEGDTAVATPYTAAVLEY

RVSIHDSEDAKFNDINMANGNGYTVFDAQHPYKANVAVKRELHTPESDRSCIHLEFDIAGSGLTYETGDH

VGVLCDNLSETVDEALRLLDMSPDTYFSLHAEKEDGTPISSSLPPPFPPCNLRTALTRYACLLSSPKKSA

LVALAAHASDPTEAERLKHLASPAGKDEYSKWVVESQRSLLEVMAEFPSAKPPLGVFFAGVAPRLQPRFY

SISSSPKIAETRIHVTCALVYEKMPTGRIHKGVCSTWMKNAVPYEKSENCSSAPIFVRQSNFKLPSDSKV

PIIMIGPGTGLAPFRGFLQERLALVESGVELGPSVLFFGCRNRRMDFIYEEELQRFVESGALAELSVAFS

REGPTKEYVQHKMMDKASDIWNMISQGAYLYVCGDAKGMARDVHRSLHTIAQEQGSMDSTKAEGFVKNLQ

TSGRYLRDVW

>CAC83301.1 cytochrome P450 reductase [Triticum aestivum] 3E-104

MDSAAAGMRDSALDLLAALLTGRAPPAAADGDQNRRLLALLATSLAVLVGCGVALLFRRSSSGAAPLAHK

SAAAKPLAAKKDQEPDPDDGRQRVALFFGTQTGTAEGFAKALAEEAKARYDKAVFKVLDLDDYAAEDEEY

EEKLKKENIAFFFLATYGDGEPTDNAARFYKWFSEGNERGEWLSNLKFGVFALGNRQYEHFNKVGKEVDQ

LLAEQGGKRIVPVGLGDDDQCIEDDFNAWKELLWPELDKLLRVEDNSSTAQSPYTAAIPQYRVVLTKPED

ATHINKSFSLSNGHVVYDSQHPCRANVAVRRELHTPASDRSCIHLEFDIAGTSLTYETGDHVGVYAENSI

ETVEEAEKLLDYSPDTYFSIYADQEDGTPLFGGSLPPPFPSPCTVRVALARYADLLNSPKKSVLLALAAH

ASDPKEAERLRHLASPAGKKEYSQWIIASQRSLLEVISEFPSAKPPLGVFFAAIAPRLQPRYYSISSSPR

MAPTRIHVTCSLVHGQTPTGRIHKGVCSTWMKNSTPLEESQECSWAPIFVRQSNFKLPADPTVPIIMVGP

GTGLAPFRGFLQERLALKETGVELGRAILFFGCRNRQMDFIYEDELNNFAESGALSELVVAFSREGPTKE

YVQHKMAEKAAELWSIVSQGGYVYVCGDAKGMARDVHRALHTIVQEQGSLDSSKAEGYVKNLQMEGRYLR

DVW

Protozoa

>XP_828830.1 NADPH--cytochrome P450 reductase [Trypanosoma brucei brucei TREU927]5E-90

MIFLVISTAIIAVLAWFVAGVFIRGGGGRGKTAAPQVVGVAQYPSQPSSRVDVRVLFGSQTGTAEMFAKT

VTREGLRLGVPMKLADVENYRPSDLAGEKYVIIICATYGEGEPTDTMVGFHEWLVDDSRAVGEELSGVKY

TVFALGDRQYKFFCREGITVDRRMSELGAQRFYPLGYGDCGNSIEEEFDNWCHNLWPVLGRALSLVLKSN

STEPVAPECRMKLWGPPEEAPLPFPKLASVLEPTQRLPSWAPVKVNKELLSNATGRSTRLIEFDTSETVI

SYQAGDHLGVLPSNPSEMVNTYLRVLGVSEQESSQVISLQNRATGKNVFPCRVSIRTALTWYIDLAGPPK

KSTLRAFAHHCTDPVEKDTLLKLLSTEPESVEAYGKLVLELRTVLGFLQRFKSMSPPLSFFLEMMPRIAP

RYFSISSDSLTHPTSVAITVAVVEGGLCTNLLQQAAVGQNIPVFVRKSNFHLPLRAKDRPIIMIGPGTGV

APFIGFLHRRSAWLEKGNKVGDALLFFGCRRREEDHIYADFMEKCLSNGALSVRDVAYSREQADKVYVQH

RLAARGKEVWEIISRGGNVYVCGDAKNMARDVERQLLDIAQKYGAMKEDEATALLEKLATDERYLKDVWT

A

>XP_646400.1 NADPH-cytochrome-P450 oxidoreductase [Dictyostelium discoideum AX4]1E-107

MEILESIDFIEVLILDNLGAIIIVAVIVGTYLYMNKPPPPPPVFNKPNNKINKEAQKPKKTITKNEDGKK

VMKIFFGTQTRTAEDFSRIIEKECKKIGIPCEVVDLESYEHEQELHSESFVMFLVATHGEGDPTDNAKEF

YLWLTNDERPTDLLNGVPFTVFGLGNKTYEHYNAVARVIDRRMEELGGKRVFERGEGDDDATLEEDFNRW

KKDMWPVVCKFLGYELKSTEDDKFVPRFRMVTLNQDSKDINDPFIKIVSTPLKPKLSTDNKVIYDMKNPY

YAEVLENRELHSNESDRSCRHIEFKLGDEVSYTTGDHLGVFPINDSKLVEQLIKRLGVNGDDMIALVPID

QEGSVIKASFGPMTIRRAFSEHLDITNPVRKSVLRALAESTTNEEEKKRLLYLATEEANEEYNKYIKNDF

RGVVDLLESFPGLQPLIAHFLEFTPRLPARMYSISSSPHNKNGVVSITSVVVNFTTGNQRAHNGVASTWL

SHLKVGDKVPLFVRESHFKLPSAATEQKPVIMVGPGTGLAPFRGFLQELQHRNHSQQQQSLLFFGCRSDT

VDYIYREELEQYHQSSVLGDLVVAFSRKTSQKVYVQNKLLEHKEKVWELLNKGAYFYVCGDGRNMSKAVQ

QALLSIIKEFGSKDDNSAQQFIDDMSSHGRYLQDVWF

https://ngphylogeny.fr/workflows/oneclick/ NGPhylogeny.fr

> FGSG_09786

MAELDTLDVIVLGVIFLGTVAYFTKGKLWGVTKDPYANGFAAGGAAKPGRTRNIVEAMEESGKNCVIFYG

SQTGTAEDYASRLAKEGKSRFGLNTMIADIEDYDFDSLDTVPNDNVVMFVLATYGEGEPTDNAVDFYEFI

TGEDATFNEGNDPPLGNLNYVAFGLGNNTYEHYNAMVRKVDQALEKFGAHRIGEAGEGDDGAGTMEEDFL

AWKDPMWESLAKKMGLEEREAVYEPIFAINERDDLSPESNEVYLGEPNKLHLEGTAKGPFNSHNPYIAPI

AESYELFSAKDRNCLHMEVDISGSNLKYETGDHIAIWPTNPGEEVNRFLDILDLSGKQHSVITVKALEPT

AKVPFPNPTTYDAILRYHLEICAPVSRQFVSTLAAFAPNDSIKAEMNRLGSDKDYFHEKTGPHYYNIARF

LSSVSKGEKWTTIPFSAFIEGLTKLQPRYYSISSSSLVQPKKISITAVVESQQIPGRDDPFRGVATNYLF

ALKQKQNGDPSPAPFGQTYELTGPRNKYDGIHVPVHVRHSNFKLPSDPGKPVIMIGPGTGVAPFRGFVQE

RAKLARDGVEVGKTLLFFGCRKPSEDFMYEKEWQEYKEALGDKFEMITAFSRESAKKVYVQHRLKERAQE

VSDLLSQKAYFYVCGDASNMAREVNTVLAQIIAEGRGVSEAKGEEIVKNMRSANQYQEDVWS

>FGSG_08413

MTNVFAADTLLSQLGPPQSIADIAALSALGVASAAYLLRGITWDKPDHYHHVWFERMGSKGGSSSSHPKA

TRDIAKKLEETGKDVVIFWGSQSGTAETFANRLSKECHLRFGLQALCADLCDYDPESIANLSQSKLAIFI

LSTYGEGDPSDNTAAFWDWLTKTPNIQIPNLRYMAFGLGNTSYRYYNRVIDVVAQHLDKYGAQRLMPVGR

ANDAQGGTEEDFLSWKDDLYTHFQENLGYQERDIPYEPSIQLIQDESLDIMDLHLGEPIQNRNGPAKVVK

QYSPIRPLAIQSSQELYTSPGRNCLHMELDISNQPELRYRTGDHLAIYPINPDYEVQLLLKAFGLEDRAE

KPLLVQTLEEGTSTKIPSPTSALALFRHYLEVAAPVSRETVGQLARFAPLSESVETLTALAKNKEAYATY

IGSNHITMGRLLHLVAPGAIWTKLPLSYVVETLPCIQARYYSISSSSTVSARRLSITVGVDKSPLQQDPS

RVIRGITTNYLYALGNALNCDTSQSVVGSDAPSYALSGPGDTLNGRKVFACIRRSNFKLPTLSSTPIIMI

GAGTGLAPFRGFILERARLQAVGKPIGKMLLFFGCRSPDQDYLYQGELAEVAQKLQGSLEIVTAFSRAEE

EPKKYVQDRVEERKTQVCHLLQEGASIYFCGRAAMARKVGNKVEESMKTQNNWTDAEARSWAESVKKGNK

WLEDVWG

>YeastNCP1

MPFGIDNTDFTVLAGLVLAVLLYVKRNSIKELLMSDDGDITAVSSGNRDIAQVVTENNKNYLVLYASQTGTAEDYAKKFSKELVAKFNLNVMCADVENYDFESLNDVPVIVSIFISTYGEGDFPDGAVNFEDFICNAEAGALSNLRYNMFGLGNSTYEFFNGAAKKAEKHLSAAGAIRLGKLGEADDGAGTTDEDYMAWKDSILEVLKDELHLDEQEAKFTSQFQYTVLNEITDSMSLGEPSAHYLPSHQLNRNADGIQLGPFDLSQPYIAPIVKSRELFSSNDRNCIHSEFDLSGSNIKYSTGDHLAVWPSNPLEKVEQFLSIFNLDPETIFDLKPLDPTVKVPFPTPTTIGAAIKHYLEITGPVSRQLFSSLIQFAPNADVKEKLTLLSKDKDQFAVEITSKYFNIADALKYLSDGAKWDTVPMQFLVESVPQMTPRYYSISSSSLSEKQTVHVTSIVENFPNPELPDAPPVVGVTTNLLRNIQLAQNNVNIAETNLPVHYDLNGPRKLFANYKLPVHVRRSNFRLPSNPSTPVIMIGPGTGVAPFRGFIRERVAFLESQKKGGNNVSLGKHILFYGSRNTDDFLYQDEWPEYAKKLDGSFEMVVAHSRLPNTKKVYVQDKLKDYEDQVFEMINNGAFIYVCGDAKGMAKGVSTALVGILSRGKSITTDEATELIKMLKTSGRYQEDVW

>Pyricularia oryzae 70-15]

MAELDTLDIIVLGLILCGTIAYFTKGKYWGVVKDPYAASFANTNGPKTGKTRNIVEKMEETGKNCVIFYG

SQTGTAEDYASRLAKEGKSRFGLETMVADLEDYDYENLDTVPSDKIVMFVLATYGEGEPTDNAVDFYEFI

TGEDVSFSEGSTLDNLNYVAFGLGNNTYEHYNSMVRNVNKALEKLGAHRIGDAGEGDDGAGTMEEDFLAW

KDPMWAALAEKMGLEEREAVYEPVFSVTEREGLTVESPEVYLGEPNKMHLEGTAKGPFNAHNPYIAPIVK

SYELFNVKDRNCLHIDVDVSGSNLTYQTGDHIAVWPTNPGEEVDCLLDVLGLTDKRDTVVSVRPLEPTAK

VPFPAPTTYDAILRYHMEICAPVSRQFIATLAAFAPDEETKAEMTKLGGDKDYFSAKISKHYLNIARVLF

NVGKGKKWNNIPFSAFIEGLTKLQPRYYSISSSSLVQPKVITITAVVEKQEIPGRDDPFRGVTTNYLLAL

KQKQNGEPHPEPFGRTFALAGPRDKYDGIKVPVHVRHSNFKLPSDSTKPIILVGPGTGVAPMRAFVQERA

KQAENGEEVGKTILFFGCRKSTEDFLYKDEWDEYKKVLGDKFELVTAFSREGPKKVYVQHRLKERAQEIN

ELLTKKAYIYVCGDAANMAREVNSVLGQIIAEQRGIPEAKAEEIVKNMRAANQYQEDVWS

>Metarhizium anisopliae

MAEFDTLDIVVLGVILLGTIAYFTKGTLWGVTKDPYANAFANANGAKAGRSRNILEKMDETGKNCVIFYG

SQTGTAEDYASRLAKEGKSRFGLETMVADLEEYDFDNLDAMPSDKVAMFILATYGEGEPTDNAVEFYEFI

TGDDVSFSEGSDPALQNLNYVAFGLGNNTYEHYNSMVRNVDKALQKLGANRIGEAGEGDDGAGTMEEDFL

AWKDPMWAALAEKMGLEEREAVYEPTFGIVDRENLTVDSPEVYLGEPNKMHLEGTAKGPFNSHNPYIAPI

AESRELFSAKDRNCIHMDVDINGSNLSYQTGDHIAIWPTNSGDEVDRFLDIIGLKEKRNNVISIKALEPT

AKVPFPTPTTYDAIVRYHLEICAPVSRQFVATLAAFAPNDEVKAEMARLGSDKEYFHSKTGPHFYNIARL

LDTVGKGEKWTNIPFSAFIEGLNKLQPRYYSISSSSLVQPKKISITAVIESQAIPGRKDPFRGVATNYLF

ALKQKQNGDPNPSPFGKTYALNGPRNKFDGIHVPVHVRHSNFKLPSDPAKPVIMVGPGTGVAPFRGFIQE

RAKQAQNGATVGRTILFFGCRKRSEDFLYESEWEEYKKALGDSLEIVTAFSRESSKKVYVQHRLKERSKE

IGELLSQKAYFYVCGDAAHMAREVNTVLAQIIAESRGVSETKGEEIVKNMRAANQYQEDVWS

>Verticillium dahliae

MAELDTLDIIVLSVILLGTAAYFTKGKYWAIEKDPYANGFASAGGPKAGKTRNIIEKLDESGKNCVIFYG

SQTGTAEDYASRLAKEGKSRFGLETMVADLEDYDFDNLDAVSTDKVVMFVLATYGEGEPTDNAVEFYEFI

TGEDVSFNEANEPALGNLNFVAFGLGNNTYEHYNSMVRNVTKALEKLGAHRIGEAGEGDDGAGTMEEDFL

AWKDPMWTALAEKMGLEEREAVYEPIFSITERDGLTPESPEVYLGEPNKMHLEGAAKGPFNSHNPYIAPI

AESRELFNVKDRNCLHVEVDVSGSNLSYQTGDHIAIWPTNPGHEVDLFLDVIGLKDKRHTVVSIKALEPT

AKVPFPTPTTYDAIVRYHLEIGAPVSRQFVSTLAAFAPNDTARAEMTRLGGDKDYFHEKTGAHYFNISRF

LSSVSGGAPWANIPFSAWIEGITKLQPRYYSISSSSLVQPKKISITAVVERQAIPGREEDPFRGVATNYL

YALKQKQNGDPNPEAFGQTYEITGPRNKYDGIHVPVHVRHSNFKLPSDPSRPVIMVGPGTGVAPFRAFIQ

ERAKQAEDGATVGPTLLFFGCRKSTEDFVYKEEFESYKKSLGDSFELITAFSREGPKKVYVQHRLRERAQ

QVNDLLQKKAYFYVCGDAANMAREVNTFLGQIIAEQRGVSEAKAEEIVKGMRAANQYQEDVWS

>Fusarium oxysporum f. sp. vasinfectum

MAELDTLDIVVLGVIFLGTVAYFTKGKLWGVTKDPYANGFAAGGASKPGRTRNIVEAMEESGKNCVVFYG

SQTGTAEDYASRLAKEGKSRFGLNTMIADLEDYDFDNLDTVPSDNIVMFVLATYGEGEPTDNAVDFYEFI

TGEDASFNEGNDPPLGNLNYVAFGLGNNTYEHYNSMVRNVNKALEKLGAHRIGEAGEGDDGAGTMEEDFL

AWKDPMWEALAKKMGLEEREAVYEPIFAINERDDLTPEANEVYLGEPNKLHLEGTAKGPFNSHNPYIAPI

AESYELFSAKDRNCLHMEIDISGSNLKYETGDHIAIWPTNPGEEVNKFLDILDLSGKQHSVVTVKALEPT

AKVPFPNPTTYDAILRYHLEICAPVSRQFVSTLAAFAPNDDIKAEMNRLGSDKDYFHEKTGPHYYNIARF

LASVSKGEKWTKIPFSAFIEGLTKLQPRYYSISSSSLVQPKKISITAVVESQQIPGRDDPFRGVATNYLF

ALKQKQNGDPNPAPFGQSYELTGPRNKYDGIHVPVHVRHSNFKLPSDPGKPIIMIGPGTGVAPFRGFVQE

RAKQARDGVEVGKTLLFFGCRKSTEDFMYQKEWQVRQHRTQSFEACTNCFSGIQGGSWR

>Aspergillus niger CBS

MAQLDTLDLVVLAVLLVGSVAYFTKGTYWAVAKDPYASTGPAMNGAAKAGKTRNIIEKMEETGKNCVIFY

GSQTGTAEDYASRLAKEGSQRFGLKTMVADLEEYDYENLDQFPEDKVAFFVLATYGEGEPTDNAVEFYQF

FTGDDVAFESGASADEKPLSKLKYVAFGLGNNTYEHYNAMVRQVDAAFQKLGAQRIGSAGEGDDGAGTME

EDFLAWKEPMWAALSESMDLQEREAVYEPVFCVTENESLSPEDETVYLGEPTQSHLQGTPKGPYSAHNPF

IAPIAESRELFTVKDRNCLHMEISIAGSNLSYQTGDHIAVWPTNAGAEVDRFLQVFGLEGKRDSVINIKG

IDVTAKVPIPTPTTYDAAVRYYMEVCAPVSRQFVATLAAFAPDEESKAEIVRLGSDKDYFHEKVTNQCFN

IAQALQSITSKPFSAVPFSLLIEGITKLQPRYYSISSSSLVQKDKISITAVVESVRLPGASHMVKGVTTN

YLLALKQKQNGDPSPDPHGLTYSITGPRNKYDGIHVPVHVRHSNFKLPSDPSRPIIMVGPGTGVAPFRGF

IQERAALAAKGEKVGPTVLFFGCRKSDEDFLYKDEWKTYQDQLGDNLKIITAFSREGPQKVYVQHRLREH

SELVSDLLKQKATFYVCGDAANMAREVNLVLGQIIAAQRGLPAEKGEEMVKHMRSSGSYQEDVWS

>Blumeria graminis

MAQLDTLDIVVLVGVLIASVAYFTKGTYWGKTEDPFAKYQLNSDKNRALGKTRNIVEKMADTGKNCVVFY

GSQTGTAEDYAARLAKEGKSRYGLETMVADLEDYDFETLDTLPDDKVAMFVVATYGEGEPTDNAVDFYQF

INSDPPEFSLGQDPPLQNLTFVAFGLGNNTYEHYNSMVRNLTSTLEKLGAHKVGAAGEGDDGAGTMEEDF

IAWKDPMWAALAEKMNLEEREAVYEPVFSIKEAENLTNDSPDVYLGEPNRLHLQGQAKGPFNSHNPFIAP

VIASRELFTNTDRNCLHLEIDLSDSGLSYQTGDHVAVWPTNSSQEVDRFLKITGLYSKRNTVVHVKPLDS

TAKIQFPTPTTYDAIARYYMEINAAVSRQFVASLAPFAPTESAKMEMIKLGNNKDYFQSKVGSQYLNVAQ

VLENVSTGVEWKSIPFSIIIEGMNKIQPRYYSISSSSLAEEKKISITATVESKSIPGRSDTLKGVTTNYL

LALKQKQNEEQDSALNGLLYNIDGPRKRYEGIRLPIHLRHSNFKLPSDPSKPIIMVGPGTGVAPFRAFVQ

ERATQAESGTKIGRMILFFGCRKESEDFLYKEEWKAHQKSLGDNFMLVTSFSRDGPKKIYVQDKMQEYSK

EINDLITKKAYFYVCGDAANMAKAVLNALTRVISEQRSIPESNAEEIVKRMRNSSQYQEDVWS

>Crucibulum laeve

MASTSSSDVAILALGVILAALYLFRDQLFAASKPKVAPIVASKSANGSGNPRDFIAKMKDGKKRLVIFYG

SQTGTAEEYAIRLAKEAKSKFGLASLVCDPEEYDFENLDQLPEDCAAFFVMATYGEGEPTDNAVQLMQNL

QDDSFEFSNGERKLDGLKYVVFGLGNKTYEHYNSIGRAVDDQLTKMGATRIGERGEGDDDKSMEEDYLEW

KDGMWEAFATAMGVEEGQGGDTADFAVSELESHPAEKVYLGEYSARALTKTKGIHDAKNPFPAPISHARE

LFEDTKDRNCVHIELNTEGSGITYQHGDHVGVWPSNPDVEVNRLLCALGLTGKKDTVIGIESLDPALAKV

PFPVPTTYGTVLRHYIDISAVAGRQILGTLSKFAPSPEAEAFMKNLNTNKEEYHRVIAGGCLKLGEVLQL

AAGNDIRAVPNSENTTAWPIPFDIIVSAIPRLQPRYYSISSSPKLHPNSIHVTVVVLKYQSIPSEHVKEK

WVYGVGSNFLLNLKNAANGETVPLISDAGEERVAMPSYEIEGPRGAYKLDTVYKAPIHVRRSTFRLPTNP

KSPVIMIGPGTGVAPFRGFVQERVALARRSIEKNGADALADWGRISLFYGCRRSTEDFLYKDEWSKYAEE

LHGKFSMHCAFSREKYKPDGSKIYVQDLIWDDREHIADAILNGKGYVYICGEAKNMSKQVEDVLARILGE

AKGGSGDVEGQAEVKLLKERSRLMLDVWS

>Agaricus bisporus

MSSTSTSDVVILALGVTLAAAYLFRDQIFSSKPKSAPVAAHSKIANGGGNPRDFIAKMKEGKKRLVIFYG

SQTGTAEEYAIRLAKEAKSKFGLASLVCDPEEYDFENLDQLPEDCVAFFVMATYGEGEPTDNAVQLMQNI

EDDSFEFSNGSHRLDGLKYVVFSLGNRTYEHYNFIGRNVDAALTKMGAQRIGERGEGDDDKSMEEDYLEW

KDGMWEAFATALGVEEGQGADSADFAVSELDSHPPEKVYLGEYSARALTKTKGIHDSKNPYPAPLCETKE

LFQSVNDRNCVHAELSIEGSGITYQHGDHLGVWPTNAELEVDRLLCALGLYNKRDTVIGIESLDPALAKV

PFPVPTTYGTVLRHYIDISAVAGRQILGPLSKYAPNPEAEALMKSLNTNKEQYLKVVSEGCLRLGEVLQL

AAGNDSSVVPTPENTTAWSIPFDLIVSAIPRLQPRYYSISSSPKLHPNSVHVTCVVLKYQNIPSDTVRQK

WIYGVSSNFLLNLRLATTNEAAPFMNDNGAQSTSFPKYHIEGPRGAYKIDNMFKAPVHVRRSTFRLPTNP

KSPVIMVGPGTGVAPFRGFVQERVALARRSIEKNGPDALQDWGRISLFYGCRREDEDFLYKDEWPLYKEE

LKGKFEMYCAFSRQNYKPDGSKIYVQDLILEQREHIADAILNGKGYVYICGEAKNMSKQVEDVLAIILGE

AKGGSAAVEGVAEVKLLKERSRLMLDVWS

>Serpula lacrymans

MAVASSSDVVILALGVTLAAAYLFRDQIFAPSKPKIVTTAPSKSENGSGNPRDFVAKMKDGKKRLVIFYG

SQTGTAEEYAIRLAKEAKTKFGITSLVCDPEEYDFENLDEVPEDCAVFFVVATYGEGEPTDNAVQLTQNL

TDDSFTFGNGEHKLPGLKYVIFSLGNKTYEHYNAIGRSYDAELTKMGAVRIGERGEGDDDKSMEEDYLEW

KDAMWEAFATAMNVEEGQGGDTADFTVTELDSHPQEKVYLGELSARALTKTKGIHDAKNPYASPVRNSRE

LFQLTGDRNCIHVELDIEGSGITYQHGDHVGVWPSNPDAEVDRLLCVLGLYDKKDSVIGIESLDPALAKV

PFPVPTTYATVLRHYIDISAVVGRQILGAMSKFAPSPEAEAFLKNLNVDKEEYAAVIGQGCFKLGEILQI

AAANDIRVPPTKDNTTPWPIPFDIIVSSIPRLQPRYYSISSSPKLHPTSIHVTAVVLKYQSATNKATQAR

WVYGVGTNFILNVKFAANGEAAPLMSGTSNTDPATVSMPSYAIQGPRGAHKQETIYKIPIHVRRSTFRLP

TNPKSPVIMVGPGTGVAPFRGFVQERVALARRTLDKNGPEALADWGKISLFYGCRKSTEDFLYKDEWPTY

TEELRGKFTMHCAFSREPPYKPDGSKIYVQDLIWQDRENIAEAILNGKGYVYICGDAKAMSKAVEEVLSR

ILGEAKGGSAEVEGAAEMKLLKERSRLMLDVWS

>Pleurotus ostreatus

MASTSSSDVLVLAVGVALAAIYLFRDQLFAQAKPKSVPVPTSKASNGSGNPRDFIAKMKEGKKRLVIFYG

SQTGTAEEYAIRLAKEAKSKFGLASLVCDPEEYDFENLDQLPEDCAVFFVMATYGEGEPTDNAVTLMQNL

EDESFEFSNGEHKLEGLKYVVFSLGNKTYEHYNKIGRDVDNVLTKMGAIRIGERGEGDDDKSMEEDYLEW

KDGMWDAFSTAMGVEEGQGGDTPDFAVTELESHPPEKVYLGELSARALTKTKGIHDAKNPFPAPISVARE

LFQSTHDRNCVHIELNTESSGISYQHGDHVGVWPSNPDVEVTRLLCALGLYEKKDNVIGIESLDPALAKV

PFPVPTTYATVLRHYIDISAVAGRQILGALSKFAPNPEAEAFLKGLSTNKEEYHTLIANGCLKLGEVLQL

AAGNDLSAAPTPENTTAWTIPFDIIVSSIPRLQPRYYSISSSPKLHPNSIHVTAVVLKYESIPNERVNGR

WIFGVGSNFLLNLKYAANGETAPLVATGSESKAASVTIPGYAIEGPRGAYKQETIYKAPIHVRRSTFRLP

TNPKSPVIMIGPGTGVAPFRGFIQERVALARRSIEKNGPDALADWGRISLFYGCRKSTEDFLYKDEWPQY

QEELRGKFTMHCAFSREVYRPDGSKIYVQDLLWDDREQVADAIINGKGYIYICGDAKSMSKAVEETLAKI

LGEAKGGSADVEGNAEVKLLKERSRLMLDVWS

>Drosophila melanogaster

MASEQTIDGAAAIPSGGGDEPFLGLLDVALLAVLIGGVTFYFLRTRKKEEEPTRSYSIQPTTVCTTSASD

NSFIKKLKASGRSLVVFYGSQTGTGEEFAGRLAKEGIRYRLKGMVADPEECDMEELLQLKDTDNSLAVFC

LATYGEGDPTDNAMEFYEWITSGDVDLSGLNYAVFGLGNKTYEHYNKVAIYVDKRLEELGANRVFELGLG

DDDANIEDDFITWKDRFWPAVCDHFGIEGGGEEVLIRQYRLLEQPDVQPDRIYTGEIARLHSIQNQRPPF

DAKNPFLAPIKVNRELHKGGGRSCMHIELSIEGSKMRYDAGDHVAMFPVNDKSLVEKLGQLCNADLDTVF

SLINTDTDSSKKHPFPCPTTYRTALTHYLEITAIPRTHILKELAEYCTDEKEKELLRSMASISPEGKEKY

QSWIQDACRNIVHILEDIKSCRPPIDHVCELLPRLQPRYYSISSSAKLHPTDVHVTAVLVEYKTPTGRIN

KGVATTYLKNKQPQGSEEVKVPVFIRKSQFRLPTKPETPIIMVGPGTGLAPFRGFIQERQFLRDEGKTVG

ESILYFGCRKRSEDYIYESELEEWVKKGTLNLKAAFSRDQGKKVYVQHLLEQDADLIWNVIGENKGHFYI

CGDAKNMAVDVRNILVKILSTKGNMSEADAVQYIKKMEAQKRYSADVWS

>Musca domestica

MSAEHVEEVVSEEPFLGTLDIALLVVLLVGATWYFMRSRKKEEAPIRSYSIQPTTVSTVSTTENSFIKKL

KASGRSLVVFYGSQTGTAEEFAGRLAKEGLRYRMKGMVADPEECDMEELLQMKDIPNSLAVFCLATYGEG

DPTDNAMEFYEWITNGEVDLTGLNYAVFGLGNKTYEHYNKVAIYVDKRLEELGATRVFELGLGDDDANIE

DDFITWKDRFWPSVCDFFGIEGSGEEVLMRQFRLLEQPDVQPDRIYTGEIARLHSMQNQRPPFDAKNPFL

ASVIVNRELHKGGGRSCMHIELDIDGSKMRYDAGDHIAMYPINDKILVEKLGKLCDANLDTVFSLINTDT

DSSKKHPFPCPTTYRTALTHYLEITAIPRTHILKELAEYCSDEKDKEFLRNMASITPEGKEKYQNWIQNS

SRNIVHILEDIKSCRPPIDHICELLPRLQPRYYSISSSSKLYPTNVHITAVLVQYETPTGRVNKGVATSY

MKEKNPSVGEVKVPVFIRKSQFRLPTKSEIPIIMVGPGTGLAPFRGFIQERQFLRDGGKVVGDTILYFGC

RKKDEDFIYREELEQYVQNGTLTLKTAFSRDQQEKIYVTHLIEQDADLIWKVIGEQKGHFYICGDAKNMA

VDVRNILVKILSTKGNMNESDAVQYIKKMEAQKRYSADVWS

>Homo sapiens

MINMGDSHVDTSSTVSEAVAEEVSLFSMTDMILFSLIVGLLTYWFLFRKKKEEVPEFTKIQTLTSSVRES

SFVEKMKKTGRNIIVFYGSQTGTAEEFANRLSKDAHRYGMRGMSADPEEYDLADLSSLPEIDNALVVFCM

ATYGEGDPTDNAQDFYDWLQETDVDLSGVKFAVFGLGNKTYEHFNAMGKYVDKRLEQLGAQRIFELGLGD

DDGNLEEDFITWREQFWLAVCEHFGVEATGEESSIRQYELVVHTDIDAAKVYMGEMGRLKSYENQKPPFD

AKNPFLAAVTTNRKLNQGTERHLMHLELDISDSKIRYESGDHVAVYPANDSALVNQLGKILGADLDVVMS

LNNLDEESNKKHPFPCPTSYRTALTYYLDITNPPRTNVLYELAQYASEPSEQELLRKMASSSGEGKELYL

SWVVEARRHILAILQDCPSLRPPIDHLCELLPRLQARYYSIASSSKVHPNSVHICAVVVEYETKAGRINK

GVATNWLRAKEPVGENGGRALVPMFVRKSQFRLPFKATTPVIMVGPGTGVAPFIGFIQERAWLRQQGKEV

GETLLYYGCRRSDEDYLYREELAQFHRDGALTQLNVAFSREQSHKVYVQHLLKQDREHLWKLIEGGAHIY

VCGDARNMARDVQNTFYDIVAELGAMEHAQAVDYIKKLMTKGRYSLDVWS

> Mus musculus

MGDSHEDTSATVPEAVAEEVSLFSTTDIVLFSLIVGVLTYWFIFKKKKEEIPEFSKIQTTAPPVKESSFV

EKMKKTGRNIIVFYGSQTGTAEEFANRLSKDAHRYGMRGMSADPEEYDLADLSSLPEIDKSLVVFCMATY

GEGDPTDNAQDFYDWLQETDVDLTGVKFAVFGLGNKTYEHFNAMGKYVDQRLEQLGAQRIFELGLGDDDG

NLEEDFITWREQFWPAVCEFFGVEATGEESSIRQYELVVHEDMDTAKVYTGEMGRLKSYENQKPPFDAKN

PFLAAVTTNRKLNQGTERHLMHLELDISDSKIRYESGDHVAVYPANDSTLVNQIGEILGADLDVIMSLNN

LDEESNKKHPFPCPTTYRTALTYYLDITNPPRTNVLYELAQYASEPSEQEHLHKMASSSGEGKELYLSWV

VEARRHILAILQDYPSLRPPIDHLCELLPRLQARYYSIASSSKVHPNSVHICAVAVEYEAKSGRVNKGVA

TSWLRTKEPAGENGRRALVPMFVRKSQFRLPFKPTTPVIMVGPGTGVAPFMGFIQERAWLREQGKEVGET

LLYYGCRRSDEDYLYREELARFHKDGALTQLNVAFSREQAHKVYVQHLLKRDKEHLWKLIHEGGAHIYVC

GDARNMAKDVQNTFYDIVAEFGPMEHTQAVDYVKKLMTKGRYSLDVWS

>Gallus gallus

MGDAGMESTVSPPEGTAQDSFLSMTDVFLISLITGLFTYWFFFRKKKEEIPDLPKIQTVSSPARDSSFIE

KMKKTGRNIVVFYGSQTGTAEEFANRLSKDAHRYGLRGMAADPEEYDLSDLSRLSEIDKSLAVFCMATYG

EGDPTDNAQDFYDWLQEADTDLSGLRFAVFGLGNKTYEHFNAMGKYVDKRLEELGAQRIFELGLGDDDGN

LEEDFITWREQFWPAVCEHFGVEATGEESSIRQYELVVHTDVNMNKVYTGEMGRLKSYENQKPPFDAKNP

FLAVVTENRKLNEGGERHLMHLELDISNSKIRYESGDHVAVYPANDASLVNQLGEILGTDLDTVMSLNNL

DEESNKKHPFPCPTSYRTALTYYLDITNPPRTNVLYELAQYATDTGEQEQLRKMASSSAEGKALYLSWVV

EARRNILAILQDMPSLRPPIDHLCELLPRLQARYYSIASSSKVHPNSIHICAVTVEYETKTGRLNKGVAT

NWLKDKVPNENGRNSLVPMYVRKSQFRLPFKPSTPVIMIGPGTGIAPFIGFIQERAWLKEQGKEVGETVL

YYGCRREREDYLYRQELARFKQEGVLTQLNVAFSRDQAEKVYVQHLLKKNKEHIWKLVNDGNAHIYVCGD

ARNMARDVQNTFYEIVSEYGNMNQSQAVDYVKKLMTKGRYSLDVWS

>Arabidopsis thaliana 1

MTSALYASDLFKQLKSIMGTDSLSDDVVLVIATTSLALVAGFVVLLWKKTTADRSGELKPLMIPKSLMAK

DEDDDLDLGSGKTRVSIFFGTQTGTAEGFAKALSEEIKARYEKAAVKDDYAADDDQYEEKLKKETLAFFC

VATYGDGEPTDNAARFYKWFTEENERDIKLQQLAYGVFALGNRQYEHFNKIGIVLDEELCKKGAKRLIEV

GLGDDDQSIEDDFNAWKESLWSELDKLLKDEDDKSVATPYTAVIPEYRVVTHDPRFTTQKSMESNVANGN

TTIDIHHPCRVDVAVQKELHTHESDRSCIHLEFDISRTGITYETGDHVGVYAENHVEIVEEAGKLLGHSL

DLVFSIHADKEDGSPLESAVPPPFPGPCTLGTGLARYADLLNPPRKSALVALAAYATEPSEAEKLKHLTS

PDGKDEYSQWIVASQRSLLEVMAAFPSAKPPLGVFFAAIAPRLQPRYYSISSSPRLAPSRVHVTSALVYG

PTPTGRIHKGVCSTWMKNAVPAEKSHECSGAPIFIRASNFKLPSNPSTPIVMVGPGTGLAPFRGFLQERM

ALKEDGEELGSSLLFFGCRNRQMDFIYEDELNNFVDQGVISELIMAFSREGAQKEYVQHKMMEKAAQVWD

LIKEEGYLYVCGDAKGMARDVHRTLHTIVQEQEGVSSSEAEAIVKKLQTEGRYLRDVW

>Arabidopsis thaliana 2

GRRSGSGNSKRVEPLKPLVIKPREEEIDDGRKKVTIFFGTQTGTAEGFAKALGEEAKARYEKTRFKIVDL

DDYAADDDEYEEKLKKEDVAFFFLATYGDGEPTDNAARFYKWFTEGNDRGEWLKNLKYGVFGLGNRQYEH

FNKVAKVVDDILVEQGAQRLVQVGLGDDDQCIEDDFTAWREALWPELDTILREEGDTAVATPYTAAVLEY

RVSIHDSEDAKFNDINMANGNGYTVFDAQHPYKANVAVKRELHTPESDRSCIHLEFDIAGSGLTYETGDH

VGVLCDNLSETVDEALRLLDMSPDTYFSLHAEKEDGTPISSSLPPPFPPCNLRTALTRYACLLSSPKKSA

LVALAAHASDPTEAERLKHLASPAGKDEYSKWVVESQRSLLEVMAEFPSAKPPLGVFFAGVAPRLQPRFY

SISSSPKIAETRIHVTCALVYEKMPTGRIHKGVCSTWMKNAVPYEKSENCSSAPIFVRQSNFKLPSDSKV

PIIMIGPGTGLAPFRGFLQERLALVESGVELGPSVLFFGCRNRRMDFIYEEELQRFVESGALAELSVAFS

REGPTKEYVQHKMMDKASDIWNMISQGAYLYVCGDAKGMARDVHRSLHTIAQEQGSMDSTKAEGFVKNLQ

TSGRYLRDVW

>Triticum aestivum

MDSAAAGMRDSALDLLAALLTGRAPPAAADGDQNRRLLALLATSLAVLVGCGVALLFRRSSSGAAPLAHK

SAAAKPLAAKKDQEPDPDDGRQRVALFFGTQTGTAEGFAKALAEEAKARYDKAVFKVLDLDDYAAEDEEY

EEKLKKENIAFFFLATYGDGEPTDNAARFYKWFSEGNERGEWLSNLKFGVFALGNRQYEHFNKVGKEVDQ

LLAEQGGKRIVPVGLGDDDQCIEDDFNAWKELLWPELDKLLRVEDNSSTAQSPYTAAIPQYRVVLTKPED

ATHINKSFSLSNGHVVYDSQHPCRANVAVRRELHTPASDRSCIHLEFDIAGTSLTYETGDHVGVYAENSI

ETVEEAEKLLDYSPDTYFSIYADQEDGTPLFGGSLPPPFPSPCTVRVALARYADLLNSPKKSVLLALAAH

ASDPKEAERLRHLASPAGKKEYSQWIIASQRSLLEVISEFPSAKPPLGVFFAAIAPRLQPRYYSISSSPR

MAPTRIHVTCSLVHGQTPTGRIHKGVCSTWMKNSTPLEESQECSWAPIFVRQSNFKLPADPTVPIIMVGP

GTGLAPFRGFLQERLALKETGVELGRAILFFGCRNRQMDFIYEDELNNFAESGALSELVVAFSREGPTKE

YVQHKMAEKAAELWSIVSQGGYVYVCGDAKGMARDVHRALHTIVQEQGSLDSSKAEGYVKNLQMEGRYLR

DVW

>Trypanosoma brucei

MIFLVISTAIIAVLAWFVAGVFIRGGGGRGKTAAPQVVGVAQYPSQPSSRVDVRVLFGSQTGTAEMFAKT

VTREGLRLGVPMKLADVENYRPSDLAGEKYVIIICATYGEGEPTDTMVGFHEWLVDDSRAVGEELSGVKY

TVFALGDRQYKFFCREGITVDRRMSELGAQRFYPLGYGDCGNSIEEEFDNWCHNLWPVLGRALSLVLKSN

STEPVAPECRMKLWGPPEEAPLPFPKLASVLEPTQRLPSWAPVKVNKELLSNATGRSTRLIEFDTSETVI

SYQAGDHLGVLPSNPSEMVNTYLRVLGVSEQESSQVISLQNRATGKNVFPCRVSIRTALTWYIDLAGPPK

KSTLRAFAHHCTDPVEKDTLLKLLSTEPESVEAYGKLVLELRTVLGFLQRFKSMSPPLSFFLEMMPRIAP

RYFSISSDSLTHPTSVAITVAVVEGGLCTNLLQQAAVGQNIPVFVRKSNFHLPLRAKDRPIIMIGPGTGV

APFIGFLHRRSAWLEKGNKVGDALLFFGCRRREEDHIYADFMEKCLSNGALSVRDVAYSREQADKVYVQH

RLAARGKEVWEIISRGGNVYVCGDAKNMARDVERQLLDIAQKYGAMKEDEATALLEKLATDERYLKDVWT

A

>Dictyostelium discoideum

MEILESIDFIEVLILDNLGAIIIVAVIVGTYLYMNKPPPPPPVFNKPNNKINKEAQKPKKTITKNEDGKKVMKIFFGTQTRTAEDFSRIIEKECKKIGIPCEVVDLESYEHEQELHSESFVMFLVATHGEGDPTDNAKEFYLWLTNDERPTDLLNGVPFTVFGLGNKTYEHYNAVARVIDRRMEELGGKRVFERGEGDDDATLEEDFNRWKKDMWPVVCKFLGYELKSTEDDKFVPRFRMVTLNQDSKDINDPFIKIVSTPLKPKLSTDNKVIYDMKNPYYAEVLENRELHSNESDRSCRHIEFKLGDEVSYTTGDHLGVFPINDSKLVEQLIKRLGVNGDDMIALVPIDQEGSVIKASFGPMTIRRAFSEHLDITNPVRKSVLRALAESTTNEEEKKRLLYLATEEANEEYNKYIKNDFRGVVDLLESFPGLQPLIAHFLEFTPRLPARMYSISSSPHNKNGVVSITSVVVNFTTGNQRAHNGVASTWLSHLKVGDKVPLFVRESHFKLPSAATEQKPVIMVGPGTGLAPFRGFLQELQHRNHSQQQQSLLFFGCRSDTVDYIYREELEQYHQSSVLGDLVVAFSRKTSQKVYVQNKLLEHKEKVWELLNKGAYFYVCGDGRNMSKAVQQALLSIIKEFGSKDDNSAQQFIDDMSSHGRYLQDVWF

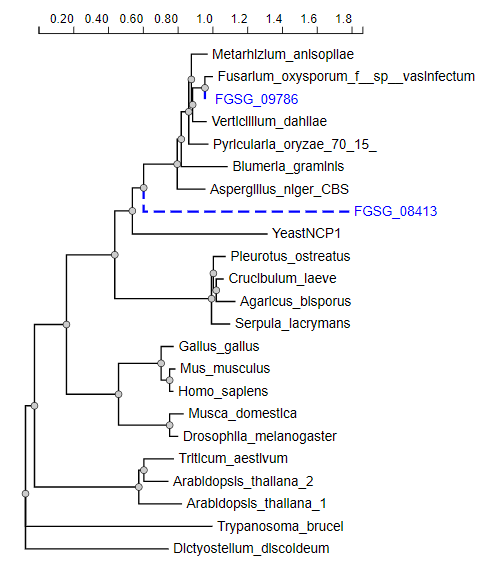

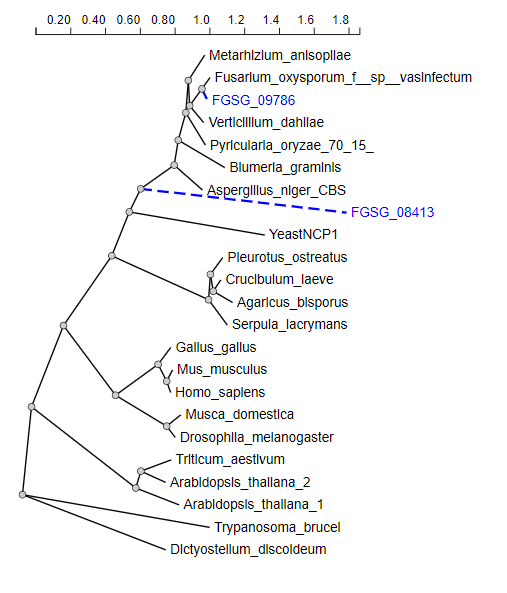

### Summary of 20 predicted sequences

Table of predicted subcelullar localizations. Use the help page for more detailed description of the output page.

#### Predicted proteins

**Musca**

**Prediction: Endoplasmic reticulum, Membrane**

| **Localization** | **Endoplasmic reticulum** | | **Mitochondrion** | | **Peroxisome** | | **Cell membrane** | **Lysosome/Vacuole** | **Golgi apparatus** | **Plastid** | **Cytoplasm** | **Nucleus** | **Extracellular** |
| --- | --- | --- | --- | --- | --- | --- | --- | --- | --- | --- | --- | --- | --- |
| **Likelihood** | 0.8887 | | 0.0812 | | 0.0102 | | 0.0097 | 0.0051 | 0.0033 | 0.0014 | 0.0002 | 0.0002 | 0 |
| **Type** | | **Soluble** | | **Membrane** | |  |  |  |  |  |  |  |  |
| **Likelihood** | | 0 | | 1 | |  |  |  |  |  |  |  |  |

**Hierarchical Tree. Donwload:** [PNG](http://www.cbs.dtu.dk/services/DeepLoc-1.0/tmp/5D01BB8C00003EF54AA12939/tree_15.png) **/** [EPS](http://www.cbs.dtu.dk/services/DeepLoc-1.0/tmp/5D01BB8C00003EF54AA12939/tree_15.eps)
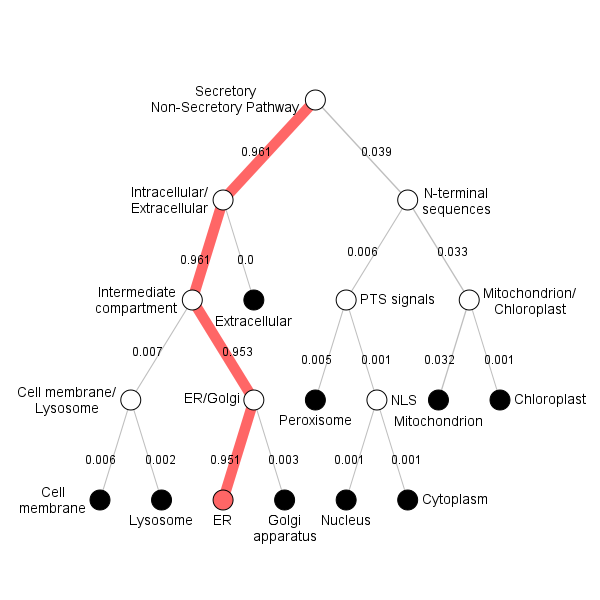

**Position Importance. Donwload:** [PNG](http://www.cbs.dtu.dk/services/DeepLoc-1.0/tmp/5D01BB8C00003EF54AA12939/alpha_15.png) **/** [EPS](http://www.cbs.dtu.dk/services/DeepLoc-1.0/tmp/5D01BB8C00003EF54AA12939/alpha_15.eps) **/** [CSV](http://www.cbs.dtu.dk/services/DeepLoc-1.0/tmp/5D01BB8C00003EF54AA12939/alpha_15.csv)
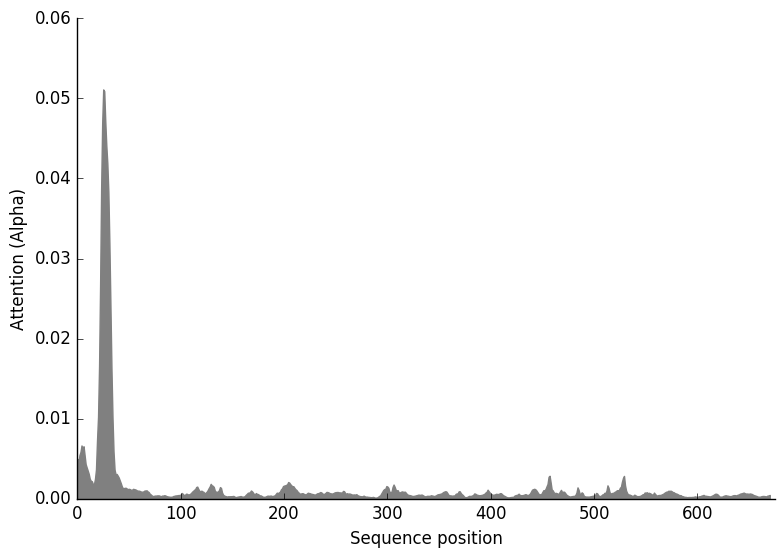

**Pleurotus**

**Prediction: Endoplasmic reticulum, Membrane**

| **Localization** | **Endoplasmic reticulum** | | **Mitochondrion** | | **Cytoplasm** | | **Cell membrane** | **Lysosome/Vacuole** | **Peroxisome** | **Golgi apparatus** | **Plastid** | **Nucleus** | **Extracellular** |
| --- | --- | --- | --- | --- | --- | --- | --- | --- | --- | --- | --- | --- | --- |
| **Likelihood** | 0.8031 | | 0.103 | | 0.0396 | | 0.0293 | 0.0106 | 0.0071 | 0.0034 | 0.0028 | 0.0011 | 0 |
| **Type** | | **Soluble** | | **Membrane** | |  |  |  |  |  |  |  |  |
| **Likelihood** | | 0.0623 | | 0.9377 | |  |  |  |  |  |  |  |  |

**Hierarchical Tree. Donwload:** [PNG](http://www.cbs.dtu.dk/services/DeepLoc-1.0/tmp/5D01BB8C00003EF54AA12939/tree_11.png) **/** [EPS](http://www.cbs.dtu.dk/services/DeepLoc-1.0/tmp/5D01BB8C00003EF54AA12939/tree_11.eps)
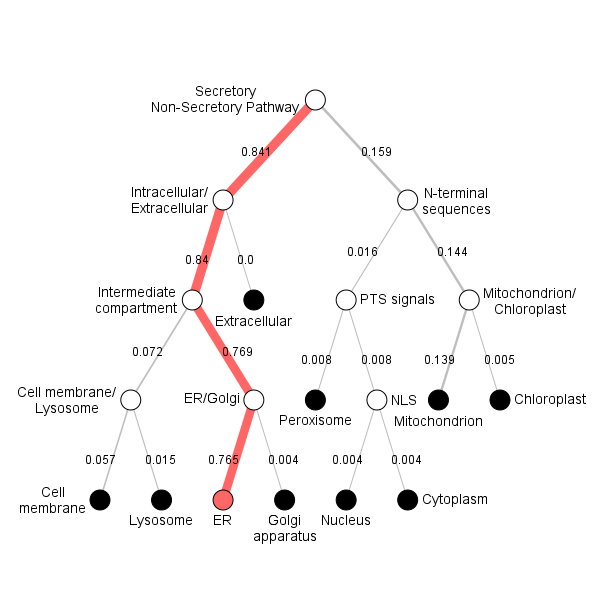

**Position Importance. Donwload:** [PNG](http://www.cbs.dtu.dk/services/DeepLoc-1.0/tmp/5D01BB8C00003EF54AA12939/alpha_11.png) **/** [EPS](http://www.cbs.dtu.dk/services/DeepLoc-1.0/tmp/5D01BB8C00003EF54AA12939/alpha_11.eps) **/** [CSV](http://www.cbs.dtu.dk/services/DeepLoc-1.0/tmp/5D01BB8C00003EF54AA12939/alpha_11.csv)
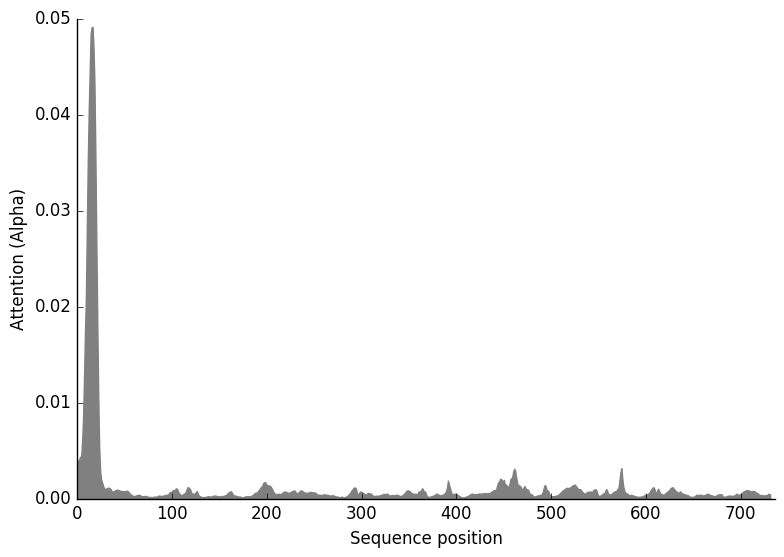

**Fusarium**

**Prediction: Endoplasmic reticulum, Membrane**

| **Localization** | **Endoplasmic reticulum** | | **Mitochondrion** | | **Cytoplasm** | | **Cell membrane** | **Lysosome/Vacuole** | **Plastid** | **Golgi apparatus** | **Peroxisome** | **Nucleus** | **Extracellular** |
| --- | --- | --- | --- | --- | --- | --- | --- | --- | --- | --- | --- | --- | --- |
| **Likelihood** | 0.7971 | | 0.109 | | 0.0392 | | 0.023 | 0.0116 | 0.0111 | 0.005 | 0.0034 | 0.0006 | 0 |
| **Type** | | **Soluble** | | **Membrane** | |  |  |  |  |  |  |  |  |
| **Likelihood** | | 0.0623 | | 0.9377 | |  |  |  |  |  |  |  |  |

**Hierarchical Tree. Donwload:** [PNG](http://www.cbs.dtu.dk/services/DeepLoc-1.0/tmp/5D01BB8C00003EF54AA12939/tree_5.png) **/** [EPS](http://www.cbs.dtu.dk/services/DeepLoc-1.0/tmp/5D01BB8C00003EF54AA12939/tree_5.eps)
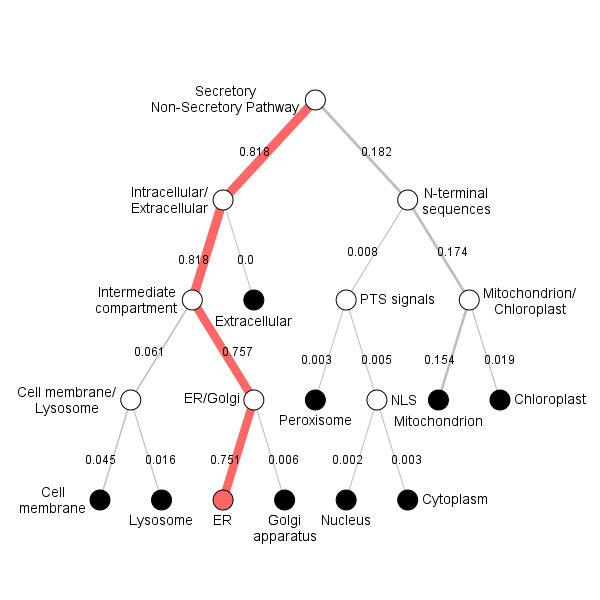

**Position Importance. Donwload:** [PNG](http://www.cbs.dtu.dk/services/DeepLoc-1.0/tmp/5D01BB8C00003EF54AA12939/alpha_5.png) **/** [EPS](http://www.cbs.dtu.dk/services/DeepLoc-1.0/tmp/5D01BB8C00003EF54AA12939/alpha_5.eps) **/** [CSV](http://www.cbs.dtu.dk/services/DeepLoc-1.0/tmp/5D01BB8C00003EF54AA12939/alpha_5.csv)
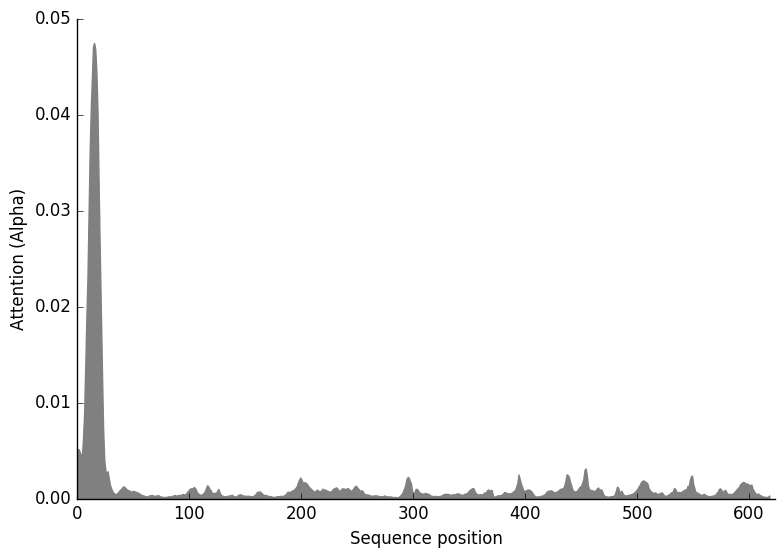

**Drosophila**

**Prediction: Endoplasmic reticulum, Membrane**

| **Localization** | **Endoplasmic reticulum** | | **Mitochondrion** | | **Cell membrane** | | **Lysosome/Vacuole** | **Golgi apparatus** | **Peroxisome** | **Plastid** | **Nucleus** | **Cytoplasm** | **Extracellular** |
| --- | --- | --- | --- | --- | --- | --- | --- | --- | --- | --- | --- | --- | --- |
| **Likelihood** | 0.9031 | | 0.055 | | 0.0238 | | 0.0083 | 0.0063 | 0.0031 | 0.0002 | 0.0001 | 0 | 0 |
| **Type** | | **Soluble** | | **Membrane** | |  |  |  |  |  |  |  |  |
| **Likelihood** | | 0 | | 1 | |  |  |  |  |  |  |  |  |

**Hierarchical Tree. Donwload:** [PNG](http://www.cbs.dtu.dk/services/DeepLoc-1.0/tmp/5D01BB8C00003EF54AA12939/tree_12.png) **/** [EPS](http://www.cbs.dtu.dk/services/DeepLoc-1.0/tmp/5D01BB8C00003EF54AA12939/tree_12.eps)
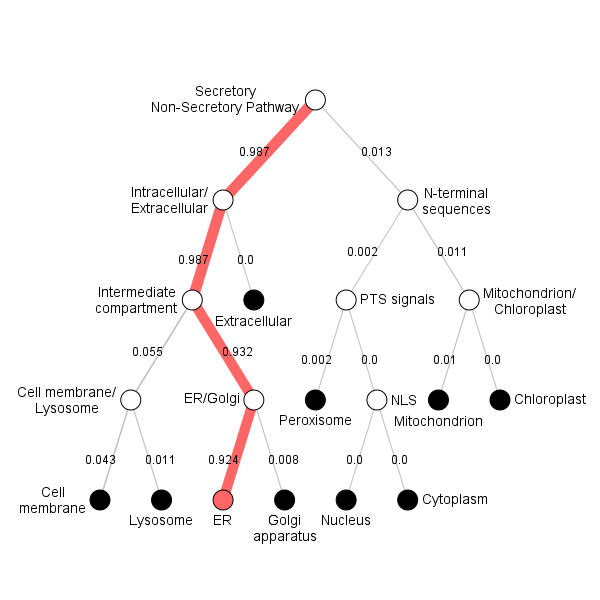

**Position Importance. Donwload:** [PNG](http://www.cbs.dtu.dk/services/DeepLoc-1.0/tmp/5D01BB8C00003EF54AA12939/alpha_12.png) **/** [EPS](http://www.cbs.dtu.dk/services/DeepLoc-1.0/tmp/5D01BB8C00003EF54AA12939/alpha_12.eps) **/** [CSV](http://www.cbs.dtu.dk/services/DeepLoc-1.0/tmp/5D01BB8C00003EF54AA12939/alpha_12.csv)
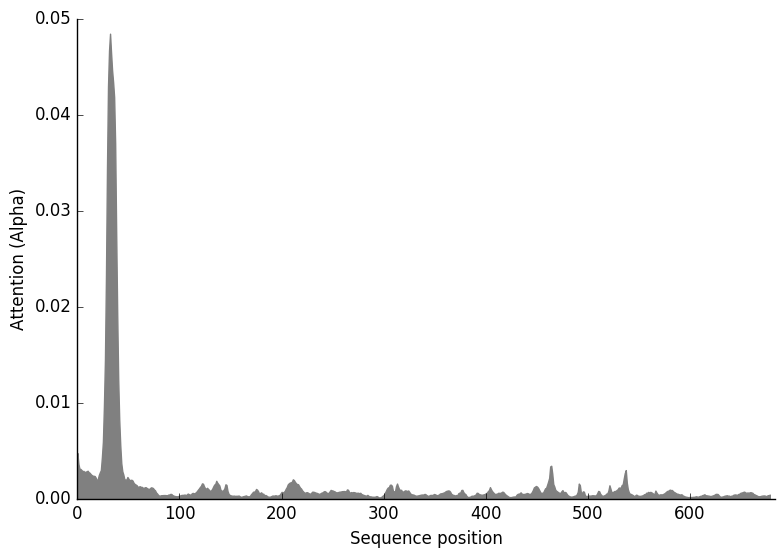

**Crucibulum**

**Prediction: Endoplasmic reticulum, Membrane**

| **Localization** | **Endoplasmic reticulum** | | **Mitochondrion** | | **Cytoplasm** | | **Cell membrane** | **Lysosome/Vacuole** | **Peroxisome** | **Golgi apparatus** | **Plastid** | **Nucleus** | **Extracellular** |
| --- | --- | --- | --- | --- | --- | --- | --- | --- | --- | --- | --- | --- | --- |
| **Likelihood** | 0.8203 | | 0.0933 | | 0.0392 | | 0.0241 | 0.0091 | 0.0053 | 0.0048 | 0.003 | 0.0008 | 0 |
| **Type** | | **Soluble** | | **Membrane** | |  |  |  |  |  |  |  |  |
| **Likelihood** | | 0.0622 | | 0.9378 | |  |  |  |  |  |  |  |  |

**Hierarchical Tree. Donwload:** [PNG](http://www.cbs.dtu.dk/services/DeepLoc-1.0/tmp/5D01BB8C00003EF54AA12939/tree_8.png) **/** [EPS](http://www.cbs.dtu.dk/services/DeepLoc-1.0/tmp/5D01BB8C00003EF54AA12939/tree_8.eps)
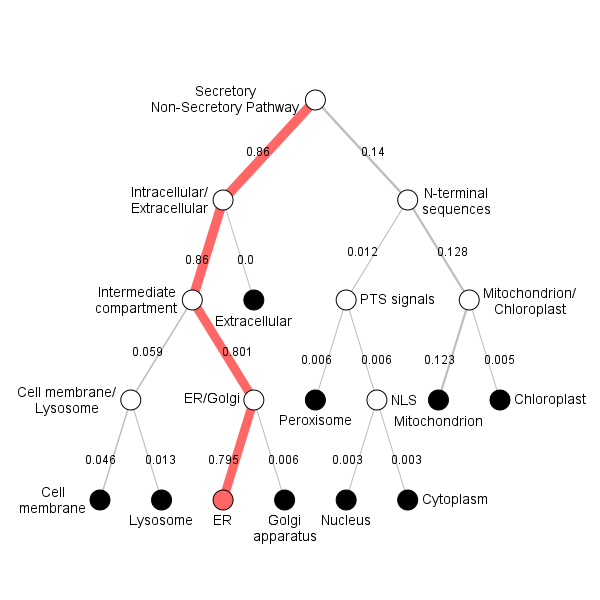

**Position Importance. Donwload:** [PNG](http://www.cbs.dtu.dk/services/DeepLoc-1.0/tmp/5D01BB8C00003EF54AA12939/alpha_8.png) **/** [EPS](http://www.cbs.dtu.dk/services/DeepLoc-1.0/tmp/5D01BB8C00003EF54AA12939/alpha_8.eps) **/** [CSV](http://www.cbs.dtu.dk/services/DeepLoc-1.0/tmp/5D01BB8C00003EF54AA12939/alpha_8.csv)
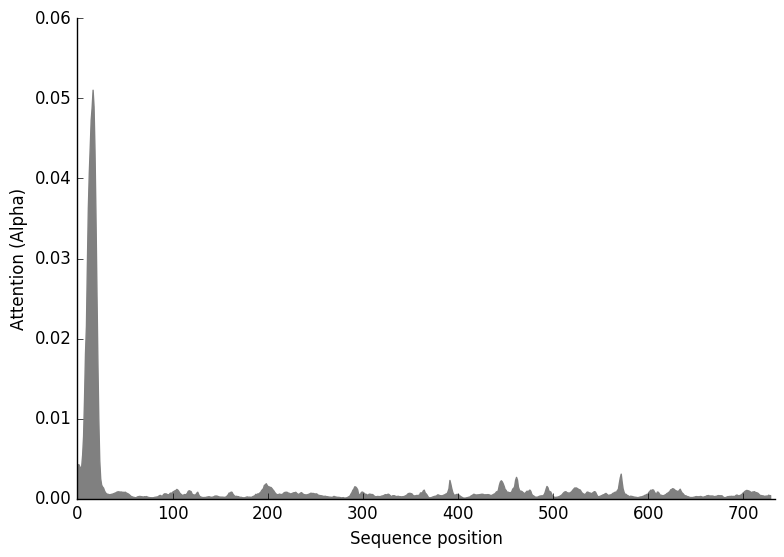

**Metarhizium**

**Prediction: Endoplasmic reticulum, Membrane**

| **Localization** | **Endoplasmic reticulum** | | **Mitochondrion** | | **Plastid** | | **Cell membrane** | **Lysosome/Vacuole** | **Golgi apparatus** | **Peroxisome** | **Cytoplasm** | **Nucleus** | **Extracellular** |
| --- | --- | --- | --- | --- | --- | --- | --- | --- | --- | --- | --- | --- | --- |
| **Likelihood** | 0.8433 | | 0.0751 | | 0.0321 | | 0.0263 | 0.0159 | 0.0047 | 0.0014 | 0.0009 | 0.0002 | 0 |
| **Type** | | **Soluble** | | **Membrane** | |  |  |  |  |  |  |  |  |
| **Likelihood** | | 0.0011 | | 0.9989 | |  |  |  |  |  |  |  |  |

**Hierarchical Tree. Donwload:** [PNG](http://www.cbs.dtu.dk/services/DeepLoc-1.0/tmp/5D01BB8C00003EF54AA12939/tree_3.png) **/** [EPS](http://www.cbs.dtu.dk/services/DeepLoc-1.0/tmp/5D01BB8C00003EF54AA12939/tree_3.eps)
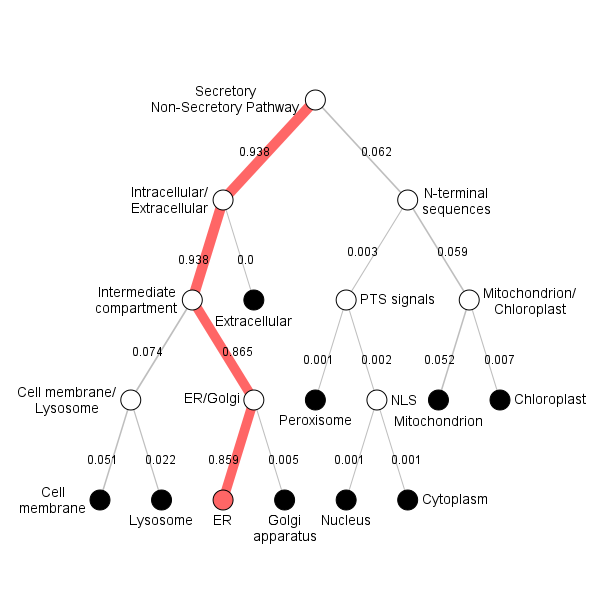

**Position Importance. Donwload:** [PNG](http://www.cbs.dtu.dk/services/DeepLoc-1.0/tmp/5D01BB8C00003EF54AA12939/alpha_3.png) **/** [EPS](http://www.cbs.dtu.dk/services/DeepLoc-1.0/tmp/5D01BB8C00003EF54AA12939/alpha_3.eps) **/** [CSV](http://www.cbs.dtu.dk/services/DeepLoc-1.0/tmp/5D01BB8C00003EF54AA12939/alpha_3.csv)
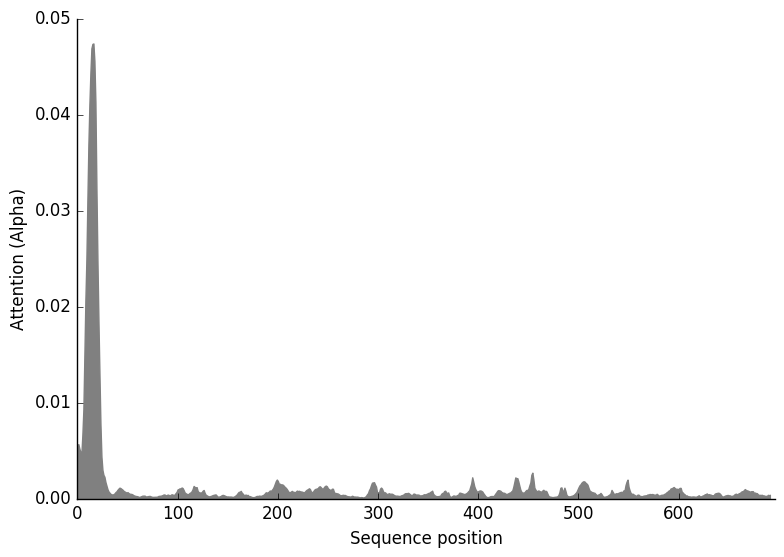

**Blumeria**

**Prediction: Endoplasmic reticulum, Membrane**

| **Localization** | **Endoplasmic reticulum** | | **Mitochondrion** | | **Cytoplasm** | | **Cell membrane** | **Lysosome/Vacuole** | **Plastid** | **Golgi apparatus** | **Peroxisome** | **Nucleus** | **Extracellular** |
| --- | --- | --- | --- | --- | --- | --- | --- | --- | --- | --- | --- | --- | --- |
| **Likelihood** | 0.8251 | | 0.0819 | | 0.0391 | | 0.0248 | 0.0146 | 0.0088 | 0.0031 | 0.0022 | 0.0003 | 0 |
| **Type** | | **Soluble** | | **Membrane** | |  |  |  |  |  |  |  |  |
| **Likelihood** | | 0.0627 | | 0.9373 | |  |  |  |  |  |  |  |  |

**Hierarchical Tree. Donwload:** [PNG](http://www.cbs.dtu.dk/services/DeepLoc-1.0/tmp/5D01BB8C00003EF54AA12939/tree_7.png) **/** [EPS](http://www.cbs.dtu.dk/services/DeepLoc-1.0/tmp/5D01BB8C00003EF54AA12939/tree_7.eps)
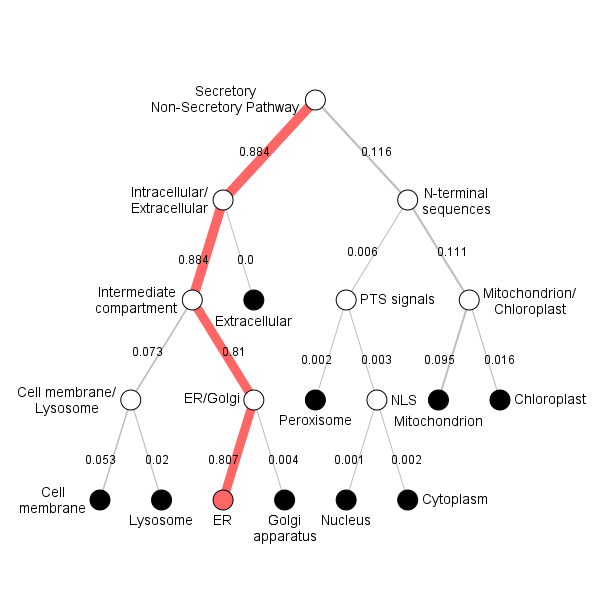

**Position Importance. Donwload:** [PNG](http://www.cbs.dtu.dk/services/DeepLoc-1.0/tmp/5D01BB8C00003EF54AA12939/alpha_7.png) **/** [EPS](http://www.cbs.dtu.dk/services/DeepLoc-1.0/tmp/5D01BB8C00003EF54AA12939/alpha_7.eps) **/** [CSV](http://www.cbs.dtu.dk/services/DeepLoc-1.0/tmp/5D01BB8C00003EF54AA12939/alpha_7.csv)
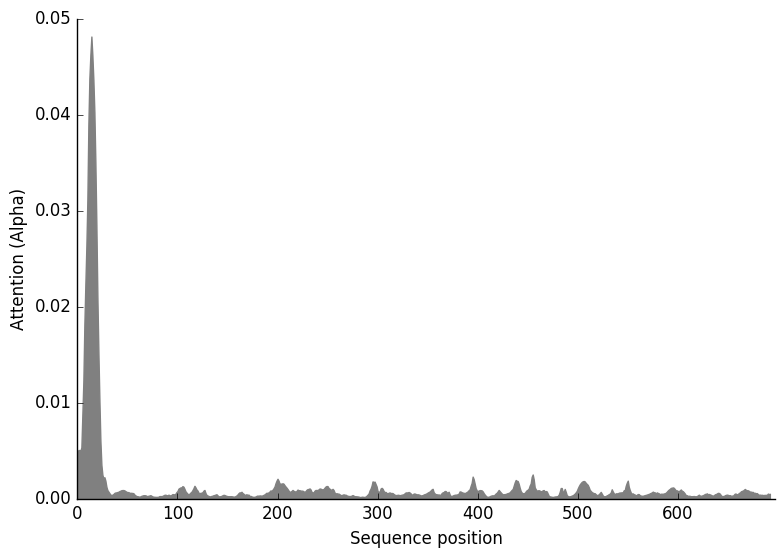

**Dictyostelium**

**Prediction: Endoplasmic reticulum, Membrane**

| **Localization** | **Endoplasmic reticulum** | | **Mitochondrion** | | **Peroxisome** | | **Plastid** | **Golgi apparatus** | **Cell membrane** | **Lysosome/Vacuole** | **Cytoplasm** | **Nucleus** | **Extracellular** |
| --- | --- | --- | --- | --- | --- | --- | --- | --- | --- | --- | --- | --- | --- |
| **Likelihood** | 0.8603 | | 0.1034 | | 0.0191 | | 0.006 | 0.0051 | 0.0025 | 0.002 | 0.0009 | 0.0007 | 0 |
| **Type** | | **Soluble** | | **Membrane** | |  |  |  |  |  |  |  |  |
| **Likelihood** | | 0.0009 | | 0.9991 | |  |  |  |  |  |  |  |  |

**Hierarchical Tree. Donwload:** [PNG](http://www.cbs.dtu.dk/services/DeepLoc-1.0/tmp/5D01BB8C00003EF54AA12939/tree_20.png) **/** [EPS](http://www.cbs.dtu.dk/services/DeepLoc-1.0/tmp/5D01BB8C00003EF54AA12939/tree_20.eps)
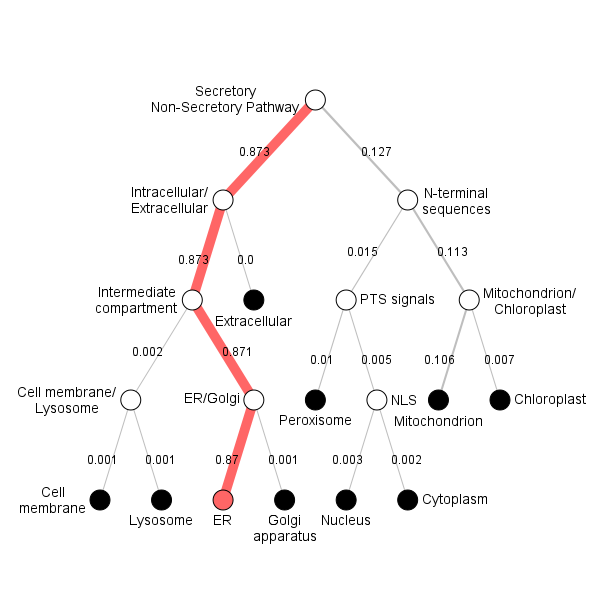

**Position Importance. Donwload:** [PNG](http://www.cbs.dtu.dk/services/DeepLoc-1.0/tmp/5D01BB8C00003EF54AA12939/alpha_20.png) **/** [EPS](http://www.cbs.dtu.dk/services/DeepLoc-1.0/tmp/5D01BB8C00003EF54AA12939/alpha_20.eps) **/** [CSV](http://www.cbs.dtu.dk/services/DeepLoc-1.0/tmp/5D01BB8C00003EF54AA12939/alpha_20.csv)
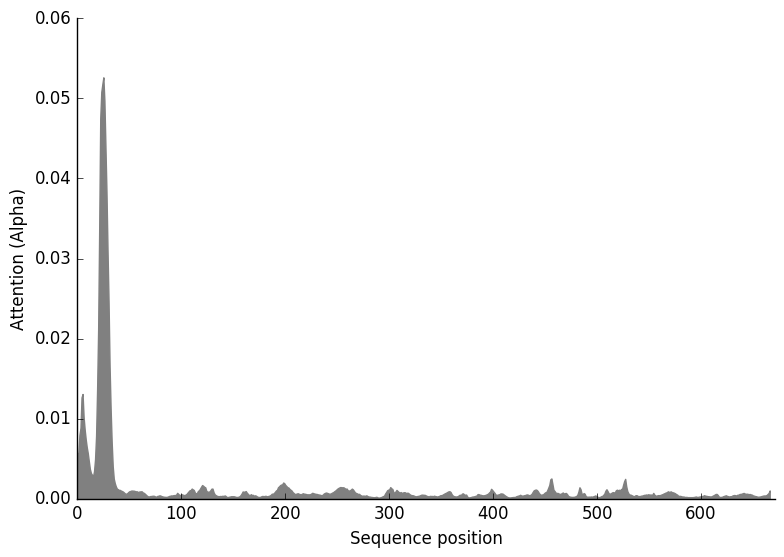

**Aspergillus**

**Prediction: Endoplasmic reticulum, Membrane**

| **Localization** | **Endoplasmic reticulum** | | **Mitochondrion** | | **Cell membrane** | | **Lysosome/Vacuole** | **Plastid** | **Golgi apparatus** | **Peroxisome** | **Cytoplasm** | **Nucleus** | **Extracellular** |
| --- | --- | --- | --- | --- | --- | --- | --- | --- | --- | --- | --- | --- | --- |
| **Likelihood** | 0.857 | | 0.0849 | | 0.0265 | | 0.0172 | 0.0079 | 0.0035 | 0.0021 | 0.0008 | 0.0003 | 0 |
| **Type** | | **Soluble** | | **Membrane** | |  |  |  |  |  |  |  |  |
| **Likelihood** | | 0.0008 | | 0.9992 | |  |  |  |  |  |  |  |  |

**Hierarchical Tree. Donwload:** [PNG](http://www.cbs.dtu.dk/services/DeepLoc-1.0/tmp/5D01BB8C00003EF54AA12939/tree_6.png) **/** [EPS](http://www.cbs.dtu.dk/services/DeepLoc-1.0/tmp/5D01BB8C00003EF54AA12939/tree_6.eps)
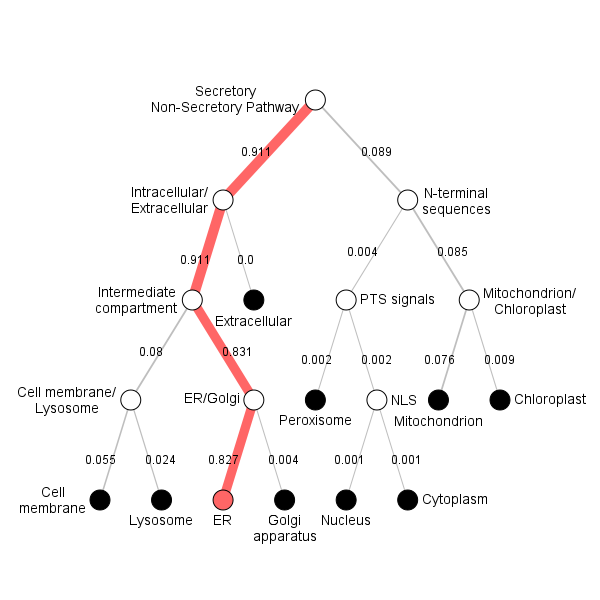

**Position Importance. Donwload:** [PNG](http://www.cbs.dtu.dk/services/DeepLoc-1.0/tmp/5D01BB8C00003EF54AA12939/alpha_6.png) **/** [EPS](http://www.cbs.dtu.dk/services/DeepLoc-1.0/tmp/5D01BB8C00003EF54AA12939/alpha_6.eps) **/** [CSV](http://www.cbs.dtu.dk/services/DeepLoc-1.0/tmp/5D01BB8C00003EF54AA12939/alpha_6.csv)
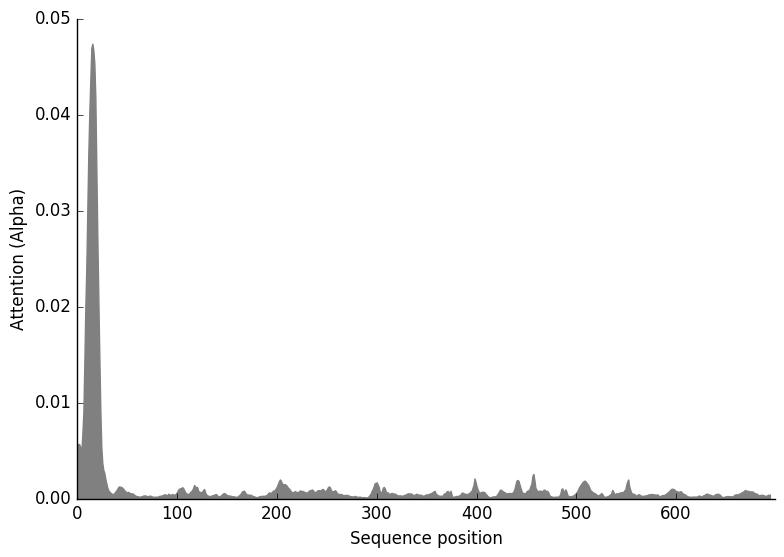

**Pyricularia**

**Prediction: Endoplasmic reticulum, Membrane**

| **Localization** | **Endoplasmic reticulum** | | **Mitochondrion** | | **Cell membrane** | | **Lysosome/Vacuole** | **Plastid** | **Golgi apparatus** | **Peroxisome** | **Cytoplasm** | **Nucleus** | **Extracellular** |
| --- | --- | --- | --- | --- | --- | --- | --- | --- | --- | --- | --- | --- | --- |
| **Likelihood** | 0.8535 | | 0.0954 | | 0.0214 | | 0.0131 | 0.0118 | 0.0024 | 0.0012 | 0.0008 | 0.0002 | 0 |
| **Type** | | **Soluble** | | **Membrane** | |  |  |  |  |  |  |  |  |
| **Likelihood** | | 0.0011 | | 0.9989 | |  |  |  |  |  |  |  |  |

**Hierarchical Tree. Donwload:** [PNG](http://www.cbs.dtu.dk/services/DeepLoc-1.0/tmp/5D01BB8C00003EF54AA12939/tree_2.png) **/** [EPS](http://www.cbs.dtu.dk/services/DeepLoc-1.0/tmp/5D01BB8C00003EF54AA12939/tree_2.eps)
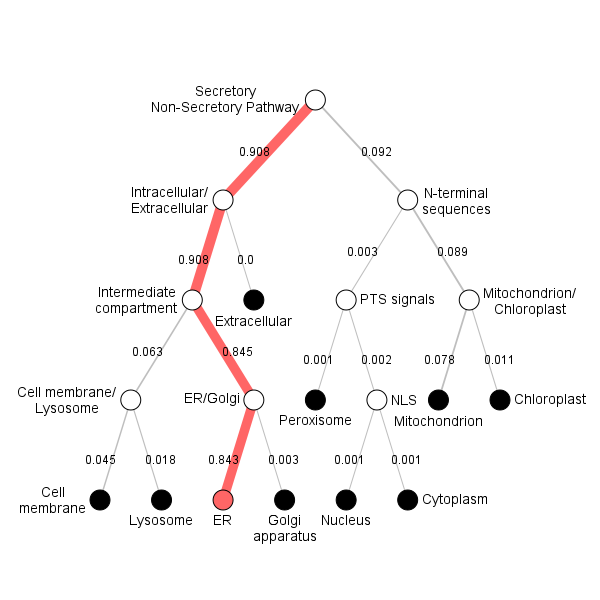

**Position Importance. Donwload:** [PNG](http://www.cbs.dtu.dk/services/DeepLoc-1.0/tmp/5D01BB8C00003EF54AA12939/alpha_2.png) **/** [EPS](http://www.cbs.dtu.dk/services/DeepLoc-1.0/tmp/5D01BB8C00003EF54AA12939/alpha_2.eps) **/** [CSV](http://www.cbs.dtu.dk/services/DeepLoc-1.0/tmp/5D01BB8C00003EF54AA12939/alpha_2.csv)
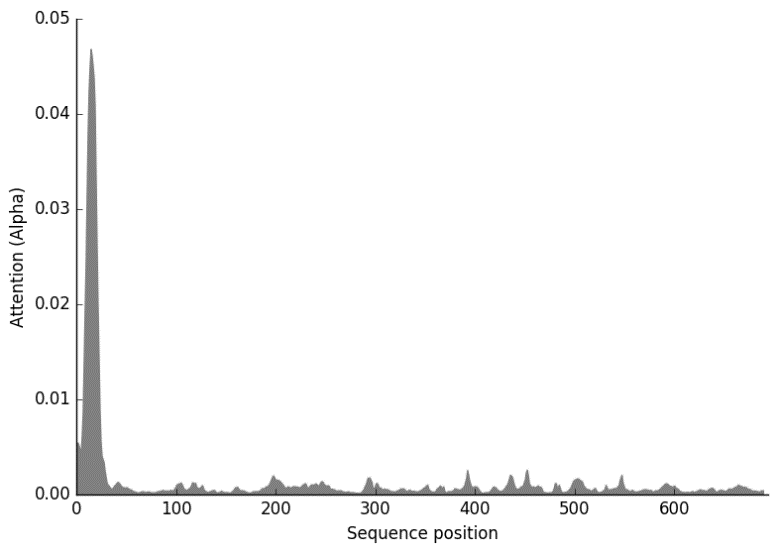

**FgPH-1**

**Prediction: Endoplasmic reticulum, Membrane**

| **Localization** | **Endoplasmic reticulum** | | **Mitochondrion** | | **Cell membrane** | | **Lysosome/Vacuole** | **Plastid** | **Golgi apparatus** | **Cytoplasm** | **Peroxisome** | **Nucleus** | **Extracellular** |
| --- | --- | --- | --- | --- | --- | --- | --- | --- | --- | --- | --- | --- | --- |
| **Likelihood** | 0.9022 | | 0.0472 | | 0.0231 | | 0.0135 | 0.0091 | 0.0035 | 0.0006 | 0.0006 | 0.0001 | 0 |
| **Type** | | **Soluble** | | **Membrane** | |  |  |  |  |  |  |  |  |
| **Likelihood** | | 0.0012 | | 0.9988 | |  |  |  |  |  |  |  |  |

**Hierarchical Tree. Donwload:** [PNG](http://www.cbs.dtu.dk/services/DeepLoc-1.0/tmp/5D01BB8C00003EF54AA12939/tree_1.png) **/** [EPS](http://www.cbs.dtu.dk/services/DeepLoc-1.0/tmp/5D01BB8C00003EF54AA12939/tree_1.eps)
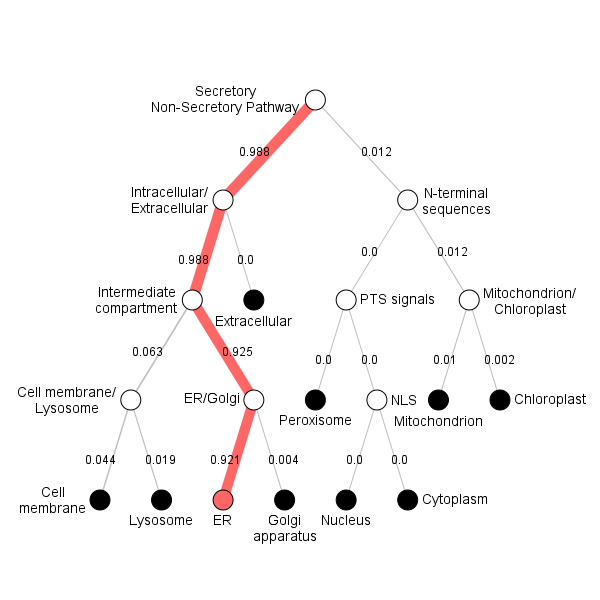

**Position Importance. Donwload:** [PNG](http://www.cbs.dtu.dk/services/DeepLoc-1.0/tmp/5D01BB8C00003EF54AA12939/alpha_1.png) **/** [EPS](http://www.cbs.dtu.dk/services/DeepLoc-1.0/tmp/5D01BB8C00003EF54AA12939/alpha_1.eps) **/** [CSV](http://www.cbs.dtu.dk/services/DeepLoc-1.0/tmp/5D01BB8C00003EF54AA12939/alpha_1.csv)
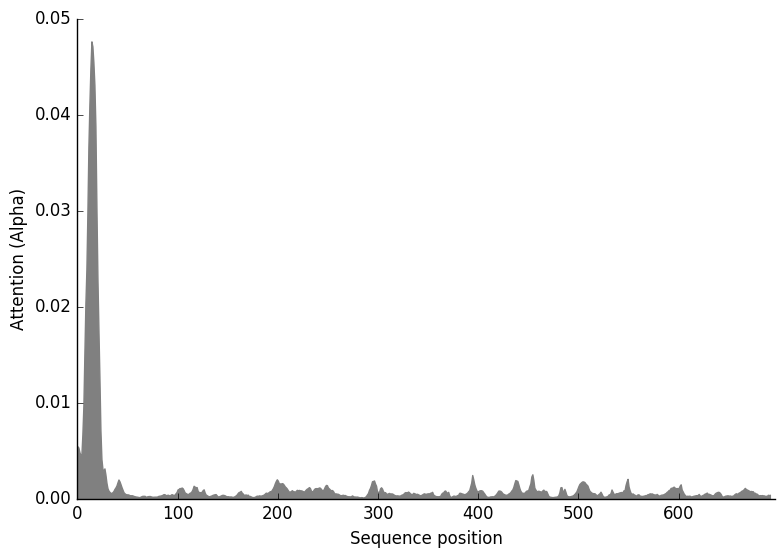

**Homo**

**Prediction: Endoplasmic reticulum, Membrane**

| **Localization** | **Endoplasmic reticulum** | | **Mitochondrion** | | **Peroxisome** | | **Golgi apparatus** | **Cell membrane** | **Lysosome/Vacuole** | **Plastid** | **Nucleus** | **Cytoplasm** | **Extracellular** |
| --- | --- | --- | --- | --- | --- | --- | --- | --- | --- | --- | --- | --- | --- |
| **Likelihood** | 0.874 | | 0.0964 | | 0.0117 | | 0.0098 | 0.0052 | 0.0015 | 0.0009 | 0.0003 | 0.0002 | 0 |
| **Type** | | **Soluble** | | **Membrane** | |  |  |  |  |  |  |  |  |
| **Likelihood** | | 0 | | 1 | |  |  |  |  |  |  |  |  |

**Hierarchical Tree. Donwload:** [PNG](http://www.cbs.dtu.dk/services/DeepLoc-1.0/tmp/5D01BB8C00003EF54AA12939/tree_13.png) **/** [EPS](http://www.cbs.dtu.dk/services/DeepLoc-1.0/tmp/5D01BB8C00003EF54AA12939/tree_13.eps)
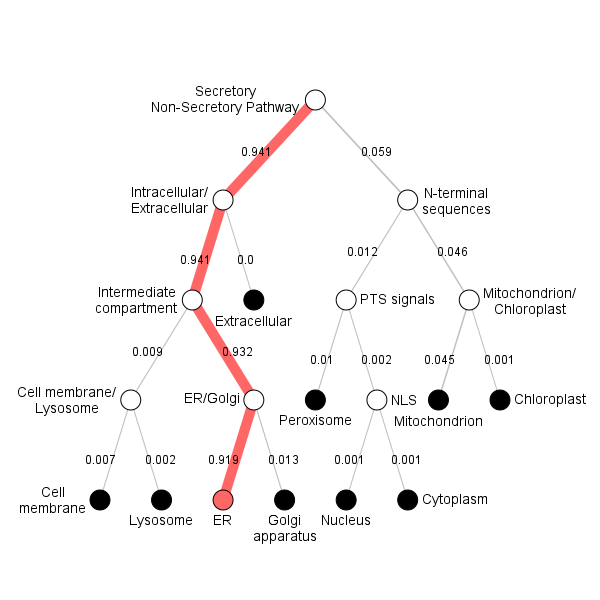

**Position Importance. Donwload:** [PNG](http://www.cbs.dtu.dk/services/DeepLoc-1.0/tmp/5D01BB8C00003EF54AA12939/alpha_13.png) **/** [EPS](http://www.cbs.dtu.dk/services/DeepLoc-1.0/tmp/5D01BB8C00003EF54AA12939/alpha_13.eps) **/** [CSV](http://www.cbs.dtu.dk/services/DeepLoc-1.0/tmp/5D01BB8C00003EF54AA12939/alpha_13.csv)
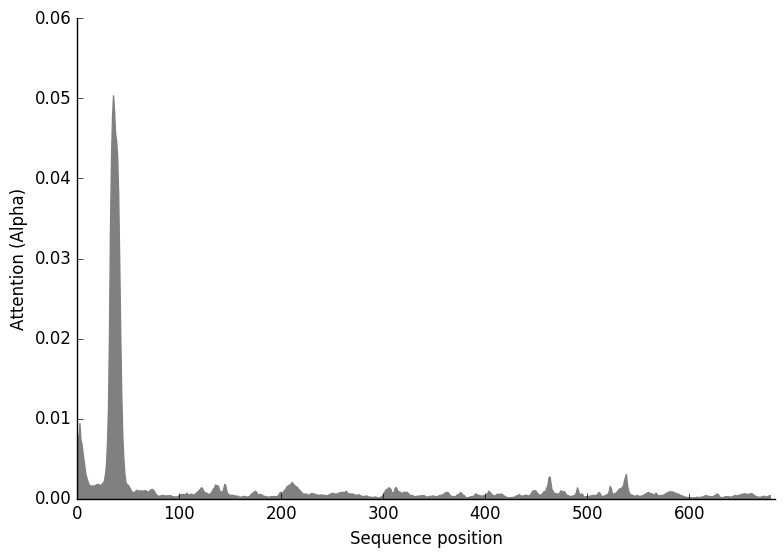

**Verticillium**

**Prediction: Endoplasmic reticulum, Membrane**

| **Localization** | **Endoplasmic reticulum** | | **Mitochondrion** | | **Cell membrane** | | **Lysosome/Vacuole** | **Plastid** | **Golgi apparatus** | **Peroxisome** | **Cytoplasm** | **Nucleus** | **Extracellular** |
| --- | --- | --- | --- | --- | --- | --- | --- | --- | --- | --- | --- | --- | --- |
| **Likelihood** | 0.8853 | | 0.0784 | | 0.0171 | | 0.0091 | 0.0054 | 0.0023 | 0.0014 | 0.0009 | 0.0002 | 0 |
| **Type** | | **Soluble** | | **Membrane** | |  |  |  |  |  |  |  |  |
| **Likelihood** | | 0.0013 | | 0.9987 | |  |  |  |  |  |  |  |  |

**Hierarchical Tree. Donwload:** [PNG](http://www.cbs.dtu.dk/services/DeepLoc-1.0/tmp/5D01BB8C00003EF54AA12939/tree_4.png) **/** [EPS](http://www.cbs.dtu.dk/services/DeepLoc-1.0/tmp/5D01BB8C00003EF54AA12939/tree_4.eps)
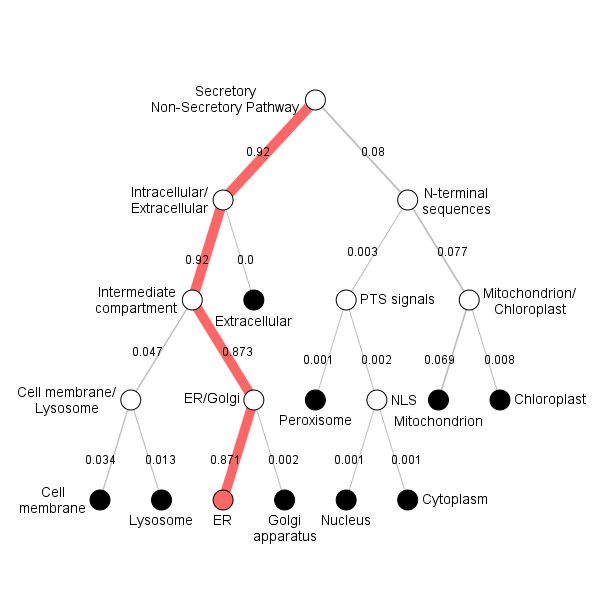

**Position Importance. Donwload:** [PNG](http://www.cbs.dtu.dk/services/DeepLoc-1.0/tmp/5D01BB8C00003EF54AA12939/alpha_4.png) **/** [EPS](http://www.cbs.dtu.dk/services/DeepLoc-1.0/tmp/5D01BB8C00003EF54AA12939/alpha_4.eps) **/** [CSV](http://www.cbs.dtu.dk/services/DeepLoc-1.0/tmp/5D01BB8C00003EF54AA12939/alpha_4.csv)
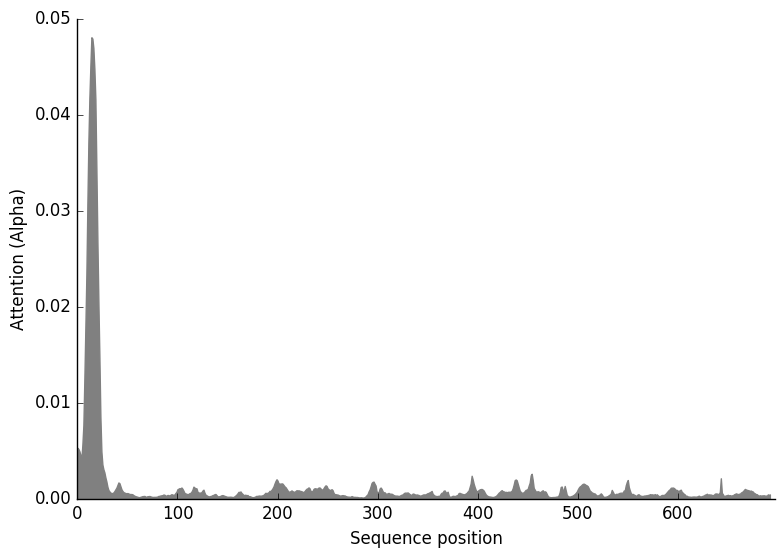

**Gallus**

**Prediction: Endoplasmic reticulum, Membrane**

| **Localization** | **Endoplasmic reticulum** | | **Mitochondrion** | | **Peroxisome** | | **Cell membrane** | **Lysosome/Vacuole** | **Golgi apparatus** | **Plastid** | **Nucleus** | **Cytoplasm** | **Extracellular** |
| --- | --- | --- | --- | --- | --- | --- | --- | --- | --- | --- | --- | --- | --- |
| **Likelihood** | 0.898 | | 0.0626 | | 0.0229 | | 0.0085 | 0.0032 | 0.0027 | 0.0013 | 0.0004 | 0.0003 | 0 |
| **Type** | | **Soluble** | | **Membrane** | |  |  |  |  |  |  |  |  |
| **Likelihood** | | 0.0001 | | 0.9999 | |  |  |  |  |  |  |  |  |

**Hierarchical Tree. Donwload:** [PNG](http://www.cbs.dtu.dk/services/DeepLoc-1.0/tmp/5D01BB8C00003EF54AA12939/tree_14.png) **/** [EPS](http://www.cbs.dtu.dk/services/DeepLoc-1.0/tmp/5D01BB8C00003EF54AA12939/tree_14.eps)
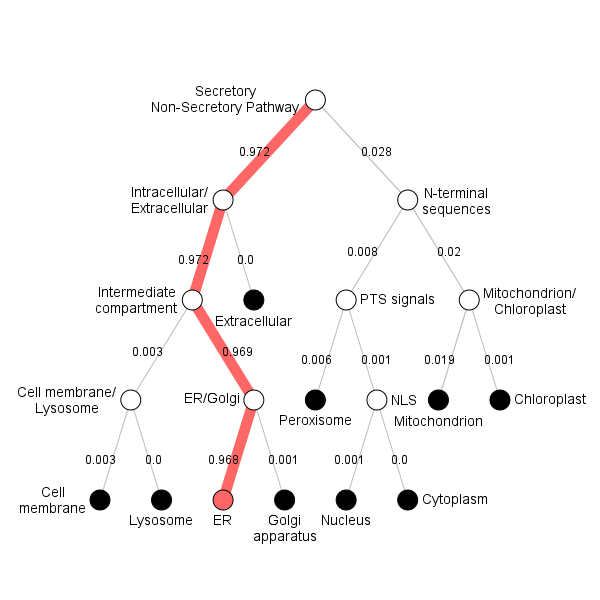

**Position Importance. Donwload:** [PNG](http://www.cbs.dtu.dk/services/DeepLoc-1.0/tmp/5D01BB8C00003EF54AA12939/alpha_14.png) **/** [EPS](http://www.cbs.dtu.dk/services/DeepLoc-1.0/tmp/5D01BB8C00003EF54AA12939/alpha_14.eps) **/** [CSV](http://www.cbs.dtu.dk/services/DeepLoc-1.0/tmp/5D01BB8C00003EF54AA12939/alpha_14.csv)
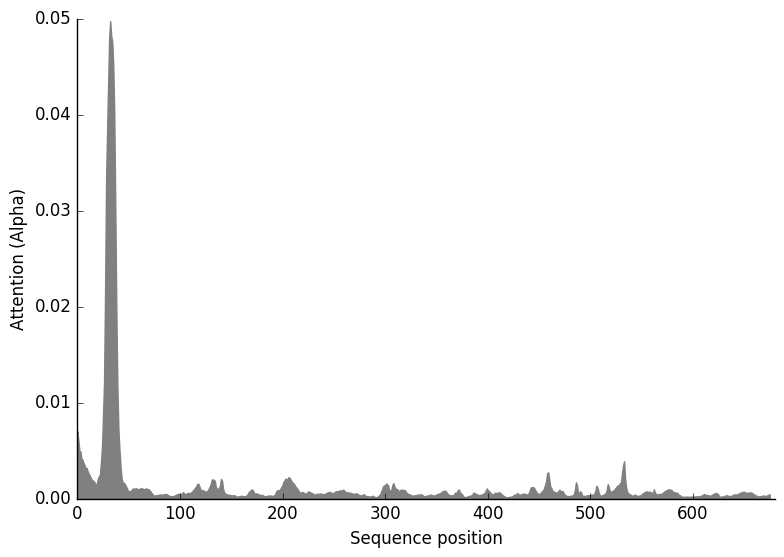

**Serpula**

**Prediction: Endoplasmic reticulum, Membrane**

| **Localization** | **Endoplasmic reticulum** | | **Mitochondrion** | | **Cytoplasm** | | **Cell membrane** | **Peroxisome** | **Lysosome/Vacuole** | **Plastid** | **Golgi apparatus** | **Nucleus** | **Extracellular** |
| --- | --- | --- | --- | --- | --- | --- | --- | --- | --- | --- | --- | --- | --- |
| **Likelihood** | 0.8591 | | 0.0711 | | 0.0323 | | 0.0163 | 0.0077 | 0.0052 | 0.0048 | 0.0027 | 0.0007 | 0 |
| **Type** | | **Soluble** | | **Membrane** | |  |  |  |  |  |  |  |  |
| **Likelihood** | | 0.048 | | 0.952 | |  |  |  |  |  |  |  |  |

**Hierarchical Tree. Donwload:** [PNG](http://www.cbs.dtu.dk/services/DeepLoc-1.0/tmp/5D01BB8C00003EF54AA12939/tree_10.png) **/** [EPS](http://www.cbs.dtu.dk/services/DeepLoc-1.0/tmp/5D01BB8C00003EF54AA12939/tree_10.eps)
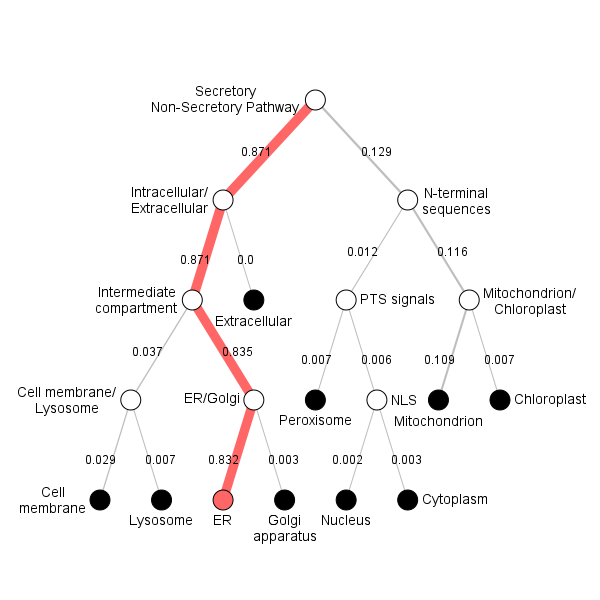

**Position Importance. Donwload:** [PNG](http://www.cbs.dtu.dk/services/DeepLoc-1.0/tmp/5D01BB8C00003EF54AA12939/alpha_10.png) **/** [EPS](http://www.cbs.dtu.dk/services/DeepLoc-1.0/tmp/5D01BB8C00003EF54AA12939/alpha_10.eps) **/** [CSV](http://www.cbs.dtu.dk/services/DeepLoc-1.0/tmp/5D01BB8C00003EF54AA12939/alpha_10.csv)
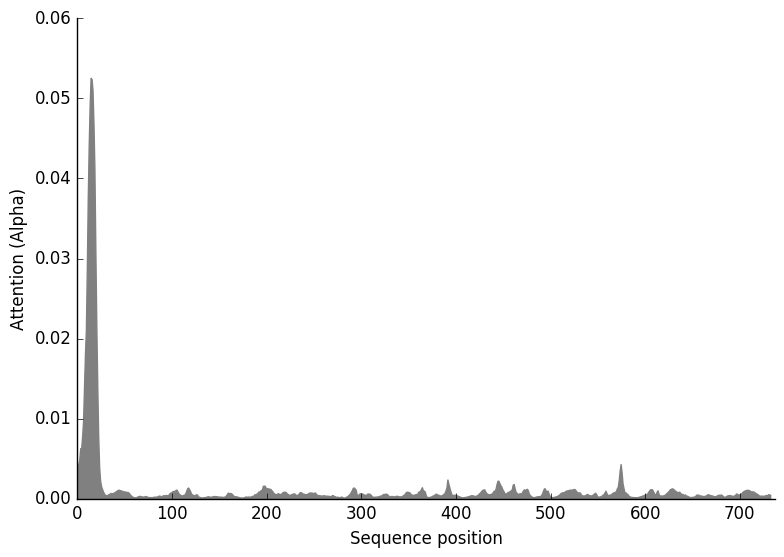

**Agaricus**

**Prediction: Endoplasmic reticulum, Membrane**

| **Localization** | **Endoplasmic reticulum** | | **Mitochondrion** | | **Cell membrane** | | **Plastid** | **Peroxisome** | **Lysosome/Vacuole** | **Golgi apparatus** | **Cytoplasm** | **Nucleus** | **Extracellular** |
| --- | --- | --- | --- | --- | --- | --- | --- | --- | --- | --- | --- | --- | --- |
| **Likelihood** | 0.8746 | | 0.0831 | | 0.0188 | | 0.0077 | 0.006 | 0.0055 | 0.0023 | 0.0018 | 0.0002 | 0 |
| **Type** | | **Soluble** | | **Membrane** | |  |  |  |  |  |  |  |  |
| **Likelihood** | | 0.0016 | | 0.9984 | |  |  |  |  |  |  |  |  |

**Hierarchical Tree. Donwload:** [PNG](http://www.cbs.dtu.dk/services/DeepLoc-1.0/tmp/5D01BB8C00003EF54AA12939/tree_9.png) **/** [EPS](http://www.cbs.dtu.dk/services/DeepLoc-1.0/tmp/5D01BB8C00003EF54AA12939/tree_9.eps)
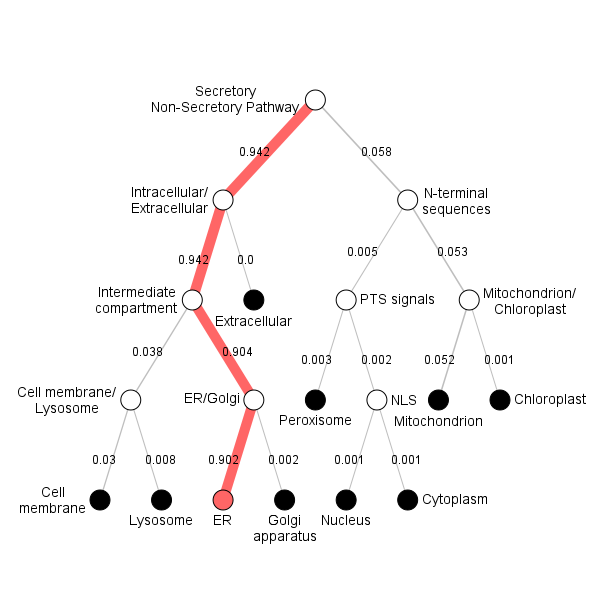

**Position Importance. Donwload:** [PNG](http://www.cbs.dtu.dk/services/DeepLoc-1.0/tmp/5D01BB8C00003EF54AA12939/alpha_9.png) **/** [EPS](http://www.cbs.dtu.dk/services/DeepLoc-1.0/tmp/5D01BB8C00003EF54AA12939/alpha_9.eps) **/** [CSV](http://www.cbs.dtu.dk/services/DeepLoc-1.0/tmp/5D01BB8C00003EF54AA12939/alpha_9.csv)
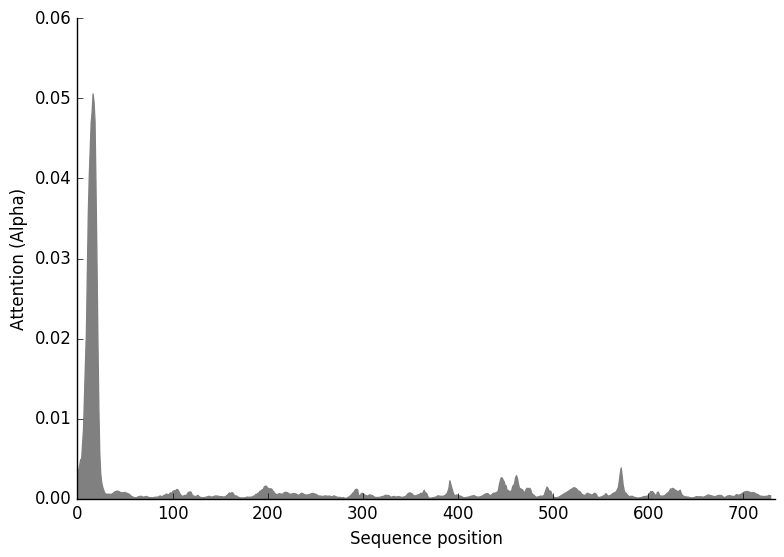

**Arabidopsis_thaliana1**

**Prediction: Endoplasmic reticulum, Membrane**

| **Localization** | **Endoplasmic reticulum** | | **Mitochondrion** | | **Peroxisome** | | **Plastid** | **Cytoplasm** | **Cell membrane** | **Golgi apparatus** | **Nucleus** | **Lysosome/Vacuole** | **Extracellular** |
| --- | --- | --- | --- | --- | --- | --- | --- | --- | --- | --- | --- | --- | --- |
| **Likelihood** | 0.4549 | | 0.3586 | | 0.1315 | | 0.0312 | 0.0107 | 0.0043 | 0.0032 | 0.003 | 0.0025 | 0 |
| **Type** | | **Soluble** | | **Membrane** | |  |  |  |  |  |  |  |  |
| **Likelihood** | | 0.0069 | | 0.9931 | |  |  |  |  |  |  |  |  |

**Hierarchical Tree. Donwload:** [PNG](http://www.cbs.dtu.dk/services/DeepLoc-1.0/tmp/5D01BB8C00003EF54AA12939/tree_16.png) **/** [EPS](http://www.cbs.dtu.dk/services/DeepLoc-1.0/tmp/5D01BB8C00003EF54AA12939/tree_16.eps)
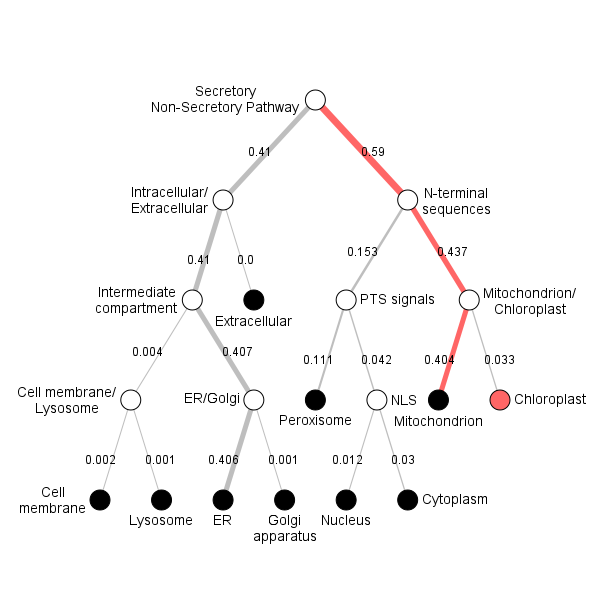

**Position Importance. Donwload:** [PNG](http://www.cbs.dtu.dk/services/DeepLoc-1.0/tmp/5D01BB8C00003EF54AA12939/alpha_16.png) **/** [EPS](http://www.cbs.dtu.dk/services/DeepLoc-1.0/tmp/5D01BB8C00003EF54AA12939/alpha_16.eps) **/** [CSV](http://www.cbs.dtu.dk/services/DeepLoc-1.0/tmp/5D01BB8C00003EF54AA12939/alpha_16.csv)
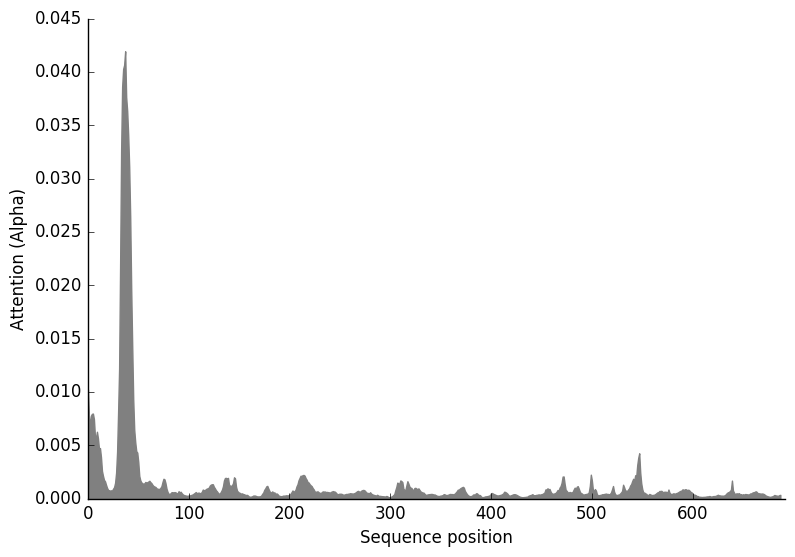

**Trypanosoma**

**Prediction: Endoplasmic reticulum, Soluble**

| **Localization** | **Endoplasmic reticulum** | | **Extracellular** | | **Plastid** | | **Mitochondrion** | **Cytoplasm** | **Lysosome/Vacuole** | **Cell membrane** | **Peroxisome** | **Nucleus** | **Golgi apparatus** |
| --- | --- | --- | --- | --- | --- | --- | --- | --- | --- | --- | --- | --- | --- |
| **Likelihood** | 0.4271 | | 0.2822 | | 0.118 | | 0.1038 | 0.0413 | 0.0169 | 0.0053 | 0.0031 | 0.0012 | 0.001 |
| **Type** | | **Soluble** | | **Membrane** | |  |  |  |  |  |  |  |  |
| **Likelihood** | | 0.6333 | | 0.3667 | |  |  |  |  |  |  |  |  |

**Hierarchical Tree. Donwload:** [PNG](http://www.cbs.dtu.dk/services/DeepLoc-1.0/tmp/5D01BB8C00003EF54AA12939/tree_19.png) **/** [EPS](http://www.cbs.dtu.dk/services/DeepLoc-1.0/tmp/5D01BB8C00003EF54AA12939/tree_19.eps)
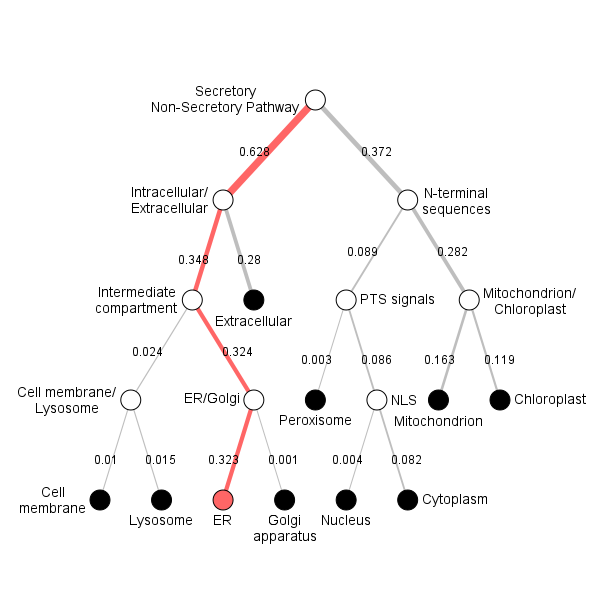

**Position Importance. Donwload:** [PNG](http://www.cbs.dtu.dk/services/DeepLoc-1.0/tmp/5D01BB8C00003EF54AA12939/alpha_19.png) **/** [EPS](http://www.cbs.dtu.dk/services/DeepLoc-1.0/tmp/5D01BB8C00003EF54AA12939/alpha_19.eps) **/** [CSV](http://www.cbs.dtu.dk/services/DeepLoc-1.0/tmp/5D01BB8C00003EF54AA12939/alpha_19.csv)
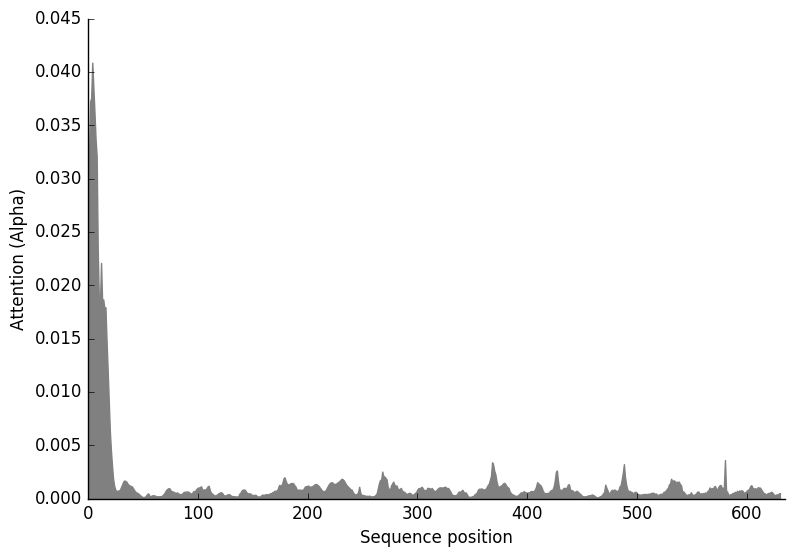

**Arabidopsis_thaliana**

**Prediction: Cytoplasm, Soluble**

| **Localization** | **Cytoplasm** | | **Mitochondrion** | | **Peroxisome** | | **Plastid** | **Cell membrane** | **Lysosome/Vacuole** | **Nucleus** | **Golgi apparatus** | **Extracellular** | **Endoplasmic reticulum** |
| --- | --- | --- | --- | --- | --- | --- | --- | --- | --- | --- | --- | --- | --- |
| **Likelihood** | 0.3794 | | 0.2456 | | 0.1414 | | 0.0569 | 0.0498 | 0.0392 | 0.0297 | 0.0282 | 0.0221 | 0.0076 |
| **Type** | | **Soluble** | | **Membrane** | |  |  |  |  |  |  |  |  |
| **Likelihood** | | 0.7908 | | 0.2092 | |  |  |  |  |  |  |  |  |

**Hierarchical Tree. Donwload:** [PNG](http://www.cbs.dtu.dk/services/DeepLoc-1.0/tmp/5D01BB8C00003EF54AA12939/tree_17.png) **/** [EPS](http://www.cbs.dtu.dk/services/DeepLoc-1.0/tmp/5D01BB8C00003EF54AA12939/tree_17.eps)
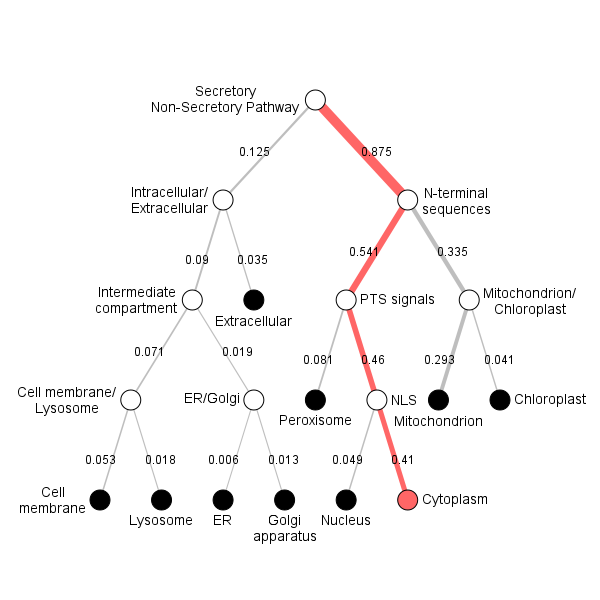

**Position Importance. Donwload:** [PNG](http://www.cbs.dtu.dk/services/DeepLoc-1.0/tmp/5D01BB8C00003EF54AA12939/alpha_17.png) **/** [EPS](http://www.cbs.dtu.dk/services/DeepLoc-1.0/tmp/5D01BB8C00003EF54AA12939/alpha_17.eps) **/** [CSV](http://www.cbs.dtu.dk/services/DeepLoc-1.0/tmp/5D01BB8C00003EF54AA12939/alpha_17.csv)
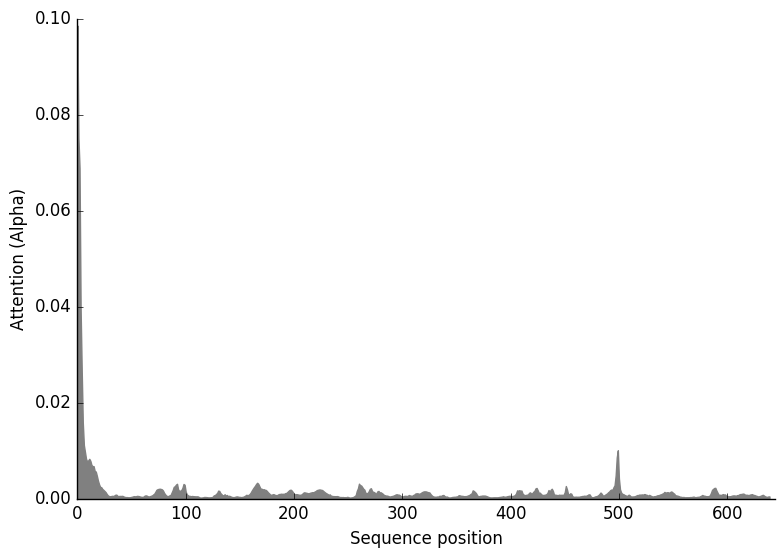

**Triticum**

**Prediction: Mitochondrion, Membrane**

| **Localization** | **Mitochondrion** | | **Plastid** | | **Peroxisome** | | **Endoplasmic reticulum** | **Cytoplasm** | **Lysosome/Vacuole** | **Golgi apparatus** | **Cell membrane** | **Nucleus** | **Extracellular** |
| --- | --- | --- | --- | --- | --- | --- | --- | --- | --- | --- | --- | --- | --- |
| **Likelihood** | 0.2445 | | 0.2228 | | 0.2215 | | 0.1738 | 0.0882 | 0.0162 | 0.0159 | 0.0094 | 0.0068 | 0.0009 |
| **Type** | | **Soluble** | | **Membrane** | |  |  |  |  |  |  |  |  |
| **Likelihood** | | 0.1726 | | 0.8274 | |  |  |  |  |  |  |  |  |

**Hierarchical Tree. Donwload:** [PNG](http://www.cbs.dtu.dk/services/DeepLoc-1.0/tmp/5D01BB8C00003EF54AA12939/tree_18.png) **/** [EPS](http://www.cbs.dtu.dk/services/DeepLoc-1.0/tmp/5D01BB8C00003EF54AA12939/tree_18.eps)
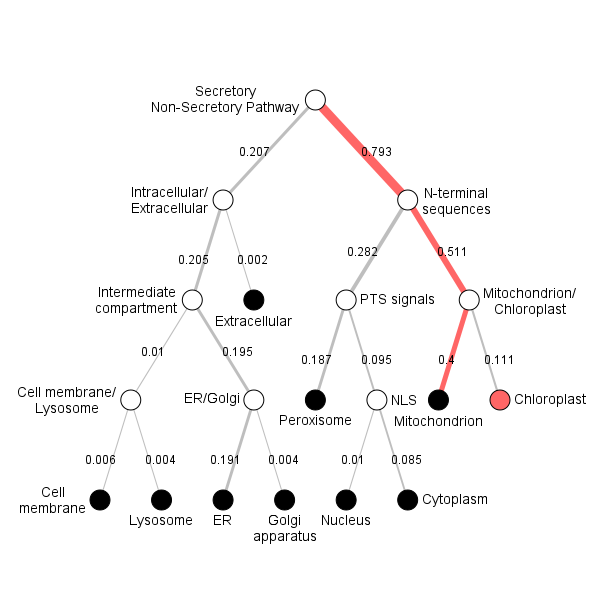

**Position Importance. Donwload:** [PNG](http://www.cbs.dtu.dk/services/DeepLoc-1.0/tmp/5D01BB8C00003EF54AA12939/alpha_18.png) **/** [EPS](http://www.cbs.dtu.dk/services/DeepLoc-1.0/tmp/5D01BB8C00003EF54AA12939/alpha_18.eps) **/** [CSV](http://www.cbs.dtu.dk/services/DeepLoc-1.0/tmp/5D01BB8C00003EF54AA12939/alpha_18.csv)
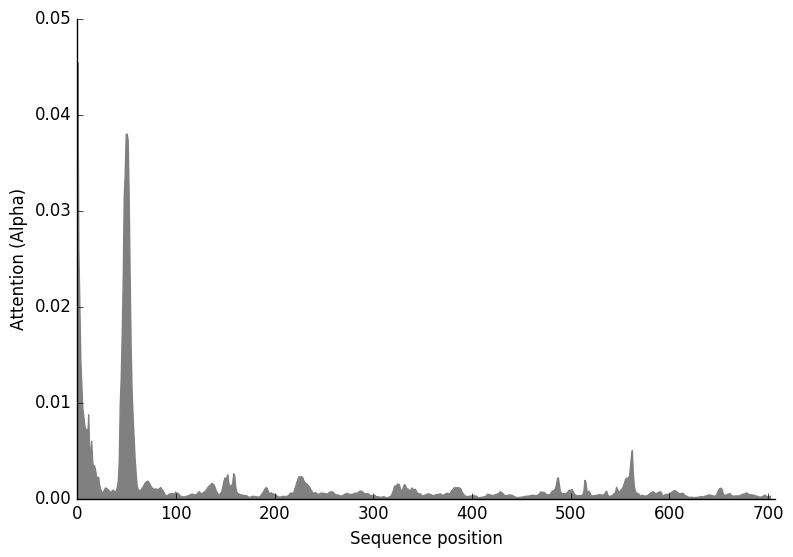

CAM prediction using CaMELS (CalModulin intEraction Learning System) <https://camels.pythonanywhere.com/> and domain structure prediction using HMMER

Ph1

Interaction prediction= 0.26 NO

Binding site Prediction

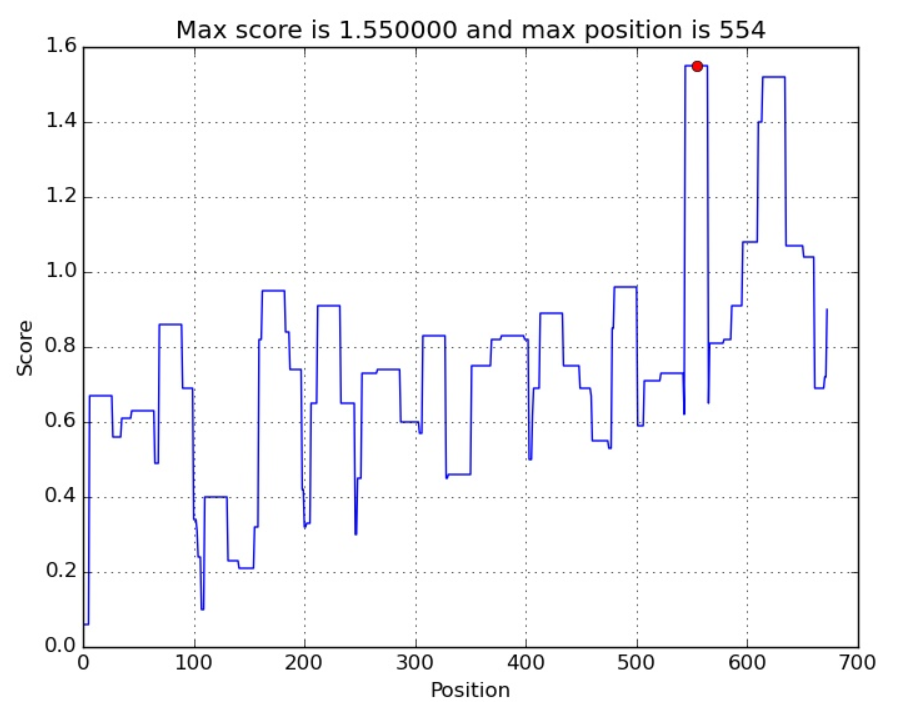

Magnaporthe oryzae

Interaction prediction= 0.33 NO

Binding site Prediction

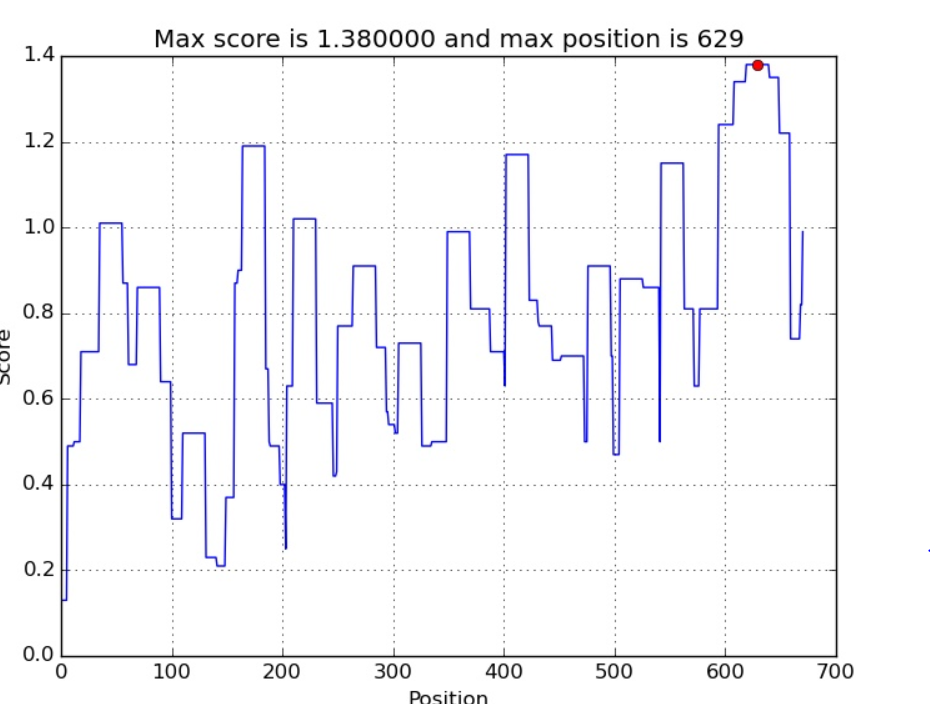

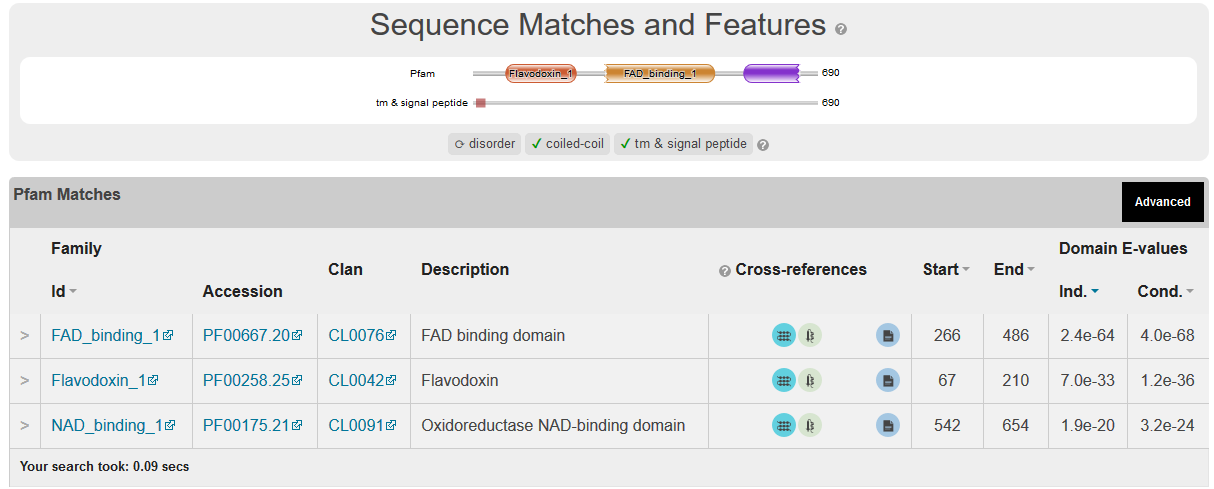

Metarhizium anisopliae

Interaction prediction=0.55 NO

Binding site Prediction

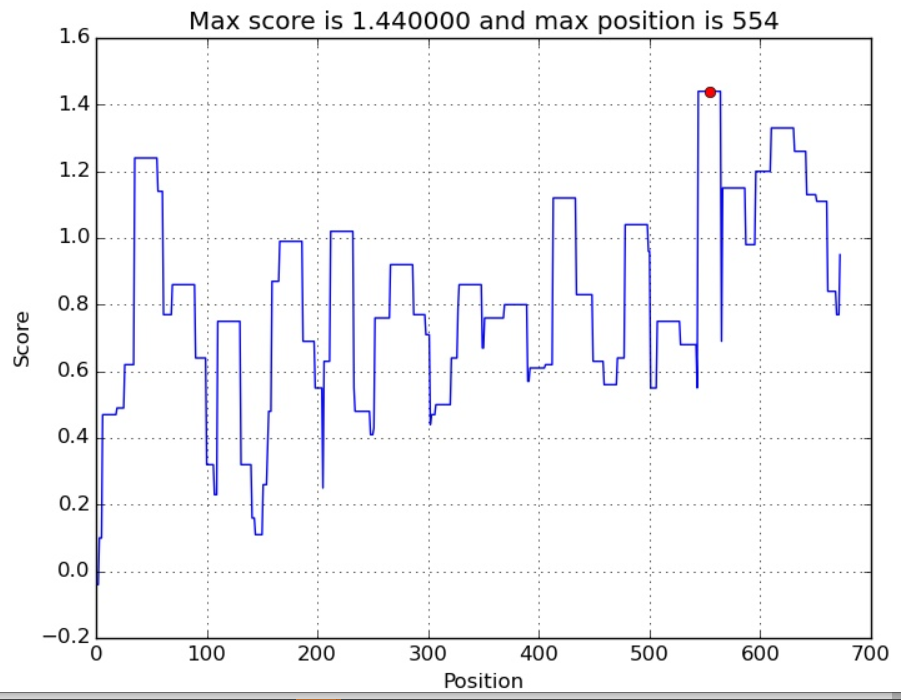

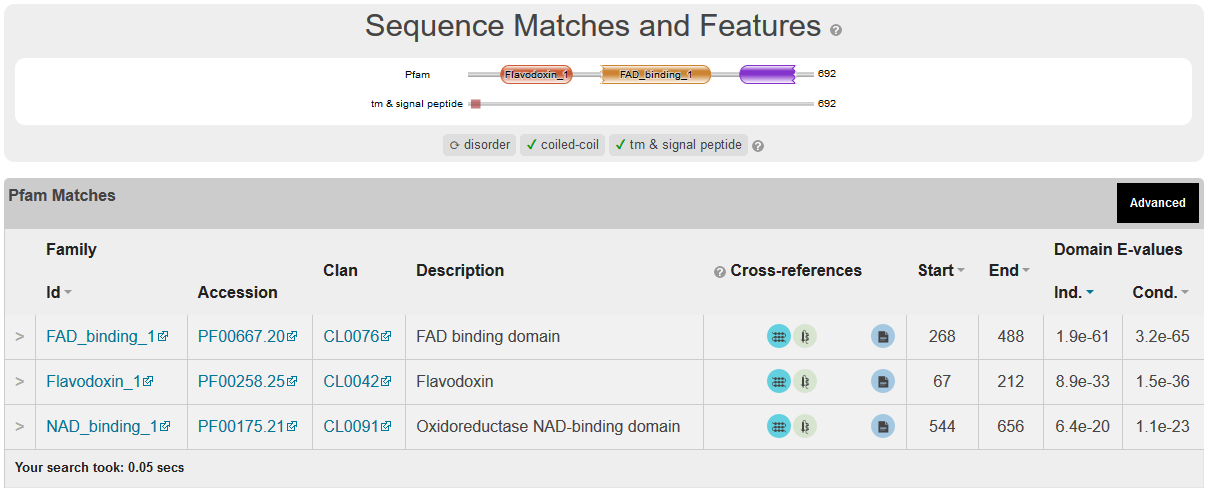

Verticillium dahliae

Interaction prediction= 0.46 NO

Binding site Prediction

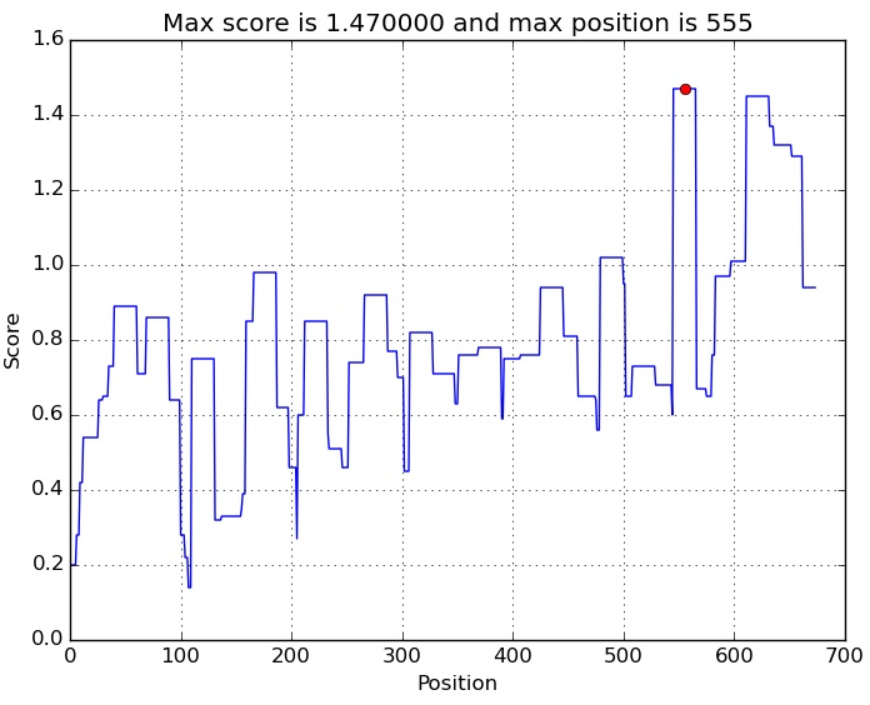

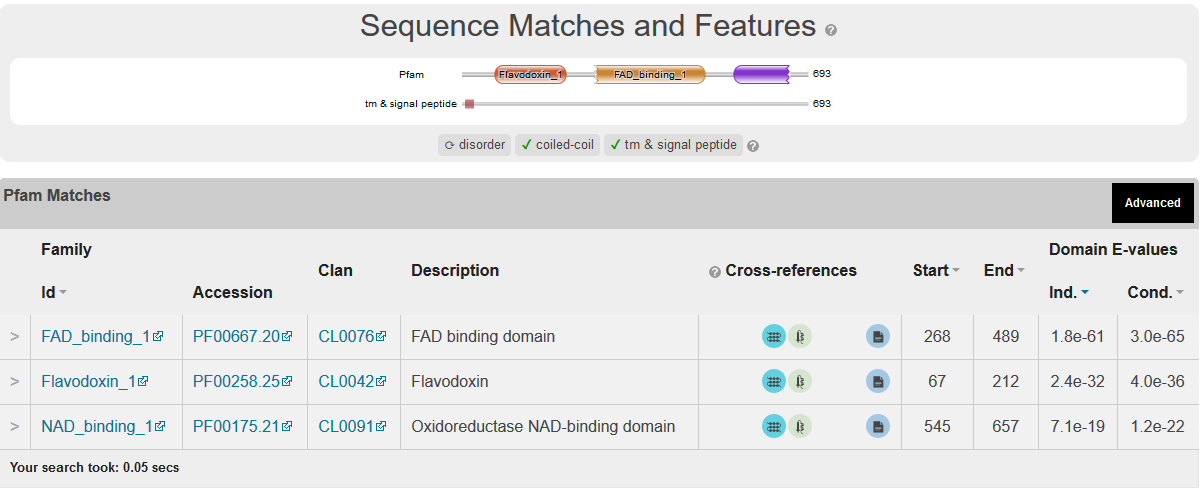

Fusarium oxysporum f. sp. vasinfectum

Interaction prediction= 0.86 NO

Binding site Prediction

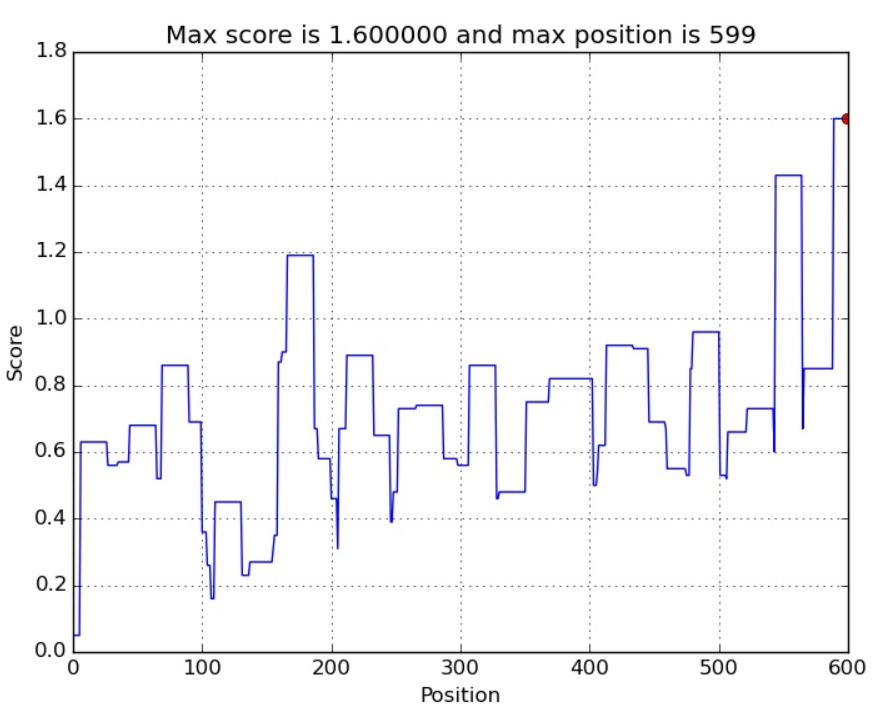

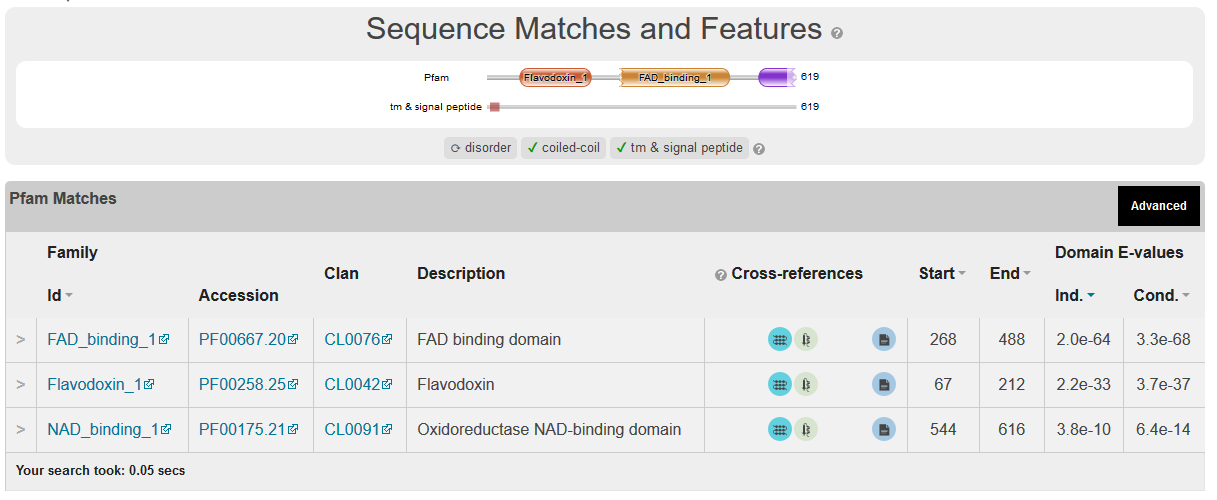

Aspergillus niger CBS

Interaction prediction= 0.29 NO

Binding site Prediction

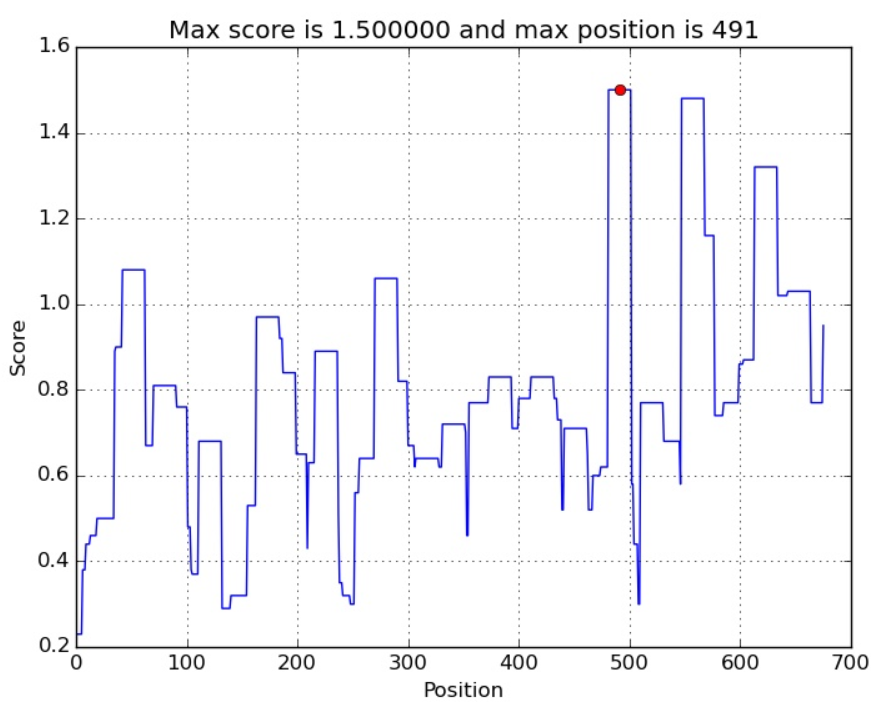

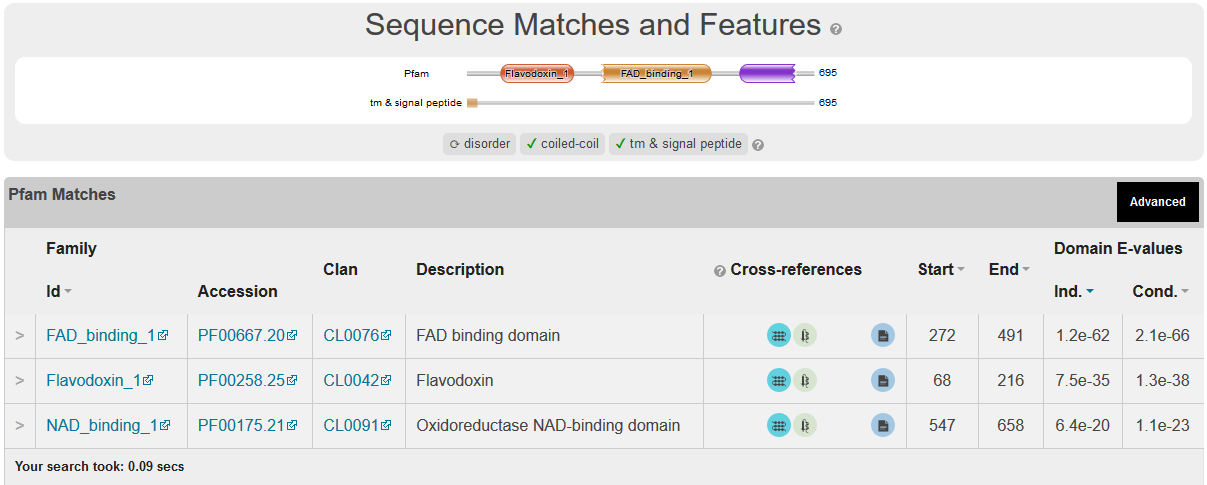

Blumeria graminis

Interaction prediction= 0.95 NO

Binding site Prediction

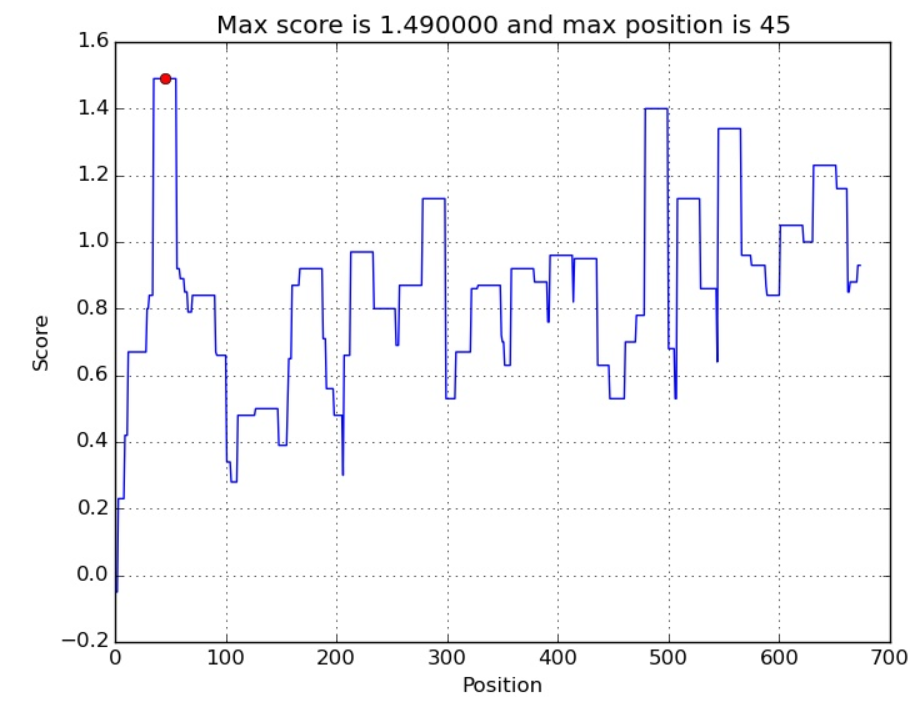

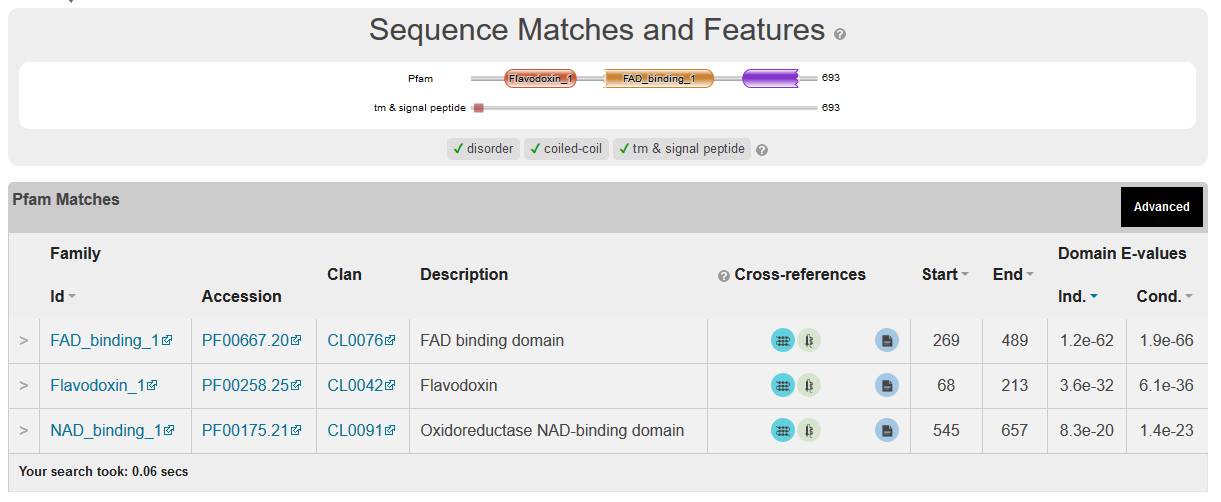

>Crucibulum leave

Interaction prediction=0.7NO

Binding site Prediction
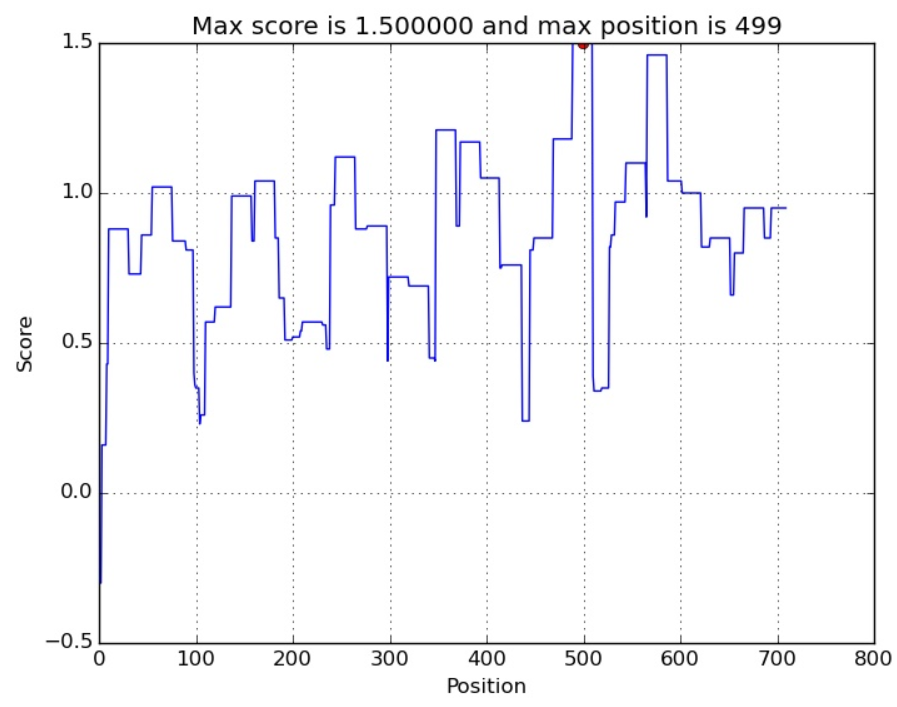

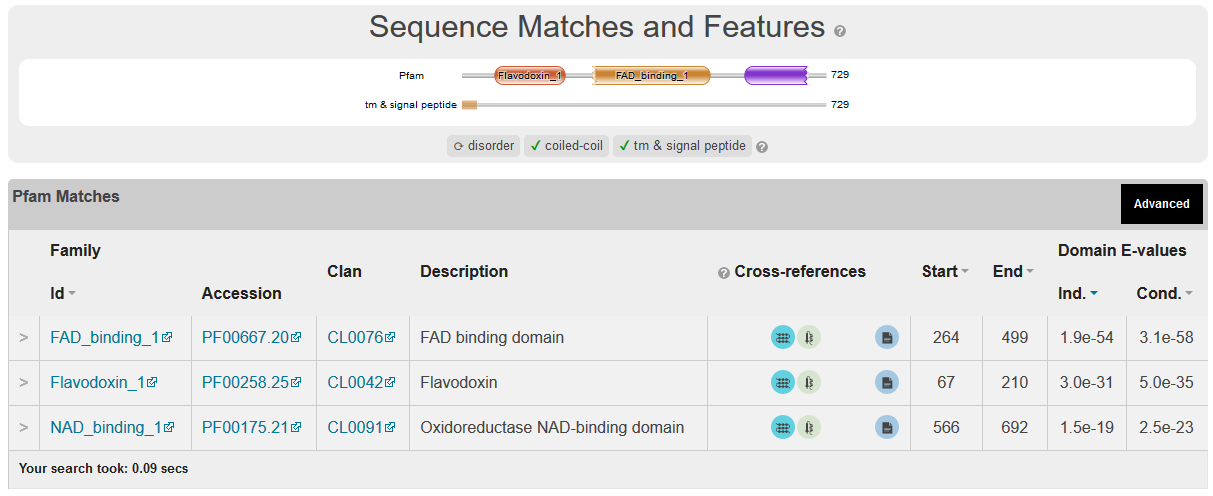

Agaricus bisporus

Interaction prediction=3.49 YES

Binding site Prediction

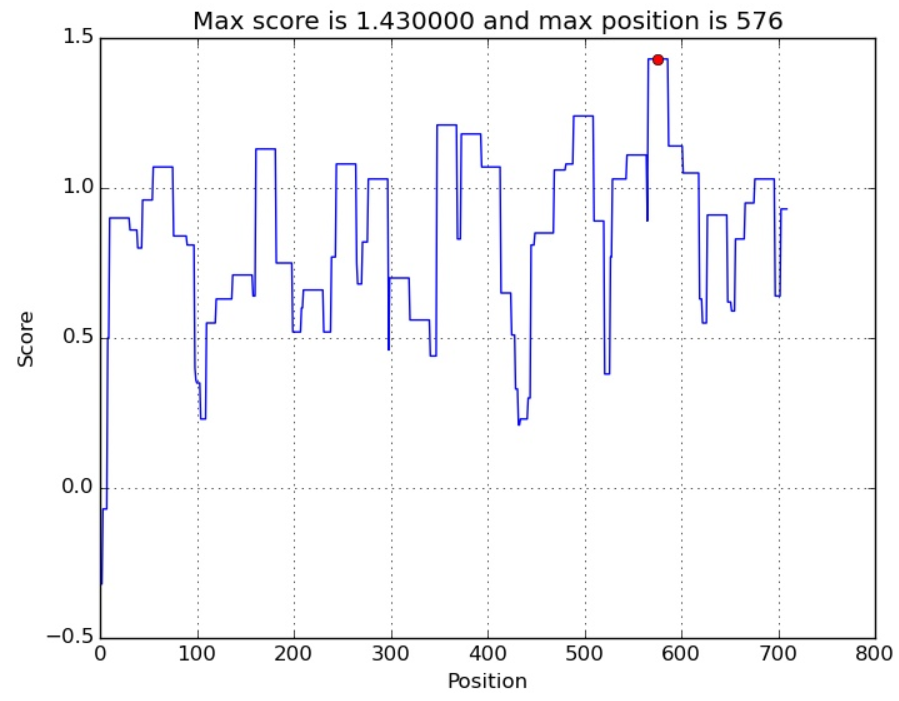

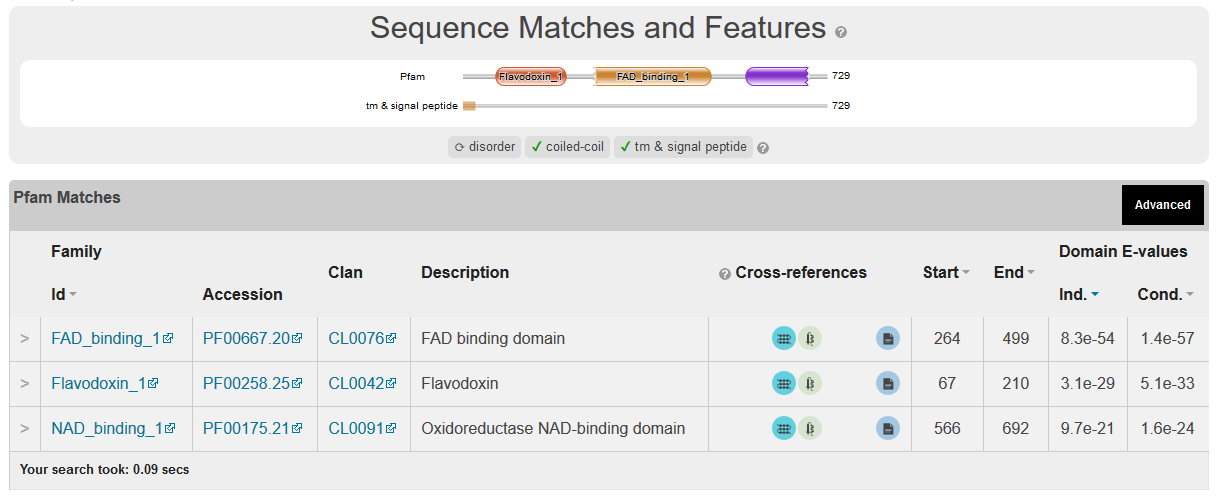

>Serpula lacrymans

Interaction prediction=2.57 YES

Binding site Prediction

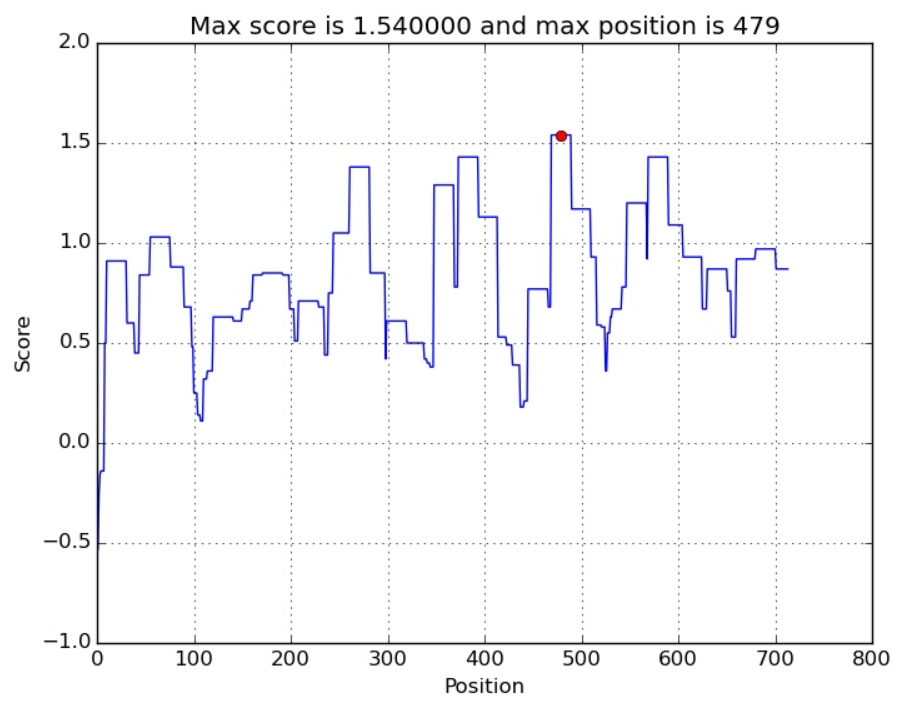

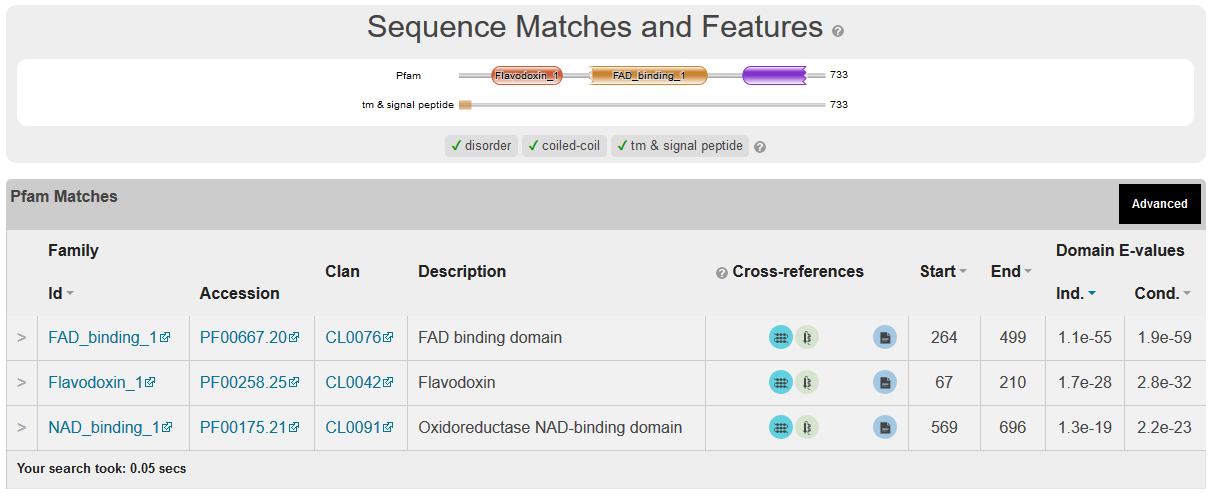

>Pleurotus ostreatus

Interaction prediction= 2.11 YES

Binding site Prediction

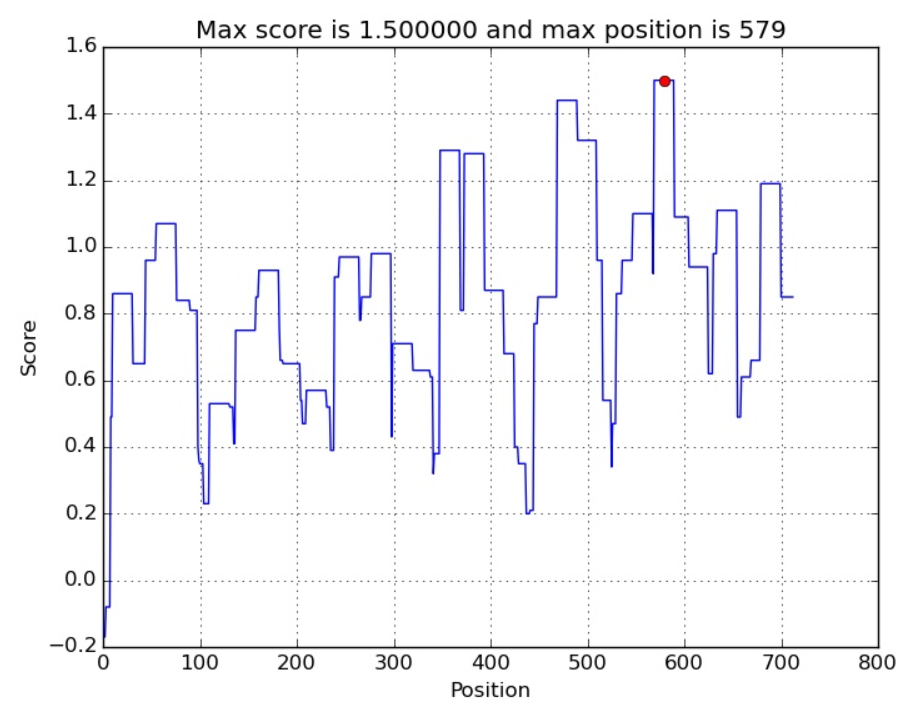

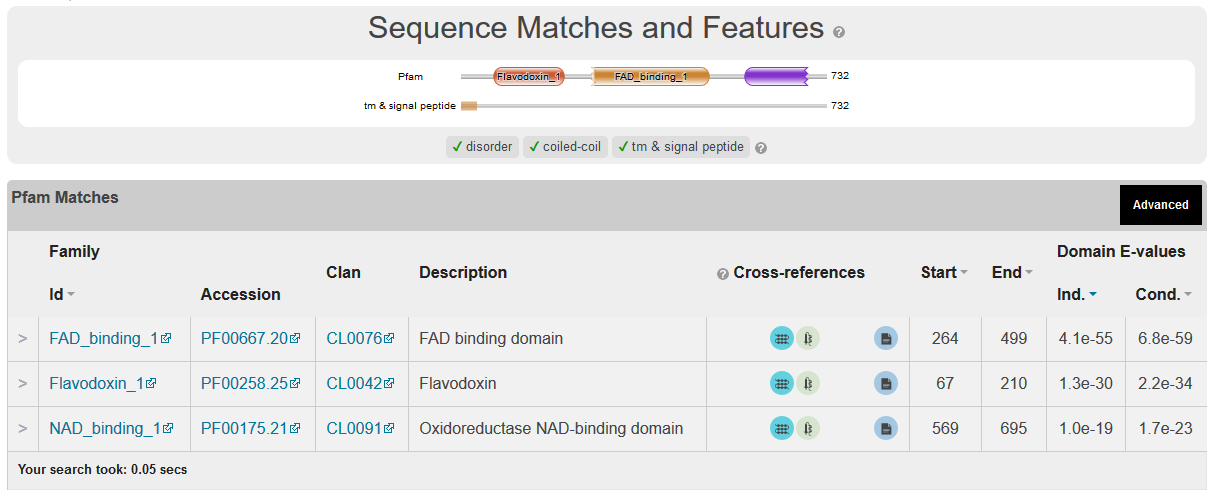

>Drosophila melanogaster

Interaction prediction= 0.36 NO

Binding site Prediction

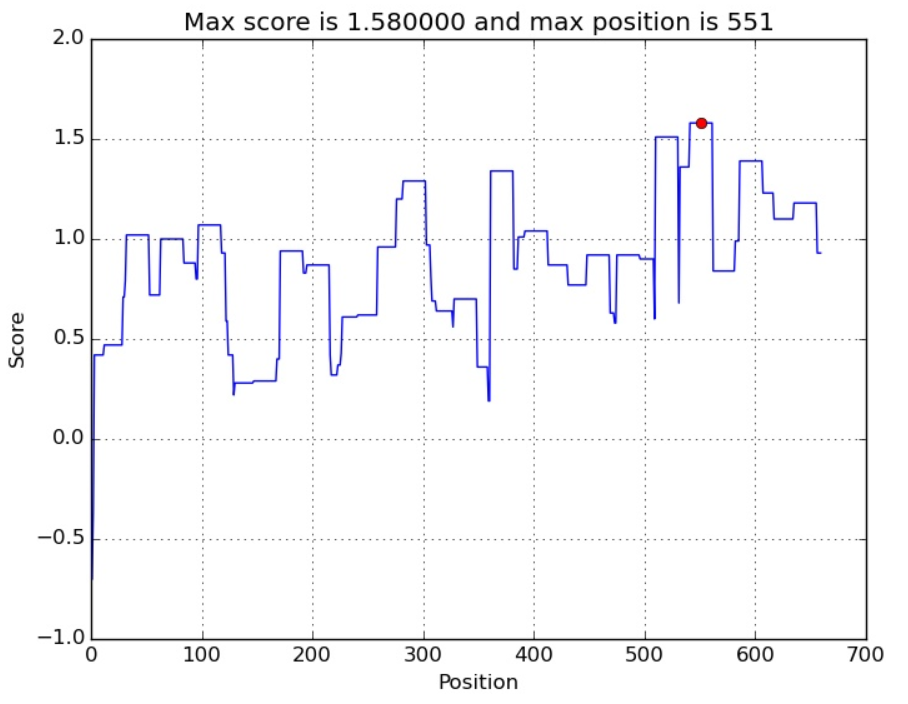

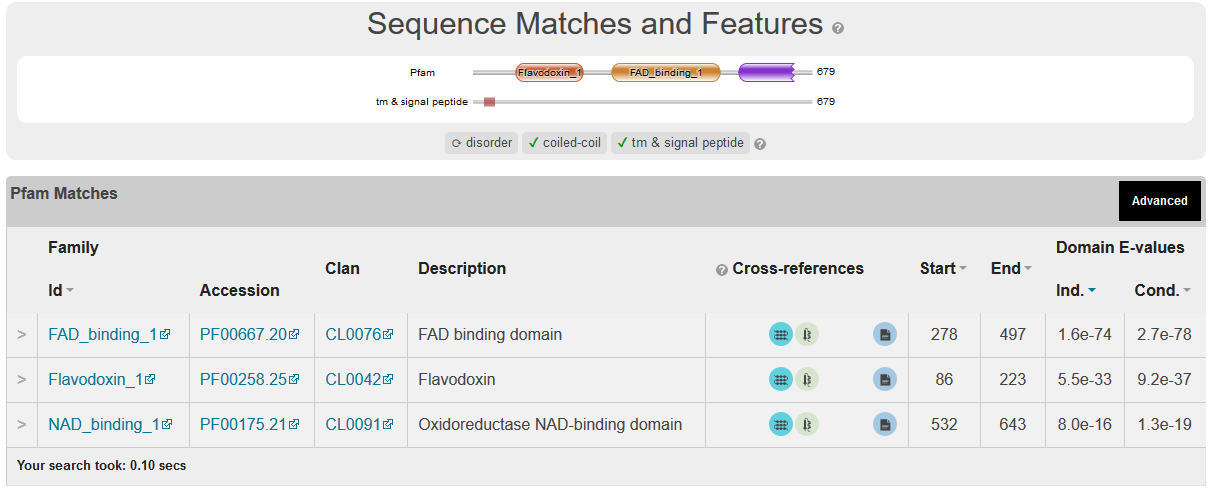

>Homo sapiens

Interaction prediction= 0.78 NO

Binding site Prediction

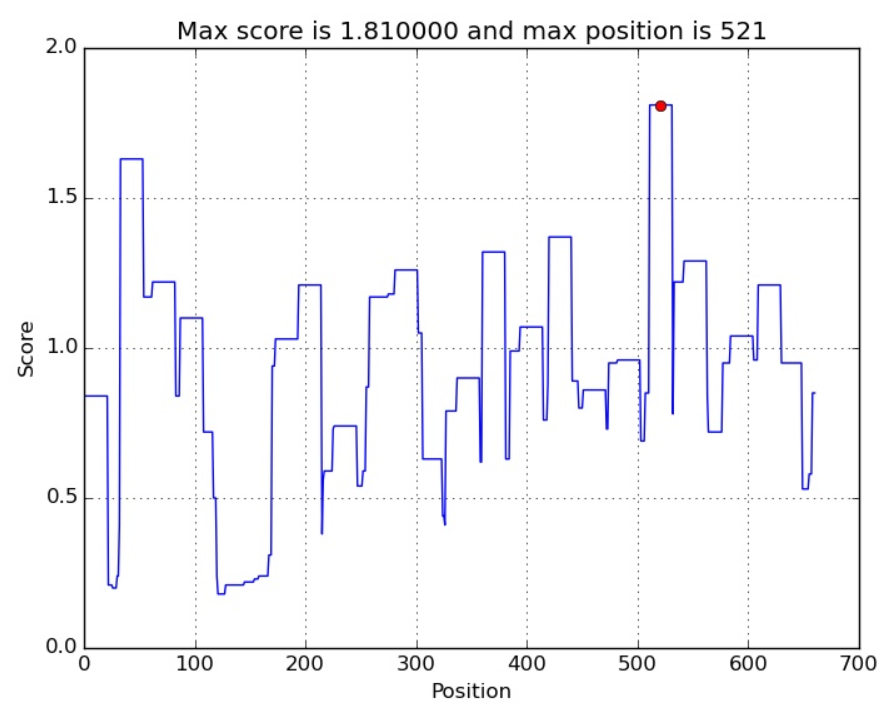

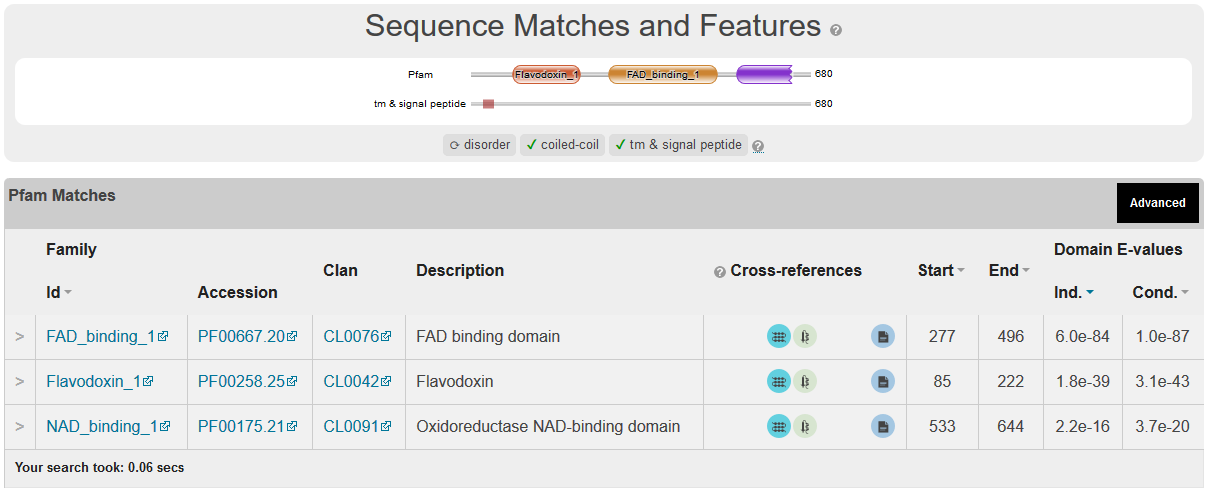

>Gallus gallus

Interaction prediction= 2.24 YES

Binding site Prediction

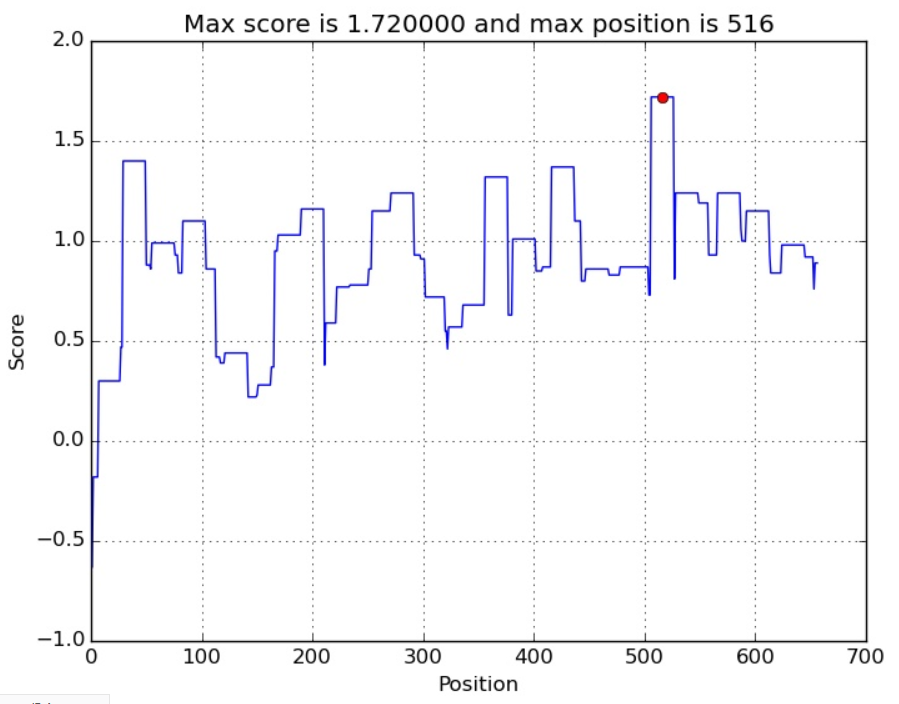

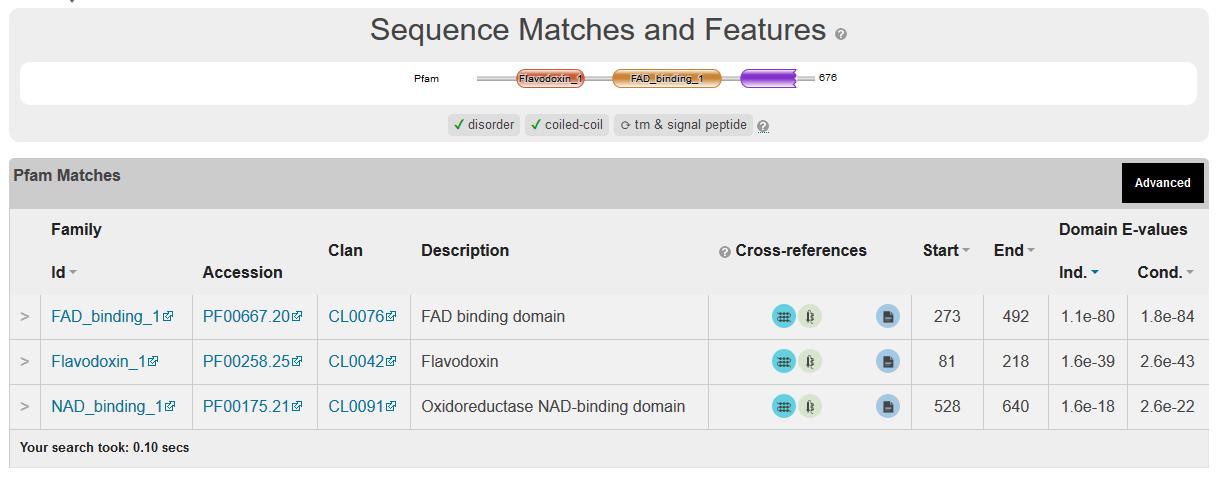

>Musca domestica

Interaction prediction= 0.78 NO

Binding site Prediction

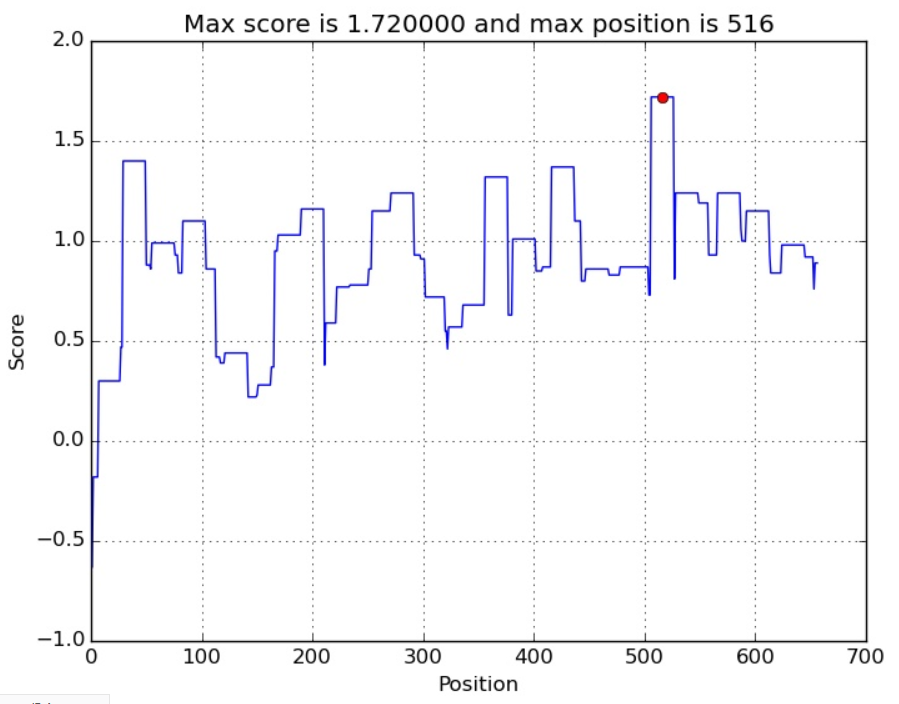

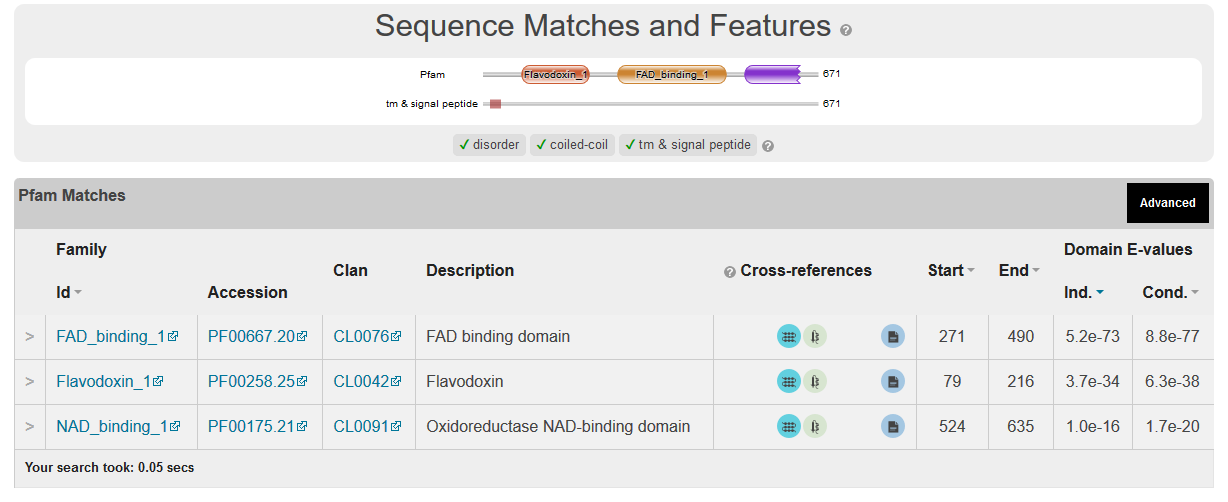

>Arabidopsis thaliana 1

Interaction prediction= 1.89 YES

Binding site Prediction

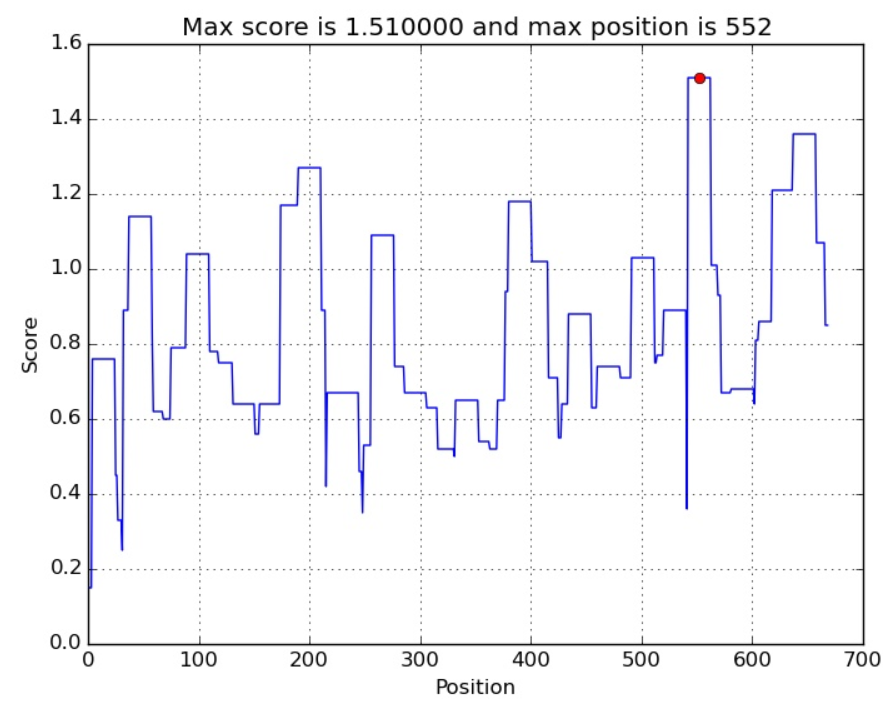

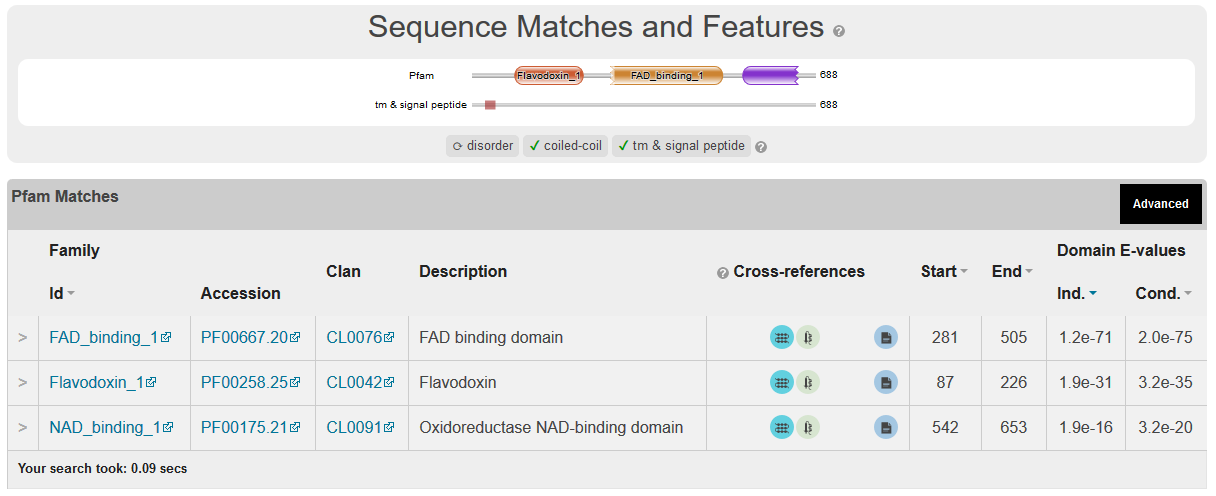

>Arabidopsis thaliana 2

Interaction prediction= 0.76 NO

Binding site Prediction

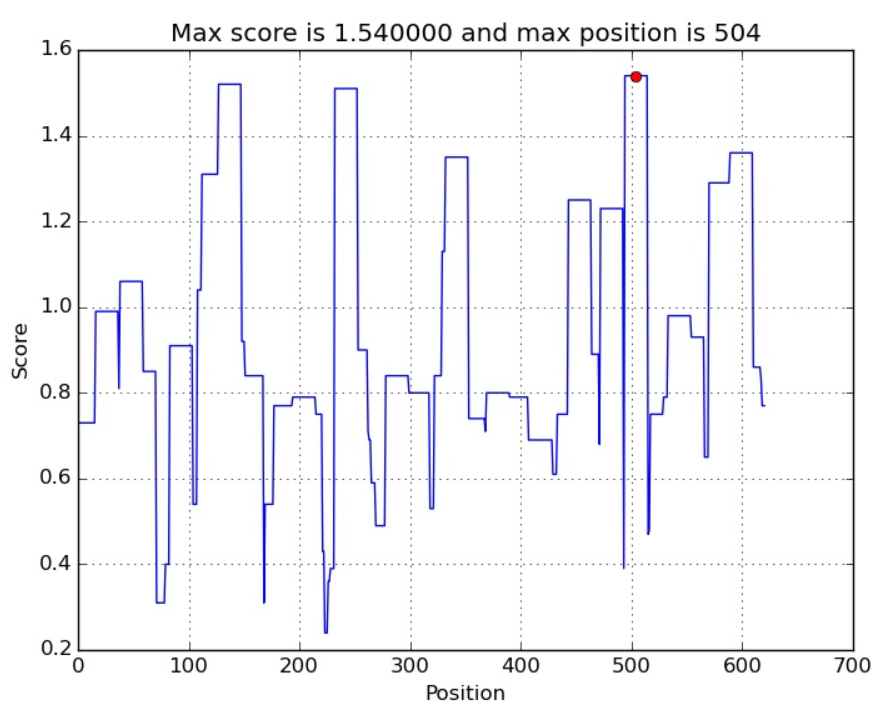

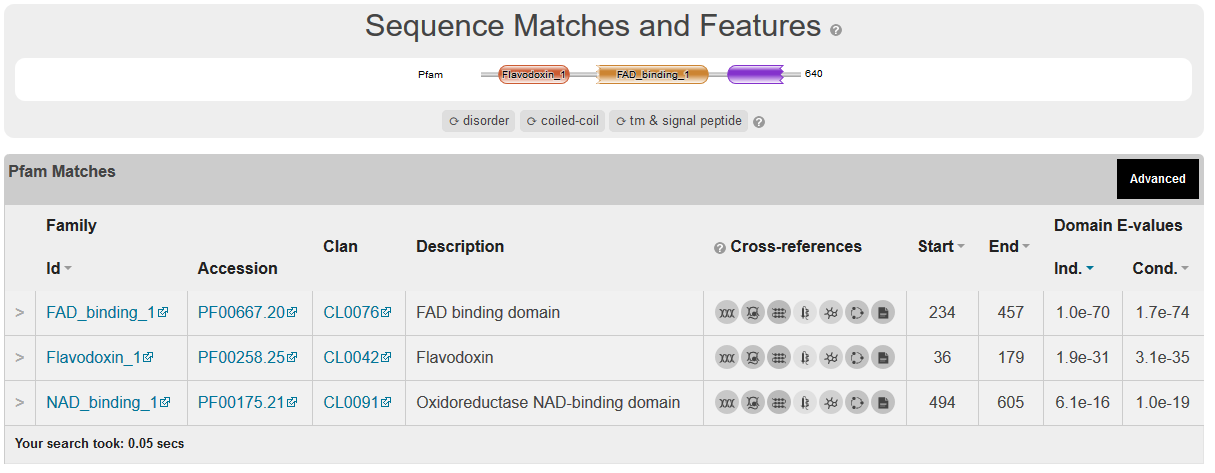

>Triticum aestivum

Interaction prediction= 1.15 YES

Binding site Prediction

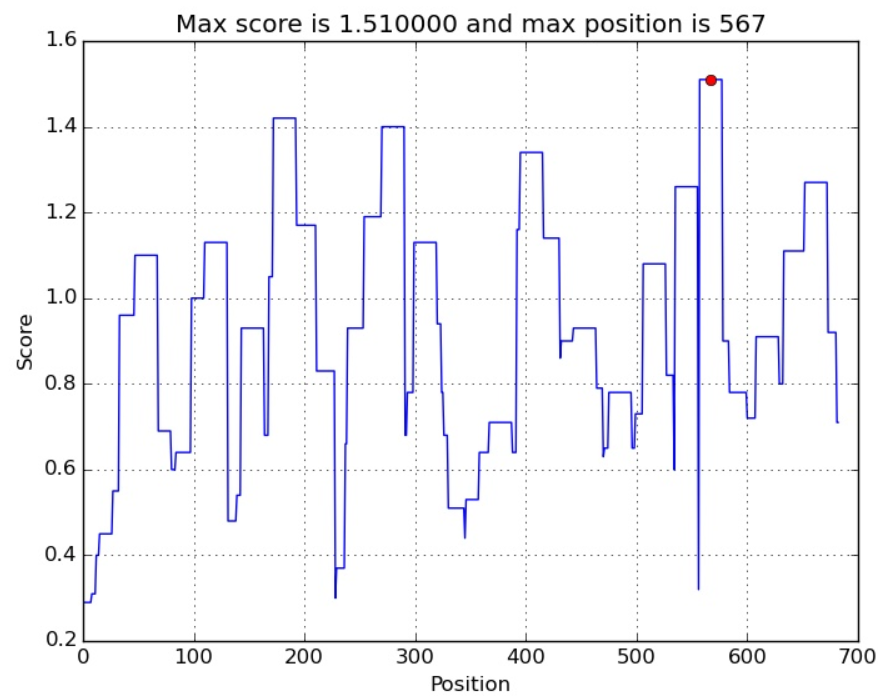

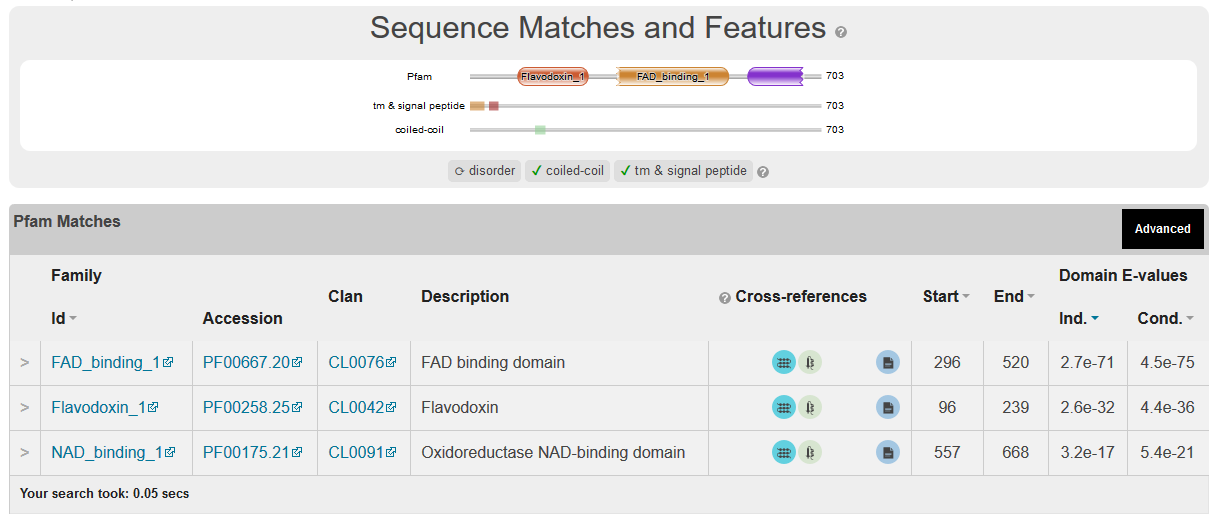

>Trypanosoma brucei

Interaction prediction= 0.26 NO

Binding site Prediction

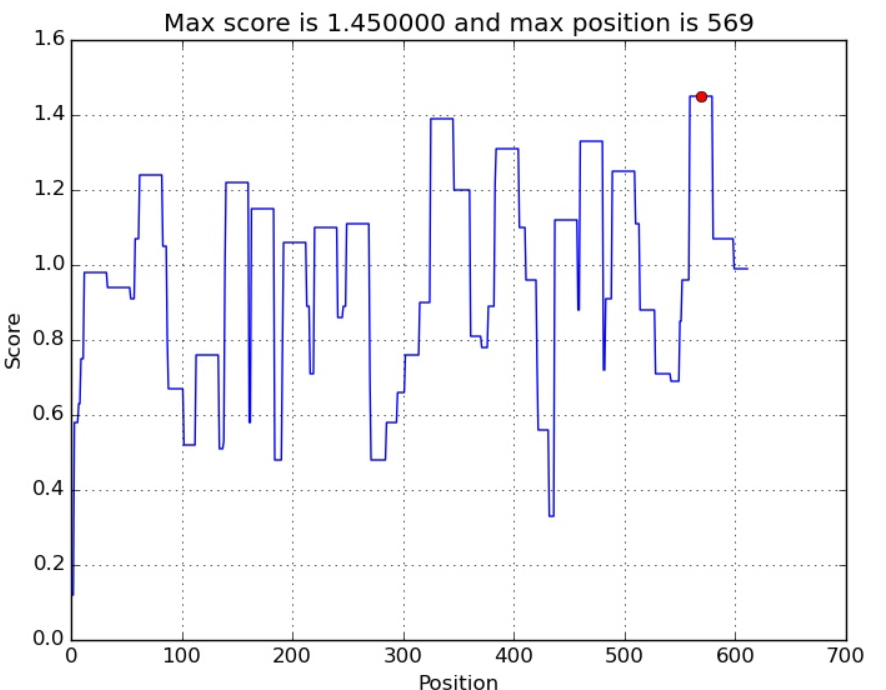

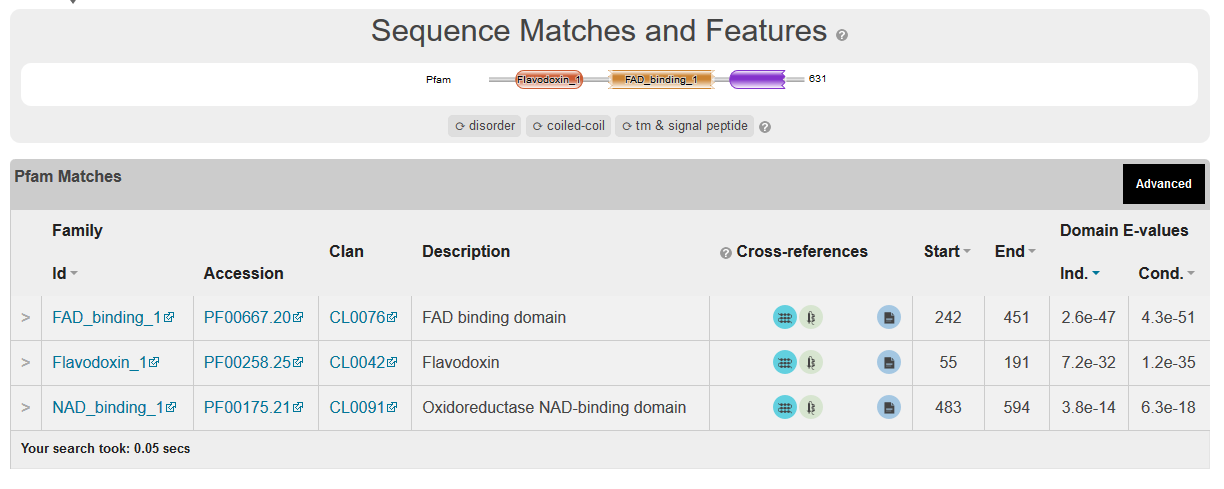

>Dictyostelium discoideum

Interaction prediction= 0.17 NO

Binding site Prediction

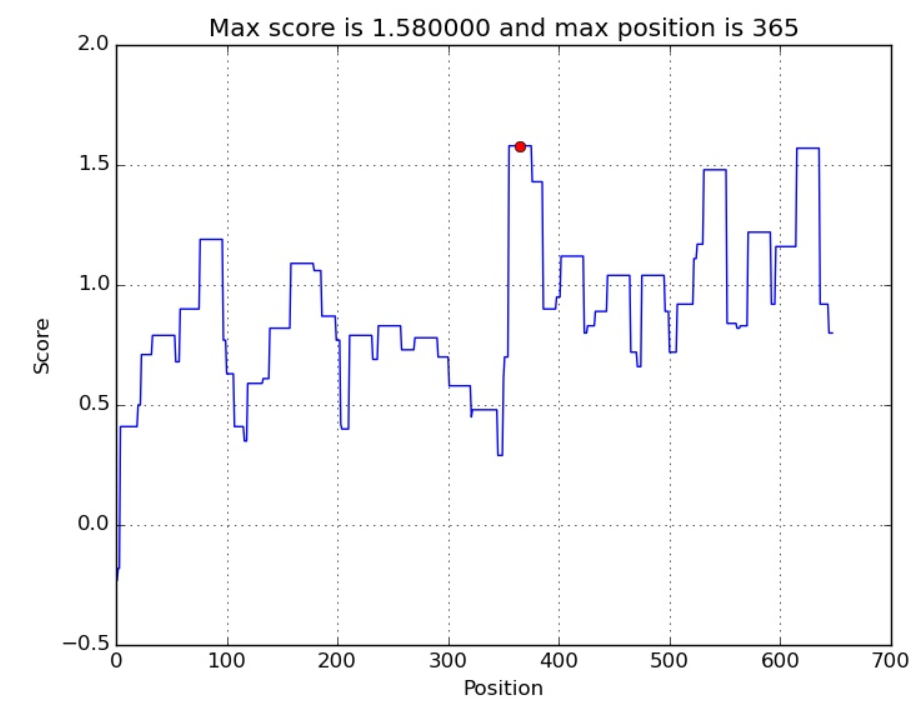

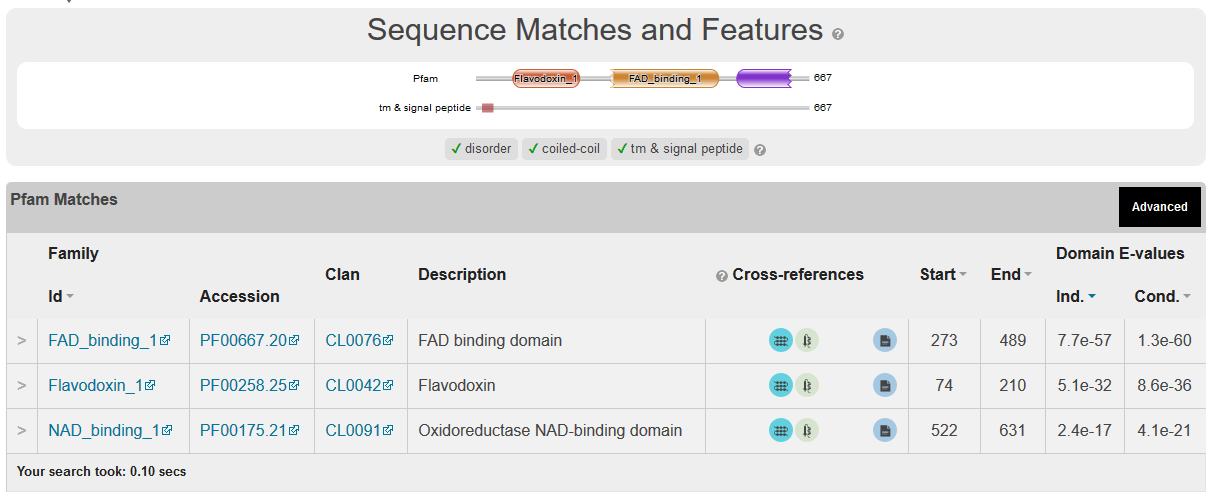

Heme reductase responsible for ergosterol production together with FGSG_01000

FGSG_08413 Has to be checked for structure (maybe be deleted then phenotype should be as FGSG_01000 when comes to endocytosis). Or a more interesting test would be to test the ratio of FGSG_08413/FGSG_01000 expression in WT and FGSG_09786 deletion. If FGSG_08413 is taking over ergosterol synthesis in FGSG_09786 deletion this ratio should be much higher in the the mutant compared to the WT.

This is the only heme-reductase that is similar to FGSG_09786

>XP_011320363.1 hypothetical protein FGSG_08413 [Fusarium graminearum PH-1]

MTNVFAADTLLSQLGPPQSIADIAALSALGVASAAYLLRGITWDKPDHYHHVWFERMGSKGGSSSSHPKATRDIAKKLEETGKDVVIFWGSQSGTAETFANRLSKECHLRFGLQALCADLCDYDPESIANLSQSKLAIFILSTYGEGDPSDNTAAFWDWLTKTPNIQIPNLRYMAFGLGNTSYRYYNRVIDVVAQHLDKYGAQRLMPVGRANDAQGGTEEDFLSWKDDLYTHFQENLGYQERDIPYEPSIQLIQDESLDIMDLHLGEPIQNRNGPAKVVKQYSPIRPLAIQSSQELYTSPGRNCLHMELDISNQPELRYRTGDHLAIYPINPDYEVQLLLKAFGLEDRAEKPLLVQTLEEGTSTKIPSPTSALALFRHYLEVAAPVSRETVGQLARFAPLSESVETLTALAKNKEAYATYIGSNHITMGRLLHLVAPGAIWTKLPLSYVVETLPCIQARYYSISSSSTVSARRLSITVGVDKSPLQQDPSRVIRGITTNYLYALGNALNCDTSQSVVGSDAPSYALSGPGDTLNGRKVFACIRRSNFKLPTLSSTPIIMIGAGTGLAPFRGFILERARLQAVGKPIGKMLLFFGCRSPDQDYLYQGELAEVAQKLQGSLEIVTAFSRAEEEPKKYVQDRVEERKTQVCHLLQEGASIYFCGRAAMARKVGNKVEESMKTQNNWTDAEARSWAESVKKGNKWLEDVWG

Interaction prediction 0.34

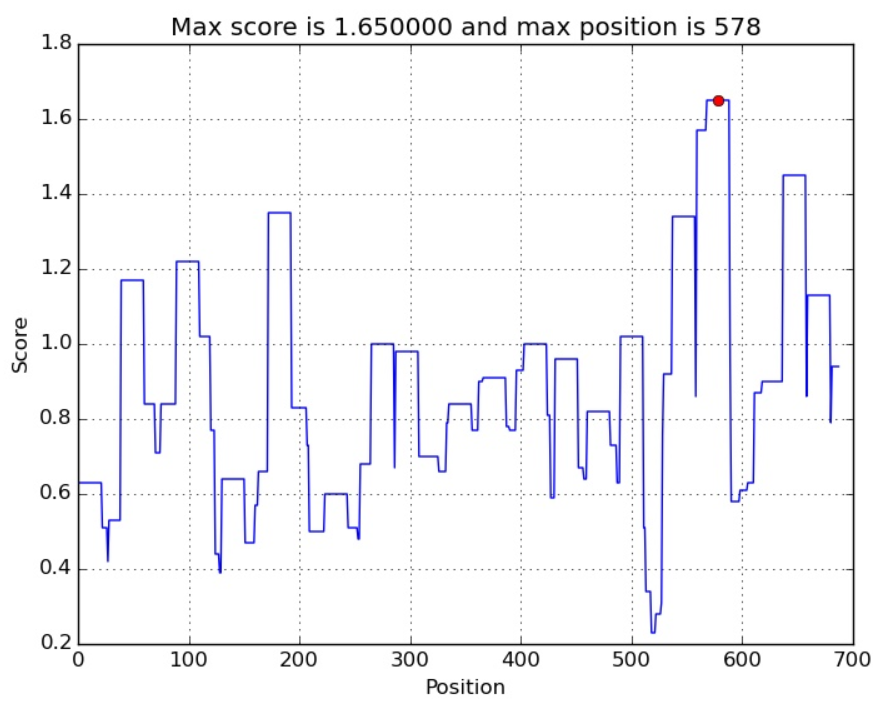

TM=20-38
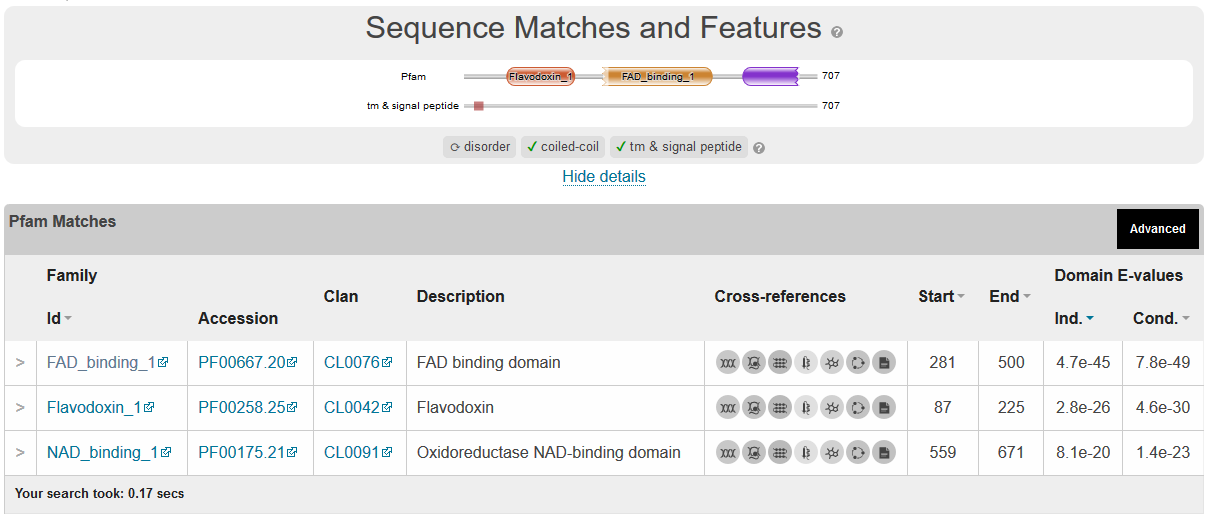

TM 20-38

The similar protein in yeast is Ergosterol ERG11 heme reductase from yeast

>YeastNCP1 YHR042W SGDID:S000001084

MPFGIDNTDFTVLAGLVLAVLLYVKRNSIKELLMSDDGDITAVSSGNRDIAQVVTENNKNYLVLYASQTGTAEDYAKKFSKELVAKFNLNVMCADVENYDFESLNDVPVIVSIFISTYGEGDFPDGAVNFEDFICNAEAGALSNLRYNMFGLGNSTYEFFNGAAKKAEKHLSAAGAIRLGKLGEADDGAGTTDEDYMAWKDSILEVLKDELHLDEQEAKFTSQFQYTVLNEITDSMSLGEPSAHYLPSHQLNRNADGIQLGPFDLSQPYIAPIVKSRELFSSNDRNCIHSEFDLSGSNIKYSTGDHLAVWPSNPLEKVEQFLSIFNLDPETIFDLKPLDPTVKVPFPTPTTIGAAIKHYLEITGPVSRQLFSSLIQFAPNADVKEKLTLLSKDKDQFAVEITSKYFNIADALKYLSDGAKWDTVPMQFLVESVPQMTPRYYSISSSSLSEKQTVHVTSIVENFPNPELPDAPPVVGVTTNLLRNIQLAQNNVNIAETNLPVHYDLNGPRKLFANYKLPVHVRRSNFRLPSNPSTPVIMIGPGTGVAPFRGFIRERVAFLESQKKGGNNVSLGKHILFYGSRNTDDFLYQDEWPEYAKKLDGSFEMVVAHSRLPNTKKVYVQDKLKDYEDQVFEMINNGAFIYVCGDAKGMAKGVSTALVGILSRGKSITTDEATELIKMLKTSGRYQEDVW

IPS 0.27

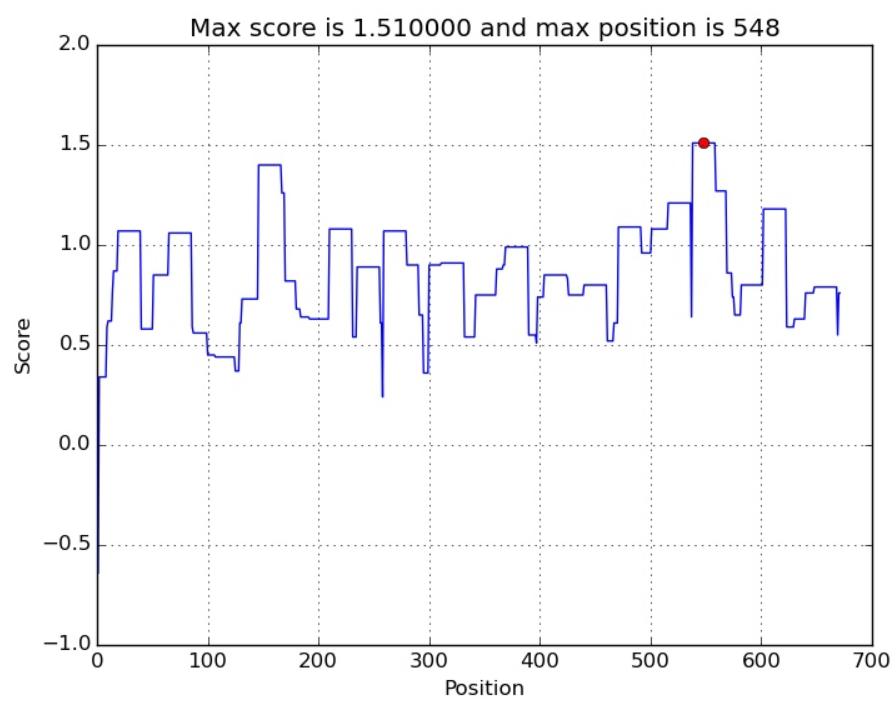

Interaction prediction=0.27 NO

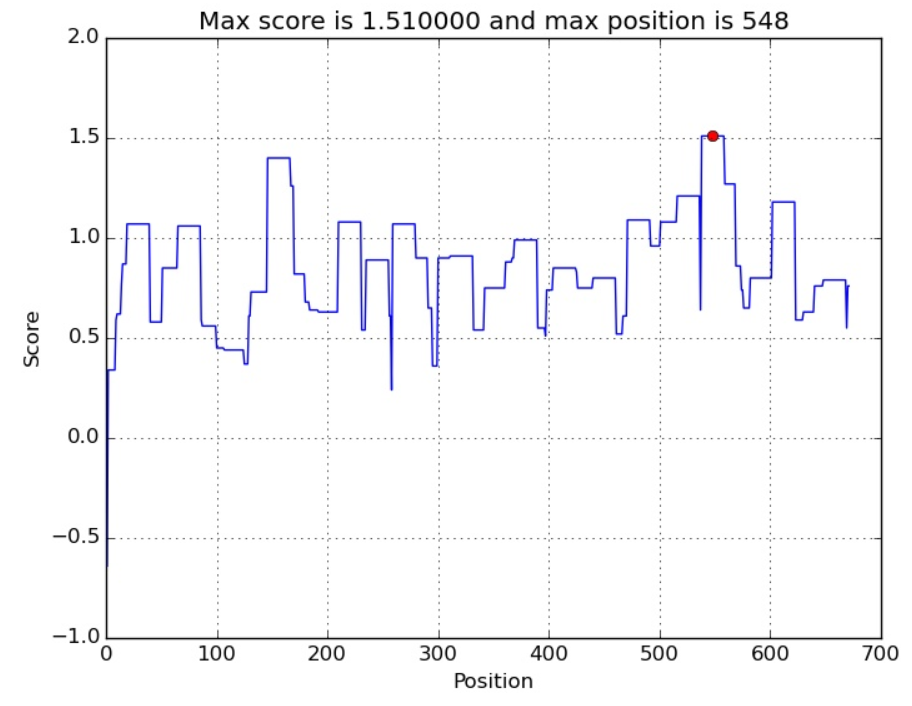

TM=1-26

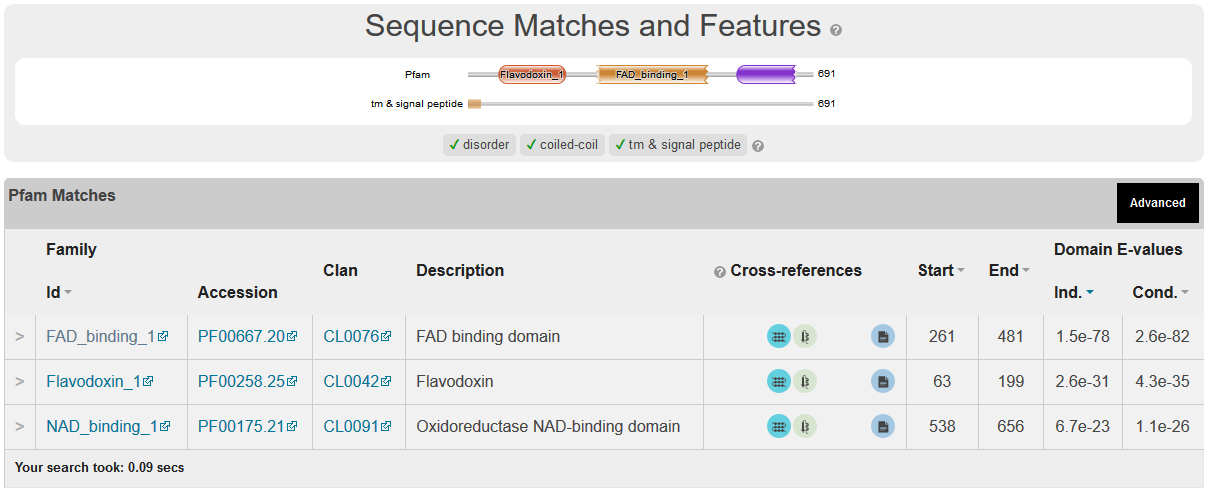

```
252 identities

XP_011320363.1      MTNVFAADTLLSQLGPPQSIADIAALSALGVASAAYLLRGITWDKPDHYHHVWFERMGSK
NCP1                --MPFGIDN-----------TDFTVLAGLVLAVLLYVKR-----------NSIKELLMSD
                        *. *.           :*::.*:.* :*   *: *           :   * : *.

XP_011320363.1      GGSSSSHPKATRDIAKKLEETGKDVVIFWGSQSGTAETFANRLSKECHLRFGLQALCADL
NCP1                DGDITAVSSGNRDIAQVVTENNKNYLVLYASQTGTAEDYAKKFSKELVAKFNLNVMCADV
                    .*. :: ....****: : *..*: ::::.**:**** :*:.:***   .*.*:.:***:

XP_011320363.1      CDYDPESIANLSQSKLAIFILSTYGEGDPSDNTAAFWDWLTKTPNIQIPNLRYMAFGLGN
NCP1                ENYDFESLNDVPVI-VSIFI-STYGEGDFPDGAVNFEDFICNAEAGALSNLRYNMFGLGN
                     :** **: ::.   ::*** ******* .*.:. * *:: ::    :.****  *****

XP_011320363.1      TSYRYYNRVIDVVAQHLDKYGAQRLMPVGRANDAQGGTEEDFLSWKDDLYTHFQENLGYQ
NCP1                STYEFFNGAAKKAEKHLSAAGAIRLGKLGEADDGAGTTDEDYMAWKDSILEVLKDELHLD
                    ::* ::* . . . :**.  ** **  :* *:*. * *:**:::***.:   ::::*  :

XP_011320363.1      ERDIPYEPSIQL-IQDESLDIMDLHLGEPIQ--------NRNGPAKVVKQYSPIRPL--A
NCP1                EQEAKFTSQFQYTVLNEITDSMS--LGEPSAHYLPSHQLNRNADGIQLGPFDLSQPYIAP
                    *.:  : ..:*  : :*  * *.  ****          ***. .  :  :.  .*   .

XP_011320363.1      IQSSQELYTSPGRNCLHMELDISNQPELRYRTGDHLAIYPINPDYEVQLLLKAFGLEDRA
NCP1                IVKSRELFSSNDRNCIHSEFDLSG-SNIKYSTGDHLAVWPSNPLEKVEQFLSIFNLD--P
                    * .*.**::* .***:* *:*:*. .::.* ******::* **  :*: :*. *.*:  .

XP_011320363.1      EKPLLVQTLEEGTSTKIPSPTSALALFRHYLEVAAPVSRETVGQLARFAPLSESVETLTA
NCP1                ETIFDLKPLDPTVKVPFPTPTTIGAAIKHYLEITGPVSRQLFSSLIQFAPNADVKEKLTL
                    *. : ::.*:  ... :*:**:  * :.****::.****: ...* .*** ::  *.**

XP_011320363.1      LAKNKEAYATYIGSNHITMGRLLHLVAPGAIWTKLPLSYVVETLPCIQARYYSISSSSTV
NCP1                LSKDKDQFAVEITSKYFNIADALKYLSDGAKWDTVPMQFLVESVPQMTPRYYSISSSSLS
                    *:*:*: :*. * *:::.:.  *: :: ** * .:*:.::**::* : .*********

XP_011320363.1      SARRLSITVGVDKSPLQQDP-SRVIRGITTNYLYALGNALNCDTSQSVVGSDAP-SYALS
NCP1                EKQTVHVTSIVENFPNPELPDAPPVVGVTTNLLRNIQLAQN---NVNIAETNLPVHYDLN
                    . . : :*  *:: *  : * :  : *:*** *  :  * *   . .:. :: *  * *.

XP_011320363.1      GPGDTLNGRKVFACIRRSNFKLPTLSSTPIIMIGAGTGLAPFRGFILERARLQAVGK---
NCP1                GPRKLFANYKLPVHVRRSNFRLPSNPSTPVIMIGPGTGVAPFRGFIRERVAFLESQKKGG
                    ** . : . *: . :*****.**: .***:****.***:******* **. :    *

XP_011320363.1      ---PIGKMLLFFGCRSPDQDYLYQGELAEVAQKLQGSLEIVTAFSRAEEEPKKYVQDRVE
NCP1                NNVSLGKHILFYGSRNTD-DFLYQDEWPEYAKKLDGSFEMVVAHSRLPNTKKVYVQDKLK
                       .:** :**:*.*..* *:***.* .* *:**:**:*:*.*.**  :  * ****.::

XP_011320363.1      ERKTQVCHLLQEGASIYFCGRA-AMARKVGNKVEESMKTQNNWTDAEARSWAESVKKGNK
NCP1                DYEDQVFEMINNGAFIYVCGDAKGMAKGVSTALVGILSRGKSITTDEATELIKMLKTSGR
                    : : **  ::::** **.** * .**. *.. :   :.  :. *  ** .  : :*....

XP_011320363.1      WLEDVWG
NCP1                YQEDVW-
                    : ****

299 identities

FgPH-1          -MAELDTLDVIVLGVIFLGTVAYFTKGKLWGV--TKDPYANGFAAGGAAKPGRTRNIVEA
NCP1            MPFGIDNTDFTVLAGLVLAVLLYVKRNSIKELLMSDDGDITAVSSG-------NRDIAQV
                    :*. *. **. :.*..: *.....:. :  :.*   ...::*       .*:*.:.

FgPH-1          MEESGKNCVIFYGSQTGTAEDYASRLAKEGKSRFGLNTMIADIEDYDFDSLDTVPNDNVV
NCP1            VTENNKNYLVLYASQTGTAEDYAKKFSKELVAKFNLNVMCADVENYDFESLNDVP--VIV
                : *..** :::*.**********..::**  :.*.**.* **:*:***:**: **   :*

FgPH-1          MFVLATYGEGEPTDNAVDFYEFITGEDATFNEGNDPPLGNLNYVAFGLGNNTYEHYNAMV
NCP1            SIFISTYGEGDFPDGAVNFEDFICNAEAG-------ALSNLRYNMFGLGNSTYEFFNGAA
                 :.::*****: .*.**:* :** . :*        .*.**.*  *****.***.:*. .

FgPH-1          RKVDQALEKFGAHRIGEAGEGDDGAGTMEEDFLAWKDPMWESLAKKMGLEEREAVYEPIF
NCP1            KKAEKHLSAAGAIRLGKLGEADDGAGTTDEDYMAWKDSILEVLKDELHLDEQEAKFTSQF
                .*.:: *.  ** *:*: **.****** :**::****.: * * .:: *:*.** : . *

FgPH-1          AINERDDLSPESNEVYLGEPNKLHL---------EGTAKGPFNSHNPYIAPIAESYELFS
NCP1            QYTVLNEI---TDSMSLGEPSAHYLPSHQLNRNADGIQLGPFDLSQPYIAPIVKSRELFS
                  .  :::   ::.: ****.  :*         :*   ***:  :******.:* ****

FgPH-1          AKDRNCLHMEVDISGSNLKYETGDHIAIWPTNPGEEVNRFLDILDLSGKQHSVITVKALE
NCP1            SNDRNCIHSEFDLSGSNIKYSTGDHLAVWPSNPLEKVEQFLSIFNL--DPETIFDLKPLD
                ::****:* *.*:****:**.****:*:**:** *:*:.**.*::*  .  ::: :*.*:

FgPH-1          PTAKVPFPNPTTYDAILRYHLEICAPVSRQFVSTLAAFAPNDSIKAEMNRLGSDKDYFHE
NCP1            PTVKVPFPTPTTIGAAIKHYLEITGPVSRQLFSSLIQFAPNADVKEKLTLLSKDKDQFAV
                **.*****.*** .* :.::*** .*****:.*:*  **** .:* ::. *..*** *

FgPH-1          KTGPHYYNIARFLSSVSKGEKWTTIPFSAFIEGLTKLQPRYYSISSSSLVQPKKISITAV
NCP1            EITSKYFNIADALKYLSDGAKWDTVPMQFLVESVPQMTPRYYSISSSSLSEKQTVHVTSI
                :  .:*:***  *. :*.* ** *:*:. ::*.:.:: *********** : :.: :*::

FgPH-1          VESQQIPGRDD--PFRGVATNYLFALKQKQNG-DPSPAPFGQTYELTGPRNKYDGIHVPV
NCP1            VENFPNPELPDAPPVVGVTTNLLRNIQLAQNNVNIAETNLPVHYDLNGPRKLFANYKLPV
                **.   *   *  *. **:** *  ::  **. : : : :   *:*.***: : . ::**

FgPH-1          HVRHSNFKLPSDPGKPVIMIGPGTGVAPFRGFVQERA------KLARDGVEVGKTLLFFG
NCP1            HVRRSNFRLPSNPSTPVIMIGPGTGVAPFRGFIRERVAFLESQKKGGNNVSLGKHILFYG
                ***.***.***:*..*****************:.**.      * . :.*.:** :**:*

FgPH-1          CRKPSEDFMYEKEWQEYKEALGDKFEMITAFSR-ESAKKVYVQHRLKERAQEVSDLLSQK
NCP1            SRN-TDDFLYQDEWPEYAKKLDGSFEMVVAHSRLPNTKKVYVQDKLKDYEDQVFEMINNG
                .*: ::**:*:.** ** : *...***:.*.**  .:****** .**:  ::* :::.:

FgPH-1          AYFYVCGDASNMAREVNTVLAQIIAEGRGVSEAKGEEIVKNMRSANQYQEDVWS
NCP1            AFIYVCGDAKGMAKGVSTALVGILSRGKSITTDEATELIKMLKTSGRYQEDVW-
                *::******..**. *.*.*. *:: *..::  :. *::* :.::..******
```

Comparisons with Mouse

>NP_001300851.1 nitric oxide synthase, inducible isoform b [Mus musculus] Highlight Haem binding region

MNPKSLTRGPRDKPTPLEELLPHAIEFINQYYGSFKEAKIEEHLARLEAVTKEIETTGTYQLTLDELIFA

TKMAWRNAPRCIGRIQWSNLQVFDARNCSTAQEMFQHICRHILYATNNGNIRSAITVFPQRSDGKHDFRL

WNSQLIRYAGYQMPDGTIRGDAATLEFTQLCIDLGWKPRYGRFDVLPLVLQADGQDPEVFEIPPDLVLEV

TMEHPKYEWFQELGLKWYALPAVANMLLEVGGLEFPACPFNGWYMGTEIGVRDFCDTQRYNILEEVGRRM

GLETHTLASLWKDRAVTEINVAVLHSFQKQNVTIMDHHTASESFMKHMQNEYRARGGCPADWIWLVPPVS

GSITPVFHQEMLNYVLSPFYYYQIEPWKTHIWQNEKLRPRRREIRF

RVLVKVVFFASMLMRKVMASRVRA

TVLFATETGKSEALARDLATLFSYAFNTKVVCMDQYKASTLEEEQLLLVVTSTFGNGDCPSNGQTLKKSL

FMLRELNHTFRYAVFGLGSSMYPQFCAFAHDIDQKLSHLGASQLAPTGEGDELSGQEDAFRSWAVQTFRA

ACETFDVRSKHHIQIPKRFTSNATWEPQQYRLIQSPEPLDLNRALSSIHAKNVFTMRLKSQQNLQSEKSS

RTTLLVQLTFEGSRGPSYLPGEHLGIFPGNQTALVQGILERVVDCPTPHQTVCLEVLDESGSYWVKDKRL

PPCSLSQALTYFLDITTPPTQLQLHKLARFATDETDRQRLEALCQPSEYNDWKFSNNPTFLEVLEEFPSL

HVPAAFLLSQLPILKPRYYSISSSQDHTPSEVHLTVAVVTYRTRDGQGPLHHGVCSTWIRNLKPQDPVPC

FVRSVSGFQLPEDPSQPCILIGPGTGIAPFRSFWQQRLHDSQHKGLKGGRMSLVFGCRHPEEDHLYQEEM

QEMVRKRVLFQVHTGYSRLPGKPKVYVQDILQKQLANEVLSVLHGEQGHLYICGDVRMARDVATTLKKLV

ATKLNLSEEQVEDYFFQLKSQKRYHEDIFGAVFSYGAKKGSALEEPKATR

>NP_035057.1 nitric oxide synthase, inducible isoform a [Mus musculus]

MACPWKFLFKVKSYQSDLKEEKDINNNVKKTPCAVLSPTIQDDPKSHQNGSPQLLTGTAQNVPESLDKLH

VTSTRPQYVRIKNWGSGEILHDTLHHKATSDFTCKSKSCLGSIMNPKSLTRGPRDKPTPLEELLPHAIEF

INQYYGSFKEAKIEEHLARLEAVTKEIETTGTYQLTLDELIFATKMAWRNAPRCIGRIQWSNLQVFDARN

CSTAQEMFQHICRHILYATNNGNIRSAITVFPQRSDGKHDFRLWNSQLIRYAGYQMPDGTIRGDAATLEF

TQLCIDLGWKPRYGRFDVLPLVLQADGQDPEVFEIPPDLVLEVTMEHPKYEWFQELGLKWYALPAVANML

LEVGGLEFPACPFNGWYMGTEIGVRDFCDTQRYNILEEVGRRMGLETHTLASLWKDRAVTEINVAVLHSF

QKQNVTIMDHHTASESFMKHMQNEYRARGGCPADWIWLVPPVSGSITPVFHQEMLNYVLSPFYYYQIEPW

KTHIWQNEKLRPRRREIRFRVLVKVVFFASMLMRKVMASRVRATVLFATETGKSEALARDLATLFSYAFN

TKVVCMDQYKASTLEEEQLLLVVTSTFGNGDCPSNGQTLKKSLFMLRELNHTFRYAVFGLGSSMYPQFCA

FAHDIDQKLSHLGASQLAPTGEGDELSGQEDAFRSWAVQTFRAACETFDVRSKHHIQIPKRFTSNATWEP

QQYRLIQSPEPLDLNRALSSIHAKNVFTMRLKSQQNLQSEKSSRTTLLVQLTFEGSRGPSYLPGEHLGIF

PGNQTALVQGILERVVDCPTPHQTVCLEVLDESGSYWVKDKRLPPCSLSQALTYFLDITTPPTQLQLHKL

ARFATDETDRQRLEALCQPSEYNDWKFSNNPTFLEVLEEFPSLHVPAAFLLSQLPILKPRYYSISSSQDH

TPSEVHLTVAVVTYRTRDGQGPLHHGVCSTWIRNLKPQDPVPCFVRSVSGFQLPEDPSQPCILIGPGTGI

APFRSFWQQRLHDSQHKGLKGGRMSLVFGCRHPEEDHLYQEEMQEMVRKRVLFQVHTGYSRLPGKPKVYV

QDILQKQLANEVLSVLHGEQGHLYICGDVRMARDVATTLKKLVATKLNLSEEQVEDYFFQLKSQKRYHED

IFGAVFSYGAKKGSALEEPKATR

Nos Isoform a

Ipred =1 YES

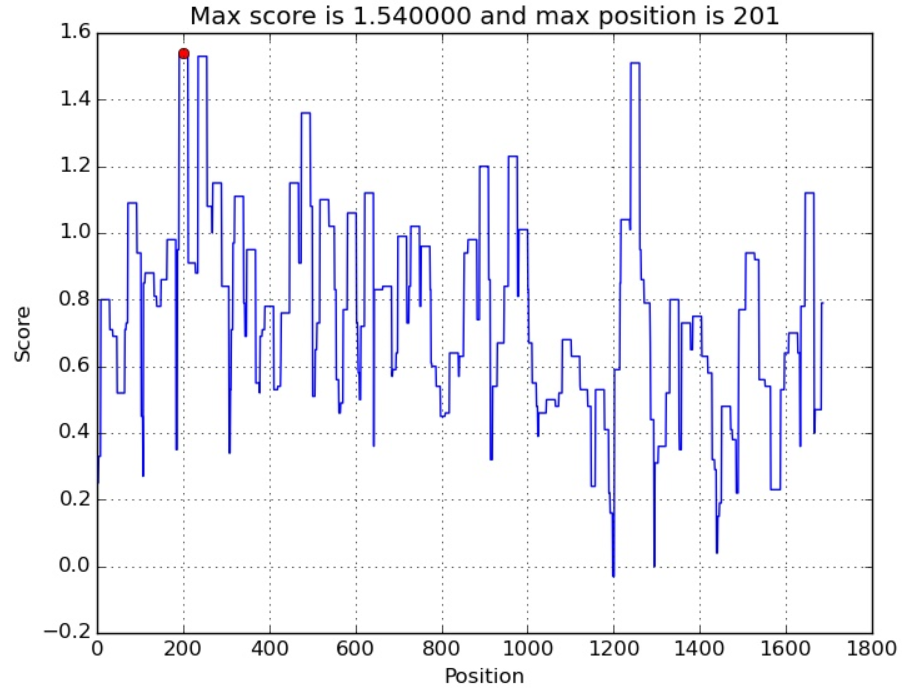

TM 509-527

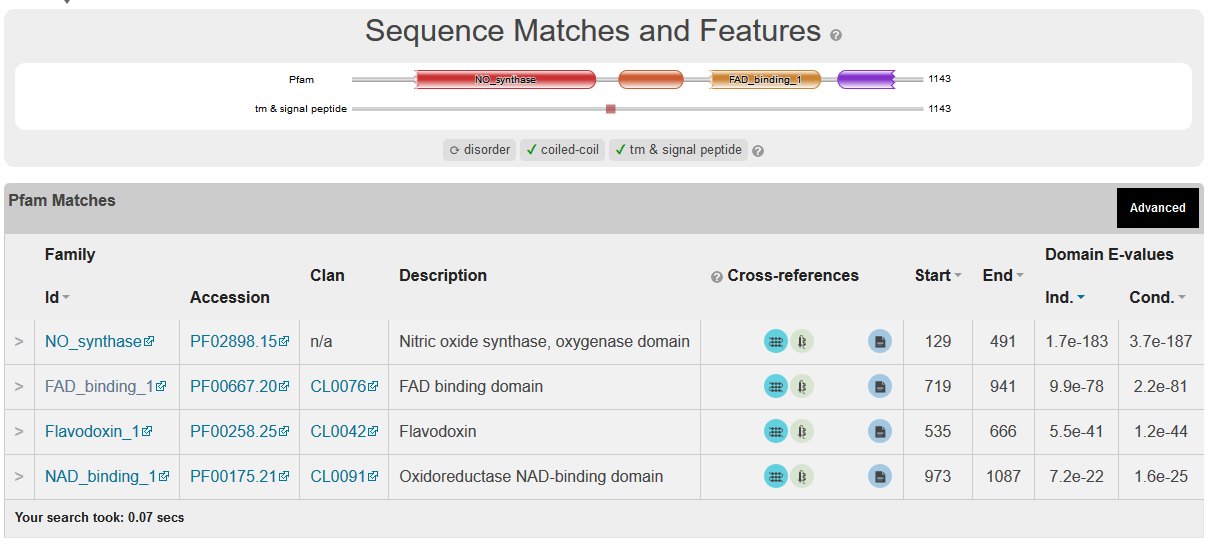

NOS Isoform b

TM=396-414

Interaction prediction=1 Yes

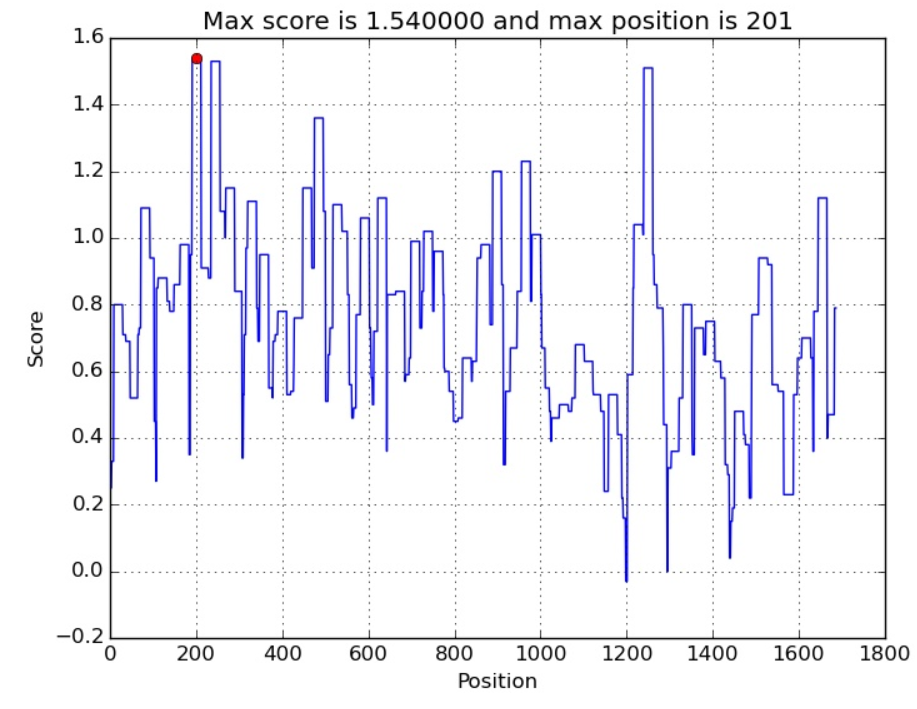

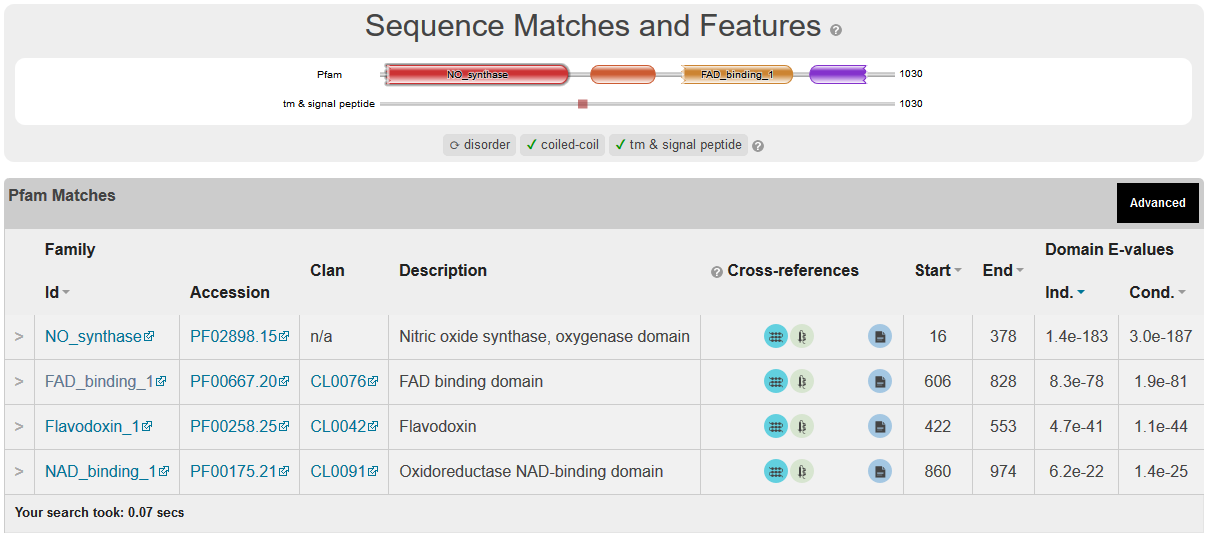

C-terminal including the TM (the reductase part)

RVLVKVVFFASMLMRKVMASRVRA

TVLFATETGKSEALARDLATLFSYAFNTKVVCMDQYKASTLEEEQLLLVVTSTFGNGDCPSNGQTLKKSL

FMLRELNHTFRYAVFGLGSSMYPQFCAFAHDIDQKLSHLGASQLAPTGEGDELSGQEDAFRSWAVQTFRA

ACETFDVRSKHHIQIPKRFTSNATWEPQQYRLIQSPEPLDLNRALSSIHAKNVFTMRLKSQQNLQSEKSS

RTTLLVQLTFEGSRGPSYLPGEHLGIFPGNQTALVQGILERVVDCPTPHQTVCLEVLDESGSYWVKDKRL

PPCSLSQALTYFLDITTPPTQLQLHKLARFATDETDRQRLEALCQPSEYNDWKFSNNPTFLEVLEEFPSL

HVPAAFLLSQLPILKPRYYSISSSQDHTPSEVHLTVAVVTYRTRDGQGPLHHGVCSTWIRNLKPQDPVPC

FVRSVSGFQLPEDPSQPCILIGPGTGIAPFRSFWQQRLHDSQHKGLKGGRMSLVFGCRHPEEDHLYQEEM

QEMVRKRVLFQVHTGYSRLPGKPKVYVQDILQKQLANEVLSVLHGEQGHLYICGDVRMARDVATTLKKLV

ATKLNLSEEQVEDYFFQLKSQKRYHEDIFGAVFSYGAKKGSALEEPKATR

Interaction Prediction =0.99

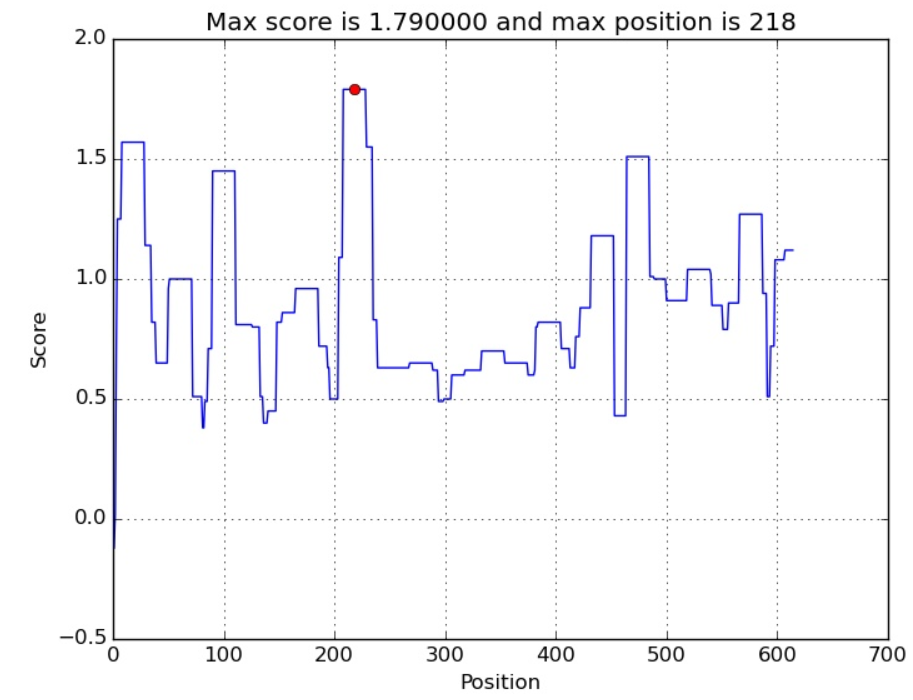

TM 1-19

>NP_032924.1 NADPH--cytochrome P450 reductase [Mus musculus] Similar to the uman gene above and to Ph1

MGDSHEDTSATVPEAVAEEVSLFSTTDIVLFSLIVGVLTYWFIFKKKKEEIPEFSKIQTTAPPVKESSFV

EKMKKTGRNIIVFYGSQTGTAEEFANRLSKDAHRYGMRGMSADPEEYDLADLSSLPEIDKSLVVFCMATY

GEGDPTDNAQDFYDWLQETDVDLTGVKFAVFGLGNKTYEHFNAMGKYVDQRLEQLGAQRIFELGLGDDDG

NLEEDFITWREQFWPAVCEFFGVEATGEESSIRQYELVVHEDMDTAKVYTGEMGRLKSYENQKPPFDAKN

PFLAAVTTNRKLNQGTERHLMHLELDISDSKIRYESGDHVAVYPANDSTLVNQIGEILGADLDVIMSLNN

LDEESNKKHPFPCPTTYRTALTYYLDITNPPRTNVLYELAQYASEPSEQEHLHKMASSSGEGKELYLSWV

VEARRHILAILQDYPSLRPPIDHLCELLPRLQARYYSIASSSKVHPNSVHICAVAVEYEAKSGRVNKGVA

TSWLRTKEPAGENGRRALVPMFVRKSQFRLPFKPTTPVIMVGPGTGVAPFMGFIQERAWLREQGKEVGET

LLYYGCRRSDEDYLYREELARFHKDGALTQLNVAFSREQAHKVYVQHLLKRDKEHLWKLIHEGGAHIYVC

GDARNMAKDVQNTFYDIVAEFGPMEHTQAVDYVKKLMTKGRYSLDVWS

Interaction prediction 0.84 NO

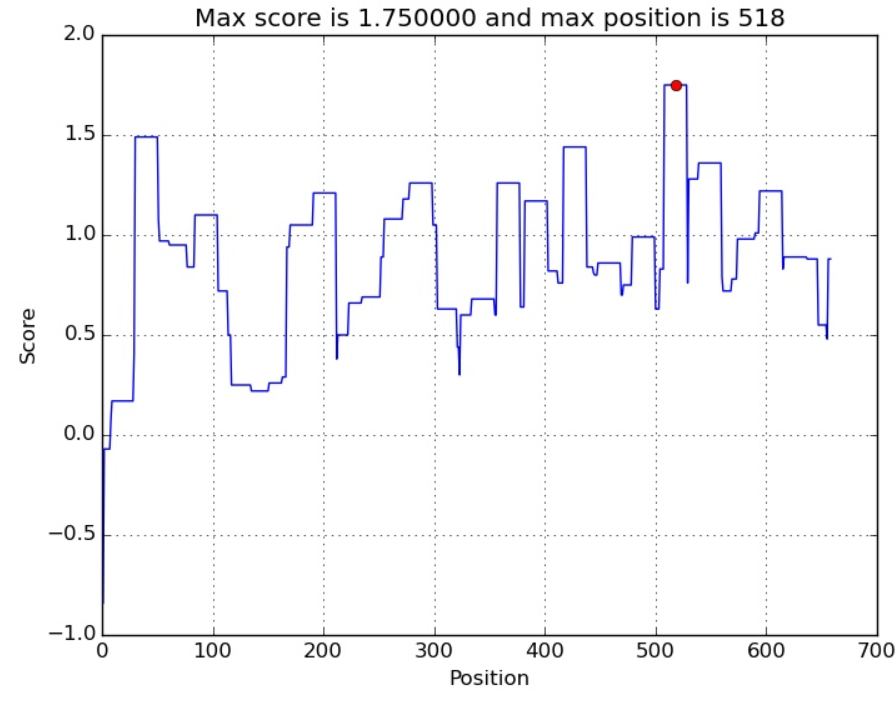

TM=20-43

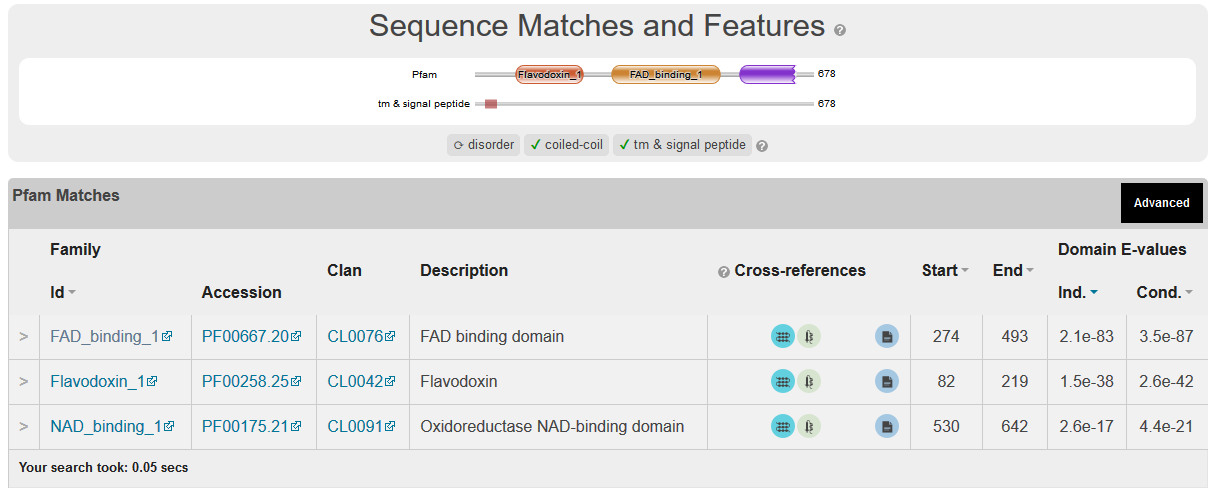

>NP_032738.1 nitric oxide synthase, brain [Mus musculus]

MEEHTFGVQQIQPNVISVRLFKRKVGGLGFLVKERVSKPPVIISDLIRGGAAEQSGLIQAGDIILAVNDRPLVDLSYDSALEVLRGIASETHVVLILRGPEGFTTHLETTFTGDGTPKTIRVTQPLGTPTKAVDLSRQPSASKDQPLAVDRVPGPSNGPQHAQGRGQGAGSVSQANGVAIDPTMKNTKANLQDSGEQDELLKEIEPVLSILTGGGKAVNRGGPAKAEMKDTGIQVDRDLDGKLHKAPPLGGENDRVFNDLWGKGNVPVVLNNPYSENEQSPASGKQSPTKNGSPSRCPRFLKVKNWETDVVLTDTLHLKSTLETGCTEQICMGSIMLPSHHIRKSEDVRTKDQLFPLAKEFLDQYYSSIKRFGSKAHMDRLEEVNKEIESTSTYQLKDTELIYGAKHAWRNASRCVGRIQWSKLQVFDARDCTTAHGMFNYICNHVKYATNKGNLRSAITIFPQRTDGKHDFRVWNSQLIRYAGYKQPDGSTLGDPANVEFTEICIQQGWKPPRGRFDVLPLLLQANGNDPELFQIPPELVLEVPIRHPKFDWFKDLGLKWYGLPAVSNMLLEIGGLEFSACPFSGWYMGTEIGVRDYCDNSRYNILEEVAKKMDLDMRKTSSLWKDQALVEINIAVLYSFQSDKVTIVDHHSATESFIKHMENEYRCRGGCPADWVWIVPPMSGSITPVFHQEMLNYRLTPSFEYQPDPWNTHVWKGTNGTPTKRRAIGFKKLAEAVKFSAKLMGQAMAKRVKATILYATETGKSQAYAKTLCEIFKHAFDAKAMSMEEYDIVHLEHEALVLVVTSTFGNGDPPENGEKFGCALMEMRHPNSVQEERKSYKVRFNSVSSYSDSRKSSGDGPDLRDNFESTGPLANVRFSVFGLGSRAYPHFCAFGHAVDTLLEELGGERILKMREGDELCGQEEAFRTWAKKVFKAACDVFCVGDDVNIEKANNSLISNDRSWKRNKFRLTYVAEAPELTQGLSNVHKKRVSAARLLSRQNLQSPKSSRSTIFVRLHTNGNQELQYQPGDHLGVFPGNHEDLVNALIERLEDAPPANHVVKVEMLEERNTALGVISNWKDESRLPPCTIFQAFKYYLDITTPPTPLQLQQFASLATNEKEKQRLLVLSKGLQEYEEWKWGKNPTMVEVLEEFPSIQMPATLLLTQLSLLQPRYYSISSSPDMYPDEVHLTVAIVSYHTRDGEGPVHHGVCSSWLNRIQADDVVPCFVRGAPSFHLPRNPQVPCILVGPGTGIAPFRSFWQQRQFDIQHKGMNPCPMVLVFGCRQSKIDHIYREETLQAKNKGVFRELYTAYSREPDRPKKYVQDVLQEQLAESVYRALKEQGGHIYVCGDVTMAADVLKAIQRIMTQQGKLSEEDAGVFISRLRDDNRYHEDIFGVTLRTYEVTNRLRSESIAFIEESKKDTDEVFSS

Interaction prediction 0.01

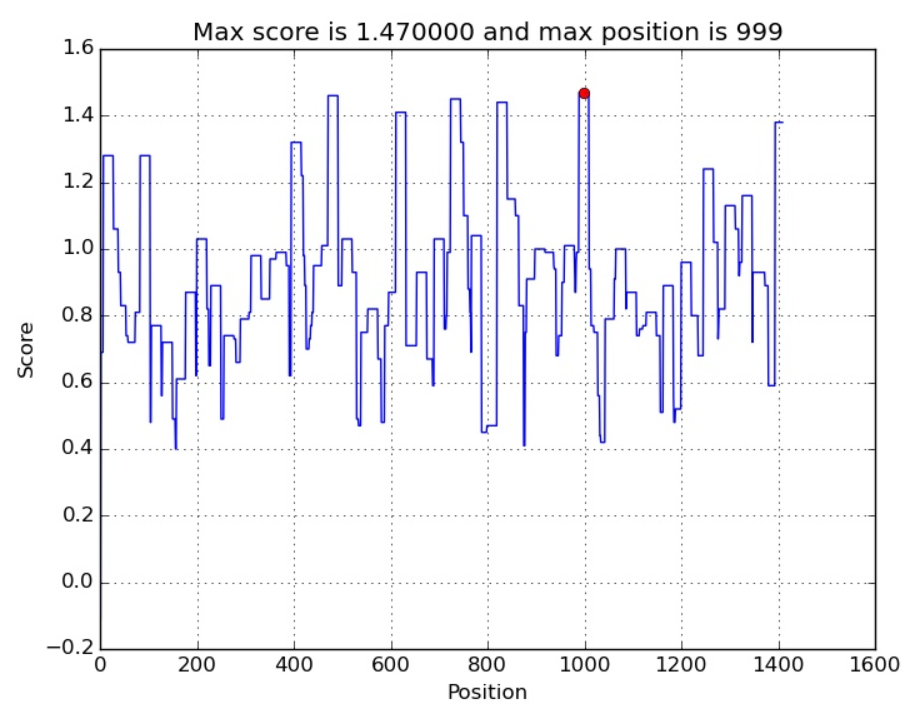

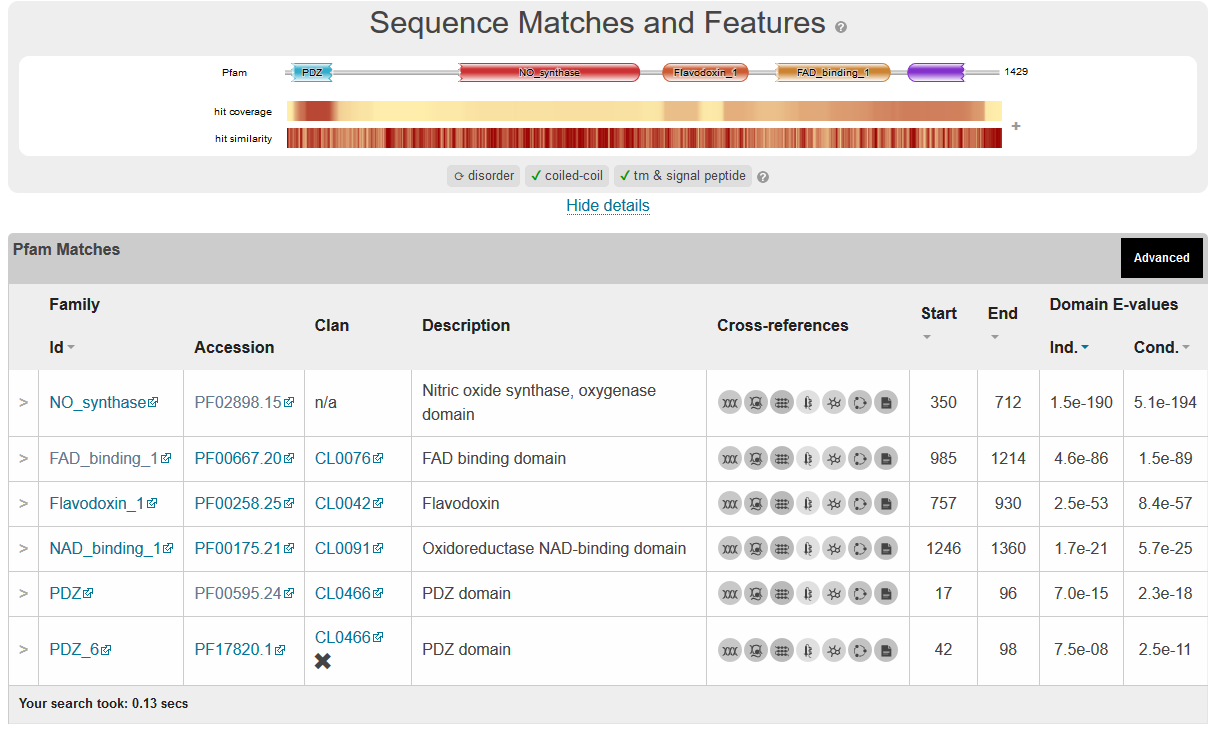
